# *rcs5-*mediated spot blotch resistance in barley is conferred by wall-associated kinases that resist pathogen manipulation

**DOI:** 10.1101/2020.04.13.040238

**Authors:** Gazala Ameen, Shyam Solanki, Thomas Drader, Lauren Sager-Bittara, Brian Steffenson, Andris Kleinhofs, Chrysafis Vogiatzis, Robert S. Brueggeman

**Affiliations:** Department of Crop and Soil Science, Washington State University, Pullman, WA 99164, USA; Department of Plant Pathology, North Dakota State University, Fargo, ND 58102, USA; Department of Plant Pathology, University of Minnesota, St. Paul, MN 55108, USA; Department of Industrial and Systems Engineering, North Carolina Agricultural and Technical State University, Greensboro, NC 27411, USA

## Abstract

Plant biotrophic pathogen disease resistances rely on immunity receptor-mediated programmed cell death (PCD) responses, but specialized necrotrophic/hemi-biotrophic pathogens hijack these mechanisms to colonize the resulting dead tissue in their necrotrophic phase. Thus, immunity receptors can become necrotrophic pathogen dominant susceptibility targets but resistance mechanisms that resist necrotroph manipulation are recessive resistance genes. The barley *rcs5* QTL imparts recessive resistance against the disease spot blotch caused by the hemi-biotrophic fungal pathogen *Bipolaris sorokiniana*. The *rcs5* genetic interval was delimited to ~0.23 cM, representing an ~234 kb genomic region containing four wall-associated kinase (WAK) genes, designated *HvWak2, Sbs1, Sbs2* (<u>s</u>usceptibility to *<u>B</u>ipolaris <u>s</u>orokiniana* <u>1</u>&<u>2</u>), and *HvWak5*. Post-transcriptional gene silencing of *Sbs1*&*2* in susceptible barley cultivars resulted in resistance showing dominant susceptibility function. Allele analysis of *Sbs1*&*2* from resistant and susceptible barley cultivars identified sequence polymorphisms associated with phenotypes in their primary coding sequence and promoter regions, suggesting differential transcriptional regulation may contribute to susceptibility. Transcript analysis of *Sbs1*&*2* showed nearly undetectable expression in resistant and susceptible cultivars prior to pathogen challenge; however, upregulation of both genes occurred specifically in susceptible cultivars post-inoculation with a virulent isolate. Apoplastic wash fluids collected from barley infected with a virulent isolate induced *Sbs1*, suggesting regulation by an apoplastic-secreted effector. Thus, *Sbs1*&2 function as *B. sorokiniana* susceptibility targets and non-functional alleles or alleles that resist induction by the pathogen mediate *rcs5-*recessive resistance. The *sbs1*&*2* alleles underlying the *rcs5* QTL that the pathogen is unable to manipulate are the first resistance genes identified against spot blotch.

**SIGNIFICANCE STATEMENT:** The *rcs5* locus in barley confers a high level of seedling resistance and a moderate level of adult plant resistance to spot blotch. It is part of a complex that has provided durable spot blotch resistance in many North American barley cultivars (cv) for more than 50 years. Genetic characterization and positional cloning of *rcs5* identified the dominant susceptibility genes, *Sbs1* and *Sbs2* (susceptibility to *Bipolaris sorokiniana* 1 and 2) as wall-associated kinases. These genes are hijacked by the hemibiotrophic pathogen in its necrotrophic phase to induce programmed cell death, facilitating disease development. We report the first spot blotch resistance/susceptibility genes cloned that function via alleles that cannot be specifically induced and hijacked by virulent isolates of the pathogen.

## INTRODUCTION

Barley ranks fourth among cereals with respect to area under production in the world (1). Although barley is not a staple food crop, it possesses unique malting characteristics that are valued for the production of beer and spirits; multi-billion dollar industries across the world. Recent studies predicted that climate change presents a threat to barley production due to higher temperatures and water deficiencies (2). Although, climate change results in drought stricken regions, others experience excess precipitation (3) and combined with elevated temperatures will provide environments more conducive to the development of fungal disease epidemics. Thus, climate change will exacerbate disease problems in areas experiencing excess precipitation, which was not addressed in the recent predictions of future barley shortages suggesting it could be worse than predicted (2).

One of the major diseases attacking barley is spot blotch, which is caused by the hemi-biotrophic/ necrotrophic ascomycete pathogen *Bipolaris sorokiniana* (teleomorph: *Cochliobolus sativus*). Spot blotch attacks both barley and wheat, causing necrotic, elongated lesions on the leaves, sheath, and stem. In addition to the foliar spot blotch disease, *B*. *sorokiniana* also causes black point on the kernels (4) and common root rot (5). Spot blotch is distributed worldwide causing yield losses exceeding 30% in barley (6) and ~25% in wheat (7) as well as lower grain quality (8). Disease resistance is the best and most sustainable strategy of managing spot blotch, thus understanding and managing resistance sources is critical.

In the Upper Midwestern United States, spot blotch in barley has been effectively managed for more than 50 years through the deployment of resistant six-rowed cultivars (9–11). This seedling resistance in the cultivar (cv) Morex was conferred predominately by the *Rcs5* gene (now referred to as *rcs5*) (11), which also imparts varying levels of adult plant resistance (9,11). An association genetics study on a large breeding panel provided a comprehensive assessment of the genetic architecture of this durable spot blotch resistance, identifying three quantitative trait loci (QTL): *Rcs-qtl-1H-11_10764*, *Rcs-qtl-3H-11_10565* and *Rcs-qtl-7H-11_20162*. Of these QTL, *Rcs-qtl-7H-11_20162* conferred the largest allelic effect across different populations (10) with several genetic studies positioning *rcs5* within the interval of this chromosome 7H QTL (9–11). Thus, the gene/s underlying the *rcs5* QTL is an important target for cloning and functional characterization, which was the objective of this study.

Millions of years of host-parasite interactions evolved diverse multi-layered plant immunity mechanisms with enough commonality in spatial and temporal pathogen detection that they have been placed, somewhat arbitrarily, into dichotomous levels. The first level of early-induced resistance responses typically rely on perception of conserved pathogen or more appropriately microbe associated molecular patterns (PAMPs or MAMPs) by transmembrane cell-surface immunity receptors. These receptors were designated pattern recognition receptors (PRRs) with either receptor-like kinase (RLK) or receptor-like protein (RLP) domain structure (12). The RLKs and RLPs have extracellular “receptor” domains and a single transmembrane domain, but RLKs also have an intracellular serine / threonine protein kinase (PK) signalling domain (12). The RLPs interact with RLKs or intracellular PKs to form receptor complexes that recognize extracellular MAMPs, which elicits cytoplasmic PAMP/MAMP-triggered immunity (PTI) signalling (13). The PTI signalling result in transcriptional reprogramming, synthesis and trafficking of defense metabolites, callose deposition, H_2_O_2_ burst, and in some instances a low amplitude PCD response (13,14).

The best characterized PTI resistance against fungi involves the perception of the major fungal MAMP, chitin, by the rice and *Arabidopsis* PRR complexes that contain homo- and heteroduplexes of RLKs and RLPs with chitin binding LysM ectodomains (15). The rice RLK/RLP heterocomplex containing OsCERK1 and OsCEBiP provides broad non-host resistance to fungal pathogens (16). The Arabidopsis AtCERK1 can act alone to bind chitin to elicit fungal defenses but may also form signalling complexes with other chitin binding RLKs like AtLYK5 (17). CERK1 orthologs in other species also play a role as a co-receptors in complexes that recognize divers MAMPs (17). The rice OsSERK1 RLK also functions in increase resistance to the blast fungus and overexpression of OsSERK1 results in PCD manifested as a disease lesion mimic mutant (18). Our *in silico* protein-protein interaction analysis reported here suggest that barley CERK1 and SERK1 orthalogs may also be a co-receptor or play a role in *Rcs5*-mediated susceptibility.

The pressure imposed by the PTI non-host resistance mechanisms resulted in the adaptation of pathogen specificity by evolving secreted effectors that suppress PTI signalling, resulting in effector triggered susceptibility (19). The second layer of the plant innate immunity system evolved to recognize pathogen effectors directly or indirectly--typically by cytosolic nucleotide binding-leucine rich repeat (NLR) resistance (R) protein receptors (19). Effector perception by NLRs activates effector triggered immunity (ETI) responses (19), typically characterized by a higher amplitude PCD called the hypersensitive response (HR) (20) that are detrimental to the biotrophic pathogen’s feeding structures (i.e. haustoria) that rely on living host cells to extract nutrient.

Research on necrotrophic pathosystems revealed that they produce small secreted necrotrophic effector (NE) proteins to induce immunity responses that result in PCD so the pathogens can colonize, feed, and complete their life cycles on the dead tissue effectively hijacking the plants immune system to cause disease (21). The identification of the host NE-target proteins showed that some fall into the NLR class of resistance (R-) genes, which elicit ETI-mediated HR responses (21,22). Characterization of the *Parastagonospora nodorum* NE SnToxA and its cognate susceptibility target *Tsn1* in wheat, determined that this NLR once triggered by SnToxA activated ETI-like HR responses (21). Thus, necrotrophic specialists elicit responses that evolved to provide immunity against biotrophs to facilitate colonization and further disease development via necrotrophic effector triggered susceptibility (NETS) (23).

The wall-associated kinases (WAKs) are plant RLKs with conserved protein domain architecture including a predicted N-terminal extracellular cell wall binding region with a wall-associated cysteine-rich galacturonan-binding (GUB-WAK), an Epidermal Growth Factor-Calcium binding domain (EGF_Ca^2+^) followed by a transmembrane domain (TM) and a C-terminal intracellular localized serine/threonine protein kinase (PK) signalling domain. The WAK proteins function in cell elongation and development, sensing abiotic stresses, and provide resistance to biotrophic pathogens (24,25). This broad functionality is due to their ability to perceive stresses at the cell-wall/plasma membrane interface which includes degradation induced by pathogen cell wall degrading enzymes (26,27).

Damage associated molecular patterns (DAMPs) are plant extracellular matrix components that are inappropriately released from compromised plant cell walls as a result of pathogen-induced degradation or damage. These oligo-galacturonide (OG) cell wall subunits can be detected as “compromised self” by the WAKs (13). The WAK extracellular “receptor” domains bind long polymers of cross-linked pectin in the cell wall as well as soluble OG pectin fragments that are inappropriately released from the cell wall during pathogen ingress. This detection of OG DAMPs by the Arabidopsis WAK1 receptor elicits PTI-like defense signalling (28,29). Both PAMP and DAMP recognition by PRRs activate intracellular defense signalling via the MAPK pathways (30,31).

The WAK class of RLKs have been implicated as dominant resistance genes that follow Flor’s gene-for-gene model (32) against biotrophic fungal pathogens. The maize genes *ZmWAK* and *Htn1* encode WAK proteins that confer resistance to *Sporisorium reilianum,* the causal agent of the disease head smut (33,34), and the hemibiotrophic pathogen *Exserohilum turcicum,* the causal agent of the disease northern corn leaf blight (35). A third WAK R-gene, designated *Stb6*, from wheat confers resistance to *Zymoseptoria tritici* isolates carrying the corresponding avirulence gene *AvrStb6* (36).

The wheat WAK gene *Snn1* has been shown to be targeted by the necrotrophic fungal pathogen *Parastagonospora nodorum* in an inverse gene-for-gene manner (37). The *Snn1* protein functions as a dominant susceptibility factor and its cognate NE, designated SnTox1, was identified and shown to directly interact with the *Snn1* WAK receptor (37). Wheat varieties carrying *Snn1* recognize *P. nodorum* isolates that produce SnTox1 eliciting PCD, which facilitates colonization and completion of its lifecycle on the resulting dead tissue. The SnTox1/Snn1 direct interaction suggests that *P. nodorum* evolved to activate *Snn1*-mediated signalling by interacting with its extracellular domain rather than the expected perception of DAMPs or OGs.

Here, we report the cloning and characterization of the *rcs5* gene/s and show that *rcs5* has a recessive resistance nature or more appropriately represents dominant susceptibility conferred by two tightly linked wall-associated kinase (WAK) proteins designated *Sbs1* and *Sbs2*. The *Sbs1* and *Sbs*2 proteins function as susceptibility targets and are specifically up regulated by virulent isolates to induce PCD facilitating disease development in the susceptible barley cultivars. Thus, *Sbs1* and *Sbs2* alleles that are either non-functional or resist induction by the pathogen provide *rcs5*-mediated resistance when present in the homozygous state.

## RESULTS

### *rcs5* recessive resistance/dominant susceptibility

Fifteen Morex (resistant) x Steptoe (susceptible) F_1_ progeny were phenotyped using the rating scale developed by Fetch and Steffenson (38) and all the F_1_ individuals assayed demonstrated an intermediate reaction of moderate resistance to moderate susceptibility to *B. sorokiniana* isolate ND85F. The F_1_s exhibited infection types (ITs) ranging from 4.5 to 5.5 with an average of 5. The evaluation of 120 Morex x Steptoe F_2_ progeny resulted in 4 resistant (scores of 1 - 3), 25 moderately resistant (scores of 3.1 - 5), 24 moderately susceptible (scores of 5.1-7) and 67 susceptible progeny (scores of 7.1 - 9). Using a cut-off score of 5 and lower as resistant and 5.1 and higher as susceptible, the progeny fit a segregation ratio of 1 resistant: 3 susceptible (χ^2^=0.0444) at p-value 0.8330 (Supp. Fig. 1) showing *rcs5* recessive resistance.

### *rcs5* genetic mapping

To resolve the *rcs5* region 1,536 Steptoe x Morex F_2_ progeny, representing 3,072 recombinant gametes, were genotyped using the cMWG773 CAPS and BF627428 markers identifying twenty-five recombinant individuals within the previously delimited ~ 2.8 cM *rcs5* interval (39). The twenty-five critical recombinant lines were allowed to self and F_2:3_ individuals were genotyped and progeny selected that contained homozygous recombinant gametes. These immortal critical recombinants were further saturated using SNP markers within the region identifying seven recombinants that delimit *rcs5*. The high-resolution mapping showed markers Sbs1 STS, Sbs2 SNP, 790 SNP, ctg_1606998_STS, 800 SNP, and BF_256735 co-segregating with *rcs5* and the SNP markers BF257002 and HvWak1 SNP delimited the gene distally and 11_20162 delimited the gene proximally (Fig. 1A).

**Figure 1.**
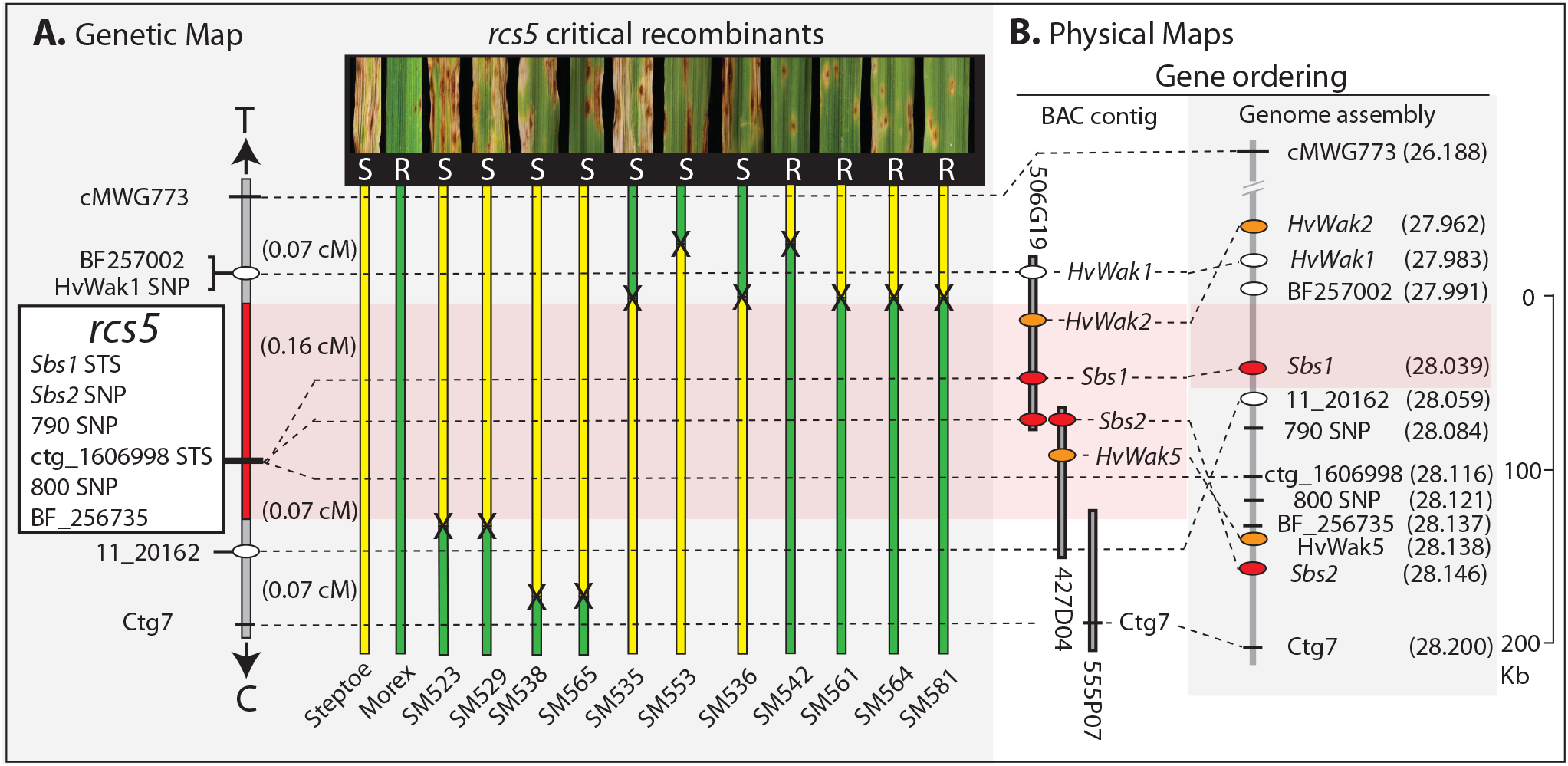
High-resolution genetic and physical mapping of the *rcs5* interval. **1A)** The vertical gray and red bar on the left depicts the high-resolution genetic map with the molecular markers labeled on the left. The white ovals indicate positions of the *rcs5* proximal and distal flanking markers and the white box contains *rcs5* cosegregating markers. The red bar shows the delimited *rcs5* region with approximate cM distances shown to the right. The arrows indicate the direction of telomere (T) and centromere (C). The yellow vertical bars represent cultivar (cv) Steptoe genotypes, whereas the green bars represent cv Morex genotypes with the black Xs representing the crossover points in the critical recombinants. The vertical dashed lines show the position of each genetic marker. The phenotypic reactions of the recombinants and parents are shown above (S = susceptible and R = resistant). The parents and recombinant nomenclatures are provided below. **1B)** Two physical maps are provided with the one on the left produced by cv Morex bacterial artificial chromosome (BAC) contig restriction mapping and sequencing and the one on the right mined from the cv Morex barley genome assembly. Vertical gray bars with black outlines represent the three cv Morex BACs representing the *rcs5* minimum-tilling path with BAC nomenclature provided. The black horizontal bars represent approximate positions of molecular markers and the colored ovals represent candidate genes. The red ovals are validated *rcs5* wall associated kinase (WAK) genes and orange ovals are WAK genes eliminated as *rcs5* candidates. The red shading depicts the delimited *rcs5* regions. For the genome physical map the vertical gray bar represents the genome sequence assembly with Mb positions provided for each gene and marker in parentheses with scale in kilobases provided on the right.

### *rcs5* physical mapping, sequencing and gene prediction

The *rcs5* flanking markers HvWak1_SNP and 11_20162 delimited *rcs5* to an ~0.23 cM genetic interval (Fig. 1A) that was present within a single BAC contig represented by three overlapping resistant cv Morex bacterial artificial chromosome (BAC) clones (Fig. 1B) (40). The sequence annotation of the three overlapping BAC clones (506G19, 427D04 and 555P07) identified five-candidate genes predicted to encode four wall-associated-kinases (WAKs); HvWak2, *Sbs1*, *Sbs2*, and HvWak5 (Fig. 1B), and a predicted non-functional truncated leucine rich-repeat (LRR) gene. Comparison of the BAC contig assembled by restriction mapping and sequencing versus the barley genome sequence (41) showed the same gene content within the *rcs5* region, yet had a different gene order (Fig. 1B). A major concern was that the discrepancies in the gene ordering between the BAC contig and genome assembly resulted in different numbers of candidate genes within the delimited region. However, due to the known accuracy of the BAC restriction mapping and the BAC contig gene order having perfect correlation with the high-resolution genetic mapping, more confidence was placed in the BAC contig assembly (Fig. 1B). Contrary to the physical map generated by the whole genome assembly, which delimited *rcs5* to a single candidate gene (*Sbs1*), the BAC contig assembly showed that the *rcs5* candidate genes were actually delimited to the four WAK genes, *HvWak2, Sbs1, Sbs2*, and *HvWak5* (Fig. 1B).

### *Rcs5* candidate gene characterization

The four candidate *rcs5* genes delimited by high-resolution mapping have conserved WAK protein domain architecture. Phylogenetic analysis using the predicted nucleotide sequences of the five WAK genes clustered at the *rcs5* region from the resistant cv Morex and susceptible cv Steptoe showed that they belong to three different clades, with *HvWak1* and *HvWak2* in subclade I, *Sbs1* alone in subclade II, and *Sbs2* and *HvWak5* closely related in subclade III (Supp. Fig. 2).

The *HvWak2* gene is encoded by 2,689 bases of gDNA in both the susceptible cv Steptoe and resistant cv Morex, with the annotated gene designated HORVU7Hr1G020660 in the barley whole genome assembly. The intron/exon structure supported by qPCR and RNAseq data contains three exons and two introns. The 2,181 nucleotide *HvWak2* mRNA is predicted to encode a 726 amino-acid (aa) protein, ~79.86 kDa with the typical WAK protein architecture containing the predicted GUB, EGF_Ca, TM and PK domains. There were no non-synonymous nucleotide polymorphisms between the cv Steptoe and cv Morex alleles; thus, *HvWak2* was not a strong *rcs5* candidate.

The resistant cv Morex *Sbs1* allele is transcribed from 1,069 bases of gDNA predicted to contain two exons and one intron producing an 870 nucleotide mRNA (Supp. Fig. 3A) predicted to encode a 289 aa protein, ~31.79 kDa, with the GUB and TM domains, and a truncated PK domain (Supp. Fig. 3B). The predicted cv Morex *Sbs1* mRNA was supported by RNAseq and cDNA sequencing (Supp. Fig. 3B). Amplicons from gDNA of the 5’ region of the cv Steptoe *Sbs1* allele were sequenced and showed that the cv Morex resistant allele contained a 635 bp deletion in exon1 adjacent to but not containing the exon1-intron1 splice junction, eliminating the EGF_Ca binding domain (Supp. Fig. 3B). The cv Morex *Sbs1* allele also had a two-nucleotide deletion in the second exon of the predicted mRNA at positions 845-846 that caused a frameshift in the coding region and was predicted to produce a truncated PK domain due to presence of an early stop codon. The cv Steptoe *Sbs1* allele was transcribed from 2,462 bases of gDNA that was predicted to produce a 2,052 nucleotide mRNA supported by qPCR and RNAseq data. The annotated mRNA is predicted to encode a 683 aa WAK protein, ~75.13 kDa, with the typical GUB, EGF_Ca, TM and PK domains (Supp. Fig. 3B). The primary sequence polymorphisms between the resistant and susceptible *Sbs1* alleles suggested that it was a strong *rcs5* candidate gene.

*Sbs2* is transcribed from 2,702 bases of gDNA containing four exons and three introns, producing a 2,160 base mRNA (Supp. Fig. 3A) predicted to encode a 719 aa protein, ~79.09 kDa with a predicted signal peptide (SP) and all the typical GUB, EGF_Ca, TM and PK WAK domains (Supp Fig. 3B). The predicted Morex allele also encodes a 719 aa protein that is polymorphic at the N-terminal coding region as compared to the cv Steptoe allele with 28 synonymous SNPs and 16 non-synonymous SNPs suggesting that it was also a strong candidate *rcs5* gene (Supp. Fig. 3B).

The *HvWak5* gene is transcribed from 2,134 bases of genomic sequence and contains three exons and two introns, producing a 1,731 nucleotide mRNA predicted to encode a 576 aa protein, ~63.36 kDa. The predicted protein contains the typical EGF_Ca, TM and PK WAK domains, but is missing the GUB domain. The predicted HvWak5 protein has identical primary aa sequence for both cv Morex and cv Steptoe alleles suggesting that it was not a strong *rcs5* candidate.

The predicted LRR gene within the delimited *rcs5* region has a stop codon 780 bp from the predicted start methionine, which resulted in a 260 aa protein containing only LRR repeats. Expression analysis via qPCR of the predicted LRR was conducted for cv Morex and cv Steptoe inoculated with *B. sorokiniana* isolate ND85F from infected leaf tissues collected at 0, 12, 24, 36, 48 and 72 hours post inoculation (hpi). The analysis showed no expression before inoculation or at any time-point during the infection process. RNAseq analysis also showed no transcripts present in the cvs Harrington and Steptoe inoculated with isolate ND85F at 72 hpi, which validated our expression analysis results. The data suggested that the LRR is a pseudogene in both resistant and susceptible cvs eliminating it as an *rcs5* candidate.

### RNAseq validation of predicted gene structures

*HvWak2*, *Sbs1* and *HvWak5* were predicted as high confidence genes, with the HORVU7Hr1G020660, HORVU7Hr1G020740 and HORVU7Hr1G020810 gene nomenclatures, respectively, in the recently released cv Morex genome sequence (41). However, the HORVU7Hr1G020740 gene prediction is not a full-length transcript for *Sbs1*, and *Sbs2* was not predicted as a high or low confidence gene. All four-candidate gene annotations were validated with RNAseq data from the non-inoculated susceptible cv Harrington and both cvs Steptoe and Harrington at 72 hpi with *B. sorokiniana* isolate ND85F. *HvWak2* was expressed pre- and post-inoculation, whereas *Sbs1*, *Sbs2* and *HvWak5* were only expressed at 72 hpi in the susceptible cvs Steptoe and Harrington, corroborating the qPCR transcript analysis described below.

### Validation of *rcs5* candidate genes by Virus Induced Gene Silencing

The *Barley Stripe Mosaic Virus*-virus induced gene silencing (BSMV-VIGS) system (42) was utilized for post-transcriptional gene silencing of the four candidate WAK genes in the resistance cv Morex and the two susceptible cvs Steptoe and Harrington. Controls inoculated with only empty BSMV constructs, FES, or *B. sorokiniana* alone were used, along with the four-candidate gene specific BSMV silencing constructs. The silencing of the candidate genes in the resistant cv Morex did not alter disease phenotypes. The seedlings remained resistant across all the treatments with average infection types not significantly different than the BSMV empty virus and non-virus FES controls, with a mean disease severity of 3 on the 1-9 scale (38) (Fig. 2A and 2D). The susceptible cvs Steptoe and Harrington both remained susceptible when infected with the empty vector control, pSL38.1 and the BSMV:*HvWak2* and BSMV:*HvWak5* silencing constructs, showing average disease scores of 6.6, 7.0, 6.0 and 6.4, 6.1, 6.2, respectively. The disease scores were not significantly different between the *HvWak2* and *HvWak5* silencing construct and the empty virus control treatments on the susceptible cvs Steptoe and Harrington (Fig. 2B-D).

**Figure 2.**
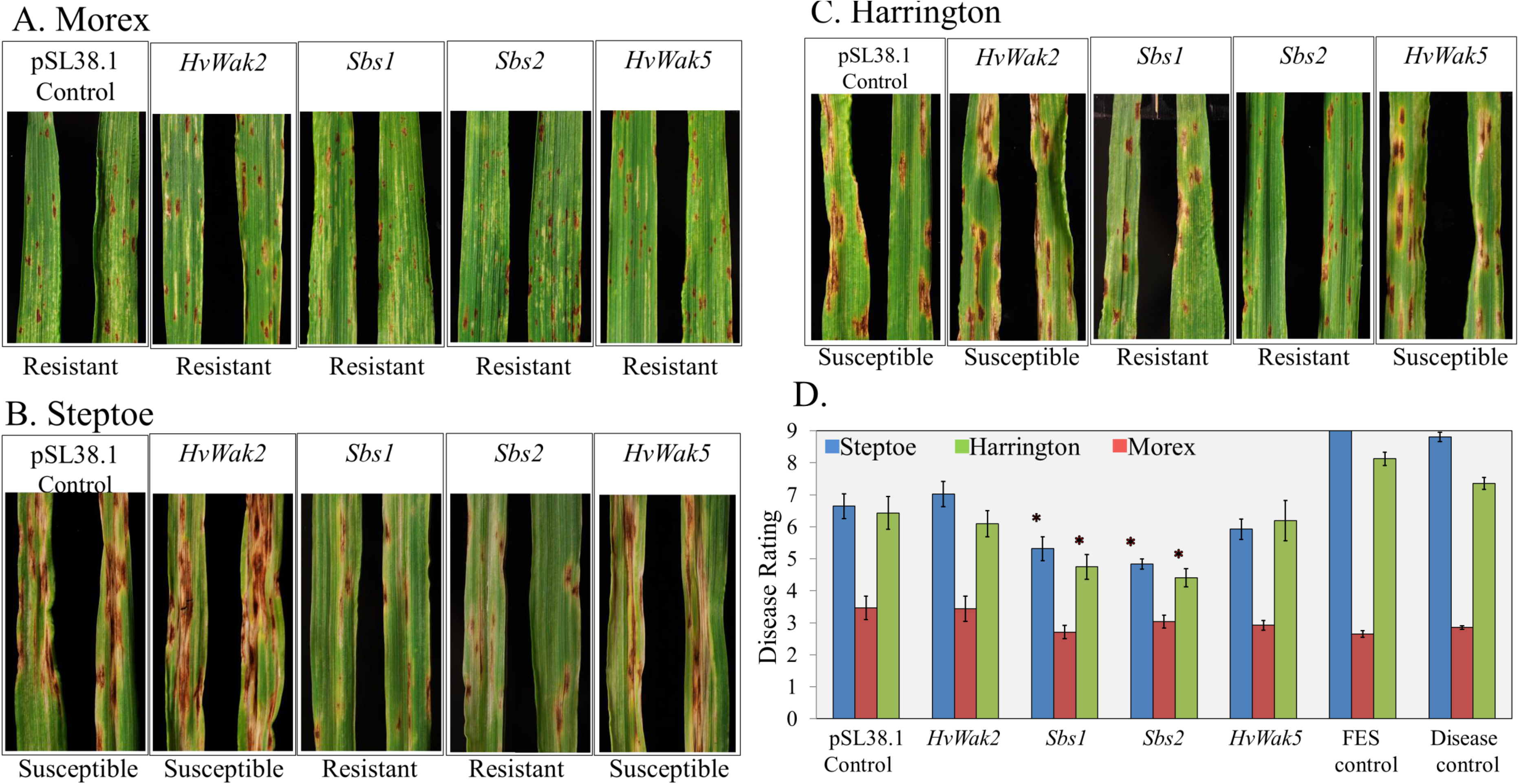
Utilization of BSMV-VIGS for the validation of the *HvWak2, Sbs1, Sbs2 and HvWak5* as *rcs5*. For the functional validation of *HvWak2, Sbs1, Sbs2* and *HvWak5* as a dominant susceptibility gene for spot blotch disease, resistant barley cultivar (cv) Morex **2A)**, and susceptible barley cvs Steptoe **2B)** and Harrington **2C)** analyzed after BSMV-VIGS-mediated post transcriptional gene silencing of the *HvWak2, Sbs1, Sbs2* and *HvWak5* genes, and the empty BSMV-VIGS control (pSL38.1). Pictures show typical reactions for each construct with general reactions labeled below. **2D)** Graphs show disease phenotyping data using a 0-9 disease rating scale (y-axis). The x-axis shows the cvs Steptoe (blue), Harrington (green) and Morex (red). For each genotype n=20 plants. The average disease rating values were calculated with standard error of mean ±1.

The BSMV:*Sbs1* and BSMV:*Sbs2* treated cvs Steptoe and Harrington seedlings had significantly lower spot blotch disease scores than the BSMV empty control, and the BSMV:*HvWak2* and BSMV:*HvWak5* experimental constructs. The BSMV:*Sbs1* treatment had mean disease scores of 5.3 for cv Steptoe and 4.75 for cv Harrington, and BSMV:*Sbs2* had mean disease scores of 4.8 for cv Steptoe and 4.4 for cv Harrington for 20 replicates (Fig. 2D). The shift from compatibility (susceptibility) towards incompatibility (resistance) in the susceptible cvs for both BSMV:*Sbs1* and BSMV:*Sbs2* silenced plants resulted in infection types as low as 2.75. This demonstrated that silencing either of the two closely linked *Sbs1* or *Sbs2* dominant susceptibility genes resulted in a significant increase of resistance in both of the susceptible cvs (Fig. 2B-D).

Quantitative PCR (qPCR) analysis was conducted to assess the specific silencing of the targeted genes based on their transcripts levels at the time points when the highest levels of gene expression were observed during *B. sorokiniana* infection. Three biological replicates of each BSMV:*Sbs1*, BSMV:*Sbs2* and BSMV:*HvWak5* treatment and the empty BSMV control were used to validate specific silencing of the targeted candidate genes. The BSMV:*Sbs1* treated seedlings had more than 85% lower *Sbs1* transcripts than were observed in comparison with the BSMV control treatments (Supp. Fig. 4A). Similarly, the BSMV:*Sbs2* treated seedlings had more than 71% lower *Sbs2* transcript levels than were observed in comparison with the BSMV control (Supp. Fig. 4B). The BSMV:*HvWak5* treatment resulted in ~ 72% lower transcript levels of *HvWak5* in comparison with the BSMV control (Supp. Fig. 4C).

### Allele analysis of *Sbs1* and *Sbs2*

For the *Sbs1* and *Sbs2* allele analysis, PCR primers were designed that produce a 4,242 bp amplicon containing the full length predicted *Sbs1* gene and 1,380 bp of the promoter region (Supp. Fig. 5A) and a 4,711 bp (based on the cv Morex allele) *Sbs2* amplicon that contained the full length predicted gene and 2,009 bp of the promoter region (Supp. Fig. 6A). The amplicons were sequenced from two susceptible and eight resistant cvs on an Ion Torrent PGM instrument to ~ 4,000 X coverage of each genotype specific *Sbs1* and *Sbs2* amplicon. Sequence comparisons showed allelic variation of both the *Sbs1* and *Sbs2* genes within the predicted coding and promoter regions that correlated with *rcs5*-mediated resistance or more accurately, *Rcs5*-mediated dominant susceptibility.

Based on the *Sbs1* allele comparative analysis, the resistant cvs were classified in resistant group-1 (RG1) including the cvs Morex, NDB112, Robust, Tradition and Pinnacle and resistant group-2 (RG2) including the cvs Bowman, ND Genesis and TR306 (Supp. 5D and 6D). The two susceptible genotypes were placed into the susceptible group (SG), which included the cvs Steptoe and Harrington. Within the *Sbs1* coding region RG2 shared higher amino acid (aa) sequence identity with the SG (99.4%) than with RG1 (74%) (Supp. Fig. 5C and 5D). Most of the divergence between RG1 and RG2 was contributed by deletions present in RG1 that eliminated the EGF-Ca domain (Supp. Fig. 5C and 5D) and truncated the PK domain (Supp. Fig. 2A). Sequence polymorphisms also included other synonymous and non-synonymous base substitutions (Supp. Fig. 5B-D). The SG and RG2 *Sbs1* alleles encode the full length 683 aa proteins, whereas the RG1 alleles encode the much smaller 289 aa predicted nonfunctional protein (Supp. Fig. 5C and 5D). The Protein Variation Effect Analyzer (PROVEAN) software predicted that the aa deletions from 150-289 and truncated PK in the RG1 allele are deleterious mutation that would result in non-functional proteins; (PROVEAN scores of −420.8 with significant PROVEAN score cutoff was - 2.5) (43). Despite the deletions that result in a predicted non-functional protein in RG1 *Sbs1* alleles, it appeared that the presence of these deletions were not solely determinant of a nonfunctional susceptibility gene because they were not present in the RG2 alleles.

There were multiple SNPs and small INDELS identified in the 1,380 bp of the 5’ untranslated region (UTR) and promoter regions sequenced from the *Sbs1* alleles showing a high level of diversity (Supp. Fig. 5A). This suggested that the differences between RG2 and SG *Sbs1* gene expression could also determine if alleles function in dominant susceptibility/ recessive resistance. However, the comparative analysis across the predicted promoter regions sequenced only identified a single polymorphic base substitution (G50A based on the nucleotide sequence provided) that perfectly correlated with the resistance –vs-susceptible alleles (Supp. Fig. 5A).

The *Sbs2* allele analysis of the ten barley genotypes revealed the same phylogenetic grouping (RG1, RG2 and SG) as that of *Sbs1* (Supp. Fig. 6D). Contrary to *Sbs1*, comparison of the *Sbs2* alleles showed that at the primary aa level RG1 and RG2 alleles were more similar to one another (99%), with RG1 and RG2 alleles having greater divergence compared to SG, 97.7% and 98.4% aa identity, respectively (Supp. Fig. 6C and 6D). Interestingly, the RG2 *Sbs2* allele appeared to be a recombinant allele with the promoter region and 5’ CDS region translating to the first 64 amino acids having high similarity to RG1 and amino acids 167 to 701 having high similarity with the SG (Supp. Fig. 6D).

The predicted protein alignments of *Sbs2* alleles identified 18 aa substitutions among the ten barley genotypes analyzed (Supp. Fig. 6C and 6D) with nine (A16F, I20A, V27T S32T, S35G, M38R R46K, K60N Q64R D227E) that could putatively contribute to resistance as they were common between RG1 and RG2 and different as compared to the SG allele (Supp. Fig. 6C and 6D). Additionally, RG1 and RG2 compared one to another had aa substitutions T127K, Y167F, E369Q, I444T, C697R and P701S (Supp. Fig. 6C and 6D). There was also a unique D536N aa substitution in the kinase domain of the RG2 as compared to RG1 and SG, which was predicted to be deleterious for the function of the *Sbs2* protein according to PROVEAN with a score of −4.6 (cutoff of −2.5) (Supp. Fig. 6D).

Multiple SNPs and small indels were present in the 2009 bp sequence upstream of the *Sbs2* coding region, representing the *Sbs2* 5’UTR and promoter region. The predicted promoter region of the RG2 alleles had a C1483G, G1504A and insertion of C at 1535 compared to RG1 and SG. The SG had A1510G, T1525C, a deletion from 1527-1539 and CC1541TT compared with both RG1 and RG2 (Supp. Fig. 6A). Thus, we speculated that differential transcriptional regulation due to promoter region polymorphism/s in the RG1 and RG2 alleles compared with the SG could contribute to the inability of the pathogen to induce RG1 *Sbs2* and RG2 *Sbs1* and *Sbs2*, a hypothesis that was tested via expression analysis.

### Expression analysis of *HvWak2*, *Sbs1*, *Sbs2*, *HvWak5*, and *HvMapk3*

The polymorphisms in the promoter regions of RG1 and RG2 vs SG *Sbs1* and *Sbs2* alleles could result in differential expression across the infection process, suggesting that the ability of the pathogen to induce expression of the WAKs could also be a determinant of resistance vs susceptibility. The qPCR analyses (Supp. Table 1) determined that *HvWak2* was constitutively expressed in both the susceptible cv Steptoe and the resistant cv Morex across the infection time course analyzed. However, in the susceptible cv Steptoe, *Sbs1* was induced five-fold at 36 hpi and remained more than two-fold upregulated at the later time-points, while in the resistant cv Morex *Sbs1* showed very low to no expression and was not induced post inoculation (Fig. 3A). Similarly, *Sbs2* was specifically upregulated six and seven-fold in susceptible cv Steptoe at 12 and 72 hpi, respectively, and had very low to no expression in resistant cv Morex. Thus, the susceptible alleles of *Sbs1* and *Sbs2* were induced by the pathogen and the resistant alleles were not (Fig. 3B). The *HvWak5* gene was specifically upregulated ~190-fold at 24 hpi in the susceptible cv Steptoe and the expression level steadily increased across the later time-points to over 1000-fold at 72 hpi (Supp. Fig. 7A). However, since *HvWak5* expression was nearly undetectable before inoculation the induced expression levels as determined by the Cq values were relatively similar to that of *Sbs1* and *Sbs2*.

**Figure 3.**
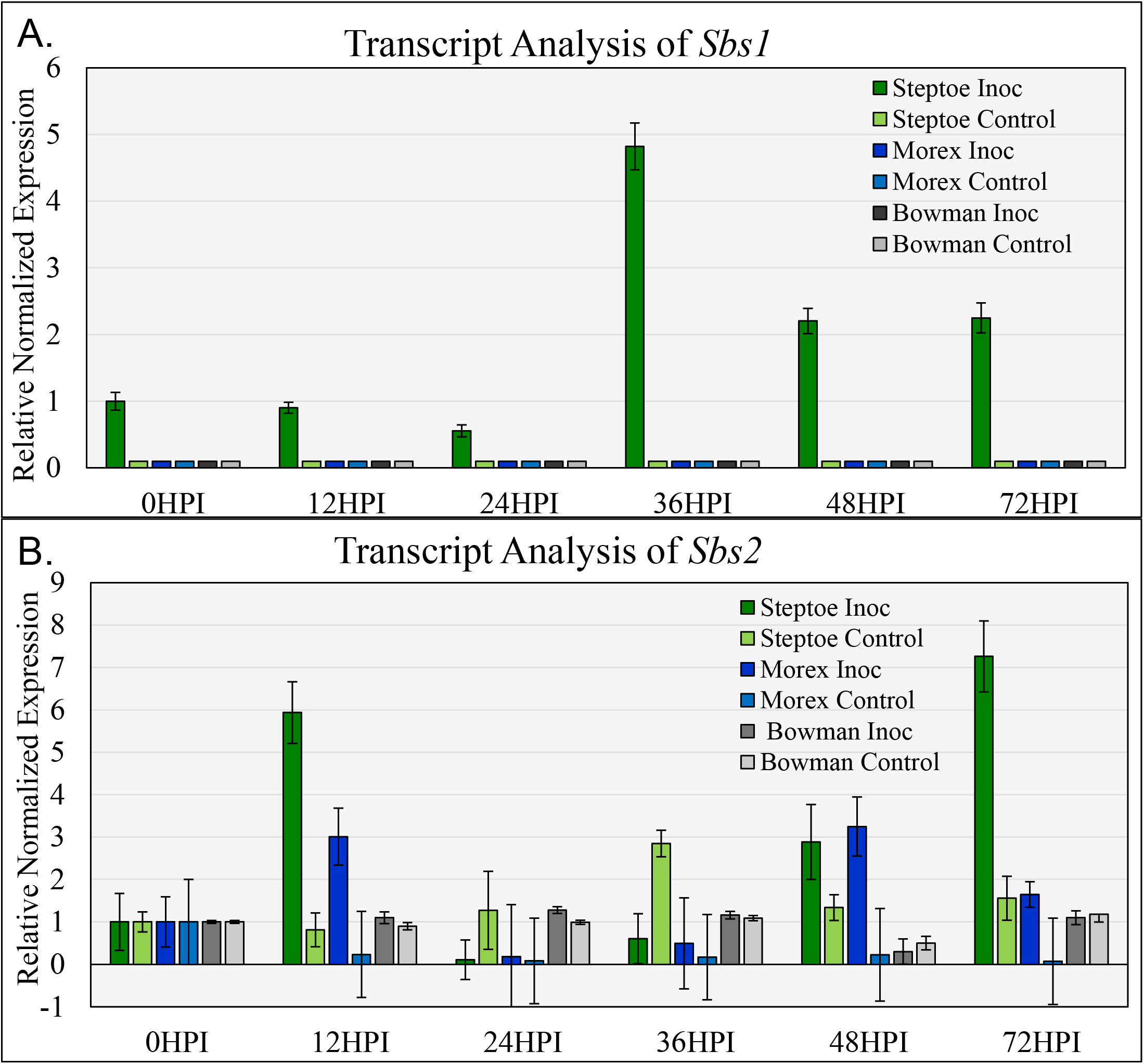
Time course qRT-PCR expression analysis of the **3A)***Sbs1* and **3B)***Sbs2* genes during the infection process with *Bipolaris sorokiniana* isolate ND85F in the barley cultivars (cvs) Steptoe, Morex, and Bowman. Pathogen inoculated susceptible cv Steptoe and resistant cvs Morex and Bowman and tween20 controls were analyzed for *Sbs1* and *Sbs2* expression. *HvSnoR14* expression was used to normalize the transcript data at each time point (X-axis). Error bars depict SEM±1(n=3). The time point 0 HPI was used as control sample for relative expression analysis (Y-axis).

WAK receptors are known to initiate Mitogen-activated protein kinases (MAPK) signalling pathways, activating host defense responses (44), so *HvMapk3* transcript expression was analysed by qPCR through the 0 to 72 hpi time course and it was found that the susceptible cv Steptoe showed approximately two-fold upregulation at 36 hpi and the transcripts remained upregulated at the later time-points during the infection process (Supp. Fig. 7B). However, *HvMapk3* expression was not induced in the resistant cv Morex at any time.

To validate the expression hypothesis, we analyzed *Sbs1* and *Sbs2* expression levels via qPCR in the inoculated and non-inoculated RG2 cv Bowman at 0, 12, 24, 36, 48 and 72 hpi and performed RNAseq at 72 hpi. The qPCR analysis showed very low to zero expression of both candidate *rcs5* genes at time point zero and no differential upregulation of either transcript in the non-inoculated versus inoculated resistant RG2 cv Bowman during the infection process (Fig. 3A and B).

### Transcriptional responses of *Sbs1* and *Sbs2* post intercellular wash fluid infiltration

Based on the post-inoculation transcript data generated for *Sbs1* and *Sbs2*, it was hypothesized that, *B. sorokiniana* triggers the upregulation of the susceptibility genes possibly by a fungal elicitor that is secreted into the apoplast of the host. The intercellular wash fluids (IWFs) collected at 12, 36 and 72 hpi, from cv Steptoe inoculated with the virulent *B. sorokiniana* isolate ND85F induced *Sbs1 ~*20 fold at 72 hours post-infiltration (Supp. Fig. 8A). A similar result was also observed for a second independent replication of the experiment, showing *Sbs1* induction, but in the second replication *Sbs1* upregulation was also observed at 12 and 36 hours post-infiltration as well as at 72 hours post infiltration with the IWFs (Supp. Fig. 8B). In both replications of the infiltration experiments, the *B. sorokiniana* avirulent isolate BS035 IWF did not induce expression of either *Sbs1* or *Sbs2* (Supp. Fig. 8A and B).

To test if the effector that putatively induces *Sbs1* in the IWFs collected from susceptible plants inoculated with the *rcs5* virulent isolate is a protein, the IWFs were treated with pronase prior to infiltration along with a non-pronase treated control. The experiment resulted in no change in upregulation of the *Sbs1* gene, providing evidence of a non-proteinaceous elicitor of the *Sbs1* susceptibility gene (Supp. Fig. 8C).

### DAB (3,3′-Diaminobenzidine) staining

Plants produce reactive oxygen species (ROS), including H_2_O_2_, which precludes PCD/ HR to defend themselves against biotrophic pathogens, however, this defense strategy is exploited by the necrotrophs that purposefully induce the host’s immunity mechanisms to tailor an environment that is conducive to their lifestyle of colonizing dead tissues (45). In the barley-*B. sorokiniana* Pathotype 1 (avirulent on *rcs5*) interaction, we observed leaves at 0, 6, 12, 18, 24, 30, 36 and 48 hpi on the resistant barley cv Morex (*rcs5*+) and the susceptible cv Steptoe (*Rcs5-*). Early and pronounced DAB staining associated with all the *B. sorokiniana* penetration sites as early as 18 hpi were apparent in the inoculated susceptible cv Steptoe. The DAB staining was not observed in the resistant cv Morex at the early time-points showing that the early ROS detection correlated with the timing of *Sbs2* upregulation during infection of the susceptible cv Steptoe (Supp. Fig. 9). During the later time-points (24-48 hpi) the DAB staining associated with penetration and colonization in the susceptible cv Steptoe, rapidly increased in the neighboring cells including the underlying mesophyll cells. However, in resistant cv Morex, the DAB staining was not observed until 24 hpi but appeared to have a higher intensity and remained limited to just a few cells adjacent to the penetration site with little expansion at the later time-points (Supp. Fig. 9).

### Transcriptome analysis post *Bipolaris sorokiniana* infection

RNAseq analysis at 72 hpi identified a total of 3,221 differentially expressed genes (DEGs) with greater than a threefold change (1,488 upregulated and 1,733 downregulated) in the susceptible cv Harrington in comparison with non-inoculated controls (Supp. Table 2). For the resistant cv Bowman, which carries RG2 alleles of *Sbs1* and *Sbs2*, 1,532 DEGs (923 upregulated and 609 downregulated) were identified (Supp. Table 3). Comparison of the upregulated gene list from cv Harrington and cv Bowman, showed that 634 genes were common, 845 Harrington-specific and 270 Bowman-specific DEGs (Supp. Table 5-9). A comparison of the downregulated genes between the susceptible and resistant genotypes showed 376 common DEGs, with 1,357 cv Harrington-specific and 233 cv Bowman-specific DEGs (Supp. Fig. 10A, B, C and D, Supp. Table 4).

The gene ontology (GO) enrichment analysis showed that *Sbs1* and *Sbs2* fall into the biotic stress related category and consistent with the qPCR data were upregulated specifically in the susceptible cv Harrington and not upregulated in cv Bowman (RG2) in response to *B. sorokiniana* infection (Supp. Fig. 10A). The GO analysis showed significant upregulation of the redox category of genes in cv Harrington suggesting that the upregulation of the genes responsible for the oxidative burst by the reactive oxygen species (ROS) an early PTI and ETI response (14) may be mediated through the WAK gene signaling pathways (Supp. Table 10).

Interestingly, nine genes that were upregulated in the susceptible cv Harrington and downregulated in the resistant cv Bowman at 72 hpi (Supp. Fig. 10B and Supp. Table 11) fell into the nitrogen compound/ steroid/ lipid metabolic and amino acid catabolic process and cell growth and morphogenesis classes using the GO biological process classification (PANTHER) (46). Using protein-protein interactions from the STRING database analysing 500 of the DEGs in the susceptible cv Harrington interaction (the top 250 upregulated and downregulated genes) at 72 hpi compared with the non-inoculated control resulted in an interaction network with 893 edges and 361 nodes connecting the differentially regulated genes. Interestingly, all the *Arabidopsis* WAK homologs from the *rcs5* QTL had edges that connected to an *Arabidopsis* chloroplastic gene AKHSDH2 (AT4G19710.2) homologous to the barley HORVU5Hr1G053950.13 gene, effectively representing an interaction hub protein (Supp. Fig. 11A) with aspartokinase/homoserine dehydrogenase activity that catalyzes the synthesis of essential amino acids.

The analysis using STRING data was unable to take into account ‘non differentially regulated genes’ that may be important for connecting the DEGs and making a more reliable interaction network. To mitigate this problem, we developed a Protein-Protein Interaction Network (PPIN) pipeline (submitted in https://github.com/Gazala-Ameen/PPIN). The PPIN analysis with the same 500 barley DEGs in the spot blotch susceptible cv Harrington (top 250 upregulated and downregulated genes) at 72 hpi compared with the non-inoculated control resulted in 361 unique Arabidopsis homolog IDs, which were used for network analysis along with the Arabidopsis homologs of the barley WAK genes that underlie the *rcs5* QTL. Using our PPIN analysis codes with *AtWak3* as an interaction focal point on the same dataset, we generated a functional protein network of 31,125 possible interaction edges (Supp. Fig. 11B; Supp. Table 12).

## DISCUSSION

High-resolution mapping delimited the *rcs5* spot blotch resistance QTL to an ~234 kb physical region of barley chromosome 7H. Four *rcs5* candidate WAK genes were identified within the region. Allele sequencing and VIGS confirmed two of the genes, *Sbs1* and *Sbs2*, as spot blotch dominant susceptibility genes in the susceptible cultivars. Allele analysis and transcript analysis across the infection process and apoplast wash fluid infiltration showed that susceptible alleles of *Sbs1* and *Sbs2* are induced by the hemibiotrophic pathogen then hijacked to promote PCD in an inverse gene-for-gene manner to colonize and complete its life cycle on the resulting dead tissues.

Alleles of host susceptibility target genes with polymorphisms that result in loss of interaction with pathogen virulence effectors or NEs in the case of necrotrophic specialists result in inverse gene-for-gene interactions genetically characterized as recessive resistance genes (22,47). Thus, the recessive allele conferring resistance must be present in the homozygous state to eliminate dominant susceptibility. Host populations pressured by pathogens containing these virulence effectors evolve to eliminate the dominant susceptibility targets through gene mutation that block manipulation by the pathogen. Some of these recessive resistances have been successfully deployed in the field including the rice *xa25* and *xa13* genes that are effective against bacterial blight, *eIF4G* effective against the rice tungro spherical virus, *mlo*-based resistance against powdery mildew in barley, and *tsn1* and *snn1* in wheat that are effective against *P. nodorum* (21,37,48–51).

Interestingly, *xa13-*mediated resistance is determined by polymorphism in the OsSweet11 gene promoter region that block *Xanthomonas oryzae* transcriptional activator-like (TAL) effector PthXo1 binding. It was hypothesized that PthXo1 evolved to induce OsSWEET11 upregulation to facilitate nutrient acquisition, thus, is a dominant susceptibility target (52). This induced transcription is similar to the *rcs5* locus as the virulent isolate of *B. sorokiniana* appears to upregulate the dominant susceptibility genes *Sbs1* and *Sbs2*.

The polymorphisms resulting in a putative nonfunctional Sbs1 protein in the resistant cv Morex and differences in expression of *Sbs1* and *Sbs2* between the cv Morex RG1 allele and the cvs Steptoe and Harrington SG alleles suggested that the pathogen induces the expression of the WAK susceptibility targets then hijack the receptors to facilitate colonization and disease development. Thus, the *rcs5*-mediated recessive resistance is conferred by the *Sbs1* and *Sbs2* alleles that have diverged and are no longer functional susceptibility proteins or are driven by regulatory elements that the pathogen can’t induce. The RG2 class of resistant genotypes have *Sbs1* alleles that closely resembled the SG alleles at the primary aa sequence representing natural allelic variation to test the expression hypothesis. The RG2 also contain recombinant-like *Sbs2* alleles that have a promoter region and N-terminal coding region similar to RG1 but a C-terminal coding region more closely related to SG, providing natural allelic variation that is very similar to a promoter swap assay that could be used to test the induction hypothesis. The allele analysis and observations that the RG2 *Sbs1* and *Sbs2* allele are not induced by the virulent isolate of *B. sorokiniana* supports the hypothesis that the pathogen evolved to induce the WAK genes to become virulent. Thus, *Sbs1* and *Sbs2* susceptible alleles have regulatory elements that the pathogen can manipulate and alleles of either one of the *Sbs* genes that block induction or are non-functional results in impaired colonization and disease suppression.

We posit that the Sbs1 and Sbs2 proteins are hijacked to induce increased levels of PCD as was shown by the DAB staining across the infection cycle (Supp. Fig. 9). Therefore, it was an attractive hypothesis that *B. sorokiniana* isolates virulent on *rcs5* secrete effectors in the apoplast that induce *Sbs1* and *Sbs2.* Intercellular wash fluids (IWFs) from virulent and avirulent isolates were used to infiltrate the susceptible cv Steptoe. Interestingly, the IWF from the *Rcs5* virulent *B. sorokiniana* isolate ND85F induced significant upregulation of *Sbs1* post infiltration at all the time points tested, suggesting that it upregulates the dominant susceptibility genes via an apoplastic secreted effector (Supp. Fig. 8A and B). The avirulent isolate IWFs did not induce *Sbs1* or *Sbs2* suggesting that they do not contain the effector.

To test if the effector/s that induced *Sbs1* was proteinaceous, the IWFs were treated with pronase prior to infiltration, which still resulted in upregulation of the *Sbs1* gene, providing preliminary evidence of a non-proteinaceous *B. sorokiniana* elicitor (Supp. Fig. 8C). A previously reported strain specific secondary metabolite (nonribosomal peptide synthetases) was shown to be a virulence factor in the *B. sorokiniana* pathotype 2 isolate ND90-Pr, which is in line with our observation that at least one of the pathotype 1 *rcs5* specific virulence effectors may be non-proteinaceous (53,54).

The *Sbs2* gene was not induced by the IWFs, suggesting that a second effector was not captured or a different concentration of the same effector is required for *Sbs2* induction. The diverse *Sbs1* and *Sbs2* promoter regions, differential timing of upregulation, and differential IWF induction indicates distinct effectors or mechanisms are eliciting the expression of the two *rcs5* susceptibility genes. Also, the necrosis or cell death phenotype was not observed after infiltration with the IWFs, suggesting that the apoplastic localized elicitor that upregulates *Sbs1* does not induce PCD. The PCD response appears to be pathogen dependent suggesting a second elicitor or effector may induce ROS and subsequent PCD responses such as the continuous production of OG DAMPs by pathogen cell wall degrading enzymes (Fig. 4). This could be an essential event for triggering the PCD through the upregulated WAK genes considering that it been shown in other pathosystems that WAK RLKs act as biotrophic resistance genes-eliciting PCD responses after OG perception (29).

**Figure 4.**
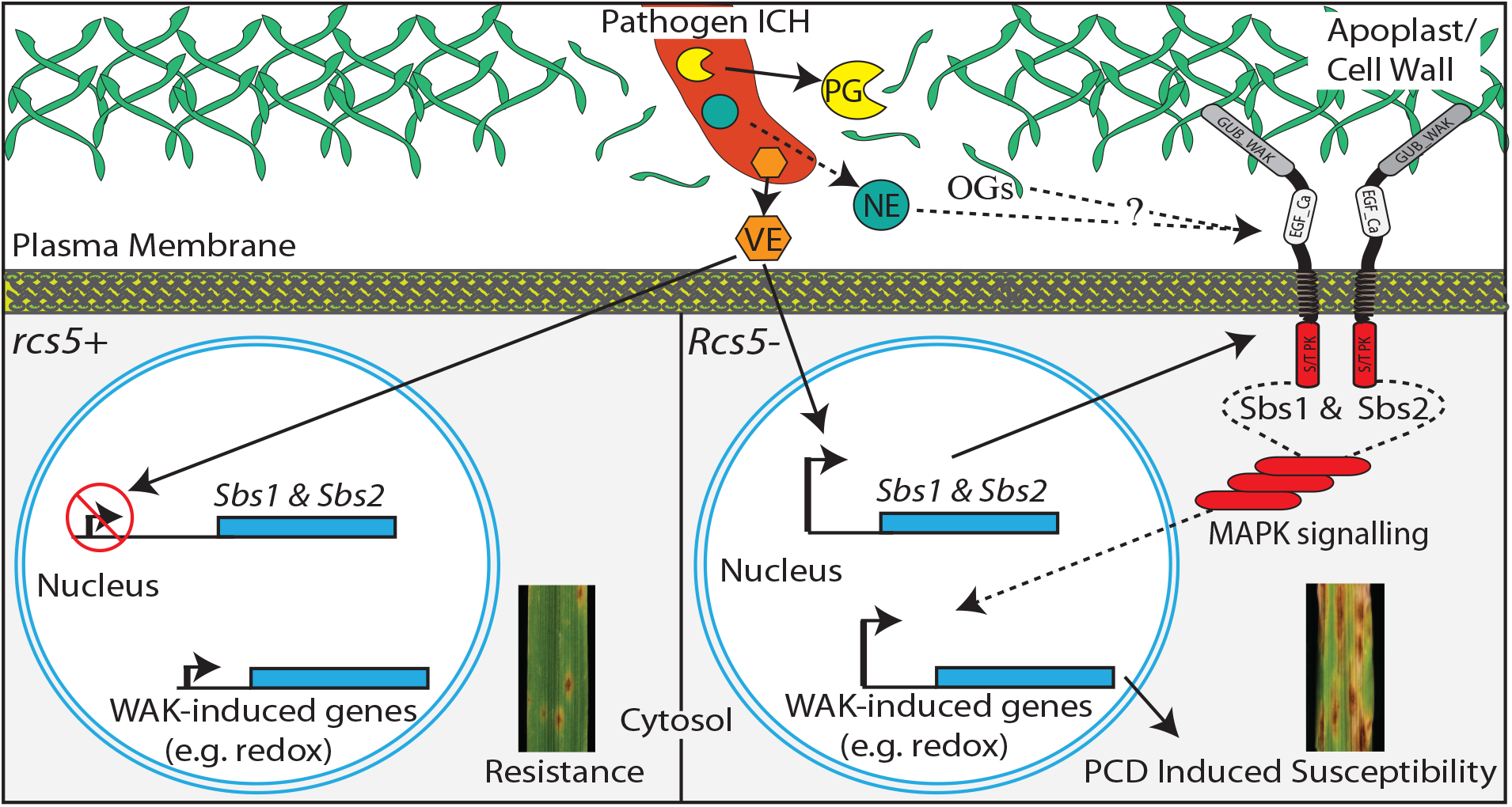
Proposed model of the spot blotch susceptibility due to the *Sbs1* and Sbs2 wall-associated kinases (WAKs) as supported by the genetic, interaction, and signaling data presented for barley– *Bipolaris sorokiniana* pathosystem. The intracellular hyphae (Pathogen ICH) of *B. sorokiniana* enter the host, ~12 hours post inoculation (hpi) and secretes unknown virulence effector/s (VE) and cell-wall degrading enzymes, including Polygalacturonases (PG) into the apoplast. The bottom right box shows the induction of the WAK receptors (susceptibility genes) *Sbs1* and *Sbs2* that are involved in the identification of oligo-galacturonide (OG) damage associated molecular patterns (DAMP) elicitors released during the infection process or by an unknown necrotrophic effector (NE) that intern induce defense responses through MAPK signaling. The defense responses induce overexpression of redox related genes, which induce reactive oxygen species (ROS), programmed cell death (PCD), and favors the necrotrophic pathogen, which hijacks the host defense machinery to induce susceptibility. The bottom left box shows the presence of sbs1 and sbs2 alleles that do not function as susceptibility targets and resist pathogen induction results in spot blotch disease resistance mediated by the *rcs5* locus.

A quantitative mass-spectrometric-based phosphoproteomic analysis identified rapidly induced cellular events in Arabidopsis using OG and Flg22 induction *in planta* identifying seven overlapping induced phosphorylation sites indicating similarity between these two signalling pathways activated by two very distinct DAMP and PAMP recognition RLKs (55). FLS2 recognition of the bacterial flagellin epitope flg22, leads to ROS production and activation of defense related proteins through MAPK signalling (56). We report specific upregulation of *HvMAPK3* in the susceptible cultivars during the infection process in the barley-*B. sorokiniana* pathosystem suggesting that both LRR-PK and WAK RLKs have conserved downstream MAPK defense signalling pathway. Interestingly the RLK CERK1 identified as an interactor with the WAKs using PPIN analysis suggests a role in this conserved signalling pathway, as it is a co receptor that acts in the identification of diverse effectors/PAMPs and subsequent conserved signalling pathways that result in PCD responses.

The upregulation of *Sbs1* and *Sbs2* may amplify the initiation of MAPK signalling, inducing ROS production, activation of defense related proteins and ultimately PCD, similar to defense responses against biotrophic pathogens. Overexpression of *Arabidopsis AtWak1* causes increased OG responses such as ROS production and callose deposition, effectively subduing the growth of biotrophic pathogens (57), but due to the lifecycle of necrotrophs this response facilitates disease development. Overexpression of OsWAK25 in rice resulted in increased susceptibility to the necrotrophic fungal pathogens *Bipolaris oryzae* (*Cochliobolus miyabeanus)* and *Rhizoctonia solani* (58). Interestingly over expression of OsWAK25 in rice also exhibited necrotic spots on leaves. These results are in line with *Sbs1* and *Sbs2* expression data suggesting an increased susceptibility for *B*. *sorokiniana* upon induction of the WAK genes by the virulent isolate.

Perturbation of cell wall structure is monitored by several receptor proteins such as WAKs, Feronia, and RLP44 (59) that initiate specific responses including the accumulation of ROS. Here we report that the barley *Sbs1* and *Sbs2* genes were upregulated at 36 and 12 hpi, respectively, by a virulent isolate of *B. sorokiniana*, which facilitates further disease development in the pathogen’s necrotrophic phase. ROS production is considered as a signal cue for activating cell death responses providing resistance against biotrophic pathogens but increasing the virulence of a necrotroph (60). DAB staining of the susceptible barley cv Steptoe that contains functional *Sbs1* and *Sbs2* alleles and the resistant cv Morex showed a different timeline of H_2_O_2_ production. Early H_2_O_2_ production at 18 HPI was identified in Steptoe corresponding with the early upregulation of the *Sbs2* gene, which we interpret as facilitating the establishment of the pathogen and the later upregulation of *Sbs1* and *Sbs2* correlates with the later expanded ROS production which facilitates further colonization and disease development in the susceptible genotypes. The ROS production identified by DAB in resistant cv Morex was delayed (24 hpi) and localized, representing a limited growth opportunity for the necrotrophic pathogen.

The GO enrichment analysis showed significant upregulation of the redox category of genes in the susceptible cv Harrington suggesting that it is in part the result of the WAK receptor-mediated signaling resulting in PCD considering that the wall localized peroxidases significantly upregulated are a major source of ROS during plant-pathogen interactions (61). In Arabidopsis, it has been shown that the OG hypersensitive mutants tend to have overexpression of peroxidases, which facilitates the constitutive production of ROS and WAKs are major OG receptors. Several studies have also shown that the peroxidase mutants have impaired ROS production and activation of defense responses post PAMP/DAMP treatment and overexpression of the peroxidases triggers the production of ROS (61,62). In the comparative transcriptome analysis, there was significant overexpression of peroxidases in the susceptible transcriptome; for example, the HORVU2Hr1G018510 peroxidase superfamily protein was upregulated 8262 folds in the susceptible cv Harrington and only 14-fold upregulated in the resistant cv Bowman (Supp. table 10). Thus, in the transcriptome of the spot blotch susceptible cultivar overexpression of the peroxidases were accompanied by higher ROS production. We hypothesize that the pathogen induces *Sbs1* and *Sbs2* enriches the cell periphery with more WAK receptors that recognize the degradation of the cell wall, possibly by detecting OGs, which triggers signalling pathways that induce the overexpression of peroxidases and ROS and ultimately PCD (63) that *B. sorokiniana* utilizes to facilitate further disease development (Fig 4).

Comparative RNAseq analysis between cvs Bowman and Harrington infected with *B. sorokiniana* identified DEG’s during the resistance and susceptibility responses. Nine genes were found to be upregulated in susceptible cv Harrington and downregulated in resistant cv Bowman, and of these nine genes *ASK3* and *DIR1* have been shown to be important in plant stress signalling. ASK3 is a cell membrane cytokine receptor (64), that functions as a histidine kinase relaying the stress signal to the downstream MAPK cascade (65). The DIR1 protein is part of a feedback regulatory loop consisting of G3P, DIR1 and AZI1 (66,67) that regulates systemic acquired resistance (SAR) and is critical for glycerolipid biosynthesis (68). Thus, we speculate that the upregulation of these genes controlling plant immune signalling and amino acid catabolism are also playing a role in defense signalling that leads to PCD. Further, STRING protein interaction analysis between DEGs identified the possible hub protein ASKDSK2 that interacts with the *Arabidopsis Sbs1* and *Sbs2* WAK homologs. ASKDSK2 which encodes a bifunctional aspartokinase/homoserine dehydrogenase, catalyzes the synthesis of the essential amino acids threonine, isoleucine and methionine. DMR1, a homolog of HSK, which functions in the same pathway as ASKDSK2, is a downy mildew and *Fusarium* recessive resistance protein (69,70) indicating that this pathway plays an important role in disease resistance signaling. These data also corresponded with the cv Harrington upregulated genes being involved in aa catabolic processes via GO annotation and the fact that they interact with the WAK receptors suggest they may play an important role during the interaction that results in dominant susceptibility.

Interestingly, a KAPP (Kinase Associated Protein Phosphatase-AT5G19280) was also identified as an interaction hub having a direct predicted interaction with three of the upregulated genes, which are predicted to encode a putative S-locus protein kinase (AT1G61420), the chitin receptor kinase CERK1 (AT3G21630) and SERK1 (AT1G71830) which are involved in defense signalling events that result in ROS production and PCD. A downregulated protein kinase like protein (AT1G61590), the homolog of the Arabidopsis RIPK was also found to directly interact with KAAP. Further, a dense string of interactions were identified at the SERK1 node and shown as a group (Supp. Fig. 11B). Interestingly, KAPP encodes for a protein phosphatase 2C 70 protein shown to be present in the cell membrane and is involved in the dephosphorylation of the *At*SERK1 receptor kinase (71). Thus, the PPIN helped to identify genes that may be important in the resistance/susceptibility signalling pathway which are not differentially regulated and yet may represent important interaction hubs for future *rcs5* signalling pathway analyses.

We report that the *Rcs5* dominant susceptibility is conferred by alleles of two wall associated kinase genes, *Sbs1* and *Sbs2*, which are being induced by the necrotrophic pathogen to hijack their function, putatively inducing programmed cell death to facilitate disease development. The presence of the barley WAK *sbs1* and *sbs2* alleles that cannot be manipulated and induced by *Rcs5* virulent isolates of *B. sorokiniana* during the infection process result in the durable *rcs5-*mediated recessive resistance. This pathosystem also represents a novel mechanisms of dominant susceptibility/recessive resistance mediated by the coordinated pathogen induced regulation and function of the two WAK genes. Preliminary characterization of the putative pathogen effector/s lead to the conclusion that it may represent a non-proteinaceous effector, possibly a secondary metabolite that induce *Sbs1* expression and provides a pathogenicity advantage to *B. sorokiniana* isolates that carry it.

## MATERIALS AND METHODS

### Barley Germplasm

The F_2_ Progeny of Steptoe/Morex were used for genetic mapping of *Rcs5*. The spot blotch susceptible and resistant cvs used in this study are Steptoe, Harrington, Morex, NDB112, Robust, Tradition, Pinnacle, Bowman, ND Genesis, and TR306.

### Spot-blotch inoculation and phenotyping

The *B. sorokiniana* isolates used in the experiments were ND85F (pathotype 1, virulent on NDB112 and Bowman differential) and BS035 (pathotype 0, avirulent). The ND85F isolate has been used to screen barley genotypes for spot blotch resistance for over 20 years (10). The BS035 isolate was isolated from *Hordeum jubatum* and determined to be pathotype 0 by screening on the barley differentials and other cultivars (2). The isolates, inoculation procedure and ratings are described in SI.

### High resolution genetic Mapping

High-resolution genetic mapping was conducted utilizing the 1,536 Steptoe/Morex F_2_ progeny (3,072 recombinant gametes), using the flanking markers BF627428 and cMWG773. This reduced the recombinants to 66, which were later screened with BF263248-BF627428 markers reducing the number to 25 recombinants. Finally, the SNP HvWak1-ctg 7 CAPS marker reduced the number of recombinants to 7 (Supp. Table 1). The Physical mapping is described in SI.

### Transcriptional Expression

The cvs. Steptoe and Morex were inoculated with *B. sorokiniana* isolate ND85F along with seedlings inoculated with Tween-20 minus fungal spores as controls. Tissue was collected from three biological replicates of the infected seedlings and controls at 0 and 12, 24, 36, 48 and 72 hpi. The qPCR materials and methods are described in the SI materials and methods section.

### Virus induced gene silencing

The post-transcriptional gene silencing was carried out utilizing *Barley Stripe Mosaic Virus*-Virus Induced Gene Silencing (*BSMV*-VIGS) following the previously described methodology (42). The gene specific BSMV-VIGS construct design are provided in the SI materials and methods section.

### Allele Analysis

*Sbs1* and *Sbs2* allele analysis was performed on ten barley cvs (Morex, NDB112, Robust, Tradition, Pinnacle, Bowman, ND Genesis, TR306, Steptoe and Harrington) to identify polymorphisms that correlate with spot blotch resistance or susceptibility. DNA was isolated using the protocol described by Tsilo et al., 2010 (72). The *Sbs1* and *Sbs2* sequences available from the cv. Morex whole genome sequence assembly and cv. Bowman draft genome assembly (41) as well as the Steptoe sequence generated by Genome Walking (described in SI), were utilized to design primers from conserved regions targeted to amplifying ~4.8kb genomic DNA covering the promoter and coding regions. The material and methods are described in the SI.

### RNAseq experiment

Susceptible cv. Harrington and resistant cv. Bowman was used to perform the gene expression profile for spot blotch resistance and susceptibility (BioProject ID PRJNA522759). Leaf tissue of 3 biological replicates of cv Bowman and cv Harrington non-inoculated control and 72hpi with *B. sorokiniana* isolate ND85F were collected. An RNAseq library was constructed and run on the Illumina NextSeq 500. The detailed materials and methods used are described in the SI.

### Network Analysis

Protein-protein interaction network analysis using the top 250 up and downregulated genes (500 DEGs) identified in the susceptible barley cv. Harrington 72hpi with *B. sorokiniana* was carried out based on interaction data from STRING-db (73) and with protein-protein interaction network (PPIN) analysis and visualization program developed with a custom python script (74), based on networkX (75). Additionally, the Arabidopsis homolog of *Sbs1*, *Sbs2* and other candidate WAKs were included. Our PPIN script (submitted in https://github.com/Gazala-Ameen/PPIN) and details are present in the SI.

### DAB staining

For visualization of ROS, five 3 cm leaf samples were collected from secondary leaves at 0, 6, 12, 18, 24, 30, 36 and 48 hpi from Morex and Steptoe and immediately transferred in 10 ml freshly prepared 1mg/ml DAB (Sigma Aldrich, MO) solution (pH 3.6) in 15 ml tubes. The detailed method is given in Supp. materials and methods section (76).

### Intercellular Wash Fluid Extraction and Infiltration

The cv Steptoe (200 plants) were inoculated with *B. sorokiniana* isolates ND85F and BS035 (~2000 spores/ml). One-third of the secondary leaves (~3 cm) were collected in 50 ml conical centrifuge tubes after 12, 36 and 72 hpi. The IWF extractions were conducted as described in Liu et al., 2015 (23), and the crude IWF extracted was ~3 ml/time-point/isolate. The IWF of each isolate (3 time-points) were mixed and infiltrated in the secondary leaves of cv Steptoe plants which were collected later for transcripts analysis. The detailed procedure is provided in SI.

