## Supplementary material for "*rcs5-*mediated spot blotch resistance in barley is conferred by wall-associated kinases that resist pathogen manipulation": SI

### SUPPLEMENTARY INFORMATION

#### MATERIALS AND METHODS

##### Barley Germplasm

The $F_2$ Progeny of Steptoe/Morex were used for genetic mapping of *Rcs5*. The spot blotch susceptible cvs used in this study are Steptoe (CIho 15229) and Harrington (SK 76333) and the resistant cvs are Morex (CIho 15773), NDB112 (CIho 11531), Robust (PI 476976), Tradition (PI 612442), Pinnacle (PI 643354), Bowman (PI 483237), ND Genesis (2ND 25276), TR306 (TR306). Plants were grown in growth chambers with 21°C temperature with 16/8 h light/dark photoperiod.

##### Spot-blotch inoculation and phenotyping

The *B. sorokiniana* isolates used were ND85F (pathotype 1, virulent on NDB112 and Bowman differential) and BS035 (pathotype 0, avirulent) in the experiments. The ND85F isolate has been used to screen barley genotypes for spot blotch resistance for over 20 years (1). The BS035 isolate was isolated from *Hordeum jubatum* by us and determined to be pathotype 0 by screening on the barley differentials and other cultivars (2). Both isolates were grown on V8-PDA at 21°C under a 12-hour photoperiod. The conidial suspension of ~2,000 conidia/ml of distilled water with a 10µl/ 100ml of Tween-20 (Polysorbate 20) to facilitate distribution and adherence of conidia was prepared. Mock inoculations were carried out by same concentration of the Tween-20 in distilled water. Inoculated barley plants were kept in humidity chambers with 100% humidity for 24-hours and later kept in the growth chambers at 21°C with 16/8 h light/dark photoperiod until rating.

##### BAC screening, Southern blot, probe hybridization, and physical map construction

BAC clones identified in Drader and Kleinhofs (2010) (2) were used to construct a minimum tiling path of BAC clones that would cover the *rcs5* interval. Individual clones were digested with HindIII and run out on a 1% agarose gel. Southern blotting and hybridization were carried out as previously described (Kleinhofs, Kilian et al. 1993 (3). Approximately 25 ng of cDNA insert was labeled with $^{32}$P dCTP (New England Nuclear, Waltham, MA) using the RTS RadPrime DNA labeling system (Gibco

BRL, Carlsbad, CA) according to the manufacturer's instructions. Membranes were exposed to X-ray film for 1–7 days depending on hybridization intensity using NEN Reflection QF intensifying screens (New England Nuclear, Waltham, MA). Unique markers were used to confirm BAC contigs and to help create a minimum tiling of BAC clones that represent the *rcs5* interval and verified using the BAC physical mapping database (http://phymap.ucdavis.edu/barley/).

**Molecular Marker development**

Primers were designed utilizing the physical map of barley cv. Morex (http://webblast.ipk-gatersleben.de/barley_ibsc/). Amplified products were sequenced from Steptoe and Morex. PCR markers were designed by comparison of Steptoe and Morex alleles and analyzed for Single Nucleotide Polymorphisms (SNP) or indels. Polymorphisms that resulted in the absence of a restriction enzyme site were used for Cleaved Amplified Polymorphic Sequence (CAPS) development or for the design of a PCR marker in case no suitable restriction enzyme site was found (Supplementary Table 1).

**Genome Walking**

High molecular mass genomic DNA was isolated using the modified CTAB method (3). Genome walking was conducted using genome walker universal kit following user guidelines (Genome Walker Universal Kit, Clontech, CA) with DraI, NruI, EcorV, PvuII and StuI restriction enzymes and amplified utilizing the adaptor primers AP1 and gene specific primer CTG35_GSP3'_1, CTG35_GSP5'_1 in both directions for *Sbs1* and CTG37_GSP3'_1 and CTG37_GSP5'_1 for *HvWak5* (Supplementary Table 1). Nested PCR was then performed on the PCR product with AP2 primer and gene specific nested primer CTG35_GSP3'_2, CTG35_GSP5'_2 in both directions for *Sbs1* and CTG37_GSP5'_2 and CTG37_GSP5'_2 for *HvWak5* (Supplementary Table 1). The subsequent walking was done using ctg35_2_GSP3', ctg35_2_GSP5', ctg35_3_GSP3 for *Sbs1* and Ctg37_2_GSP5'_1, Ctg37_2_GSP5'_2 for *HvWak5* gene in both directions following the genome walker universal kit user guidelines (Supplementary Table 1).

**Transcriptional Expression**

The primers used were HvuSnoR14 -F1 and HvuSnoR14 -R1; WAK2Q-F1 and WAK2Q-R1; WAK3Q-F1 and WAK3Q-R1; WAK4Q-F1 and WAK4Q-R1 and W5-QF1 and W5-QR1 for *HvSnoR14, HvWak2*, *Sbs1*, *Sbs2*, and *HvWak5*, respectively (Supp. Table 1). A 20 µl qPCR reaction was prepared using 0.8 µl of 100 nM of each gene specific forward and reverse primer, 4 µl of diluted cDNA template, 10 µl SsoAdvanced SYBR green supermix (BIO-RAD), and 4.4 µl qPCR grade ultrapure water (Ambion, MA) in a hard-shell 96 well plate (BioRad, CA) and sealed with Microseal 'B' PCR plate seal (BioRad, CA). qPCR reactions were denatured for 30 sec at 95°C, followed by 45 cycles of denaturation at 95°C for 15 sec and annealing at primer at 58°C for 60 sec. Then a melt-curve was generated at temperatures from 65°C to 95°C with 0.5°C increment (2-5 sec per step). All representative cDNA fragments for each candidate gene and the control reference gene *HvSnoR14* were sequenced to ensure for targeted amplicon.

**Virus induced gene silencing**

The post-transcriptional gene silencing was carried out utilizing *Barley Stripe Mosaic Virus*-Virus Induced Gene Silencing (*BSMV*-VIGS) following the methodology as described by (4). To design gene specific *BSMV*-VIGS constructs for each *rcs5* candidate gene, with minimal off target silencing, 150-200 bp fragments were identified from the predicted mRNA transcript of each gene that contained the highest unique nucleotide sequence (< 85% identity with other genes in the barley genome and avoiding regions that contain continuous stretches of 20 bp of similarity with the predicted cDNA of the wall-associated-kinases *HvWak2, Sbs1*, *Sbs2* and *HvWak5*. In the end, 181, 157, 156, and 165 bp specific sequences were identified for *HvWak2, Sbs1*, *Sbs2* and *HvWak5*, respectively. These amplicons were generated by PCR, using primers GA-WAK2KD-FP, GA-WAK2KD-RP and GA-WAK2KD-REV-FP, GA-WAK2KD-REV-RP for *HvWak2* GA-WAK3KD-FP, GA-WAK3KD-RP and GA-WAK3KD-REV-FP, GA-WAK3KD-REV-RP for *Sbs1* GA-WAK4KD2-FP,GA-WAK4KD2-RP and GA-WAK4KD2-REV-FP, GA-WAK4KD2-REV-RP for *Sbs2* GA-WAK5KD-FP, GA-WAK5KD-RP1 and GA-WAK5KD-REV-FP, GA-WAK5KD-REV-RP1 for *HvWak5* with Not1 and Pac1 sites in sense and anti-sense directions, from the cultivar Morex and Steptoe cDNA (Supplementary Table 1). The amplicons were then sequenced to confirm similarity in Morex and Steptoe. Once confirmed, these cDNA fragments were cloned into the *BSMV* gamma genome based vector (pSL38.1) using NotI and PacI in both sense and anti-sense directions. RNA was synthesized using the mMessage mMachine T7 transcription kit (Ambion, MA), according to the manufacturer's guidelines for the alpha, beta and gamma genome of *BSMV* and gamma genome construct of each candidate gene. RNA of *BSMV* sense and antisense fragments were rub-inoculated onto 10-day-old seedlings of Morex, Steptoe and Harrington (20 replicates for each) with a carrier, FES (0.1M Glycine, 0.06M $K_2HPO_4$, 1% Bentonite w/v, 1% Celite w/v). *BSMV* only, FES only and *B. sorokiniana* only inoculations were also included as controls. After inoculation plants were kept in humidity chambers with 100% humidity for 24 h and later transferred to the growth chambers with 21°C temperature with 16/8 h light/dark photoperiod. After 14 days, when *BSMV* symptoms were systemic in the entire plant, *B. sorokiniana* isolate ND85F was inoculated as described above. After 6 and 10 dpi, the plants were evaluated as a double blind test for disease reaction on the 0-9 scale as described above.

The disease severity rating were analysed and an LSD test applied at $\alpha$= 0.05 using SAS 9.4 (SAS Institute, Cary NC). The mean disease severity for the 20 replicates were calculated for each cultivar for all of the treatments and a t-test and one-way ANOVA was used to compare the group means at $\alpha$= 0.05. Tissue samples were collected from three biological replicates of each *BSMV*: Sbs1, *BSMV*: Sbs2 and *BSMV*: HvWak5 and the control empty *BSMV* treatments at 12, 36 and 72 hpi with *B. sorokiniana* for RNA extraction and RT-PCR was conducted using primers WAK2Q-F1and WAK2Q-R1; WAK3Q-F1 and WAK3Q-R1; WAK4Q-F1and WAK4Q-R1; W5-QF1 and W5-QR1 for *HvWak2*, *Sbs1*, *Sbs2* and *HvWak5,* respectively (Supplementary Table 1). For quantification of the transcripts post inoculation BSMV-VIGS, A 20 μl qPCR reaction was prepared using 0.8 μl of 100 nM of each gene specific forward and reverse primer, 4 μl of diluted cDNA template, 10 μl Sso Advanced supermix, and 4.4 μl qPCR grade ultrapure water (Ambion®) in a hard-shell 96 well plate (BioRad) and sealed with Microseal 'B' PCR plate seal (BioRad). qPCR reactions was denaturation for 30 sec at 95°C, followed by 45 cycles of denaturation at 95°C for 15 sec and annealing at 58 °C for 60 sec. Then, a melt-curve was generated for temperatures from 65°C to 95°C with 0.5°C increment (2-5 sec per step). All the samples were normalized with the reference *SnoR14* transcript using primers HvuSnoR14 -F1 and HvuSnoR14 -R1and same qPCR conditions as described above.

**Allele Analysis**

The Allele analysis were carried out by amplifying the genomic fragments with primers Sbs1_F4 and Sbs1_PRO_R1 for *Sbs1* and Sbs2_F3 and Sbs2_R5 for *Sbs2* (Supplementary Table 1). The PCR reactions contained 2 µl (~20ng/ µl) of template gDNA, 200 µM dNTPs, 0.5 µM of each forward and reverse primer, 5 µl of 5X Q5 reaction buffer, 5 µl of 5X Q5 high GC enhancer, 0.02 unit/ µl final concentration of the Q5 high fidelity DNA polymerase and nuclease free water up to a 25 µl reaction volume. The PCR cycle parameters had an initial denaturation of 95°C for 5 min, followed by 35 cycles of 95°C for 60 sec, 60°C for 180 sec, and 72°C for 60 sec followed by a final extension of 72°C for 10 min. PCR products were visualized on a 1% agarose gel containing GelRed (Biotium, CA) and subsequently purified using E.Z.N.A. Cycle Pure Kit (Omega Bio-Tek, Norcross, GA) following the manufacturer's standard protocol. The PCR amplicons were used to create barcoded Ion Torrent sequencing libraries using the NEBNext Fast DNA fragmentation and Library Prep Set for Ion Torrent following the manufacturer's standard protocol (Life Technologies, CA). Approximately 3 pg of the final multiplexed barcoded library was sequenced on the Ion Torrent PGM instrument utilizing the Ion PGM Template OT2 200 kit, Ion PGM Sequencing 200 Kit v2 and an Ion 314 Chip (Life Technologies), following the manufacturer's standard protocols. Barcoded libraries of each allele PCR fragment was sequenced resulting in a mean read length of 223 and an average of 240,000 reads per sample. The data analysis was conducted with CLC Genomics workbench 8.0.3 software (QIAGEN Bioinformatics, CA). Trimmed reads were aligned to a subset of reference file containing *Sbs1* (4347 bp) and *Sbs2* (4773 bp) genomic DNA sequence. 90-99 % read alignment was achieved for each genotype using our defined parameters of mismatch cost of 2, Insertion/ Deletion cost of 3, Insertion/ Deletion open cost of 6, Insertion/Deletion extend cost of 1, length fraction = 0.5 and similarity fraction = 0.8. Consensus sequence was extracted using aligned read to either *HvSbs1* or *HvSbs2* with minimum of 300 reads per base pair and PHRED quality score >30 for each base call. *HvSbs1* and *HvSbs2* consensus sequence was used for *in silico* analysis of promoter region on FGENESH (5).

**RNAseq experiment**

Susceptible cv. Harrington and resistant cv. Bowman was used to perform the gene expression profile for spot blotch resistance and susceptibility (BioProject ID PRJNA522759). Leaf tissue of three biological replicates of Bowman (BioSample Acc SAMN10953628, SAMN10953629, SAMN10953630 as non-inoculated control and SAMN10953631, SAMN10953632 and SAMN10953633 were 72hpi) Harrington (BioSample Acc SAMN10953637, SAMN10953638, SAMN10953639 as non-inoculated control and SAMN10953634, SAMN10953635 and SAMN10953636 were 72hpi that were inoculated with *B. sorokiniana* isolate ND85F were collected at 72 hours post-inoculation. An RNAseq library was constructed using TruSeq RNA library prep kit v2 (Illumina) and run on the Illumina NextSeq 500. The total reads obtained are summarized in Fig. 8C. Quality of reads were checked in FastQC v0.11.5 (6). The data analysis and comparisons were carried out in the CLC Genomics workbench 8.0.3 software (QIAGEN Bioinformatics, CA). The reads of the 12 samples were aligned using barley concatenated reference high confidence and low confidence gene list provided in the barley IBSC IPK 2016 database. Alignment was performed using mismatch cost 2, insertion and deletion cost 3, length and similarity fraction 0.9 and reads were aligned for both strand specificity and maximum number of hits for a read was set at 10. Expression was calculated using RPKM. Empirical analysis of DGE (EDGE) was performed to run "Exact test" for two group comparisons (7). fdr corrected $P< 0.05$ and EDGE $\geq 3$ fold regulation values were used to get the differentially regulated genes between the non-inoculated and inoculated samples for both genotypes to create the final list of differentially regulated transcripts for spot blotch susceptibility and resistance. The final list was then compared between for the specific upregulated and downregulated genes during susceptibility and resistance.

**Network Analysis**

A protein-protein interaction network (PPIN) analysis with the top 250 up and downregulated genes (500 DEGs) identified in the susceptible barley cv. Harrington 72hpi with *B. sorokiniana* was carried out using data from STRING-db (8) and on a PPIN (protein-protein interaction network) analysis and visualization program developed on our own custom python script (9), based on NetworkX (14). Additionally, the Arabidopsis homolog of *Sbs1*, *Sbs2* and other candidate WAKs were included. Using our PPIN script (submitted in https://github.com/Gazala-Ameen/PPIN), candidate WAKs underlying the *Rcs5* QTL were denoted by yellow color, upregulated genes as red and downregulated genes as green. The protein-protein interaction network of *Arabidopsis thaliana* was created based on the interactions database in String-DB v10.5, a curated database (8) combining interactions from different venues with their corresponding probabilities of occurrence (10). We then modeled identifying maximum reliability pathways connecting our up-regulated, down-regulated, and important genes as a Steiner Tree (11) problem on the largest connected component of the obtained PPIN. Our approach is novel as it allows for a selection of multiple genes to connect and identify their pathways and also include the genes which are not provided in the list but are important to connect and built a more reliable network. The Steiner Tree problem is NP-hard (12), and as such it would prove impossible to solve for the 24,265 vertices and 5,318,667 interactions of the largest connected component within the *Arabidopsis thaliana* proteome. Therefore, we solved it with the Kou approximation algorithm (13), using NetworkX (14) for graph processing and visualization purposes in our PPIN script.

**Dab staining**

For visualization of ROS, five 3 cm leaf samples were collected from secondary leaves at 0, 6, 12, 18, 24, 30, 36 and 48 hpi from Morex and Steptoe and immediately transferred in 10 ml freshly prepared 1mg/ml DAB (Sigma Aldrich, MO) solution (pH 3.6) in 15 ml tubes. Plastic mesh on the top of the tube was used to ensure the proper dipping of leaf bits. Samples were DAB stained for 6 hours on an orbital shaker (VWR) at room temperature at 120 rpm shaking. Later DAB solution was removed and samples were washed twice with anhydrous Farmer's fixative (3-ethanol:1- glacial acetic acid) and then cleared and fixed simultaneously in 30 ml Farmers' fixative (FF) for 12 hours transferring in 50 ml tubes.

Farmer's fixative was changed once after 12 hours and samples were cleared for an additional 3 hours. Cleared samples were stored in fresh 45 ml Farmer's fixative solution in the 50 ml tube in dark until visualized under microscopy. To visualize the germination and growth of *B. sorokiniana* conidial spores and their association with DAB staining, we used wide-field florescence on a Zeiss Axio Imager M2 epiflorescence upright microscope. Emission was 435-485, chromatic beam splitter was set at FT420 and excitation filter BP379-401 and Axio Visison rel. 4.8 software was used to visualize pathogen associated DAB.

**Intercellular Wash Fluid Extraction and Infiltration**

The respective *B. sorokiniana* isolates ND85F and BS035 (inoculum concentration of ~2000 spores/ml) were inoculated onto about 200 Steptoe plants as described in the above protocol. Then, about one-third of the secondary leaves (~3 cm long pieces) were collected in 50 ml conical centrifuge tubes after 12, 36 and 72 hpi. The IWF extractions were conducted as described in Liu et al., 2015 (15), and the crude IWF extracted was about ~3 ml from each time-point per isolate. The IWF of the three time-points were mixed and infiltrated into secondary leaves of Steptoe plants which were then returned to the growth chambers with 21°C temperature and 16/8 h light/dark photoperiod until collection of the infiltrated tissue in liquid nitrogen at 12, 36 and 72 hours post-infiltration in three replicates for RNA extraction and subsequently qPCR assay to determine the targeted transcripts. This experiment was run twice. The Pronase treatment was applied to 5 ml of IWF of the ND85F and BS035 isolates, where the final concentration of Pronase was 1mg/ml and kept at room temperature for 4 hours along with the IWF without Pronase treatments and water-only treatments as control (16). These five treatments were then infiltrated into secondary leaves of Steptoe and infiltrated leaf tissues were collected at 72 hours post infiltration in three replicates in liquid nitrogen for subsequent transcript analysis. The primers used were HvuSnoR14 -F1 and HvuSnoR14 -R1; WAK3Q-F1 and WAK3Q-R1; WAK4Q-F1 and WAK4Q-R1 for *HvSnoR14, Sbs1, Sbs2* respectively (Supplementary Table 1). The qPCR reaction is described in supplementary materials and method section.

**DATA AND MATERIALS AVAILABILITY**

The *HvWak2* gene has GenBank Accession (acc) MK182575, *Sbs1* gene has GenBank acc MK182577 of the resistant cv Morex, *Sbs1* gene of the susceptible cv Steptoe has GenBank acc MK182585, the *Sbs2* gene of Morex has GenBank acc MK182595, the *Sbs2* of Steptoe allele has GenBank acc MK182587 and *HvWak5* gene GenBank acc MK182576 for both Morex and Steptoe. The Allele of the *Sbs1* comparative analysis, the resistant cvs were classified in resistant group-1 (RG1) including the cvs Morex, NDB112, Robust, Tradition and Pinnacle (GenBank acc MK182577, MK182578, MK182579, MK182580 and MK182581, respectively) and resistant group-2 (RG2) including the cvs Bowman, ND Genesis and TR306 (GenBank accMK182582, MK182583 and MK182584, respectively). The two susceptible genotypes were placed into the susceptible group (SG), which included the cvs Steptoe and Harrington (GenBank acc MK182585 and MK182586, respectively). The *Sbs2* allele analysis of the ten barley genotypes revealed the same phylogenetic grouping (RG1, RG2 and SG) as that of *Sbs1* with RG1 consisting of cvs Morex, NDB112, Robust, Tradition and Pinnacle (GenBank acc MK182587, MK182590, MK182591, MK182593 and MK182594, respectively) and RG2 including the cvs Bowman, ND Genesis and TR306 (GenBank acc MK182588, MK182589 and MK182592, respectively). The two susceptible cvs Steptoe and Harrington (GenBank acc MK182595 and MK182596, respectively).

The RNAseq transcript were deposited as BioProject ID PRJNA522759 and has three biological replicates of Bowman (BioSample Acc SAMN10953628, SAMN10953629, SAMN10953630 as non-inoculated control and SAMN10953631, SAMN10953632 and SAMN10953633 were 72hpi that were inoculated with *B. sorokiniana* isolate ND85F. The Harrington transcripts were deposited as BioSample Acc SAMN10953637, SAMN10953638, SAMN10953639 as non-inoculated control and SAMN10953634, SAMN10953635 and SAMN10953636 were 72hpi that were inoculated with *B. sorokiniana* isolate ND85F. The Harrington-*B. sorokiniana* susceptibility transcriptome was used on a PPIN (protein-protein interaction network) analysis and visualization program developed on our own custom python script (9), based on NetworkX (14), which can be accessed using our PPIN script (submitted in https://github.com/Gazala-Ameen/PPIN).


#### SUPPLEMENTARY FIGURE LEGENDS

**Supplementary Figure 1.** Frequency distribution for the mean of the spot blotch disease response of *Bipolaris sorokiniana* isolate ND85F as observed on the 120 $F_2$ progeny of the Morex /Steptoe population, along with the corresponding rating scale developed by Steffenson and Fetch.

**Supplementary Figure 2.** Homology of the five spot blotch disease susceptibility candidates Wall-associated Kinases (WAKs) in barley**.**

**Supplementary Figure 3.** The *Sbs1* and *Sbs2* predicted gene and protein structures from spot blotch susceptible and resistant barley cultivars. **3A)** The genomic and cDNA gene structures for *Sbs1* and *Sbs2* with cultivar (cv) Steptoe representing the susceptibility group (SG), and resistant cvs Morex and Bowman representing resistant group 1 (RG1) and resistant group 2 (RG2), respectively. The genomic and predicted mRNA structures are represented to scale with exon (black bars), intron (black Vs), UTR (white bars), inducible promoter (yellow bars), noninducible promoter (green bars) transcription start site (TSS), start codon (ATG), and stop codon (TGA). The position of the TSS, start codon and stop codon are provided reletive to the start codon. The positions of the BSMV-VIGS and qPCR amplicons are represented as red bars labeled BV and qP, respectively. **3B)** The Sbs1 and Sbs2 protein structures are shown for cvs Steptoe (SG),  Morex (RG1)and Bowman (RG2). The white bar represents the full-length predicted proteins with blue, grey, red and green bar represents the GUB (galacturonic acid binding), EGF_Ca (epidermal growth factor –Calcium binding), TM (transmembrane) and STKc_IRAK (Serine/Threonine kinases, Interleukin-1 Receptor Associated Kinases) domains, respectively. The Sbs1 in Steptoe and Bowman encodes a full length protein, however Sbs1 in Morex encodes a protein with missing EGF_Ca domain and a truncated STPKc_IRAK. The Sbs2 in all three cultivars encode a full length wall-associated kinase protein.

**Supplementary Figure 4.** Transcript analysis of **4A)** *Sbs1*, **4B)** *Sbs2* and **4C)** *HvWak5* was analyzed at 36, 12 and 36 hours post inoculation (hpi), respectively, during the infection process of *Bipolaris sorokiniana* in the BSMV:*Sbs1*, BSMV:*Sbs2* and BSMV:*HvWak5* silenced plants of cultivar Steptoe along with the BSMV-VIGS control (n=3). The reference gene *HvSnoR14* expression in each sample at each time point was used to normalize the transcripts (X-axis). Error bars depict SEM±1(n=3). BSMV-VIGS control was used as control sample for relative expression analysis (Y-axis).

**Supplementary Figure 5A.** The Allele analysis of *Sbs1* gene in the eight resistant cultivars, grouped as Group-1 resistant (G1R) and Group-2 resistant (G2R) and two susceptible cultivars (SG) with the promoter and the exon, intron and protein domains represented in the figure. **5B)** The CDS of the *Sbs1* gene in the ten cultivars are represented in the figure along with the domain architecture of the predicted protein. **5C)** The predicted protein structure of the ten barley cultivars along with the domains. **5D)** The SBS1 protein analysis in the eight resistant cultivars, grouped as Group-1 resistant (G1R) and Group-2 resistant (G2R) and two susceptible cultivars (SG). Polymorphism between the G1R, G2R and SG is shown along with the numbers representing the corresponding amino-acid positions, where dots (.) represents same, (-) represents gaps and the alphabets represents the amino acid. The pairwise comparisons between the ten genotypes is represented with GAPS in the x-axis and differences in amino-acid in the y-axis.

**Supplementary Figure 6A.** The Allele analysis of *Sbs2* gene in the eight resistant cultivars, grouped as Group-1 resistant (G1R) and Group-2 resistant (G2R) and two susceptible cultivars (SG) with the promoter region and the exon, intron and protein domains represented in the figure. **6B)** The CDS of the *Sbs2* gene in the ten cultivars are represented in the figure along with the domain architecture of the predicted protein. **6C)** The predicted protein structure of the ten barley cultivars along with the domains. **6D)** The SBS2 protein analysis of the eight resistant cultivars, grouped as Group-1 resistant (G1R) and Group-2 resistant (G2R) and two susceptible cultivars (SG). Polymorphism between the G1R, G2R and SG is shown along with the numbers representing the corresponding amino-acid positions, where dots (.) represents same, (-) represents gaps and the alphabets represents the amino acid. The pairwise comparisons between the ten genotypes is represented with GAPS in the x-axis and differences in amino-acid in the y-axis.

**Supplementary Figure 7.** Time course qRT-PCR transcript analysis of the spot blotch susceptible cultivar Steptoe and resistant cultivar Morex inoculated with the ND85F isolate of *Bipolaris sorokiniana* and their tween control were analyzed for the expression analysis. **7A)** *HvWak5* transcript were analyzed during the spot blotch susceptibility in the barley cultivar Steptoe and Morex. **7B)** *HvMAPK3* was required during the infection process of *Bipolaris sorokiniana* in barley. The reference gene *HvSnoR14* expression at each time point was used to normalize the transcripts (X-axis). Error bars depict SEM±1(n=3). Time point 0 HPI was used as control sample for relative expression analysis (Y-axis).

**Supplementary figure 8A and B.** Two run of the time course qRT-PCR transcript analysis of the *Sbs1* post infiltration of the extracted Intercellular Fluid Wash (IWF) from avirulent isolate BS035 and virulent isolate ND85F of *Bipolaris sorokiniana* inoculated on cultivar Steptoe. The reference gene *HvSnoR14* expression at each time point was used to normalize the transcripts (X-axis). Error bars depict SEM±1(n=3). Time point 0 HPI was used as control sample for relative expression analysis (Y-axis). **8C)** The qRT-PCR transcript analysis of the *Sbs1* post infiltration of the Pronase treated and non-treated extracted Intercellular Fluid Wash (IWF) from avirulent isolate BS035 and virulent isolate ND85F of *Bipolaris sorokiniana* inoculated on cultivar Steptoe. The reference gene *HvSnoR14* expression at each time point was used to normalize the transcripts (X-axis). Error bars depict SEM±1(n=3). Water infiltrated Steptoe was used as control sample for relative expression analysis (Y-axis).

**Supplementary Figure 9.** DAB staining to characterize the *Bipolaris sorokiniana* infection timeline, and ROS production in the barley cultivar Steptoe and Morex. The panels show *B. sorokiniana* isolate ND85F infection on Steptoe and Morex from 6 to 36 hours post inoculation (hpi).

**Supplementary Figure 10.** Differentially expressed genes (DEGs) during the infection process of *Bipolaris sorokiniana* in resistant cultivar Bowman and susceptible cultivar Harrington. **10A)** A graphical representation of the DEGs based on their biological function category in the resistant and susceptibility of spot blotch disease. **10B)** Venn- diagram showing the common and/or unique DEGs in the yellow and red ellipse representing the susceptibility upregulated (Harrington_up) and downregulated (Harrington_Down); and blue and green ellipse representing resistant upregulated (Bowman_up) and downregulated (Bowman_Down) at 72 hpi with *B. sorokiniana* normalized with their non-inoculated controls DEGs (n=3). **10C)** Total reads obtained and mapped to the reference genome of barley in each sample of the DEGs analysis. **10D)** Heat Map represents the Log transformed fold changes of the DEGs in the susceptibility and resistance to spot blotch disease.

**Supplementary Figure 11A.** Protein-protein interaction re-analysis of Arabidopsis homologs of 500 differentially regulated (250 upregulated and 250 downregulated transcripts in Harrington at 72hpi with

*Bipolaris sorokiniana*) genes with STRING database. The green color represents upregulated and red color represents downregulated genes in the susceptible cultivar Harrington during spot blotch susceptibility. Line thickness of network edges represent strength of data support and kmeans network clustering method with three cluster group was specified for grouping. The candidate WAKs homologs are represented in the far left in the network. Arabidopsis gene identifiers on each bubble represents the interacting genes in the network. **11B)** Prediction of protein-protein interaction of 361 Arabidopsis orthologs for 500 differentially regulated (250 upregulated and 250 downregulated genes in susceptible cultivar Harrington at 72 hpi with *Bipolaris sorokiniana*) along with the Arabidopsis WAK homologs of candidate WAKs in the *rcs5* locus. Yellow color represents the Arabidopsis WAK homologs of *Rcs5* region barley WAK genes added to the list for WAK specific interaction network, green color represents up-regulated, red represents down-regulated, and blue represents not differentially regulated genes introduced in the network for making necessary connections by PPIN analysis from Arabidopsis protein database. Arabidopsis gene identifiers on each bubble represents the interacting genes in the network.

**Supplementary Table 1.** The list of primers used in this study along with their target, amplicon size and physical position in the whole genome of Barley.

**Supplementary Table 2.** The differentially expressed genes in the barley susceptible cultivar Harrington at 72 hpi with *Bipolaris sorokiniana* isolate ND85F. The transcripts were normalized with the non-inoculated DEGs in Harrington (n=3).

**Supplementary Table 3.** The differentially expressed genes in the barley resistant cultivar Bowman at 72 hpi with *Bipolaris sorokiniana* isolate ND85F. The transcripts were normalized with the non-inoculated DEGs in Bowman (n=3).

**Supplementary Table 4.** The exclusively upregulated genes in the susceptible cultivar Harrington at 72 hpi with *Bipolaris sorokiniana* isolate ND85F.

**Supplementary Table 5.** The exclusively downregulated genes in the barley susceptible cultivar Harrington at 72 hpi with *Bipolaris sorokiniana* isolate ND85F.

**Supplementary Table 6.** The exclusively upregulated genes in the barley resistant cultivar Bowman at 72 hpi with *Bipolaris sorokiniana* isolate ND85F.

**Supplementary Table 7.** The exclusively downregulated genes in the barley resistant cultivar Bowman at 72 hpi with *Bipolaris sorokiniana* isolate ND85F.

**Supplementary Table 8.** The common genes upregulated in barley susceptible cultivar Harrington and resistant cultivar Bowman at 72 hpi with *Bipolaris sorokiniana* isolate ND85F.

**Supplementary Table 9.** The common genes downregulated in barley susceptible cultivar Harrington and resistant cultivar Bowman at 72 hpi with *Bipolaris sorokiniana* isolate ND85F.

**Supplementary Table 10.** The gene list along with the transcriptional expression (fold changes) of the Peroxidases super family involved in the spot blotch susceptibility in barley compared with their expression in spot blotch resistance.

**Supplementary Table 11.** The unique genes downregulated in barley susceptible cultivar Harrington and upregulated in the resistant cultivar Bowman at 72 hpi with *Bipolaris sorokiniana* isolate ND85F.

**Supplementary Table 12.** The unique genes upregulated in barley susceptible cultivar Harrington and downregulated in the resistant cultivar Bowman at 72 hpi with *Bipolaris sorokiniana* isolate ND85F.

**Supplementary Table 13.** The list of proteins in the Predicted network of protein-protein interaction of 361 Arabidopsis orthologs for 500 differentially regulated (250 upregulated and 250 downregulated genes in the barley susceptible cultivar Harrington at 72 hpi with *Bipolaris sorokiniana*) along with the Arabidopsis WAK homologs of candidate WAKs in the *Rcs5* locus. The table shows the protein nodes and their identity based on the Arabidopsis protein-protein interaction database known till date.

Supplementary Figure 1.

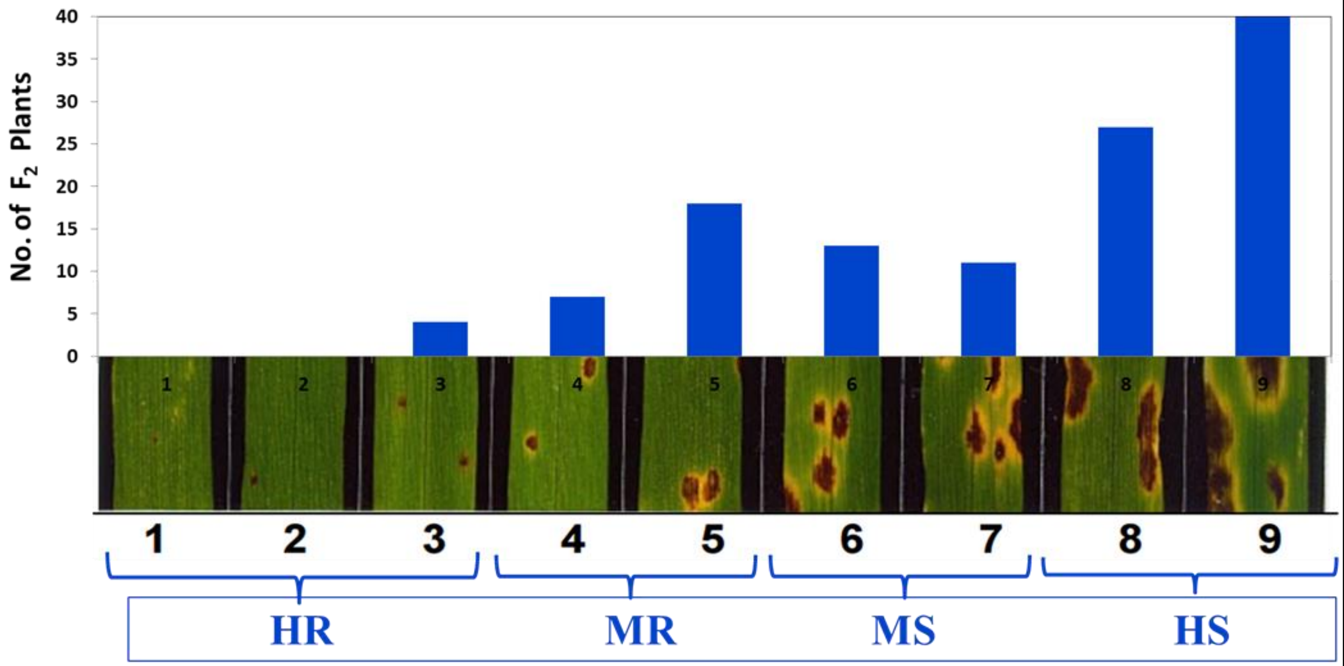


Disease Rating Scale (0-9; Fetch and Steffenson)

120 $F_2$ (1R: 3S) $\chi^2$= 0.0444, p value= 0.8330

Supplementary Figure 2.

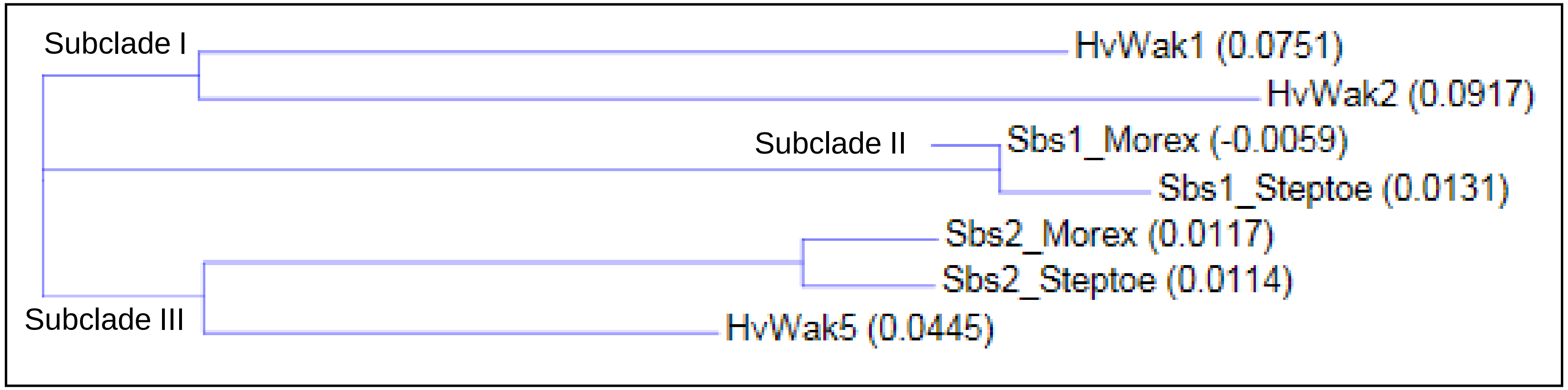

Supplimentary Figure 3

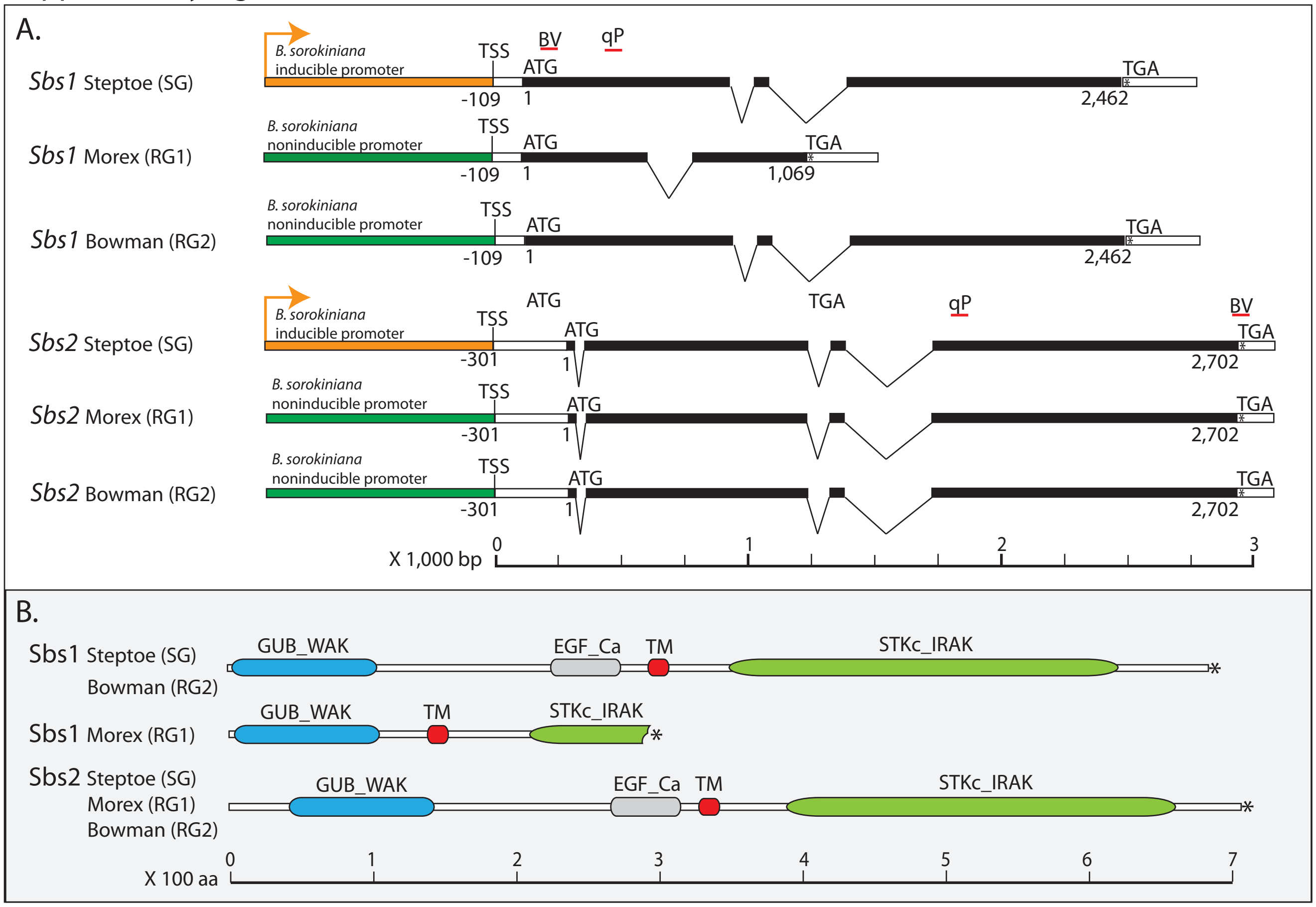

Supplementary Figure 4.

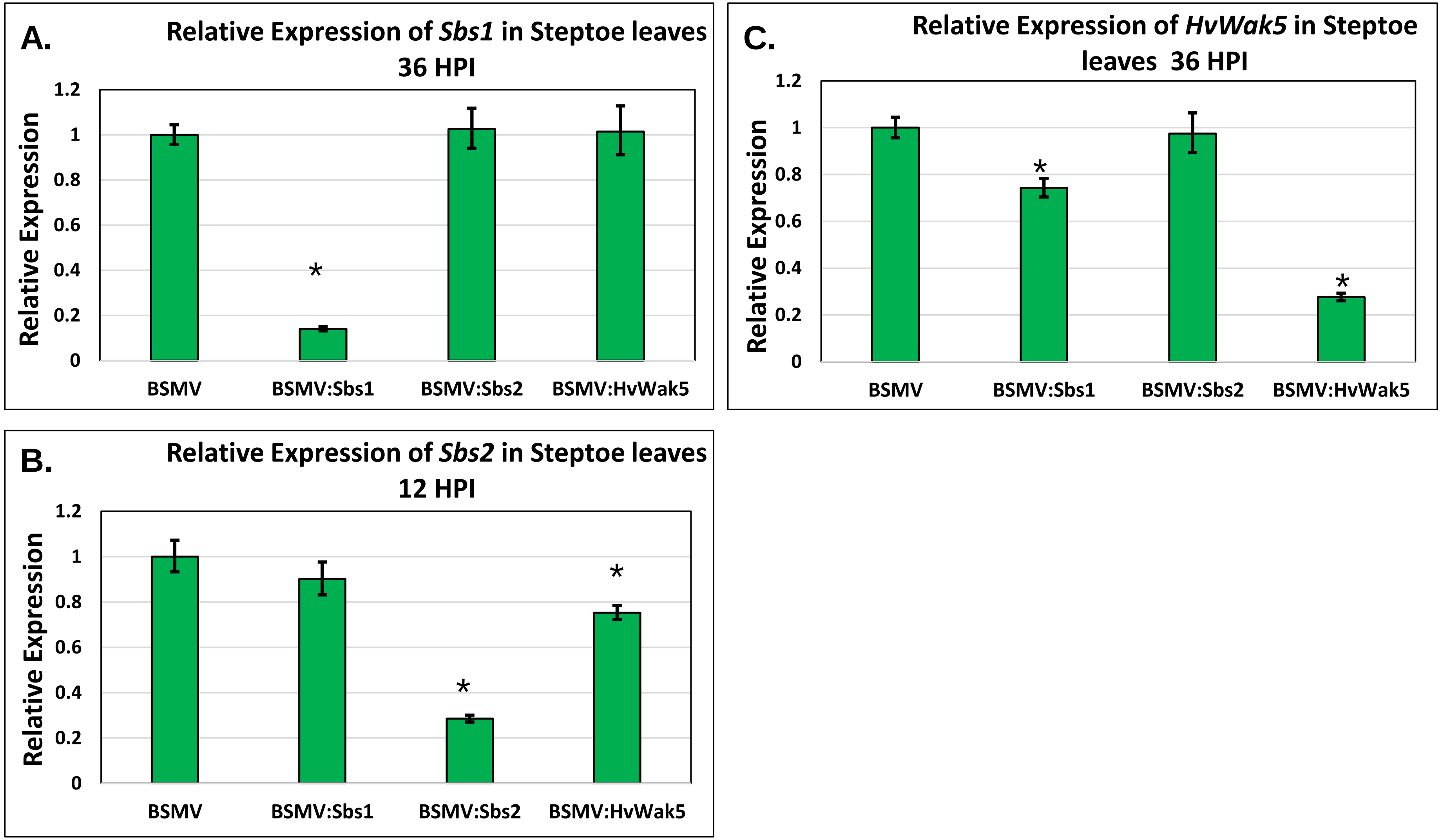

### Supplementary Figure 5A

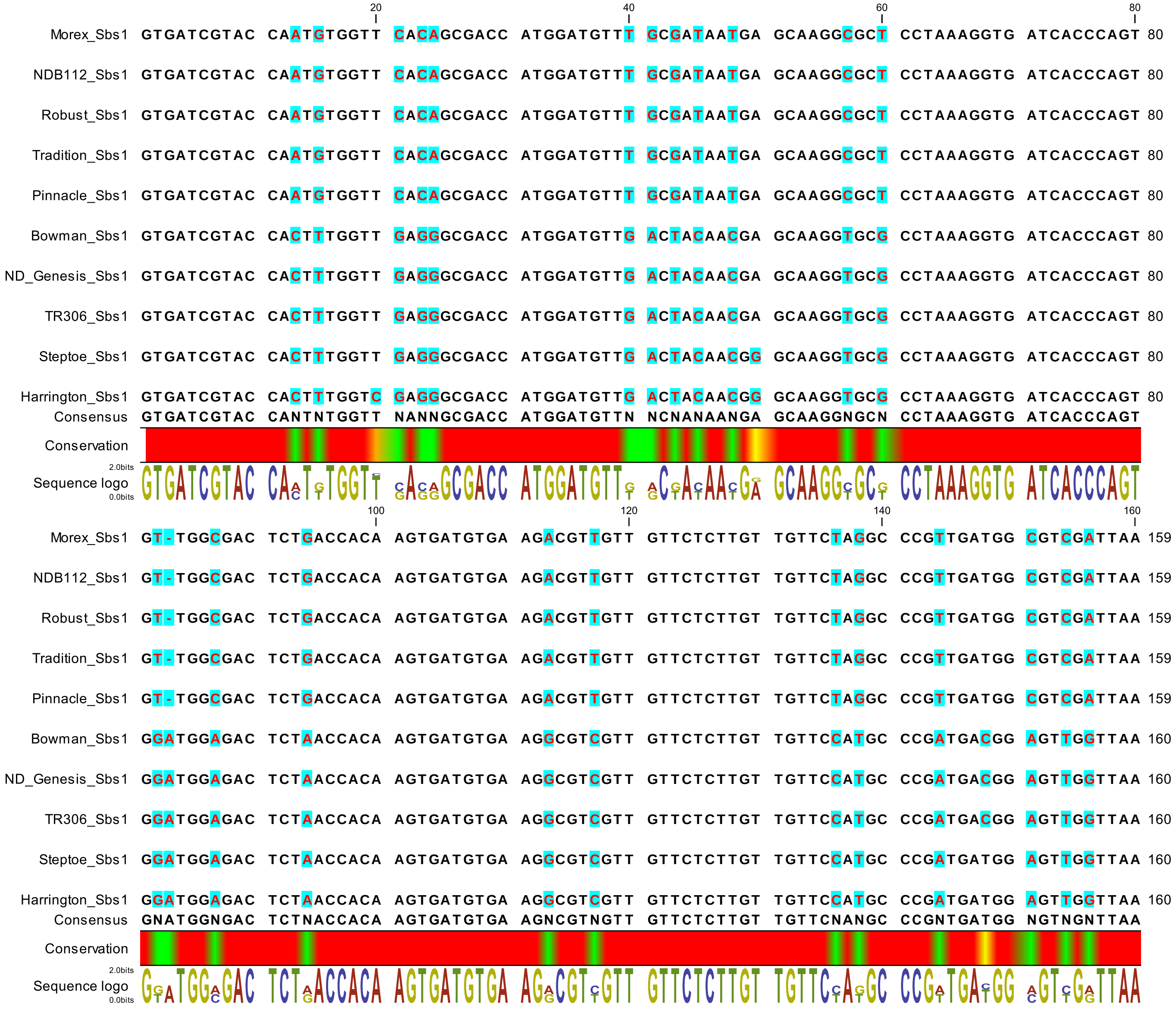

Supplementary Figure 5A

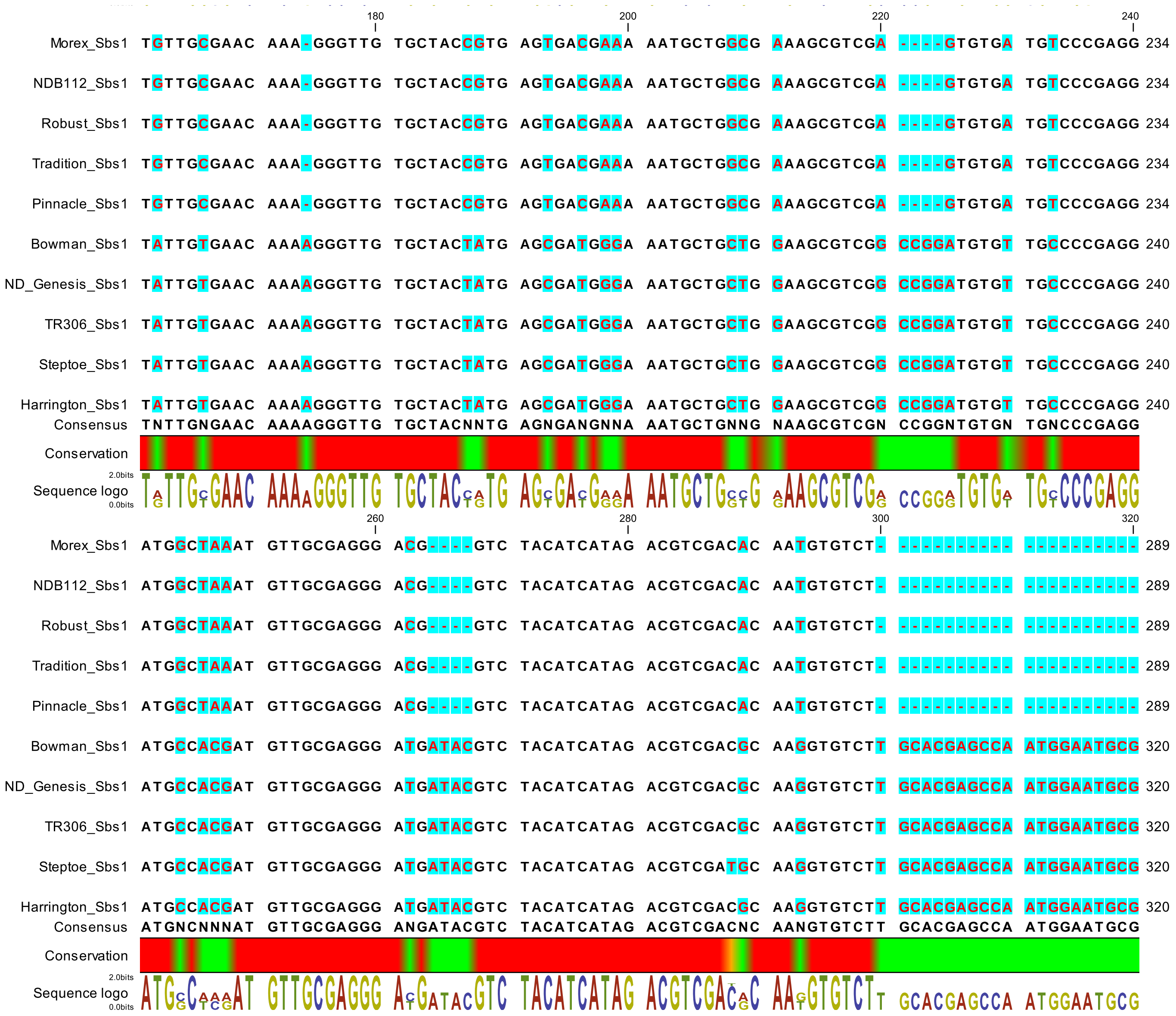

Supplementary Figure 5A

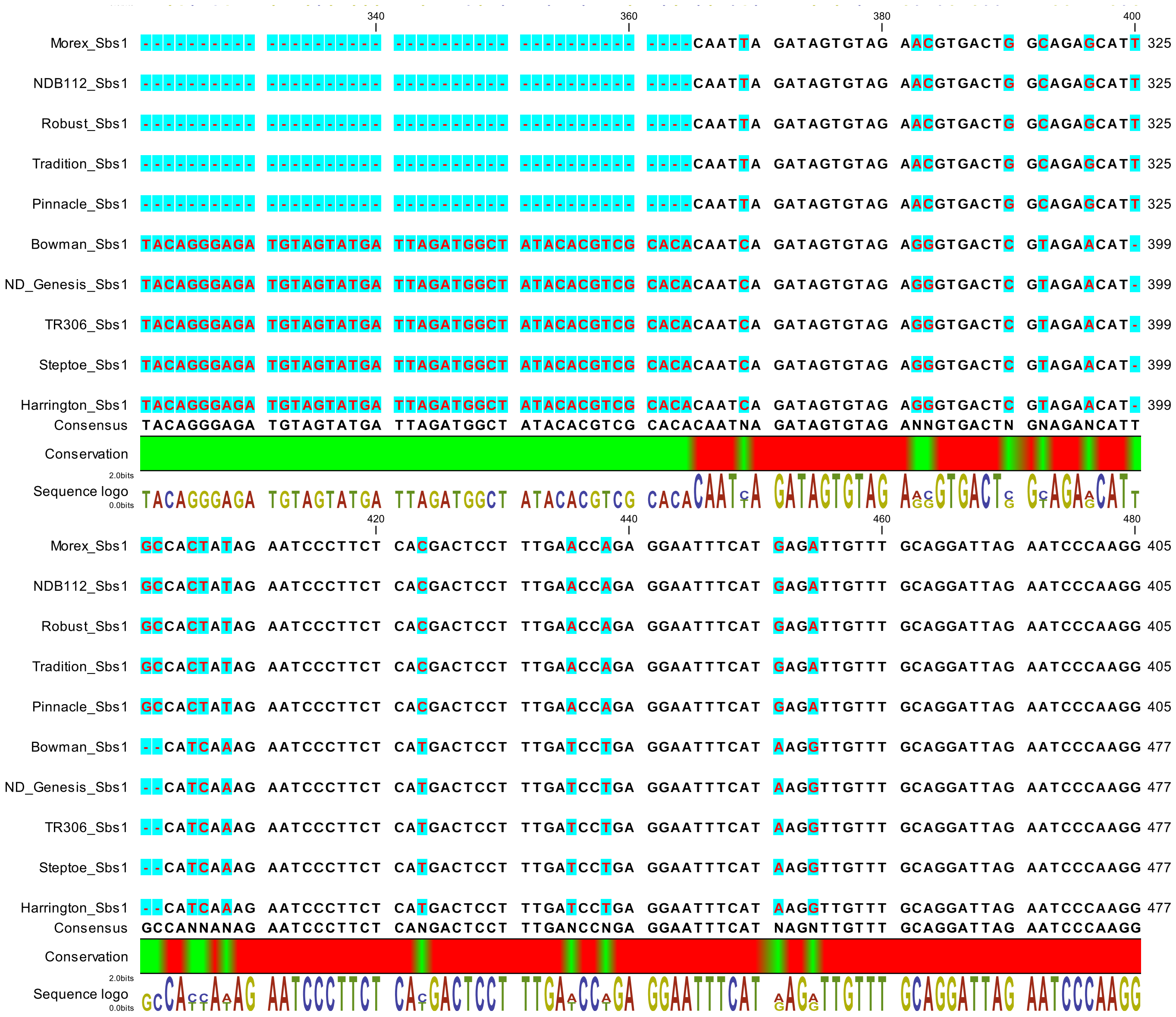

Supplementary Figure 5A

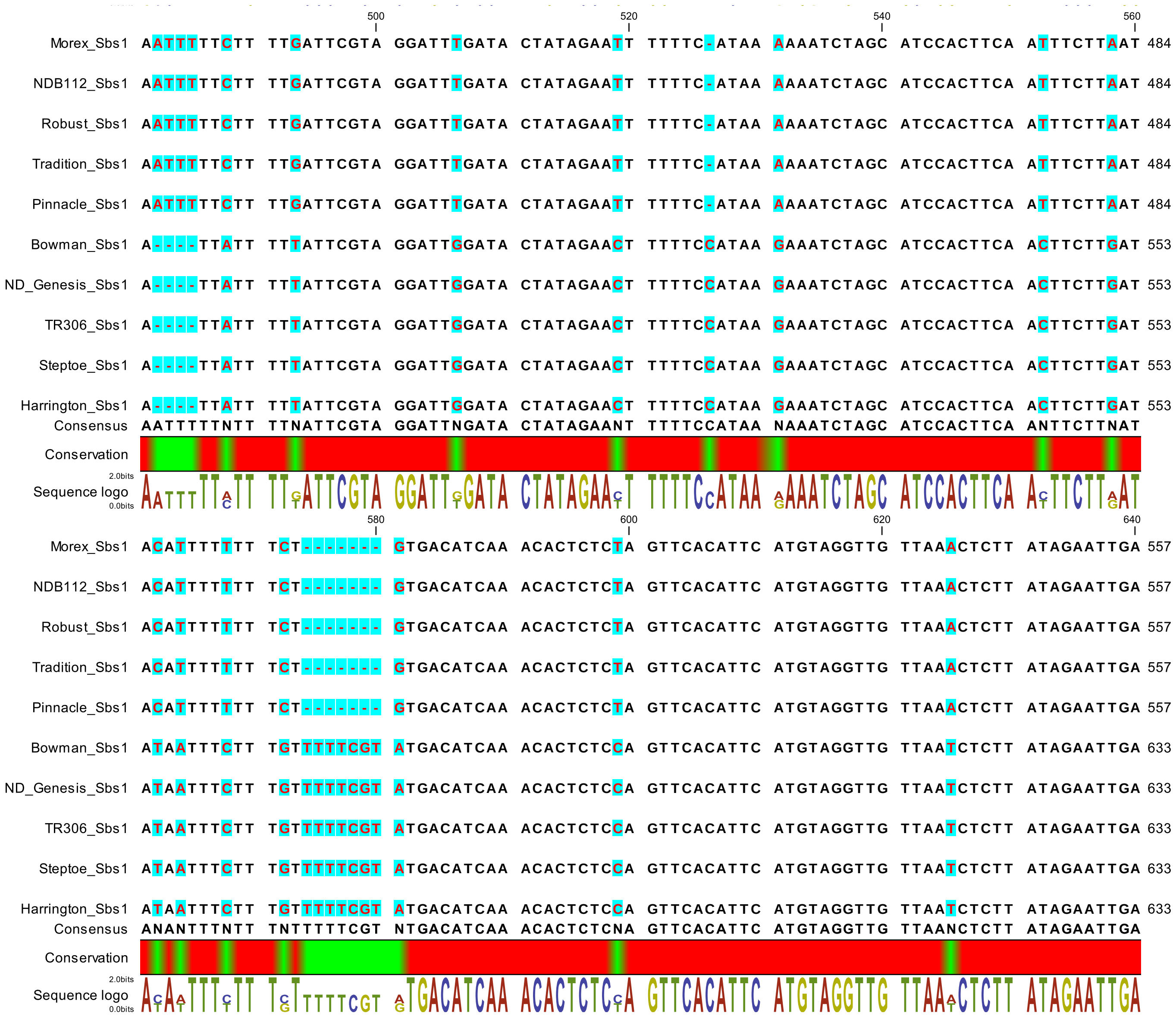

### Supplementary Figure 5A

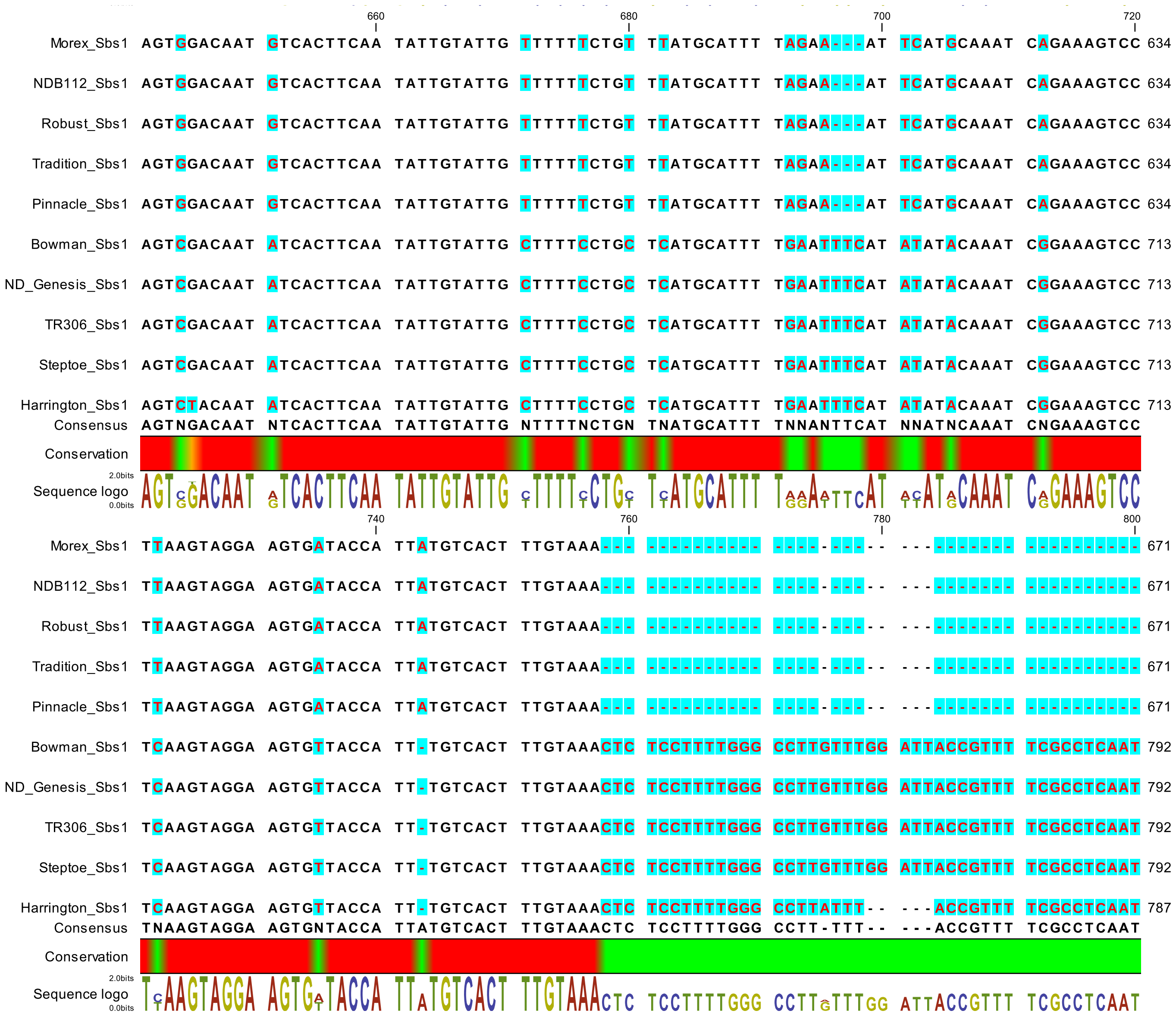

### Supplementary Figure 5A

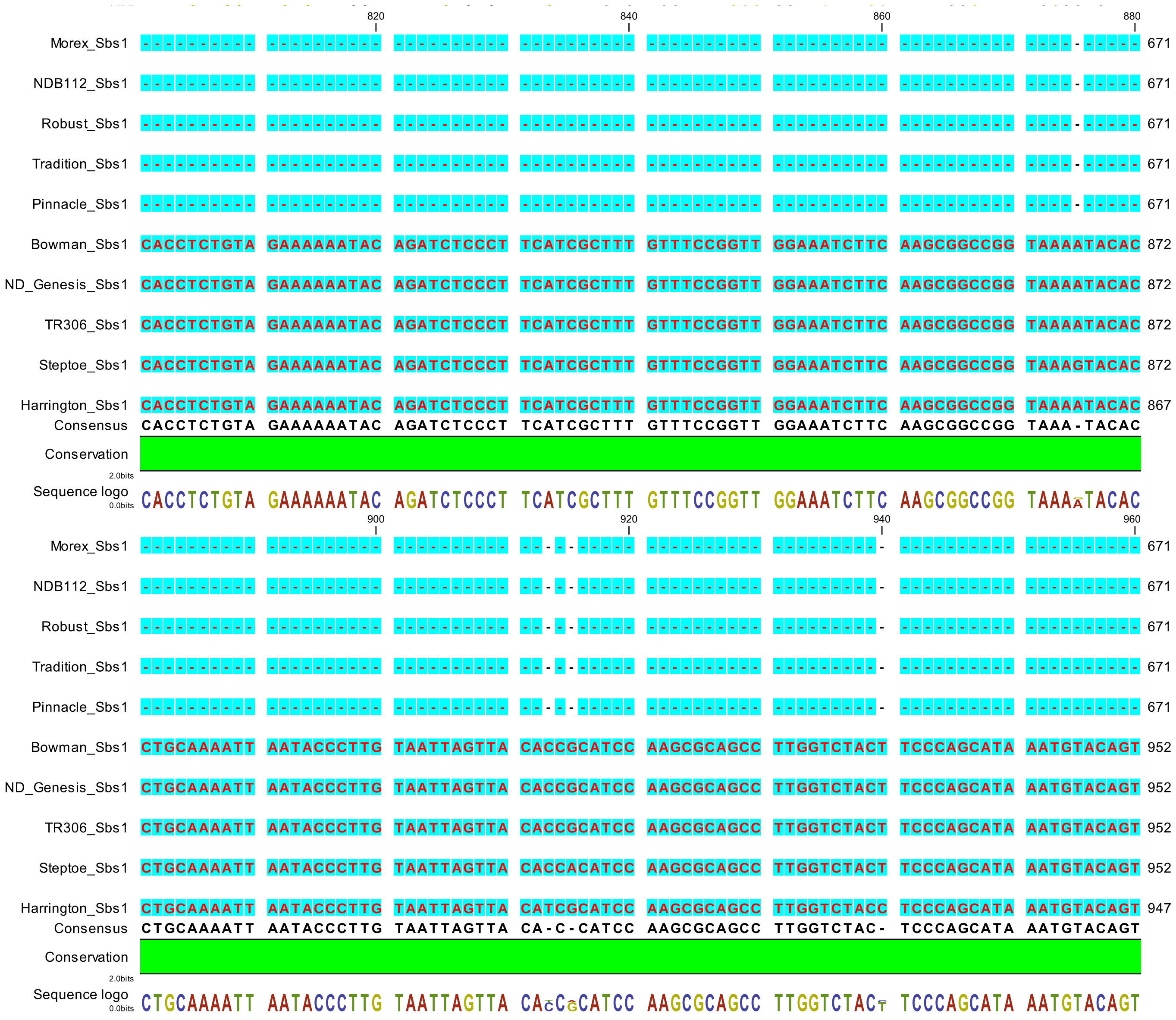

### Supplementary Figure 5A

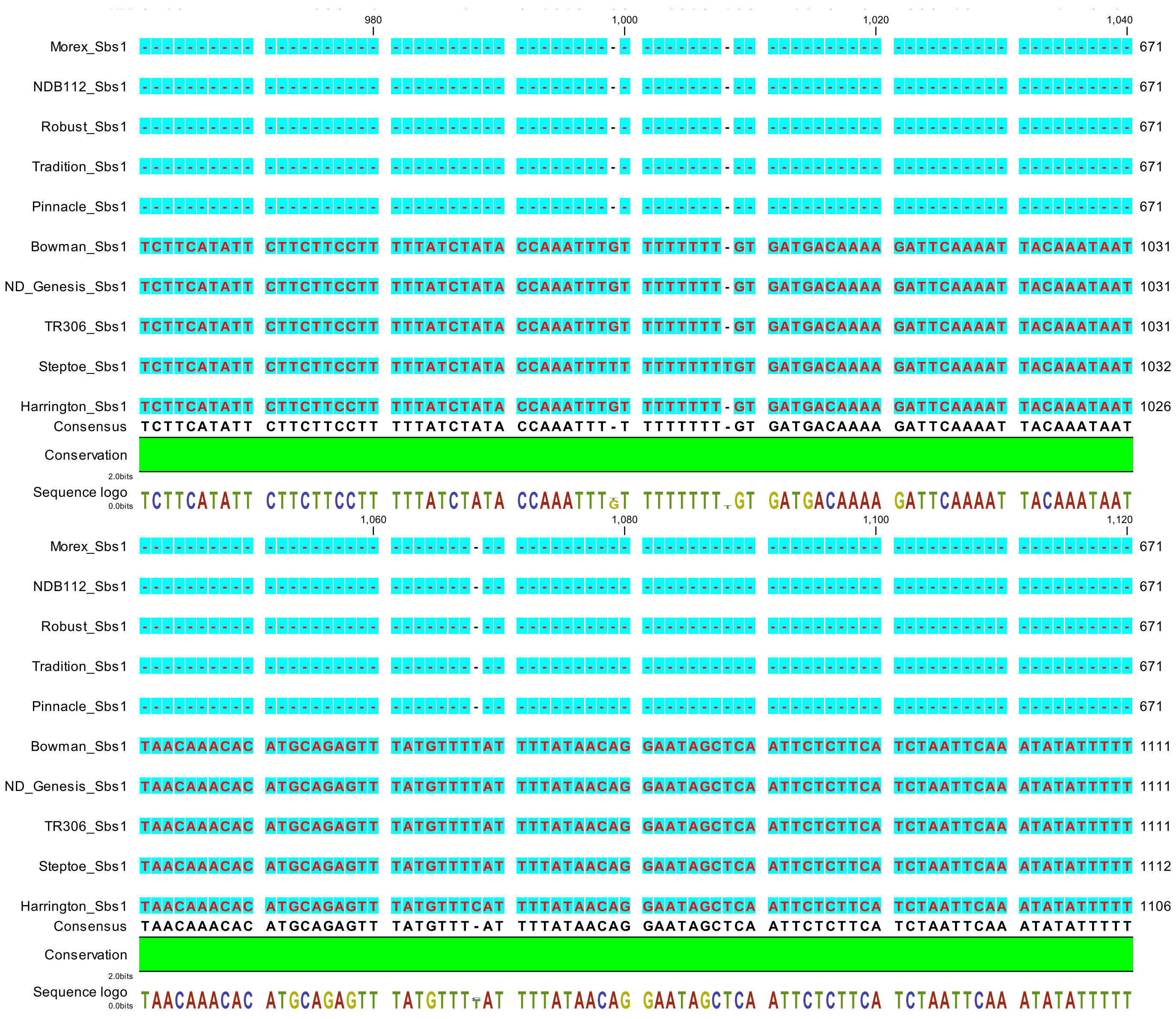

### Supplementary Figure 5A

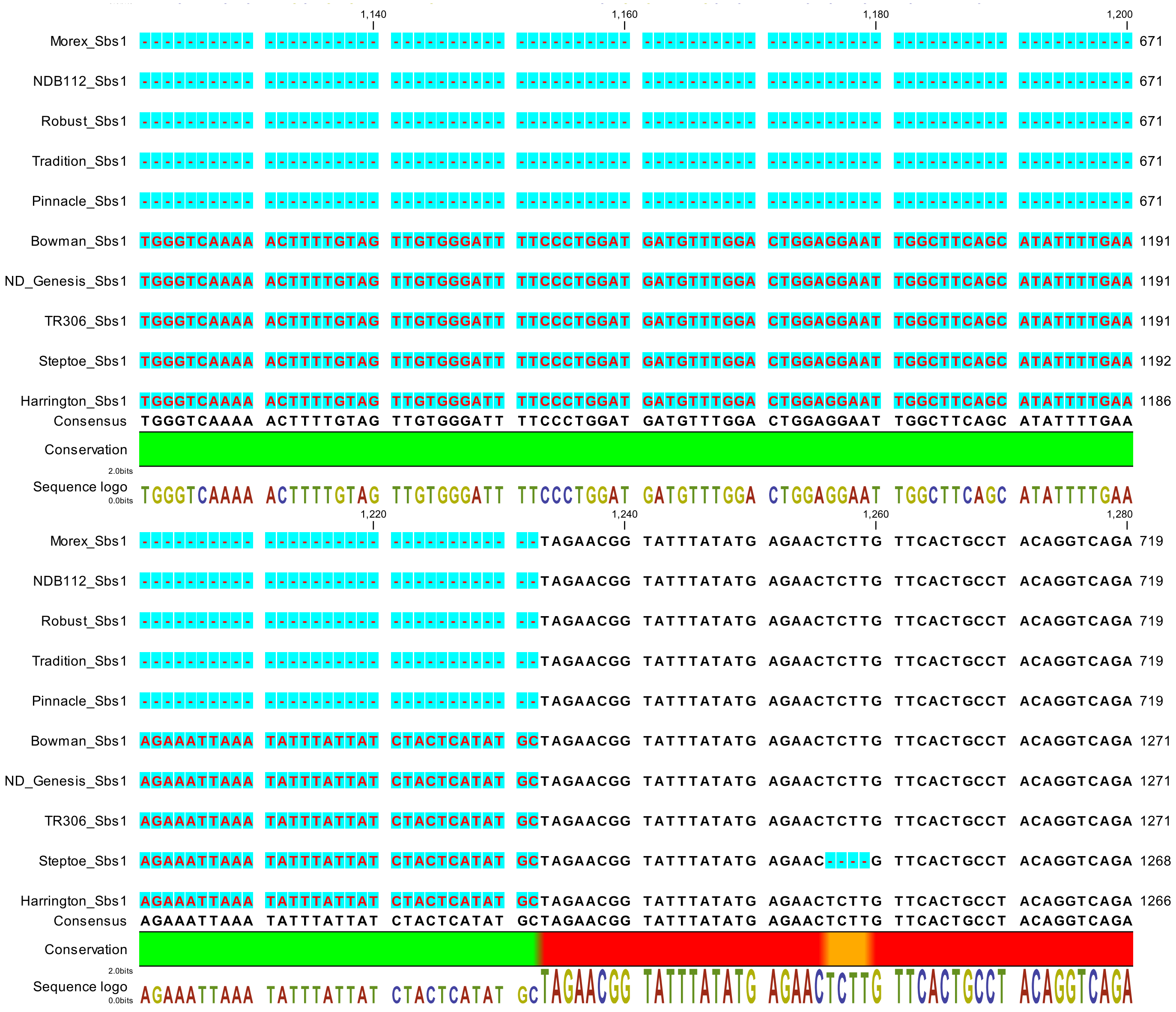

### Supplementary Figure 5A

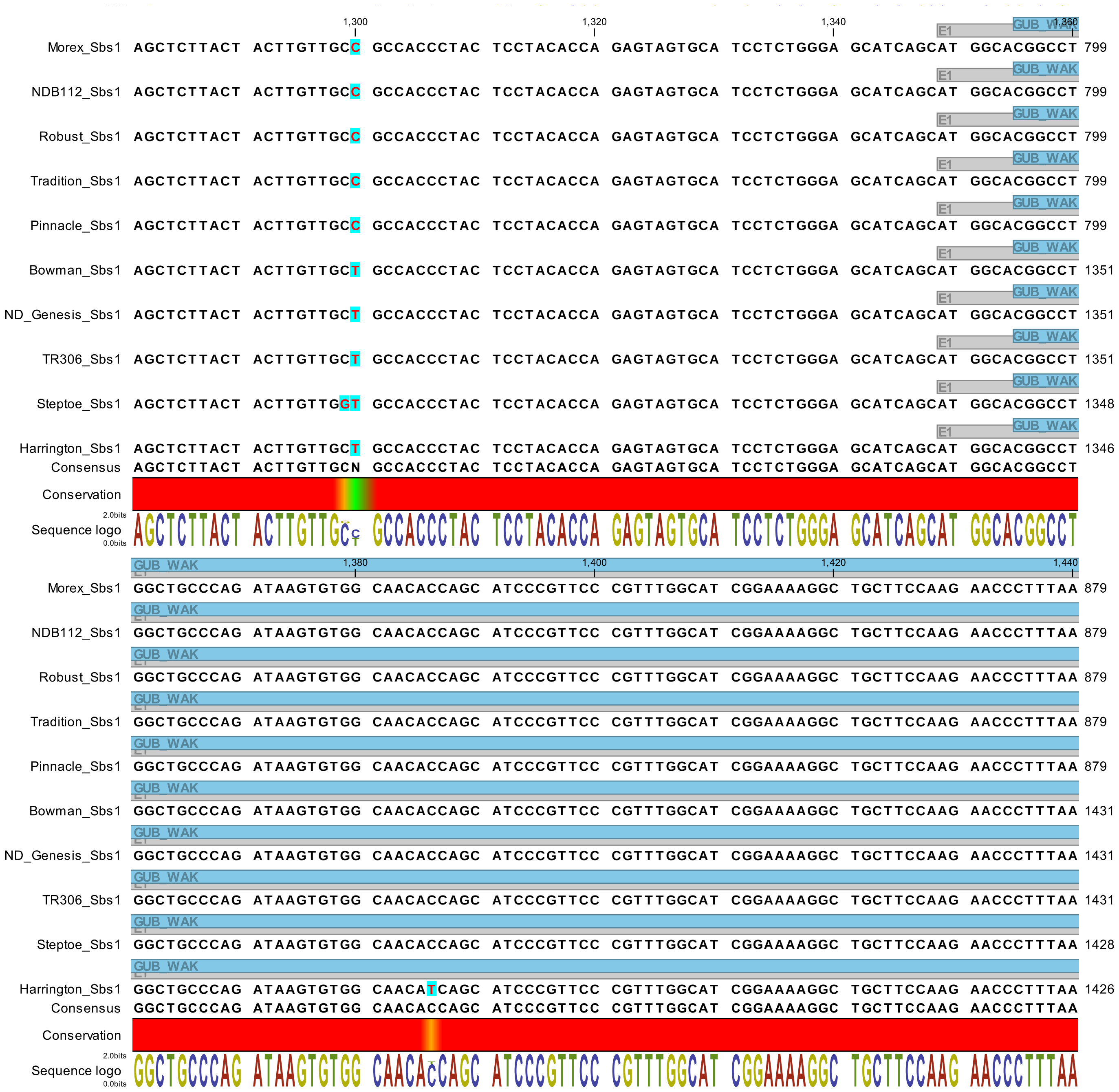

### Supplementary Figure 5A

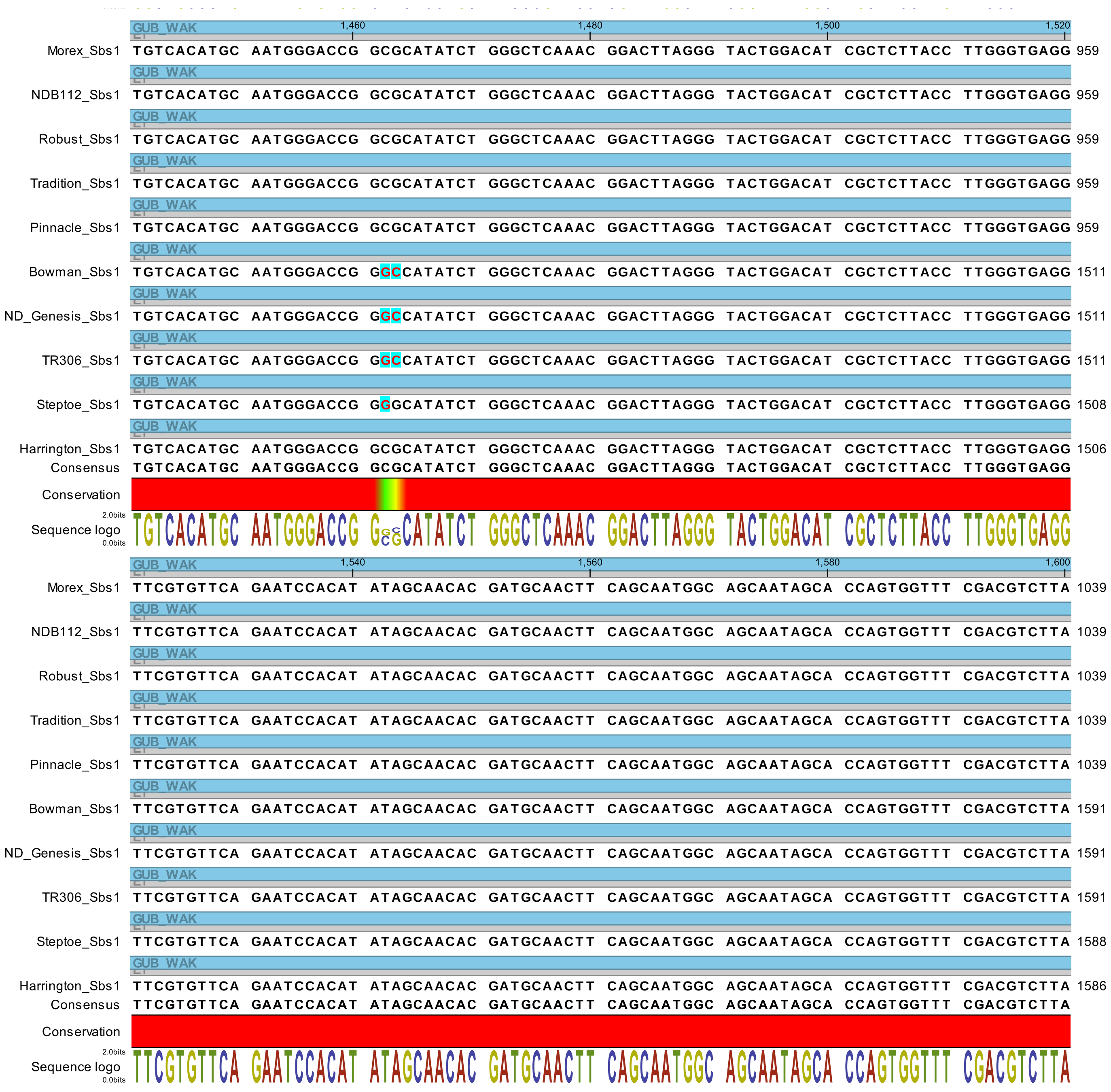

Supplementary Figure 5A

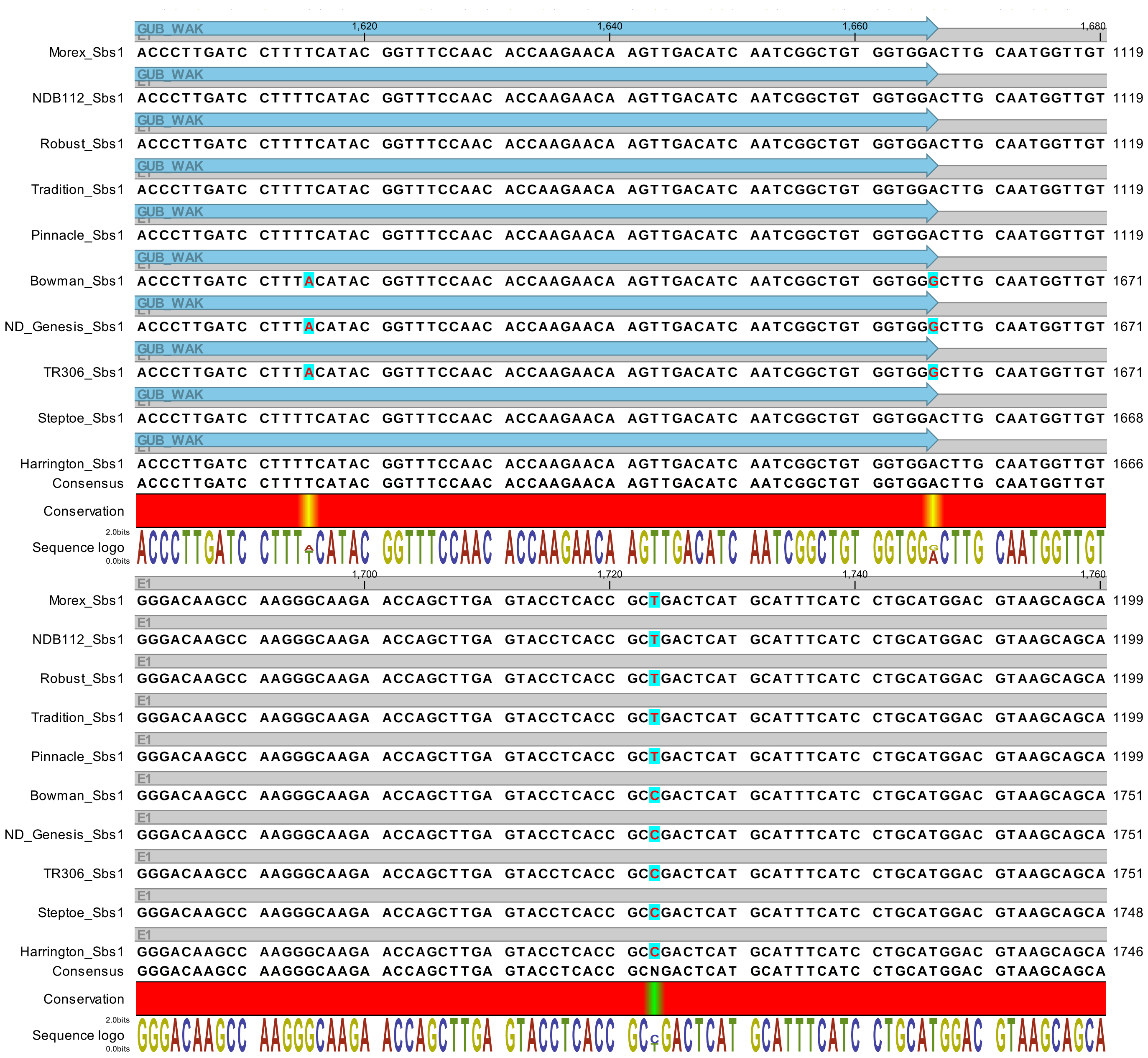

### Supplementary Figure 5A

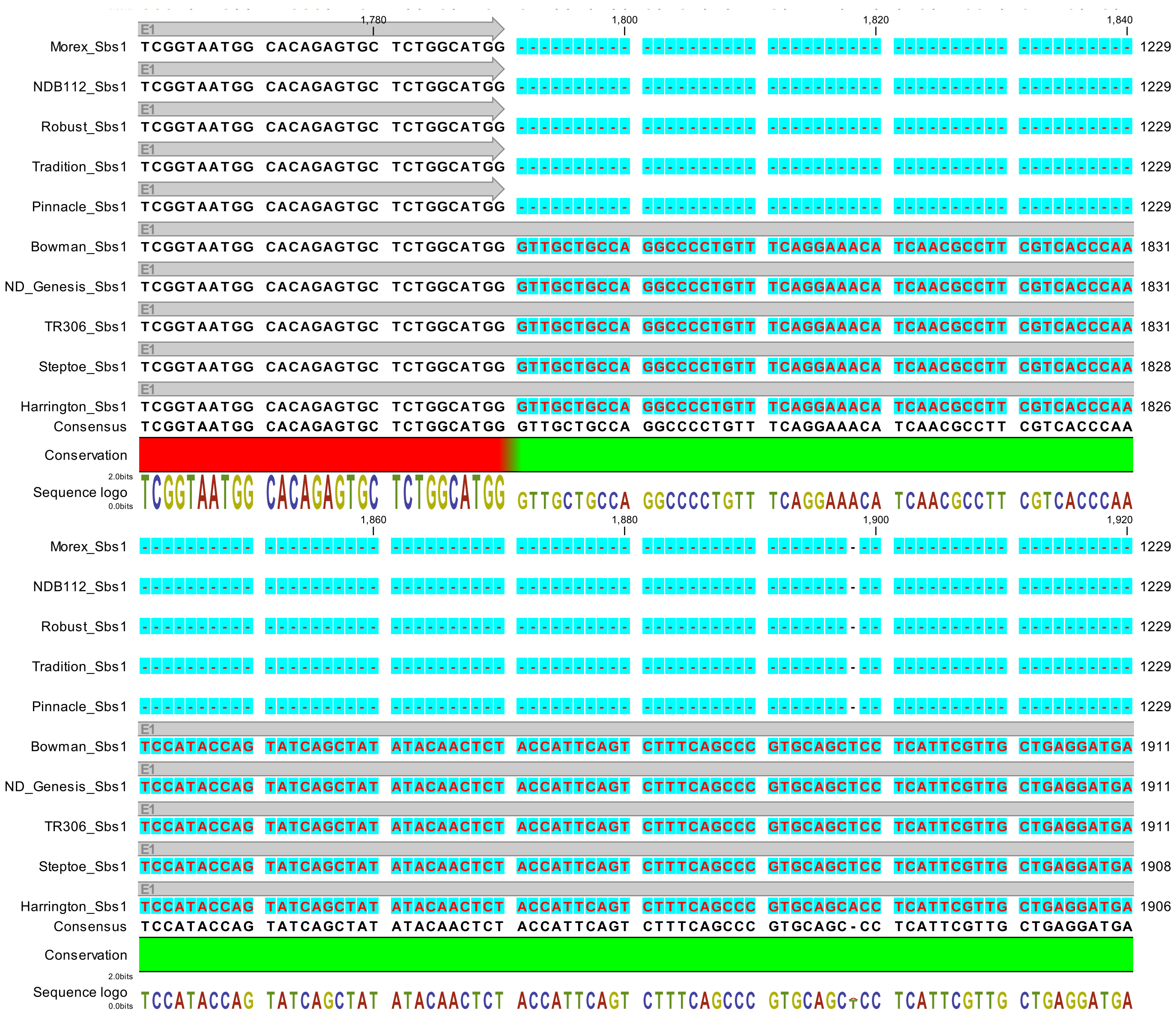

### Supplementary Figure 5A

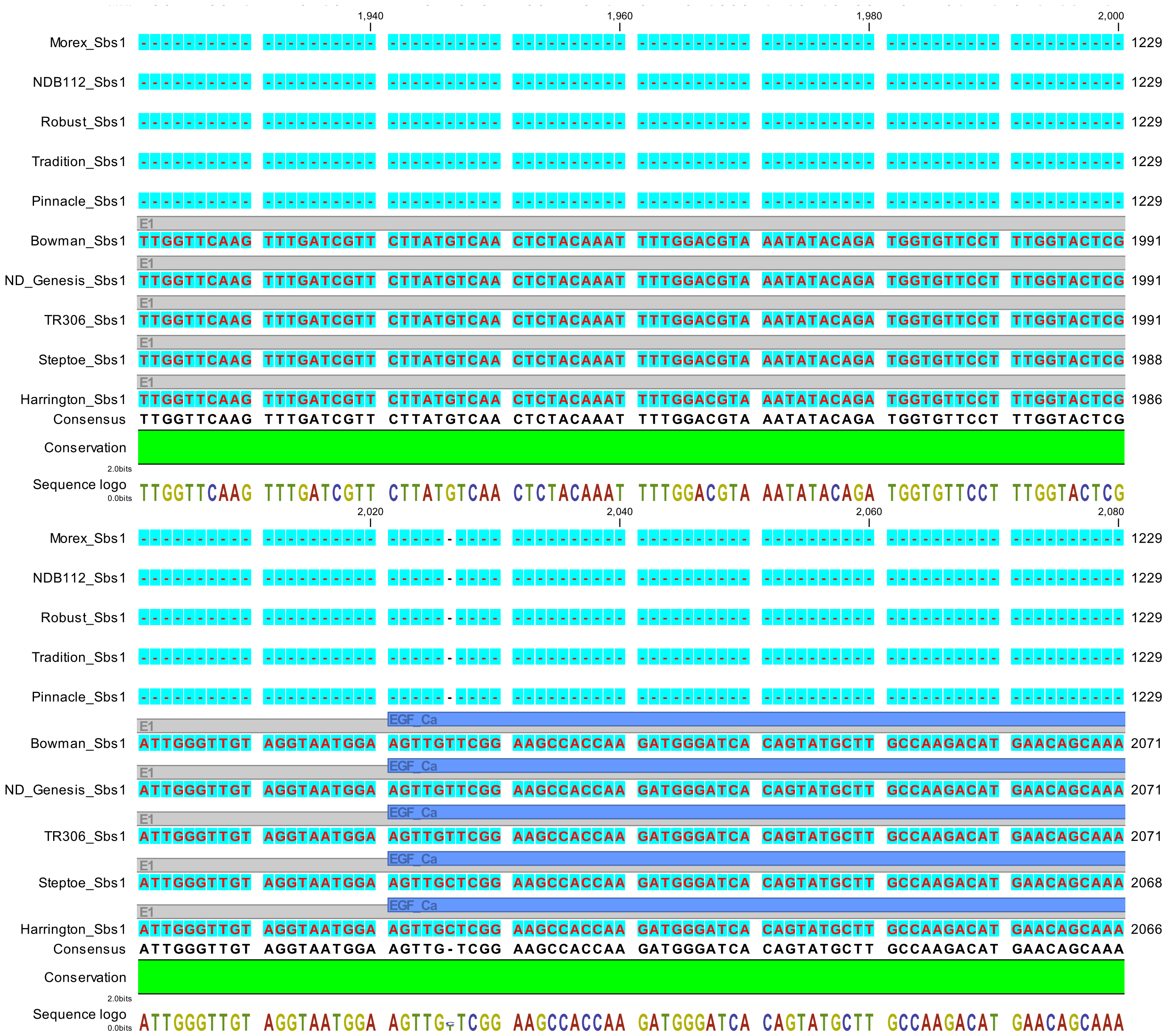

### Supplementary Figure 5A

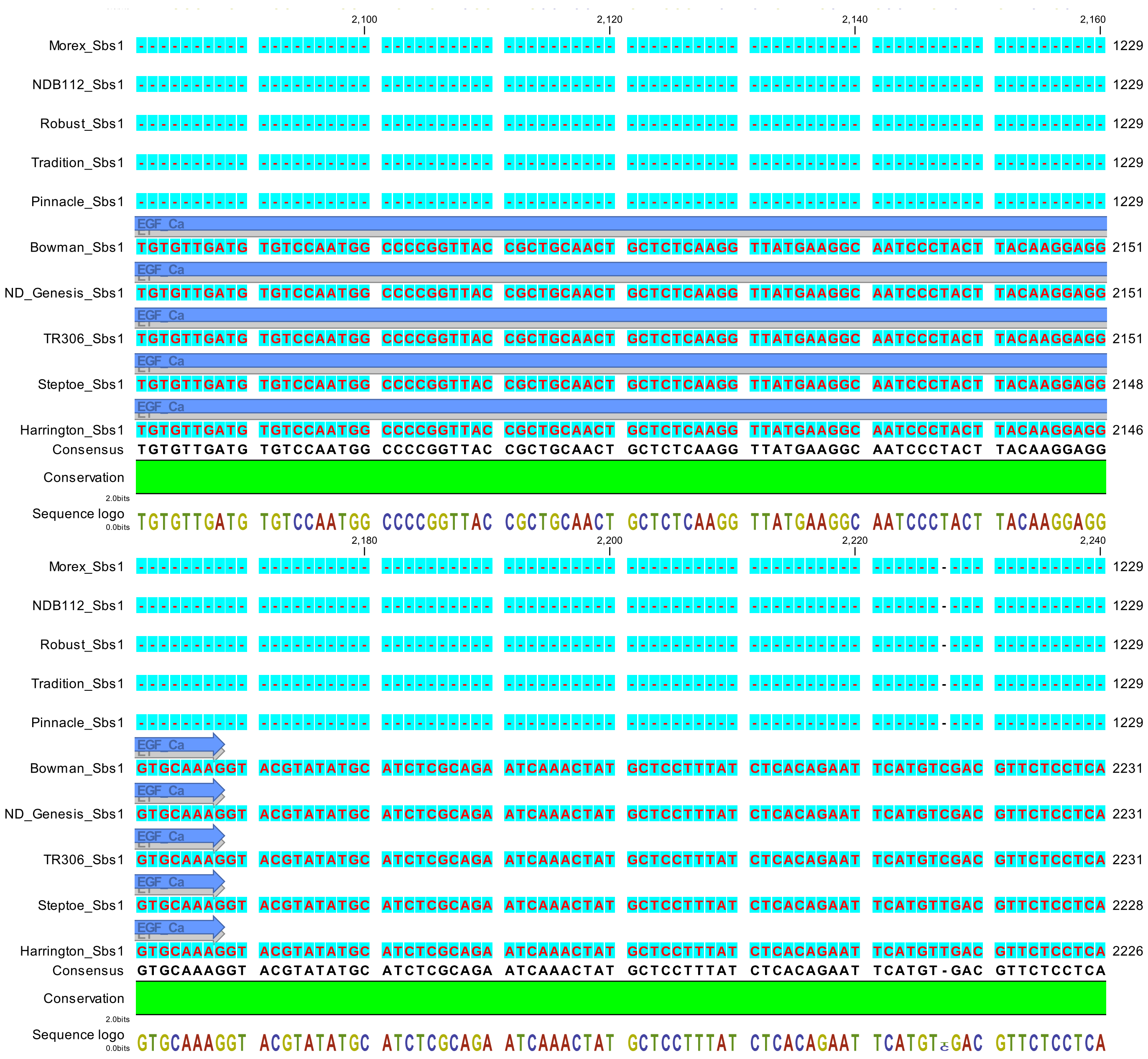

### Supplementary Figure 5A

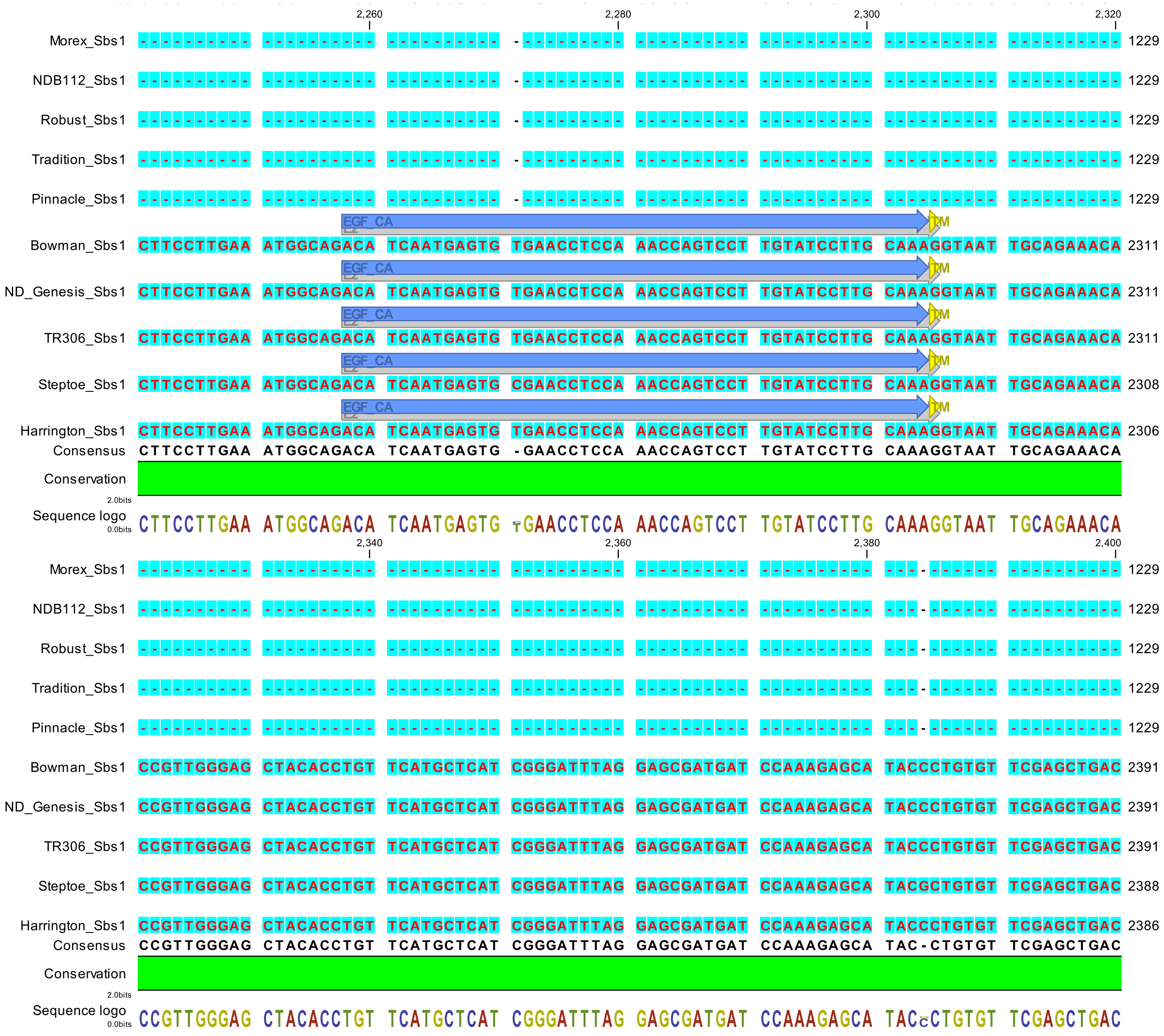

### Supplementary Figure 5A

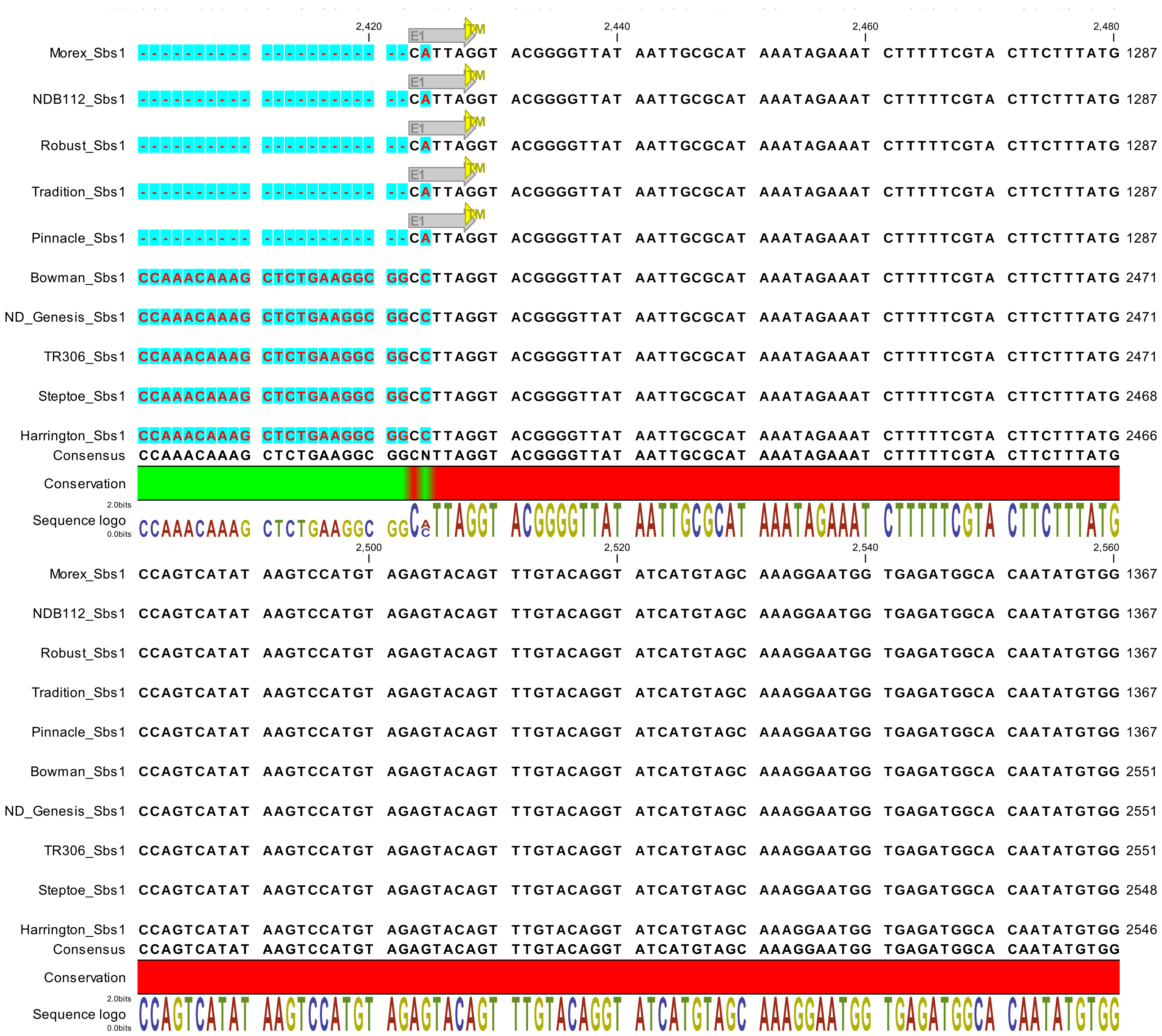

### Supplementary Figure 5A

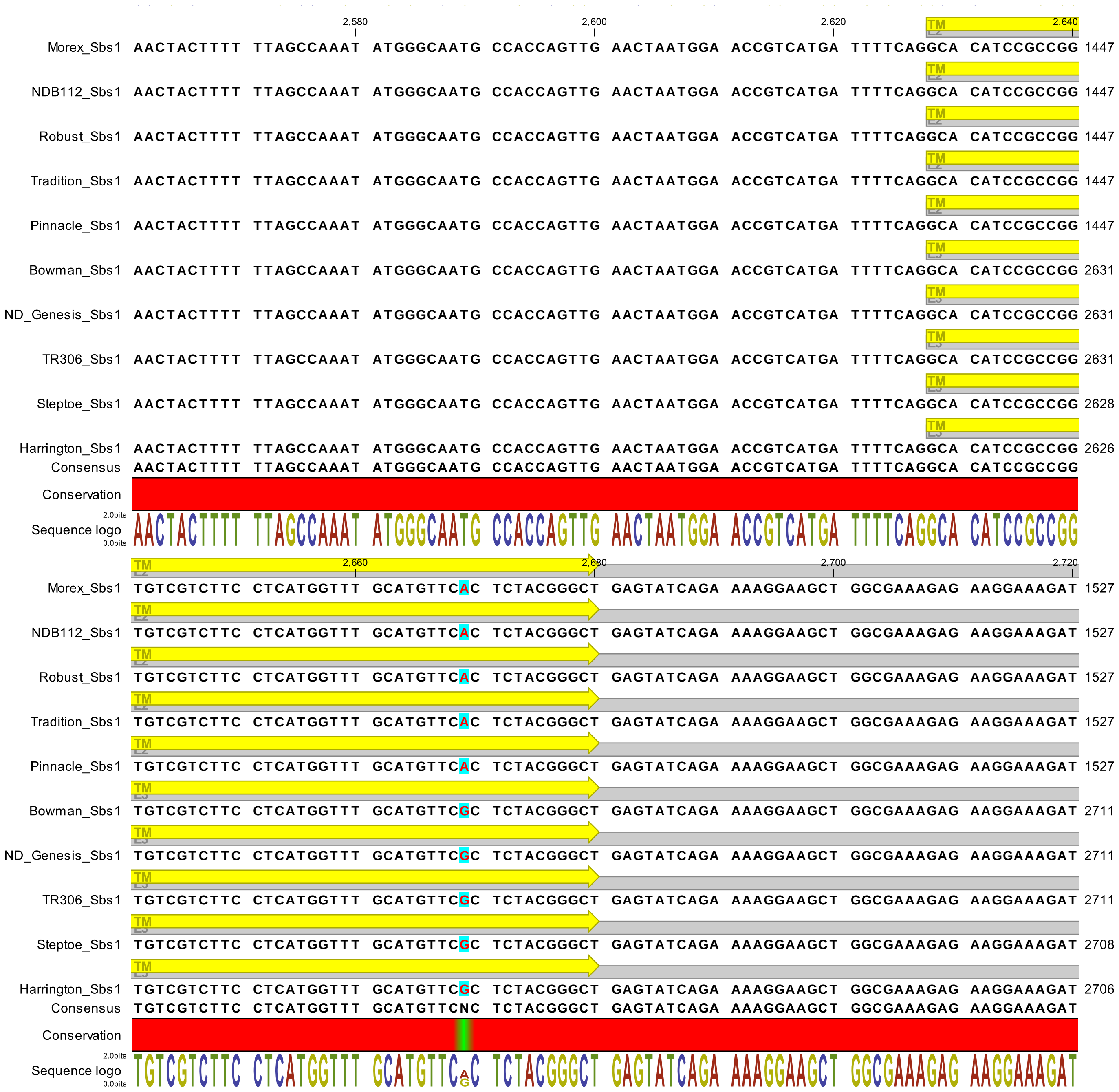

### Supplementary Figure 5A

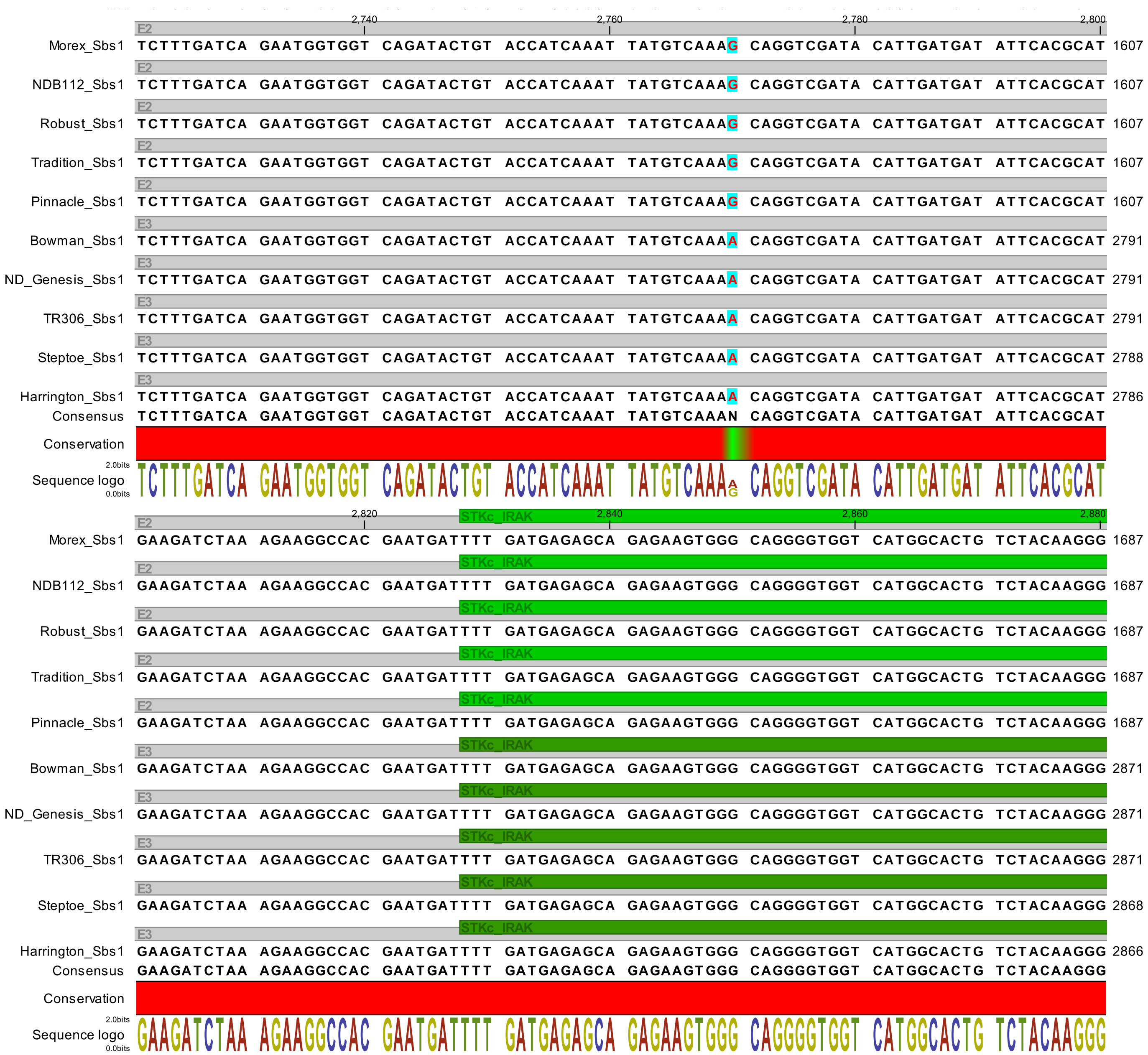

Supplementary Figure 5A

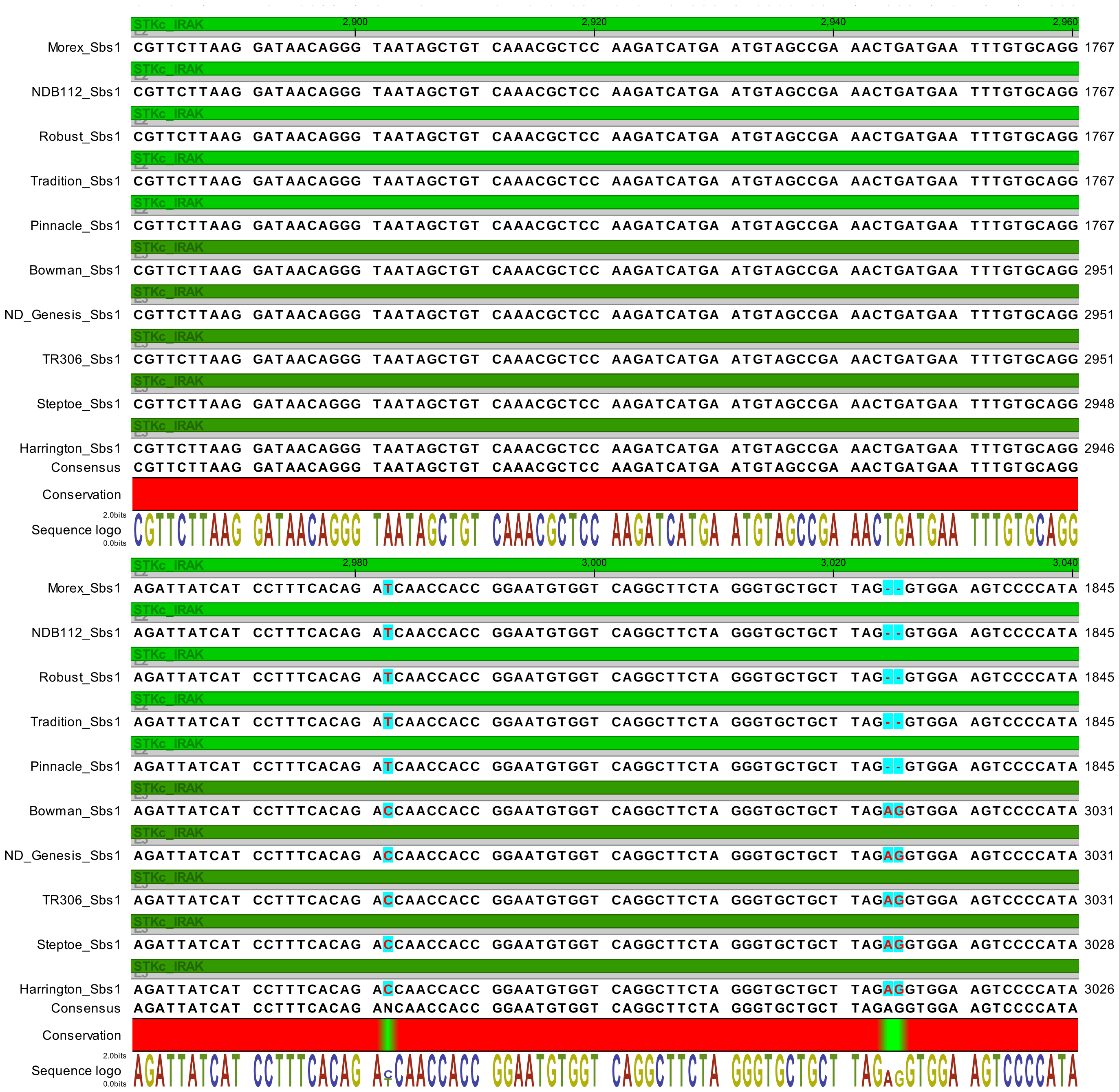

### Supplementary Figure 5A

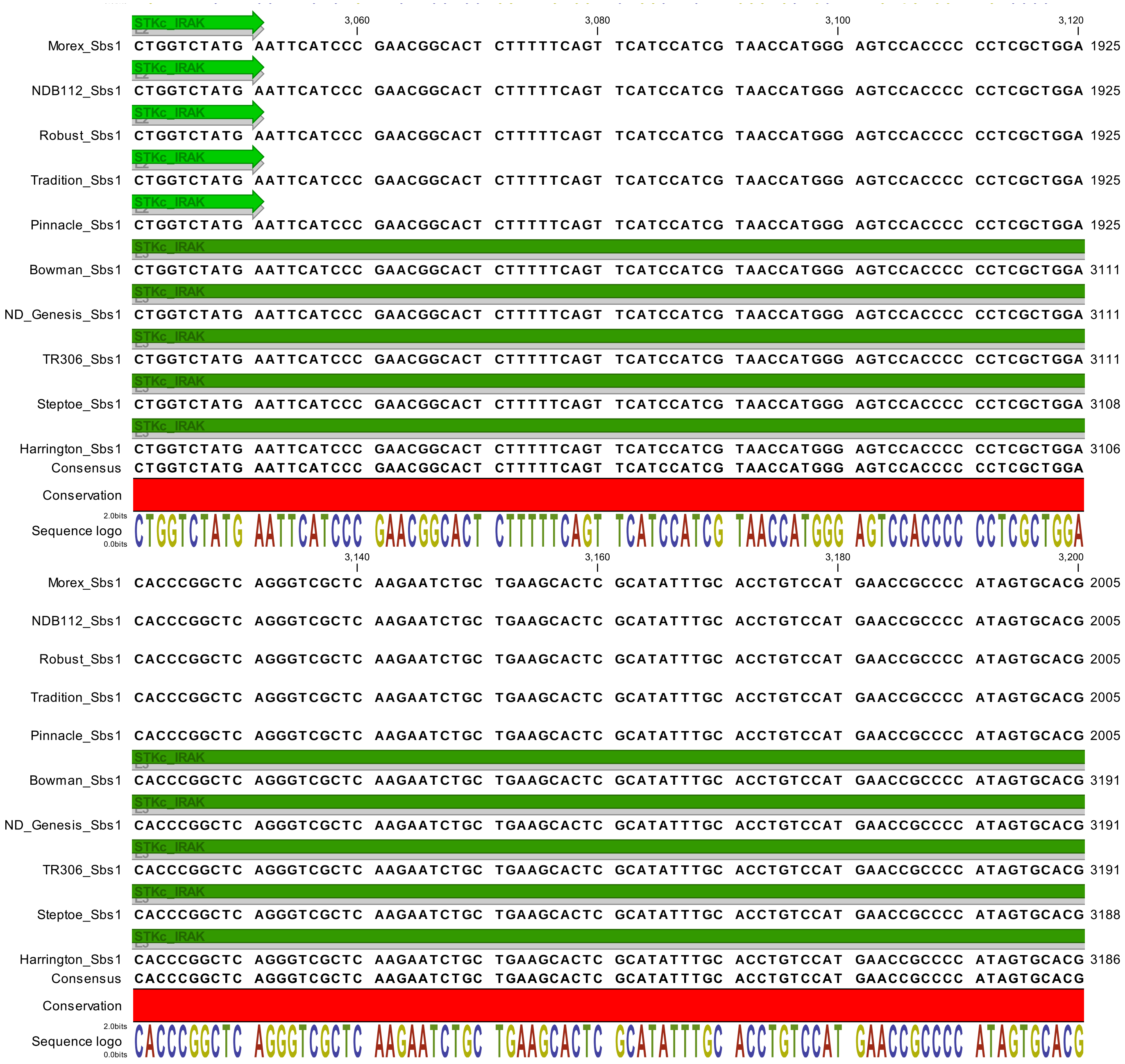

### Supplementary Figure 5A

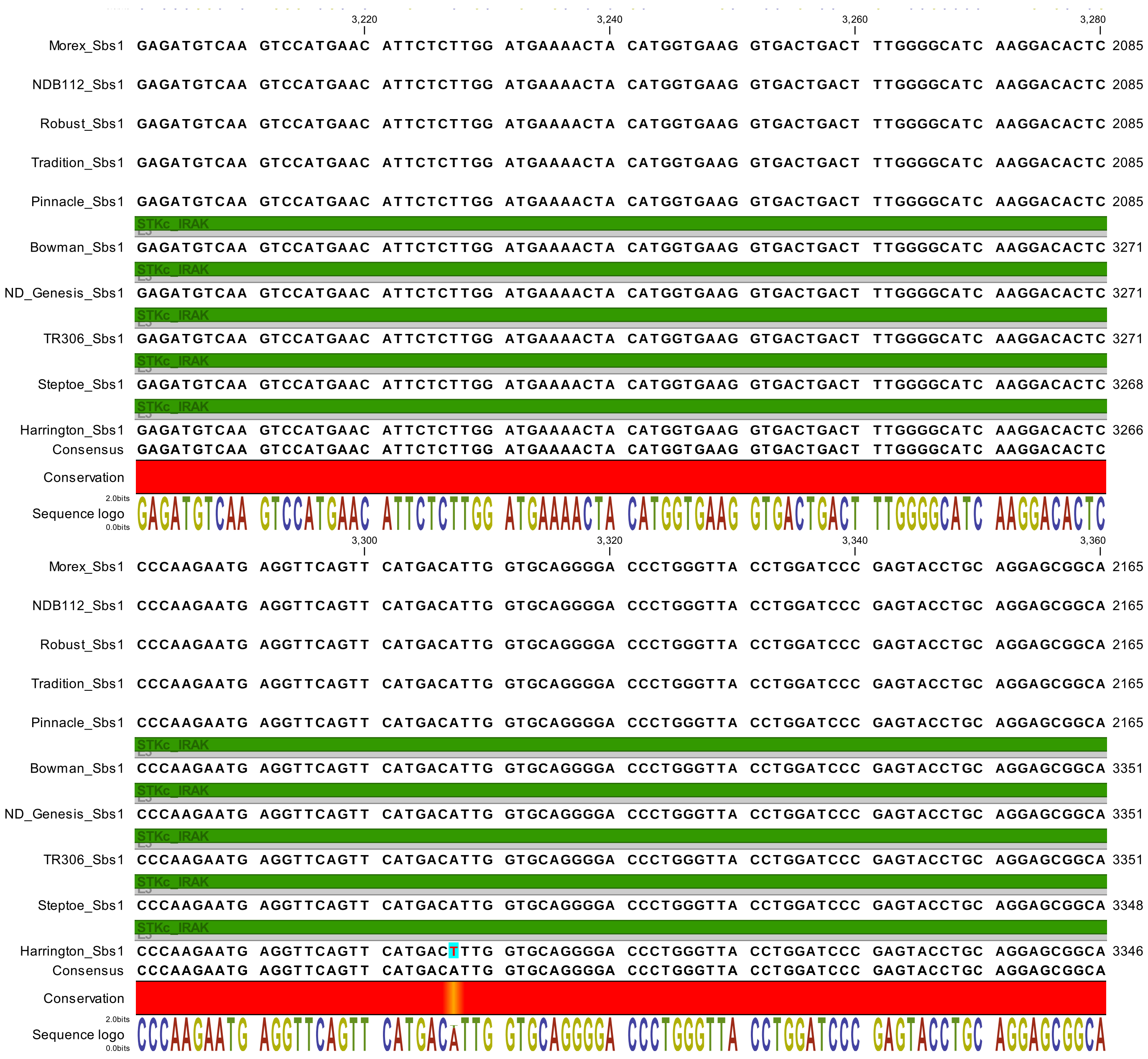

### Supplementary Figure 5A

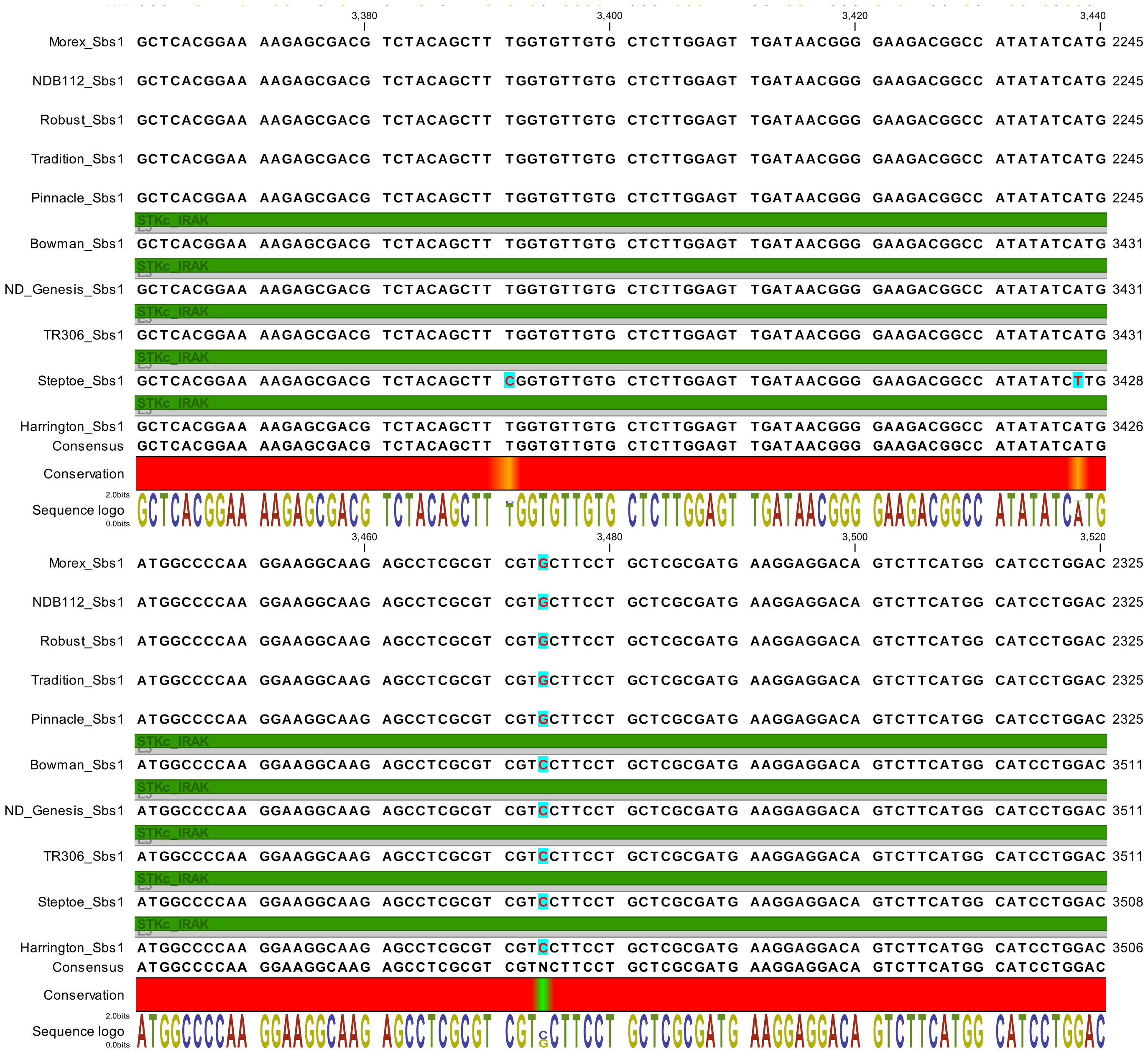

### Supplementary Figure 5A

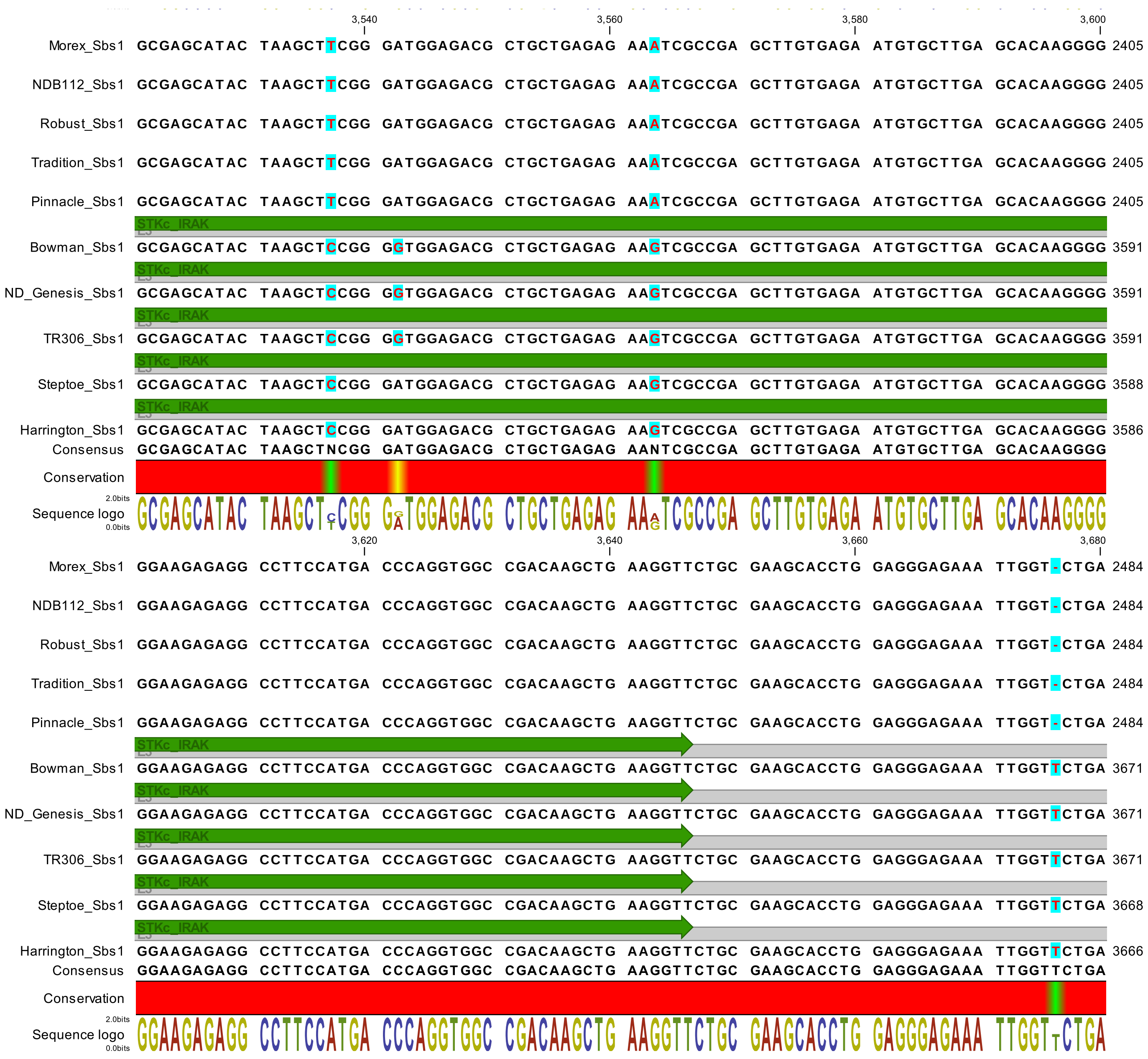

### Supplementary Figure 5A

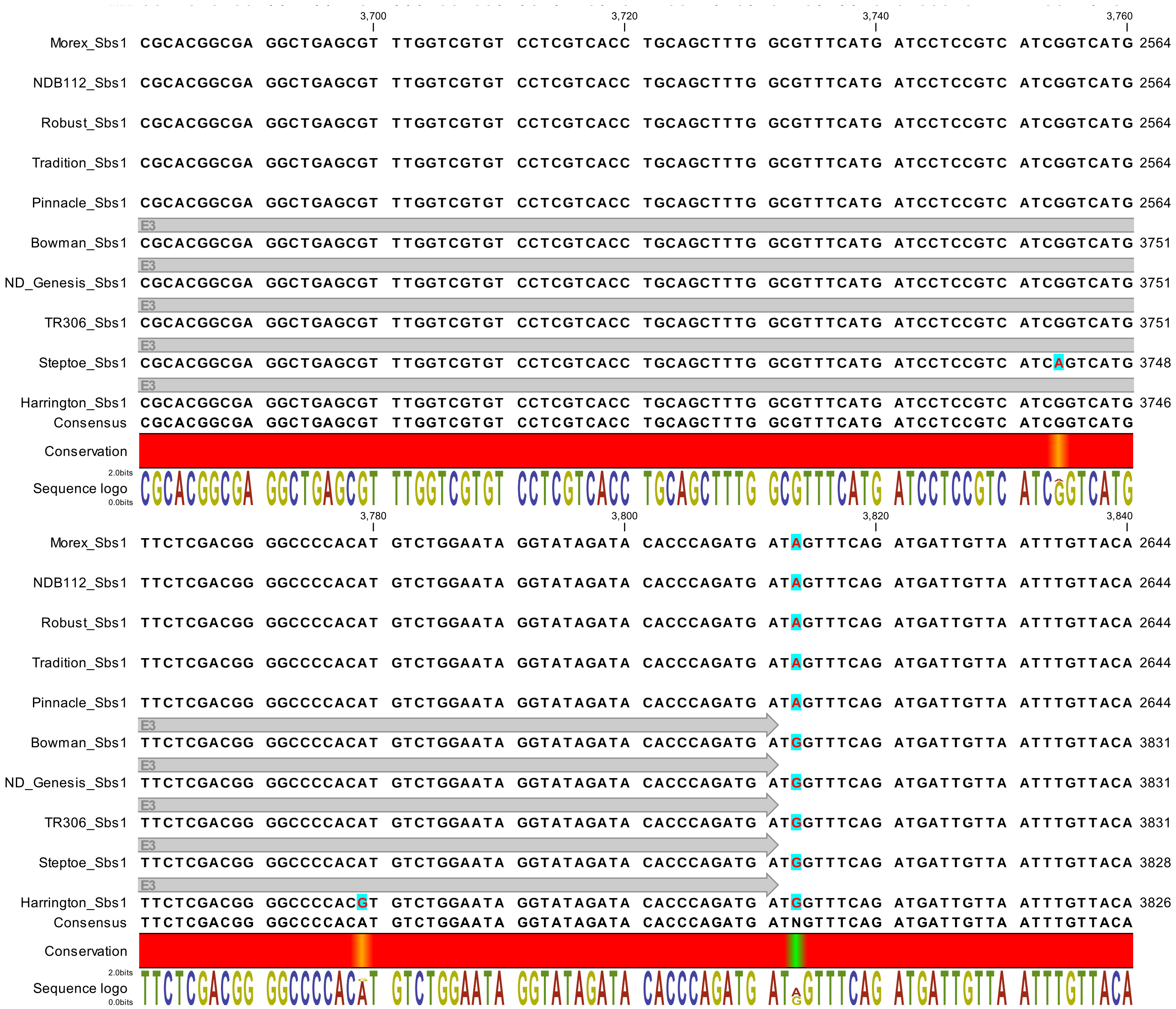

### Supplementary Figure 5A

| | 3,860 | | 3,880 | | 3,900 | | 3,920 | | |
|---|---|---|---|---|---|---|---|---|---|
| Morex_Sbs1 | CCATGCCAAG | AGTATGGCTT | GCGGTATGCT | GTCCACATTT | TGTCCTGTAT | TTGTGGTACC | AAGTACTATT | ATTAGTGTAG | 2724 |
| NDB112_Sbs1 | CCATGCCAAG | AGTATGGCTT | GCGGTATGCT | GTCCACATTT | TGTCCTGTAT | TTGTGGTACC | AAGTACTATT | ATTAGTGTAG | 2724 |
| Robust_Sbs1 | CCATGCCAAG | AGTATGGCTT | GCGGTATGCT | GTCCACATTT | TGTCCTGTAT | TTGTGGTACC | AAGTACTATT | ATTAGTGTAG | 2724 |
| Tradition_Sbs1 | CCATGCCAAG | AGTATGGCTT | GCGGTATGCT | GTCCACATTT | TGTCCTGTAT | TTGTGGTACC | AAGTACTATT | ATTAGTGTAG | 2724 |
| Pinnacle_Sbs1 | CCATGCCAAG | AGTATGGCTT | GCGGTATGCT | GTCCACATTT | TGTCCTGTAT | TTGTGGTACC | AAGTACTATT | ATTAGTGTAG | 2724 |
| Bowman_Sbs1 | CCATGCCAAG | AGTATGGCTT | GCGGTATGCT | GTCCACATTT | TGTCCTGTAT | TTGTGGTACC | AAGTACTATT | ATTAGTGTAG | 3911 |
| ND_Genesis_Sbs1 | CCATGCCAAG | AGTATGGCTT | GCGGTATGCT | GTCCACATTT | TGTCCTGTAT | TTGTGGTACC | AAGTACTATT | ATTAGTGTAG | 3911 |
| TR306_Sbs1 | CCATGCCAAG | AGTATGGCTT | GCGGTATGCT | GTCCACATTT | TGTCCTGTAT | TTGTGGTACC | AAGTACTATT | ATTAGTGTAG | 3911 |
| Steptoe_Sbs1 | CCATGCCAAG | AGTATGGCTT | GCGGTATGCT | GTCCACATTT | TGTCCTGTAT | TTGTGGTACC | AAGTACTATT | ATTAGTGTAG | 3908 |
| Harrington_Sbs1 | CCATGCCAAG | AGTATGGCTT | GCGGTATGCT | GTCCACATTT | TGTCCTGTAT | TTGTGGTACC | AAGTACTATT | ATTAGTGTAG | 3906 |
| Consensus | CCATGCCAAG | AGTATGGCTT | GCGGTATGCT | GTCCACATTT | TGTCCTGTAT | TTGTGGTACC | AAGTACTATT | ATTAGTGTAG | |
| Conservation | | | | | | | | | |
| Sequence logo (2.0bits / 0.0bits) | CCATGCCAAG | AGTATGGCTT | GCGGTATGCT | GTCCACATTT | TGTCCTGTAT | TTGTGGTACC | AAGTACTATT | ATTAGTGTAG | |

| | 3,940 | | 3,960 | | 3,980 | | 4,000 | | |
|---|---|---|---|---|---|---|---|---|---|
| Morex_Sbs1 | AAGTAGGTAT | TGGTTATTGC | AACGTATATG | CCCTATGTTT | GTTACGGAAT | ATTTGACCTT | TGTTTCTTGT | GTCTTGTTGC | 2804 |
| NDB112_Sbs1 | AAGTAGGTAT | TGGTTATTGC | AACGTATATG | CCCTATGTTT | GTTACGGAAT | ATTTGACCTT | TGTTTCTTGT | GTCTTGTTGC | 2804 |
| Robust_Sbs1 | AAGTAGGTAT | TGGTTATTGC | AACGTATATG | CCCTATGTTT | GTTACGGAAT | ATTTGACCTT | TGTTTCTTGT | GTCTTGTTGC | 2804 |
| Tradition_Sbs1 | AAGTAGGTAT | TGGTTATTGC | AACGTATATG | CCCTATGTTT | GTTACGGAAT | ATTTGACCTT | TGTTTCTTGT | GTCTTGTTGC | 2804 |
| Pinnacle_Sbs1 | AAGTAGGTAT | TGGTTATTGC | AACGTATATG | CCCTATGTTT | GTTACGGAAT | ATTTGACCTT | TGTTTCTTGT | GTCTTGTTGC | 2804 |
| Bowman_Sbs1 | AAGTAGGTAT | TGGTTATTGC | AACGTATATG | CCCTATGTTT | GTTACGGAAT | ATTTGACCTT | TGTTTCTTGT | GTCTTGTTGC | 3991 |
| ND_Genesis_Sbs1 | AAGTAGGTAT | TGGTTATTGC | AACGTATATG | CCCTATGTTT | GTTACGGAAT | ATTTGACCTT | TGTTTCTTGT | GTCTTGTTGC | 3991 |
| TR306_Sbs1 | AAGTAGGTAT | TGGTTATTGC | AACGTATATG | CCCTATGTTT | GTTACGGAAT | ATTTGACCTT | TGTTTCTTGT | GTCTTGTTGC | 3991 |
| Steptoe_Sbs1 | AAGTAGGTAT | TGGTTATTGC | AACGTATATG | CCCTATGTTT | GTTACGGAAT | ATTTGACCTT | TGTTTCTTGT | GTCTTGTTGC | 3988 |
| Harrington_Sbs1 | AAGTAGGTAT | TGGTTATTGC | AACGTATATG | CCCTATGTTT | GTTACGGAAT | ATTTGACCTT | TGTTTCTTGT | GTCTTGTTGC | 3986 |
| Consensus | AAGTAGGTAT | TGGTTATTGC | AACGTATATG | CCCTATGTTT | GTTACGGAAT | ATTTGACCTT | TGTTTCTTGT | GTCTTGTTGC | |
| Conservation | | | | | | | | | |
| Sequence logo (2.0bits / 0.0bits) | AAGTAGGTAT | TGGTTATTGC | AACGTATATG | CCCTATGTTT | GTTACGGAAT | ATTTGACCTT | TGTTTCTTGT | GTCTTGTTGC | |

### Supplementary Figure 5A

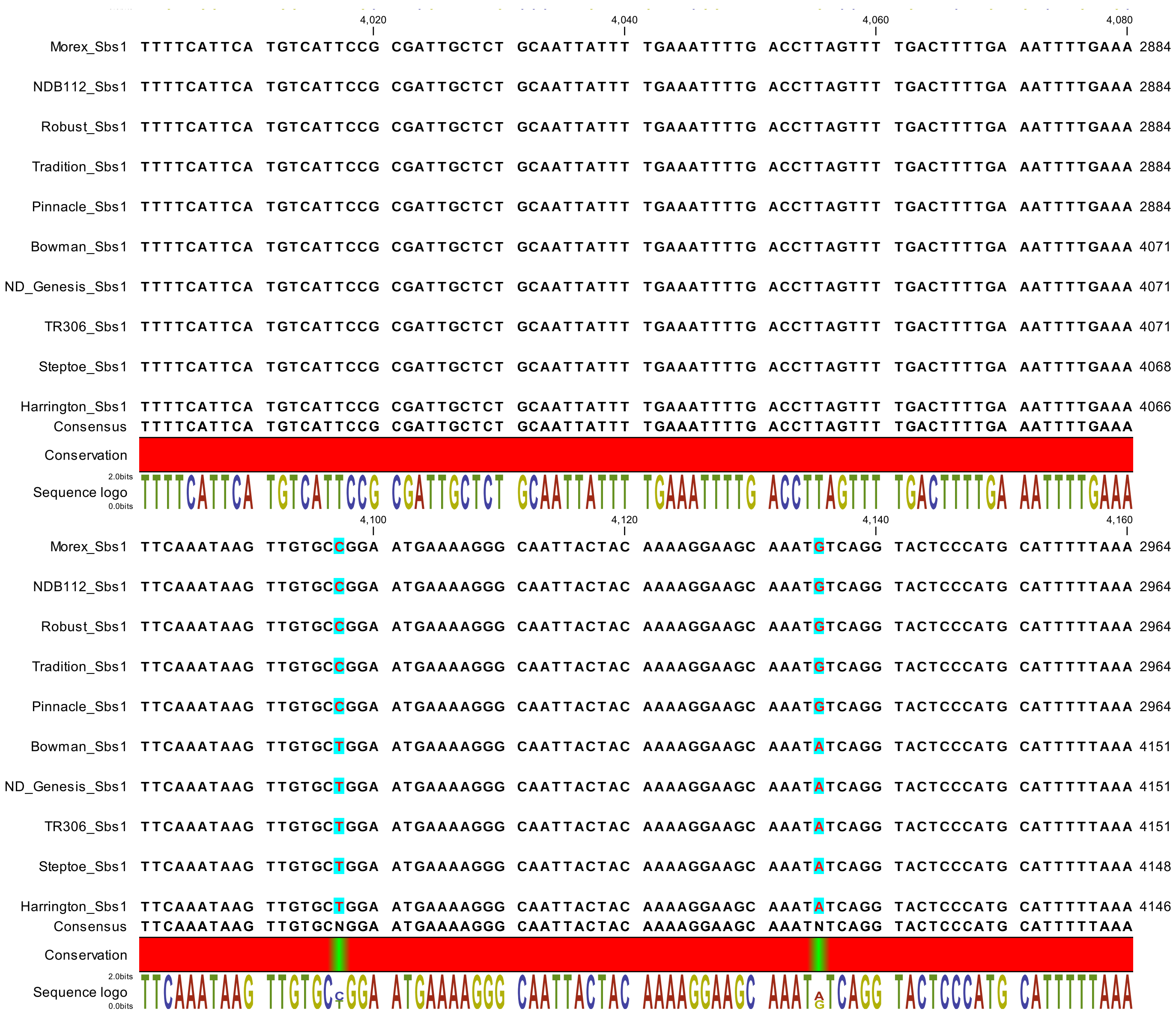

Supplementary Figure 5A

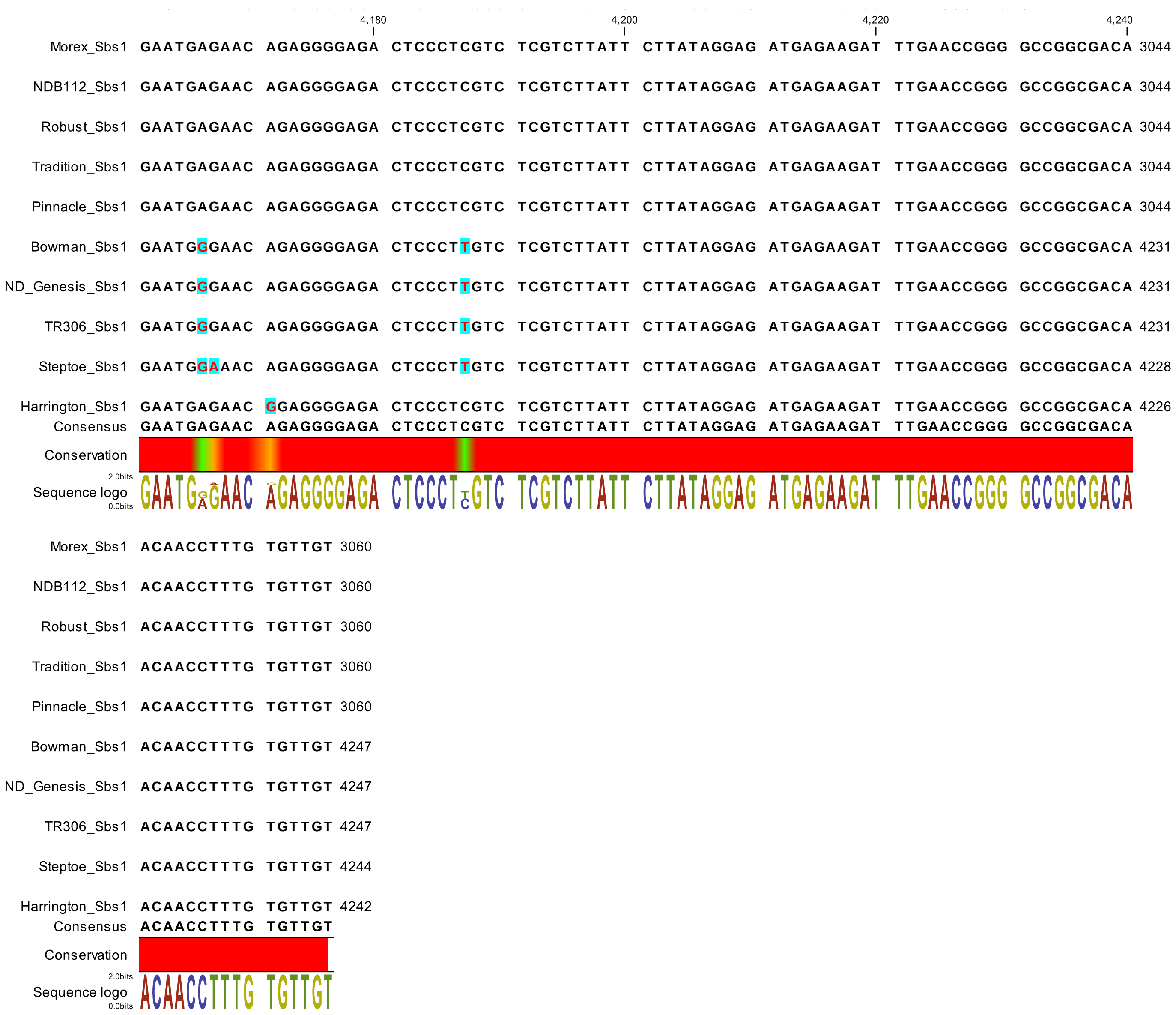

### Supplementary Figure 5B

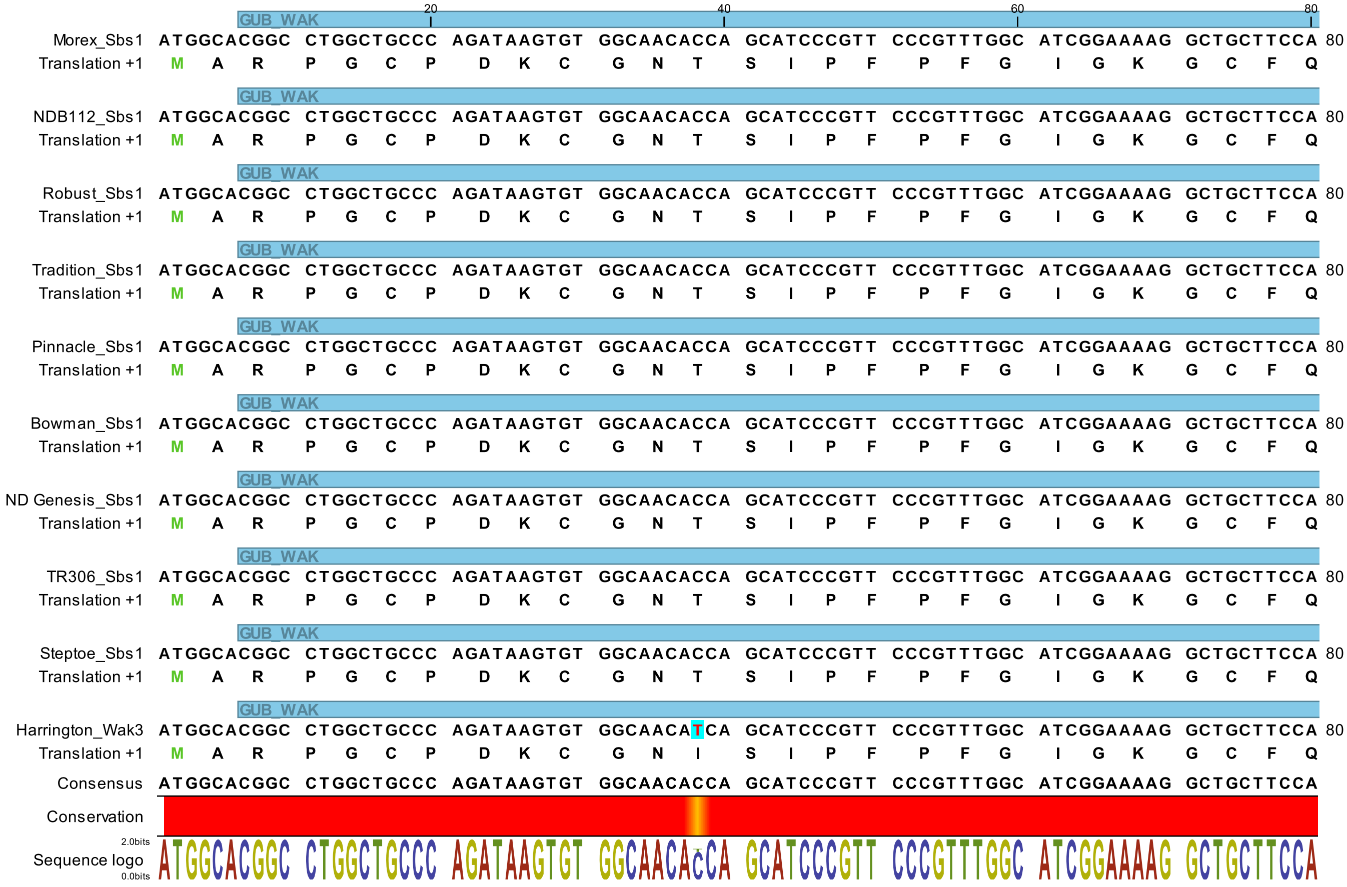

Supplementary Figure 5B

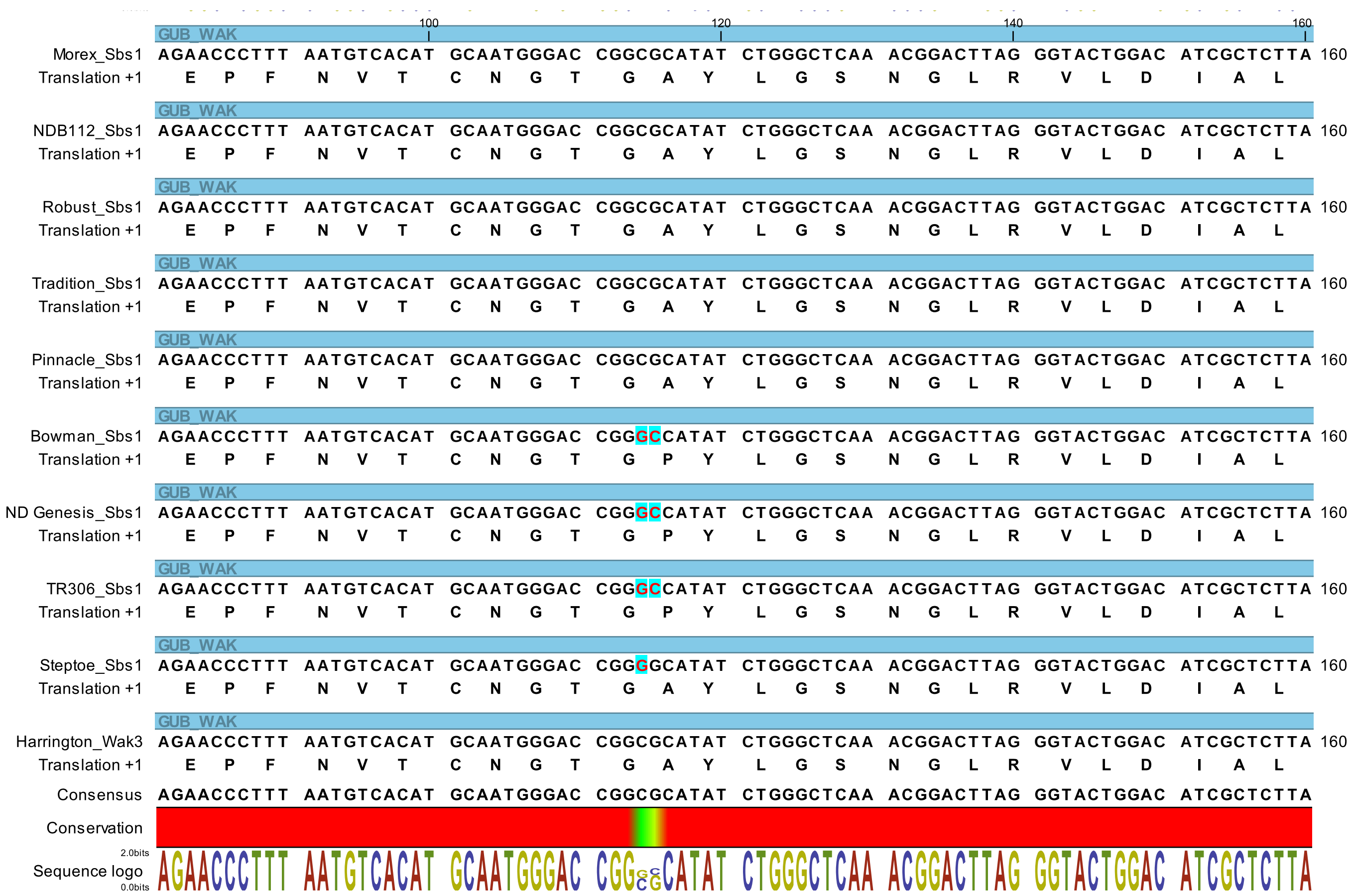

### Supplementary Figure 5B

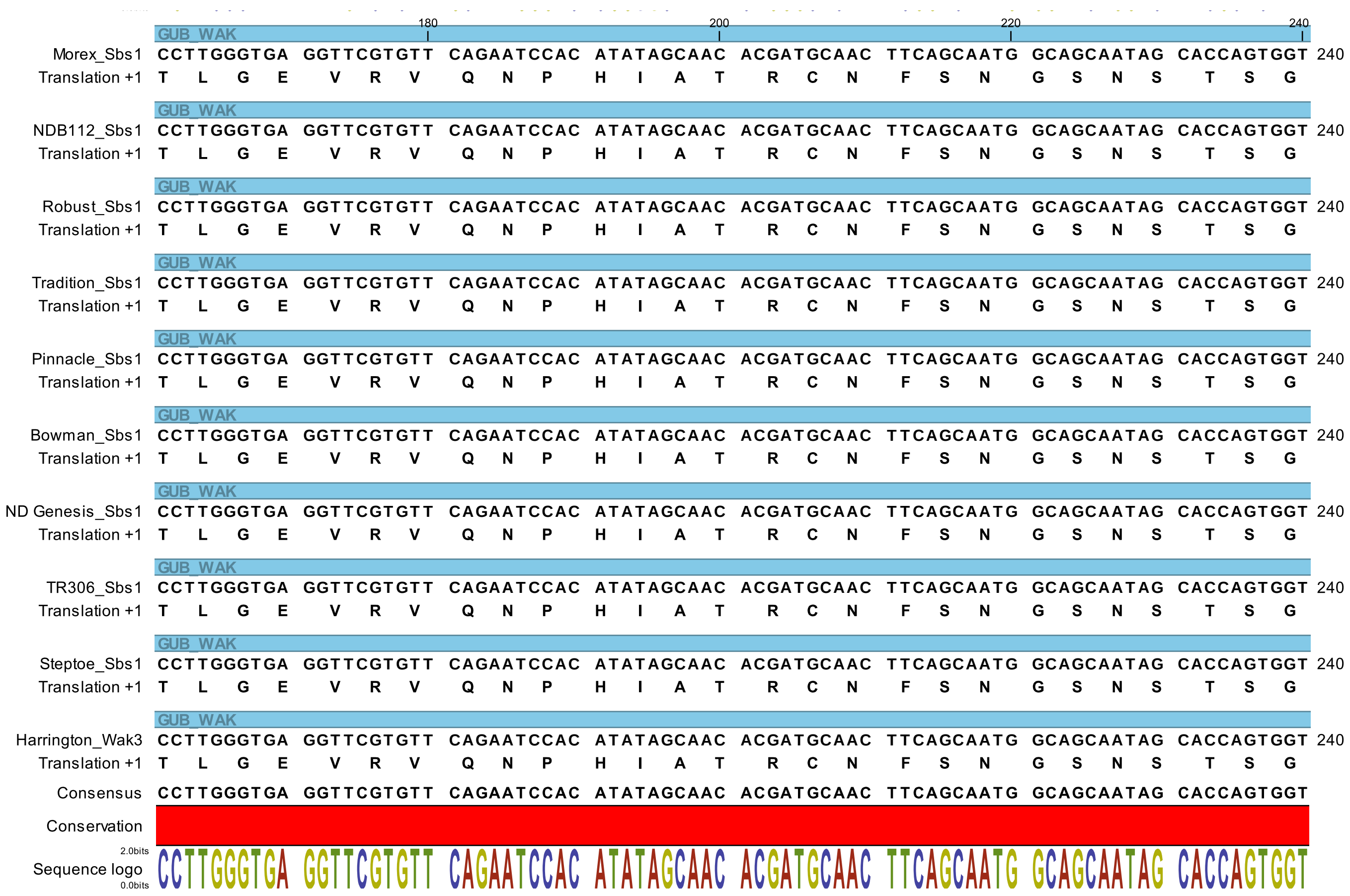

### Supplementary Figure 5B

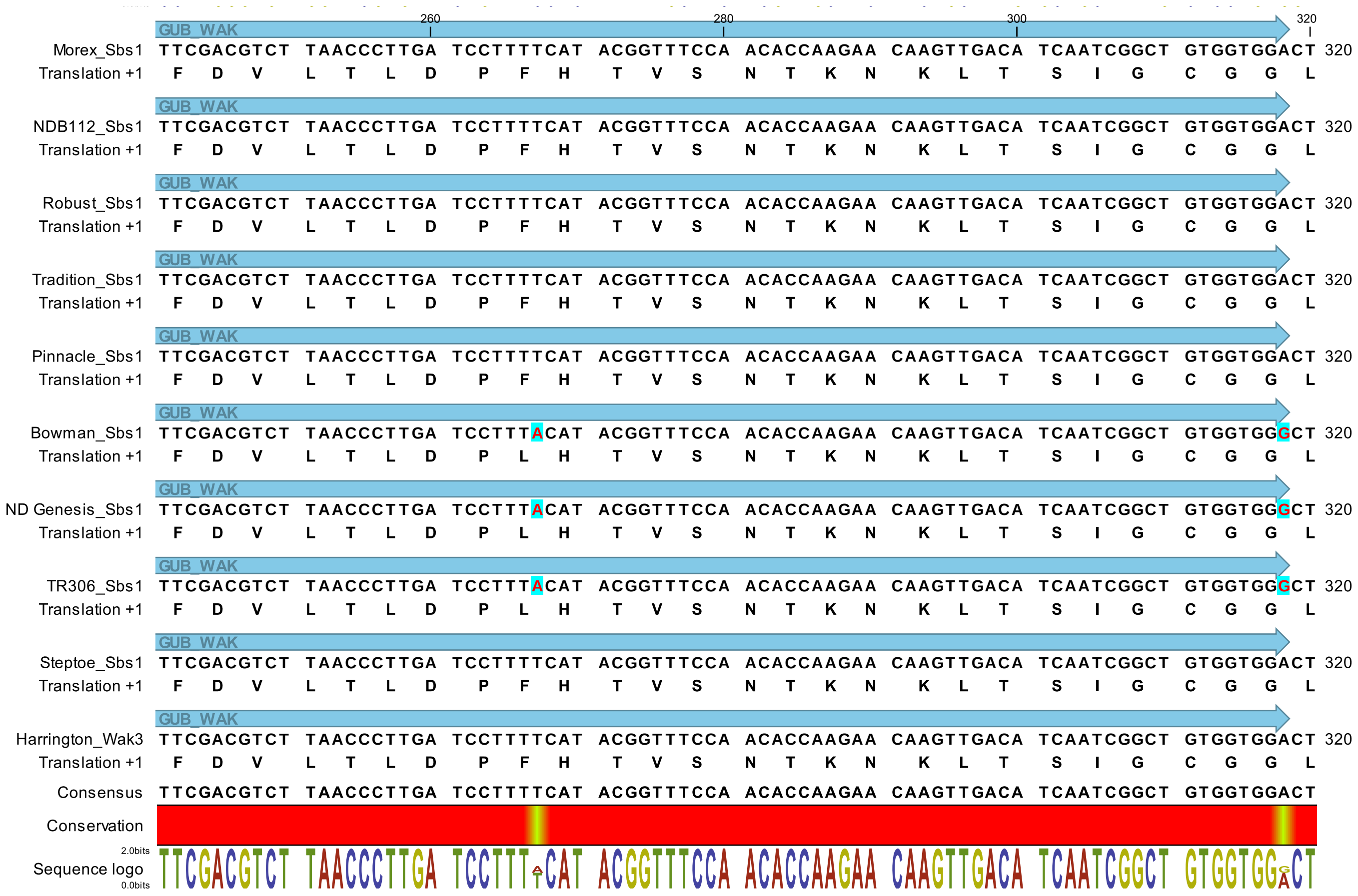

### Supplementary Figure 5B

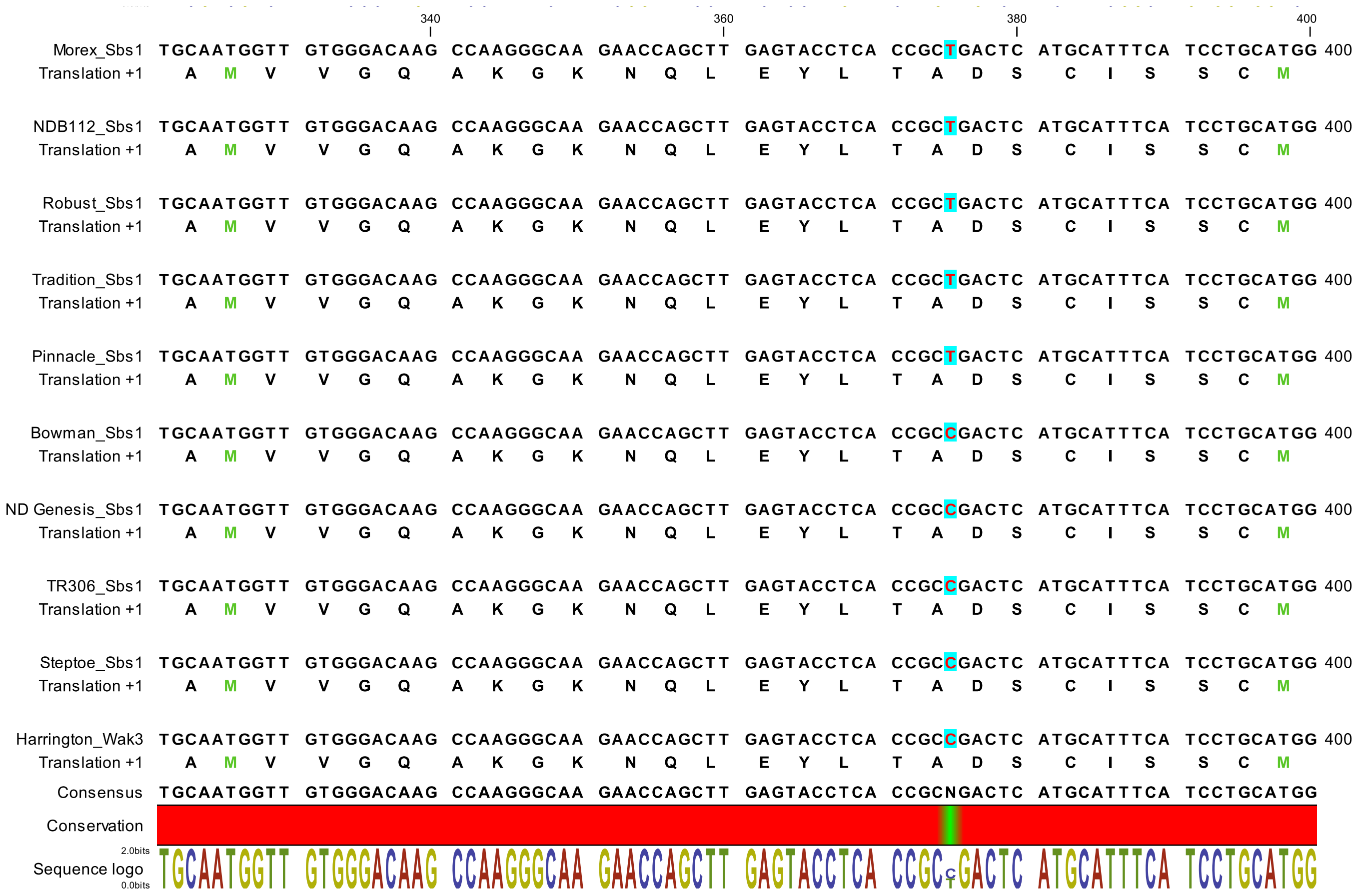

### Supplementary Figure 5B

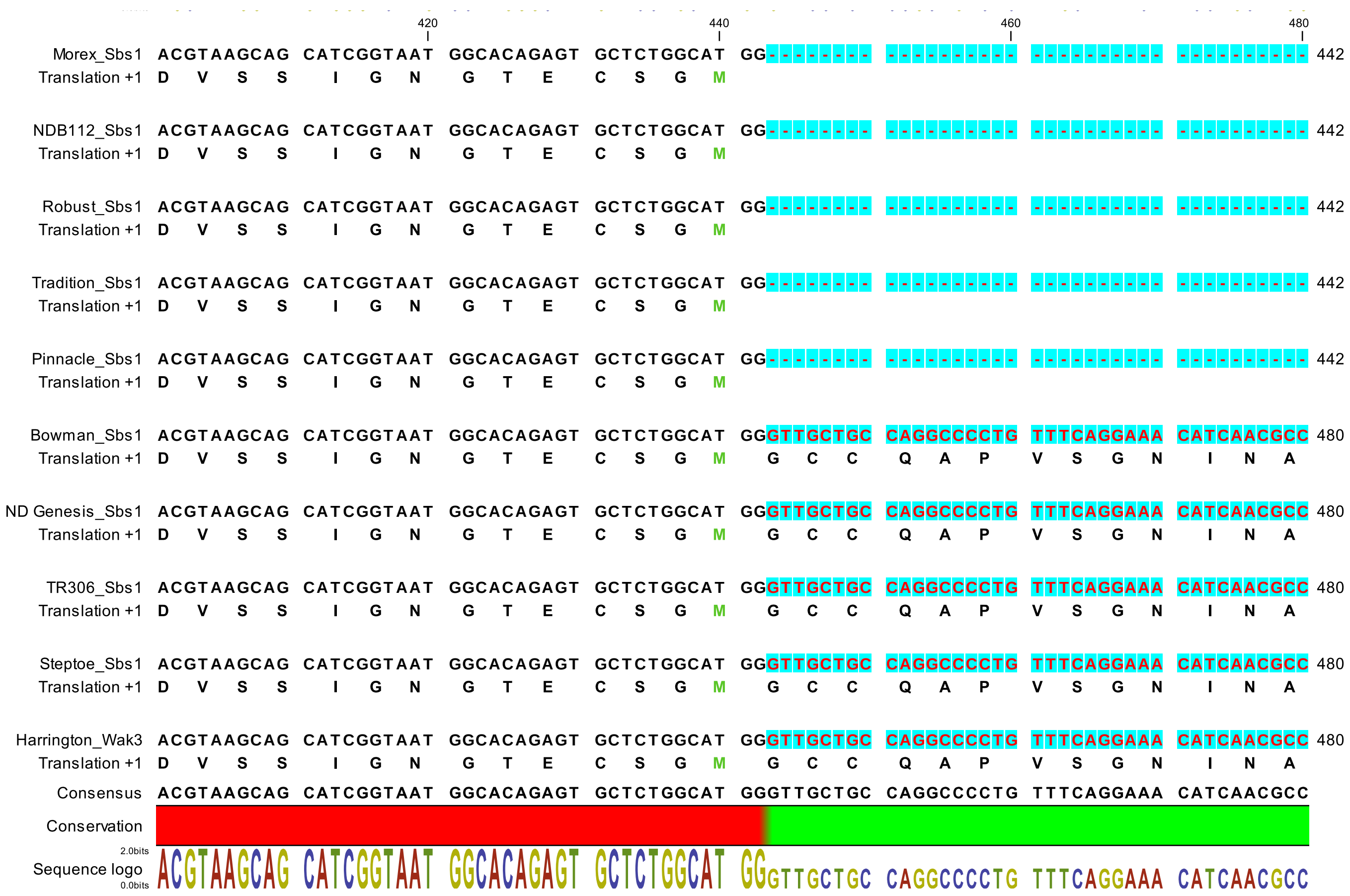

### Supplementary Figure 5B

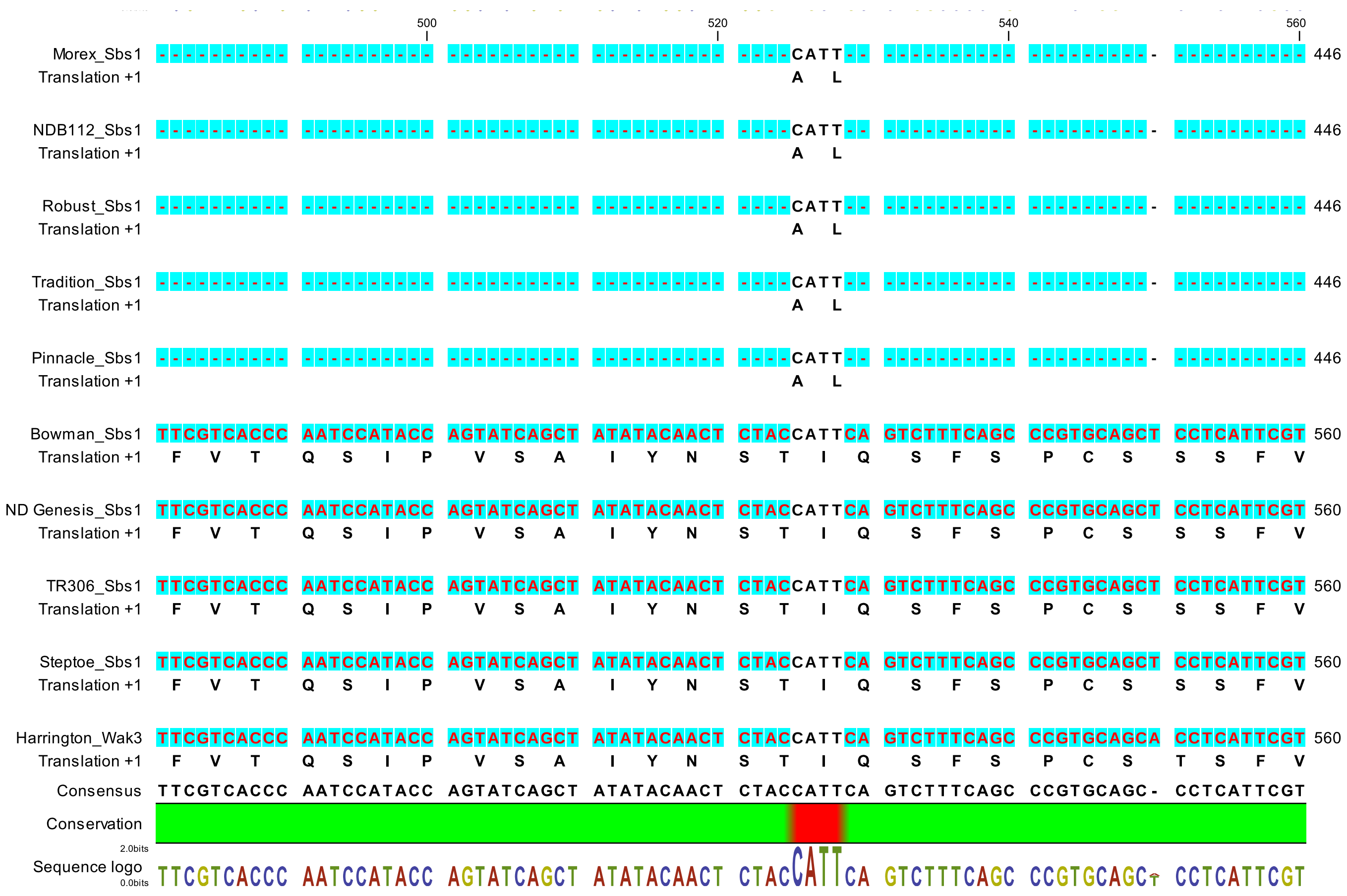

### Supplementary Figure 5B

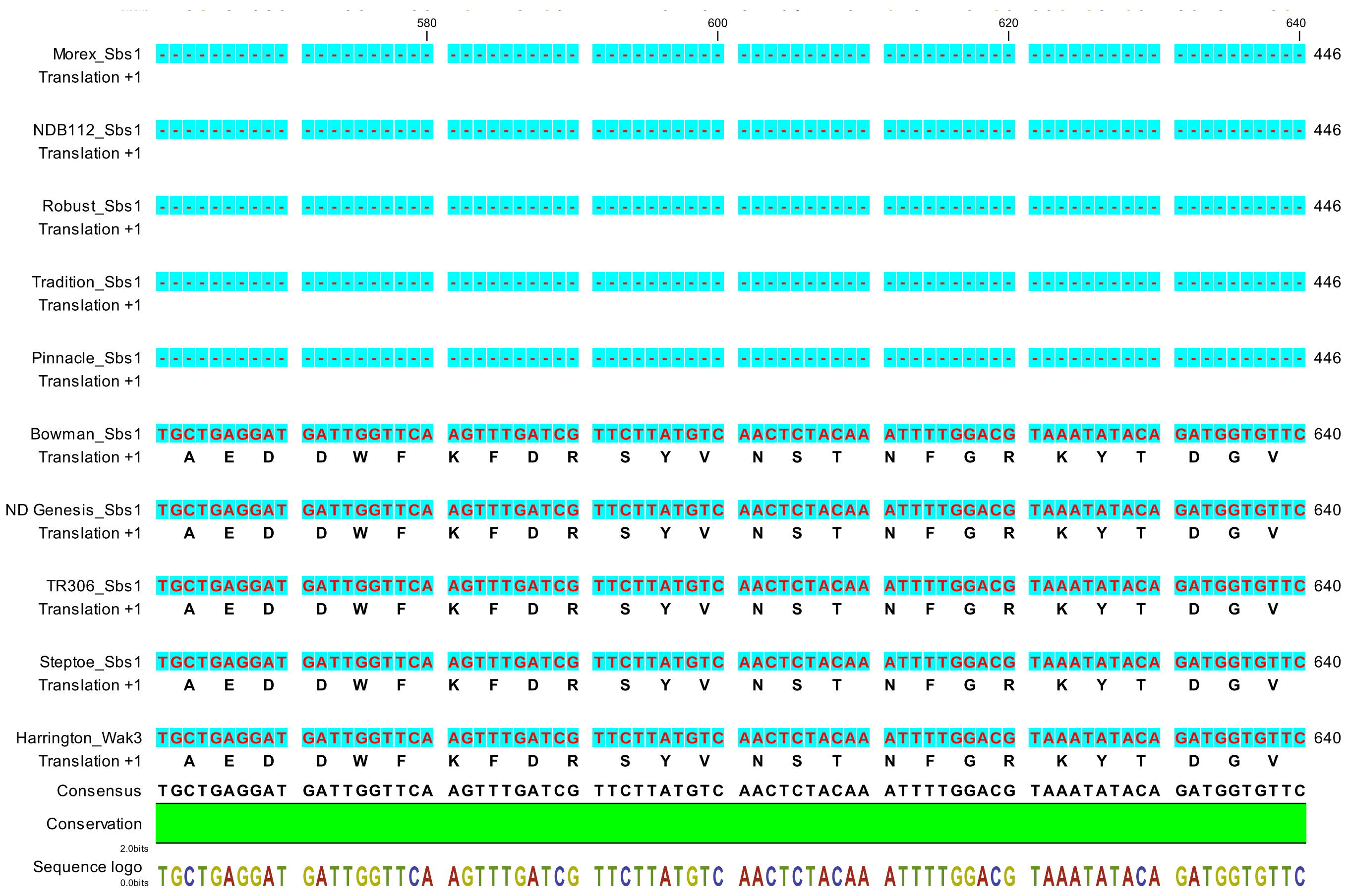

Supplementary Figure 5B

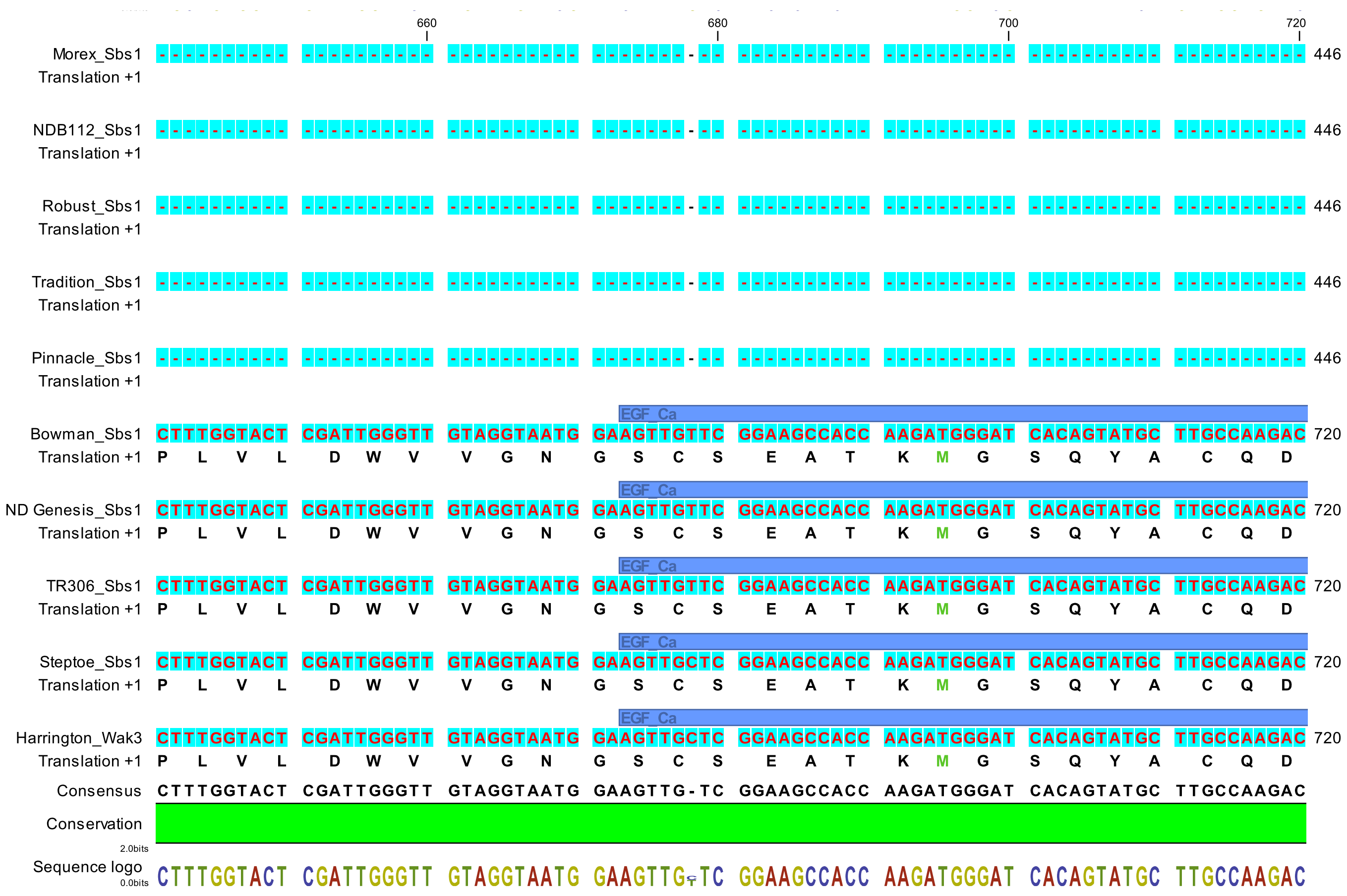

### Supplementary Figure 5B

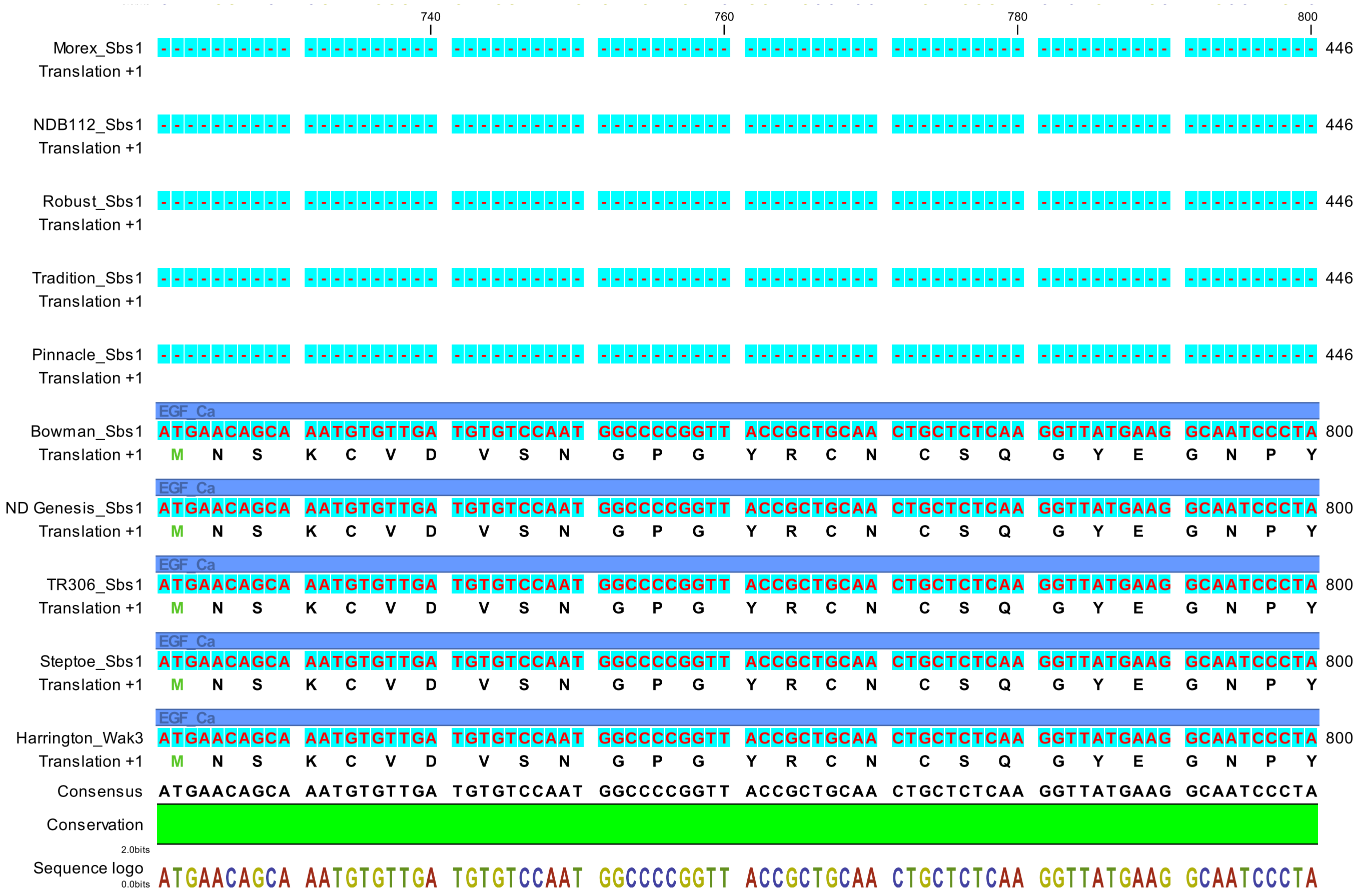

### Supplementary Figure 5B

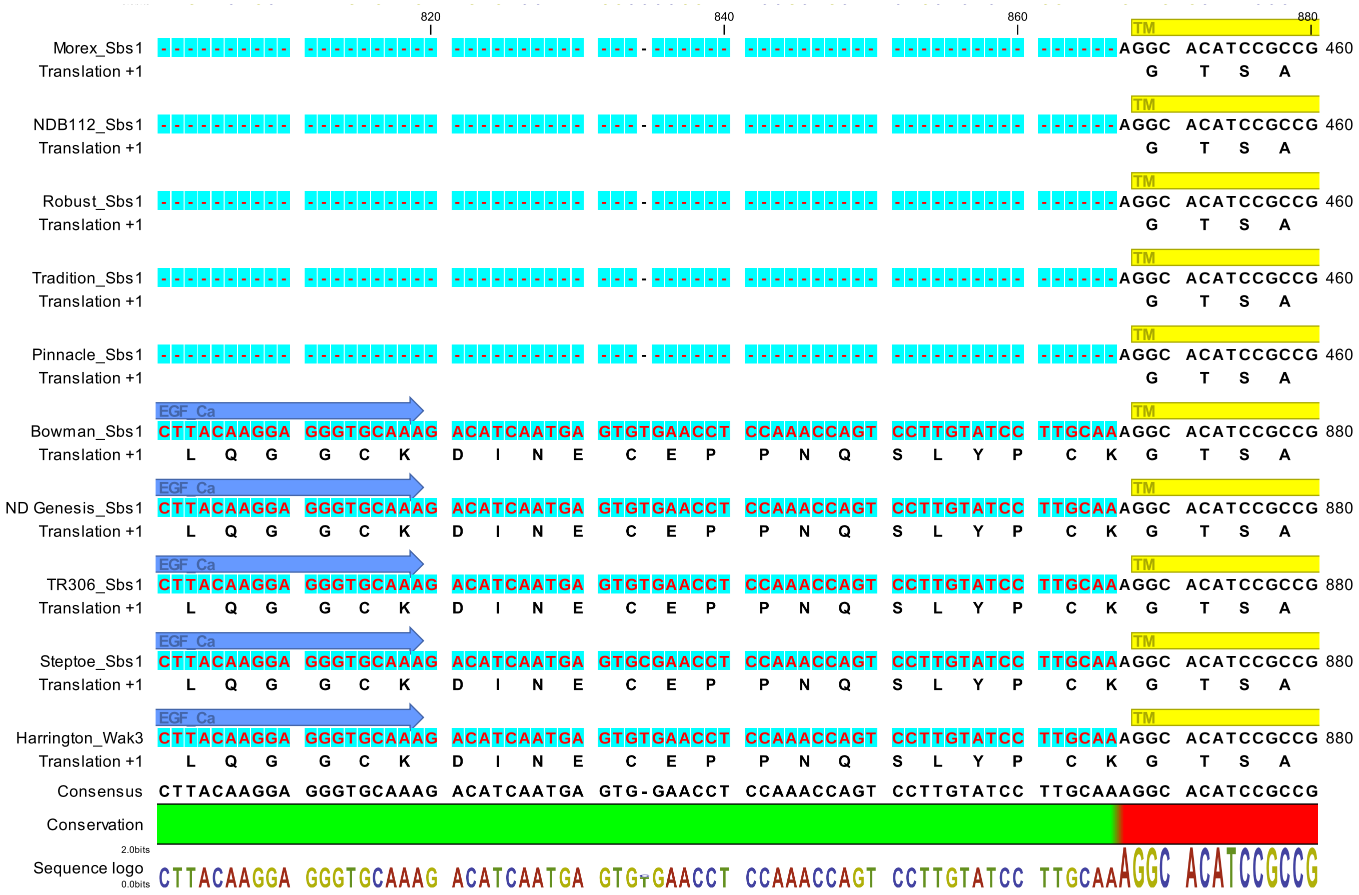

### Supplementary Figure 5B

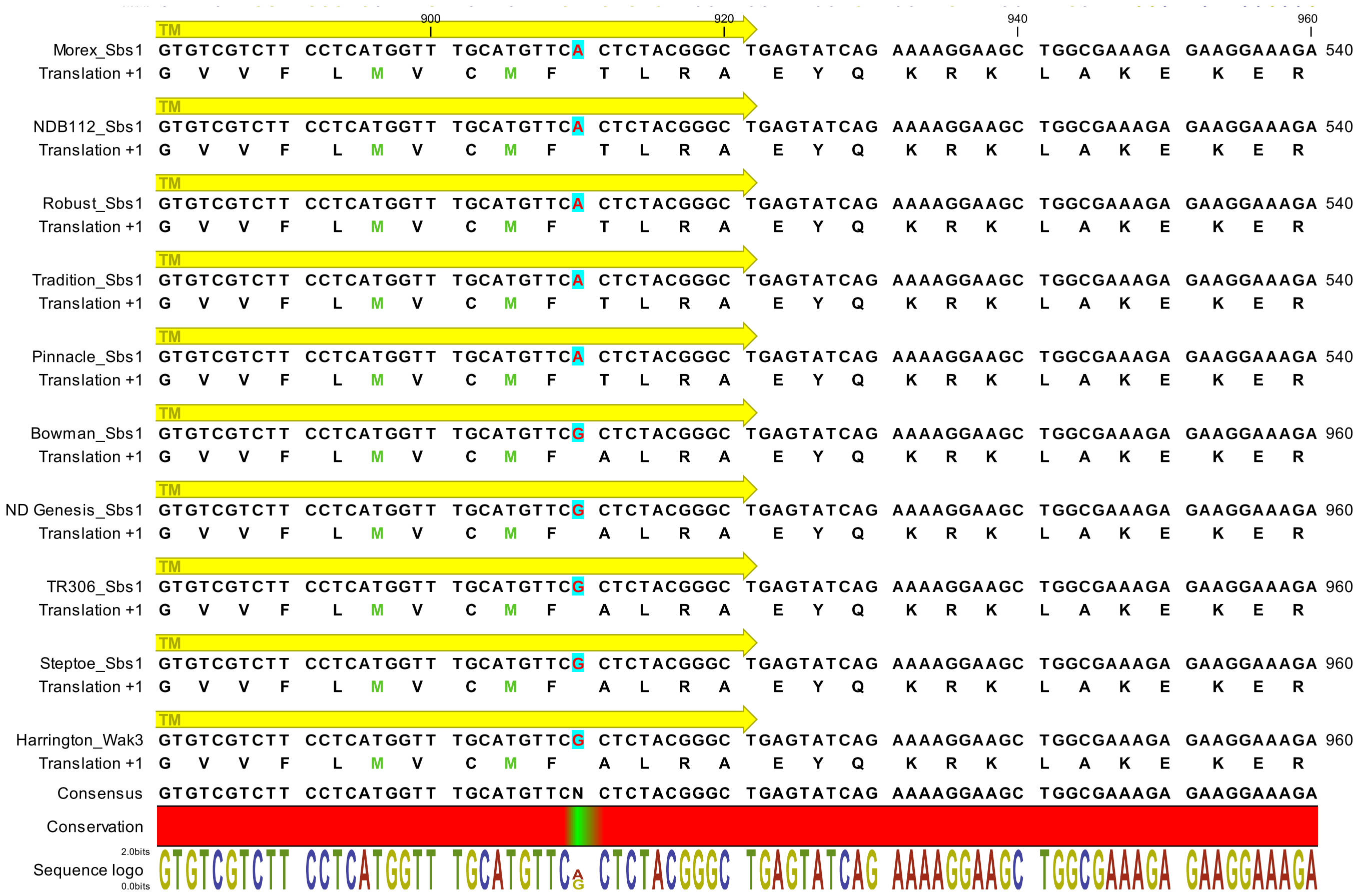

### Supplementary Figure 5B

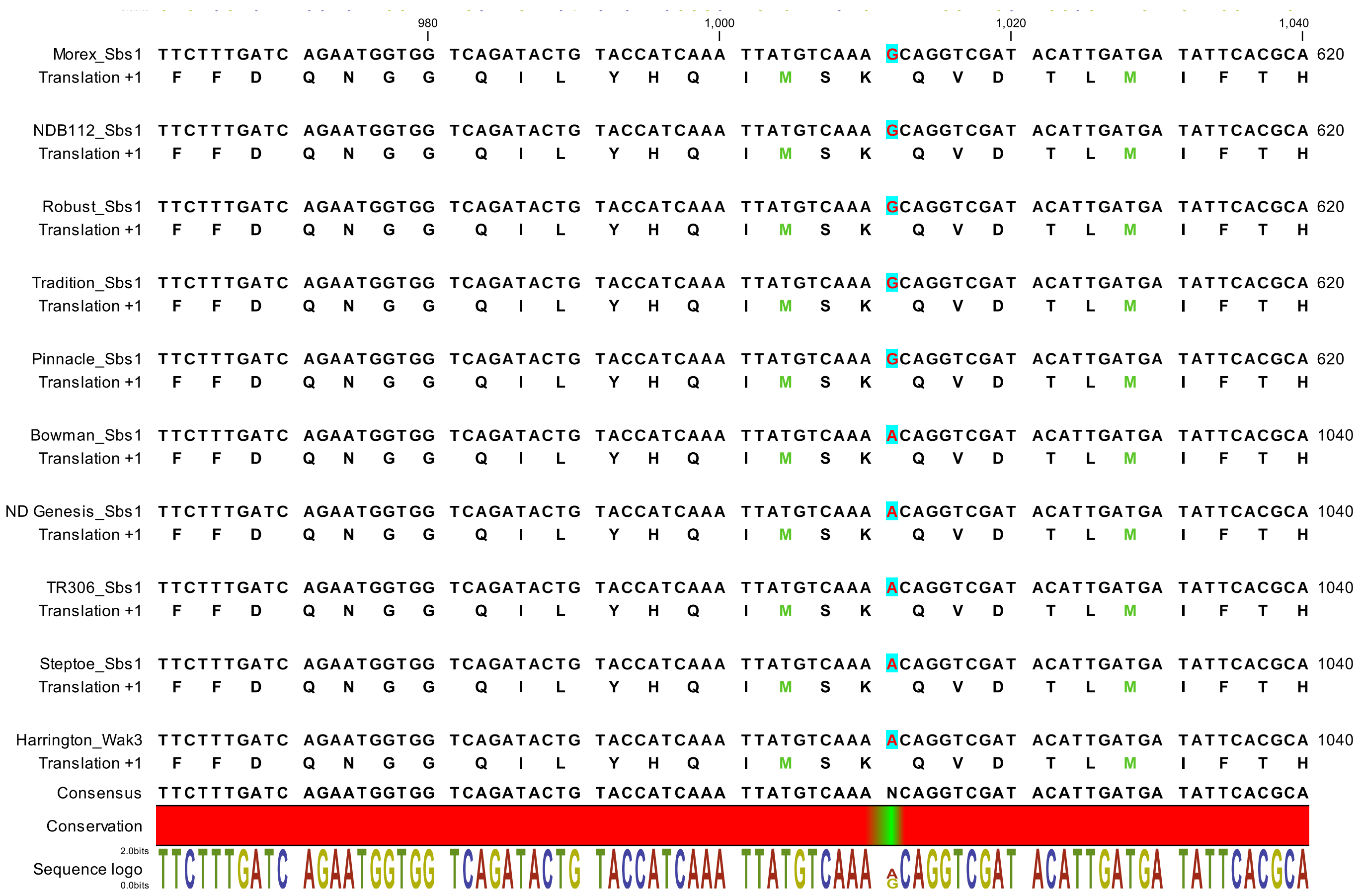

### Supplementary Figure 5B

```
                                              1,060                 1,080                 1,100                 1,120
                                                         STKc_IRAK
Morex_Sbs1        TGAAGATCTA  AAGAAGGCCA  CGAATGATTT  TGATGAGAGC  AGAGAAGTGG  GCAGGGGTGG  TCATGGCACT  GTCTACAAGG  700
Translation +1      E  D  L     K  K  A     T  N  D  F   D  E  S     R  E  V     G  R  G  G   H  G  T     V  Y  K
                                                         STKc_IRAK
NDB112_Sbs1       TGAAGATCTA  AAGAAGGCCA  CGAATGATTT  TGATGAGAGC  AGAGAAGTGG  GCAGGGGTGG  TCATGGCACT  GTCTACAAGG  700
Translation +1      E  D  L     K  K  A     T  N  D  F   D  E  S     R  E  V     G  R  G  G   H  G  T     V  Y  K
                                                         STKc_IRAK
Robust_Sbs1       TGAAGATCTA  AAGAAGGCCA  CGAATGATTT  TGATGAGAGC  AGAGAAGTGG  GCAGGGGTGG  TCATGGCACT  GTCTACAAGG  700
Translation +1      E  D  L     K  K  A     T  N  D  F   D  E  S     R  E  V     G  R  G  G   H  G  T     V  Y  K
                                                         STKc_IRAK
Tradition_Sbs1    TGAAGATCTA  AAGAAGGCCA  CGAATGATTT  TGATGAGAGC  AGAGAAGTGG  GCAGGGGTGG  TCATGGCACT  GTCTACAAGG  700
Translation +1      E  D  L     K  K  A     T  N  D  F   D  E  S     R  E  V     G  R  G  G   H  G  T     V  Y  K
                                                         STKc_IRAK
Pinnacle_Sbs1     TGAAGATCTA  AAGAAGGCCA  CGAATGATTT  TGATGAGAGC  AGAGAAGTGG  GCAGGGGTGG  TCATGGCACT  GTCTACAAGG  700
Translation +1      E  D  L     K  K  A     T  N  D  F   D  E  S     R  E  V     G  R  G  G   H  G  T     V  Y  K
                                                         STKc_IRAK
Bowman_Sbs1       TGAAGATCTA  AAGAAGGCCA  CGAATGATTT  TGATGAGAGC  AGAGAAGTGG  GCAGGGGTGG  TCATGGCACT  GTCTACAAGG  1120
Translation +1      E  D  L     K  K  A     T  N  D  F   D  E  S     R  E  V     G  R  G  G   H  G  T     V  Y  K
                                                         STKc_IRAK
ND Genesis_Sbs1   TGAAGATCTA  AAGAAGGCCA  CGAATGATTT  TGATGAGAGC  AGAGAAGTGG  GCAGGGGTGG  TCATGGCACT  GTCTACAAGG  1120
Translation +1      E  D  L     K  K  A     T  N  D  F   D  E  S     R  E  V     G  R  G  G   H  G  T     V  Y  K
                                                         STKc_IRAK
TR306_Sbs1        TGAAGATCTA  AAGAAGGCCA  CGAATGATTT  TGATGAGAGC  AGAGAAGTGG  GCAGGGGTGG  TCATGGCACT  GTCTACAAGG  1120
Translation +1      E  D  L     K  K  A     T  N  D  F   D  E  S     R  E  V     G  R  G  G   H  G  T     V  Y  K
                                                         STKc_IRAK
Steptoe_Sbs1      TGAAGATCTA  AAGAAGGCCA  CGAATGATTT  TGATGAGAGC  AGAGAAGTGG  GCAGGGGTGG  TCATGGCACT  GTCTACAAGG  1120
Translation +1      E  D  L     K  K  A     T  N  D  F   D  E  S     R  E  V     G  R  G  G   H  G  T     V  Y  K
                                                         STKc_IRAK
Harrington_Wak3   TGAAGATCTA  AAGAAGGCCA  CGAATGATTT  TGATGAGAGC  AGAGAAGTGG  GCAGGGGTGG  TCATGGCACT  GTCTACAAGG  1120
Translation +1      E  D  L     K  K  A     T  N  D  F   D  E  S     R  E  V     G  R  G  G   H  G  T     V  Y  K
Consensus         TGAAGATCTA  AAGAAGGCCA  CGAATGATTT  TGATGAGAGC  AGAGAAGTGG  GCAGGGGTGG  TCATGGCACT  GTCTACAAGG
```

Conservation

Sequence logo

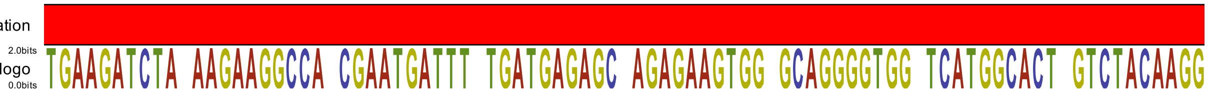

### Supplementary Figure 5B

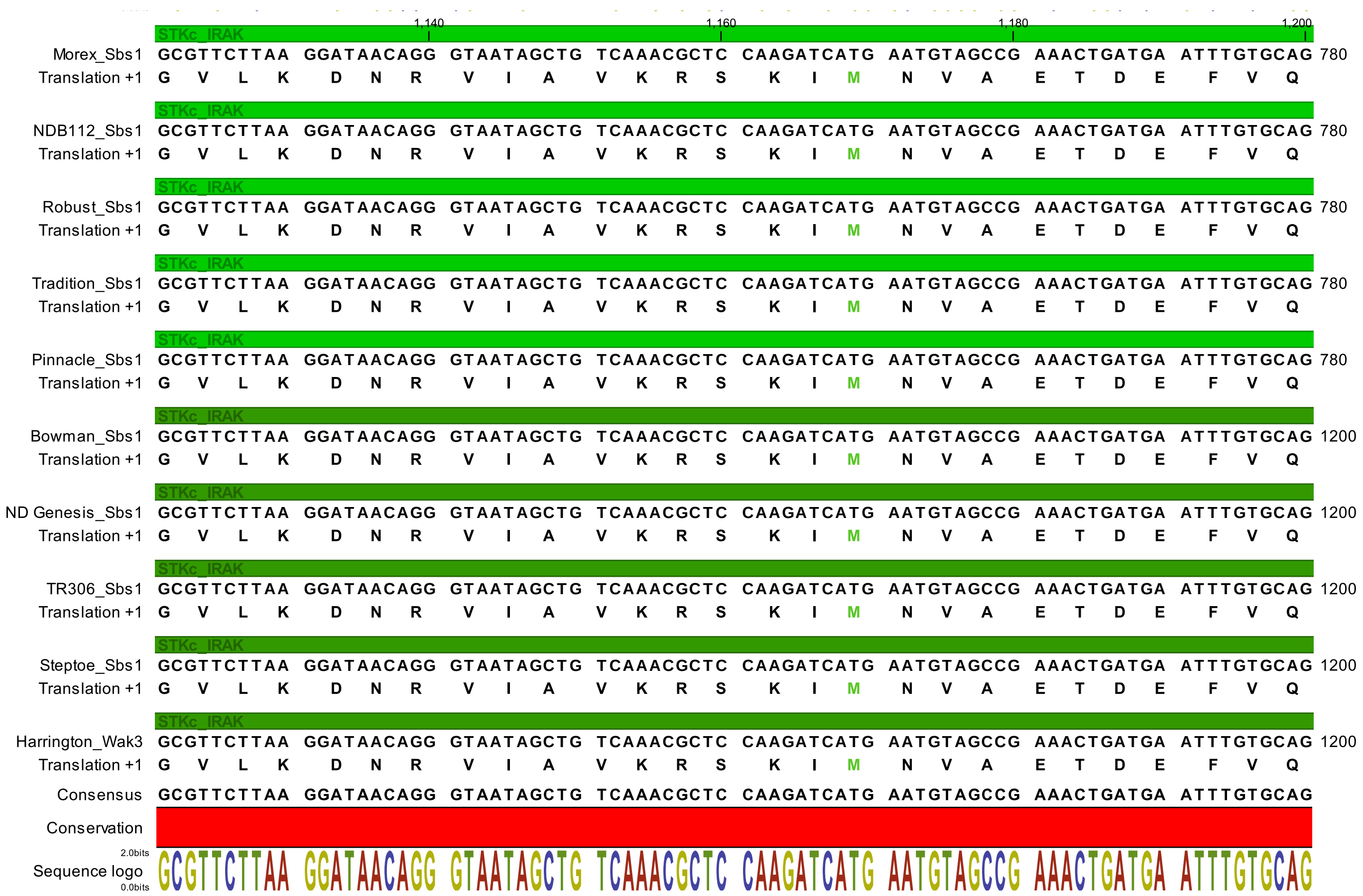

### Supplementary Figure 5B

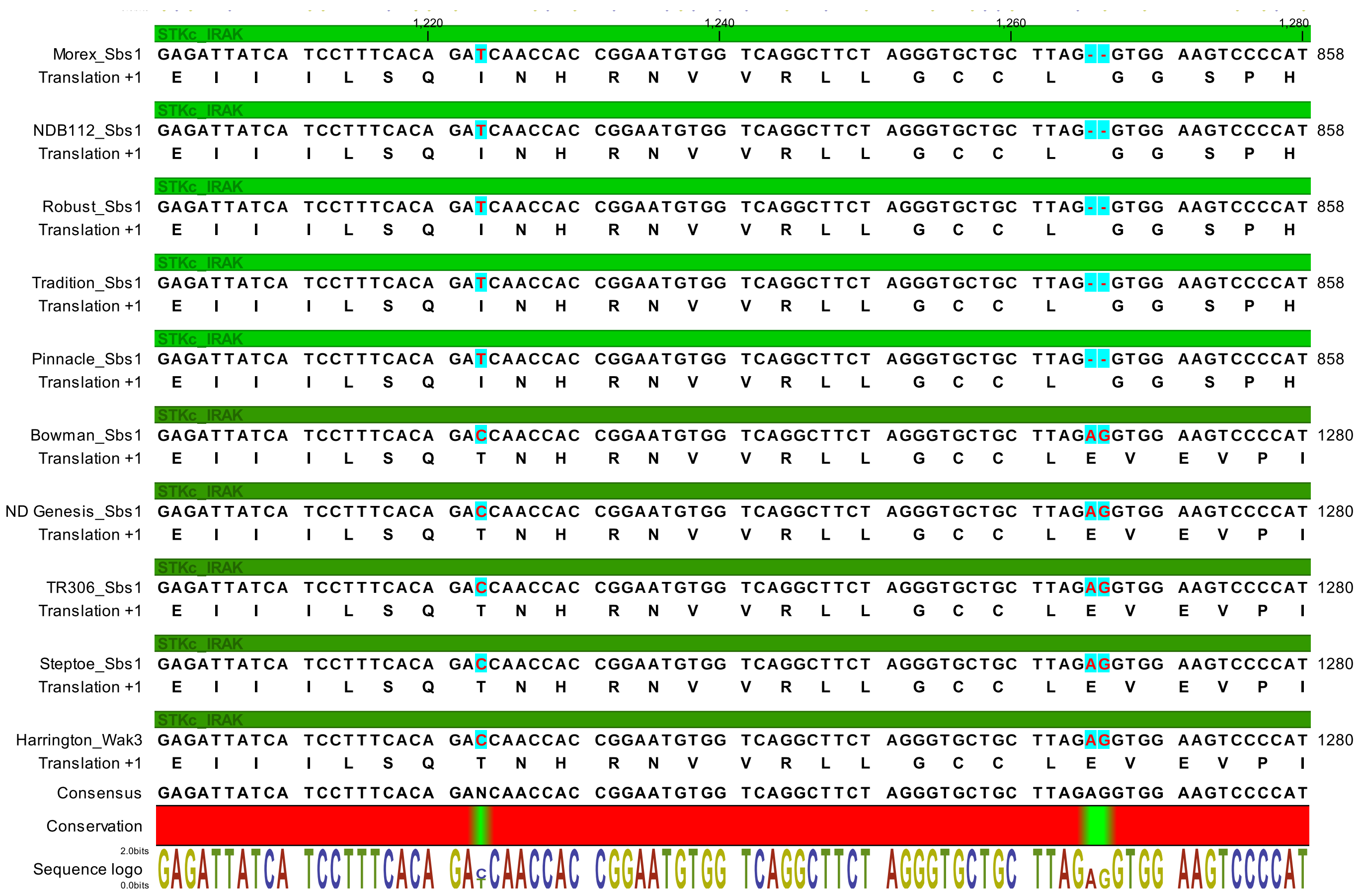

### Supplementary Figure 5B

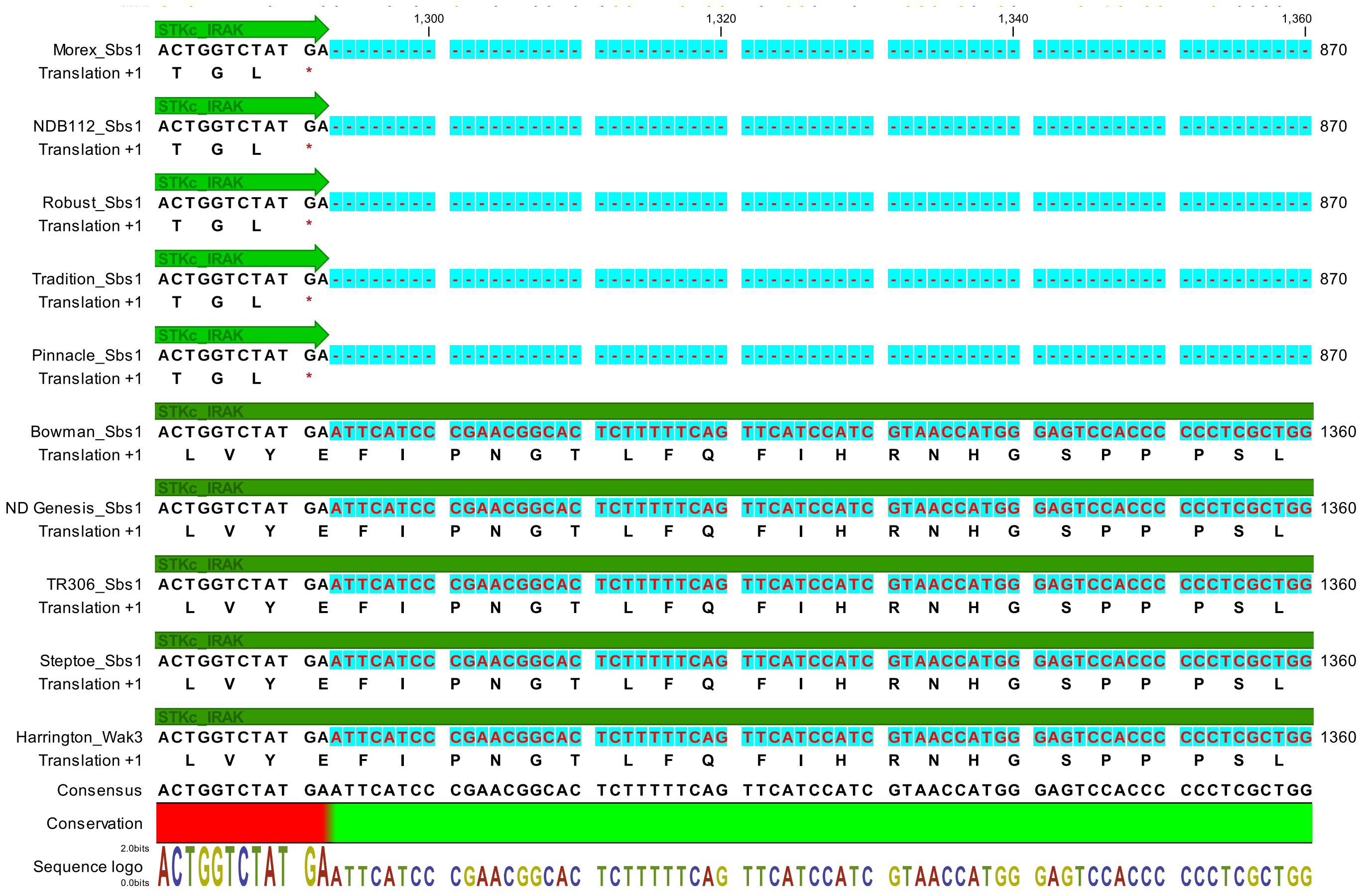

### Supplementary Figure 5B

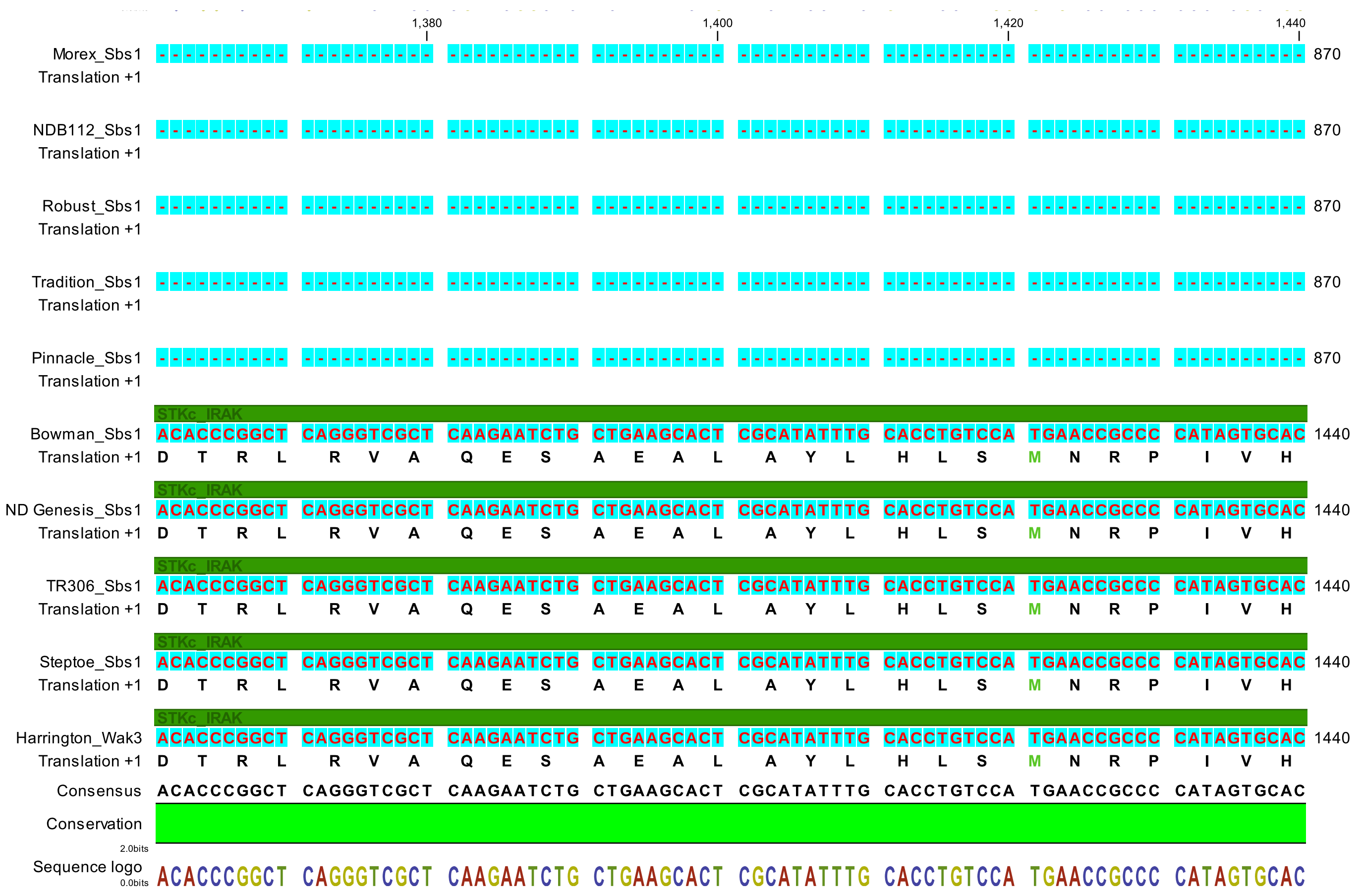

### Supplementary Figure 5B

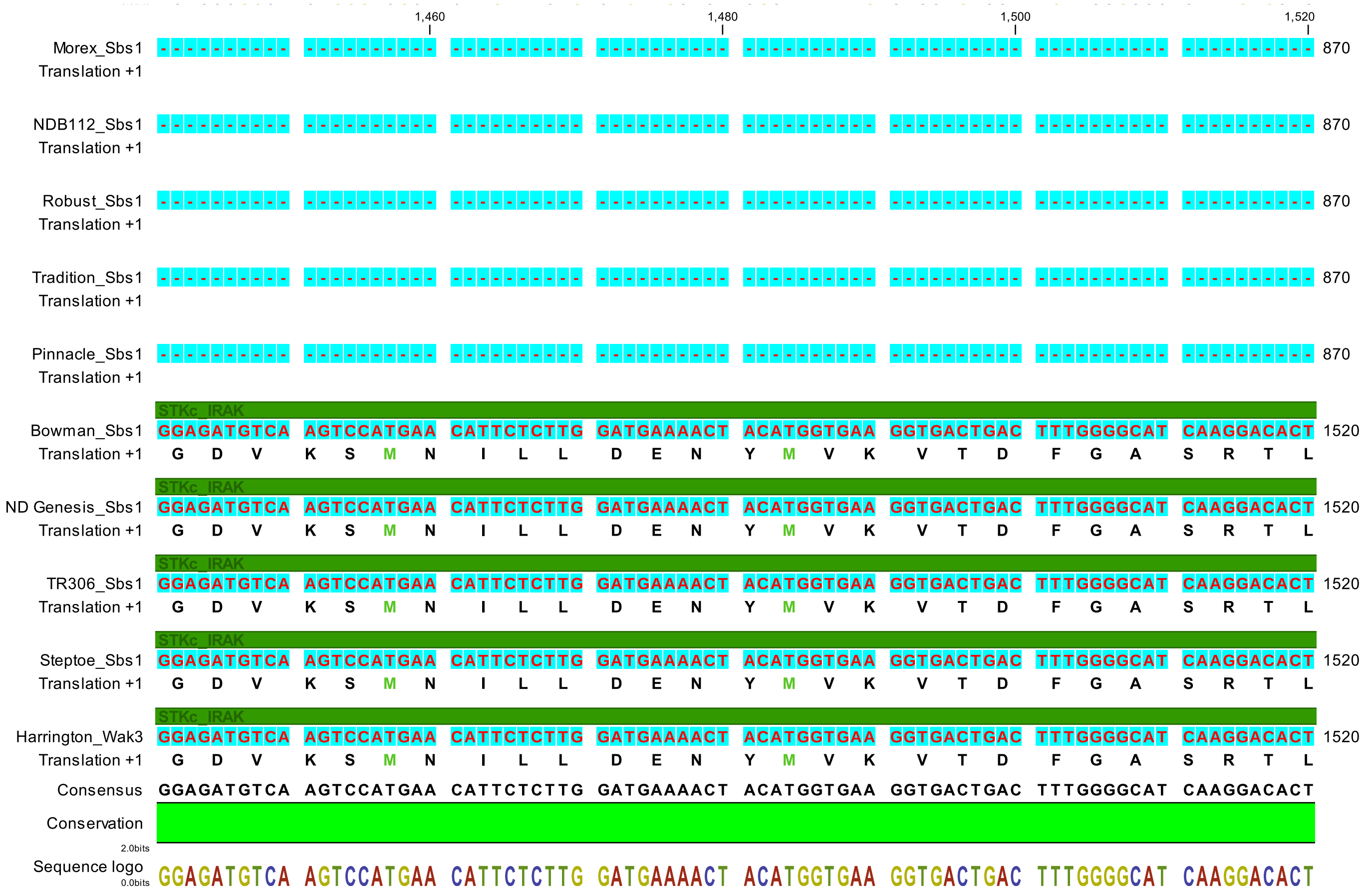

Supplementary Figure 5B

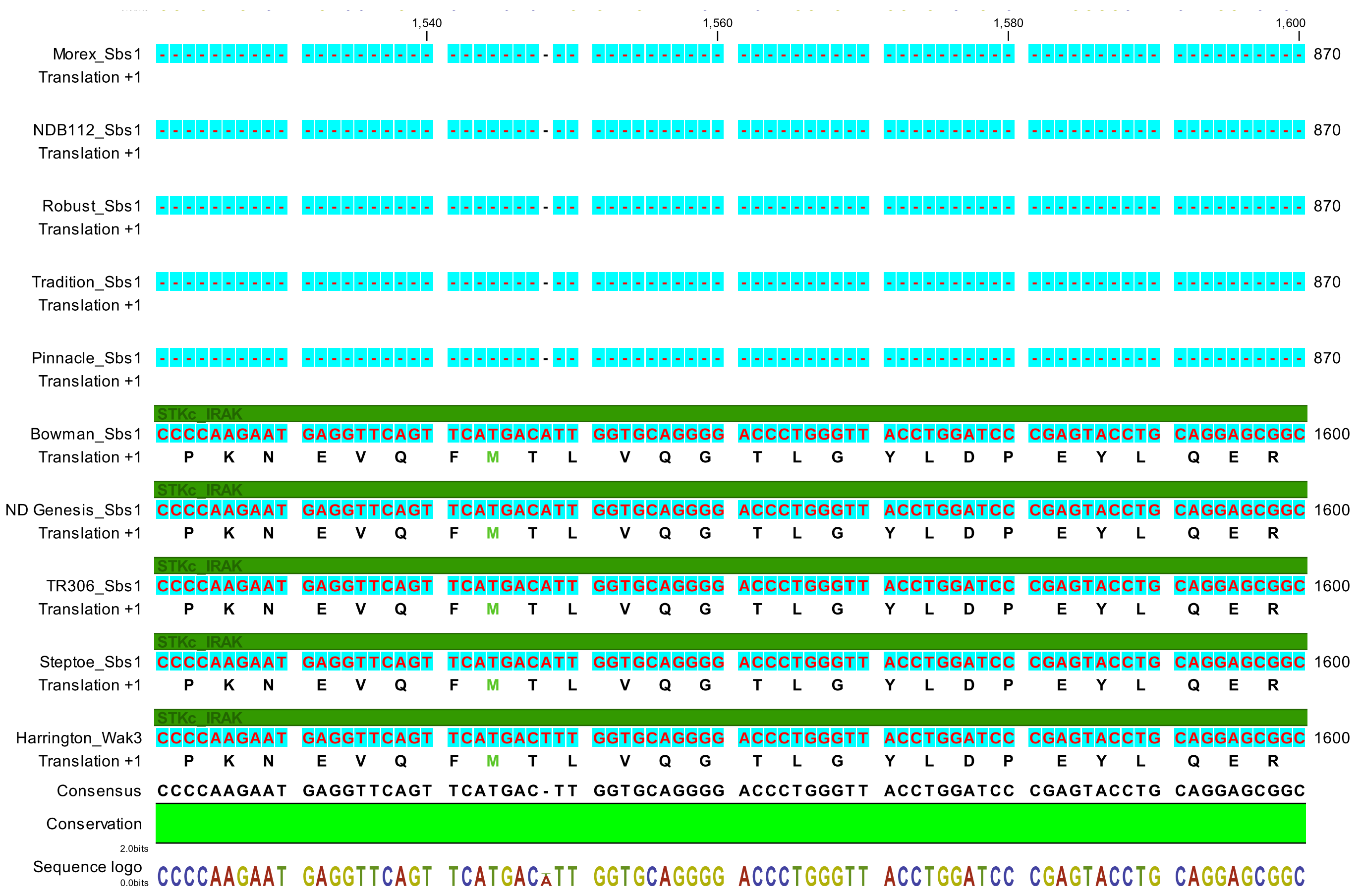

### Supplementary Figure 5B

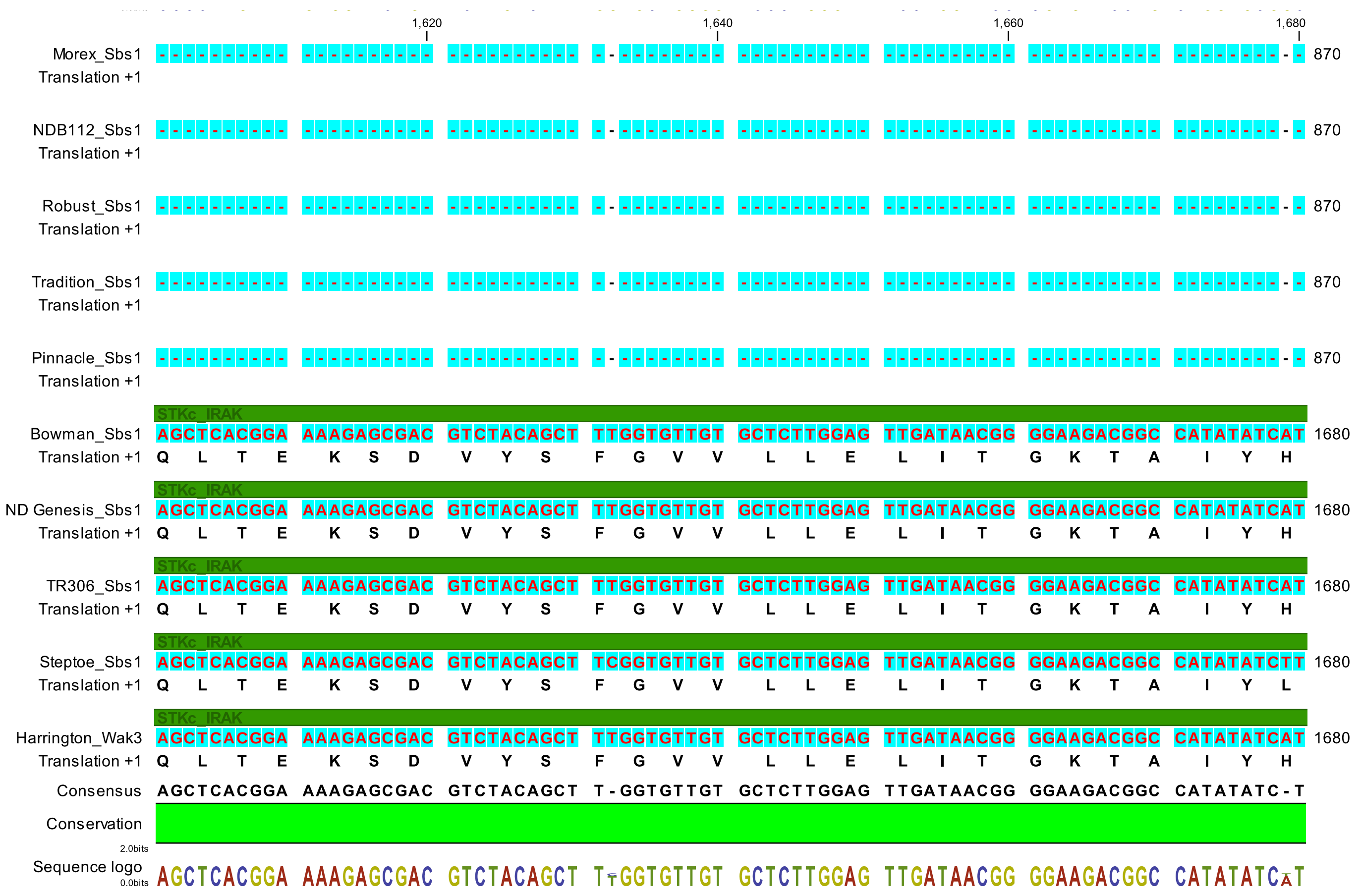

Supplementary Figure 5B

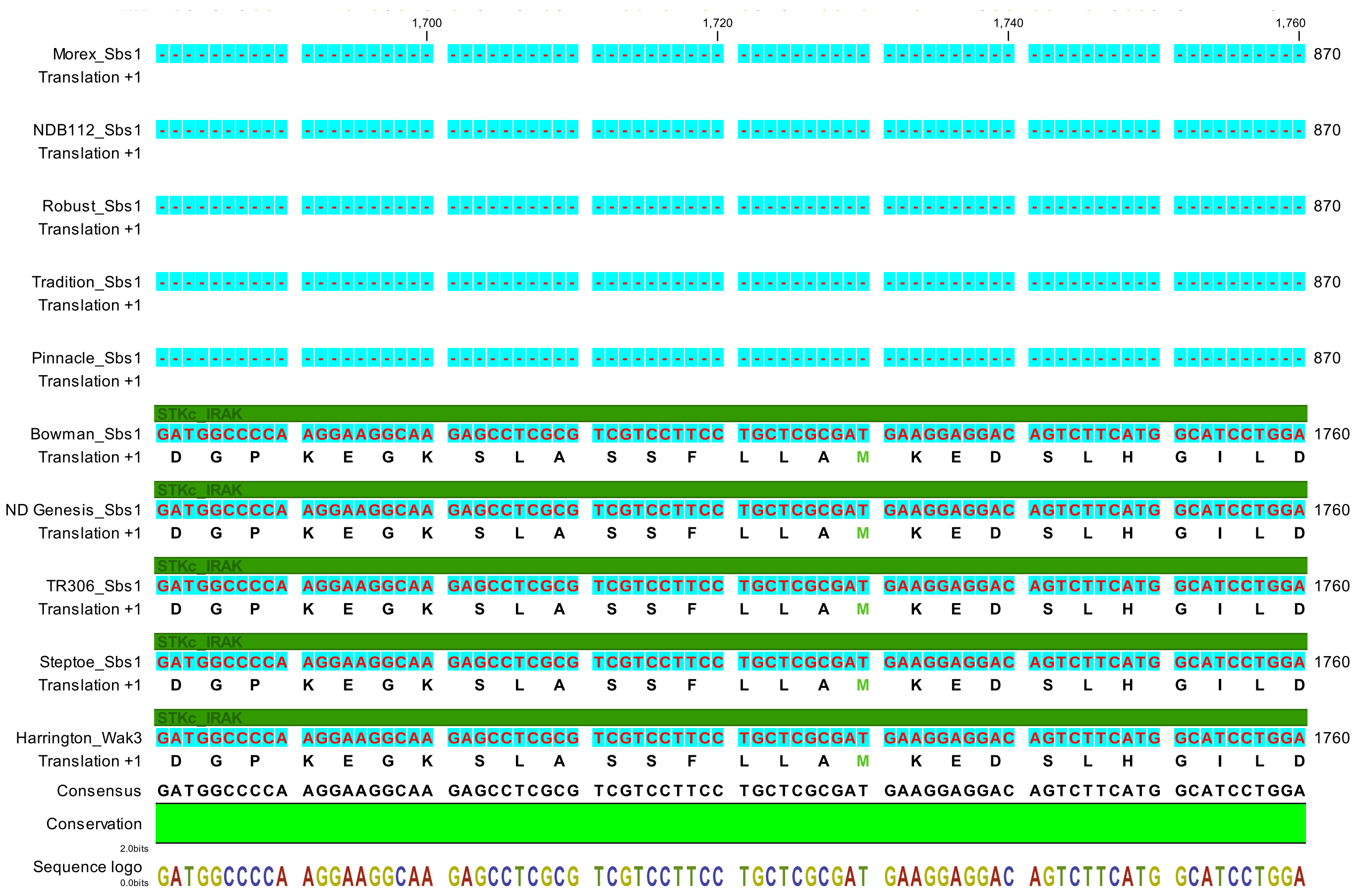

Supplementary Figure 5B

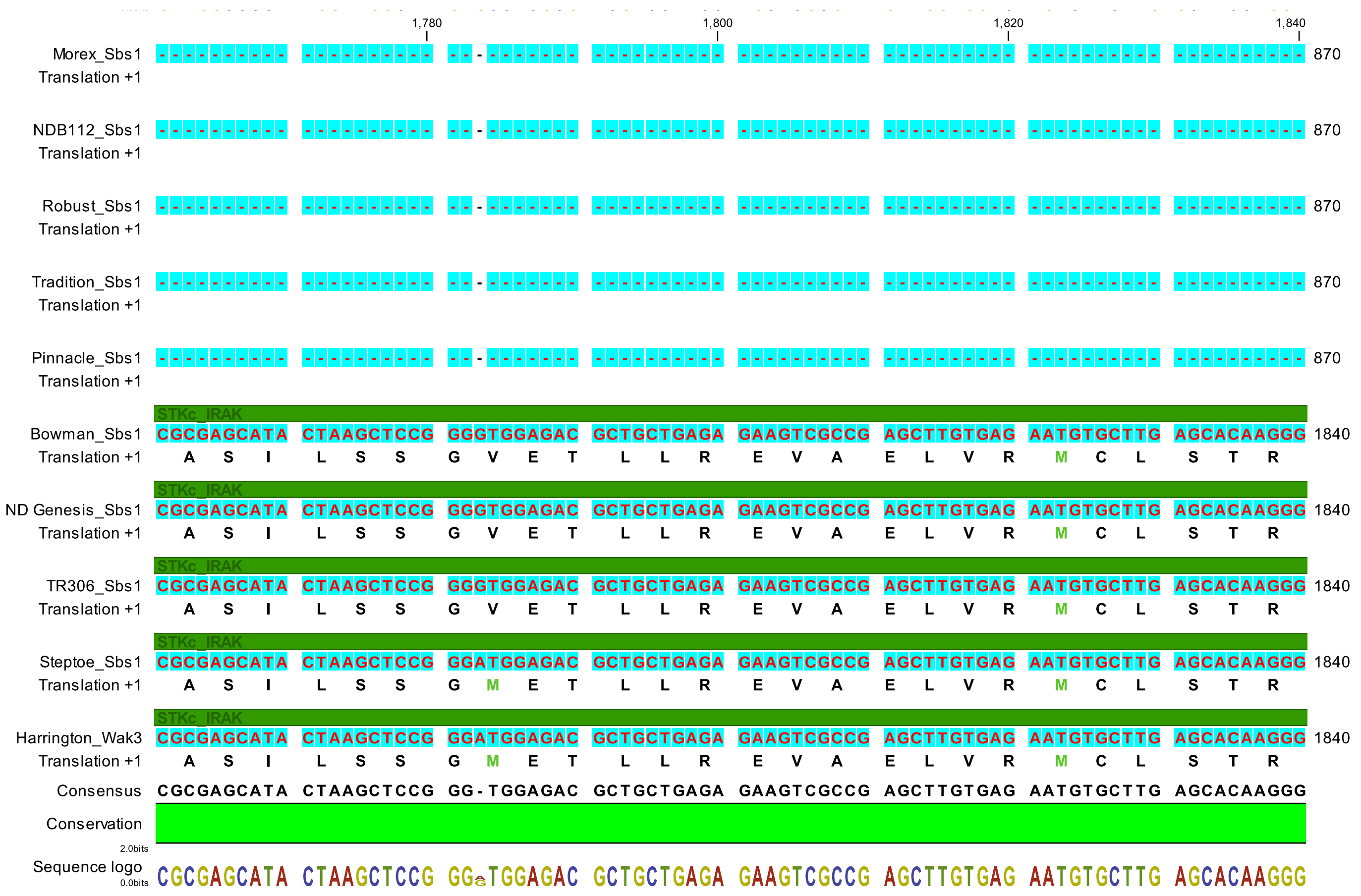

### Supplementary Figure 5B

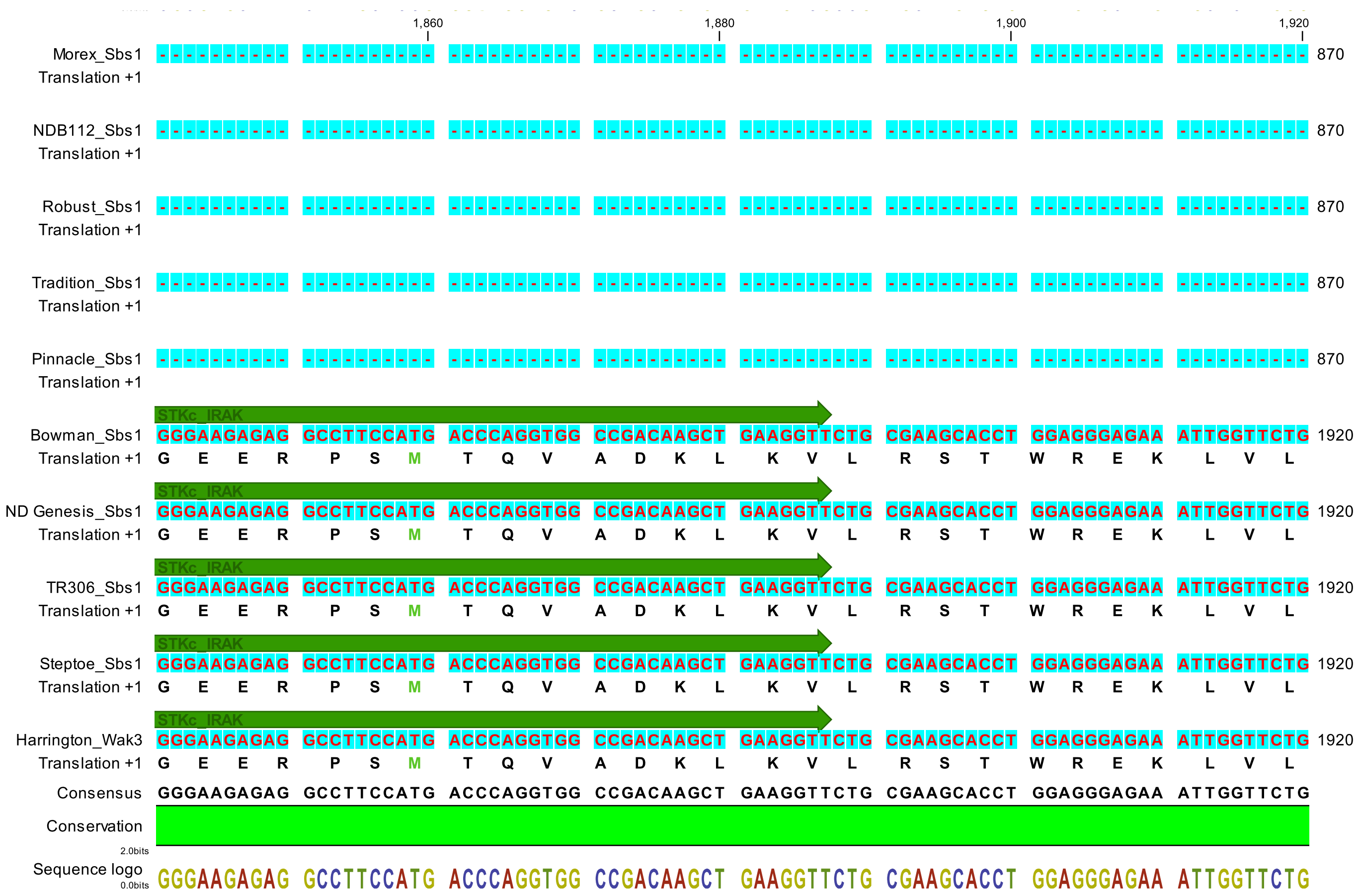

### Supplementary Figure 5B

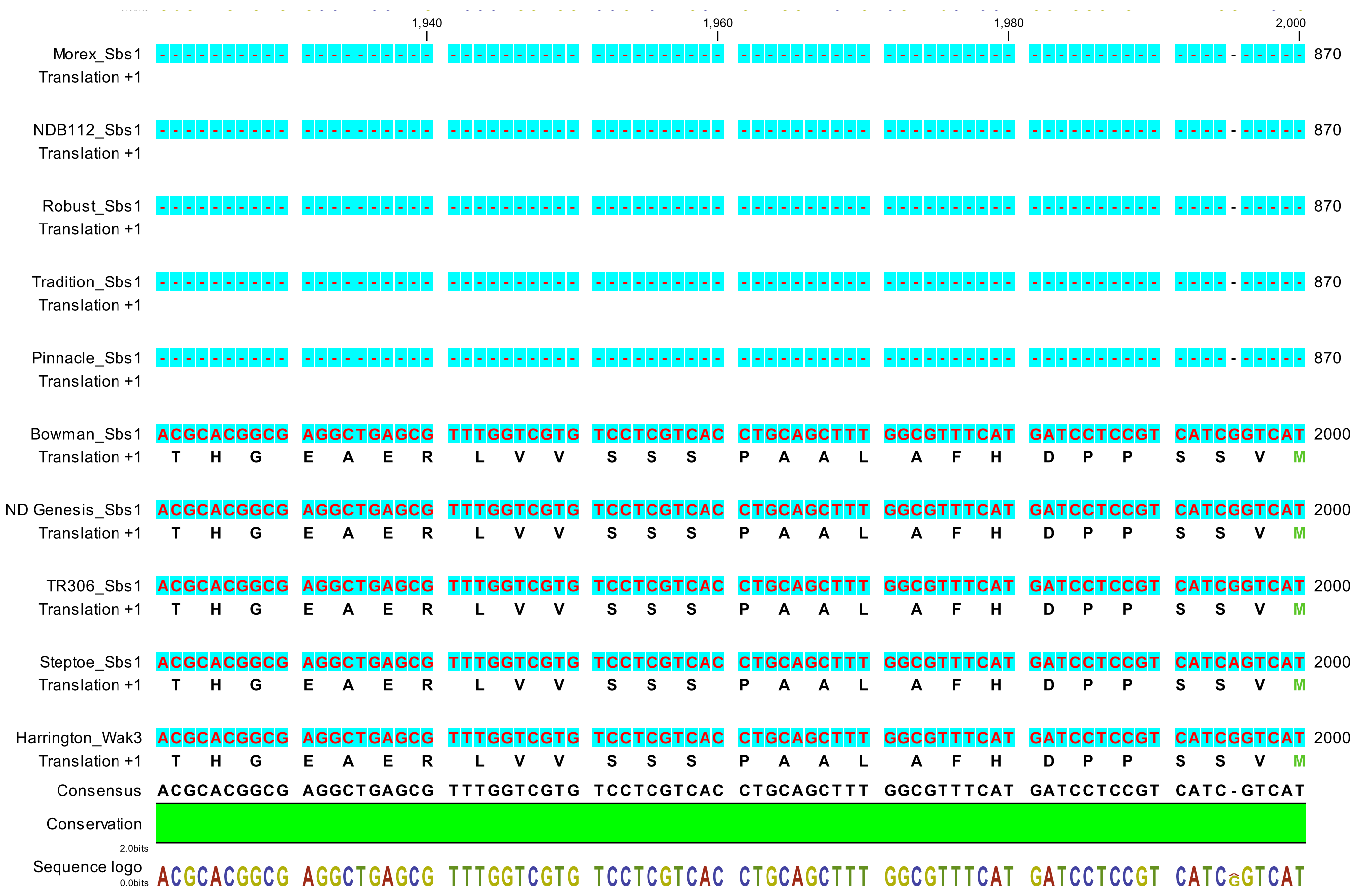

### Supplementary Figure 5B

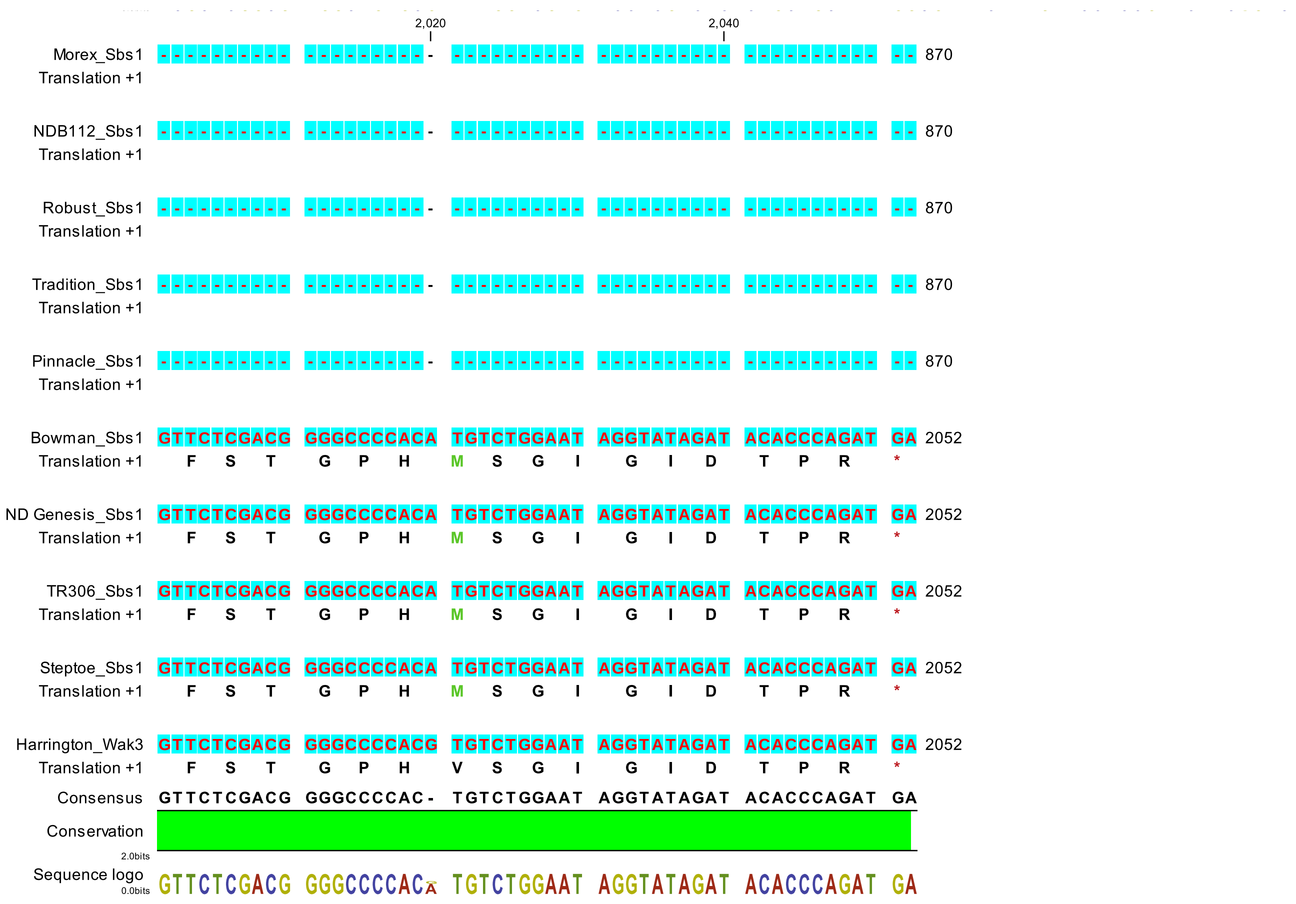

### Supplementary Figure 5C

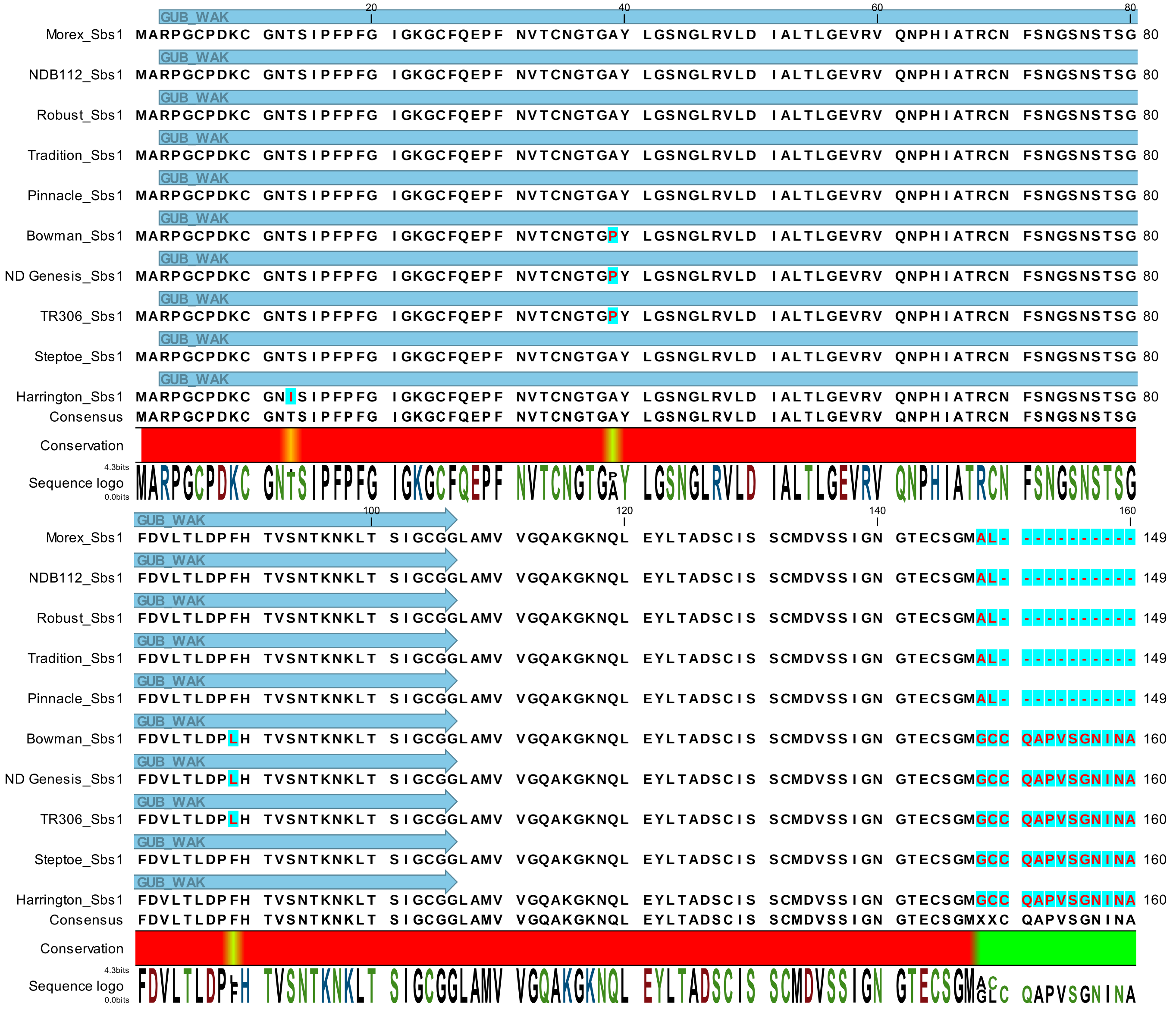

Supplementary Figure 5C

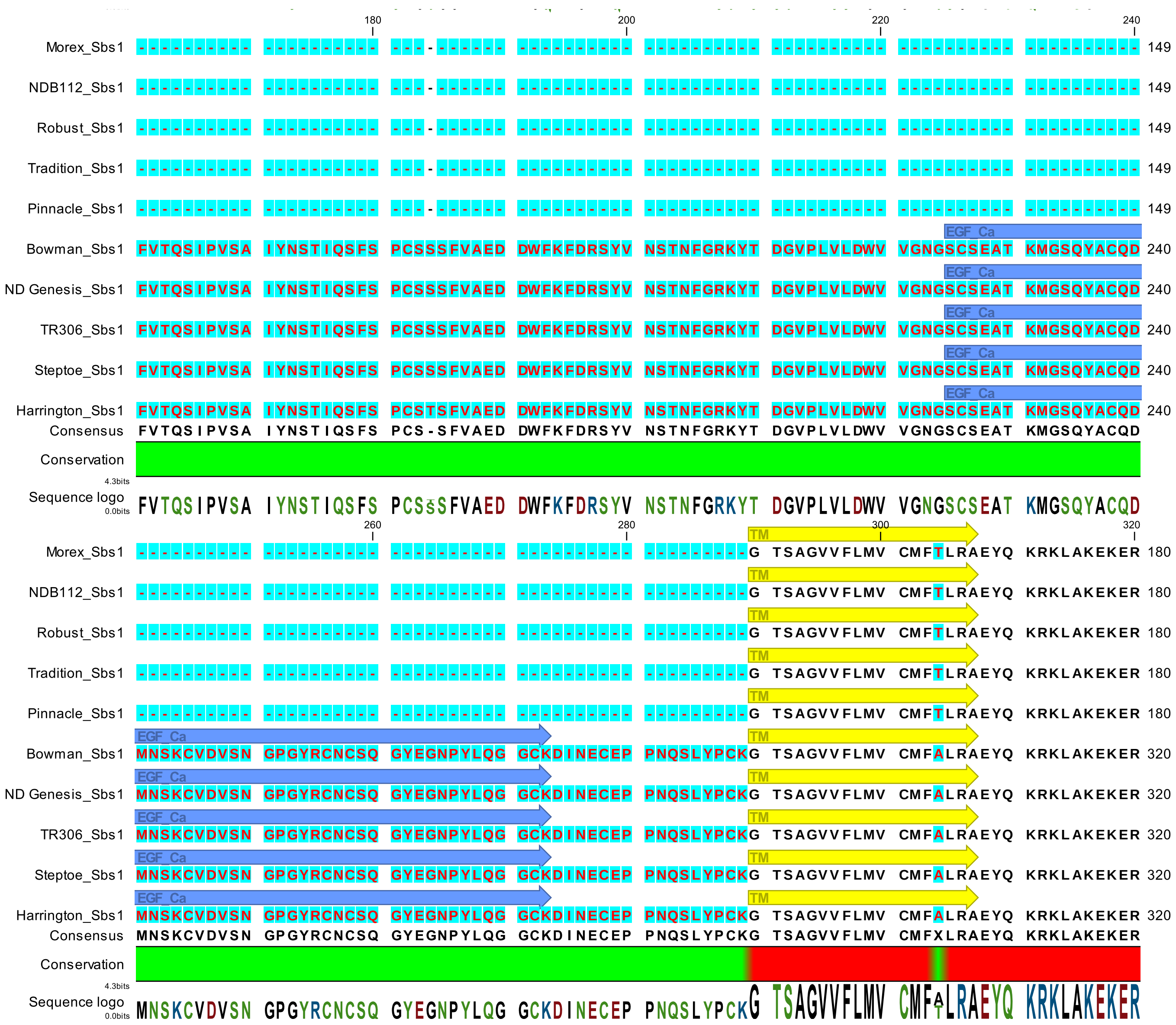

Supplementary Figure 5C

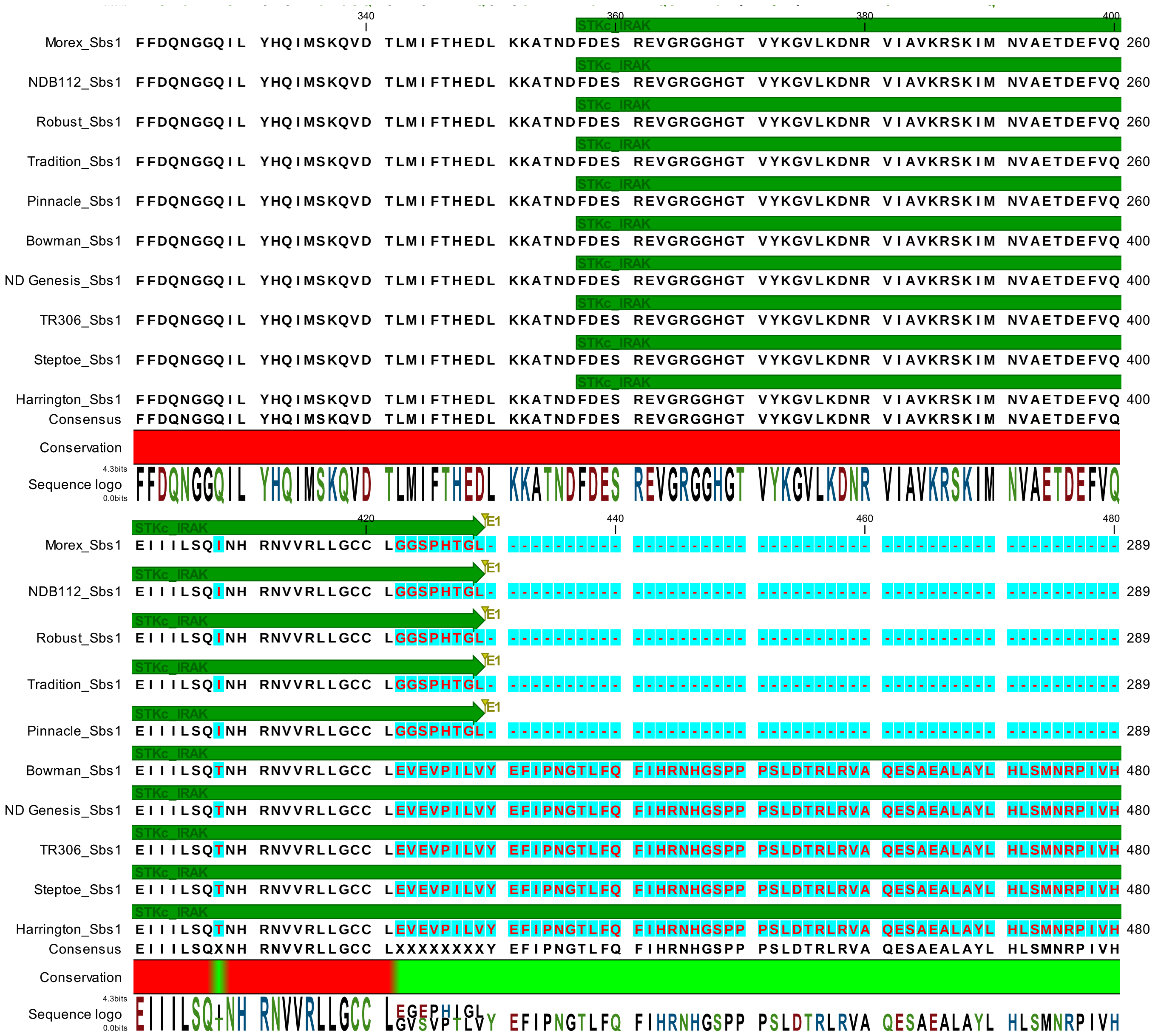

### Supplementary Figure 5C

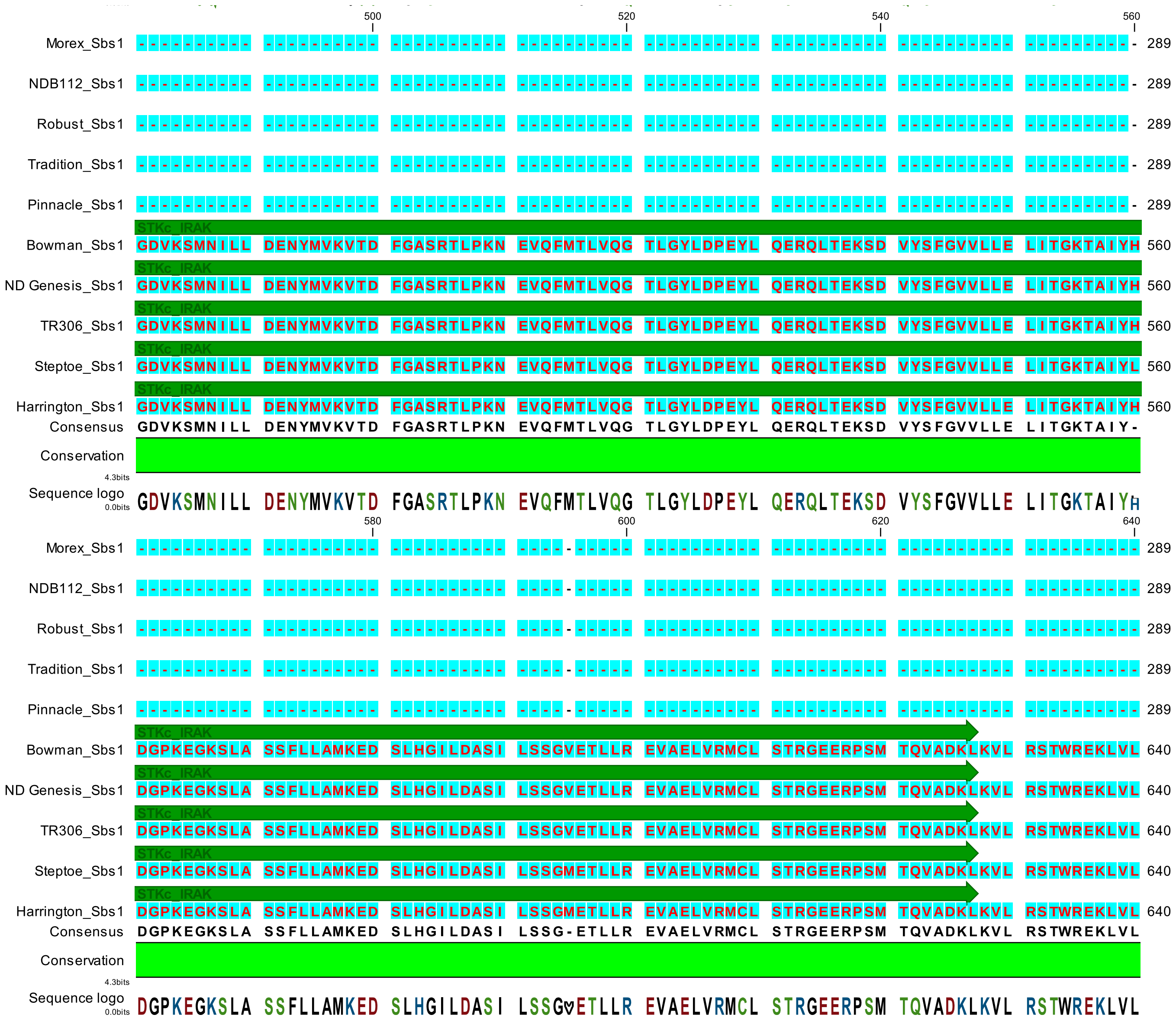

Supplementary Figure 5C

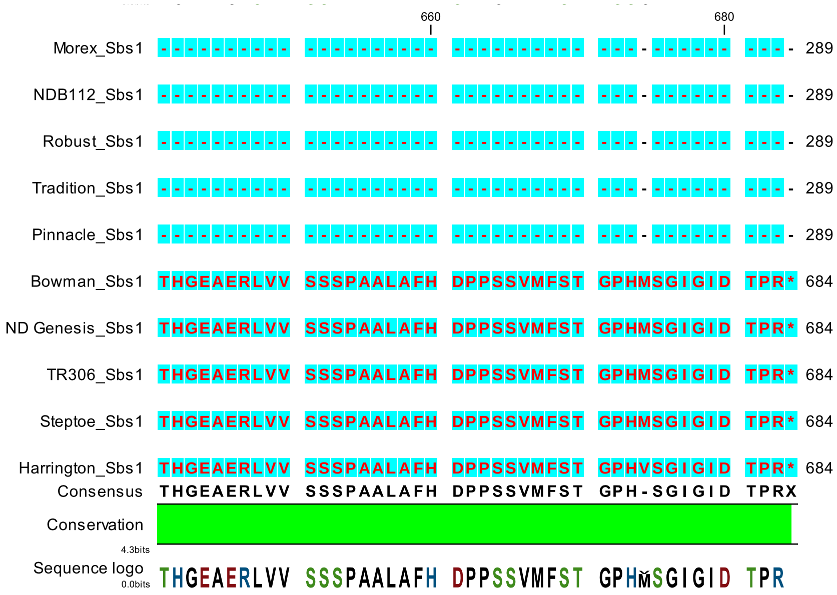

### Supplementary Figure 5D.

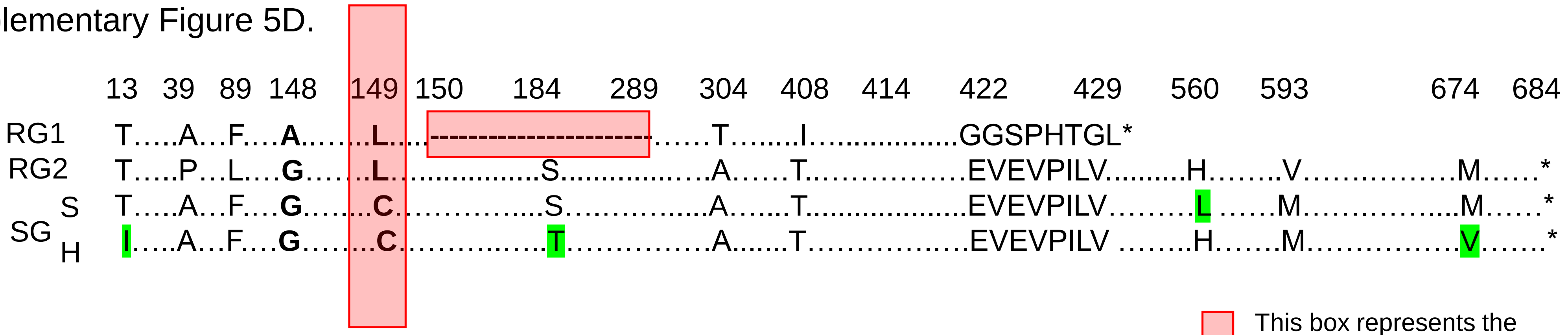


This box represents the deleterious changes in the protein (PROVEAN)

Gaps

| Group | Line | | 1 | 2 | 3 | 4 | 5 | 6 | 7 | 8 | 9 | 10 |
|---|---|---|---|---|---|---|---|---|---|---|---|---|
| RG1 | Morex_Sbs1 | 1 | | 0 | 0 | 0 | 0 | 633 | 633 | 633 | 633 | 633 |
| RG1 | NDB112_Sbs1 | 2 | 0 | | 0 | 0 | 0 | 633 | 633 | 633 | 633 | 633 |
| RG1 | Robust_Sbs1 | 3 | 0 | 0 | | 0 | 0 | 633 | 633 | 633 | 633 | 633 |
| RG1 | Tradition_Sbs1 | 4 | 0 | 0 | 0 | | 0 | 633 | 633 | 633 | 633 | 633 |
| RG1 | Pinnacle_Sbs1 | 5 | 0 | 0 | 0 | 0 | | 633 | 633 | 633 | 633 | 633 |
| RG2 | Bowman_Sbs1 | 6 | 649 | 649 | 649 | 649 | 649 | | 0 | 0 | 0 | 0 |
| RG2 | ND Genesis_Sbs1 | 7 | 649 | 649 | 649 | 649 | 649 | 0 | | 0 | 0 | 0 |
| RG2 | TR306_Sbs1 | 8 | 649 | 649 | 649 | 649 | 649 | 0 | 0 | | 0 | 0 |
| SG | Steptoe_Sbs1 | 9 | 647 | 647 | 647 | 647 | 647 | 9 | 9 | 9 | | 0 |
| SG | Harrington_Wak3 | 10 | 646 | 646 | 646 | 646 | 646 | 10 | 10 | 10 | 9 | |

Differences

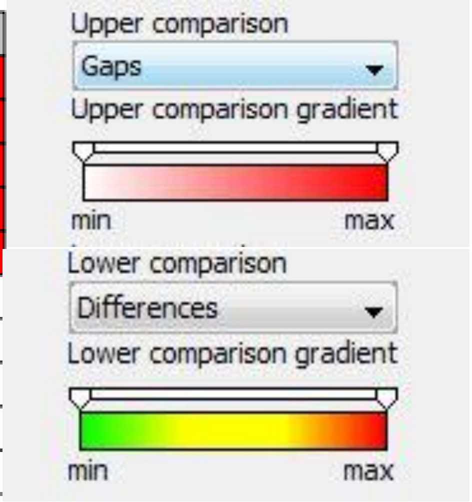

### Supplementary Figure 6A

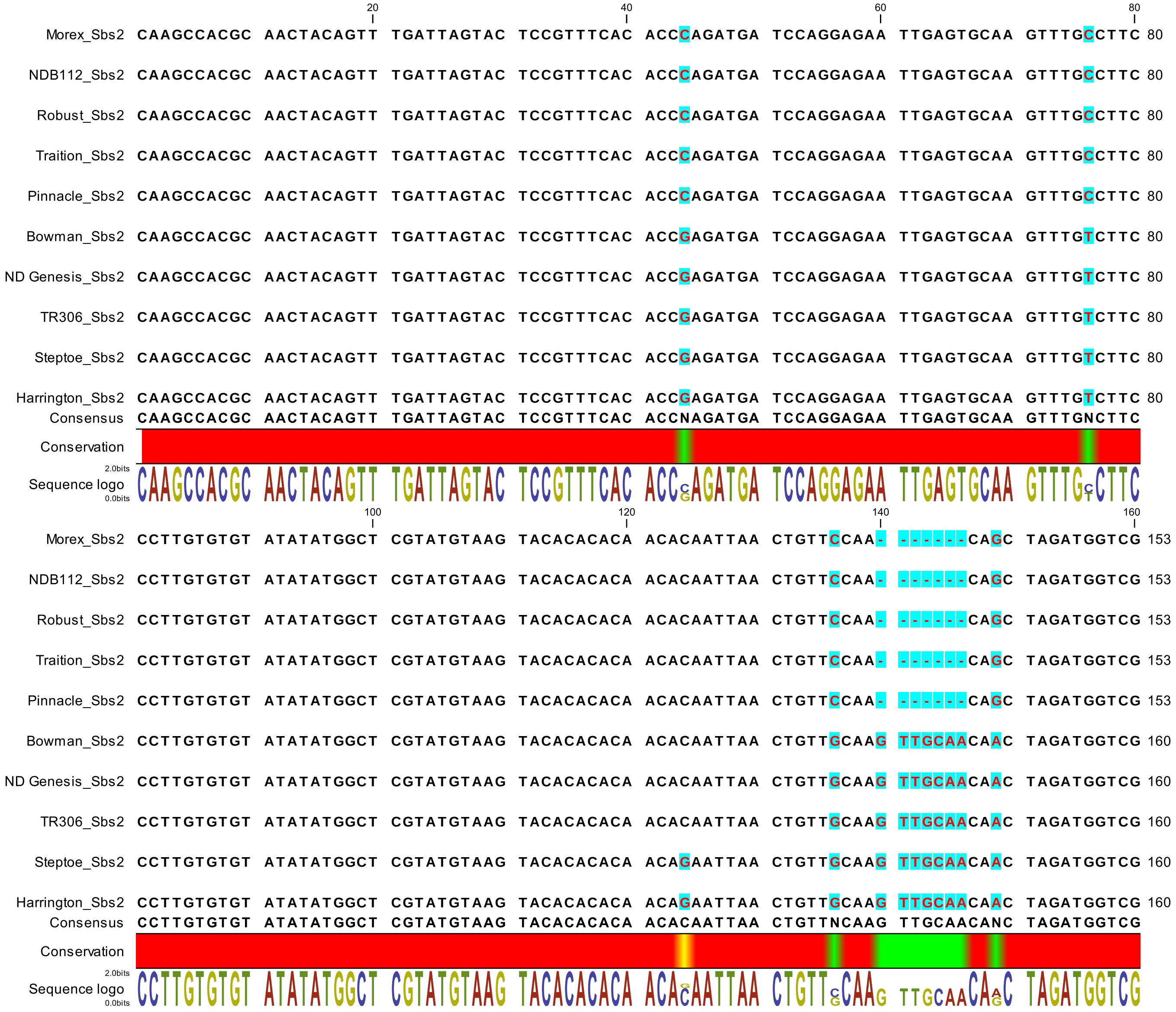

Supplementary Figure 6A

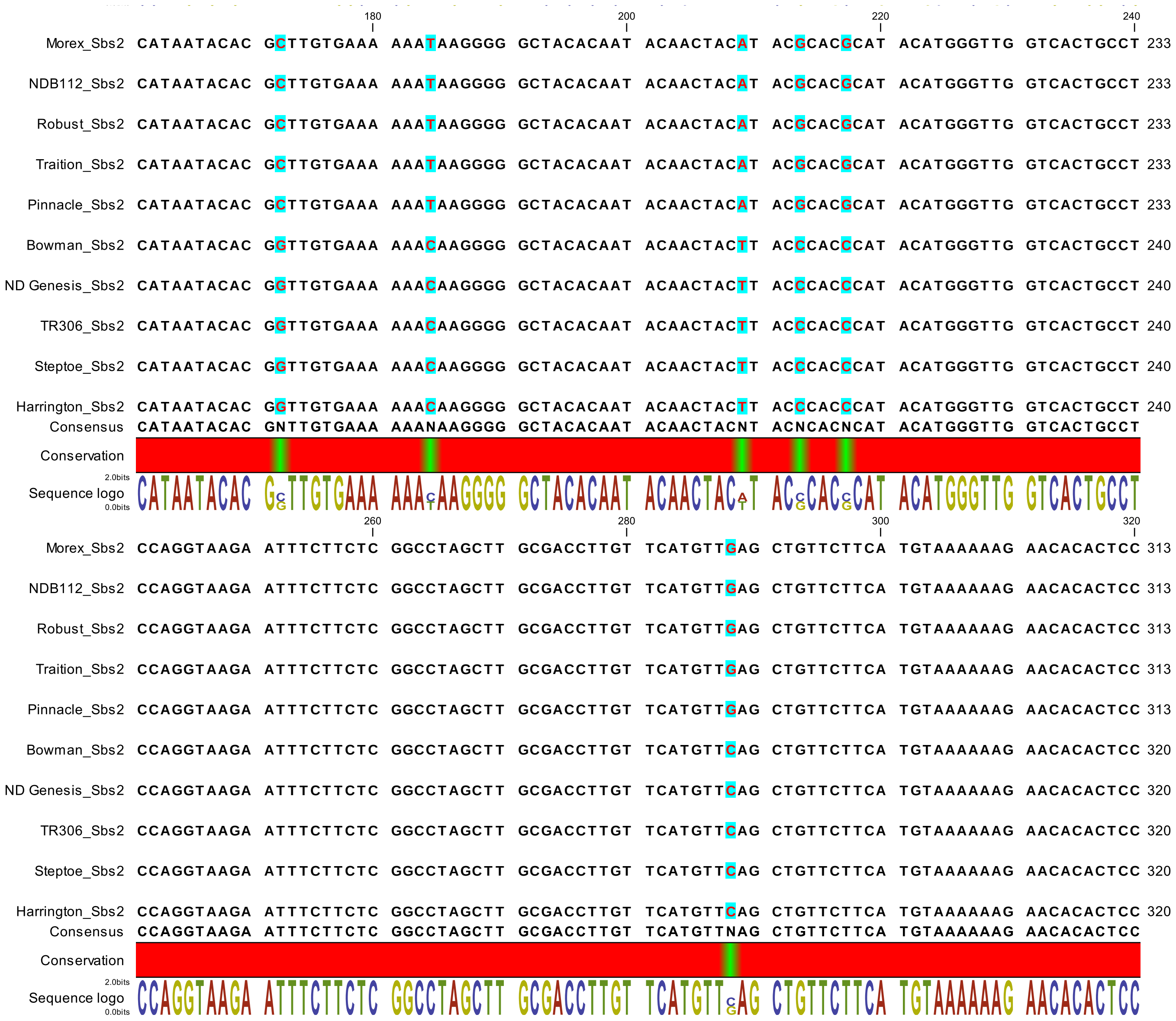

### Supplementary Figure 6A

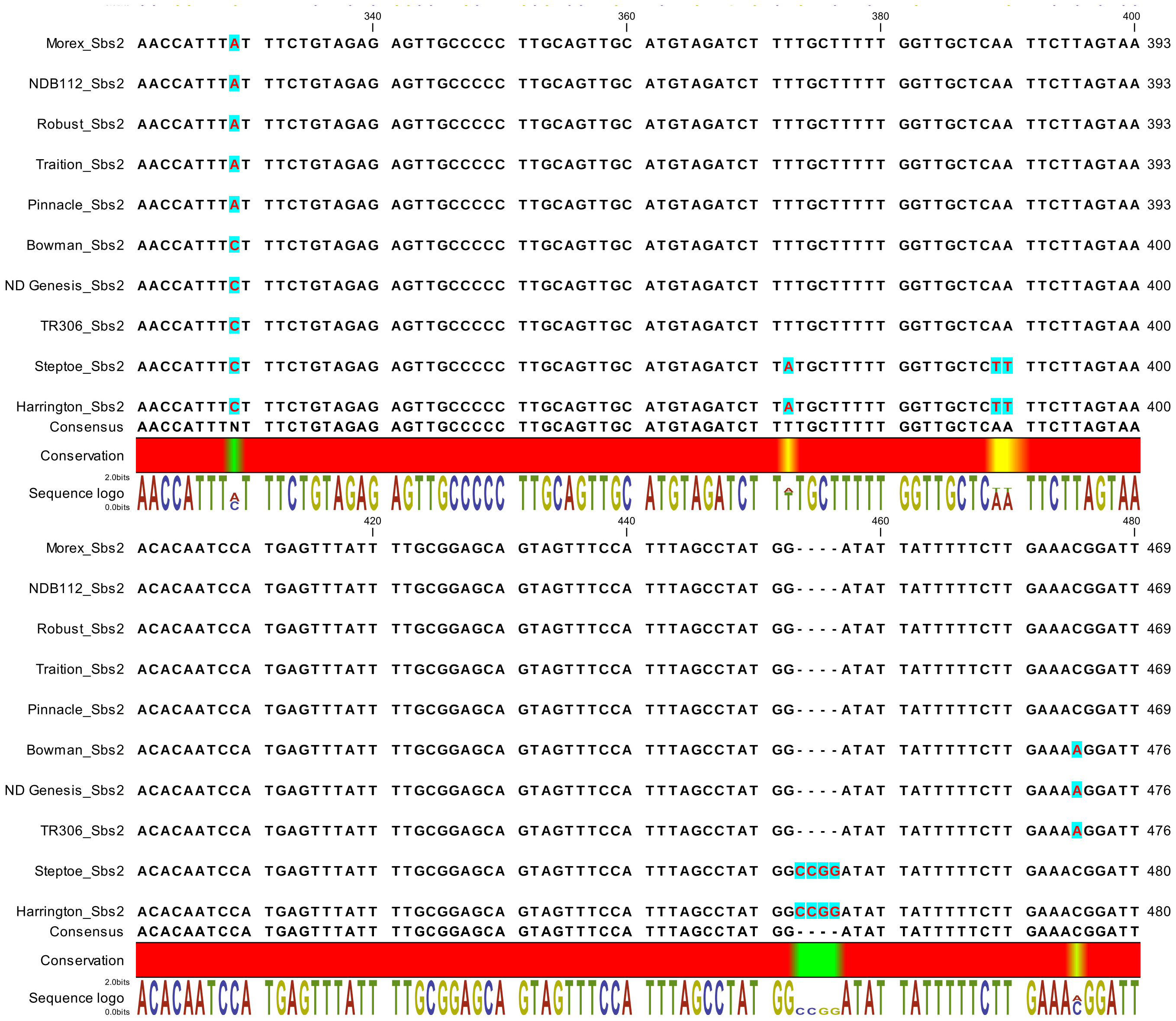

### Supplementary Figure 6A

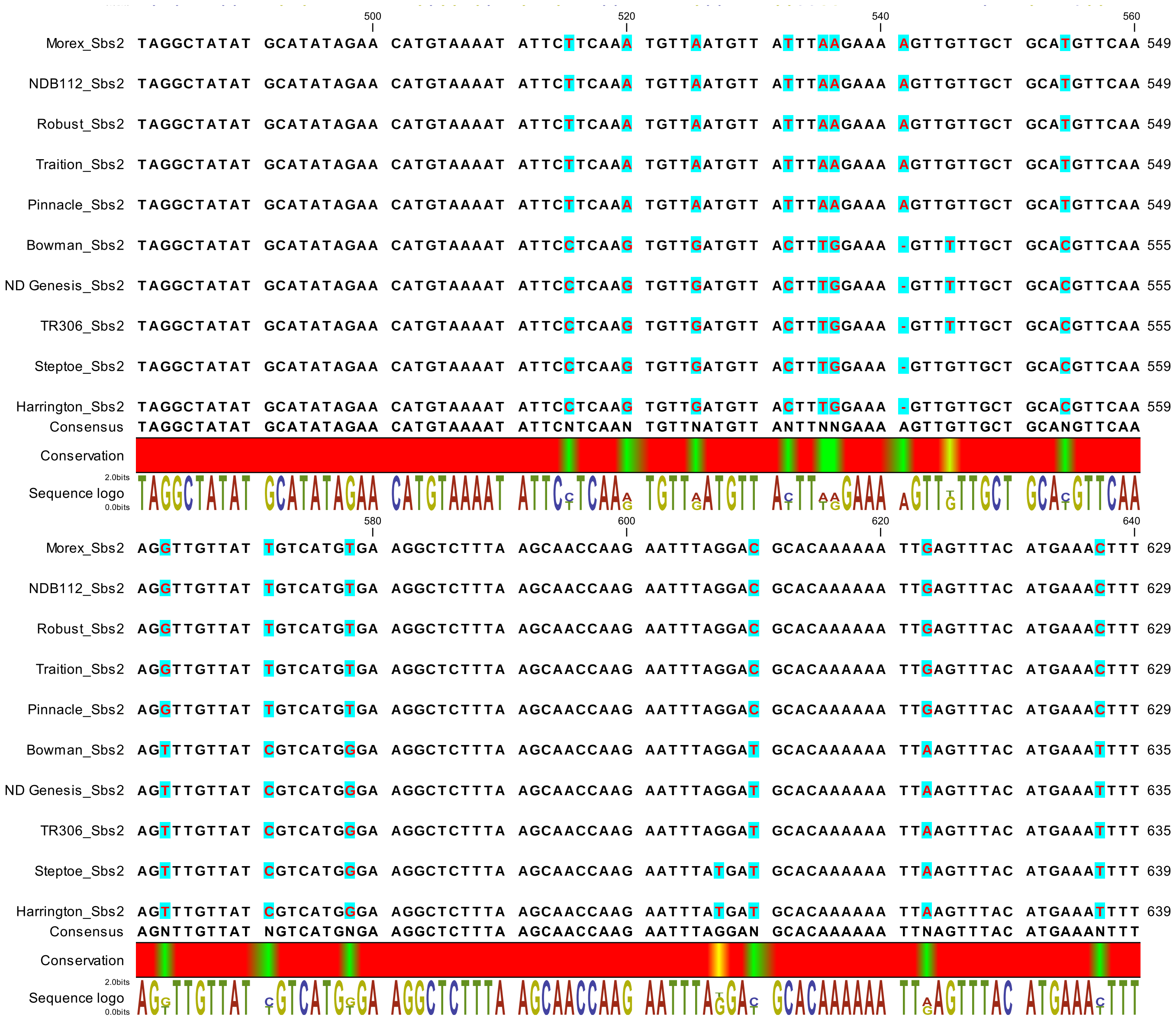

### Supplementary Figure 6A

```
                                       660                 680                 700                 720
                                        |                   |                   |                   |
Morex_Sbs2        CACCTATTTC ATCGCACCAT CATACTATTA AGCTTGATGA TAGCTTCTCA GAACCCCCCC CCCCCC--TC ATGCGAAATA 707
NDB112_Sbs2       CACCTATTTC ATCGCACCAT CATACTATTA AGCTTGATGA TAGCTTCTCA GAACCCCCCC CCCCCCCCTC ATGCGAAATA 709
Robust_Sbs2       CACCTATTTC ATCGCACCAT CATACTATTA AGCTTGATGA TAGCTTCTCA GAACCCCCCC CCCC----TC ATGCGAAATA 705
Traition_Sbs2     CACCTATTTC ATCGCACCAT CATACTATTA AGCTTGATGA TAGCTTCTCA GAACCCCCCC CCCCCC--TC ATGCGAAATA 707
Pinnacle_Sbs2     CACCTATTTC ATCGCACCAT CATACTATTA AGCTTGATGA TAGCTTCTCA GAACCCCCCC CCCCCC--TC ATGCGAAATA 707
Bowman_Sbs2       CACCTAATTT GTCTCACCAT CATACTATTA ACCTTGATGA TAGCTTGTCA GAATCCCCCC CCCCCTCCTC ATGCGAAGTA 715
ND Genesis_Sbs2   CACCTAATTT GTCTCACCAT CATACTATTA ACCTTGATGA TAGCTTGTCA GAATCCCCCC CCCCCTCCTC ATGCGAAGTA 715
TR306_Sbs2        CACCTAATTT GTCTCACCAT CATACTATTA ACCTTGATGA TAGCTTGTCA GAATCCCCCC CCCCCTCCTC ATGCGAAGTA 715
Steptoe_Sbs2      CACCTATTTT GTCTCACCAT CATACTATTA ACCTTGATGA TAGCTTGTCA GAATCCCCCC CCCCTCCCTC ATGCGAAGTA 719
Harrington_Sbs2   CACCTATTTT GTCTCACCAT CATACTATTA ACCTTGATGA TAGCTTGTCA GAATCCCCCC CCCCTCCCTC ATGCGAAGTA 719
Consensus         CACCTATTTN NTCNCACCAT CATACTATTA ANCTTGATGA TAGCTTNTCA GAANCCCCCC CCCCCCCCTC ATGCGAANTA
Conservation
Sequence logo

                                       740                 760                 780                 800
                                        |                   |                   |                   |
Morex_Sbs2        GCAATATTCA CATTTTATAC ACCC-GAGGG ATA--TTCGA GCATGAACTT CACCCCTTGC TCCCCGAGGT TAATGTGCTT 784
NDB112_Sbs2       GCAATATTCA CATTTTATAC ACCC-GAGGG ATA--TTCGA GCATGAACTT CACCCCTTGC TCCCCGAGGT TAATGTGCTT 786
Robust_Sbs2       GCAATATTCA CATTTTATAC ACCC-GAGGG ATA--TTCGA GCATGAACTT CACCCCTTGC TCCCCGAGGT TAATGTGCTT 782
Traition_Sbs2     GCAATATTCA CATTTTATAC ACCC-GAGGG ATA--TTCGA GCATGAACTT CACCCCTTGC TCCCCGAGGT TAATGTGCTT 784
Pinnacle_Sbs2     GCAATATTCA CATTTTATAC ACCC-GAGGG ATA--TTCGA GCATGAACTT CACCCCTTGC TCCCCGAGGT TAATGTGCTT 784
Bowman_Sbs2       GCAATATTCA CATTTTATAC GCCCCGAGGG ATAATTTCGA GCATGA-CTT CACCCCCTGC TCCCCGAGGT TAATGCATTT 794
ND Genesis_Sbs2   GCAATATTCA CATTTTATAC GCCCCGAGGG ATAATTTCGA GCATGA-CTT CACCCCCTGC TCCCCGAGGT TAATGCATTT 794
TR306_Sbs2        GCAATATTCA CATTTTATAC GCCCCGAGGG ATAATTTCGA GCATGA-CTT CACCCCCTGC TCCCCGAGGT TAATGCATTT 794
Steptoe_Sbs2      GCAATATTCA CATTTTATAC ACCCCGAGGG ATAATTTCGA GCATGA-CTT CACCCCCTGC TCCCCGAGGT TACTGCATTT 798
Harrington_Sbs2   GCAATATTCA CATTTTATAC ACCCCGAGGG ATAATTTCGA GCATGA-CTT CACCCCCTGC TCCCCGAGGT TACTGCATTT 798
Consensus         GCAATATTCA CATTTTATAC ACCCCGAGGG ATAATTTCGA GCATGAACTT CACCCCNTGC TCCCCGAGGT TAATGNNNTT
Conservation
Sequence logo
```

### Supplementary Figure 6A

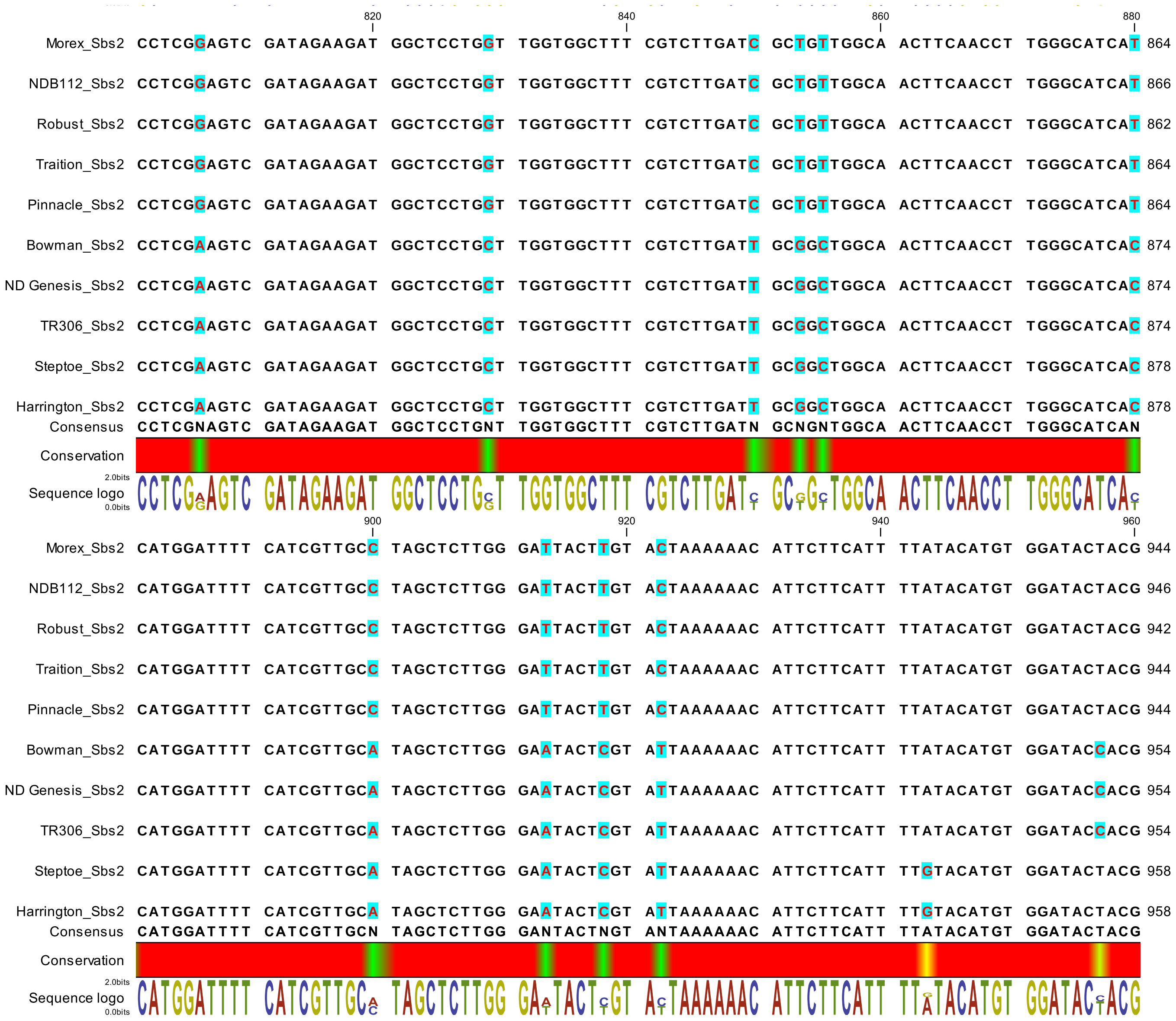

### Supplementary Figure 6A

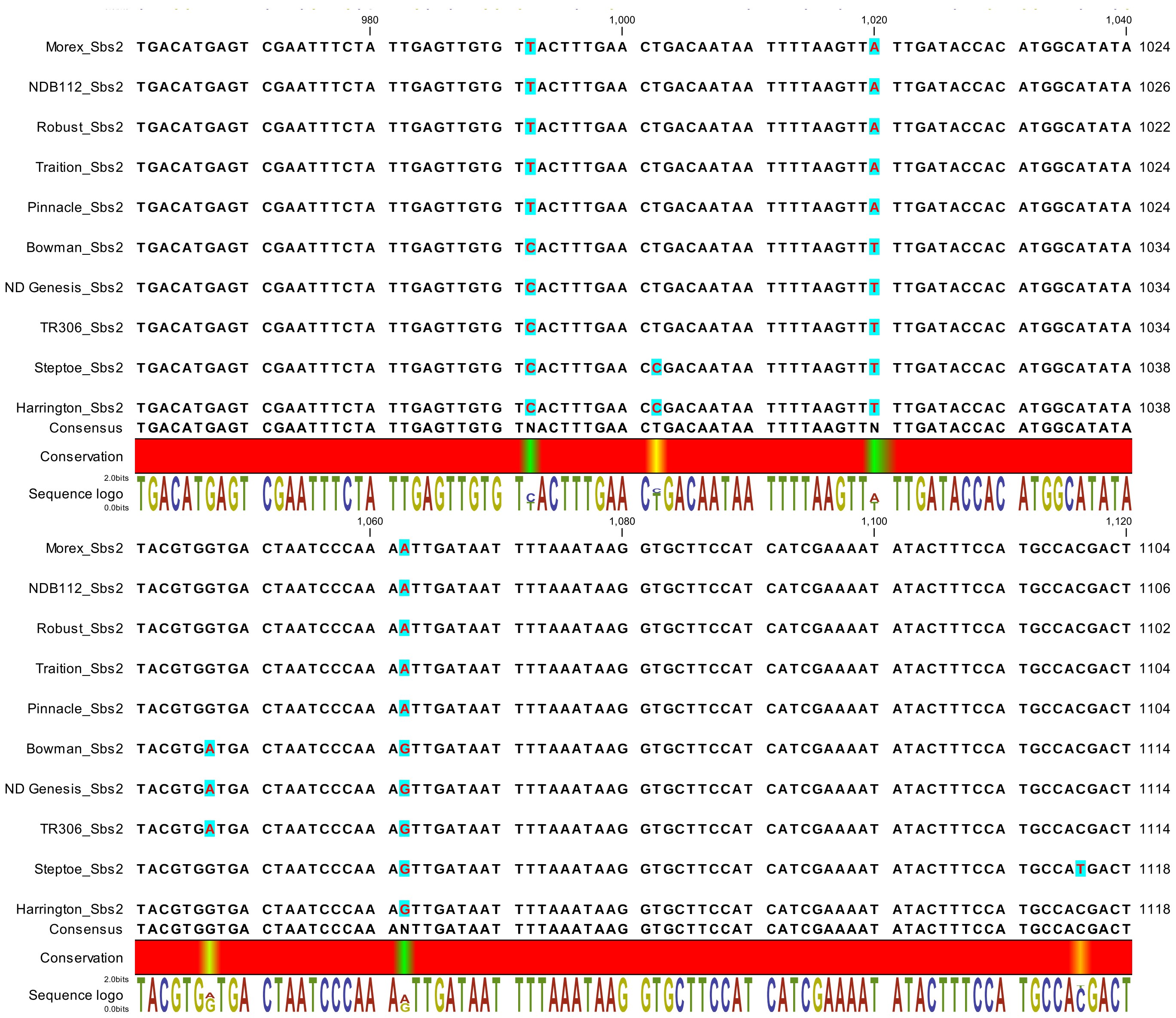

### Supplementary Figure 6A

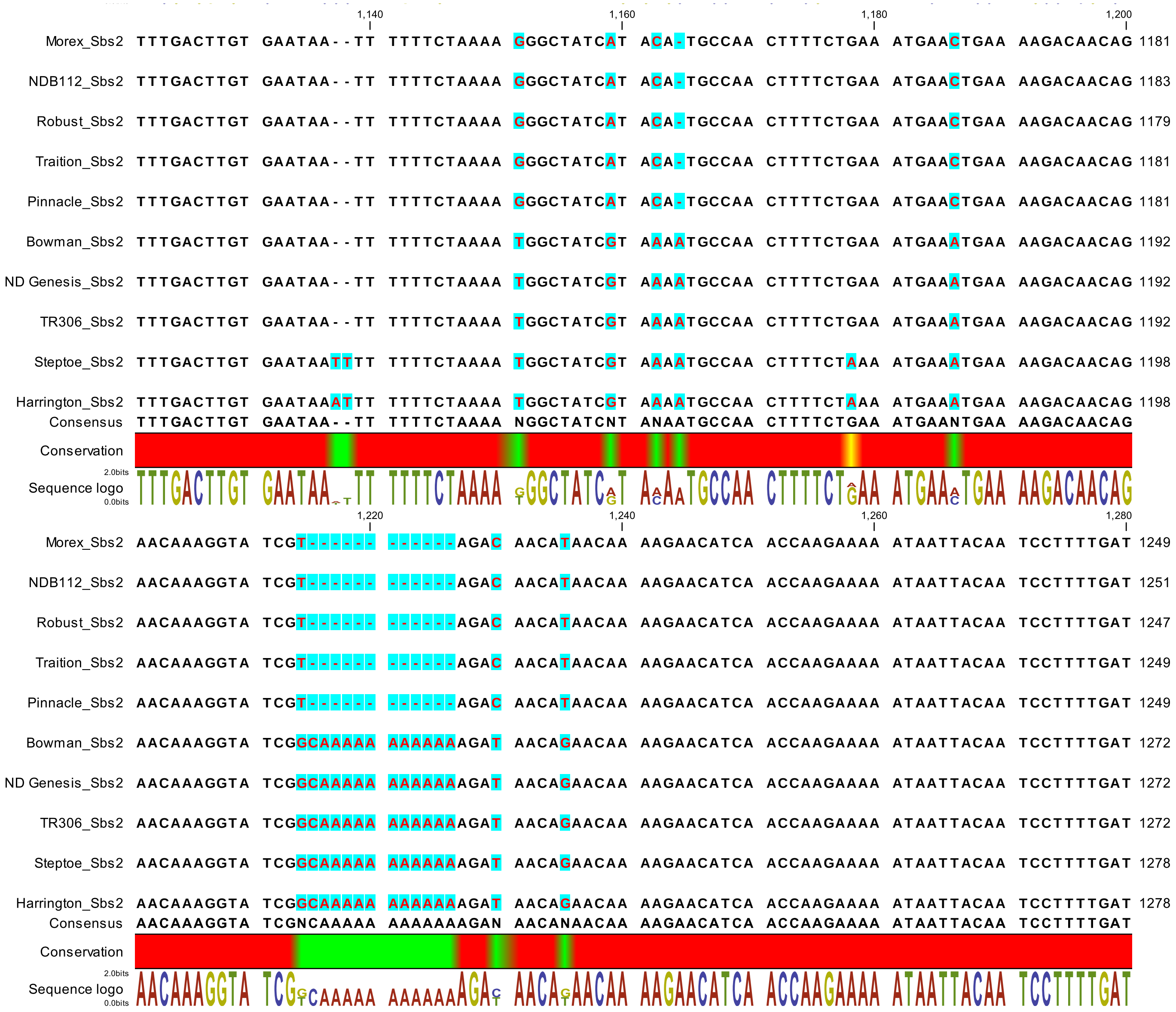

### Supplementary Figure 6A

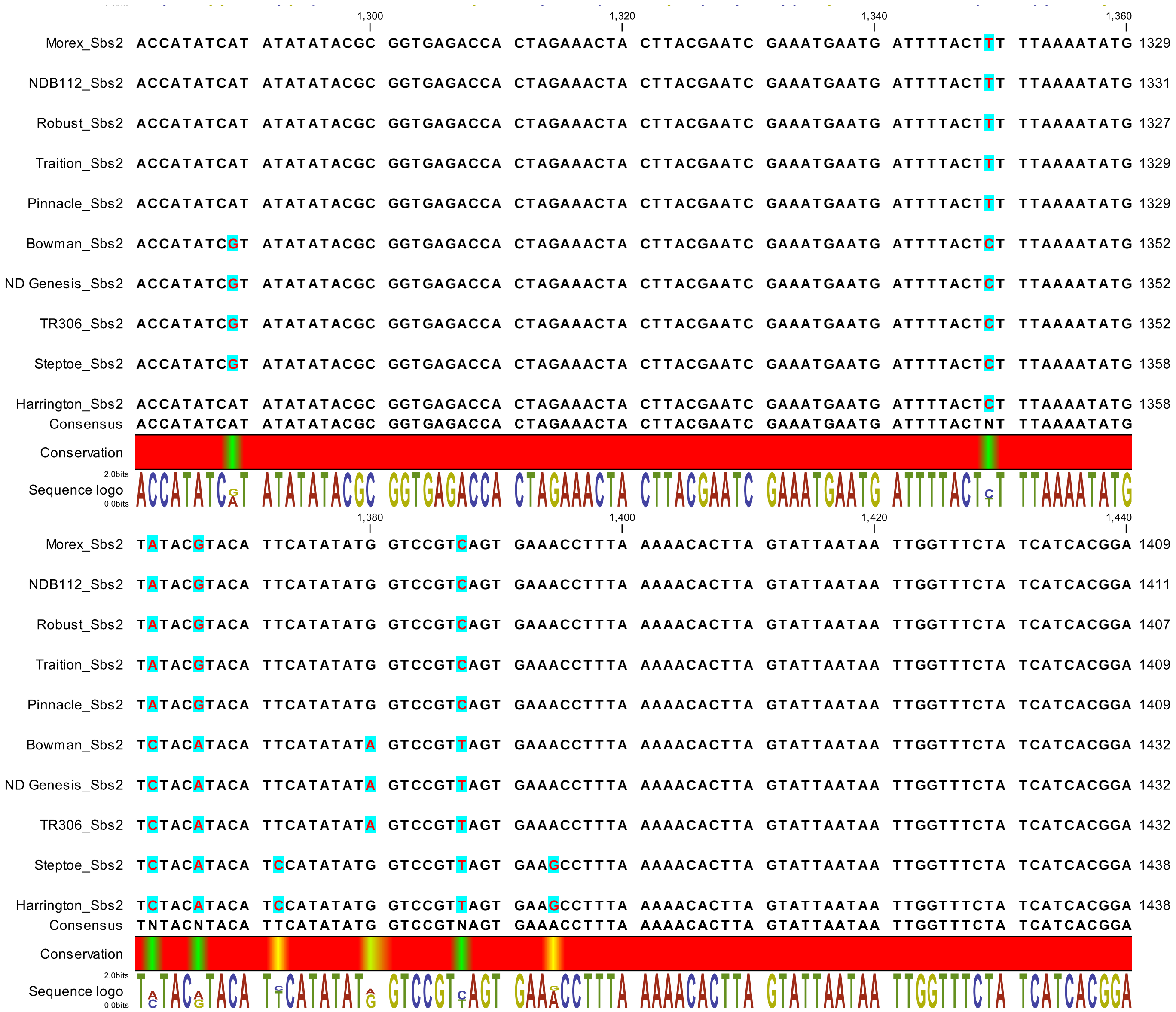

### Supplementary Figure 6A

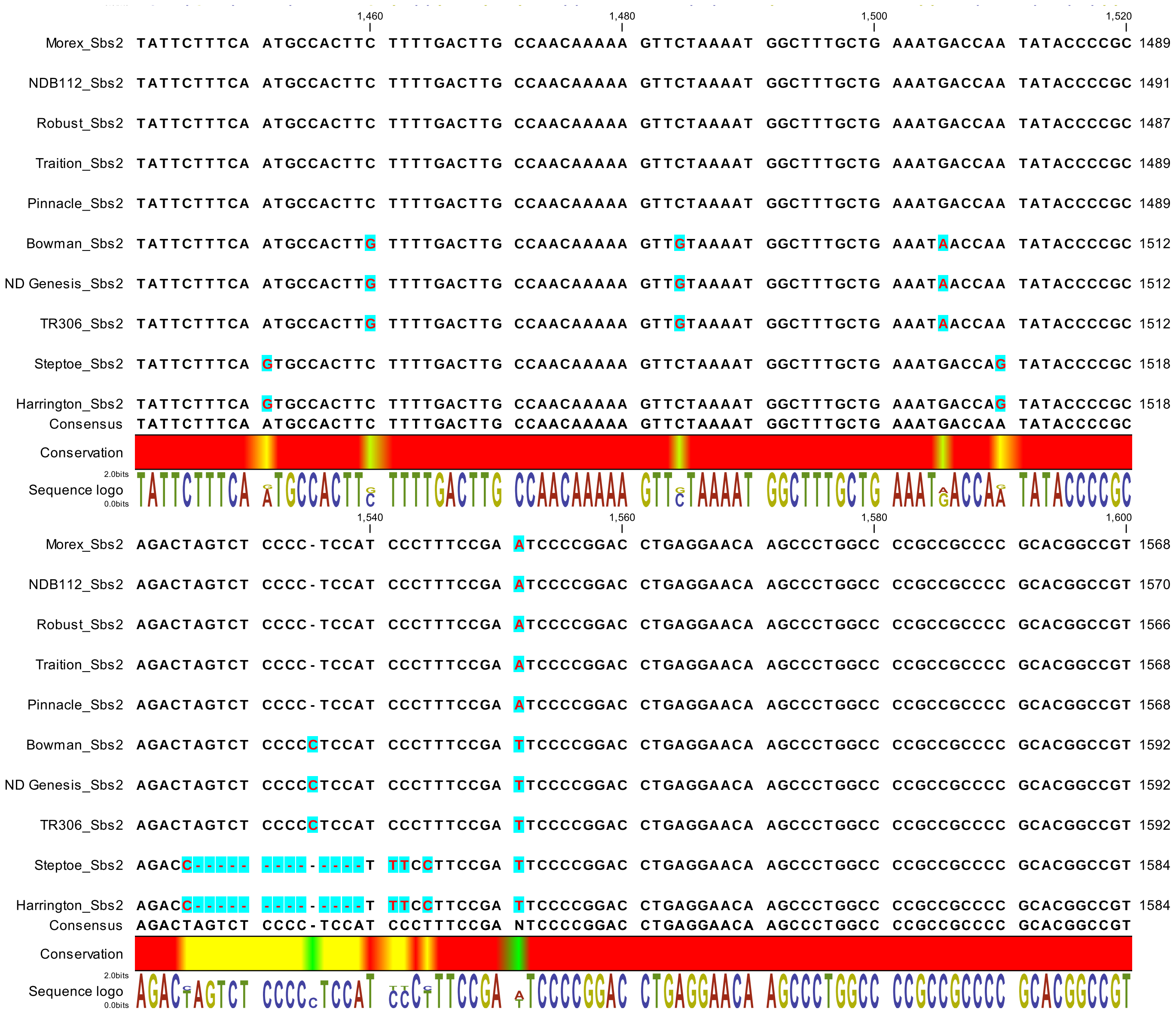

### Supplementary Figure 6A

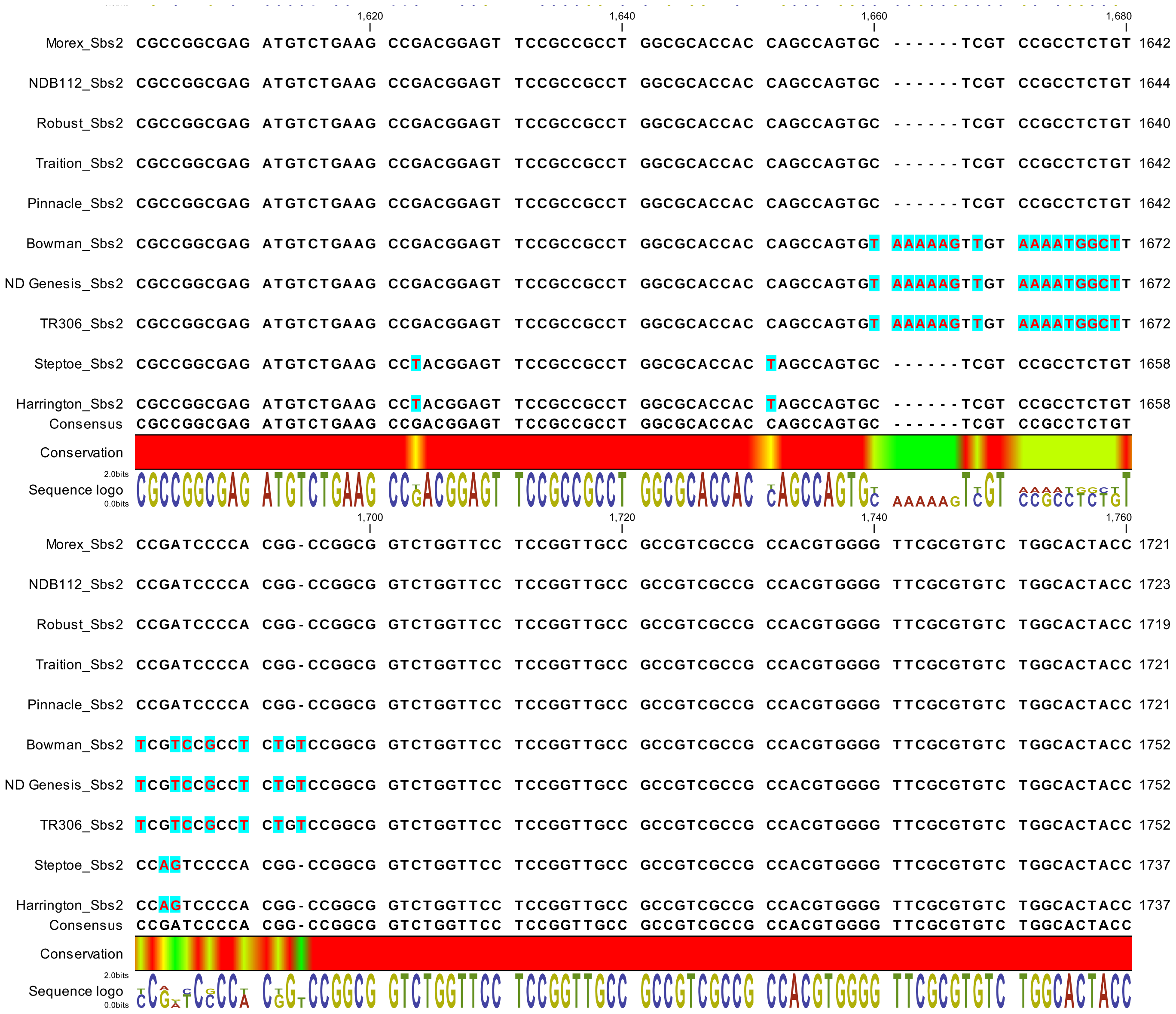

### Supplementary Figure 6A

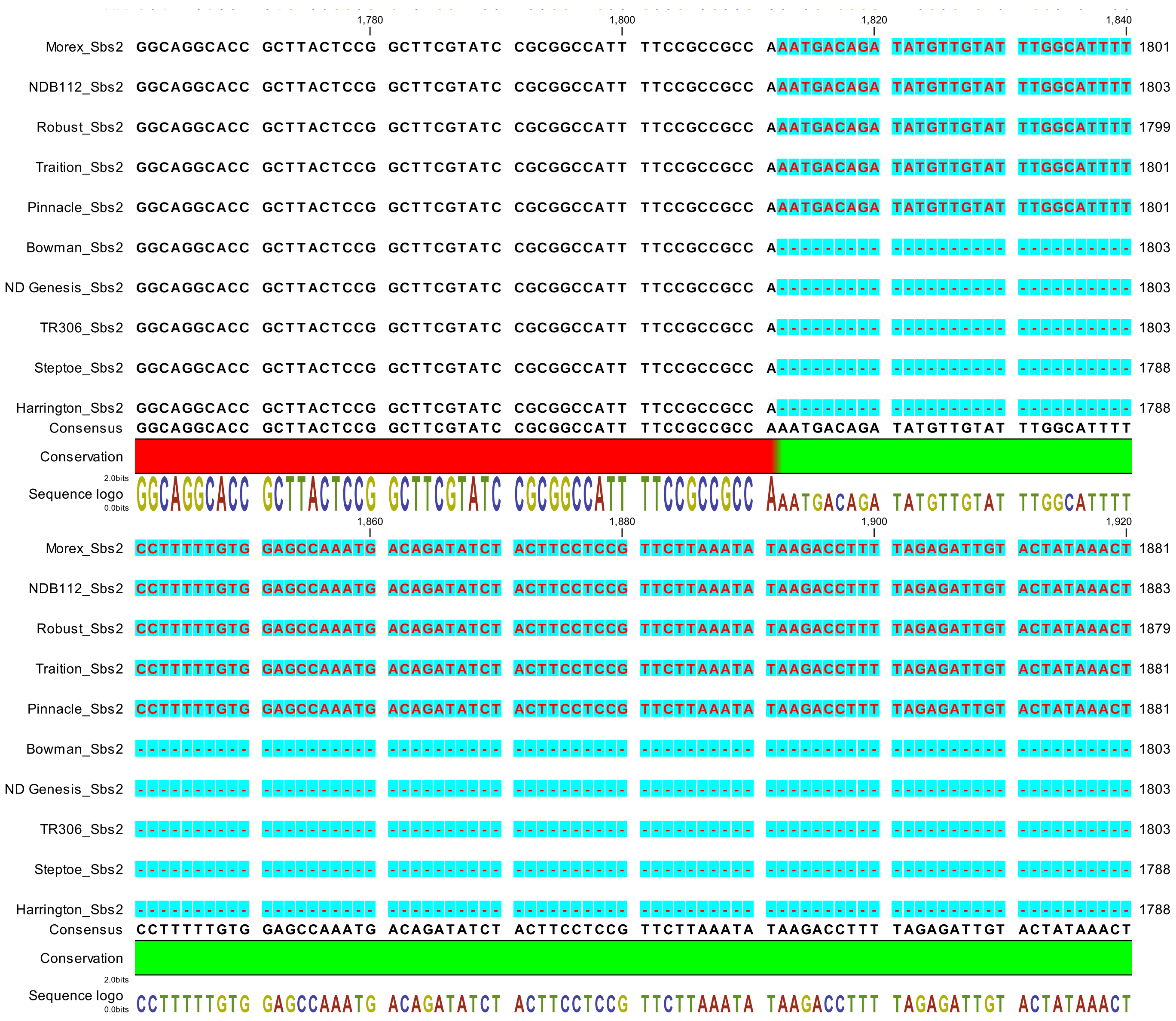

### Supplementary Figure 6A

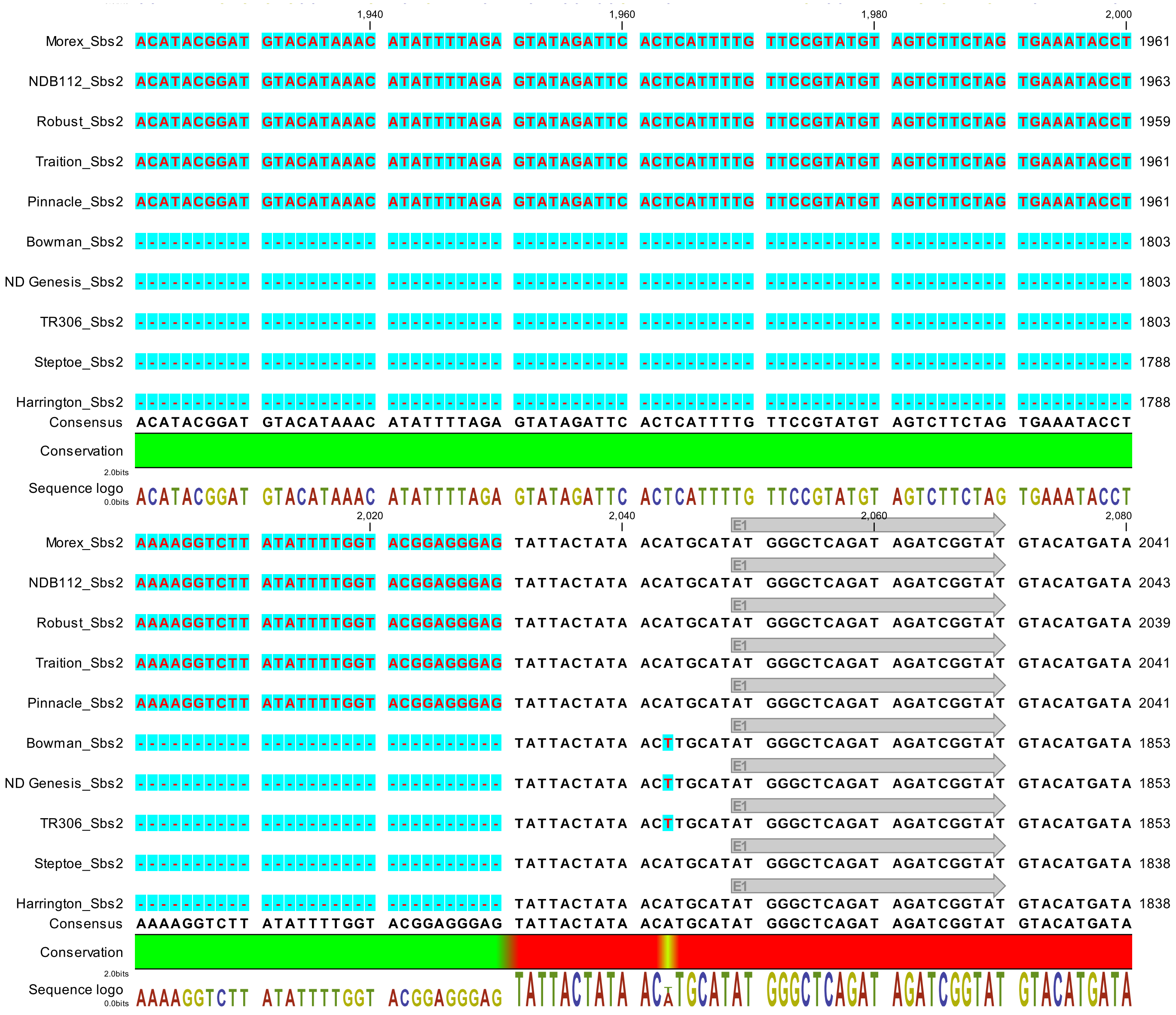

Supplementary Figure 6A

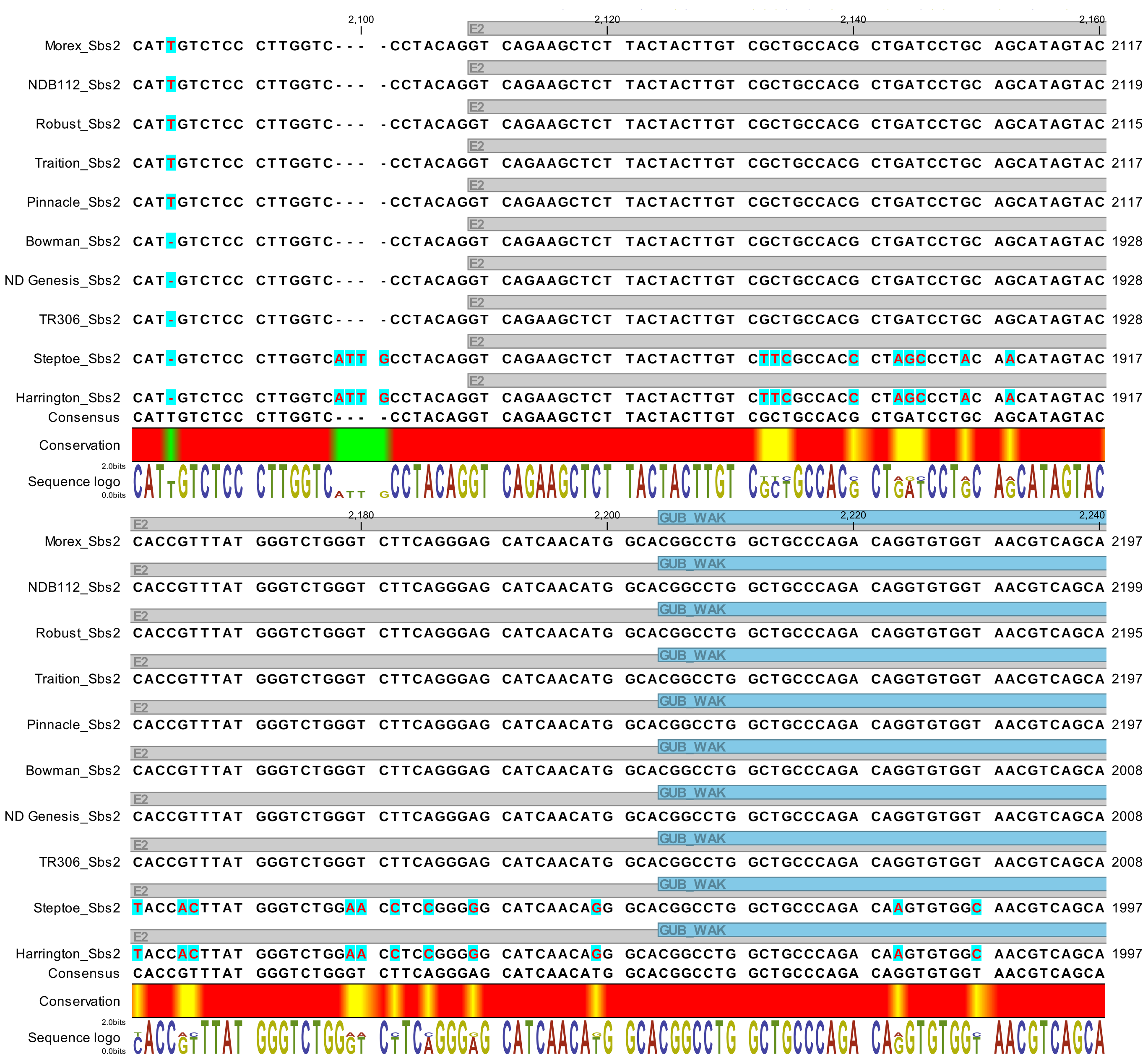

### Supplementary Figure 6A

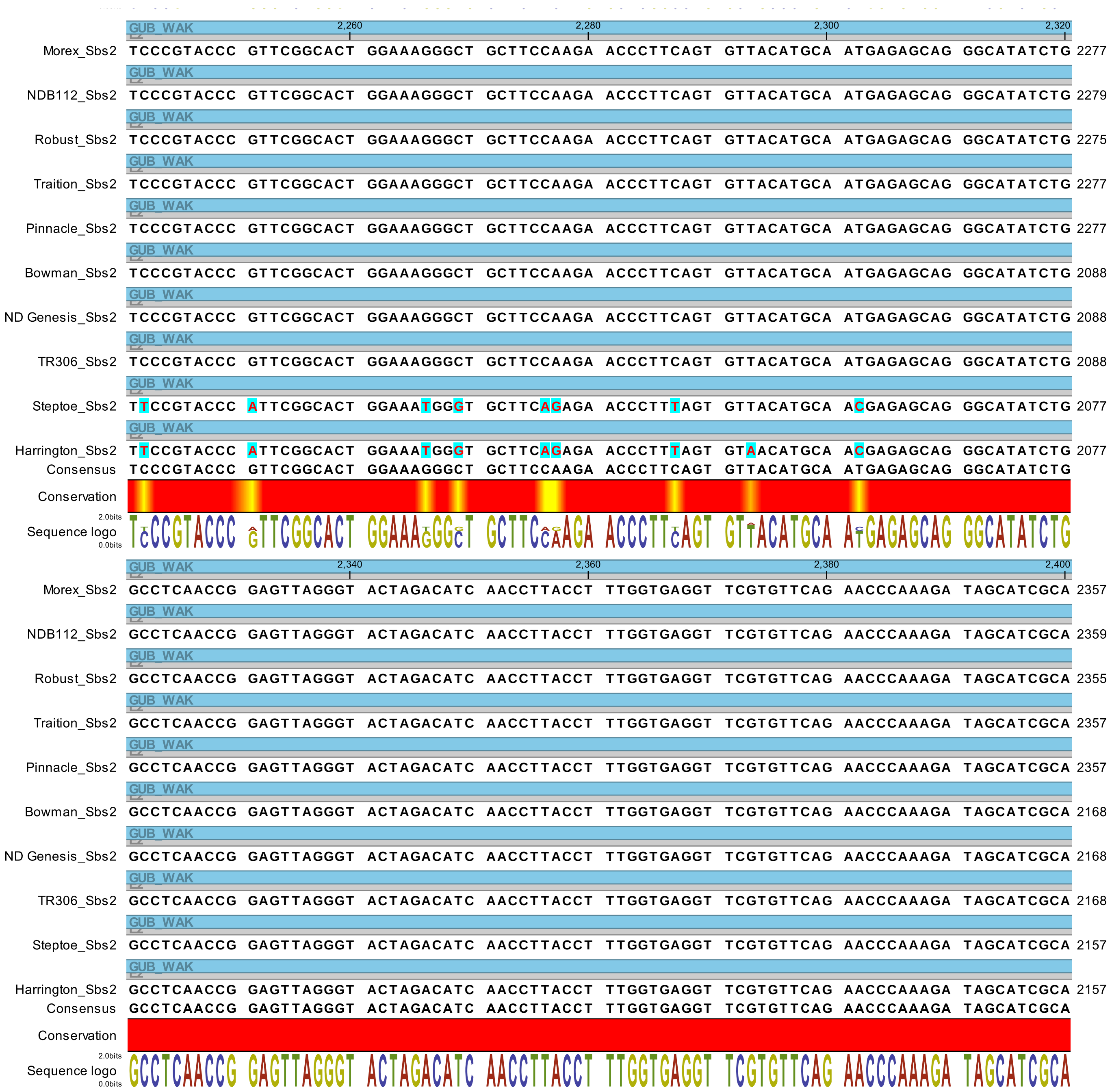

Supplementary Figure 6A

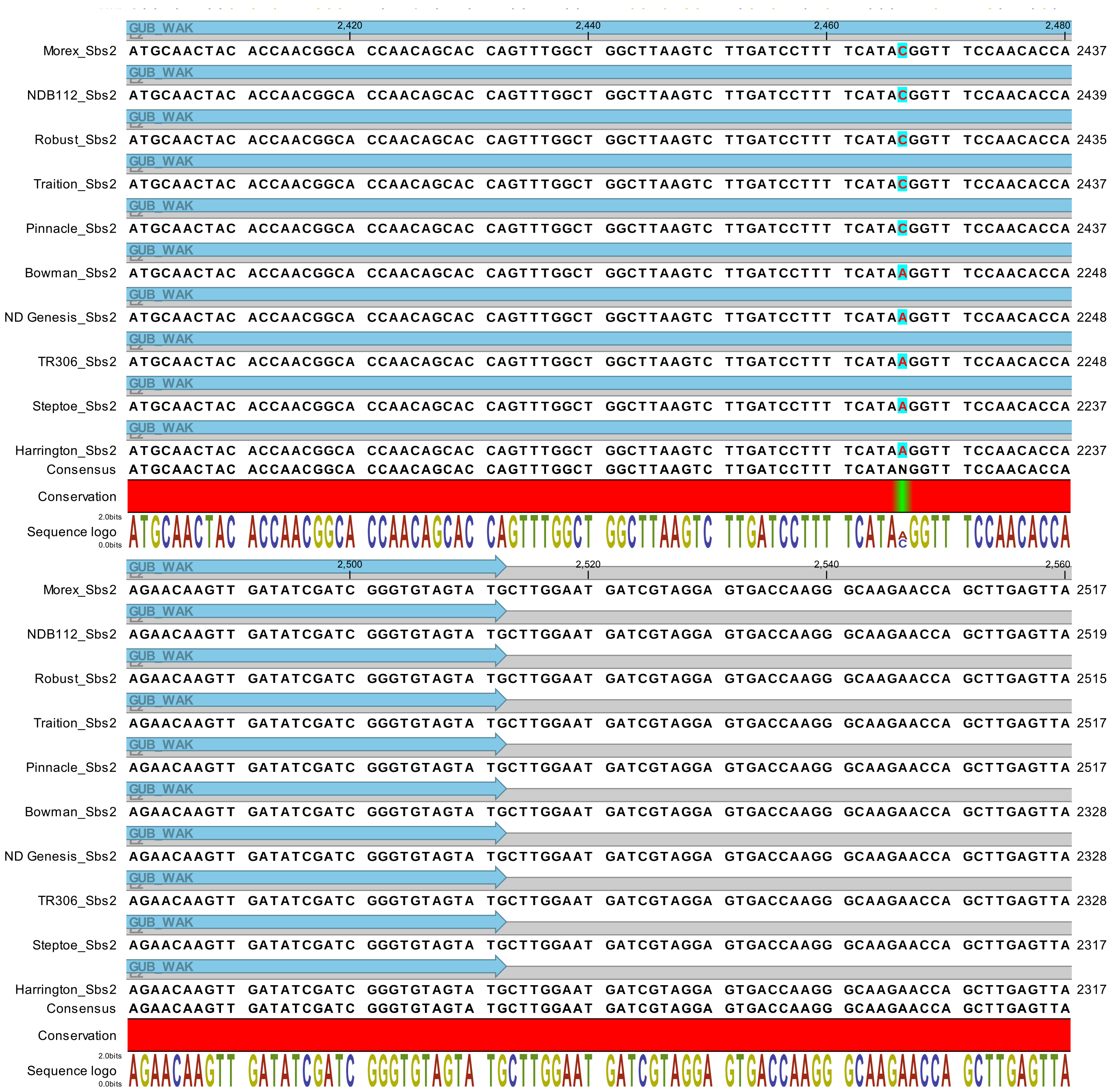

### Supplementary Figure 6A

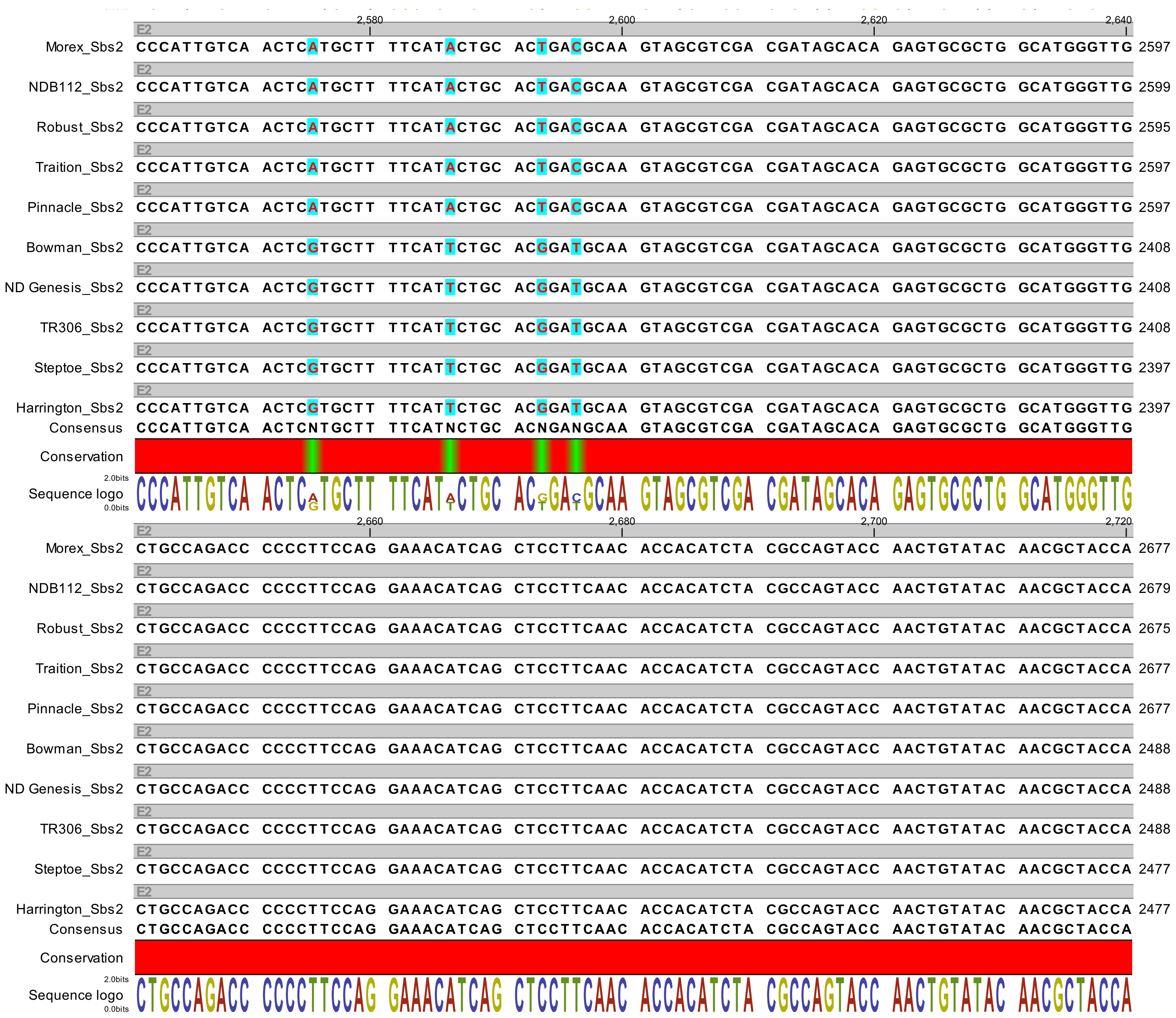

Supplementary Figure 6A

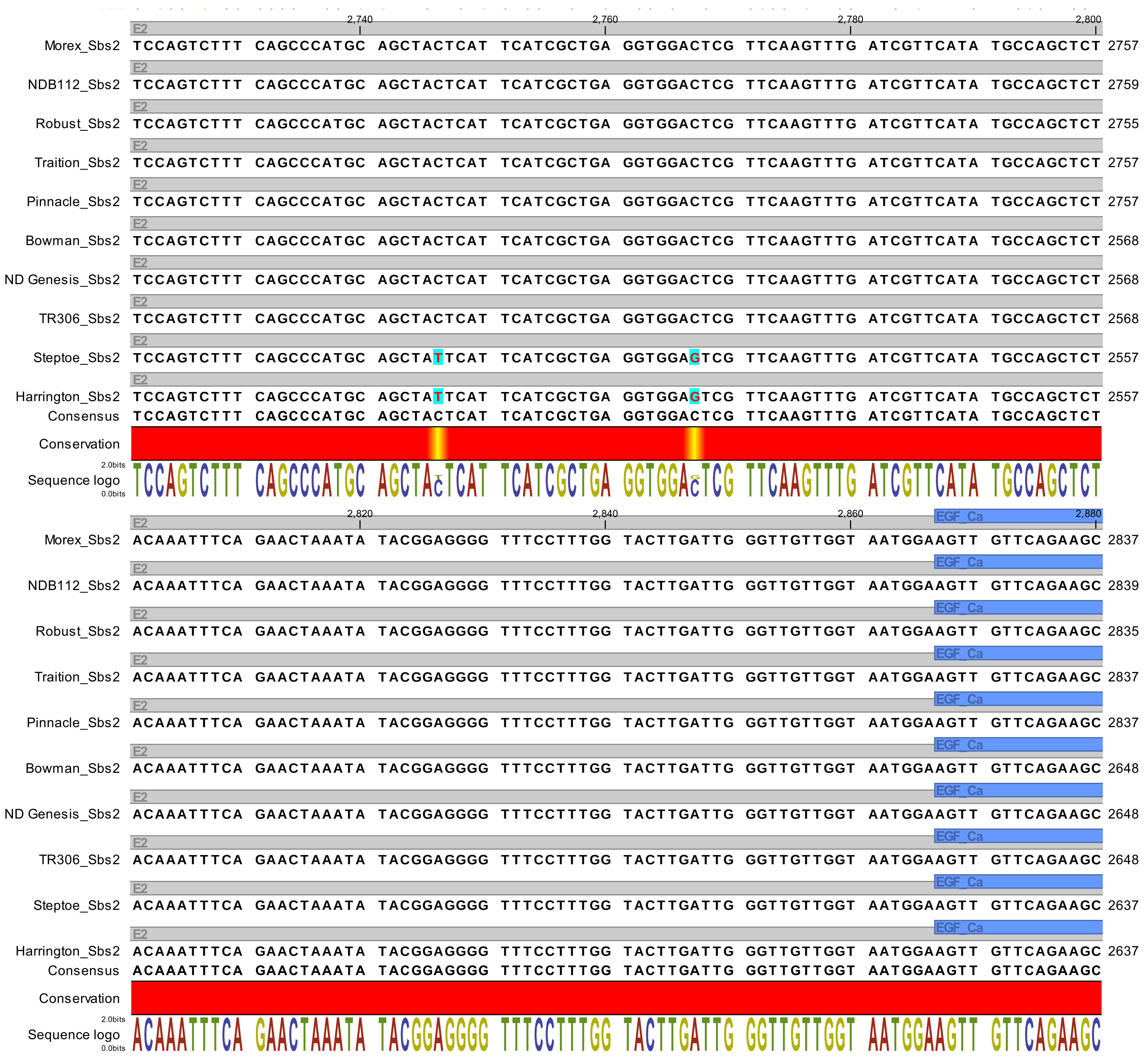

### Supplementary Figure 6A

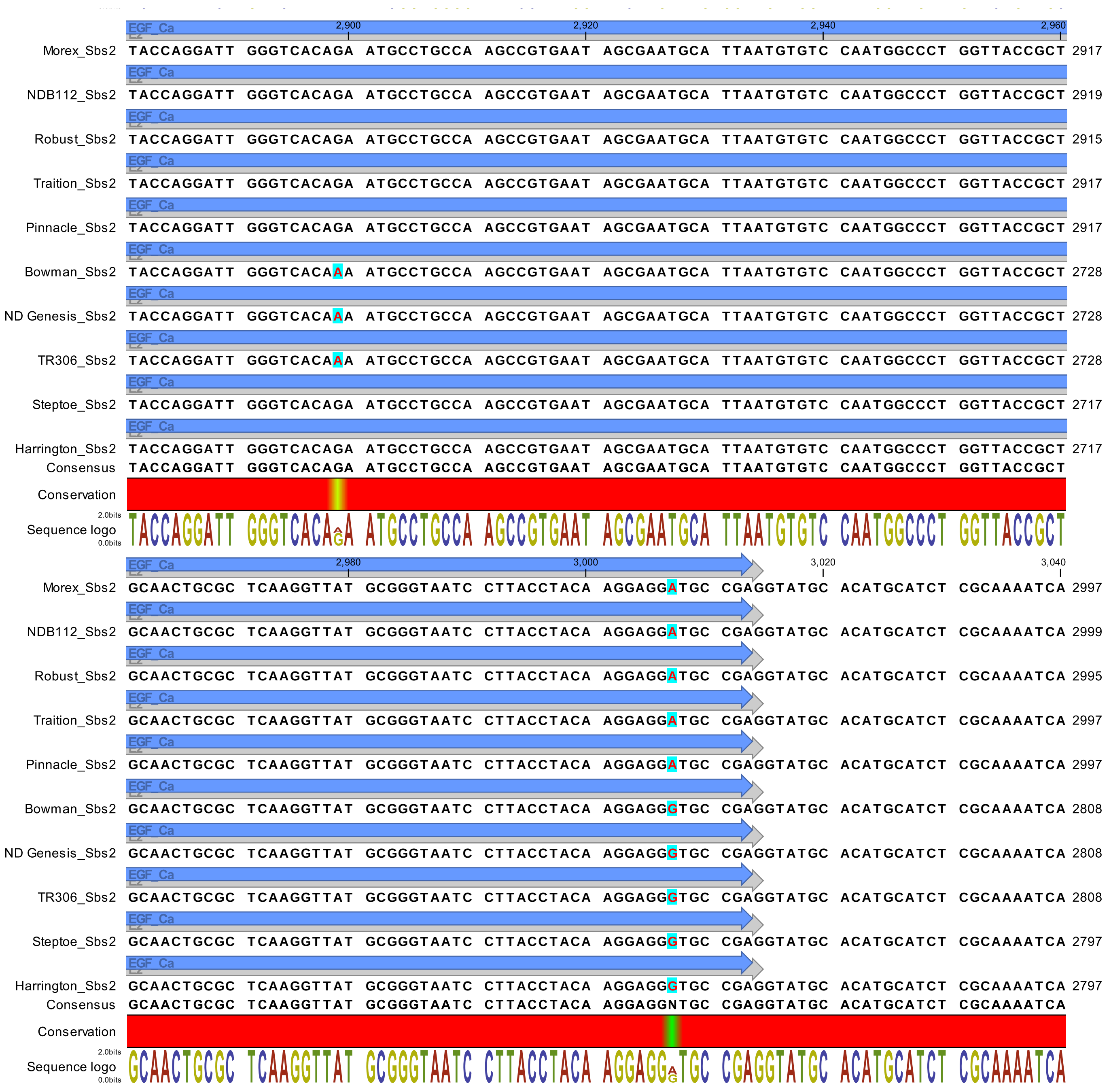

### Supplementary Figure 6A

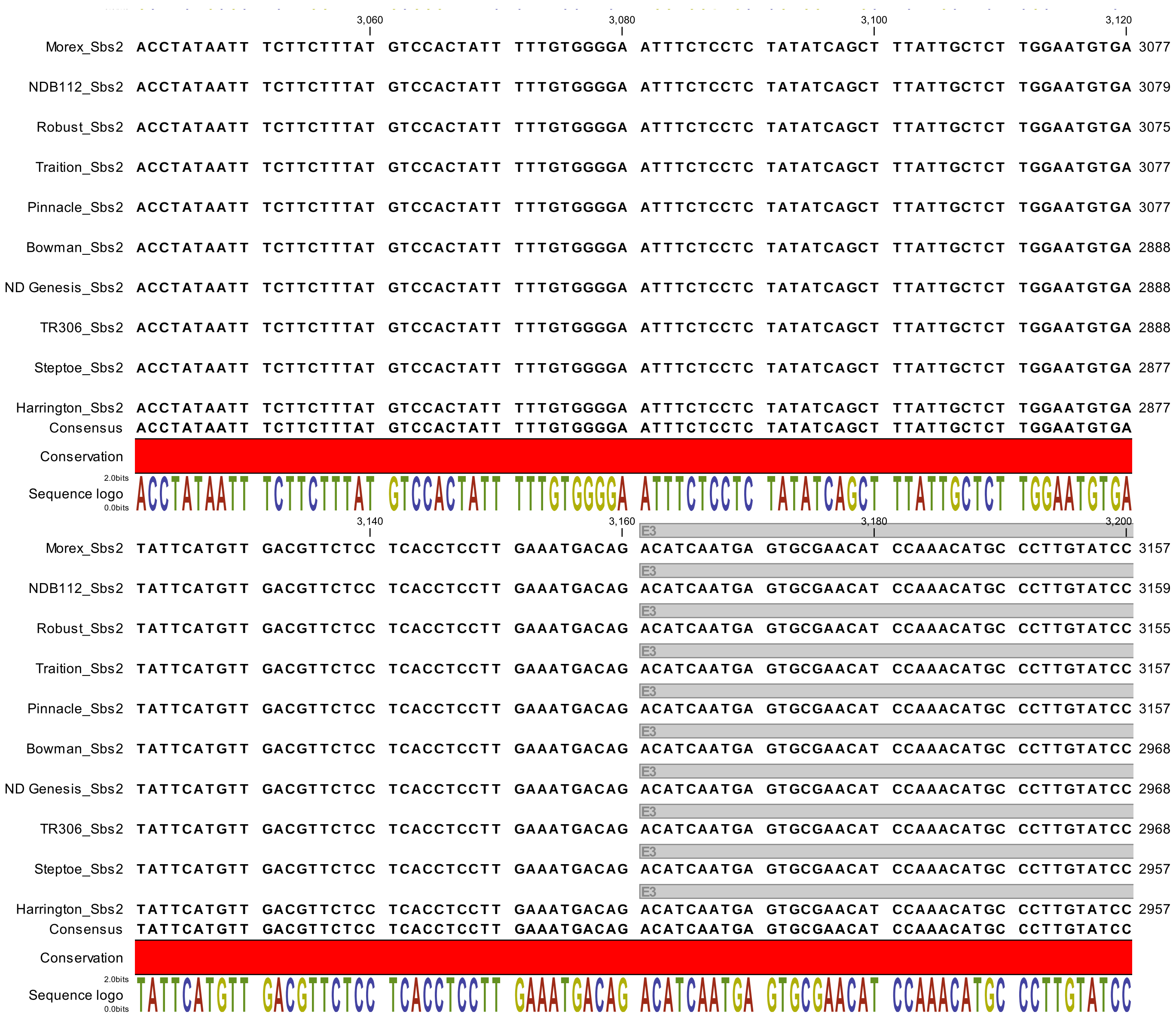

Supplementary Figure 6A

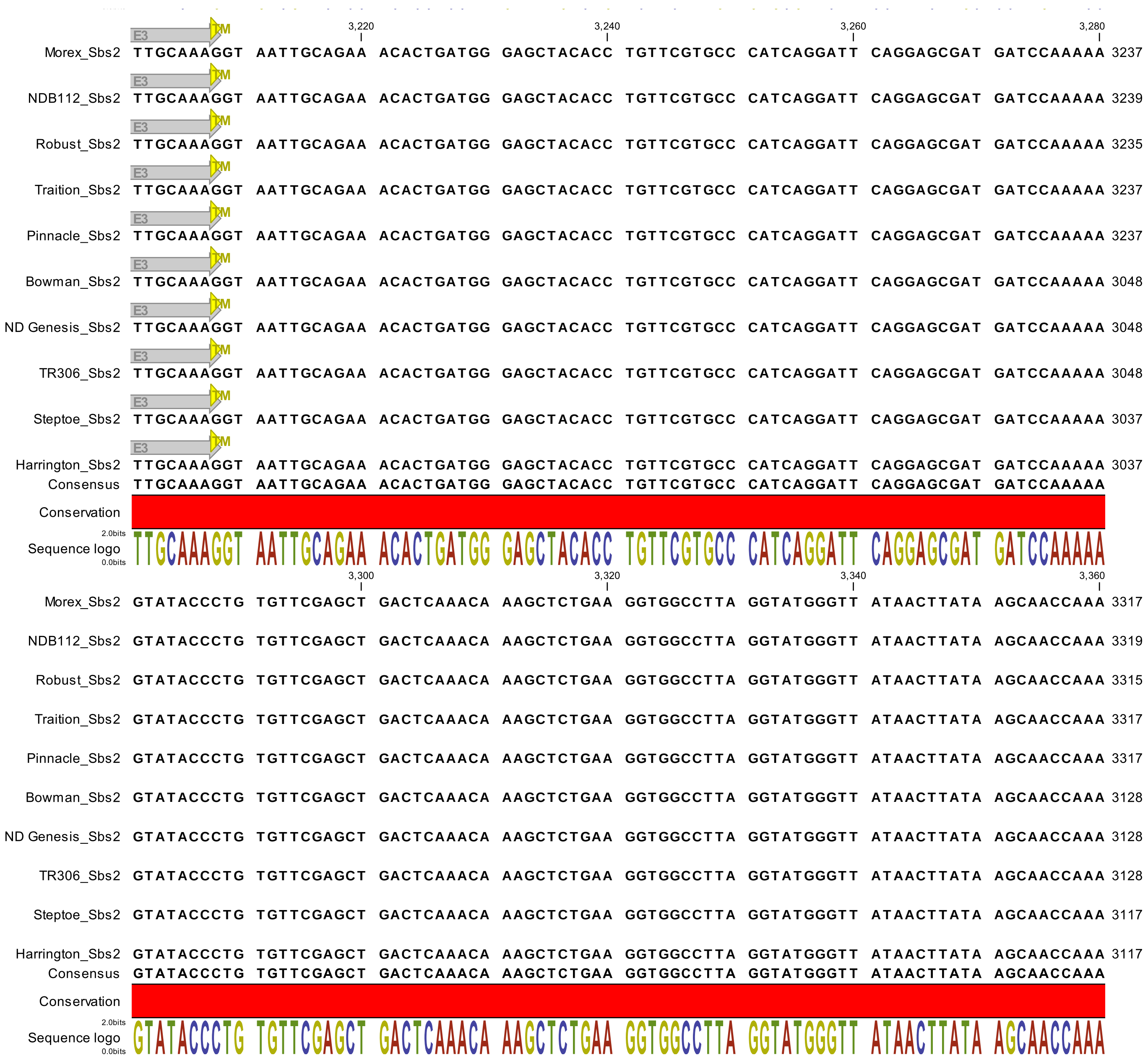

### Supplementary Figure 6A

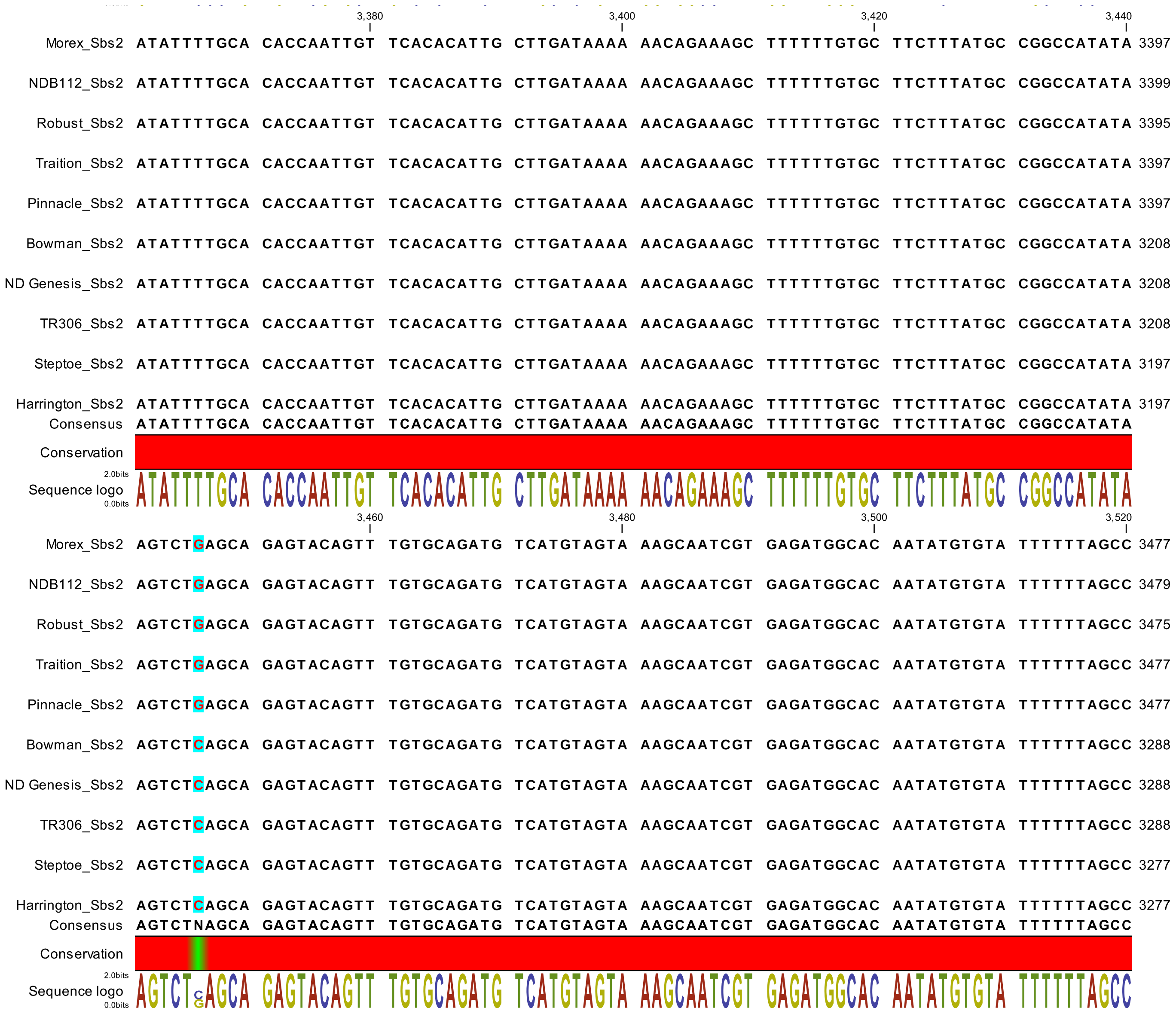

### Supplementary Figure 6A

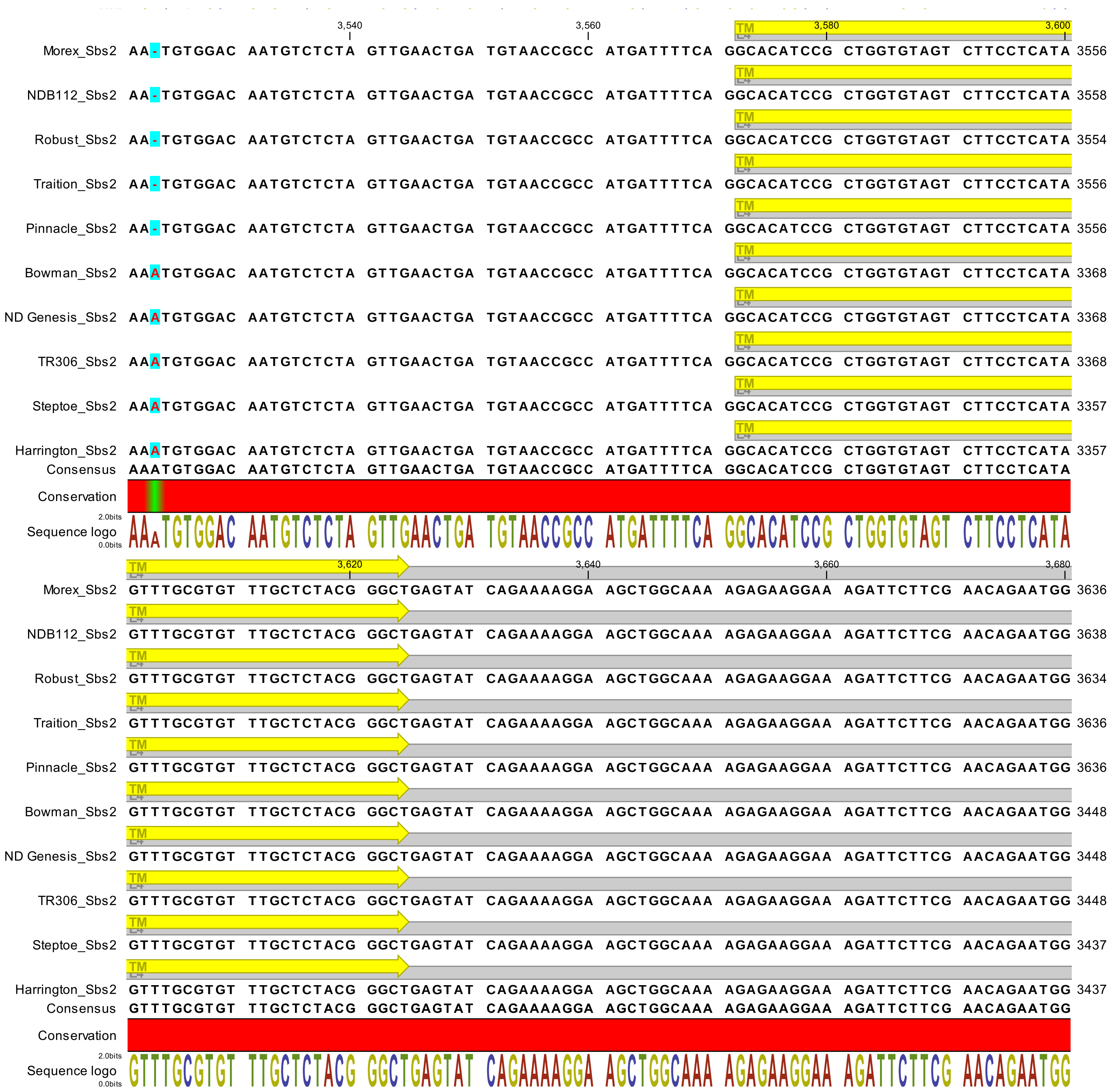

### Supplementary Figure 6A

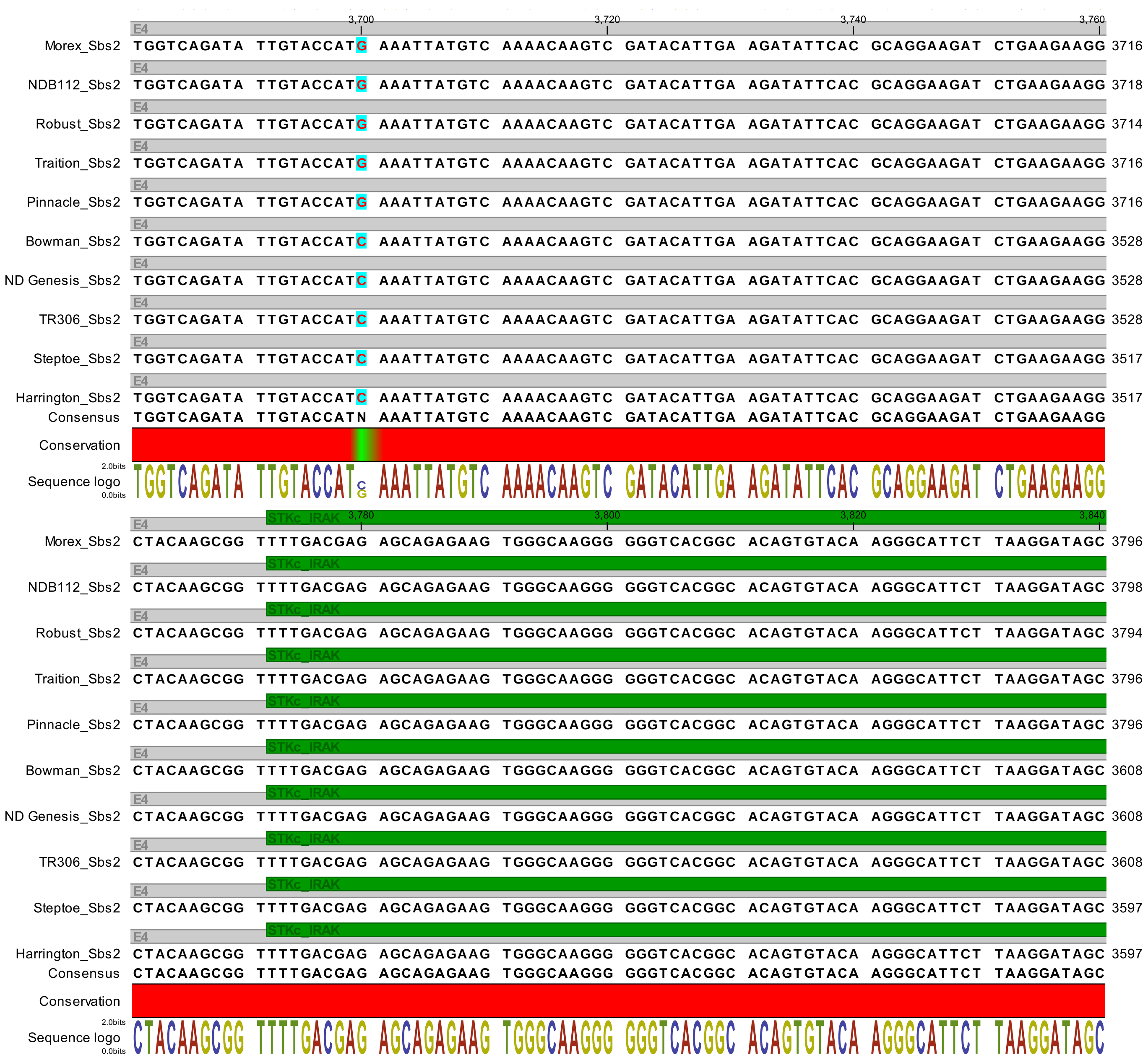

### Supplementary Figure 6A

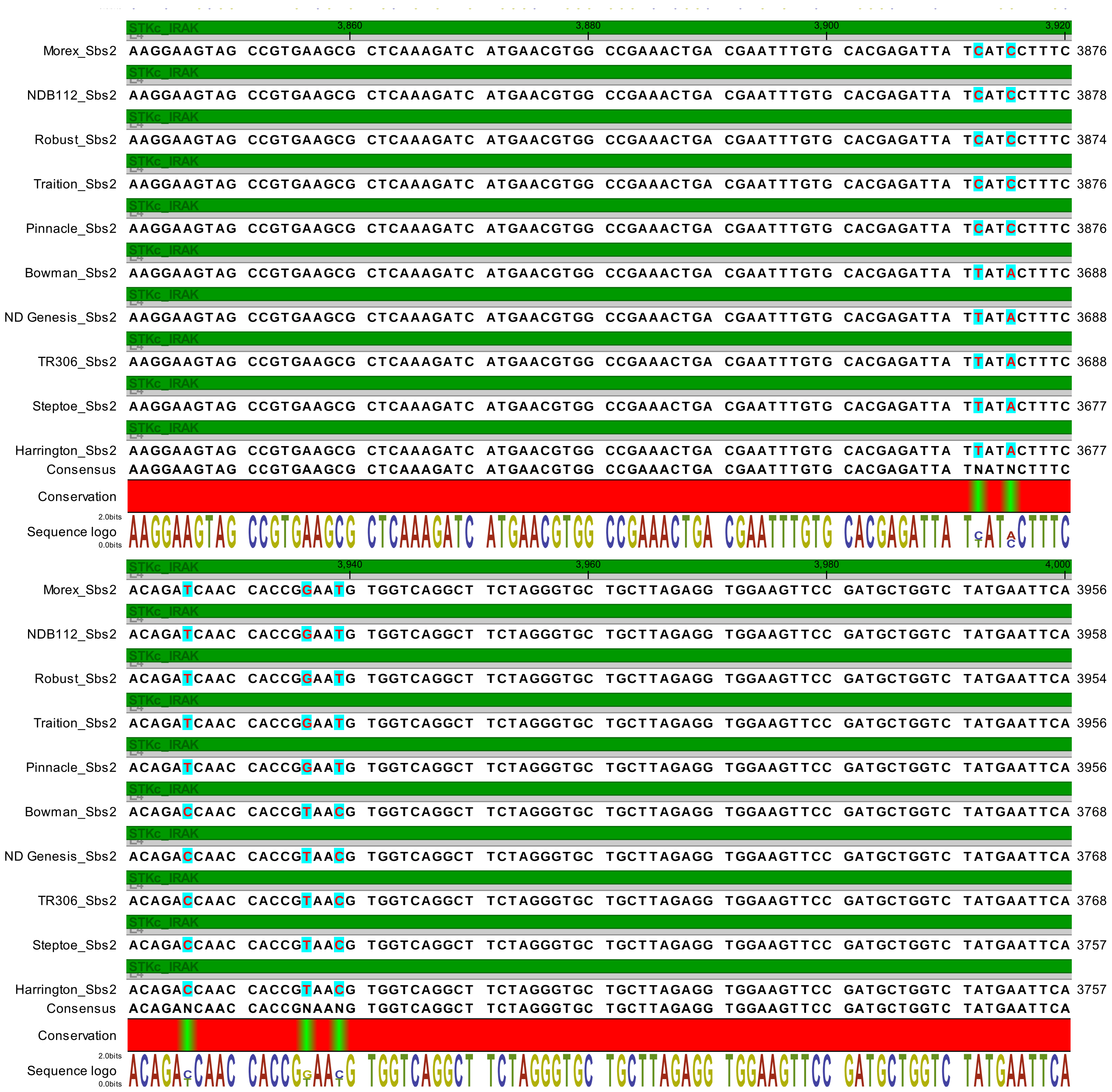

### Supplementary Figure 6A

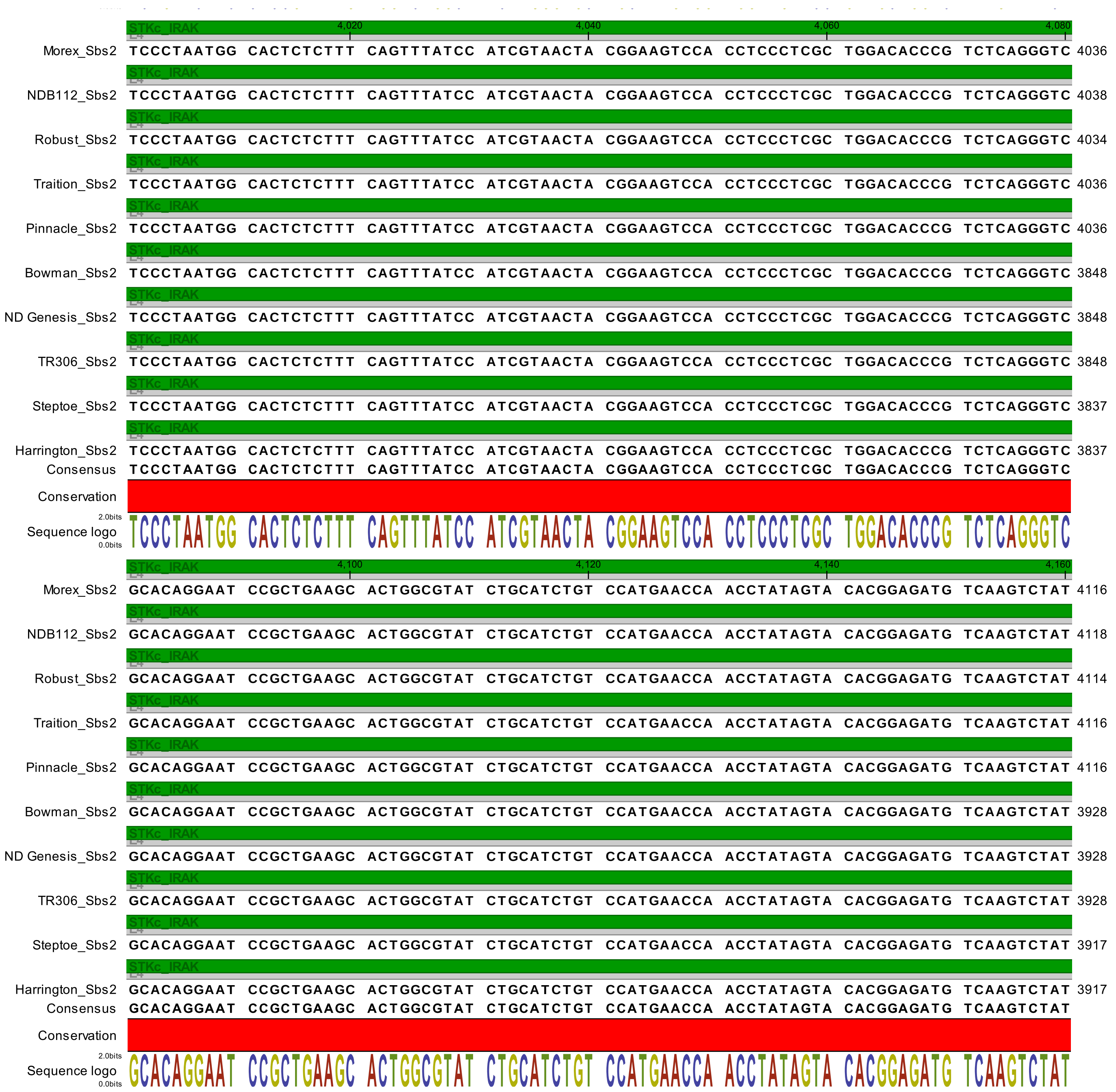

### Supplementary Figure 6A

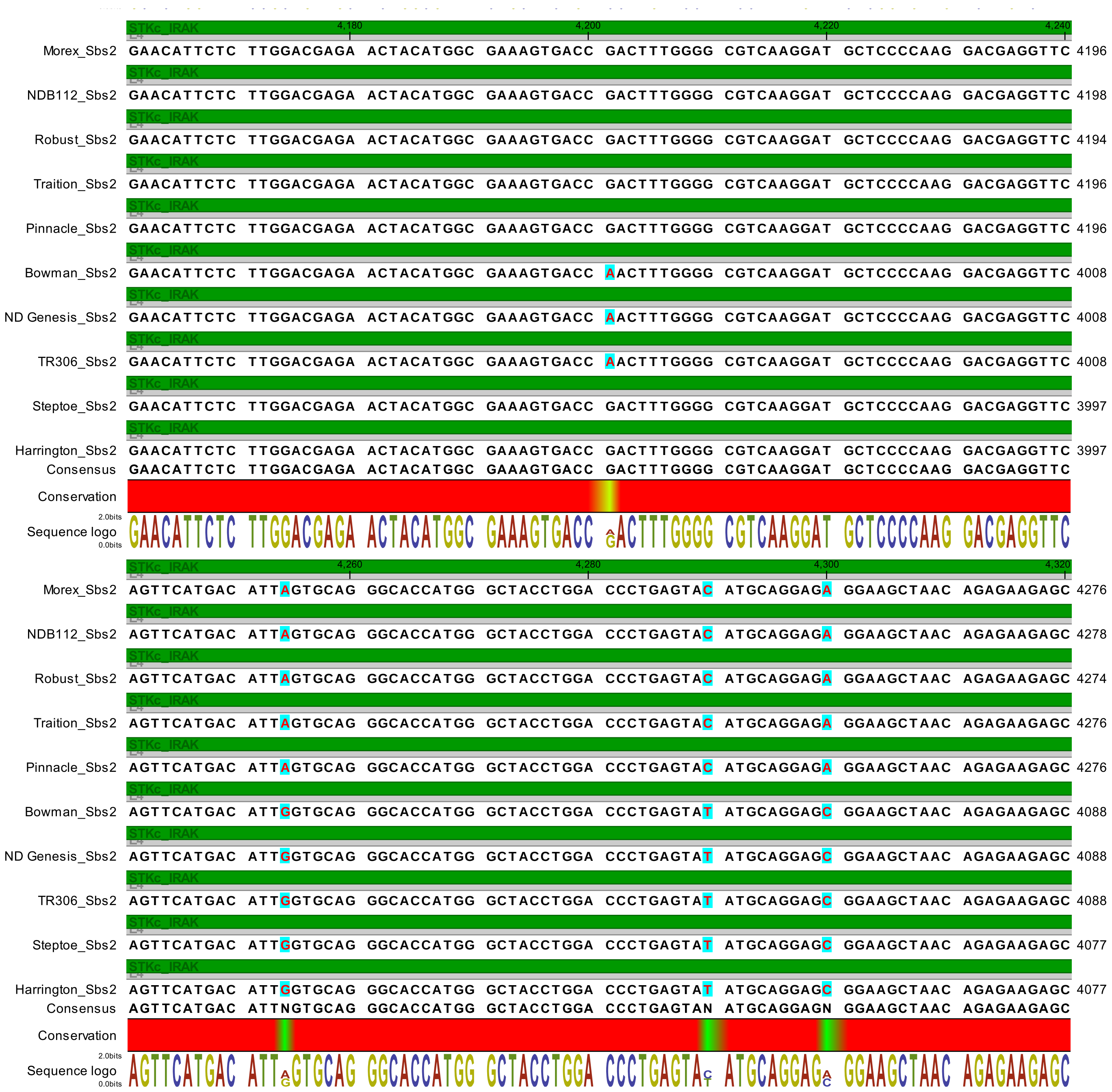

### Supplementary Figure 6A

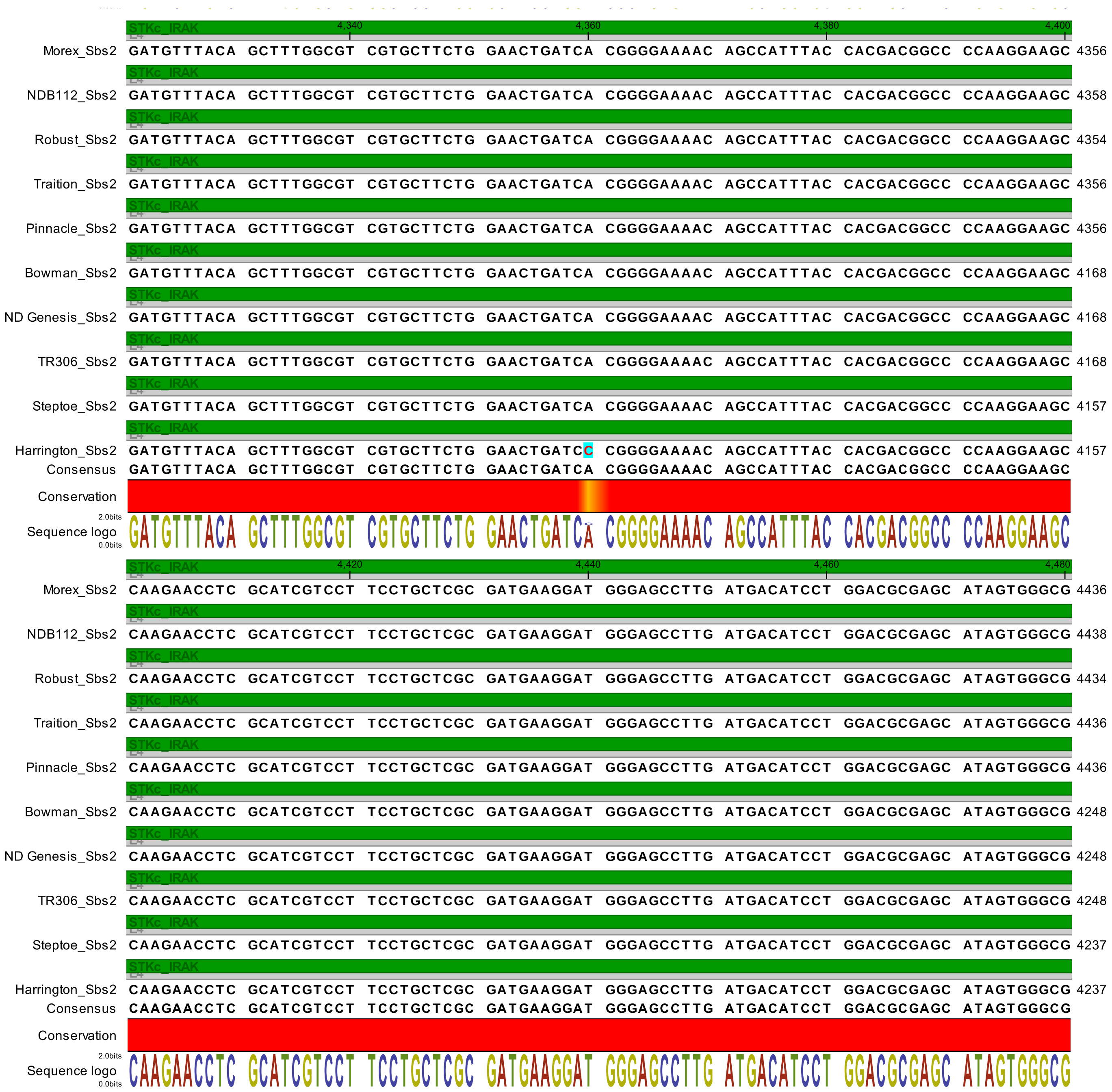

### Supplementary Figure 6A

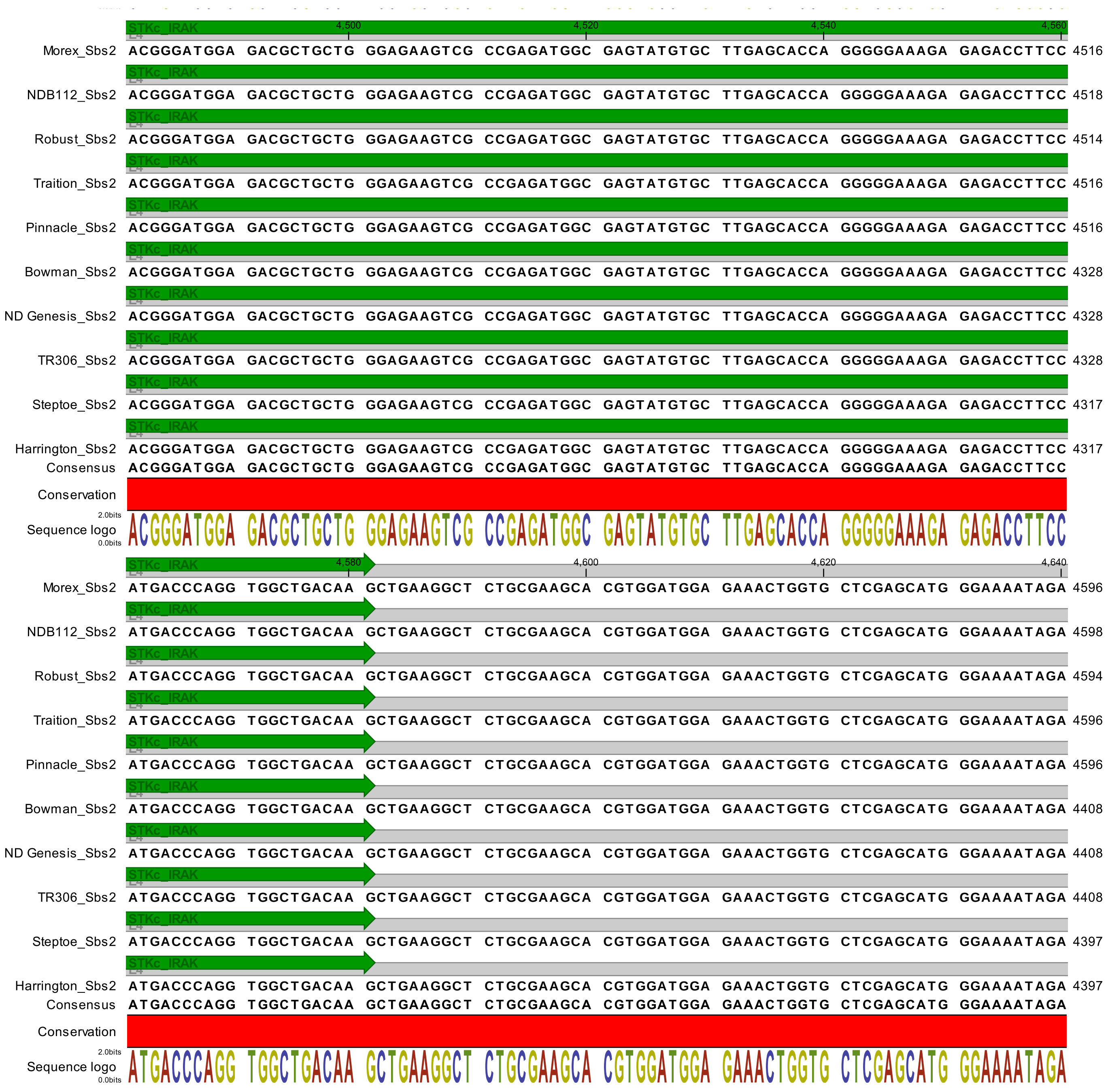

### Supplementary Figure 6A

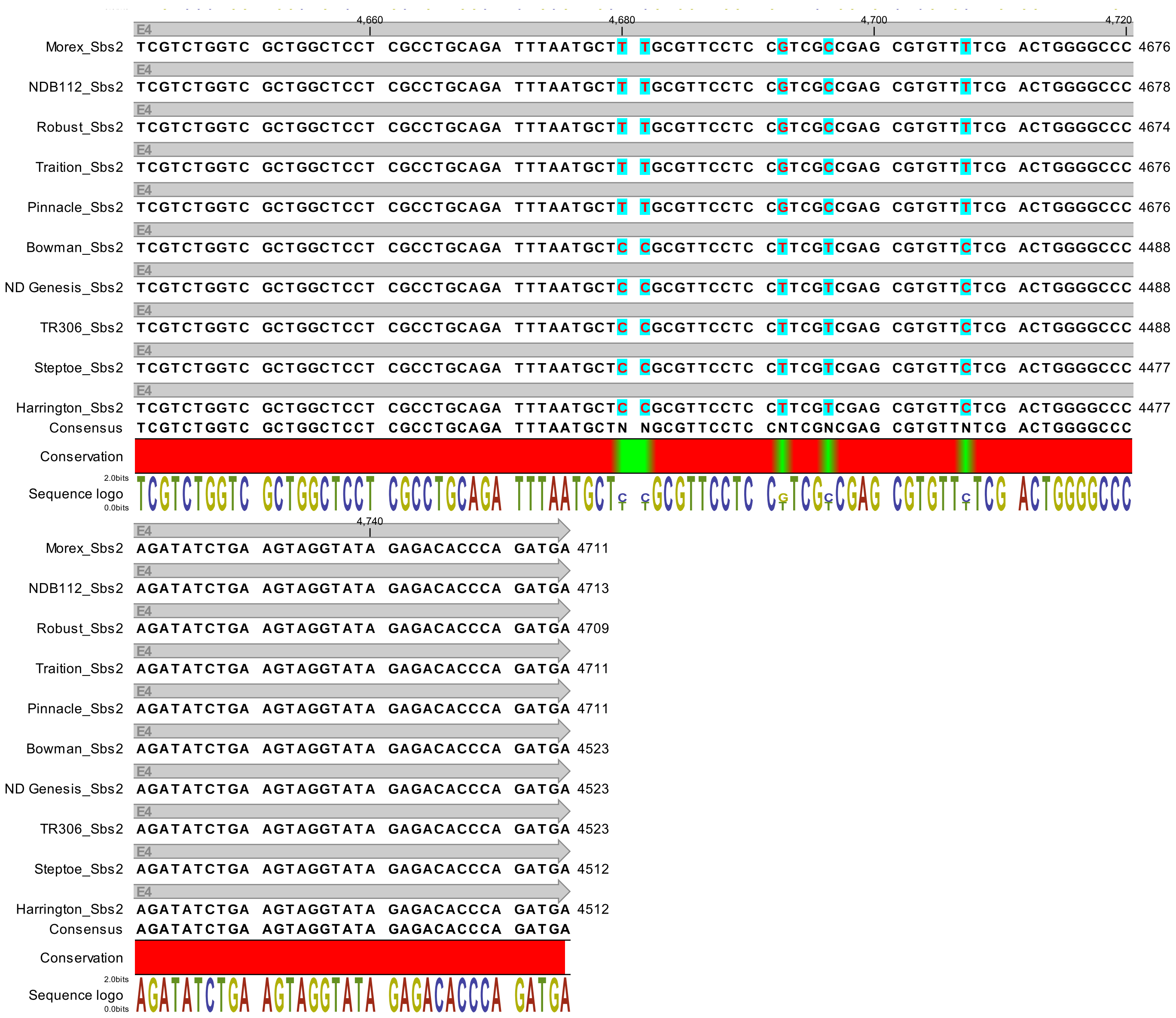

### Supplementary Figure 6B

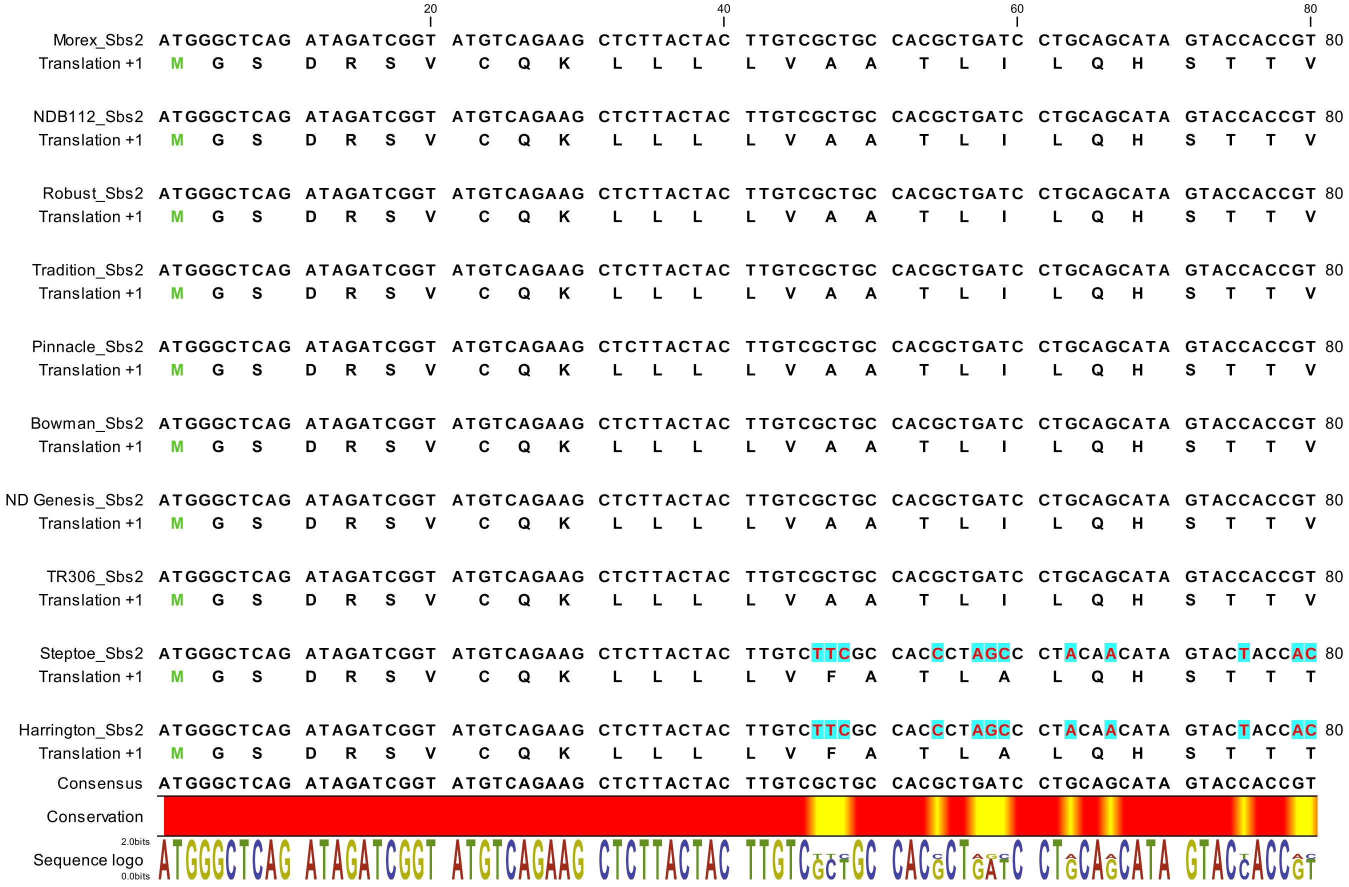

Supplementary Figure 6B

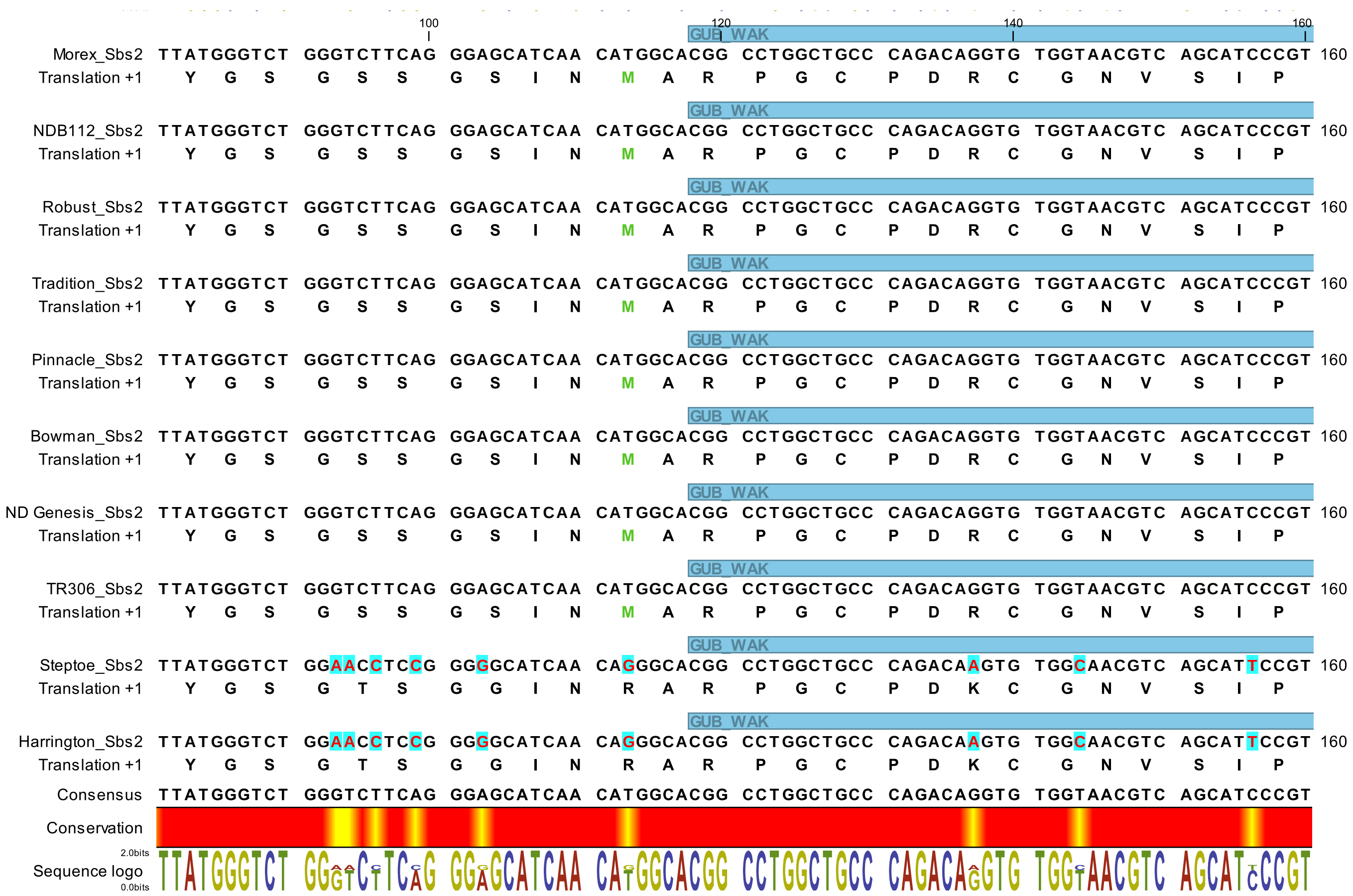

### Supplementary Figure 6B

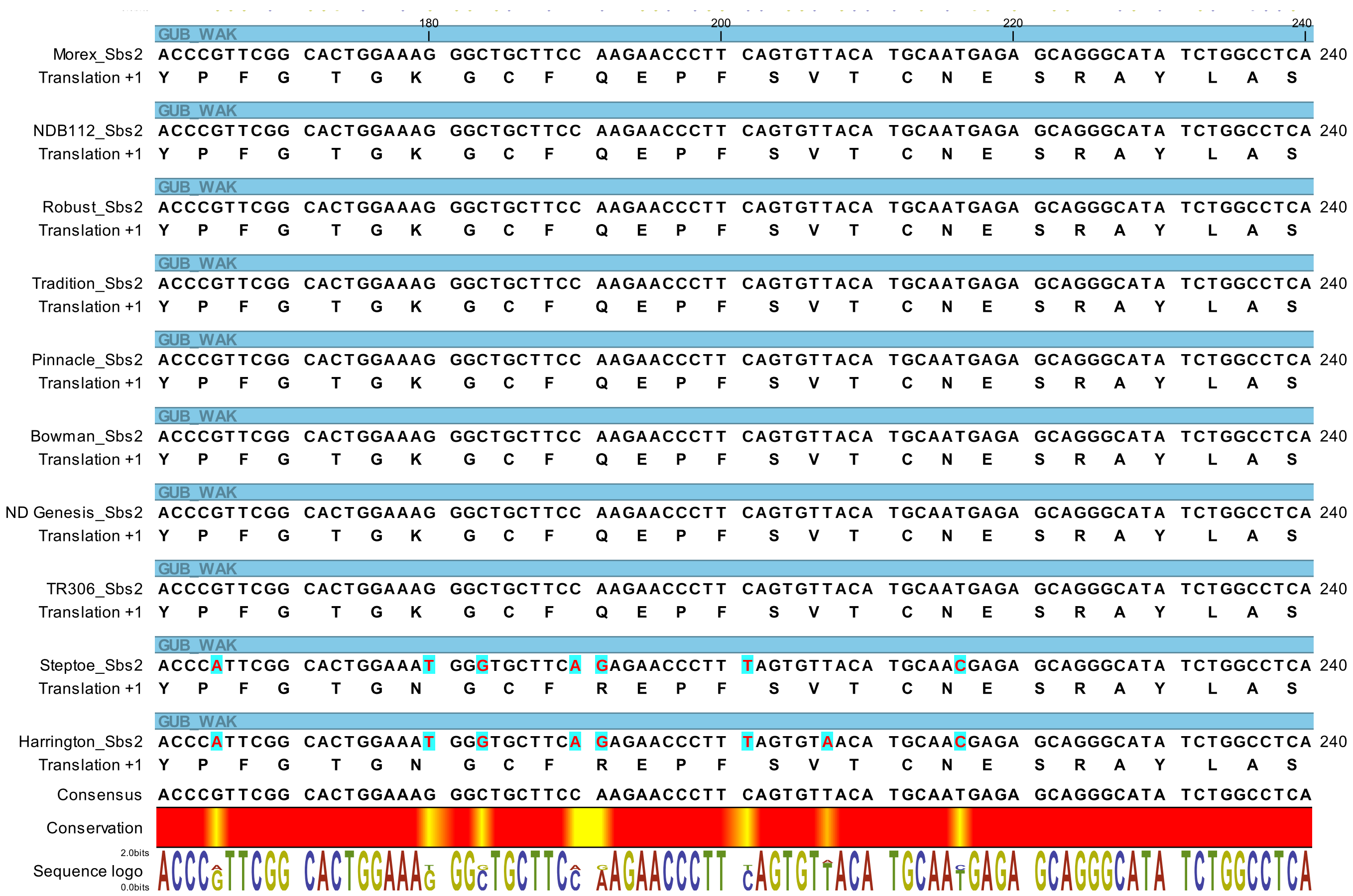

### Supplementary Figure 6B

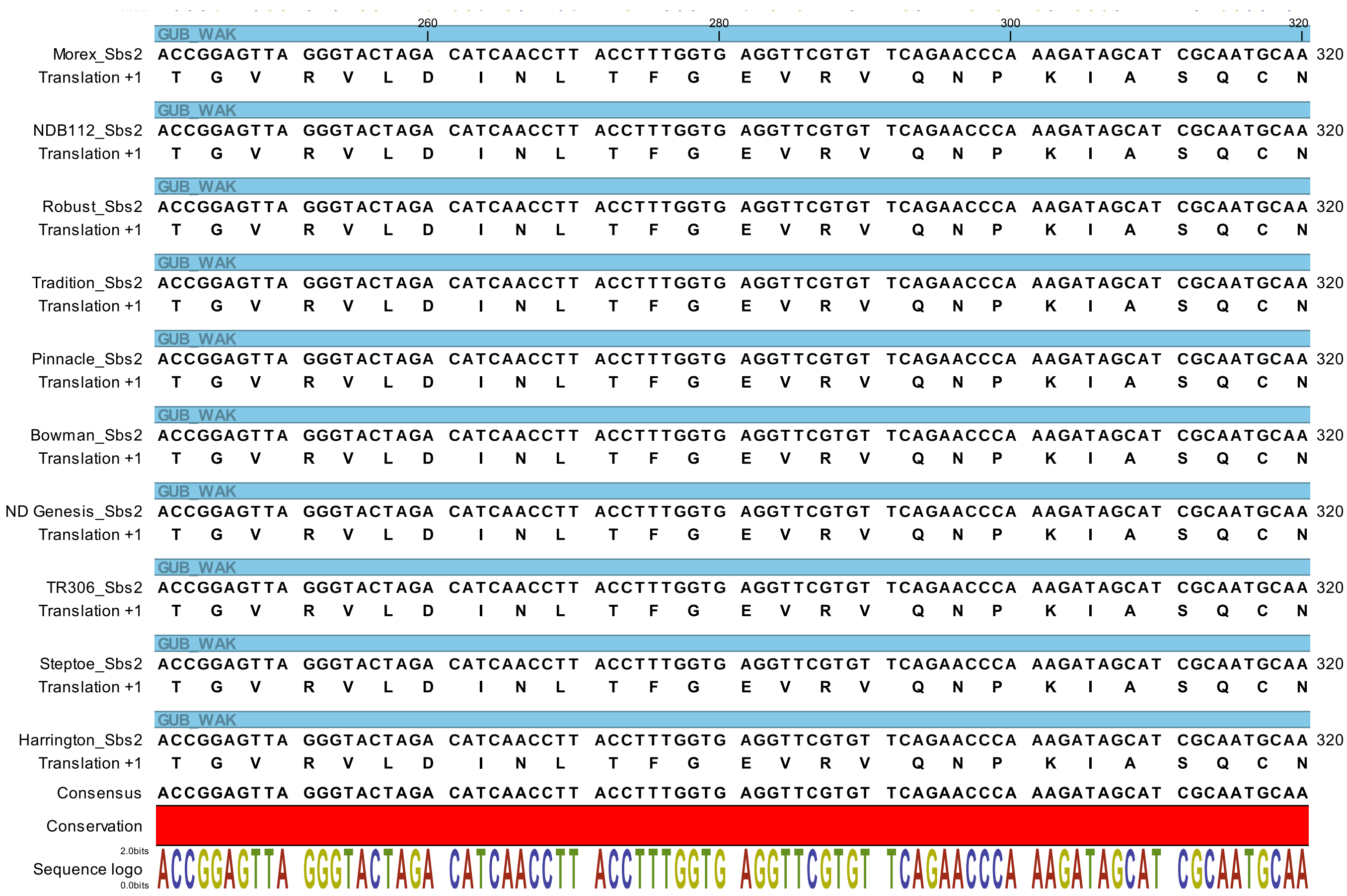

### Supplementary Figure 6B

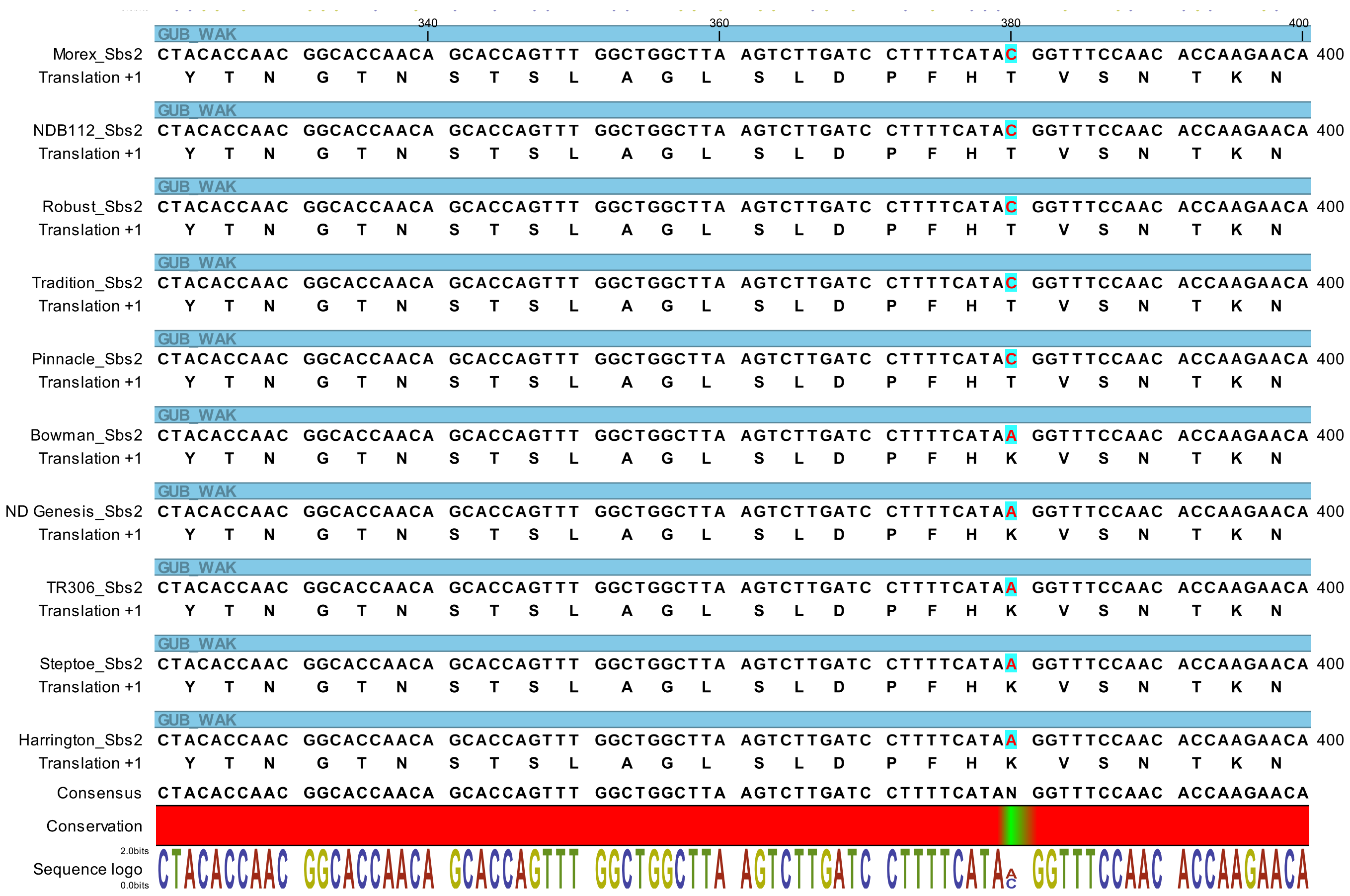

### Supplementary Figure 6B

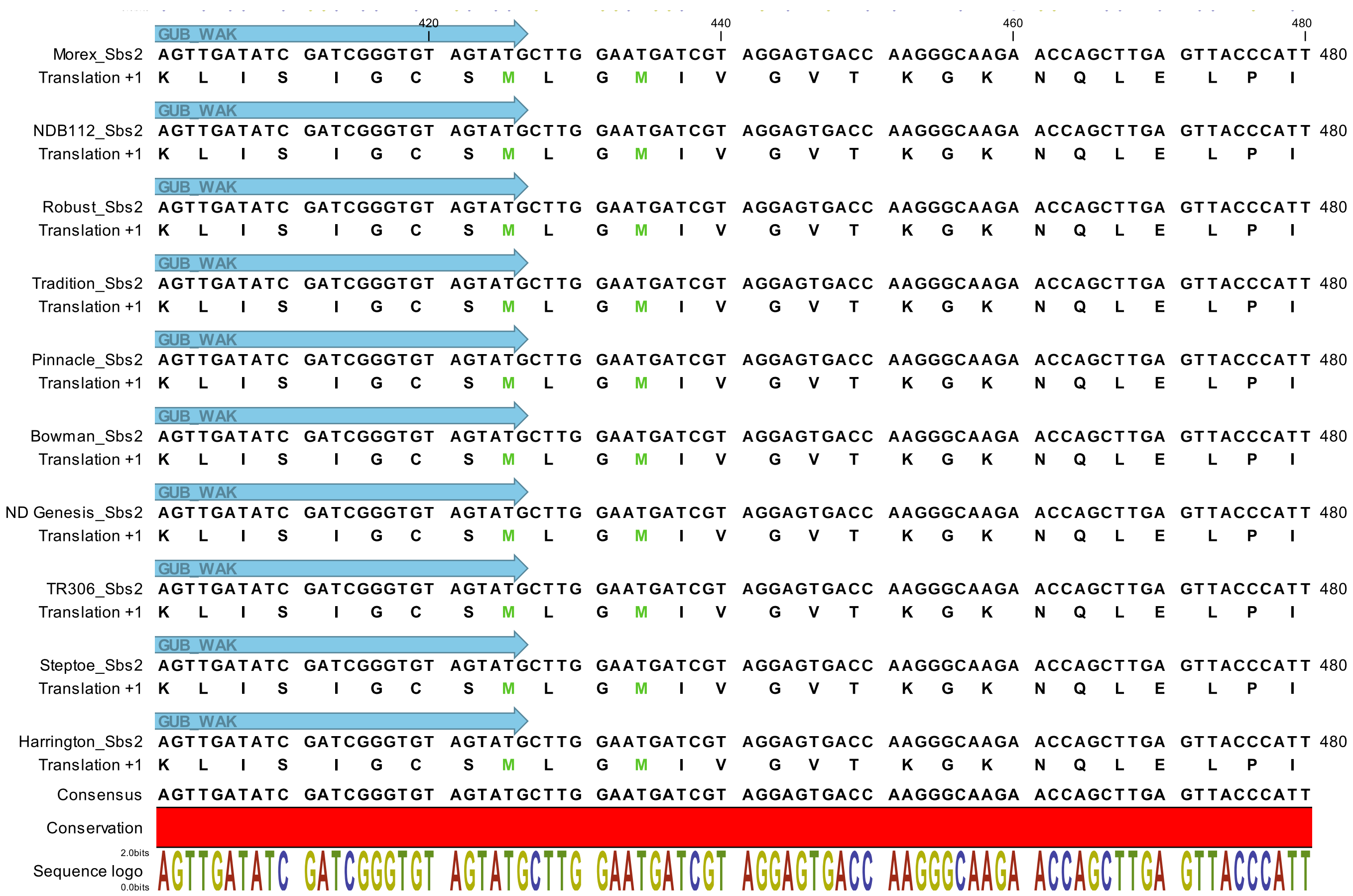

### Supplementary Figure 6B

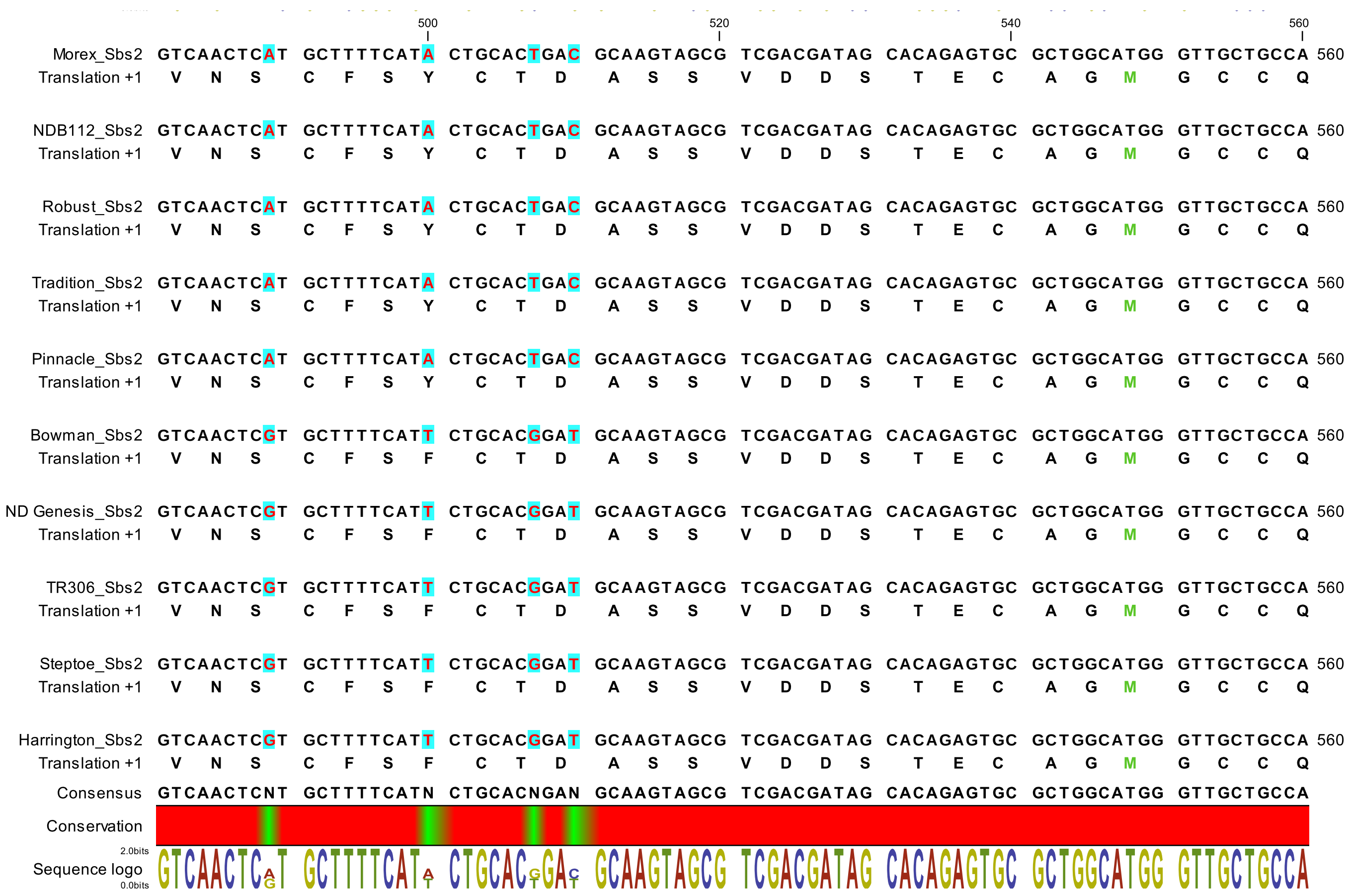

### Supplementary Figure 6B

| | 580 | | 600 | | 620 | | 640 | | |
|---|---|---|---|---|---|---|---|---|---|
| Morex_Sbs2 | GACCCCCCTT | CCAGGAAACA | TCAGCTCCTT | CAACACCACA | TCTACGCCAG | TACCAACTGT | ATACAACGCT | ACCATCCAGT | 640 |
| Translation +1 | T P L | P G N | I S S F | N T T | S T P | V P T V | Y N A | T I Q | |
| NDB112_Sbs2 | GACCCCCCTT | CCAGGAAACA | TCAGCTCCTT | CAACACCACA | TCTACGCCAG | TACCAACTGT | ATACAACGCT | ACCATCCAGT | 640 |
| Translation +1 | T P L | P G N | I S S F | N T T | S T P | V P T V | Y N A | T I Q | |
| Robust_Sbs2 | GACCCCCCTT | CCAGGAAACA | TCAGCTCCTT | CAACACCACA | TCTACGCCAG | TACCAACTGT | ATACAACGCT | ACCATCCAGT | 640 |
| Translation +1 | T P L | P G N | I S S F | N T T | S T P | V P T V | Y N A | T I Q | |
| Tradition_Sbs2 | GACCCCCCTT | CCAGGAAACA | TCAGCTCCTT | CAACACCACA | TCTACGCCAG | TACCAACTGT | ATACAACGCT | ACCATCCAGT | 640 |
| Translation +1 | T P L | P G N | I S S F | N T T | S T P | V P T V | Y N A | T I Q | |
| Pinnacle_Sbs2 | GACCCCCCTT | CCAGGAAACA | TCAGCTCCTT | CAACACCACA | TCTACGCCAG | TACCAACTGT | ATACAACGCT | ACCATCCAGT | 640 |
| Translation +1 | T P L | P G N | I S S F | N T T | S T P | V P T V | Y N A | T I Q | |
| Bowman_Sbs2 | GACCCCCCTT | CCAGGAAACA | TCAGCTCCTT | CAACACCACA | TCTACGCCAG | TACCAACTGT | ATACAACGCT | ACCATCCAGT | 640 |
| Translation +1 | T P L | P G N | I S S F | N T T | S T P | V P T V | Y N A | T I Q | |
| ND Genesis_Sbs2 | GACCCCCCTT | CCAGGAAACA | TCAGCTCCTT | CAACACCACA | TCTACGCCAG | TACCAACTGT | ATACAACGCT | ACCATCCAGT | 640 |
| Translation +1 | T P L | P G N | I S S F | N T T | S T P | V P T V | Y N A | T I Q | |
| TR306_Sbs2 | GACCCCCCTT | CCAGGAAACA | TCAGCTCCTT | CAACACCACA | TCTACGCCAG | TACCAACTGT | ATACAACGCT | ACCATCCAGT | 640 |
| Translation +1 | T P L | P G N | I S S F | N T T | S T P | V P T V | Y N A | T I Q | |
| Steptoe_Sbs2 | GACCCCCCTT | CCAGGAAACA | TCAGCTCCTT | CAACACCACA | TCTACGCCAG | TACCAACTGT | ATACAACGCT | ACCATCCAGT | 640 |
| Translation +1 | T P L | P G N | I S S F | N T T | S T P | V P T V | Y N A | T I Q | |
| Harrington_Sbs2 | GACCCCCCTT | CCAGGAAACA | TCAGCTCCTT | CAACACCACA | TCTACGCCAG | TACCAACTGT | ATACAACGCT | ACCATCCAGT | 640 |
| Translation +1 | T P L | P G N | I S S F | N T T | S T P | V P T V | Y N A | T I Q | |
| Consensus | GACCCCCCTT | CCAGGAAACA | TCAGCTCCTT | CAACACCACA | TCTACGCCAG | TACCAACTGT | ATACAACGCT | ACCATCCAGT | |

Conservation

Sequence logo

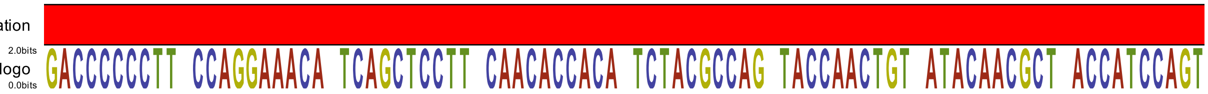

### Supplementary Figure 6B

| | 660 | | 680 | | 700 | | 720 | |
|---|---|---|---|---|---|---|---|---|
| Morex_Sbs2 | CTTTCAGCCC ATGCAGCTAC | TCATTCATCG | CTGAGGTGGA | CTCGTTCAAG | TTTGATCGTT | CATATGCCAG | CTCTACAAAT | 720 |
| Translation +1 | S F S P C S Y | S F I | A E V D | S F K | F D R | S Y A S | S T N | |
| NDB112_Sbs2 | CTTTCAGCCC ATGCAGCTAC | TCATTCATCG | CTGAGGTGGA | CTCGTTCAAG | TTTGATCGTT | CATATGCCAG | CTCTACAAAT | 720 |
| Translation +1 | S F S P C S Y | S F I | A E V D | S F K | F D R | S Y A S | S T N | |
| Robust_Sbs2 | CTTTCAGCCC ATGCAGCTAC | TCATTCATCG | CTGAGGTGGA | CTCGTTCAAG | TTTGATCGTT | CATATGCCAG | CTCTACAAAT | 720 |
| Translation +1 | S F S P C S Y | S F I | A E V D | S F K | F D R | S Y A S | S T N | |
| Tradition_Sbs2 | CTTTCAGCCC ATGCAGCTAC | TCATTCATCG | CTGAGGTGGA | CTCGTTCAAG | TTTGATCGTT | CATATGCCAG | CTCTACAAAT | 720 |
| Translation +1 | S F S P C S Y | S F I | A E V D | S F K | F D R | S Y A S | S T N | |
| Pinnacle_Sbs2 | CTTTCAGCCC ATGCAGCTAC | TCATTCATCG | CTGAGGTGGA | CTCGTTCAAG | TTTGATCGTT | CATATGCCAG | CTCTACAAAT | 720 |
| Translation +1 | S F S P C S Y | S F I | A E V D | S F K | F D R | S Y A S | S T N | |
| Bowman_Sbs2 | CTTTCAGCCC ATGCAGCTAC | TCATTCATCG | CTGAGGTGGA | CTCGTTCAAG | TTTGATCGTT | CATATGCCAG | CTCTACAAAT | 720 |
| Translation +1 | S F S P C S Y | S F I | A E V D | S F K | F D R | S Y A S | S T N | |
| ND Genesis_Sbs2 | CTTTCAGCCC ATGCAGCTAC | TCATTCATCG | CTGAGGTGGA | CTCGTTCAAG | TTTGATCGTT | CATATGCCAG | CTCTACAAAT | 720 |
| Translation +1 | S F S P C S Y | S F I | A E V D | S F K | F D R | S Y A S | S T N | |
| TR306_Sbs2 | CTTTCAGCCC ATGCAGCTAC | TCATTCATCG | CTGAGGTGGA | CTCGTTCAAG | TTTGATCGTT | CATATGCCAG | CTCTACAAAT | 720 |
| Translation +1 | S F S P C S Y | S F I | A E V D | S F K | F D R | S Y A S | S T N | |
| Steptoe_Sbs2 | CTTTCAGCCC ATGCAGCTAT | TCATTCATCG | CTGAGGTGGA | GTCGTTCAAG | TTTGATCGTT | CATATGCCAG | CTCTACAAAT | 720 |
| Translation +1 | S F S P C S Y | S F I | A E V E | S F K | F D R | S Y A S | S T N | |
| Harrington_Sbs2 | CTTTCAGCCC ATGCAGCTAT | TCATTCATCG | CTGAGGTGGA | GTCGTTCAAG | TTTGATCGTT | CATATGCCAG | CTCTACAAAT | 720 |
| Translation +1 | S F S P C S Y | S F I | A E V E | S F K | F D R | S Y A S | S T N | |
| Consensus | CTTTCAGCCC ATGCAGCTAC | TCATTCATCG | CTGAGGTGGA | CTCGTTCAAG | TTTGATCGTT | CATATGCCAG | CTCTACAAAT | |

Conservation

Sequence logo

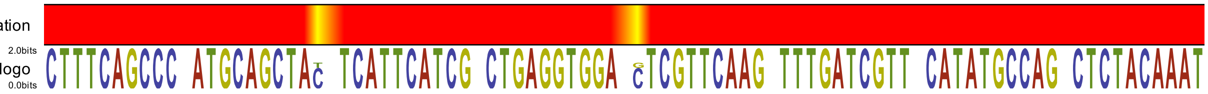

### Supplementary Figure 6B

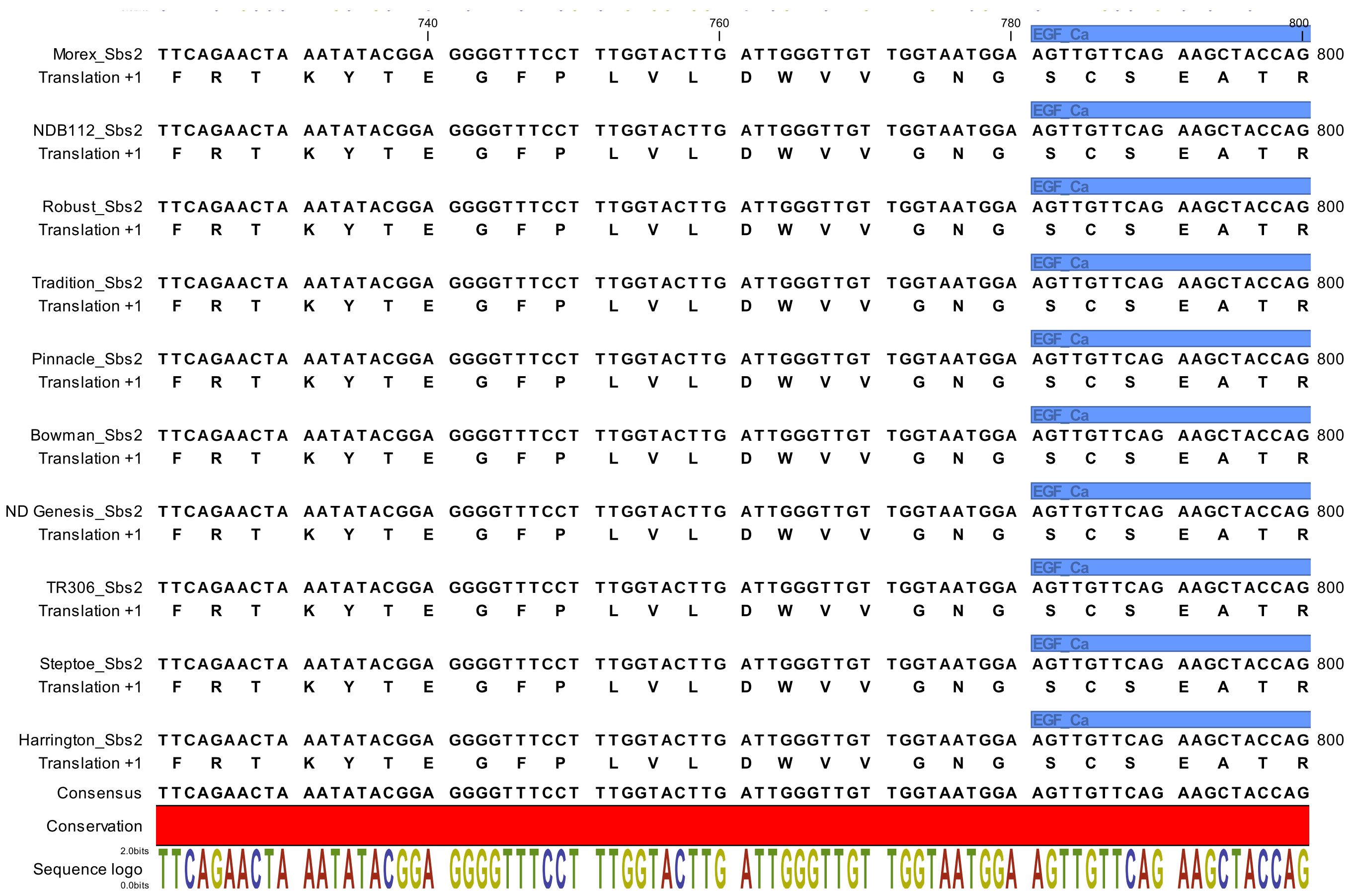

Supplementary Figure 6B

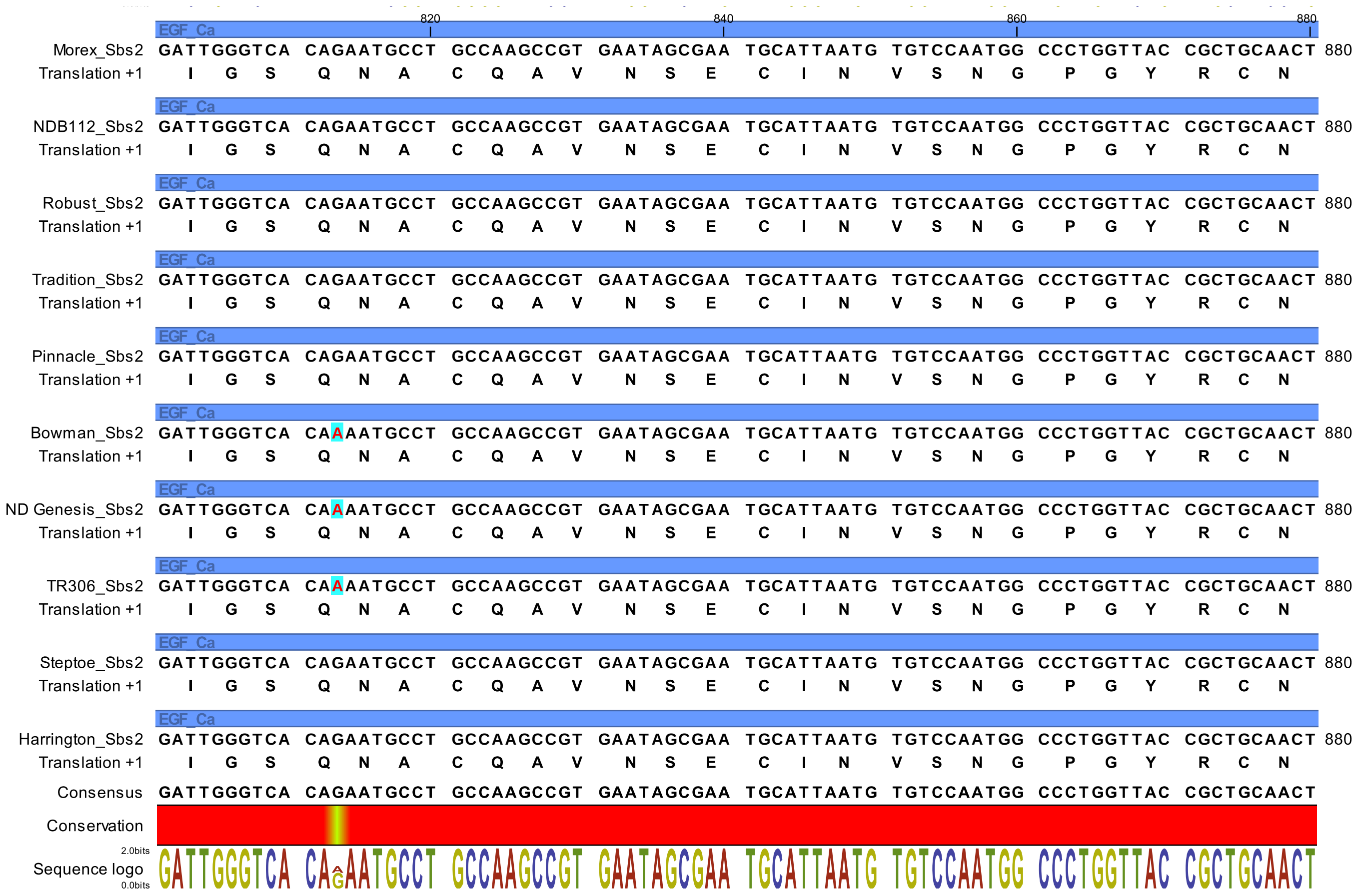

### Supplementary Figure 6B

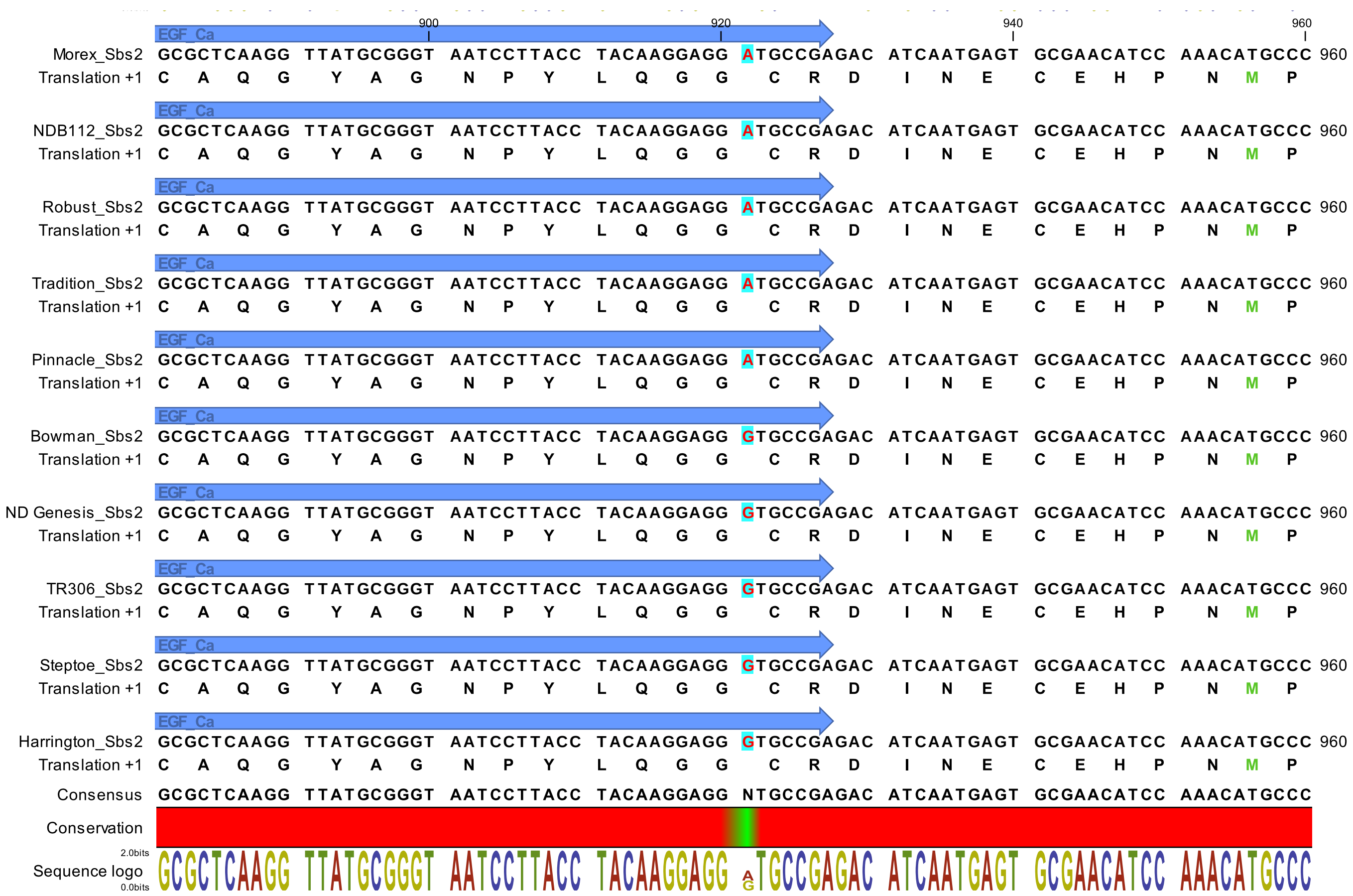

### Supplementary Figure 6B

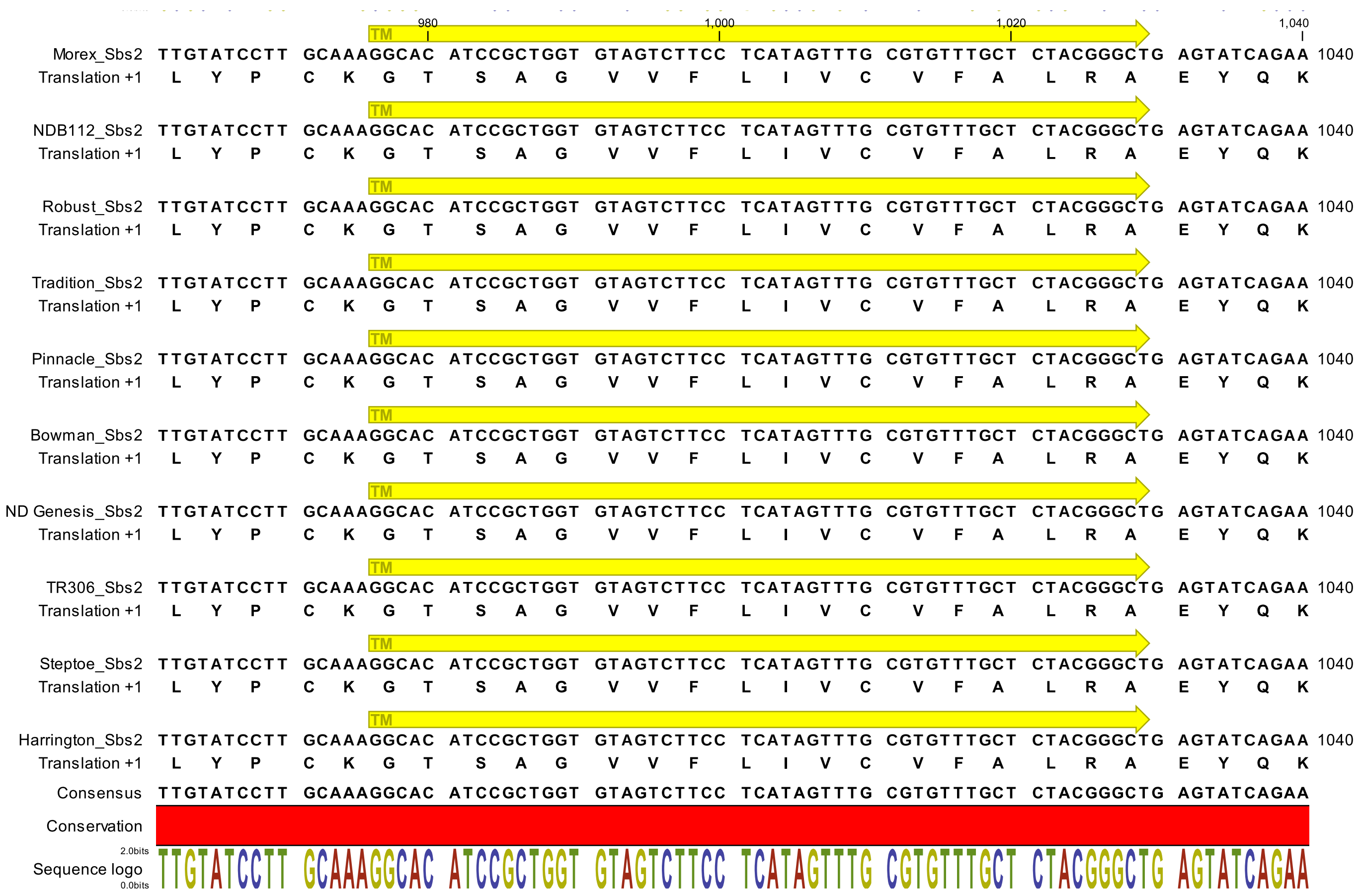

Supplementary Figure 6B

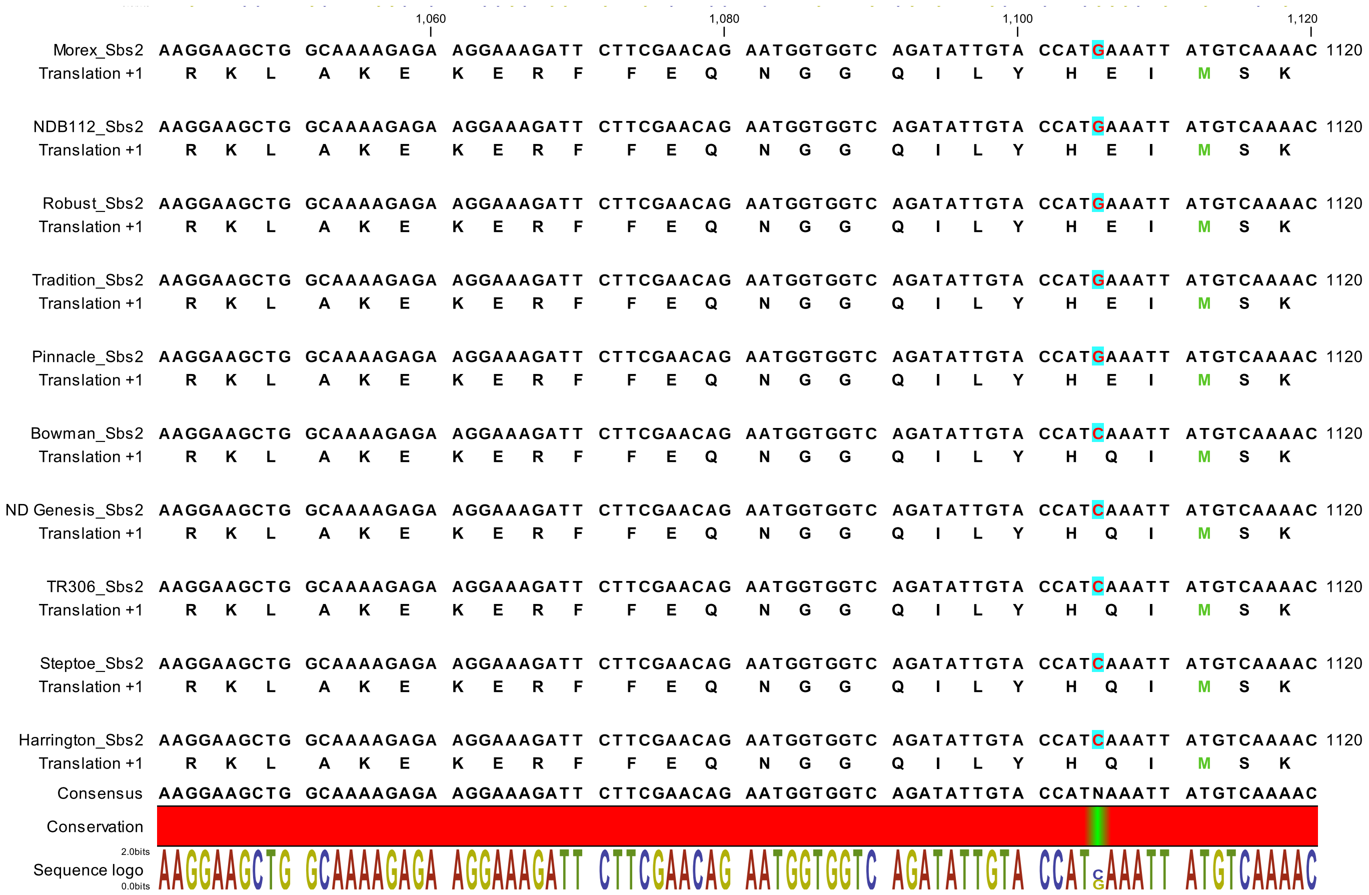

### Supplementary Figure 6B

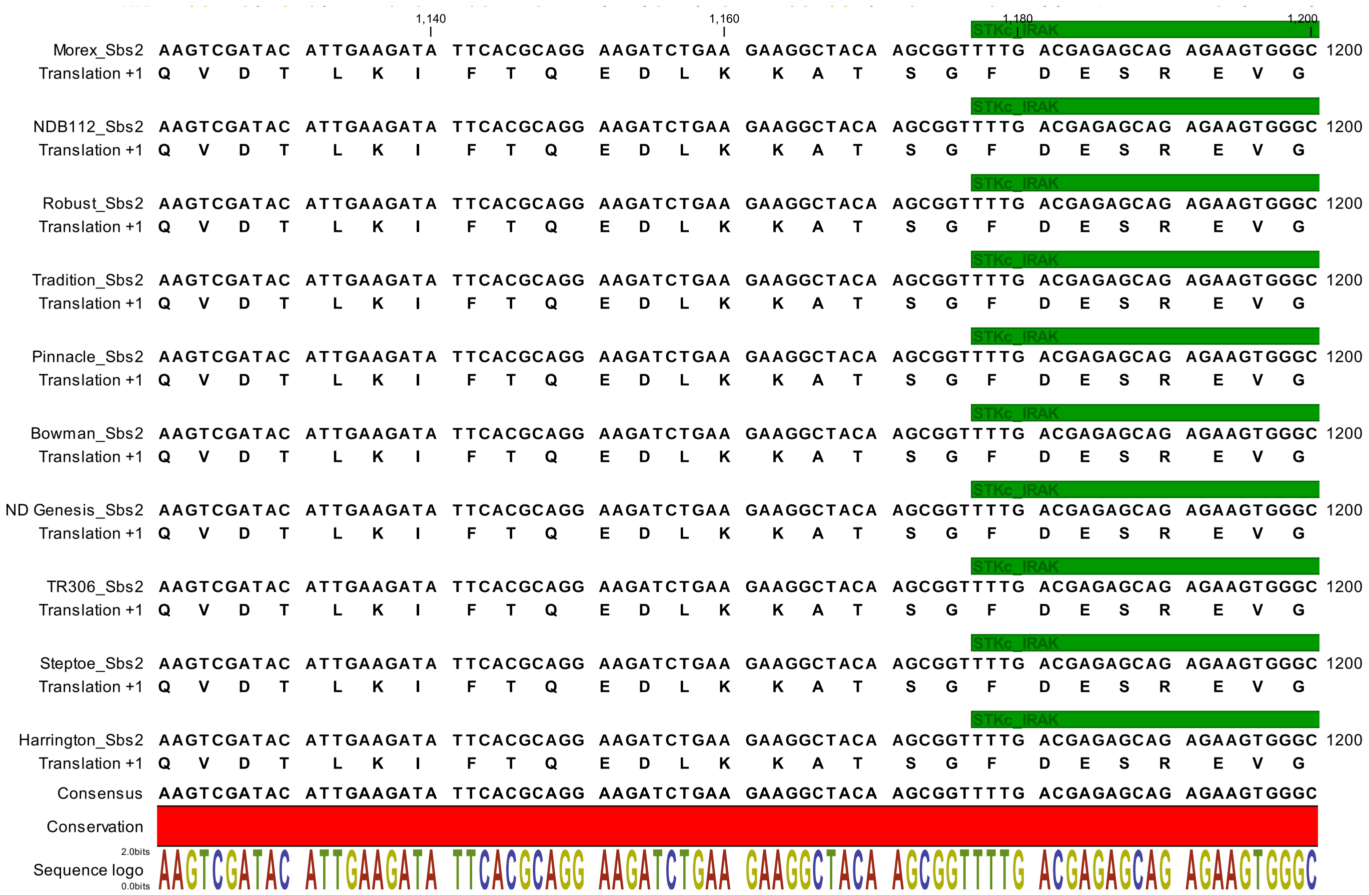

### Supplementary Figure 6B

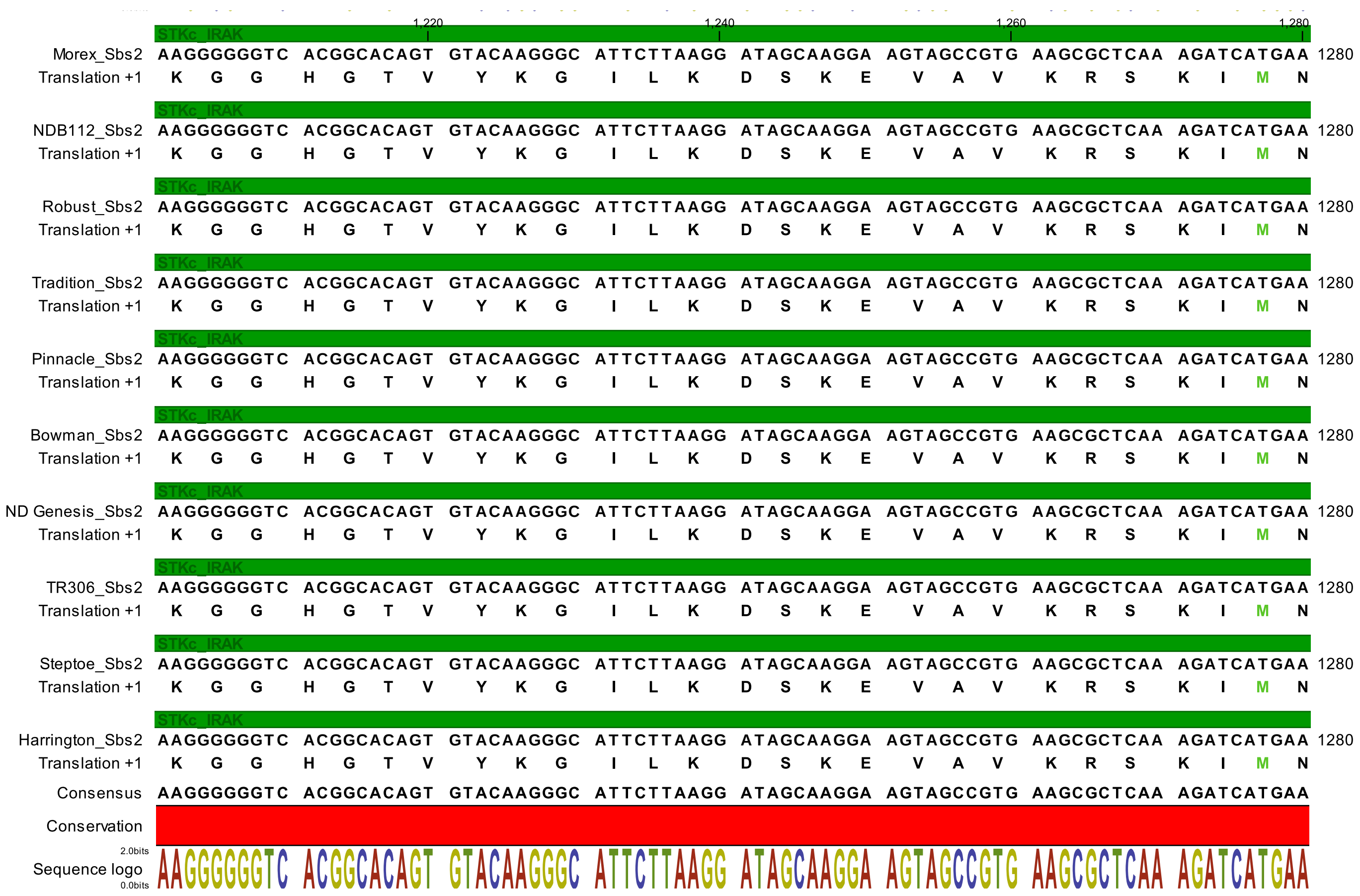

### Supplementary Figure 6B

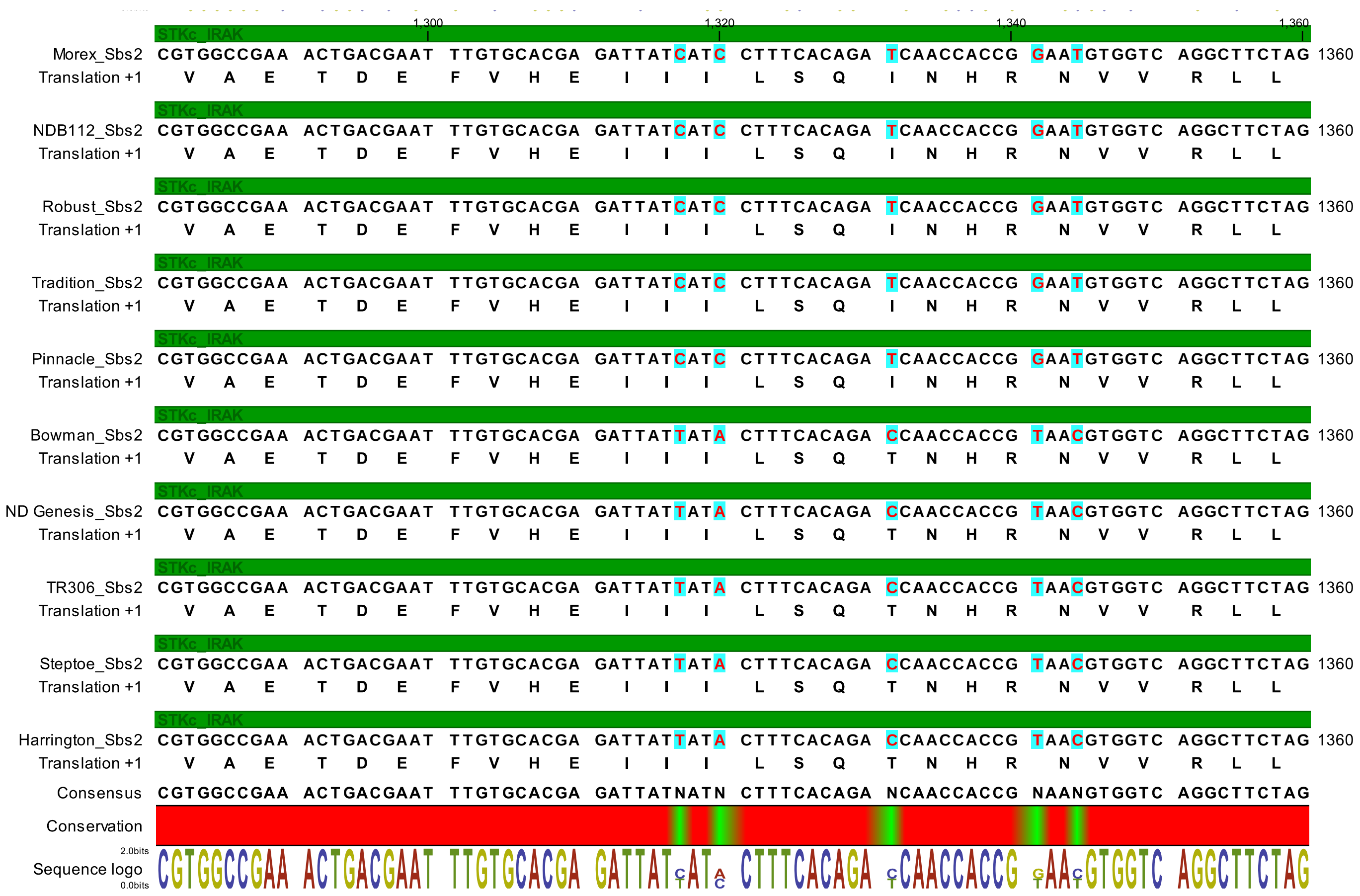

### Supplementary Figure 6B

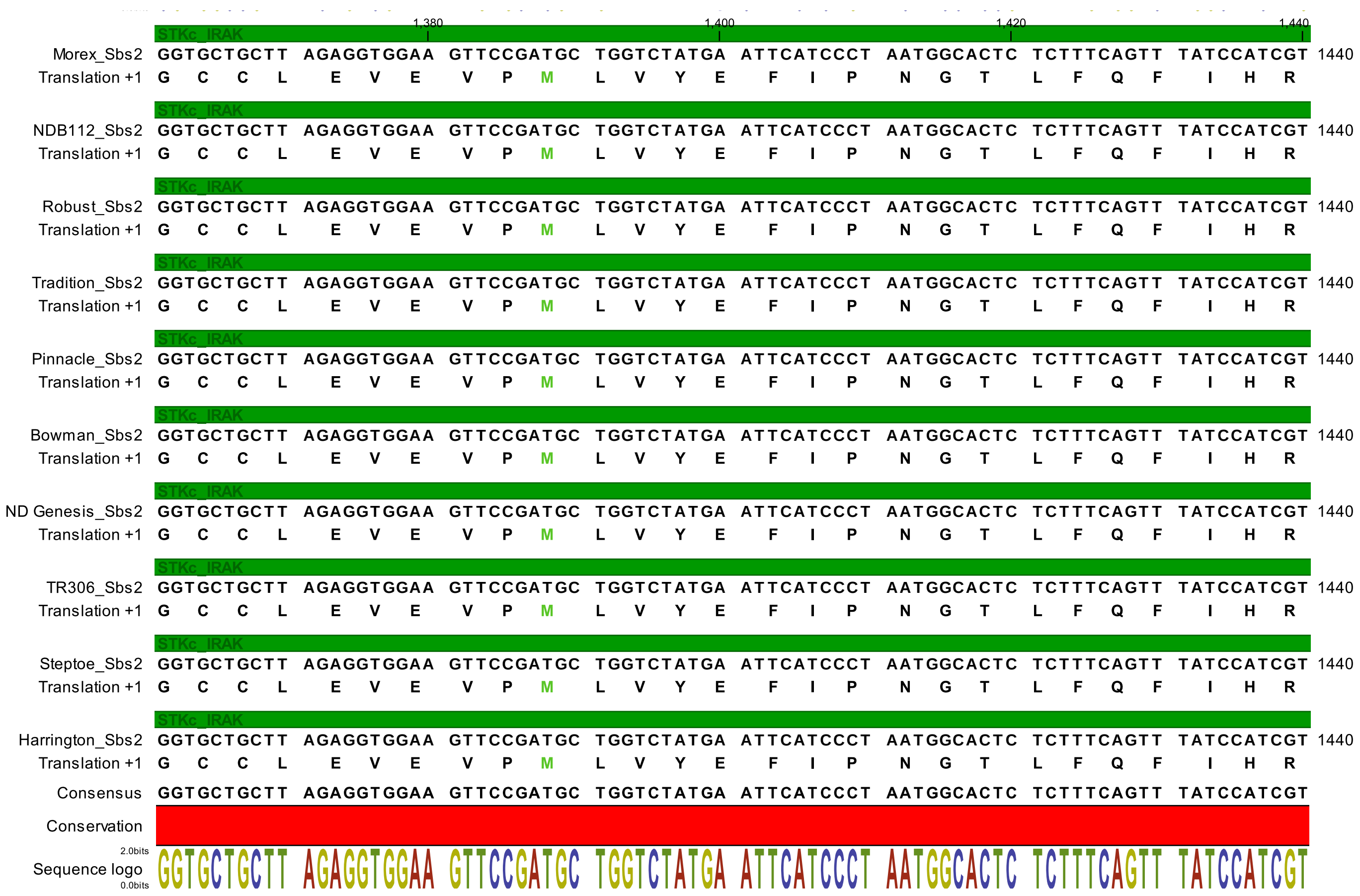

### Supplementary Figure 6B

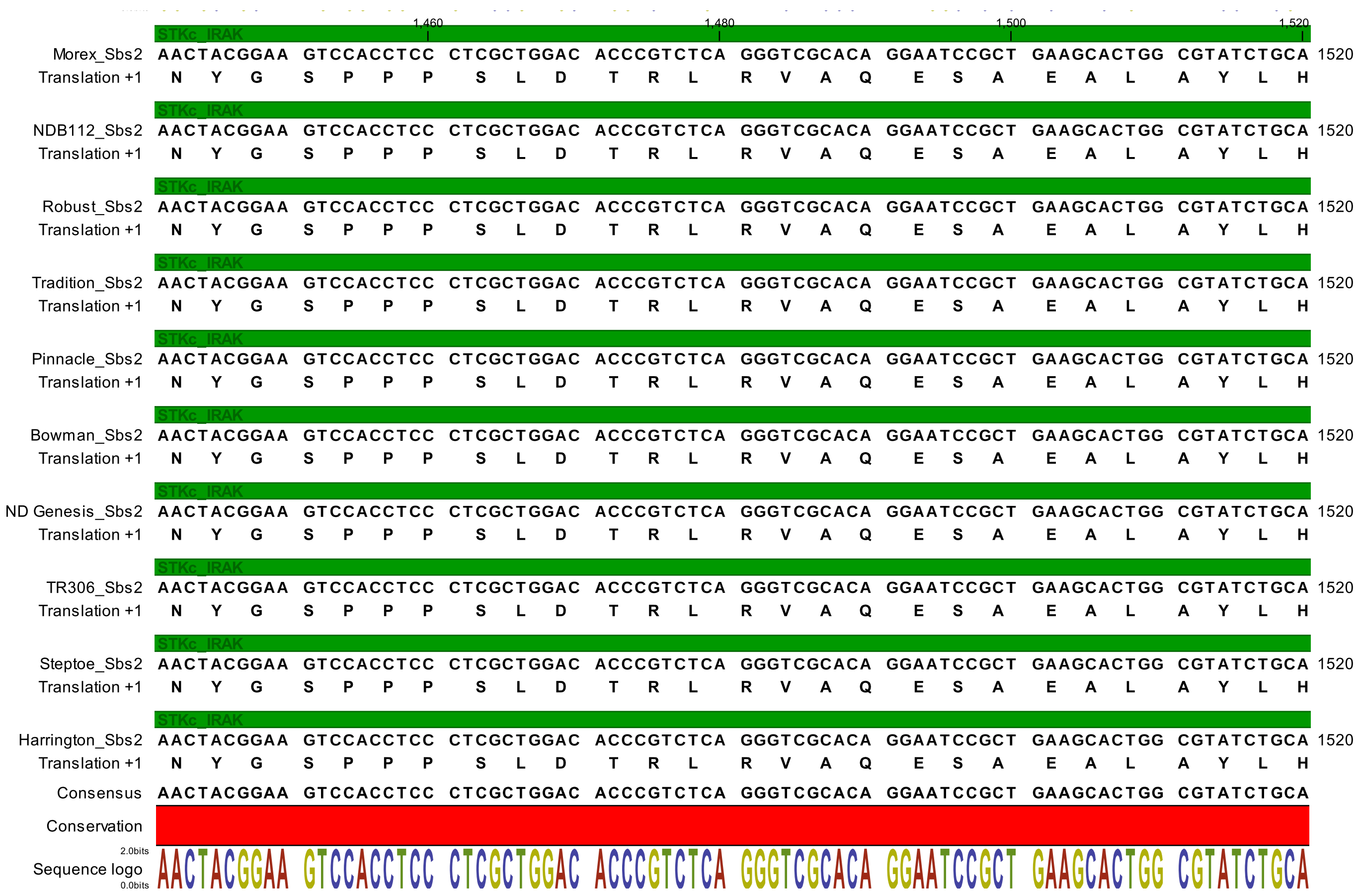

### Supplementary Figure 6B

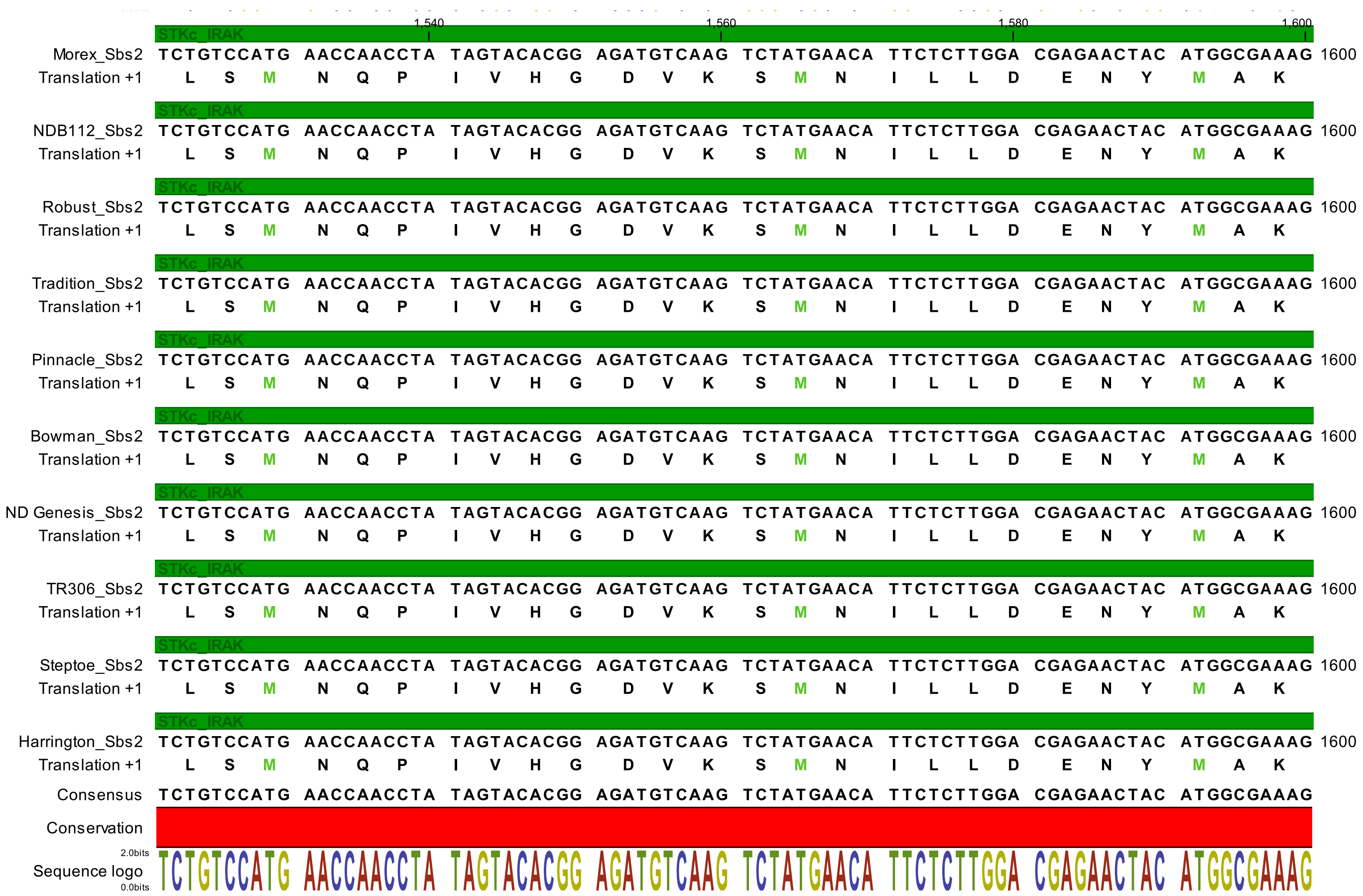

### Supplementary Figure 6B

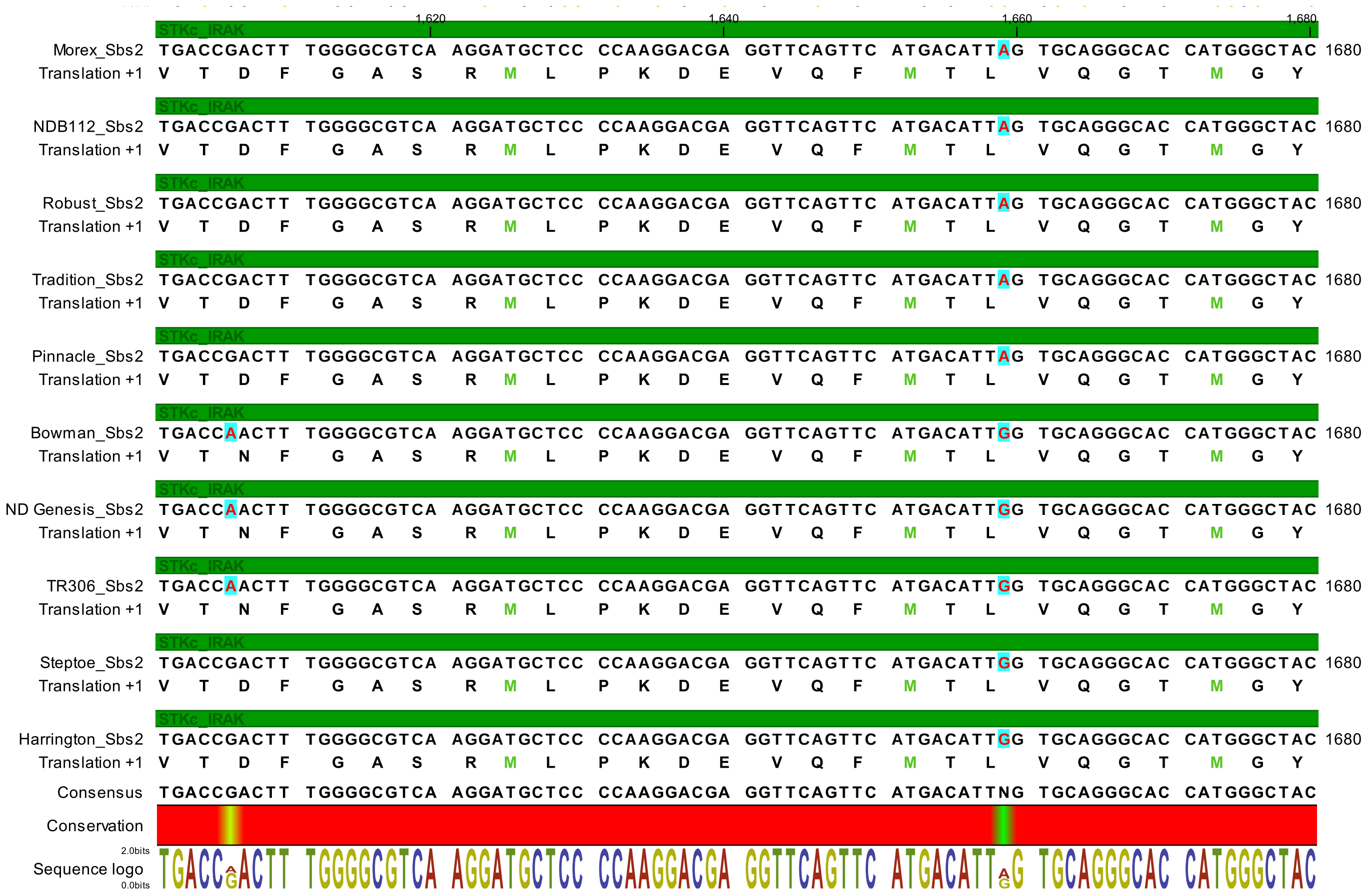

### Supplementary Figure 6B

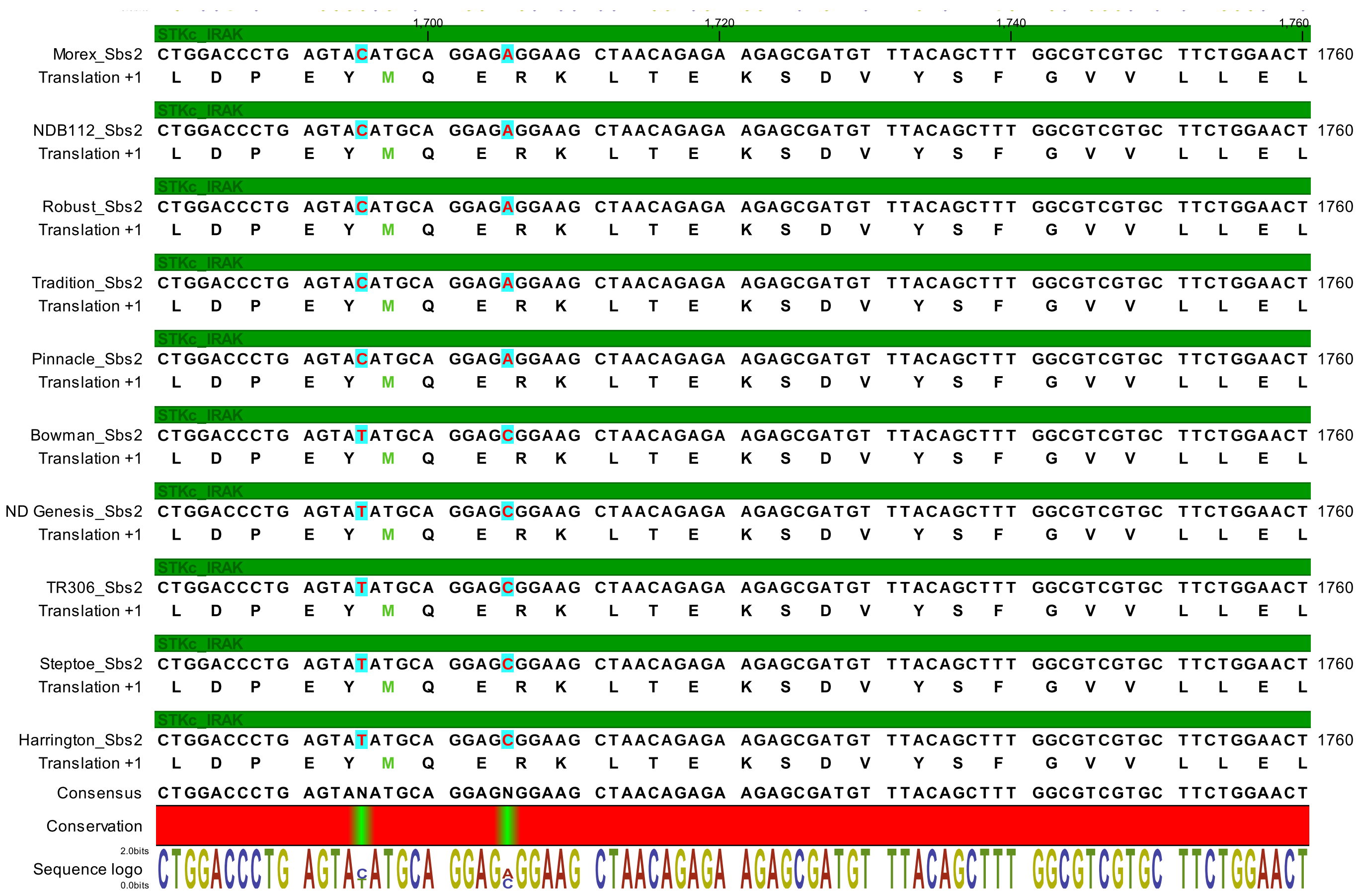

### Supplementary Figure 6B

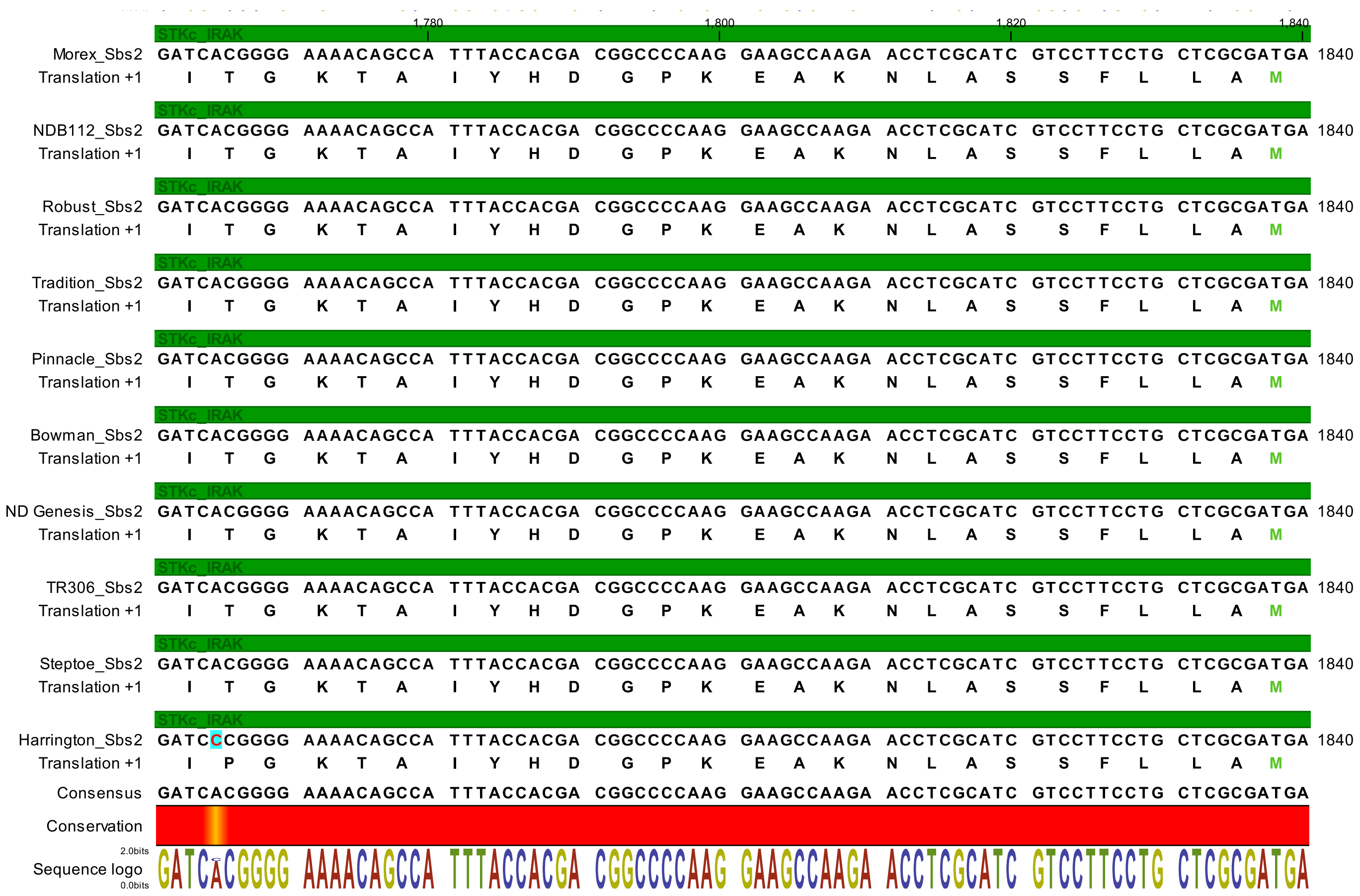

### Supplementary Figure 6B

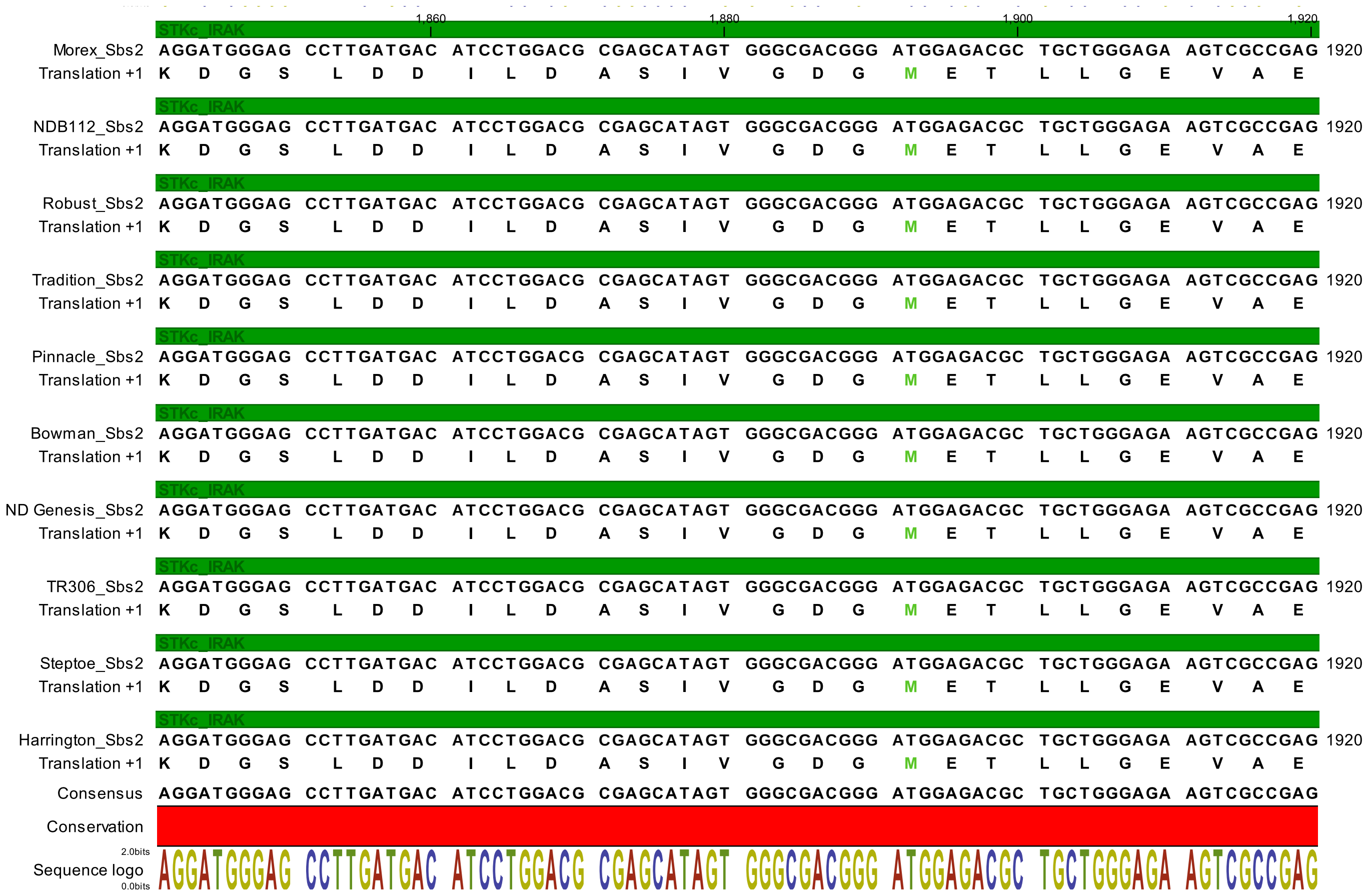

### Supplementary Figure 6B

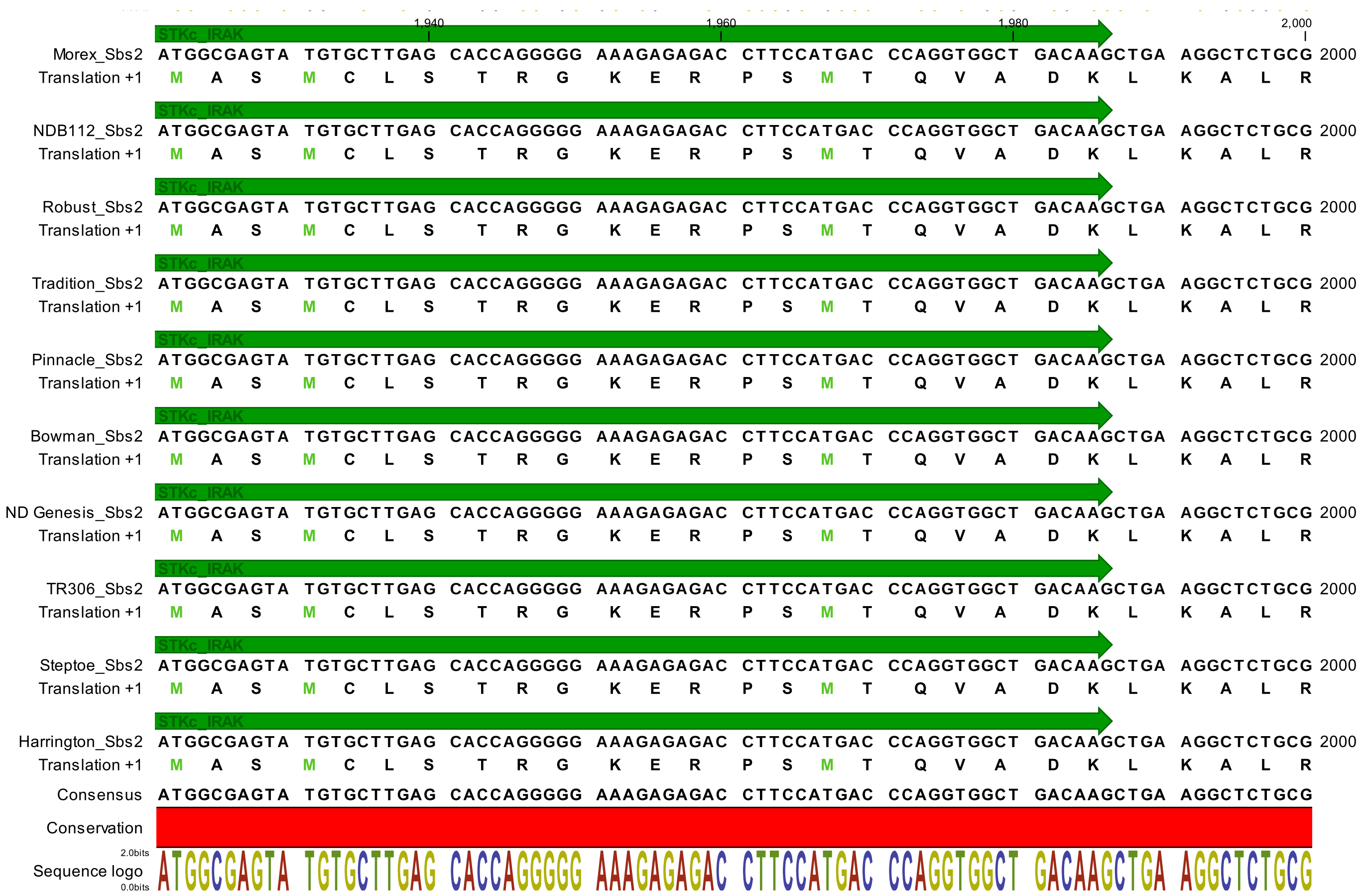

### Supplementary Figure 6B

| | | | | | | | | | |
|---|---|---|---|---|---|---|---|---|---|
| | | | 2,020 | | 2,040 | | 2,060 | | 2,080 |
| Morex_Sbs2 | AAGCACGTGG | ATGGAGAAAC | TGGTGCTCGA | GCATGGGAAA | ATAGATCGTC | TGGTCGCTGG | CTCCTCGCCT | GCAGATTTAA | 2080 |
| Translation +1 | S T W | M E K | L V L E | H G K | I D R | L V A G | S S P | A D L | |
| NDB112_Sbs2 | AAGCACGTGG | ATGGAGAAAC | TGGTGCTCGA | GCATGGGAAA | ATAGATCGTC | TGGTCGCTGG | CTCCTCGCCT | GCAGATTTAA | 2080 |
| Translation +1 | S T W | M E K | L V L E | H G K | I D R | L V A G | S S P | A D L | |
| Robust_Sbs2 | AAGCACGTGG | ATGGAGAAAC | TGGTGCTCGA | GCATGGGAAA | ATAGATCGTC | TGGTCGCTGG | CTCCTCGCCT | GCAGATTTAA | 2080 |
| Translation +1 | S T W | M E K | L V L E | H G K | I D R | L V A G | S S P | A D L | |
| Tradition_Sbs2 | AAGCACGTGG | ATGGAGAAAC | TGGTGCTCGA | GCATGGGAAA | ATAGATCGTC | TGGTCGCTGG | CTCCTCGCCT | GCAGATTTAA | 2080 |
| Translation +1 | S T W | M E K | L V L E | H G K | I D R | L V A G | S S P | A D L | |
| Pinnacle_Sbs2 | AAGCACGTGG | ATGGAGAAAC | TGGTGCTCGA | GCATGGGAAA | ATAGATCGTC | TGGTCGCTGG | CTCCTCGCCT | GCAGATTTAA | 2080 |
| Translation +1 | S T W | M E K | L V L E | H G K | I D R | L V A G | S S P | A D L | |
| Bowman_Sbs2 | AAGCACGTGG | ATGGAGAAAC | TGGTGCTCGA | GCATGGGAAA | ATAGATCGTC | TGGTCGCTGG | CTCCTCGCCT | GCAGATTTAA | 2080 |
| Translation +1 | S T W | M E K | L V L E | H G K | I D R | L V A G | S S P | A D L | |
| ND Genesis_Sbs2 | AAGCACGTGG | ATGGAGAAAC | TGGTGCTCGA | GCATGGGAAA | ATAGATCGTC | TGGTCGCTGG | CTCCTCGCCT | GCAGATTTAA | 2080 |
| Translation +1 | S T W | M E K | L V L E | H G K | I D R | L V A G | S S P | A D L | |
| TR306_Sbs2 | AAGCACGTGG | ATGGAGAAAC | TGGTGCTCGA | GCATGGGAAA | ATAGATCGTC | TGGTCGCTGG | CTCCTCGCCT | GCAGATTTAA | 2080 |
| Translation +1 | S T W | M E K | L V L E | H G K | I D R | L V A G | S S P | A D L | |
| Steptoe_Sbs2 | AAGCACGTGG | ATGGAGAAAC | TGGTGCTCGA | GCATGGGAAA | ATAGATCGTC | TGGTCGCTGG | CTCCTCGCCT | GCAGATTTAA | 2080 |
| Translation +1 | S T W | M E K | L V L E | H G K | I D R | L V A G | S S P | A D L | |
| Harrington_Sbs2 | AAGCACGTGG | ATGGAGAAAC | TGGTGCTCGA | GCATGGGAAA | ATAGATCGTC | TGGTCGCTGG | CTCCTCGCCT | GCAGATTTAA | 2080 |
| Translation +1 | S T W | M E K | L V L E | H G K | I D R | L V A G | S S P | A D L | |
| Consensus | AAGCACGTGG | ATGGAGAAAC | TGGTGCTCGA | GCATGGGAAA | ATAGATCGTC | TGGTCGCTGG | CTCCTCGCCT | GCAGATTTAA | |

Conservation

Sequence logo

2.0bits

0.0bits

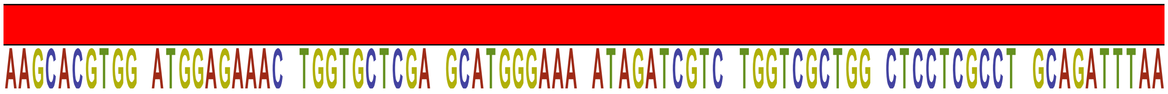

### Supplementary Figure 6B

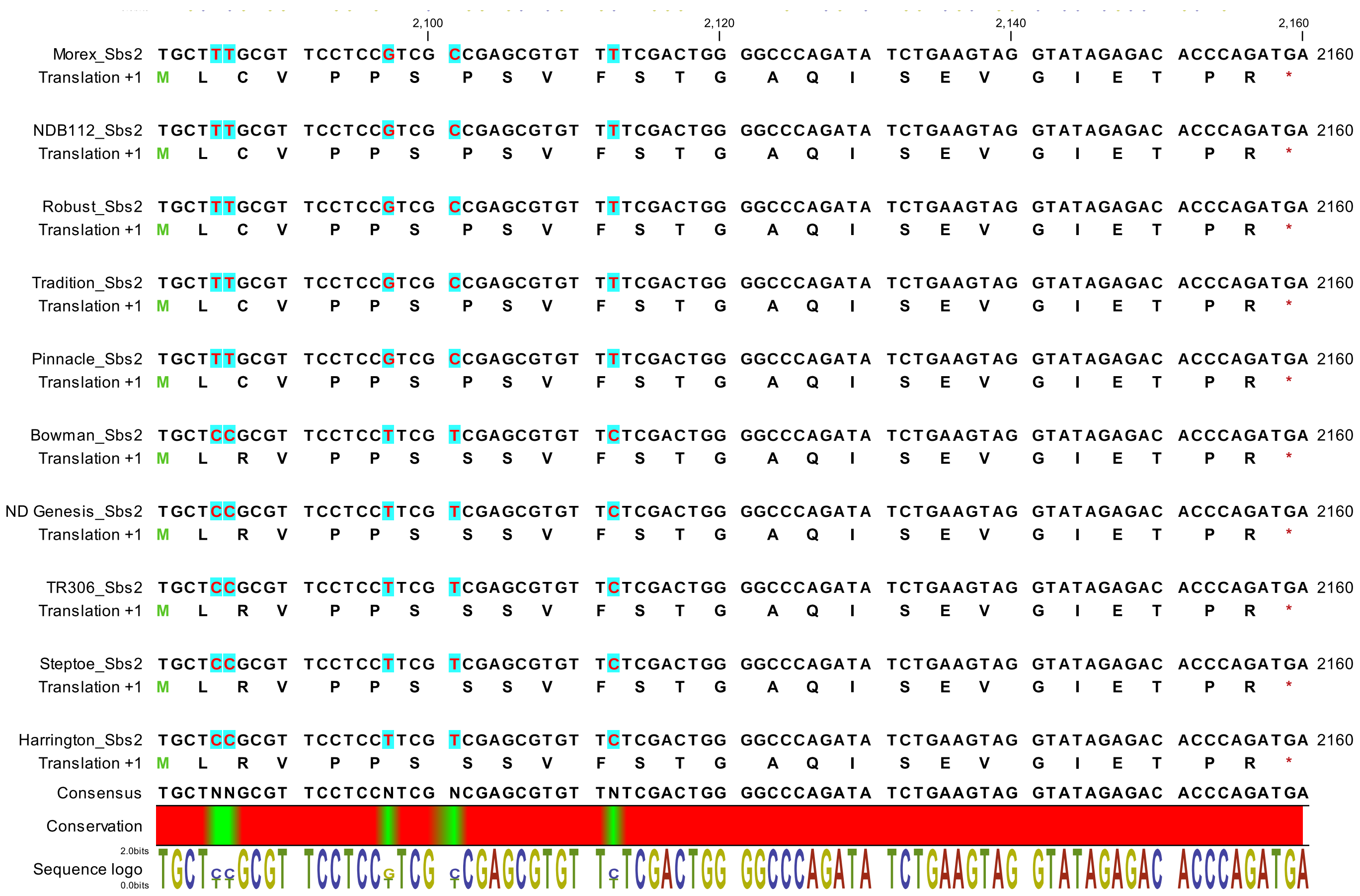

Supplementary Figure 6C

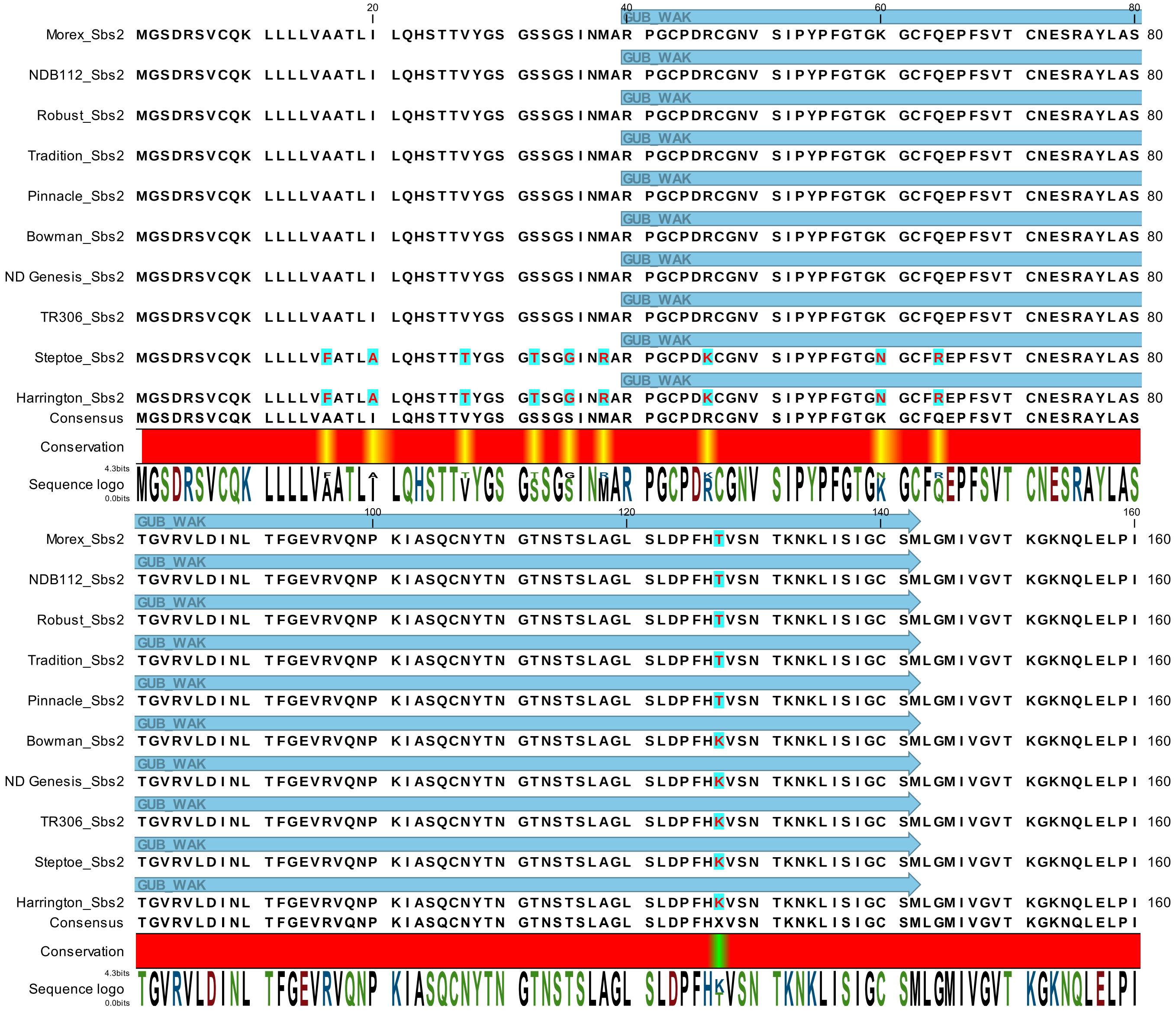

### Supplementary Figure 6C

```
                                              180         200         220         240
Morex_Sbs2       VNSCFSYCTD ASSVDDSTEC AGMGCCQTPL PGNISSFNTT STPVPTVYNA TIQSFSPCSY SFIAEVDSFK FDRSYASSTN 240
NDB112_Sbs2      VNSCFSYCTD ASSVDDSTEC AGMGCCQTPL PGNISSFNTT STPVPTVYNA TIQSFSPCSY SFIAEVDSFK FDRSYASSTN 240
Robust_Sbs2      VNSCFSYCTD ASSVDDSTEC AGMGCCQTPL PGNISSFNTT STPVPTVYNA TIQSFSPCSY SFIAEVDSFK FDRSYASSTN 240
Tradition_Sbs2   VNSCFSYCTD ASSVDDSTEC AGMGCCQTPL PGNISSFNTT STPVPTVYNA TIQSFSPCSY SFIAEVDSFK FDRSYASSTN 240
Pinnacle_Sbs2    VNSCFSYCTD ASSVDDSTEC AGMGCCQTPL PGNISSFNTT STPVPTVYNA TIQSFSPCSY SFIAEVDSFK FDRSYASSTN 240
Bowman_Sbs2      VNSCFSFCTD ASSVDDSTEC AGMGCCQTPL PGNISSFNTT STPVPTVYNA TIQSFSPCSY SFIAEVDSFK FDRSYASSTN 240
ND Genesis_Sbs2  VNSCFSFCTD ASSVDDSTEC AGMGCCQTPL PGNISSFNTT STPVPTVYNA TIQSFSPCSY SFIAEVDSFK FDRSYASSTN 240
TR306_Sbs2       VNSCFSFCTD ASSVDDSTEC AGMGCCQTPL PGNISSFNTT STPVPTVYNA TIQSFSPCSY SFIAEVDSFK FDRSYASSTN 240
Steptoe_Sbs2     VNSCFSFCTD ASSVDDSTEC AGMGCCQTPL PGNISSFNTT STPVPTVYNA TIQSFSPCSY SFIAEVESFK FDRSYASSTN 240
Harrington_Sbs2  VNSCFSFCTD ASSVDDSTEC AGMGCCQTPL PGNISSFNTT STPVPTVYNA TIQSFSPCSY SFIAEVESFK FDRSYASSTN 240
Consensus        VNSCFSXCTD ASSVDDSTEC AGMGCCQTPL PGNISSFNTT STPVPTVYNA TIQSFSPCSY SFIAEVDSFK FDRSYASSTN
Conservation
Sequence logo    VNSCFS[F/Y]CTD ASSVDDSTEC AGMGCCQTPL PGNISSFNTT STPVPTVYNA TIQSFSPCSY SFIAEV[E/D]SFK FDRSYASSTN

                                        260         280         300         320
                                        EGF_Ca (positions ~261-306)
Morex_Sbs2       FRTKYTEGFP LVLDWVVGNG SCSEATRIGS QNACQAVNSE CINVSNGPGY RCNCAQGYAG NPYLQGGCRD INECEHPNMP 320
NDB112_Sbs2      FRTKYTEGFP LVLDWVVGNG SCSEATRIGS QNACQAVNSE CINVSNGPGY RCNCAQGYAG NPYLQGGCRD INECEHPNMP 320
Robust_Sbs2      FRTKYTEGFP LVLDWVVGNG SCSEATRIGS QNACQAVNSE CINVSNGPGY RCNCAQGYAG NPYLQGGCRD INECEHPNMP 320
Tradition_Sbs2   FRTKYTEGFP LVLDWVVGNG SCSEATRIGS QNACQAVNSE CINVSNGPGY RCNCAQGYAG NPYLQGGCRD INECEHPNMP 320
Pinnacle_Sbs2    FRTKYTEGFP LVLDWVVGNG SCSEATRIGS QNACQAVNSE CINVSNGPGY RCNCAQGYAG NPYLQGGCRD INECEHPNMP 320
Bowman_Sbs2      FRTKYTEGFP LVLDWVVGNG SCSEATRIGS QNACQAVNSE CINVSNGPGY RCNCAQGYAG NPYLQGGCRD INECEHPNMP 320
ND Genesis_Sbs2  FRTKYTEGFP LVLDWVVGNG SCSEATRIGS QNACQAVNSE CINVSNGPGY RCNCAQGYAG NPYLQGGCRD INECEHPNMP 320
TR306_Sbs2       FRTKYTEGFP LVLDWVVGNG SCSEATRIGS QNACQAVNSE CINVSNGPGY RCNCAQGYAG NPYLQGGCRD INECEHPNMP 320
Steptoe_Sbs2     FRTKYTEGFP LVLDWVVGNG SCSEATRIGS QNACQAVNSE CINVSNGPGY RCNCAQGYAG NPYLQGGCRD INECEHPNMP 320
Harrington_Sbs2  FRTKYTEGFP LVLDWVVGNG SCSEATRIGS QNACQAVNSE CINVSNGPGY RCNCAQGYAG NPYLQGGCRD INECEHPNMP 320
Consensus        FRTKYTEGFP LVLDWVVGNG SCSEATRIGS QNACQAVNSE CINVSNGPGY RCNCAQGYAG NPYLQGGCRD INECEHPNMP
Conservation
Sequence logo    FRTKYTEGFP LVLDWVVGNG SCSEATRIGS QNACQAVNSE CINVSNGPGY RCNCAQGYAG NPYLQGGCRD INECEHPNMP
```

Sequence logo scale: 4.3bits – 0.0bits

### Supplementary Figure 6C

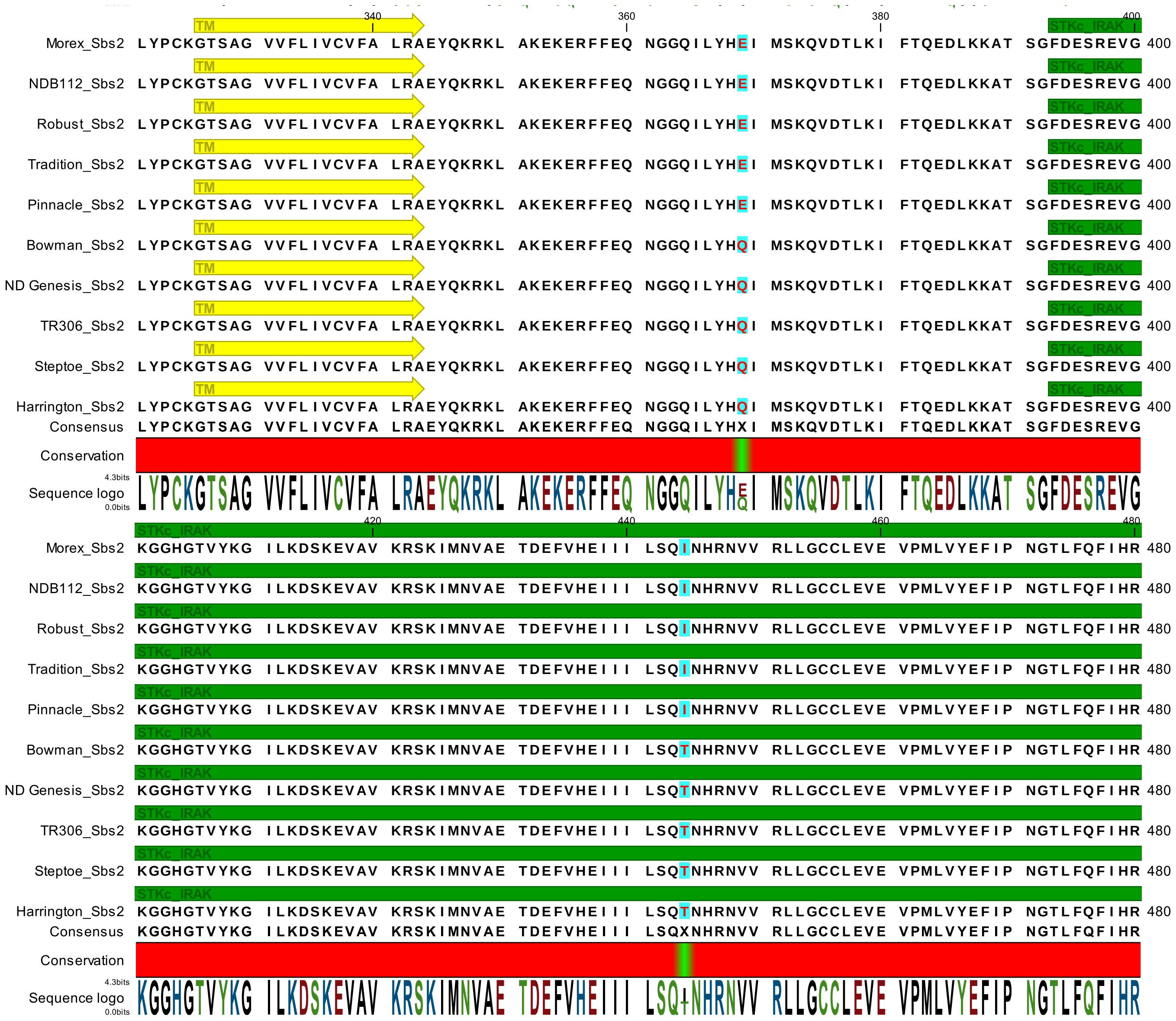

### Supplementary Figure 6C

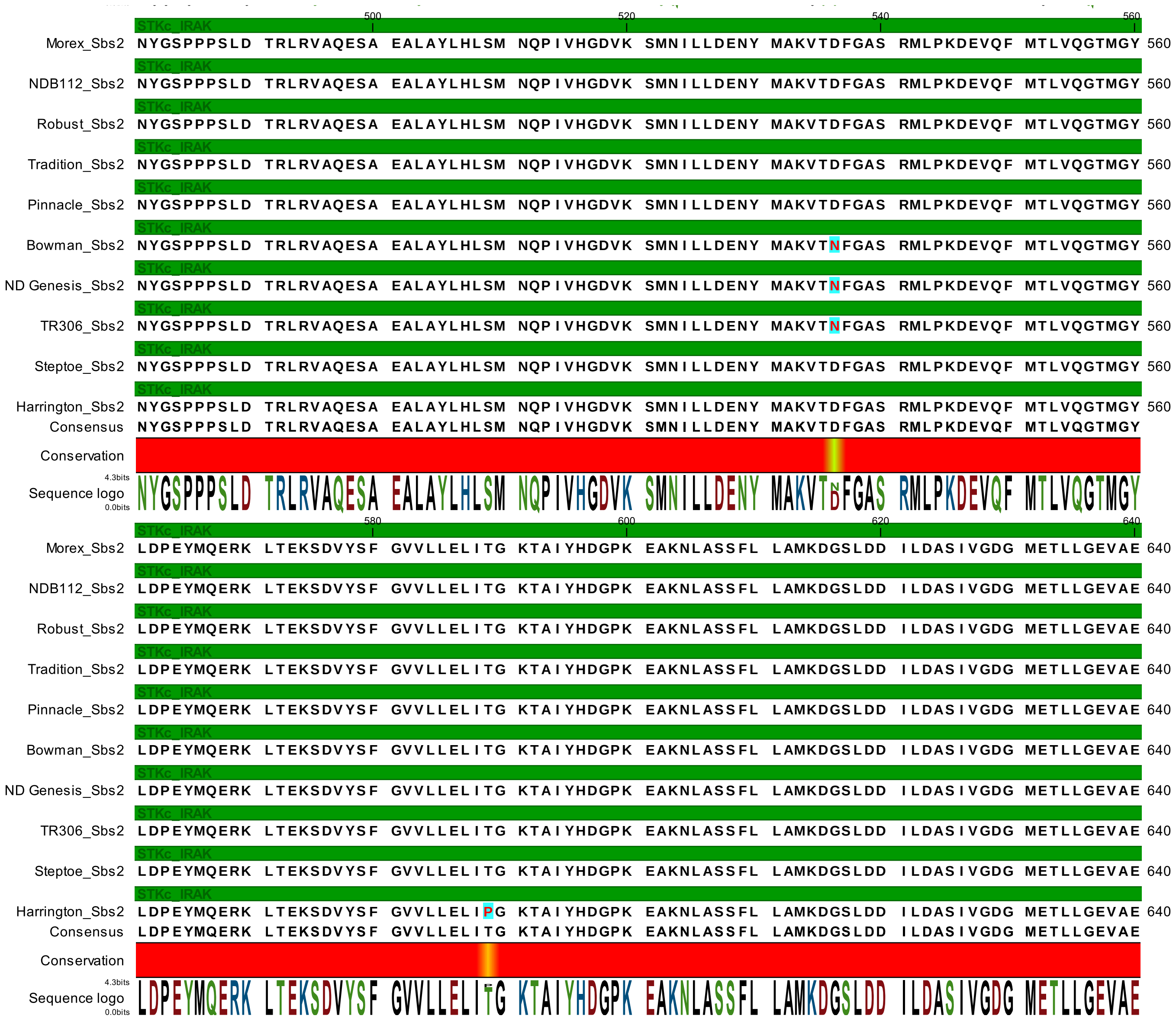

### Supplementary Figure 6C

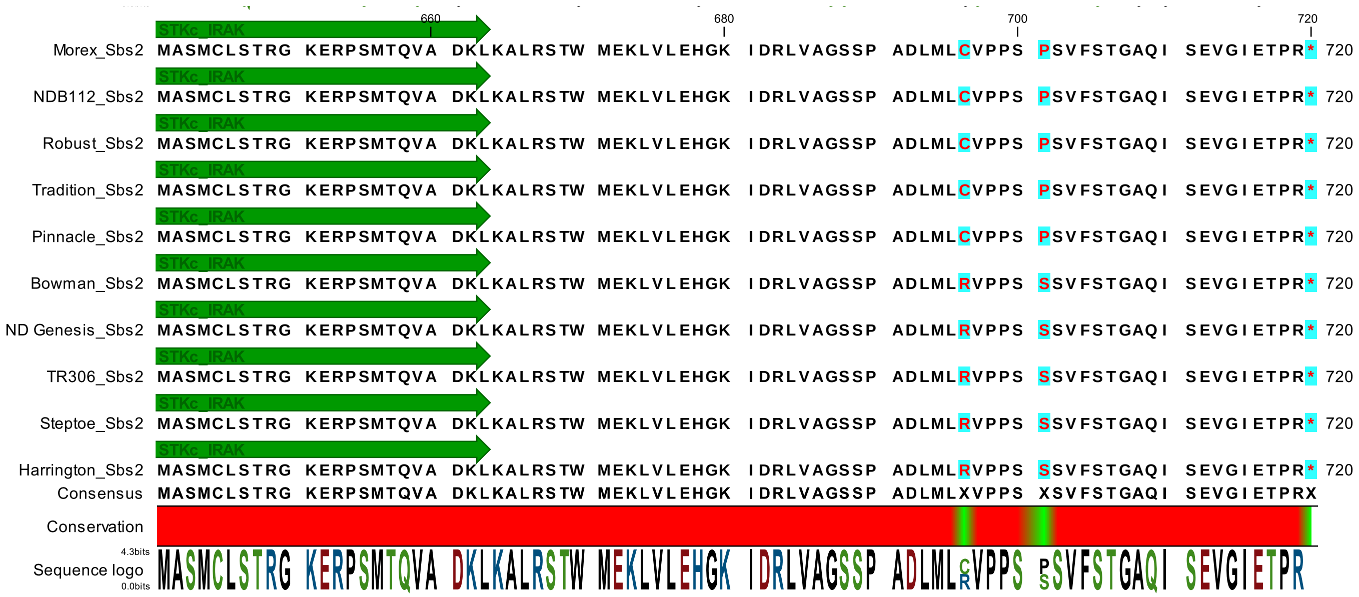

Supplementary Figure 6D.

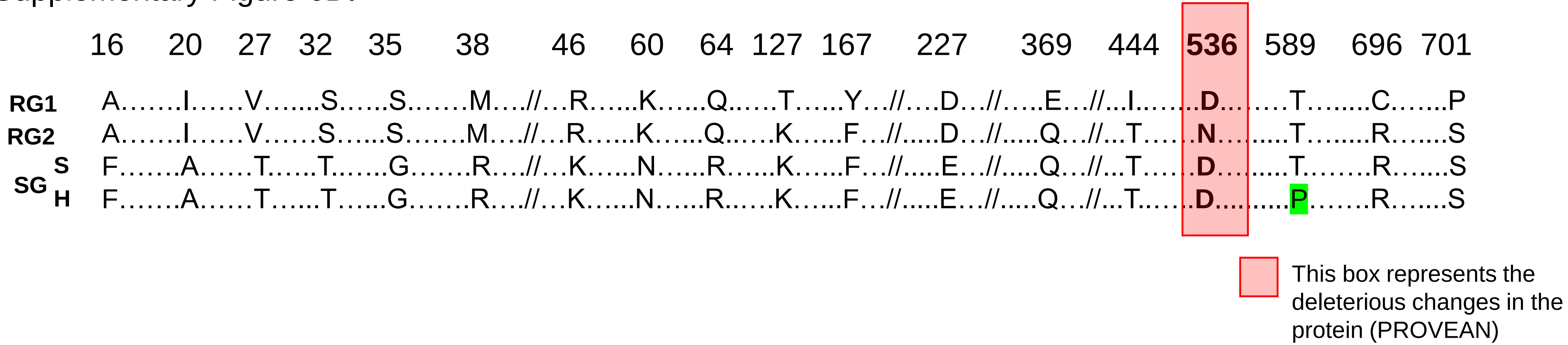


Gaps

| Differences | | | 1 | 2 | 3 | 4 | 5 | 6 | 7 | 8 | 9 | 10 |
|---|---|---|---|---|---|---|---|---|---|---|---|---|
| RG1 | Morex_Sbs2 | 1 | | 0 | 0 | 0 | 0 | 0 | 0 | 0 | 0 | 0 |
| RG1 | NDB112_Sbs2 | 2 | 0 | | 0 | 0 | 0 | 0 | 0 | 0 | 0 | 0 |
| RG1 | Robust_Sbs2 | 3 | 0 | 0 | | 0 | 0 | 0 | 0 | 0 | 0 | 0 |
| RG1 | Tradition_Sbs2 | 4 | 0 | 0 | 0 | | 0 | 0 | 0 | 0 | 0 | 0 |
| RG1 | Pinnacle_Sbs2 | 5 | 0 | 0 | 0 | 0 | | 0 | 0 | 0 | 0 | 0 |
| RG2 | Bowman_Sbs2 | 6 | 22 | 22 | 22 | 22 | 22 | | 0 | 0 | 0 | 0 |
| RG2 | ND Genesis_Sbs2 | 7 | 22 | 22 | 22 | 22 | 22 | 0 | | 0 | 0 | 0 |
| RG2 | TR306_Sbs2 | 8 | 22 | 22 | 22 | 22 | 22 | 0 | 0 | | 0 | 0 |
| SG | Steptoe_Sbs2 | 9 | 50 | 50 | 50 | 50 | 50 | 32 | 32 | 32 | | 0 |
| SG | Harrington_Sbs2 | 10 | 52 | 52 | 52 | 52 | 52 | 34 | 34 | 34 | 2 | |

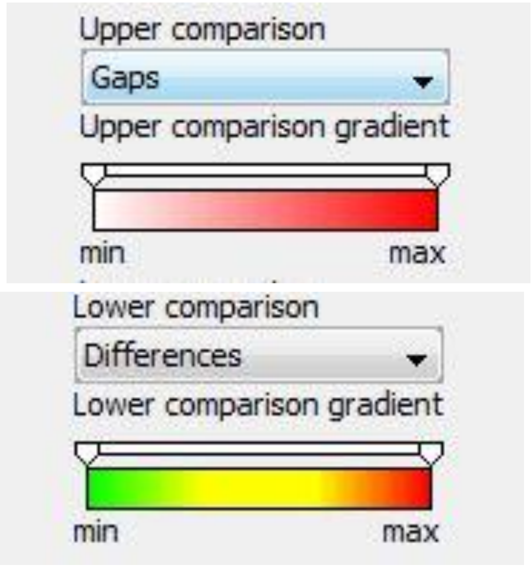

Supplementary Figure 7.

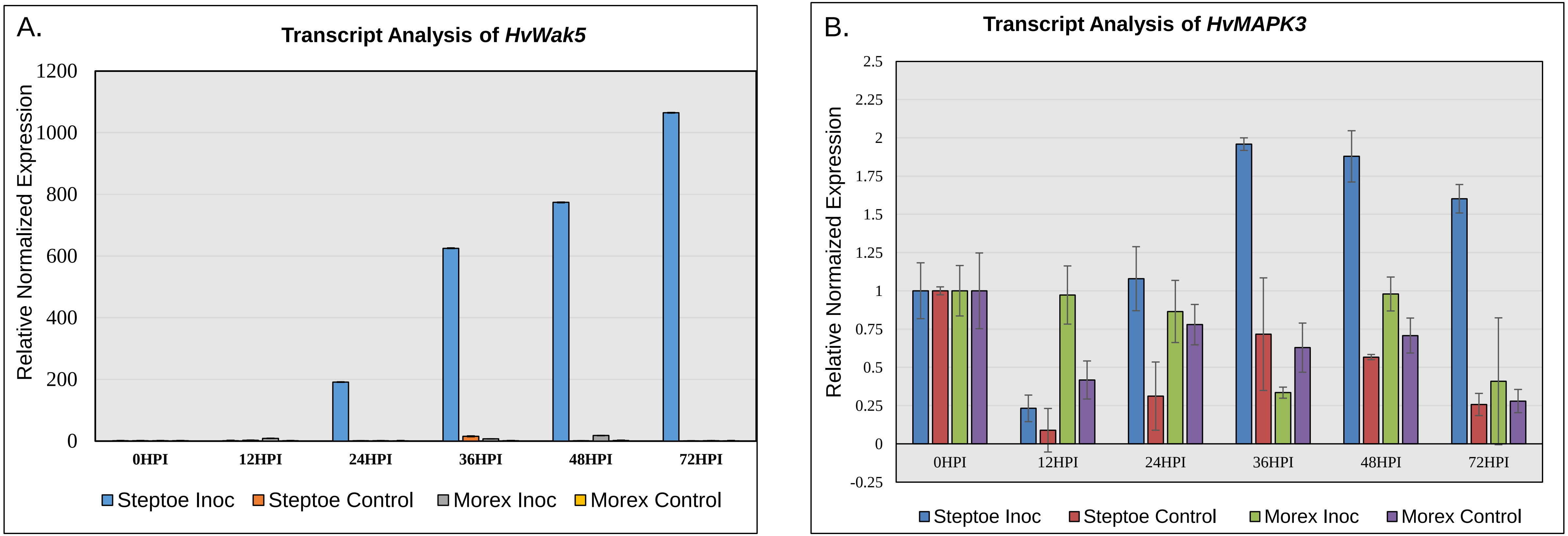

Supplementary Figure 8.

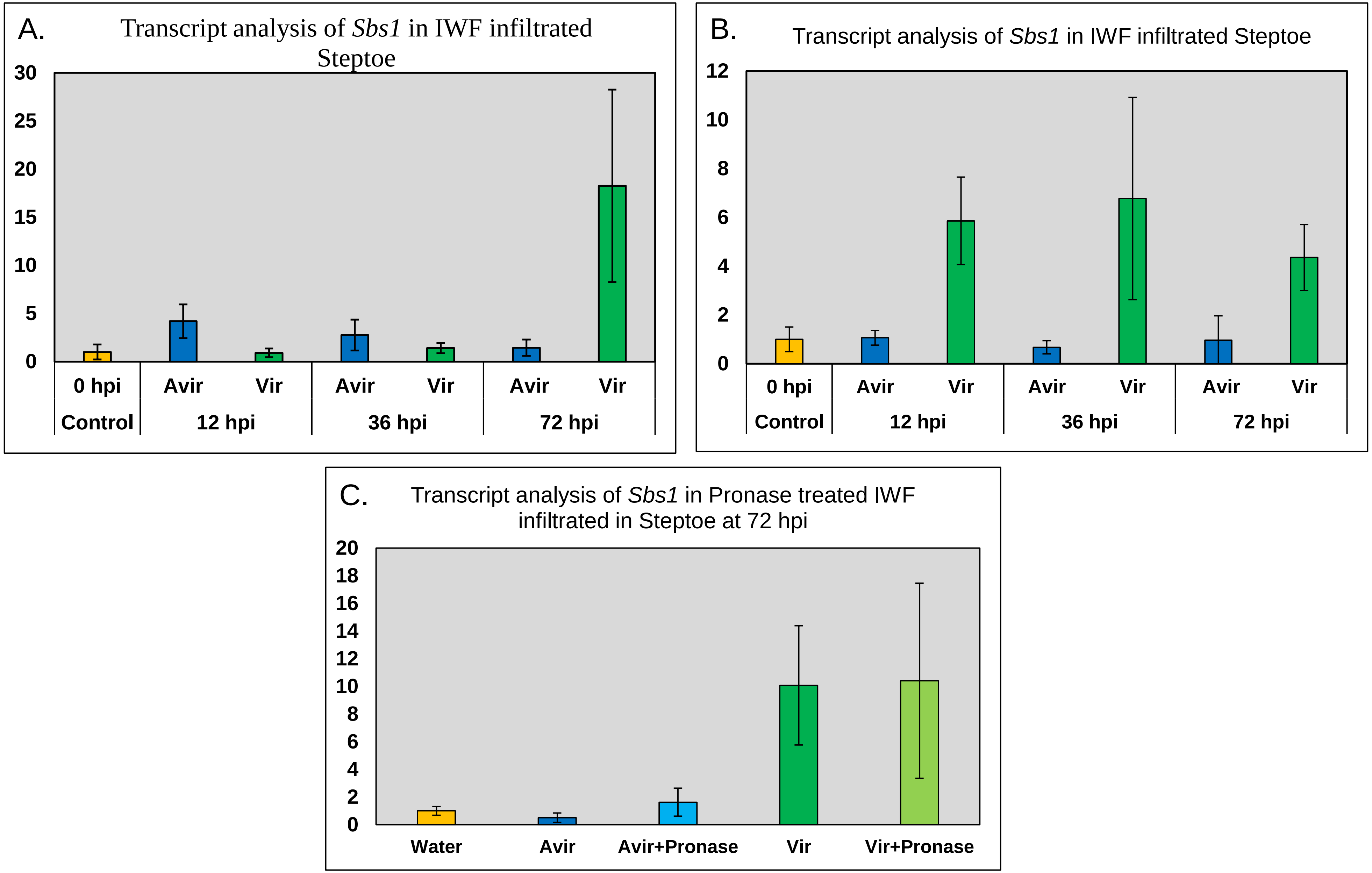

Supplementary Figure 9.

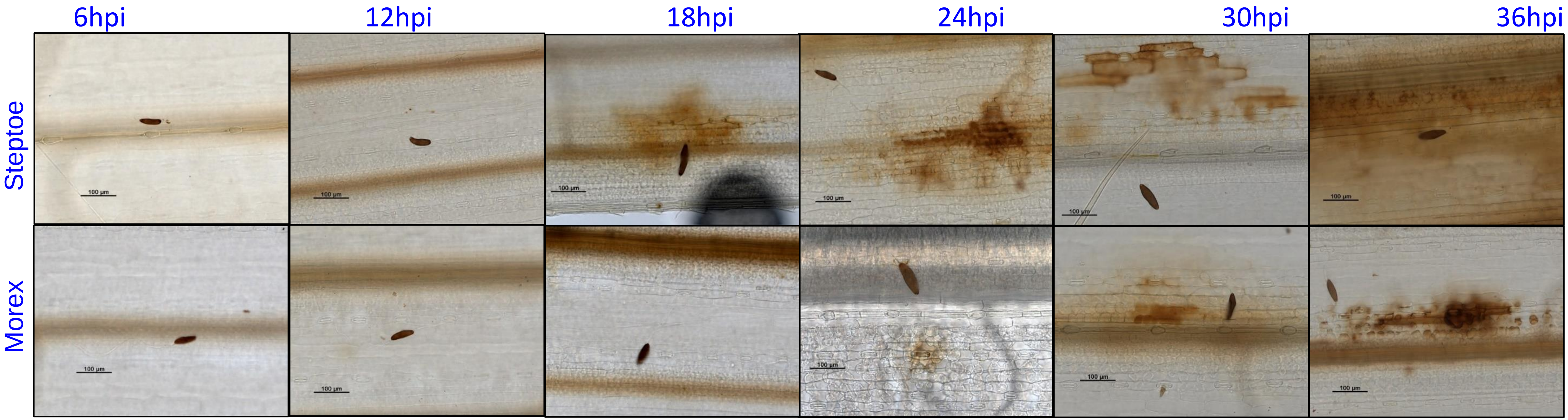

Supplementary Figure 10.

A.

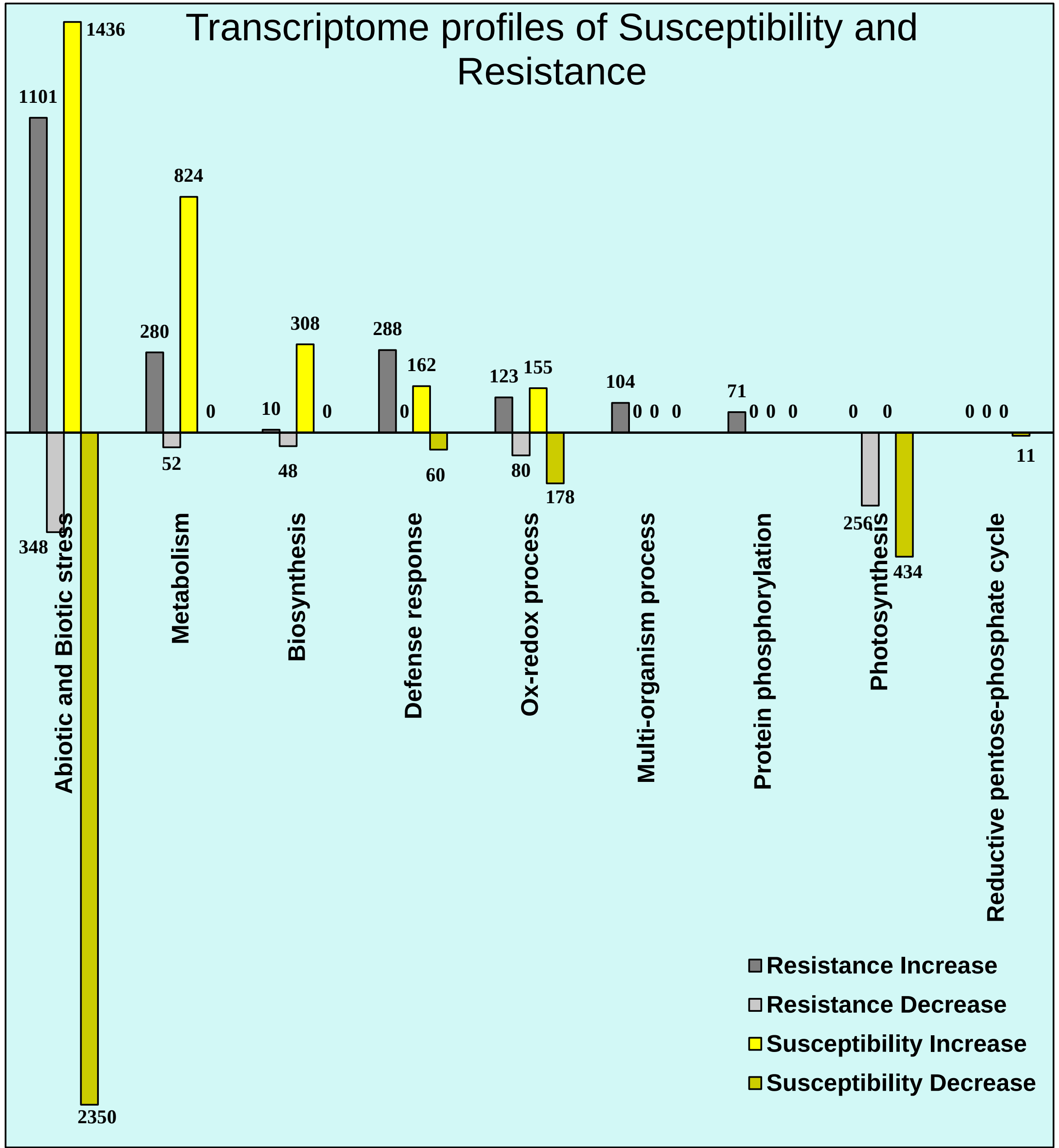


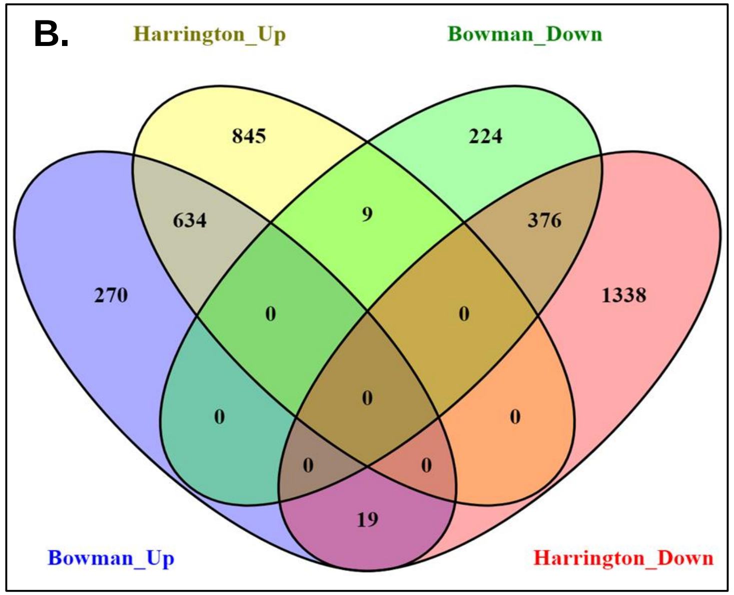


C.

| Genotype | Treatment | Replicate | Total Reads | Reads mapped to barley genome: Reads | Reads mapped to barley genome: Rate |
|---|---|---|---|---|---|
| Harrington | Non-Inoc | 1 | 56711792 | 47189882 | 83.21 |
| Harrington | Non-Inoc | 2 | 53235817 | 44968295 | 84.47 |
| Harrington | Non-Inoc | 3 | 41307906 | 35272821 | 85.39 |
| Harrington | 72 hpl | 1 | 52518290 | 41883336 | 79.75 |
| Harrington | 72 hpl | 2 | 54534806 | 42640765 | 78.19 |
| Harrington | 72 hpl | 3 | 63496548 | 49501909 | 77.96 |
| Bowman | Non-Inoc | 1 | 56508226 | 46726652 | 82.69 |
| Bowman | Non-Inoc | 2 | 52549749 | 44241634 | 84.19 |
| Bowman | Non-Inoc | 3 | 61817167 | 50881710 | 82.31 |
| Bowman | 72 hpl | 1 | 48707010 | 42034150 | 86.3 |
| Bowman | 72 hpl | 2 | 60002329 | 53720085 | 89.53 |
| Bowman | 72 hpl | 3 | 60830236 | 51255557 | 84.26 |

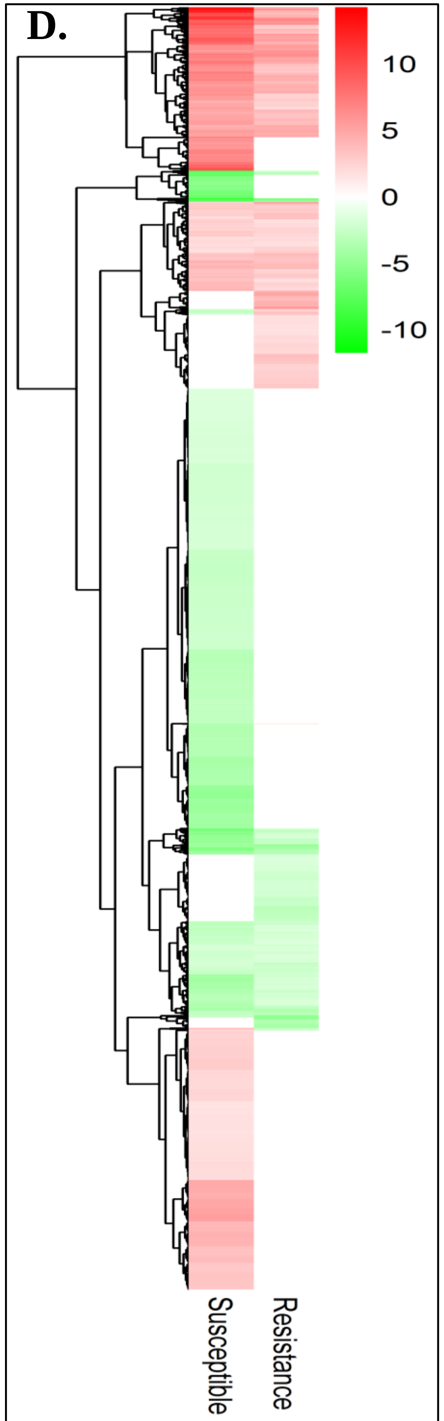

### Supplementary Figure 11A.

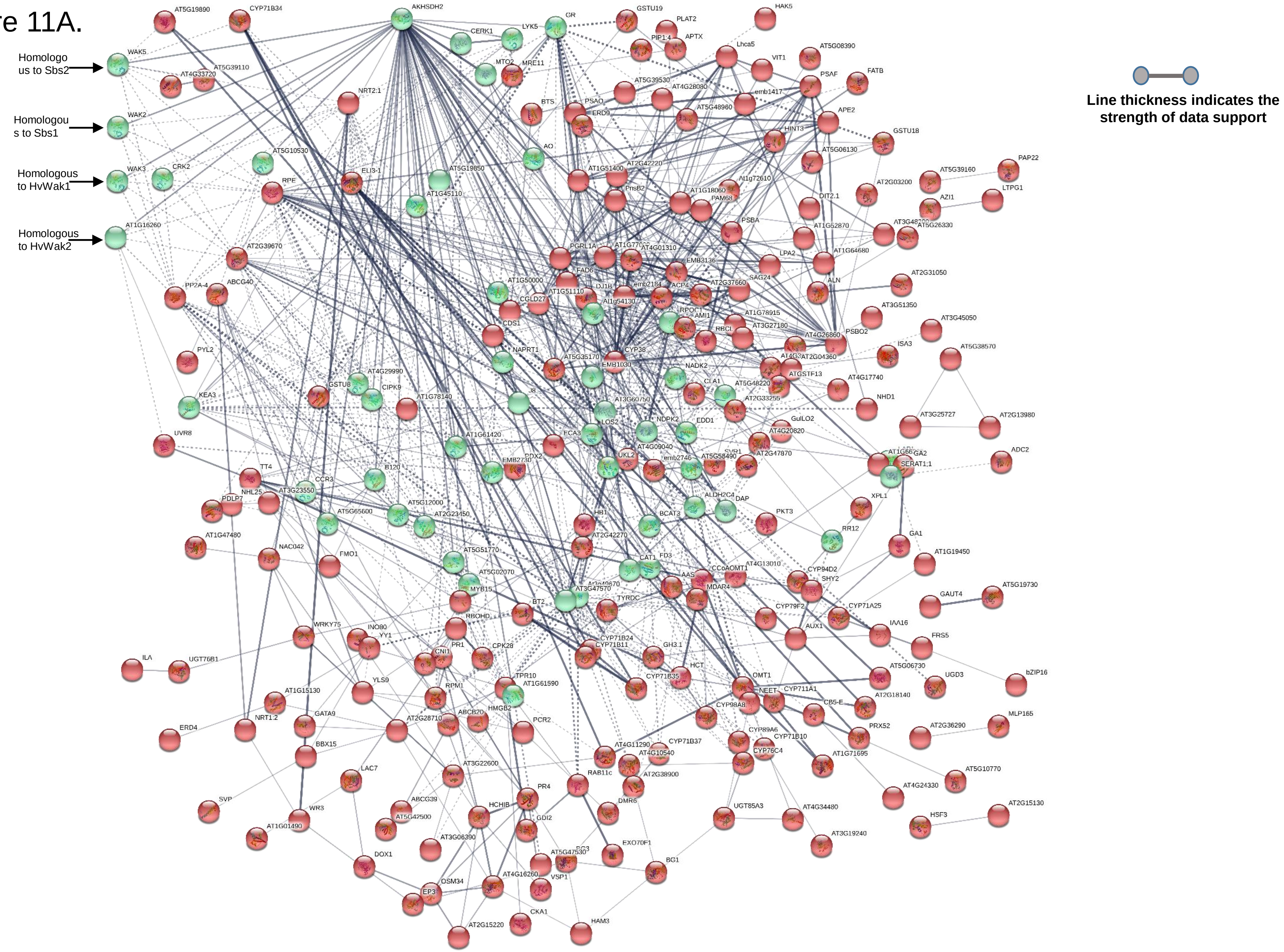

Supplementary Figure 11B.

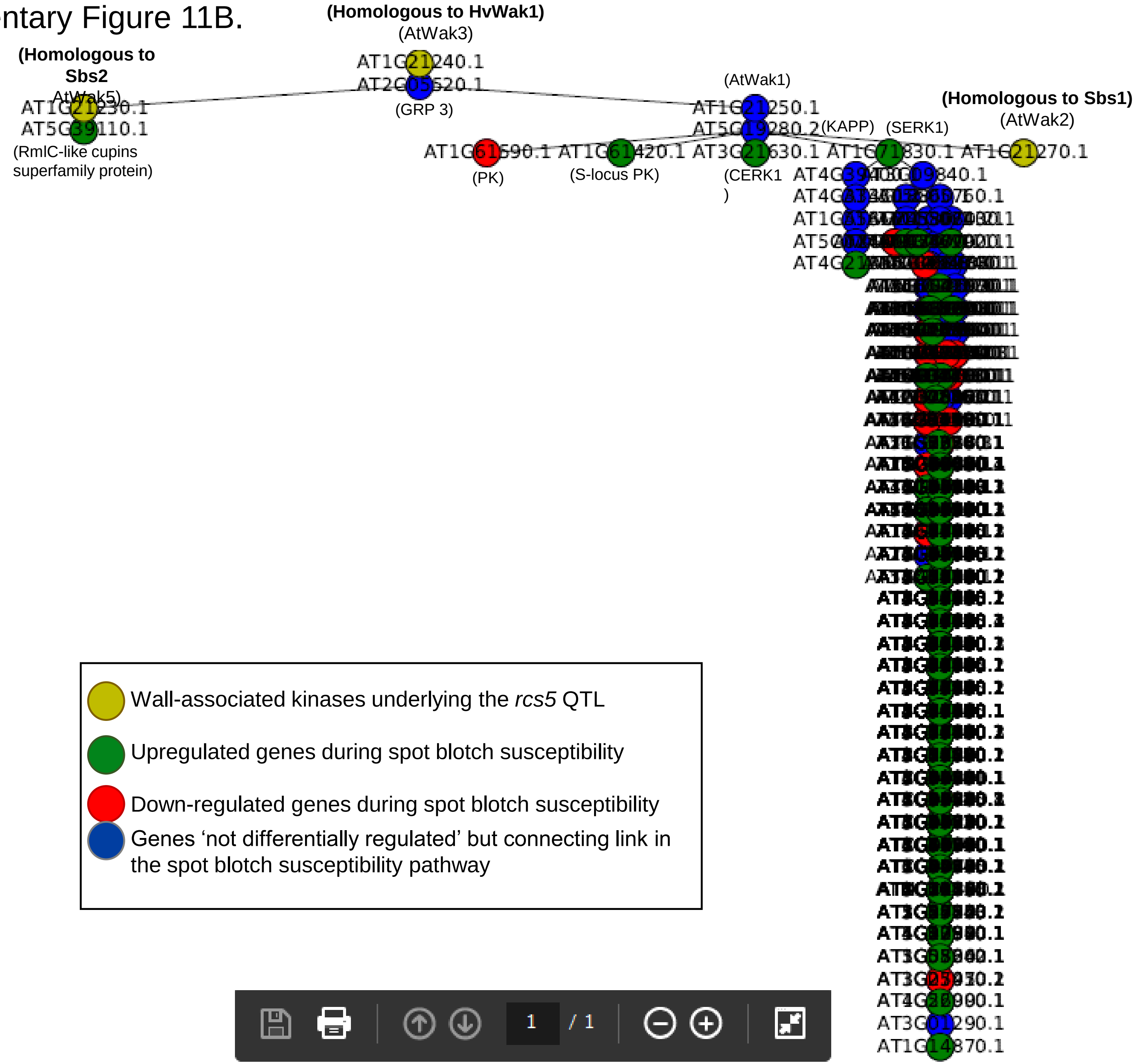

**Supplementary Table 1**

| Target | full length gene | Primer 5' - 3' | Amplicon size (bp) | Physical Position |
|---|---|---|---|---|
| HvWak2 | GA_WAK2_F1 | GGGTCAAAACGTTTGTACTAGTGAT | 1022 | 27959229 |
| | GA_WAK2_R1 | AGAGGAACTCCATCTGCATATTTAC | | 27960250 |
| | GA_WAK2_F2 | AGTTCCGTTAAAACCACATCCTTAC | 1092 | 27960081 |
| | GA_WAK2_R2_Gdna | CTTTTCTGATACTCAGCCCATAGAG | | 27961172 |
| | GA_WAK2_F3_Gdna | TTATCAGTCTGCATTCATACCATGC | 895 | 27960974 |
| | GA_WAK2_R3_Gdna | AAGCTGTAGACGTCGCTCTTCTCT | | 27961868 |
| | GA_WAK2_F4_Gdna | GAACATCCTCTTGGACGACAAGTA | 1001 | 27961695 |
| | GA_WAK2_R4_Gdna | GCACGGGTCTTTTGCTAGTATTTAT | | 27962695 |
| Sbs1 | GA_WAK3_F1_Gdna | ATGGAGCGCACAGTATAGATAATTC | 931 | 28039259 |
| | GA_WAK3_R1_Gdna | GATGTCAACTTGTTCTTGGTGTTG | | 28038693 |
| | GA_WAK3_F2_Gdna | GTTCAGAATCCACATATAGCAACAC | 890 (Morex); 1525 (Steptoe) | 28038817 |
| | GA_WAK3_R2_Gdna | AATTCATAGACCAGTATGGGGACTT | | 28037928 |
| | GA_WAK3_F3_Gdna | GGATAACAGGGTAATAGCTGTCAAA | 567 | 28038090 |
| | GA_WAK3_R3_Gdna | CATCTGAAACTATCATCTGGGTGTA | | n/a |
| Sbs2 | GA_WAK4_F0_Gdna | TCACCAAAGGTAAGGTTGATGTCTA | 863 | 28145586 |
| | GA_WAK4_R0_Gdna | CTTTGCTGAAATGACCAATATACCC | | 28144605 |
| | GA_WAK4_F1_Gdna | GCATCCTCCTTGTAGGTAAGGATTA | 903 | 28146229 |
| | GA_WAK4_R1_Gdna | GGTCAGAAGCTCTTACTACTTGTCG | | 28145327 |
| | GA_WAK4_F2_Gdna | ATCCTTAAGAATGCCCTTGTACACT | 979 | 28147055 |
| | GA_WAK4_R2_Gdna | GGTAATGGAAGTTGTTCAGAAGCTA | | 28146077 |
| | GA_WAK4_F3_Gdna | GACTGAAACTATCATCTGGGTGTCT | 1097 | 28148112 |
| | GA_WAK4_R3_Gdna | GAACAGAATGGTGGTCAGATATTGT | | 28146888 |
| HvWak5 | GA_WAK5_F1_Gdna | TACATGTCTTGATTGTTGCCTACAG | 916 | 28135129 |
| | GA_WAK5_R1_Gdna | TGTAGGTATGGATTGCCCTCATAA | | 28136044 |
| | GA_WAK5_F2_Gdna | TTGATCGTTCATATGTCAGCTCTAC | 1039 | 28135828 |
| | GA_WAK5_R2_Gdna | CCTTCTTAAGGTCTTCTTGTGTGAA | | 28136821 |
| | GA_WAK5_F3_Gdna | CGAGTATCAGAAAAGGAAGTTGAAG | 1129 | 28136685 |
| | GA_WAK5_R3_Gdna | CCTGGGTGTTTCTATACTTATTCCA | | 28137813 |
| | **Walking** | | | |
| | AP1 | GTAATACGACTCACTATAGGGC | Adaptor primer | |
| | AP2 | ACTATAGGGCACGCGTGGT | | |
| Sbs2 | ctg35_2_GSP3' | ACCGTCATGATTTTCAGGCACATCCG | second Walk | 28038369 |
| | ctg35_2_GSP5' | CACTGCCTACAGGTCAGAAGCTCTTA CTA | second Walk | 28039080 |
| | CTG35_GSP3'_1 | GTGCGAACCTCCAAACCAGTCCTTGT ATCC | First Walk | 27960562 |
| | CTG35_GSP3'_2 | CACCGTTGGGAGCTACACCTGTTCATG CTC | nested to first | 27960613 |
| | CTG35_GSP5'_1 | CAAGCTGGTTCTTGCCCTTGGCTTGTC CC | First Walk | 28038634 |

| | | | | |
|---|---|---|---|---|
| | CTG35_GSP5'_2 | CCATTGCAAGTCCACCACAGCCGATTGATG | nested to first | 28038667 |
| | ctg35_3_GSP3 | CCCGAACGGCACTCTTTTTCAGTTCATCCA | Third Walk | 28037924 |
| HvWak5 | Ctg37_2_GSP5'_1 | TCCATCCCGGCGCTAACTATGCTTGCATCC | second Walk | 28137551 |
| | Ctg37_2_GSP5'_2 | GGATGGCATCAAGGCTTCCATCCTTCATCG | nested to second | 28137520 |
| | CTG37_GSP3'_1 | TCCATCGACGAGCACGGTCTCGACTG | First Walk | 28137750 |
| | CTG37_GSP3'_2 | CATGTCTGGAATAAGTATAGAAACACCCAGGT | nested to first | 28137783 |
| | CTG37_GSP5'_1 | CAGGTGAGGAGCACATGACCAGACGCTCTG | First Walk | 28137727 |
| | CTG37_GSP5'_2 | TCTCAAGCACGAGTTTCGCCCTCCAGGTGC | nested to first | 28137688 |
| | **Markers** | | | |
| BF627428 | BF627428_F | GGTATGACAGGAAAAGAGAGACAGT | 1031 | 30199300 |
| | BF627428_R | CTATTGTTACAACTTGTGTGGTTGC | | 30200330 |
| cMWG773 | cMWG773-5_F | CTGAATGAATCTCGTCGAAACTACT | 865 | 26188406 |
| | cMWG773-5_R | ACGAATCTGGACTTCATCTTGGAC | | 26189270 |
| BF627428_Steptoe | GA_Step_BF627428_F1 | GTTGCTGCATTCGGTTTCAGTTTC | 1031, 727 | steptoe specific |
| | BF627428_R | CTATTGTTACAACTTGTGTGGTTGC | | 30200330 |
| BF627428_Morex | BF627428_F | GGTATGACAGGAAAAGAGAGACAGT | 1031, 355 | 30199300 |
| | GA_Mor_BF627428_R1 | TCATGACCGATCATGTGATAATTT | | 30199654 |
| cMWG773_Steptoe | GA_Step_CMWG773-5_F1 | ACGATGAGCCGCGAGCCTCGAC | 865, 217 | steptoe specific |
| | cMWG773-5_R | ACGAATCTGGACTTCATCTTGGAC | | 26189270 |
| cMWG773_Morex | cMWG773-5_F | CTGAATGAATCTCGTCGAAACTACT | 865, 689 | 26188406 |
| | GA_MOR_CMWG773-5_R1 | GCTCTTCAGGAGGAGAATGGCATTG | | 26189094 |
| HvWak1 SNP | GA_rcs5_1584_F1 | GTCTTAGGCATATCCTTCAGTACCAT | 771 | 27972826 |
| | GA_rcs5_1584_R1 | GAAACTGTACACGTCGCTCTTCTC | | 27973591 |
| Ctg7 | Ctg7_CAPS | TGTACCGGAGCATCCATCTTGATAC | 487 | 28203453 |
| | Ctg7_CAPS | GCAGTTTCTTGGGACATCGACGAAC | | 28203940 |
| BF263248 | GA_BF263248_F1 | GAGGTCGACTTCGCCAACTACTTCT | 1265 | 27861430 |
| | GA_BF263248_R1 | TGTCACATTTTTCATACCGAAAGCA | | 27862695 |
| BF257002 | GA_BF257002_F1 | CTTCTTCTCCACCCTCGGTAAGC | 781 | 27991817 |
| | GA_BF257002_R1 | CCTCCTTTTACCCAACATGTCTTCC | | 27991036 |
| 11_20162 | 11_20162_F1 | GAACTTGCTGCGCATTATTTGATCT | 614 | 28121569 |
| | 11_20162_R1 | AAAAGCACACTGCTGCTGCTCTACT | | 28122183 |
| 790 SNP | GA_790_F1 | ATGGCGTGGCAAATAAAGAAAGTCT | 2249 | 28115641 |
| | GA_790_R1 | CAGTGAAGTGTGTGAGGCAAGAAGA | | 28113392 |
| ctg_1606998 STS | GA_Mor_ctg_1606998_F1 | TCAAAAGCCAACACTCCTTCCATTA | 489 | 28116556 |
| | GA_Mor_ctg_1606998_R1 | CATGCATGTCCTATACACTTAGACGAA | | 28116067 |
| 800 SNP | GA_800_F1 | CCCAAGGAAGTGAAGTGAACTGGTA | 1514 | 28122194 |

| | | | | |
|---|---|---|---|---|
| | GA_800_R1 | CCAAAACTCCACGTTCCTTTTCTCT | | 28123708 |
| BF256735 | GA_BF256735_F1 | GGAATCGTGCCATTGATATTGTTGT | 1373 | 28140935 |
| | GA_BF256735_R1 | GTTGCTTTGCATCGTTTGGAGTATC | | 28139562 |
| W4_SNP | GA_W4_SNP_F1 | TTGACAATGGGTAACTCAAGCTGGT | 518 | 28145790 |
| | GA_W4_SNP_R1 | ATGGGCTCAGATAGATCGGTATGTA | | 28145272 |
| | **VIGS knock-down Primers** | | | |
| HvWak2 | GA-WAK2KD-FP | AGTTTAATTAAAGCAAACATCAGTTC CGTTAAAAC | 181 | 27960071 |
| | GA-WAK2KD-RP | AGTGCGGCCGCCAGAGGAACTCCATC TGCATATTTAC | | 27960252 |
| | GA-WAK2KD-REV-FP | AGTGCGGCCGCAGCAAACATCAGTTC CGTTAAAAC | 181 | 27960071 |
| | GA-WAK2KD-REV-RP | AGTTTAATTAACAGAGGAACTCCATC TGCATATTTAC | | 27960252 |
| Sbs1 | GA-WAK3KD-FP | AGTTTAATTAAATATCTGGGCTCAAAC GGACTTAG | 157 | 28038878 |
| | GA-WAK3KD-RP | AGTGCGGCCGCCGTATGAAAAGGATC AAGGGTTAAG | | 28038721 |
| | GA-WAK3KD-REV-FP | AGTGCGGCCGCATATCTGGGCTCAAA CGGACTTAG | 157 | 28038878 |
| | GA-WAK3KD-REV-RP | AGTTTAATTAACGTATGAAAAGGATC AAGGGTTAAG | | 28038721 |
| Sbs2 | GA-WAK4KD2-FP | AGTTTAATTAATGAAGGTGTAATTATG GACTGAAAC | 156 | 28148131 |
| | GA-WAK4KD2-RP | AGTGCGGCCGCCATGGGAAAATAGAT CGTCTGGTC | | 28147975 |
| | GA-WAK4KD2-REV-FP | AGTGCGGCCGCTGAAGGTGTAATTAT GGACTGAAAC | 156 | 28148131 |
| | GA-WAK4KD2-REV-RP | AGTTTAATTAACATGGGAAAATAGAT CGTCTGGTC | | 28147975 |
| HvWak5 | GA-WAK5KD-FP | AGTTTAATTAACTCTGCGAAGCACCTG GAGGGCGAAAC | 165 | 28137650 |
| | GA-WAK5KD-RP1 | AGTGCGGCCGCACCTGGGTGTTTCTAT ACTTATTCC | | 28137815 |
| | GA-WAK5KD-REV-FP | AGTGCGGCCGCCTCTGCGAAGCACCT GGAGGGCGAAAC | 165 | 28137650 |
| | GA-WAK5KD-REV-RP1 | AGTTTAATTAAACCTGGGTGTTTCTAT ACTTATTCC | | 28137815 |
| | **VIGS-q-PCR Primers** | | | |
| HvuSnoR14 | HvuSnoR14 -F1 | GATGTTTATGTATGATAGTCTGTC | 67 | 34228595, 34229909 |
| | HvuSnoR14 -R1 | GTCGGGATGTATGCGTGTC | | 34228529, 34229843 |
| Sbs1 | SS-W3-QFP1 | CTTGAGTACCTCACCGCTGACTCAT | 63 | 28038637 |
| | SS-W3-QRP1 | ATTACCGATGCTGCTTACGTCCAT | | 28038575 |
| Sbs2 | WAK4_QPCR_F6 | TCCGCTGGTGTAGTCTTCCTCATAG | 197 | 28146795 |
| | WAK4_QPCR_R6 | AAACCGCTTGTAGCCTTCTTCAGAT | | 28146991 |
| HvWak5 | W5-QF1 | CCGCGCCAGTACCAACTATATACAA | 97 | 28135738 |
| | W5-QR1 | CGATCAAACTTGAACAAGTCCACCT | | 28135834 |
| | **time-course q-PCR Primers** | | | |
| HvWak2 | WAK2Q-F1 | GGCATAGAGACGCCCCGATGATCAG | 180 | 27962272 |
| | WAK2Q-R1 | AAGCTCAGATGCCGCATAGTCAGAC | | 27962455 |

| | | | | |
|---|---|---|---|---|
| Sbs1 | WAK3Q-F1 | ATATCTGGGCTCAAACGGACTTAG | 157 | 28038878 |
| | WAK3Q-R1 | CGTATGAAAAGGATCAAGGGTTAAG | | 28038722 |
| Sbs2 | WAK4Q-F1 | GATGAAGGTGTAATTATGGACTGAA | 158 | 28148133 |
| | WAK4Q-R1 | CATGGGAAAATAGATCGTCTGGTC | | 28147975 |
| HvWak5 | W5-QF1 | CCGCGCCAGTACCAACTATATACAA | 97 | 28135738 |
| | W5-QR1 | CGATCAAACTTGAACAAGTCCACCT | | 28135834 |
| | **Allele Analysis Primers** | | | |
| Sbs1 | Sbs1-F4 | CTCAAGTGTATGGTGAAACCTAGTGC | 4347 | 28169296 |
| | Sbs1_Pro_R1 | GAAACAGAGCAAGTAGATGGGAGCA | | n/a |
| Sbs2 | Sbs2_F3 | GACTGAAACTATCATCTGGGTGTCT | 4885 | 28148112 |
| | Sbs2_R5 | TCCTCCTGCTGTCCAAATTAAACAA | | 28142992 |

SUPPLEMENTARY TABLE 2

| S No | Feature ID | Barley Annotation | EDGE test: Noninoc_Harr vs B.s.72Hr_Harr- Fold change | Ara. homology |
|---|---|---|---|---|
| 1 | HORVU5Hr1G051970 | Pathogenesis-related thaumatin superfamily protein | 19346.11 | AT4G11650.1 |
| 2 | HORVU7Hr1G033530 | Cysteine-rich venom protein | 13346.17 | AT2G14610.1 |
| 3 | HORVU3Hr1G111600 | Pathogenesis-related protein | 11819.76 | AT1G64790.1 |
| 4 | HORVU1Hr1G089540 | Chalcone synthase 2 | 11360.32 | AT5G13930.1 |
| 5 | HORVU1Hr1G089520 | O-methyltransferase family protein | 10060.05 | AT4G35160.1 |
| 6 | HORVU1Hr1G065150 | undescribed protein | 8959.19 | AT2G43550.1 |
| 7 | HORVU3Hr1G113120 | Wound-induced protein | 8509.44 | AT3G04720.1 |
| 8 | HORVU7Hr1G122120 | Thaumatin-like pathogenesis-related protein 1 | 8398.99 | AT4G11650.1 |
| 9 | HORVU5Hr1G055950 | Cysteine-rich venom protein | 8287.05 | AT4G33720.1 |
| 10 | HORVU2Hr1G018510 | Peroxidase superfamily protein | 8262.48 | AT5G05340.1 |
| 11 | HORVU5Hr1G056040 | Cysteine-rich venom protein | 6896.29 | AT4G33720.1 |
| 12 | HORVU3Hr1G113620 | Wound-induced protein | 6399.76 | AT3G04720.1 |
| 13 | HORVU3Hr1G111620 | undescribed protein | 5502.03 | AT5G65090.3 |
| 14 | HORVU5Hr1G106010 | Cysteine-rich secretory protein 3 | 5380.60 | AT3G19690.1 |
| 15 | HORVU3Hr1G105630 | Glucan endo-1,3-beta-glucosidase GI | 4735.11 | AT4G16260.1 |
| 16 | HORVU5Hr1G005230 | Unknown | 3889.00 | #N/A |
| 17 | HORVU2Hr1G028670 | Chymotrypsin inhibitor | 3220.06 | AT5G43580.1 |
| 18 | HORVU1Hr1G089620 | O-methyltransferase 1 | 3205.48 | AT5G54160.1 |
| 19 | HORVU2Hr1G044340 | Peroxidase superfamily protein | 3094.16 | AT5G05340.1 |
| 20 | HORVU1Hr1G065670 | undescribed protein | 2872.94 | AT5G47910.1 |
| 21 | HORVU1Hr1G065660 | undescribed protein | 2397.53 | AT4G19140.1 |
| 22 | HORVU2Hr1G076320 | undescribed protein | 2212.32 | AT4G34600.1 |

| | | | | |
|---|---|---|---|---|
| 23 | HORVU3Hr1G006440 | Xylanase inhibitor | 2061.09 | AT1G03230.1 |
| 24 | HORVU1Hr1G089700 | O-methyltransferase 1 | 1985.02 | AT5G54160.1 |
| 25 | HORVU3Hr1G006450 | Xylanase inhibitor | 1514.40 | AT1G03230.1 |
| 26 | HORVU3Hr1G003980 | Bowman-Birk type trypsin inhibitor | 1395.88 | ATCG00180.1 |
| 27 | HORVU1Hr1G019750 | Laccase-19 | 1345.50 | AT3G09220.1 |
| 28 | HORVU5Hr1G067760 | Kiwellin | 1295.61 | #N/A |
| 29 | HORVU5Hr1G010130 | undescribed protein | 1246.50 | AT4G16120.1 |
| 30 | HORVU2Hr1G004550 | Cytochrome P450 superfamily protein | 1222.78 | AT1G64940.1 |
| 31 | HORVU4Hr1G005380 | germin-like protein 4 | 1151.80 | AT1G18980.1 |
| 32 | HORVU6Hr1G014600 | Bowman-Birk type trypsin inhibitor | 1150.36 | AT1G61420.1 |
| 33 | HORVU1Hr1G065650 | undescribed protein | 1079.97 | AT2G05635.2 |
| 34 | HORVU5Hr1G023720 | Pathogenesis-related protein STH-2 | 1041.15 | AT1G35260.1 |
| 35 | HORVU6Hr1G001850 | undescribed protein | 1030.78 | AT2G29780.1 |
| 36 | HORVU0Hr1G005300 | 2-oxoglutarate (2OG) and Fe(II)-dependent oxygenase superfamily protein | 1014.24 | AT5G24530.1 |
| 37 | HORVU2Hr1G118020 | undescribed protein | 982.87 | AT1G09580.1 |
| 38 | HORVU2Hr1G118010 | Leucine-rich receptor-like protein kinase family protein | 942.08 | AT3G47570.1 |
| 39 | HORVU5Hr1G122520 | undescribed protein | 882.93 | AT3G25727.1 |
| 40 | HORVU3Hr1G105560 | Glucan endo-1,3-beta-glucosidase GI | 853.01 | AT4G16260.1 |
| 41 | HORVU4Hr1G005440 | germin-like protein 4 | 852.12 | AT1G18980.1 |
| 42 | HORVU0Hr1G000840 | undescribed protein | 850.04 | AT5G47910.1 |
| 43 | HORVU2Hr1G004600 | Cytochrome P450 superfamily protein | 848.71 | AT3G26310.1 |
| 44 | HORVU2Hr1G004530 | Cytochrome P450 superfamily protein | 847.84 | AT5G25120.1 |
| 45 | HORVU2Hr1G004640 | Cytochrome P450 superfamily protein | 795.77 | AT3G26300.1 |
| 46 | HORVU3Hr1G100190 | Jasmonate O-methyltransferase | 793.81 | AT5G04370.1 |
| 47 | HORVU2Hr1G004540 | KS protein | 765.98 | AT1G79460.1 |
| 48 | HORVU7Hr1G085450 | Double-strand break repair protein MRE11 | 751.32 | AT5G54260.1 |
| 49 | HORVU2Hr1G018430 | Peroxidase superfamily protein | 729.65 | AT5G05340.1 |
| 50 | HORVU6Hr1G088440 | 2-oxoglutarate (2OG) and Fe(II)-dependent oxygenase superfamily protein | 659.14 | AT5G20400.1 |
| 51 | HORVU7Hr1G011930 | undescribed protein | 648.70 | AT1G45474.2 |
| 52 | HORVU3Hr1G001950 | undescribed protein | 635.14 | AT4G39970.1 |
| 53 | HORVU2Hr1G117940 | Heavy metal transport/detoxification superfamily protein | 624.95 | AT1G01490.2 |
| 54 | HORVU0Hr1G009470 | germin-like protein 2 | 611.78 | AT5G39110.1 |
| 55 | HORVU2Hr1G114390 | L-tyrosine decarboxylase | 603.79 | AT2G20340.1 |
| 56 | HORVU6Hr1G066380 | germin-like protein 1 | 593.23 | AT1G72610.1 |
| 57 | HORVU2Hr1G004620 | Copalyl diphosphate synthase 2 | 593.07 | AT4G02780.1 |
| 58 | HORVU3Hr1G109350 | receptor kinase 1 | 588.64 | AT4G03230.3 |
| 59 | HORVU3Hr1G105600 | Glucan endo-1,3-beta-glucosidase GI | 559.07 | AT4G16260.1 |
| 60 | HORVU4Hr1G002640 | Pathogenesis-related thaumatin superfamily protein | 550.05 | AT4G11650.1 |

| | | | | |
|---|---|---|---|---|
| 61 | HORVU7Hr1G011940 | undescribed protein | 544.93 | AT5G60540.1 |
| 62 | HORVU2Hr1G095460 | Glutathione S-transferase family protein | 542.73 | AT3G62760.1 |
| 63 | HORVU5Hr1G023730 | Pathogenesis-related protein STH-21 | 524.53 | AT1G35260.1 |
| 64 | HORVU4Hr1G071370 | Membrane protein | 520.85 | AT5G47530.1 |
| 65 | HORVU7Hr1G012240 | undescribed protein | 507.69 | AT5G63840.2 |
| 66 | HORVU0Hr1G011720 | Major pollen allergen Bet v 1-B | 507.17 | AT1G35260.1 |
| 67 | HORVU6Hr1G053710 | High-affinity nitrate transporter 3.1 | 504.55 | AT5G50200.3 |
| 68 | HORVU4Hr1G002650 | Pathogenesis-related thaumatin superfamily protein | 495.88 | AT4G11650.1 |
| 69 | HORVU3Hr1G071470 | ABC transporter G family member 32 | 486.24 | AT1G15520.1 |
| 70 | HORVU2Hr1G114440 | L-tyrosine decarboxylase | 478.18 | AT2G20340.1 |
| 71 | HORVU2Hr1G028660 | Chymotrypsin inhibitor | 453.27 | AT2G38900.2 |
| 72 | HORVU6Hr1G094790 | receptor-like protein kinase 1 | 449.17 | AT3G21630.1 |
| 73 | HORVU5Hr1G124650 | ABC transporter G family member 48 | 447.49 | AT1G66950.1 |
| 74 | HORVU7Hr1G121260 | Flavin-containing monooxygenase family protein | 430.35 | AT1G19250.1 |
| 75 | HORVU2Hr1G028720 | Chymotrypsin inhibitor | 405.31 | AT5G43580.1 |
| 76 | HORVU3Hr1G105480 | 0 | 398.95 | #N/A |
| 77 | HORVU5Hr1G023660 | unknown function | 396.17 | AT5G51770.1 |
| 78 | HORVU5Hr1G009940 | undescribed protein | 388.82 | AT2G33150.1 |
| 79 | HORVU1Hr1G001220 | receptor kinase 3 | 387.96 | AT4G23180.3 |
| 80 | HORVU2Hr1G083570 | Cytochrome P450 superfamily protein | 385.65 | AT3G26330.1 |
| 81 | HORVU3Hr1G064470 | Chitinase family protein | 385.21 | AT3G12500.1 |
| 82 | HORVU5Hr1G072520 | FAD-binding Berberine family protein | 383.95 | AT4G20820.1 |
| 83 | HORVU5Hr1G010120 | Protein WIR1A | 370.99 | AT2G23450.2 |
| 84 | HORVU3Hr1G105420 | Glucan endo-1,3-beta-glucosidase GI | 368.52 | AT3G57240.1 |
| 85 | HORVU3Hr1G091780 | undescribed protein | 368.10 | AT2G18140.1 |
| 86 | HORVU3Hr1G099760 | L-gulonolactone oxidase 2 | 361.55 | AT2G46750.1 |
| 87 | HORVU7Hr1G002740 | laccase 7 | 347.36 | AT3G09220.1 |
| 88 | HORVU3Hr1G032370 | Isoflavone reductase-like protein | 346.99 | AT4G39230.1 |
| 89 | HORVU3Hr1G105290 | Glucan endo-1,3-beta-glucosidase GI | 346.83 | AT3G57240.1 |
| 90 | HORVU3Hr1G104940 | Leucine-rich repeat receptor-like protein kinase family protein | 343.65 | AT1G71830.1 |
| 91 | HORVU2Hr1G114450 | L-tyrosine decarboxylase | 333.20 | AT2G20340.1 |
| 92 | HORVU3Hr1G095730 | Glutathione S-transferase family protein | 328.32 | AT1G10360.1 |
| 93 | HORVU7Hr1G019380 | Cytochrome P450 superfamily protein | 325.48 | AT3G26300.1 |
| 94 | HORVU5Hr1G118800 | Glutathione S-transferase family protein | 319.93 | AT3G62760.1 |
| 95 | HORVU3Hr1G091740 | Peroxidase superfamily protein | 316.99 | AT5G06730.1 |
| 96 | HORVU4Hr1G079620 | Cytochrome P450 superfamily protein | 316.39 | AT2G26170.1 |
| 97 | HORVU2Hr1G018480 | Peroxidase superfamily protein | 316.07 | AT5G05340.1 |
| 98 | HORVU3Hr1G112060 | undescribed protein | 312.59 | AT2G42270.1 |
| 99 | HORVU3Hr1G097860 | Laccase-19 | 310.87 | AT3G09220.1 |

| | | | | |
|---|---|---|---|---|
| 100 | HORVU1Hr1G009490 | Bifunctional inhibitor/lipid-transfer protein/seed storage 2S albumin superfamily protein | 309.84 | AT3G22600.1 |
| 101 | HORVU3Hr1G094340 | Chromosome 3B, genomic scaffold, cultivar Chinese Spring | 302.23 | AT1G14590.1 |
| 102 | HORVU4Hr1G074780 | Protein of unknown function (DUF1997) | 297.15 | AT5G39530.1 |
| 103 | HORVU1Hr1G062380 | CASP-like protein 1 | 286.71 | AT3G06390.1 |
| 104 | HORVU5Hr1G106020 | Cysteine-rich venom protein | 285.11 | AT3G19690.1 |
| 105 | HORVU2Hr1G109640 | Cytochrome P450 superfamily protein | 277.03 | AT2G45550.1 |
| 106 | HORVU7Hr1G084920 | Chitinase 1 | 276.30 | AT1G14430.1 |
| 107 | HORVU1Hr1G049280 | Glutathione S-transferase family protein | 276.11 | AT1G10370.1 |
| 108 | HORVU3Hr1G002900 | receptor kinase 2 | 273.36 | AT1G70520.1 |
| 109 | HORVU1Hr1G010120 | L-tyrosine decarboxylase | 271.34 | AT2G20340.1 |
| 110 | HORVU1Hr1G090930 | unknown function | 263.36 | AT2G31900.4 |
| 111 | HORVU5Hr1G123550 | receptor kinase 2 | 262.20 | AT4G21390.1 |
| 112 | HORVU7Hr1G017640 | Enolase | 261.44 | AT2G36530.1 |
| 113 | HORVU5Hr1G052150 | Indole-3-glycerol phosphate synthase | 258.86 | AT5G48220.1 |
| 114 | HORVU1Hr1G043910 | Plant basic secretory protein (BSP) family protein | 258.15 | AT2G15130.1 |
| 115 | HORVU1Hr1G052430 | Chitinase family protein | 250.79 | AT3G12500.1 |
| 116 | HORVU7Hr1G081700 | lipid phosphate phosphatase 3 | 249.44 | AT3G02600.9 |
| 117 | HORVU3Hr1G099530 | UDP-Glycosyltransferase superfamily protein | 247.29 | AT3G11340.1 |
| 118 | HORVU1Hr1G090860 | unknown function | 239.77 | AT3G62650.2 |
| 119 | HORVU6Hr1G008640 | catalase 1 | 229.19 | AT1G20630.1 |
| 120 | HORVU5Hr1G122410 | undescribed protein | 229.15 | AT1G69850.1 |
| 121 | HORVU3Hr1G091750 | Peroxidase superfamily protein | 226.99 | AT5G06730.1 |
| 122 | HORVU1Hr1G008360 | laccase 7 | 225.60 | AT3G09220.1 |
| 123 | HORVU4Hr1G056890 | LOB domain-containing protein 1 | 225.53 | AT1G07900.1 |
| 124 | HORVU2Hr1G112060 | undescribed protein | 214.70 | AT1G49560.1 |
| 125 | HORVU5Hr1G009930 | undescribed protein | 207.62 | AT5G37660.1 |
| 126 | HORVU7Hr1G085420 | undescribed protein | 204.05 | AT3G29830.2 |
| 127 | HORVU1Hr1G009800 | Bifunctional inhibitor/lipid-transfer protein/seed storage 2S albumin superfamily protein | 203.95 | AT3G22600.1 |
| 128 | HORVU2Hr1G126150 | 1-deoxy-D-xylulose 5-phosphate synthase 1 | 202.30 | AT4G15560.1 |
| 129 | HORVU7Hr1G008830 | Glutathione S-transferase family protein | 202.26 | AT1G10360.1 |
| 130 | HORVU2Hr1G119500 | Late embryogenesis abundant (LEA) hydroxyproline-rich glycoprotein family | 202.01 | AT5G36970.1 |
| 131 | HORVU7Hr1G075760 | L-gulonolactone oxidase 2 | 200.61 | AT2G46750.1 |
| 132 | HORVU5Hr1G115870 | PLANT CADMIUM RESISTANCE 2 | 196.48 | AT1G14870.1 |
| 133 | HORVU5Hr1G119640 | undescribed protein | 187.67 | AT1G33600.2 |
| 134 | HORVU3Hr1G077880 | Chromosome 3B, genomic scaffold, cultivar Chinese Spring | 187.34 | AT3G20340.1 |
| 135 | HORVU3Hr1G104990 | undescribed protein | 186.79 | #N/A |

| | | | | |
|---|---|---|---|---|
| 136 | HORVU6Hr1G004770 | unknown function | 186.42 | AT1G21230.1 |
| 137 | HORVU3Hr1G112050 | undescribed protein | 185.93 | AT3G55640.3 |
| 138 | HORVU5Hr1G083340 | purple acid phosphatase 15 | 185.01 | AT3G07130.1 |
| 139 | HORVU2Hr1G101150 | xyloglucan endotransglucosylase/hydrolase 13 | 179.39 | AT5G57540.1 |
| 140 | HORVU6Hr1G069710 | receptor kinase 2 | 179.04 | AT4G21390.1 |
| 141 | HORVU5Hr1G081620 | Two-component response regulator-like PRR95 | 177.28 | AT2G46790.3 |
| 142 | HORVU7Hr1G116650 | Glutathione S-transferase family protein | 176.36 | AT3G62760.1 |
| 143 | HORVU4Hr1G077300 | Bowman-Birk type trypsin inhibitor | 173.76 | AT1G18197.1 |
| 144 | HORVU7Hr1G010840 | L-tyrosine decarboxylase | 166.40 | AT4G28680.4 |
| 145 | HORVU2Hr1G124300 | Glutathione S-transferase family protein | 165.37 | AT1G78380.1 |
| 146 | HORVU3Hr1G013380 | Leucine-rich repeat protein kinase family protein | 161.83 | AT4G29990.2 |
| 147 | HORVU1Hr1G062030 | Chitinase family protein | 160.17 | AT3G12500.1 |
| 148 | HORVU2Hr1G001780 | WRKY DNA-binding protein 75 | 158.55 | AT5G13080.1 |
| 149 | HORVU5Hr1G000940 | receptor kinase 2 | 158.16 | AT5G10530.1 |
| 150 | HORVU6Hr1G061420 | Cytochrome b5 | 157.71 | AT5G53560.1 |
| 151 | HORVU7Hr1G011950 | undescribed protein | 157.31 | AT5G35620.3 |
| 152 | HORVU6Hr1G013690 | undescribed protein | 154.39 | AT3G51350.1 |
| 153 | HORVU3Hr1G002000 | germin-like protein 9 | 141.37 | AT5G38960.1 |
| 154 | HORVU4Hr1G071300 | LL-diaminopimelate aminotransferase | 137.71 | AT4G33680.1 |
| 155 | HORVU5Hr1G009840 | RAB GDP dissociation inhibitor 2 | 134.35 | AT3G59920.1 |
| 156 | HORVU6Hr1G020540 | Cytochrome P450 superfamily protein | 133.75 | AT3G26230.1 |
| 157 | HORVU0Hr1G008950 | Branched-chain-amino-acid aminotransferase 6 | 131.78 | AT3G49680.1 |
| 158 | HORVU3Hr1G084210 | ferredoxin 3 | 129.03 | AT2G27510.1 |
| 159 | HORVU2Hr1G102710 | Protein NRT1/ PTR FAMILY 8.3 | 128.37 | AT2G02040.1 |
| 160 | HORVU6Hr1G005720 | high affinity nitrate transporter 2.6 | 127.85 | AT1G08090.1 |
| 161 | HORVU5Hr1G096850 | Cytochrome P450 superfamily protein | 127.08 | AT5G57260.1 |
| 162 | HORVU2Hr1G122630 | early nodulin-like protein 12 | 126.46 | AT5G26330.1 |
| 163 | HORVU7Hr1G017810 | Protein kinase superfamily protein | 126.38 | AT5G60900.1 |
| 164 | HORVU3Hr1G074960 | Peroxidase superfamily protein | 126.01 | AT4G11290.1 |
| 165 | HORVU7Hr1G010990 | L-tyrosine decarboxylase | 124.38 | AT2G20340.1 |
| 166 | HORVU2Hr1G039670 | Disease resistance protein (TIR-NBS class) | 123.44 | AT1G52900.1 |
| 167 | HORVU3Hr1G099550 | Eukaryotic aspartyl protease family protein | 122.91 | AT1G03230.1 |
| 168 | HORVU3Hr1G086840 | C2H2-like zinc finger protein | 122.76 | AT2G28710.1 |
| 169 | HORVU7Hr1G107360 | SAUR-like auxin-responsive protein family | 119.45 | AT1G56150.1 |
| 170 | HORVU3Hr1G096280 | Sec14p-like phosphatidylinositol transfer family protein | 119.39 | AT1G01630.1 |
| 171 | HORVU1Hr1G071380 | unknown function | 119.26 | AT1G74540.1 |
| 172 | HORVU7Hr1G080190 | ADP-ribosylation factor 3 | 118.48 | AT3G62560.1 |
| 173 | HORVU3Hr1G011870 | Cytochrome P450 superfamily protein | 118.16 | AT5G57260.1 |
| 174 | HORVU7Hr1G007480 | receptor-like protein kinase 2 | 117.76 | AT3G47570.1 |
| 175 | HORVU1Hr1G022430 | Leucine aminopeptidase 1 | 117.62 | AT2G24200.3 |

| | | | | |
|---|---|---|---|---|
| 176 | HORVU1Hr1G091960 | undescribed protein | 115.99 | AT1G06475.1 |
| 177 | HORVU2Hr1G124960 | Peroxidase superfamily protein | 115.12 | AT1G71695.1 |
| 178 | HORVU7Hr1G009310 | 2-oxoglutarate (2OG) and Fe(II)-dependent oxygenase superfamily protein | 115.08 | AT5G20400.1 |
| 179 | HORVU2Hr1G102720 | Protein NRT1/ PTR FAMILY 8.3 | 115.02 | AT2G02040.1 |
| 180 | HORVU3Hr1G096360 | HXXXD-type acyl-transferase family protein | 114.43 | AT3G62160.1 |
| 181 | HORVU1Hr1G089530 | sugar transporter protein 7 | 113.65 | AT1G19450.1 |
| 182 | HORVU5Hr1G099460 | NAC domain protein, | 113.54 | AT2G43000.1 |
| 183 | HORVU2Hr1G071570 | Potassium transporter family protein | 113.10 | AT4G13420.1 |
| 184 | HORVU0Hr1G019300 | Glutathione S-transferase family protein | 113.02 | AT3G09270.1 |
| 185 | HORVU7Hr1G043540 | Cytochrome P450 superfamily protein | 112.84 | AT3G48280.1 |
| 186 | HORVU6Hr1G089460 | dihydroflavonol 4-reductase-like1 | 112.56 | AT1G15950.3 |
| 187 | HORVU6Hr1G012640 | Chymotrypsin inhibitor | 110.29 | AT5G43580.1 |
| 188 | HORVU3Hr1G026330 | Subtilisin-like protease | 109.25 | AT4G10540.1 |
| 189 | HORVU7Hr1G118770 | Disease resistance-responsive (dirigent-like protein) family protein | 109.23 | AT5G42500.1 |
| 190 | HORVU2Hr1G122610 | Blue copper protein | 109.00 | AT2G31050.1 |
| 191 | HORVU7Hr1G008140 | receptor-like protein kinase 4 | 108.39 | AT5G10530.1 |
| 192 | HORVU2Hr1G063690 | MATE efflux family protein | 106.73 | AT3G23550.1 |
| 193 | HORVU3Hr1G094250 | Chromosome 3B, genomic scaffold, cultivar Chinese Spring | 105.88 | AT4G19975.1 |
| 194 | HORVU6Hr1G021870 | Homeobox-leucine zipper protein family | 105.77 | AT5G06710.1 |
| 195 | HORVU5Hr1G052760 | Heavy metal transport/detoxification superfamily protein | 104.18 | AT3G20180.3 |
| 196 | HORVU6Hr1G013710 | Plant basic secretory protein (BSP) family protein | 102.92 | AT2G15220.1 |
| 197 | HORVU7Hr1G073340 | early nodulin-like protein 15 | 101.76 | AT2G32300.2 |
| 198 | HORVU3Hr1G022780 | Protein kinase family protein | 101.54 | AT5G02070.1 |
| 199 | HORVU7Hr1G108490 | Leucine-rich receptor-like protein kinase family protein | 101.53 | AT3G47570.1 |
| 200 | HORVU6Hr1G033060 | alpha/beta-Hydrolases superfamily protein | 100.54 | AT2G36290.1 |
| 201 | HORVU2Hr1G118030 | undescribed protein | 100.30 | AT1G48980.3 |
| 202 | HORVU2Hr1G111780 | receptor-like protein kinase 4 | 99.91 | AT4G32300.1 |
| 203 | HORVU2Hr1G125250 | receptor kinase 2 | 99.46 | AT4G23180.3 |
| 204 | HORVU3Hr1G067510 | WRKY DNA-binding protein 50 | 99.06 | AT5G26170.1 |
| 205 | HORVU3Hr1G105400 | Glucan endo-1,3-beta-glucosidase GV | 99.02 | AT3G57270.1 |
| 206 | HORVU4Hr1G011760 | UDP-Glycosyltransferase superfamily protein | 98.97 | AT3G11340.1 |
| 207 | HORVU1Hr1G020720 | Peroxidase superfamily protein | 98.88 | AT5G19890.1 |
| 208 | HORVU5Hr1G115750 | PLANT CADMIUM RESISTANCE 2 | 98.29 | AT1G14870.1 |
| 209 | HORVU7Hr1G002820 | alpha dioxygenase | 97.62 | AT3G01420.1 |
| 210 | HORVU1Hr1G044000 | Plant basic secretory protein (BSP) family protein | 96.29 | AT2G15220.1 |
| 211 | HORVU2Hr1G034670 | Auxin-responsive GH3 family protein | 96.03 | AT2G14960.1 |
| 212 | HORVU3Hr1G112140 | Abscisic stress-ripening protein 1 | 95.78 | #N/A |

| | | | | |
|---|---|---|---|---|
| 213 | HORVU1Hr1G049230 | Glutathione S-transferase family protein | 94.42 | AT1G10360.1 |
| 214 | HORVU3Hr1G091730 | Peroxidase superfamily protein | 94.04 | AT5G06730.1 |
| 215 | HORVU7Hr1G121290 | flavin-dependent monooxygenase 1 | 94.00 | AT1G19250.1 |
| 216 | HORVU1Hr1G041720 | Nucleotide/sugar transporter family protein | 93.51 | AT2G28315.1 |
| 217 | HORVU2Hr1G002750 | receptor kinase 2 | 93.39 | AT5G10530.1 |
| 218 | HORVU5Hr1G080090 | serine carboxypeptidase-like 17 | 93.17 | AT1G33540.2 |
| 219 | HORVU4Hr1G018100 | undescribed protein | 92.37 | AT1G77400.1 |
| 220 | HORVU1Hr1G049290 | unknown function | 91.07 | AT5G63420.1 |
| 221 | HORVU3Hr1G087020 | alpha/beta-Hydrolases superfamily protein | 91.06 | AT2G39420.1 |
| 222 | HORVU6Hr1G001520 | protein kinase family protein | 90.80 | AT4G29990.1 |
| 223 | HORVU7Hr1G092510 | ABC transporter G family member 44 | 90.14 | AT1G15520.1 |
| 224 | HORVU2Hr1G085260 | Protein NRT1/ PTR FAMILY 4.3 | 89.49 | AT1G59740.1 |
| 225 | HORVU2Hr1G119250 | Lipase/lipooxygenase, PLAT/LH2 family protein | 89.05 | AT2G22170.1 |
| 226 | HORVU7Hr1G122110 | 0 | 88.80 | #N/A |
| 227 | HORVU6Hr1G002530 | receptor kinase 2 | 88.26 | AT1G21230.1 |
| 228 | HORVU5Hr1G099470 | NAC domain protein, | 87.17 | AT2G43000.1 |
| 229 | HORVU5Hr1G075540 | Oxidoreductase, zinc-binding dehydrogenase family protein | 85.83 | AT4G13010.1 |
| 230 | HORVU2Hr1G067710 | undescribed protein | 85.35 | AT4G34480.1 |
| 231 | HORVU2Hr1G028780 | Chymotrypsin inhibitor | 83.92 | AT2G38900.2 |
| 232 | HORVU3Hr1G023230 | UDP-Glycosyltransferase superfamily protein | 83.23 | AT3G53150.1 |
| 233 | HORVU3Hr1G111640 | Glutathione S-transferase family protein | 83.05 | AT1G10360.1 |
| 234 | HORVU7Hr1G040780 | undescribed protein | 82.79 | AT4G06634.2 |
| 235 | HORVU4Hr1G071330 | Cytochrome b561 and DOMON domain-containing protein | 82.38 | AT3G59070.1 |
| 236 | HORVU5Hr1G012040 | receptor kinase 2 | 82.02 | AT5G65600.1 |
| 237 | HORVU2Hr1G124320 | Glutathione S-transferase family protein | 81.87 | AT1G78380.1 |
| 238 | HORVU3Hr1G098610 | Leucine-rich repeat receptor-like protein kinase family protein | 80.79 | AT5G06860.1 |
| 239 | HORVU0Hr1G008830 | MLO-like protein 1 | 80.33 | AT4G02600.2 |
| 240 | HORVU2Hr1G122270 | Glutathione S-transferase family protein | 80.29 | AT1G10360.1 |
| 241 | HORVU6Hr1G066420 | HXXXD-type acyl-transferase family protein | 80.12 | AT5G48930.1 |
| 242 | HORVU5Hr1G099390 | NAC domain containing protein 42 | 79.91 | AT2G43000.1 |
| 243 | HORVU1Hr1G088550 | undescribed protein | 79.59 | AT4G10930.2 |
| 244 | HORVU2Hr1G085270 | Chitinase | 77.97 | AT3G54420.1 |
| 245 | HORVU7Hr1G034270 | Major pollen allergen Bet v 1-F/I | 77.83 | AT2G40330.1 |
| 246 | HORVU2Hr1G104240 | germin-like protein 2 | 77.53 | AT5G39160.1 |
| 247 | HORVU7Hr1G010850 | L-tyrosine decarboxylase | 76.89 | AT2G20340.1 |
| 248 | HORVU3Hr1G001840 | undescribed protein | 75.53 | #N/A |
| 249 | HORVU7Hr1G009760 | undescribed protein | 75.42 | AT4G26440.1 |
| 250 | HORVU4Hr1G007190 | Basic blue protein | 75.36 | AT2G02850.1 |
| 251 | HORVU5Hr1G118590 | ammonium transporter 2 | 75.17 | AT2G38290.1 |

| | | | | |
|---|---|---|---|---|
| 252 | HORVU3Hr1G111550 | undescribed protein | 72.80 | AT5G43210.1 |
| 253 | HORVU3Hr1G011990 | germin-like protein 2 | 72.60 | AT5G39100.1 |
| 254 | HORVU0Hr1G005160 | L-allo-threonine aldolase | 72.52 | AT1G08630.6 |
| 255 | HORVU5Hr1G066250 | Vacuolar-processing enzyme | 72.37 | AT2G25940.1 |
| 256 | HORVU2Hr1G112040 | undescribed protein | 71.99 | #N/A |
| 257 | HORVU5Hr1G079680 | UDP-glycosyltransferase 74 F1 | 70.91 | AT2G43840.1 |
| 258 | HORVU1Hr1G014450 | undescribed protein | 70.82 | AT5G23430.1 |
| 259 | HORVU7Hr1G011860 | undescribed protein | 70.01 | AT3G04960.2 |
| 260 | HORVU2Hr1G004480 | Cytochrome P450 superfamily protein | 69.89 | AT3G26300.1 |
| 261 | HORVU4Hr1G065000 | Peroxidase superfamily protein | 69.60 | AT5G06730.1 |
| 262 | HORVU7Hr1G024360 | Non-specific lipid-transfer protein | 69.16 | AT5G59310.1 |
| 263 | HORVU3Hr1G112080 | undescribed protein | 69.12 | AT2G14680.4 |
| 264 | HORVU2Hr1G067700 | undescribed protein | 68.64 | AT2G19820.1 |
| 265 | HORVU5Hr1G011540 | 2-oxoglutarate (2OG) and Fe(II)-dependent oxygenase superfamily protein | 68.53 | AT2G36690.2 |
| 266 | HORVU3Hr1G109430 | receptor kinase 2 | 67.78 | AT4G03230.3 |
| 267 | HORVU2Hr1G079580 | unknown function | 67.68 | AT2G14095.1 |
| 268 | HORVU3Hr1G006620 | Cysteine-rich receptor-like protein kinase 25 | 67.63 | AT4G23160.1 |
| 269 | HORVU1Hr1G063740 | NAC domain protein, | 67.32 | AT1G01720.1 |
| 270 | HORVU3Hr1G006140 | GDSL esterase/lipase | 67.28 | AT3G26430.1 |
| 271 | HORVU4Hr1G020230 | rRNA N-glycosidase | 67.12 | AT3G56810.1 |
| 272 | HORVU2Hr1G085280 | Chitinase | 66.83 | AT3G54420.1 |
| 273 | HORVU7Hr1G040740 | Cysteine-rich venom protein | 66.51 | AT5G57625.1 |
| 274 | HORVU2Hr1G112120 | Anthranilate N-benzoyltransferase protein 1 | 66.43 | AT5G48930.1 |
| 275 | HORVU6Hr1G077180 | undescribed protein | 66.22 | AT3G17100.2 |
| 276 | HORVU7Hr1G019390 | receptor kinase 2 | 65.88 | AT2G37710.1 |
| 277 | HORVU5Hr1G010880 | germin-like protein 2 | 65.58 | AT5G39100.1 |
| 278 | HORVU4Hr1G014670 | Calcineurin-like metallo-phosphoesterase superfamily protein | 65.22 | AT1G18480.1 |
| 279 | HORVU3Hr1G026270 | Subtilisin-like protease | 65.10 | AT4G10520.1 |
| 280 | HORVU5Hr1G114700 | UDP-Glycosyltransferase superfamily protein | 64.67 | AT3G11340.1 |
| 281 | HORVU4Hr1G053150 | Plant protein of unknown function (DUF247) | 64.49 | AT3G50170.1 |
| 282 | HORVU2Hr1G008220 | G-type lectin S-receptor-like serine/threonine-protein kinase | 64.40 | AT4G21380.2 |
| 283 | HORVU4Hr1G001250 | Caffeic acid 3-O-methyltransferase | 64.23 | AT5G54160.1 |
| 284 | HORVU6Hr1G055000 | Chalcone synthase 1 | 63.58 | AT5G13930.1 |
| 285 | HORVU4Hr1G010740 | undescribed protein | 63.30 | AT1G72060.1 |
| 286 | HORVU7Hr1G003630 | receptor kinase 2 | 62.89 | AT4G21390.1 |
| 287 | HORVU4Hr1G087840 | Protein of unknown function (DUF1218) | 62.40 | AT5G17210.1 |
| 288 | HORVU4Hr1G085820 | UDP-galactose transporter 2 | 62.21 | AT4G23010.1 |
| 289 | HORVU3Hr1G004050 | myb domain protein 62 | 62.08 | AT1G68320.1 |
| 290 | HORVU2Hr1G071270 | Ethylene-responsive transcription factor 1 | 61.56 | AT5G13330.1 |

| | | | | |
|---|---|---|---|---|
| 291 | HORVU1Hr1G092650 | UDP-Glycosyltransferase superfamily protein | 61.31 | AT3G16520.3 |
| 292 | HORVU6Hr1G038470 | unknown function | 61.12 | AT2G25735.1 |
| 293 | HORVU6Hr1G094480 | Flavin-containing monooxygenase family protein | 61.09 | AT1G12160.1 |
| 294 | HORVU4Hr1G009190 | Sec14p-like phosphatidylinositol transfer family protein | 60.74 | AT1G75170.2 |
| 295 | HORVU5Hr1G056420 | Flavin-containing monooxygenase family protein | 60.70 | AT1G19250.1 |
| 296 | HORVU7Hr1G024930 | Subtilisin-like protease | 60.24 | AT4G10540.1 |
| 297 | HORVU2Hr1G017440 | unknown function | 60.17 | AT2G39050.1 |
| 298 | HORVU2Hr1G008130 | Serine/threonine-protein kinase BRI1-like 1 | 60.08 | AT4G21380.2 |
| 299 | HORVU5Hr1G124810 | receptor kinase 1 | 60.06 | AT4G23230.1 |
| 300 | HORVU2Hr1G098710 | undescribed protein | 59.82 | AT3G49845.2 |
| 301 | HORVU3Hr1G108360 | Orotidine 5'-phosphate decarboxylase | 59.77 | AT3G54470.1 |
| 302 | HORVU3Hr1G077790 | Polyphenol oxidase, chloroplastic | 59.28 | AT3G02210.1 |
| 303 | HORVU5Hr1G077920 | beta glucosidase 11 | 59.12 | AT1G02850.2 |
| 304 | HORVU2Hr1G003230 | Chromosome 3B, genomic scaffold, cultivar Chinese Spring | 58.84 | AT3G14470.2 |
| 305 | HORVU3Hr1G059480 | Acidic endochitinase | 58.57 | AT5G24090.1 |
| 306 | HORVU4Hr1G054870 | Major pollen allergen Bet v 1-F/I | 58.46 | AT2G26040.1 |
| 307 | HORVU6Hr1G000130 | Rhodanese-like domain-containing protein 17 | 58.40 | AT5G66170.3 |
| 308 | HORVU4Hr1G066240 | Non-symbiotic hemoglobin | 58.06 | AT2G16060.1 |
| 309 | HORVU2Hr1G012140 | Leucine-rich repeat receptor-like protein kinase family protein | 57.54 | AT1G74190.1 |
| 310 | HORVU7Hr1G091880 | undescribed protein | 57.44 | AT4G37290.1 |
| 311 | HORVU0Hr1G011730 | unknown function | 57.05 | AT5G65440.2 |
| 312 | HORVU3Hr1G002830 | Glutathione S-transferase family protein | 57.02 | AT1G10360.1 |
| 313 | HORVU7Hr1G007520 | Leucine-rich repeat receptor-like protein kinase family protein | 56.78 | AT5G44700.1 |
| 314 | HORVU7Hr1G019400 | receptor kinase 1 | 56.19 | AT2G37710.1 |
| 315 | HORVU5Hr1G102900 | Cysteine-rich receptor-like protein kinase 25 | 56.02 | AT5G48540.1 |
| 316 | HORVU7Hr1G025930 | undescribed protein | 55.97 | #N/A |
| 317 | HORVU7Hr1G028710 | 2-oxoglutarate (2OG) and Fe(II)-dependent oxygenase superfamily protein | 55.70 | AT5G20400.1 |
| 318 | HORVU3Hr1G098340 | methyl esterase 1 | 55.61 | AT2G23620.1 |
| 319 | HORVU1Hr1G061140 | Pyrophosphate-energized proton pump 1 | 55.54 | AT1G15690.1 |
| 320 | HORVU3Hr1G112040 | Peroxidase superfamily protein | 55.48 | AT1G71695.1 |
| 321 | HORVU5Hr1G041660 | protein kinase family protein | 54.68 | AT1G51890.1 |
| 322 | HORVU6Hr1G001530 | protein kinase family protein | 54.29 | AT1G49100.1 |
| 323 | HORVU5Hr1G123540 | receptor kinase 1 | 54.11 | AT4G23180.1 |
| 324 | HORVU7Hr1G082180 | undescribed protein | 53.86 | AT1G16030.1 |
| 325 | HORVU1Hr1G021140 | Glutathione S-transferase family protein | 53.21 | AT3G62760.1 |
| 326 | HORVU5Hr1G057800 | L-ascorbate oxidase | 53.13 | AT4G39830.4 |
| 327 | HORVU4Hr1G005540 | germin-like protein 4 | 52.94 | AT1G18970.1 |

| | | | | |
|---|---|---|---|---|
| 328 | HORVU3Hr1G080100 | NAC domain containing protein 102 | 52.65 | AT5G63790.1 |
| 329 | HORVU6Hr1G077960 | UDP-Glycosyltransferase superfamily protein | 52.57 | AT1G22360.2 |
| 330 | HORVU2Hr1G103180 | L-lactate dehydrogenase | 52.09 | AT4G18360.4 |
| 331 | HORVU3Hr1G098620 | Chromosome 3B, genomic scaffold, cultivar Chinese Spring | 51.08 | AT4G08350.1 |
| 332 | HORVU2Hr1G044360 | Peroxidase superfamily protein | 51.03 | AT5G05340.1 |
| 333 | HORVU7Hr1G085480 | undescribed protein | 50.45 | #N/A |
| 334 | HORVU5Hr1G014170 | Basic-leucine zipper (bZIP) transcription factor family protein | 50.31 | AT5G28770.2 |
| 335 | HORVU7Hr1G031650 | sugar transporter 4 | 50.12 | AT4G36670.1 |
| 336 | HORVU7Hr1G091800 | germin-like protein 2 | 50.11 | AT5G39150.1 |
| 337 | HORVU4Hr1G067450 | sugar transporter protein 7 | 49.60 | AT5G26340.1 |
| 338 | HORVU3Hr1G006130 | GDSL esterase/lipase | 49.55 | AT3G26430.1 |
| 339 | HORVU5Hr1G111240 | receptor kinase 1 | 49.28 | AT4G21410.3 |
| 340 | HORVU4Hr1G004000 | Major facilitator superfamily protein | 49.23 | AT2G39210.1 |
| 341 | HORVU1Hr1G049070 | Glutathione S-transferase family protein | 48.41 | AT1G10370.1 |
| 342 | HORVU3Hr1G016860 | Chymotrypsin inhibitor | 48.19 | AT2G38870.1 |
| 343 | HORVU1Hr1G084900 | 1-deoxy-D-xylulose 5-phosphate synthase 1 | 48.06 | AT4G15560.1 |
| 344 | HORVU3Hr1G079620 | Protein RETICULATA-RELATED 3, chloroplastic | 47.84 | AT3G08640.1 |
| 345 | HORVU7Hr1G029460 | undescribed protein | 47.18 | AT2G17780.4 |
| 346 | HORVU7Hr1G114660 | Indole-3-glycerol phosphate synthase | 47.09 | AT2G04400.1 |
| 347 | HORVU2Hr1G110230 | Protein kinase superfamily protein | 46.97 | AT1G78290.3 |
| 348 | HORVU3Hr1G074230 | Auxin-responsive GH3 family protein | 46.93 | AT4G37390.1 |
| 349 | HORVU5Hr1G115900 | PLANT CADMIUM RESISTANCE 2 | 46.68 | AT1G14870.1 |
| 350 | HORVU7Hr1G074640 | lysine histidine transporter 1 | 46.66 | AT5G40780.1 |
| 351 | HORVU7Hr1G035980 | undescribed protein | 46.53 | AT5G22870.1 |
| 352 | HORVU7Hr1G091860 | L-tyrosine decarboxylase | 45.99 | AT2G20340.1 |
| 353 | HORVU1Hr1G014500 | undescribed protein | 45.94 | AT1G66730.1 |
| 354 | HORVU7Hr1G042120 | Protein NRT1/ PTR FAMILY 3.1 | 45.26 | AT1G68570.1 |
| 355 | HORVU3Hr1G018980 | basic helix-loop-helix (bHLH) DNA-binding family protein | 45.16 | AT5G56960.1 |
| 356 | HORVU4Hr1G012750 | Adenine/guanine permease AZG2 | 45.07 | AT5G50300.2 |
| 357 | HORVU7Hr1G076120 | Peroxidase superfamily protein | 44.90 | AT4G33420.1 |
| 358 | HORVU2Hr1G122620 | Blue copper protein | 44.75 | AT2G31050.1 |
| 359 | HORVU2Hr1G003210 | Cytochrome P450 superfamily protein | 44.45 | AT1G11680.1 |
| 360 | HORVU7Hr1G121950 | Retrotransposon protein, putative, unclassified | 44.28 | AT4G29090.1 |
| 361 | HORVU7Hr1G049580 | Protein kinase family protein | 44.20 | AT1G21230.1 |
| 362 | HORVU1Hr1G084720 | unknown function | 44.19 | AT3G20630.1 |
| 363 | HORVU3Hr1G090220 | Protein kinase superfamily protein | 44.17 | AT2G19130.1 |
| 364 | HORVU1Hr1G019500 | sucrose-phosphatase 1 | 44.05 | AT2G35840.4 |
| 365 | HORVU7Hr1G110840 | Nodulin-like / Major Facilitator Superfamily protein | 43.97 | AT2G39210.1 |
| 366 | HORVU7Hr1G116310 | Cytochrome P450 superfamily protein | 43.88 | AT3G14660.1 |

| | | | | |
|---|---|---|---|---|
| 367 | HORVU1Hr1G051580 | PLAC8 family protein | 43.87 | AT1G63830.3 |
| 368 | HORVU4Hr1G064070 | U-box domain-containing protein 21 | 43.67 | AT5G37490.1 |
| 369 | HORVU7Hr1G107670 | Two-component response regulator ORR21 | 43.59 | AT1G14600.1 |
| 370 | HORVU2Hr1G033610 | dihydroflavonol 4-reductase | 43.48 | AT4G27250.1 |
| 371 | HORVU0Hr1G021480 | Arabidopsis thaliana protein of unknown function (DUF821) | 43.42 | AT5G23850.1 |
| 372 | HORVU2Hr1G088660 | Caleosin-related family protein | 43.42 | AT4G26740.1 |
| 373 | HORVU4Hr1G074080 | undescribed protein | 43.34 | AT2G30620.2 |
| 374 | HORVU2Hr1G006100 | receptor kinase 2 | 43.32 | AT3G53810.1 |
| 375 | HORVU1Hr1G038780 | unknown protein; BEST Arabidopsis thaliana protein match is: unknown protein . | 42.86 | AT1G07040.1 |
| 376 | HORVU2Hr1G044270 | receptor kinase 1 | 42.81 | AT4G05200.2 |
| 377 | HORVU7Hr1G056250 | undescribed protein | 42.74 | AT4G28460.1 |
| 378 | HORVU2Hr1G076490 | unknown function | 42.57 | AT3G19380.1 |
| 379 | HORVU7Hr1G092800 | Tryptophan synthase beta chain 2 | 42.40 | AT5G38530.1 |
| 380 | HORVU1Hr1G003210 | beta-galactosidase 7 | 42.26 | AT5G20710.1 |
| 381 | HORVU5Hr1G124120 | gibberellin 20 oxidase 2 | 42.24 | AT4G25420.1 |
| 382 | HORVU2Hr1G121170 | Cytochrome P450 superfamily protein | 42.21 | AT3G61040.2 |
| 383 | HORVU4Hr1G057550 | pseudo-response regulator 7 | 42.06 | AT5G02810.1 |
| 384 | HORVU3Hr1G086680 | zinc-finger protein 3 | 41.91 | AT2G28710.1 |
| 385 | HORVU3Hr1G021140 | gigantea protein (GI) | 41.51 | AT1G22770.1 |
| 386 | HORVU6Hr1G073990 | early nodulin-like protein 19 | 41.50 | AT2G32300.2 |
| 387 | HORVU5Hr1G066460 | unknown protein; BEST Arabidopsis thaliana protein match is: unknown protein . | 41.43 | AT1G23110.7 |
| 388 | HORVU2Hr1G113180 | D-3-phosphoglycerate dehydrogenase | 41.32 | AT4G34200.1 |
| 389 | HORVU4Hr1G028780 | myb domain protein 112 | 41.31 | AT5G49620.1 |
| 390 | HORVU6Hr1G073160 | unknown function | 41.20 | AT2G18690.1 |
| 391 | HORVU6Hr1G004300 | unknown function | 41.03 | AT1G21240.2 |
| 392 | HORVU1Hr1G010210 | Bidirectional sugar transporter N3 | 40.89 | AT3G48740.1 |
| 393 | HORVU5Hr1G074030 | N-terminal protein myristoylation | 40.85 | AT5G66580.1 |
| 394 | HORVU2Hr1G113080 | Protein kinase family protein | 40.66 | AT1G16260.2 |
| 395 | HORVU1Hr1G045580 | undescribed protein | 40.02 | AT1G55020.1 |
| 396 | HORVU3Hr1G031020 | Cytochrome P450 superfamily protein | 39.76 | AT3G48320.1 |
| 397 | HORVU3Hr1G025810 | serine/threonine protein kinase 1 | 39.72 | AT5G45810.1 |
| 398 | HORVU7Hr1G028370 | Pheophorbide a oxygenase, chloroplastic | 39.63 | AT4G25650.1 |
| 399 | HORVU5Hr1G000910 | Serine/threonine-protein kinase | 39.43 | AT4G32300.1 |
| 400 | HORVU3Hr1G006760 | Chromosome 3B, genomic scaffold, cultivar Chinese Spring | 39.15 | AT3G14460.1 |
| 401 | HORVU3Hr1G006730 | Cysteine-rich receptor-like protein kinase 10 | 39.02 | AT4G23180.1 |
| 402 | HORVU3Hr1G024820 | Chromosome 3B, genomic scaffold, cultivar Chinese Spring | 38.82 | AT5G06570.4 |
| 403 | HORVU2Hr1G044440 | glucan endo-1,3-beta-glucosidase-like protein 3 | 38.63 | AT2G39640.1 |
| 404 | HORVU3Hr1G035680 | flowering promoting factor 1 | 38.38 | AT5G24860.1 |

| | | | | |
|---|---|---|---|---|
| 405 | HORVU3Hr1G001740 | Chromosome 3B, genomic scaffold, cultivar Chinese Spring | 38.37 | AT5G22050.1 |
| 406 | HORVU2Hr1G062730 | Protein NRT1/ PTR FAMILY 4.5 | 38.29 | AT1G27040.2 |
| 407 | HORVU5Hr1G122510 | Leucine-rich receptor-like protein kinase family protein | 38.23 | AT3G47570.1 |
| 408 | HORVU6Hr1G088420 | 2-oxoglutarate (2OG) and Fe(II)-dependent oxygenase superfamily protein | 38.09 | AT1G49390.1 |
| 409 | HORVU3Hr1G001990 | germin-like protein 9 | 37.71 | AT3G05950.1 |
| 410 | HORVU6Hr1G090600 | Glutathione S-transferase family protein | 37.50 | AT1G10370.1 |
| 411 | HORVU7Hr1G100810 | 2-oxoglutarate (2OG) and Fe(II)-dependent oxygenase superfamily protein | 37.35 | AT1G06620.1 |
| 412 | HORVU5Hr1G071140 | cellulose synthase like E1 | 37.23 | AT1G55850.1 |
| 413 | HORVU7Hr1G006940 | receptor kinase 1 | 37.12 | AT1G56130.2 |
| 414 | HORVU1Hr1G090820 | unknown function | 36.79 | AT2G25625.2 |
| 415 | HORVU2Hr1G083020 | Cytochrome P450 superfamily protein | 36.67 | AT3G48270.1 |
| 416 | HORVU2Hr1G076470 | U-box domain-containing protein | 36.58 | AT5G64660.1 |
| 417 | HORVU2Hr1G030660 | unknown function | 36.40 | AT5G42900.2 |
| 418 | HORVU2Hr1G004500 | Cytochrome P450 superfamily protein | 36.09 | AT2G30770.1 |
| 419 | HORVU7Hr1G024350 | Long-chain-fatty-acid--CoA ligase 1 | 36.05 | AT4G23850.1 |
| 420 | HORVU6Hr1G088960 | Cytochrome P450 superfamily protein | 35.93 | AT3G14690.1 |
| 421 | HORVU3Hr1G006800 | receptor kinase 2 | 35.83 | AT4G23280.1 |
| 422 | HORVU6Hr1G001420 | Protein kinase family protein | 35.75 | AT1G21230.1 |
| 423 | HORVU1Hr1G025530 | Lactoylglutathione lyase / glyoxalase I family protein | 35.63 | AT1G15380.2 |
| 424 | HORVU5Hr1G093090 | cationic amino acid transporter 5 | 35.62 | AT4G21120.1 |
| 425 | HORVU4Hr1G051450 | Phospholipid-transporting ATPase 1 | 35.45 | AT5G04930.1 |
| 426 | HORVU4Hr1G043960 | Eukaryotic aspartyl protease family protein | 35.41 | AT2G03200.1 |
| 427 | HORVU6Hr1G077940 | unknown function | 35.36 | AT1G66170.1 |
| 428 | HORVU3Hr1G028620 | Cytochrome P450 61 | 35.34 | AT2G34500.1 |
| 429 | HORVU2Hr1G096380 | Branched-chain-amino-acid aminotransferase 6 | 35.30 | AT5G65780.1 |
| 430 | HORVU5Hr1G066720 | Family of unknown function (DUF716) | 35.28 | AT1G55230.1 |
| 431 | HORVU7Hr1G035190 | RNA ligase/cyclic nucleotide phosphodiesterase family protein | 35.27 | AT4G18930.1 |
| 432 | HORVU3Hr1G069320 | WRKY DNA-binding protein 65 | 35.23 | AT1G29280.1 |
| 433 | HORVU7Hr1G040170 | Benzoate O-methyltransferase | 35.14 | AT3G21950.1 |
| 434 | HORVU6Hr1G069980 | receptor kinase 2 | 35.02 | AT5G06740.1 |
| 435 | HORVU6Hr1G074060 | Chalcone synthase A | 34.79 | AT5G13930.1 |
| 436 | HORVU4Hr1G063590 | Chaperone protein DnaJ | 34.66 | AT2G17880.1 |
| 437 | HORVU3Hr1G117480 | 2-oxoglutarate (2OG) and Fe(II)-dependent oxygenase superfamily protein | 34.29 | AT4G16765.6 |
| 438 | HORVU5Hr1G047730 | ferritin 4 | 34.27 | AT2G40300.1 |
| 439 | HORVU5Hr1G106000 | RING/U-box superfamily protein | 33.99 | AT3G05200.1 |
| 440 | HORVU1Hr1G072690 | Aldose reductase A | 33.85 | AT2G37790.1 |

| | | | | |
|---|---|---|---|---|
| 441 | HORVU6Hr1G088400 | Leucine-rich receptor-like protein kinase family protein | 33.76 | AT3G47570.1 |
| 442 | HORVU4Hr1G064480 | ethylene responsive element binding factor 5 | 33.34 | AT5G19790.1 |
| 443 | HORVU4Hr1G072880 | phosphoserine aminotransferase | 33.31 | AT4G35630.1 |
| 444 | HORVU3Hr1G018690 | receptor kinase 2 | 33.24 | AT2G37710.1 |
| 445 | HORVU2Hr1G008140 | receptor kinase 3 | 32.89 | AT4G27300.1 |
| 446 | HORVU3Hr1G021150 | gigantea protein (GI) | 32.87 | AT1G22770.1 |
| 447 | HORVU3Hr1G028020 | GDSL esterase/lipase | 32.76 | AT1G28570.2 |
| 448 | HORVU1Hr1G001570 | O-methyltransferase family protein | 32.76 | AT4G35160.1 |
| 449 | HORVU3Hr1G072210 | undescribed protein | 32.61 | AT3G63150.1 |
| 450 | HORVU3Hr1G094370 | Chromosome 3B, genomic scaffold, cultivar Chinese Spring | 32.59 | AT4G19975.1 |
| 451 | HORVU6Hr1G074370 | undescribed protein | 32.57 | AT1G44970.1 |
| 452 | HORVU6Hr1G077930 | UDP-Glycosyltransferase superfamily protein | 32.52 | AT1G22380.1 |
| 453 | HORVU3Hr1G056110 | Cytochrome P450 superfamily protein | 32.41 | AT3G14690.1 |
| 454 | HORVU3Hr1G087570 | 2-oxoglutarate (2OG) and Fe(II)-dependent oxygenase superfamily protein | 32.18 | AT5G05600.2 |
| 455 | HORVU6Hr1G053700 | undescribed protein | 31.97 | AT3G27473.1 |
| 456 | HORVU3Hr1G026050 | delta(3), delta(2)-enoyl CoA isomerase 1 | 31.68 | AT4G14440.1 |
| 457 | HORVU4Hr1G051290 | NAC domain protein, | 31.53 | AT1G69490.1 |
| 458 | HORVU3Hr1G011850 | Cytochrome P450 superfamily protein | 31.42 | AT3G48270.1 |
| 459 | HORVU1Hr1G015090 | tyrosine aminotransferase 3 | 31.40 | AT5G53970.1 |
| 460 | HORVU4Hr1G001040 | Leucine-rich repeat receptor-like protein kinase family protein | 31.39 | AT2G34930.1 |
| 461 | HORVU2Hr1G045350 | Syntaxin-132 | 31.16 | AT3G03800.2 |
| 462 | HORVU4Hr1G078370 | undescribed protein | 30.95 | AT1G78880.1 |
| 463 | HORVU7Hr1G101300 | undescribed protein | 30.95 | AT5G04220.2 |
| 464 | HORVU1Hr1G016870 | Peroxidase superfamily protein | 30.94 | AT1G05260.1 |
| 465 | HORVU3Hr1G109510 | undescribed protein | 30.93 | AT4G20020.2 |
| 466 | HORVU7Hr1G113290 | unknown function | 30.82 | AT2G38465.1 |
| 467 | HORVU3Hr1G024660 | Acetyl esterase | 30.69 | AT5G06570.4 |
| 468 | HORVU2Hr1G109440 | Cytochrome P450 superfamily protein | 30.44 | AT4G36220.1 |
| 469 | HORVU6Hr1G083050 | lysine-ketoglutarate reductase/saccharopine dehydrogenase bifunctional enzyme | 30.34 | AT4G33150.3 |
| 470 | HORVU1Hr1G092680 | FAD-binding Berberine family protein | 30.30 | AT5G44390.1 |
| 471 | HORVU1Hr1G090730 | 3-ketoacyl-CoA synthase 11 | 30.14 | AT2G26640.1 |
| 472 | HORVU2Hr1G018260 | serine/threonine protein kinase 1 | 30.13 | AT5G58380.1 |
| 473 | HORVU1Hr1G016840 | Peroxidase superfamily protein | 30.06 | AT1G05260.1 |
| 474 | HORVU5Hr1G096210 | UDP-Glycosyltransferase superfamily protein | 29.98 | AT3G02100.1 |
| 475 | HORVU7Hr1G024940 | Subtilisin-like protease | 29.81 | AT4G10520.1 |
| 476 | HORVU3Hr1G002910 | 0 | 29.62 | #N/A |
| 477 | HORVU5Hr1G086670 | Flavin-binding monooxygenase family protein | 29.45 | AT1G19250.1 |
| 478 | HORVU3Hr1G064680 | Protein transport protein Sec61 subunit alpha | 29.44 | AT2G34250.3 |

| | | | | |
|---|---|---|---|---|
| 479 | HORVU3Hr1G108100 | carotenoid cleavage dioxygenase 1 | 29.42 | AT3G63520.1 |
| 480 | HORVU3Hr1G020210 | WRKY DNA-binding protein 72 | 29.14 | AT5G15130.1 |
| 481 | HORVU2Hr1G044890 | receptor kinase 3 | 29.07 | AT4G05200.1 |
| 482 | HORVU4Hr1G056240 | Asparagine synthetase [glutamine-hydrolyzing] 1 | 29.05 | AT5G65010.1 |
| 483 | HORVU2Hr1G084130 | malate synthase | 29.03 | AT5G03860.2 |
| 484 | HORVU6Hr1G079950 | undescribed protein | 28.99 | #N/A |
| 485 | HORVU7Hr1G021260 | WRKY DNA-binding protein 61 | 28.99 | AT5G15130.1 |
| 486 | HORVU1Hr1G058520 | beta glucosidase 11 | 28.96 | AT1G02850.5 |
| 487 | HORVU5Hr1G122060 | Basic blue protein | 28.92 | AT2G02850.1 |
| 488 | HORVU5Hr1G014130 | Tuliposide A-converting enzyme b3, amyloplastic | 28.89 | AT1G47480.1 |
| 489 | HORVU4Hr1G066200 | Non-symbiotic hemoglobin | 28.75 | AT2G16060.1 |
| 490 | HORVU5Hr1G050940 | cinnamoyl coa reductase | 28.73 | AT1G15950.3 |
| 491 | HORVU2Hr1G044450 | Glucan endo-1,3-beta-glucosidase 3 | 28.70 | AT5G55180.2 |
| 492 | HORVU7Hr1G098810 | Heat shock 70 kDa protein C | 28.57 | AT5G42020.1 |
| 493 | HORVU7Hr1G106160 | Protein NRT1/ PTR FAMILY 5.1 | 28.56 | AT2G40460.1 |
| 494 | HORVU3Hr1G007590 | Protein kinase superfamily protein | 28.44 | AT1G18390.2 |
| 495 | HORVU3Hr1G085890 | ABC transporter C family member 3 | 28.43 | AT3G13080.4 |
| 496 | HORVU7Hr1G114790 | Laccase | 28.36 | AT3G09510.1 |
| 497 | HORVU0Hr1G023570 | HXXXD-type acyl-transferase family protein | 28.31 | AT5G48930.1 |
| 498 | HORVU3Hr1G018740 | sugar transporter 1 | 28.19 | AT5G61520.1 |
| 499 | HORVU2Hr1G123590 | Protein kinase family protein | 28.11 | AT1G21210.1 |
| 500 | HORVU2Hr1G042550 | receptor kinase 1 | 28.04 | AT4G23180.2 |
| 501 | HORVU6Hr1G073510 | Potassium transporter family protein | 27.98 | AT4G13420.1 |
| 502 | HORVU1Hr1G065630 | undescribed protein | 27.84 | AT5G50930.1 |
| 503 | HORVU4Hr1G022270 | Peroxidase superfamily protein | 27.80 | AT5G05340.1 |
| 504 | HORVU2Hr1G104090 | undescribed protein | 27.80 | AT1G16630.1 |
| 505 | HORVU1Hr1G013480 | sugar transporter 1 | 27.75 | AT1G34580.1 |
| 506 | HORVU2Hr1G027480 | Cytochrome P450 superfamily protein | 27.75 | AT2G46950.1 |
| 507 | HORVU3Hr1G109590 | Glycosyltransferase family 61 protein | 27.52 | AT2G41640.1 |
| 508 | HORVU2Hr1G126120 | Protein NRT1/ PTR FAMILY 5.10 | 27.46 | AT1G22570.2 |
| 509 | HORVU2Hr1G019790 | Rubber elongation factor, putative isoform 2 | 27.40 | AT2G47780.1 |
| 510 | HORVU4Hr1G061120 | Anthranilate synthase component 1 | 27.10 | AT5G05730.1 |
| 511 | HORVU1Hr1G090460 | undescribed protein | 27.10 | AT1G78410.2 |
| 512 | HORVU7Hr1G107640 | 0 | 27.08 | #N/A |
| 513 | HORVU5Hr1G115310 | Mannan endo-1,4-beta-mannosidase 4 | 27.03 | AT5G66460.1 |
| 514 | HORVU4Hr1G089030 | Protein of unknown function, DUF584 | 27.00 | AT3G15040.1 |
| 515 | HORVU7Hr1G028750 | F-box domain containing protein, expressed | 26.80 | AT3G06240.1 |
| 516 | HORVU2Hr1G092440 | undescribed protein | 26.71 | AT2G32240.1 |
| 517 | HORVU2Hr1G093660 | Ethylene-responsive transcription factor 1B | 26.67 | AT3G23240.1 |
| 518 | HORVU5Hr1G061810 | Protein kinase superfamily protein | 26.60 | AT2G05940.1 |

| | | | | |
|---|---|---|---|---|
| 519 | HORVU1Hr1G091030 | Auxin efflux carrier family protein | 26.56 | AT1G70940.1 |
| 520 | HORVU6Hr1G008730 | catalase 3 | 26.56 | AT1G20620.4 |
| 521 | HORVU2Hr1G083010 | undescribed protein | 26.53 | AT4G12120.1 |
| 522 | HORVU6Hr1G089560 | U-box domain-containing protein | 26.51 | AT2G35930.1 |
| 523 | HORVU4Hr1G088930 | P-loop containing nucleoside triphosphate hydrolases superfamily protein | 26.39 | AT2G46620.1 |
| 524 | HORVU6Hr1G092020 | Leucine-rich repeat receptor-like protein kinase family protein | 26.36 | AT5G20480.2 |
| 525 | HORVU3Hr1G096860 | Potassium transporter family protein | 26.31 | AT4G13420.1 |
| 526 | HORVU7Hr1G001520 | undescribed protein | 26.20 | AT4G00300.1 |
| 527 | HORVU0Hr1G021610 | F-box protein PP2-B7 | 26.03 | AT2G02360.1 |
| 528 | HORVU1Hr1G071390 | unknown function | 26.03 | AT1G74540.1 |
| 529 | HORVU6Hr1G067910 | Protein NRT1/ PTR FAMILY 8.3 | 25.90 | AT2G02040.1 |
| 530 | HORVU3Hr1G053350 | ABC transporter G family member 36 | 25.72 | AT1G15520.1 |
| 531 | HORVU4Hr1G003450 | undescribed protein | 25.67 | AT1G22680.1 |
| 532 | HORVU5Hr1G106850 | ABC transporter G family member 48 | 25.45 | AT1G66950.1 |
| 533 | HORVU2Hr1G042540 | unknown function | 25.43 | AT1G32660.1 |
| 534 | HORVU5Hr1G061580 | Secretory carrier-associated membrane protein 4 | 25.38 | AT2G20840.1 |
| 535 | HORVU7Hr1G117850 | O-methyltransferase family protein | 25.37 | AT4G35160.1 |
| 536 | HORVU3Hr1G068700 | Ribose-phosphate pyrophosphokinase 4 | 25.30 | AT1G10700.1 |
| 537 | HORVU6Hr1G077120 | U-box domain-containing protein 21 | 25.30 | AT5G37490.1 |
| 538 | HORVU3Hr1G105920 | Amino-acid permease BAT1 homolog | 25.13 | AT2G01170.1 |
| 539 | HORVU3Hr1G057180 | unknown function | 25.10 | AT5G62770.1 |
| 540 | HORVU2Hr1G122580 | Blue copper protein | 24.96 | AT2G32300.1 |
| 541 | HORVU2Hr1G046370 | 18.8 kDa class V heat shock protein | 24.85 | AT4G21870.1 |
| 542 | HORVU6Hr1G001370 | undescribed protein | 24.84 | #N/A |
| 543 | HORVU2Hr1G080890 | Anthranilate synthase | 24.82 | AT1G25220.1 |
| 544 | HORVU7Hr1G101310 | Calcium-dependent lipid-binding (CaLB domain) family protein | 24.81 | AT5G04220.2 |
| 545 | HORVU2Hr1G124930 | Peroxidase superfamily protein | 24.81 | AT1G71695.1 |
| 546 | HORVU6Hr1G004310 | unknown function | 24.70 | AT1G12120.1 |
| 547 | HORVU6Hr1G074310 | undescribed protein | 24.63 | AT3G57020.2 |
| 548 | HORVU7Hr1G026640 | FAR1-related sequence 5 | 24.58 | AT4G38180.1 |
| 549 | HORVU7Hr1G084610 | 3-ketoacyl-CoA synthase 11 | 24.49 | AT2G26640.1 |
| 550 | HORVU5Hr1G070400 | ABC transporter G family member 11 | 24.38 | AT1G17840.1 |
| 551 | HORVU6Hr1G067980 | Protein NRT1/ PTR FAMILY 8.3 | 24.32 | AT2G02040.1 |
| 552 | HORVU2Hr1G061670 | undescribed protein | 24.27 | AT2G26170.1 |
| 553 | HORVU7Hr1G107150 | undescribed protein | 24.25 | AT2G29690.2 |
| 554 | HORVU3Hr1G091120 | amino acid permease 3 | 24.24 | AT5G09220.1 |
| 555 | HORVU1Hr1G077420 | Inositol-tetrakisphosphate 1-kinase 4 | 24.19 | AT5G16760.1 |
| 556 | HORVU6Hr1G089590 | U-box domain-containing protein | 23.94 | AT2G35930.1 |
| 557 | HORVU6Hr1G057310 | undescribed protein | 23.88 | AT1G24062.1 |

| | | | | |
|---|---|---|---|---|
| 558 | HORVU2Hr1G004510 | Cytochrome P450 superfamily protein | 23.84 | AT5G25120.1 |
| 559 | HORVU5Hr1G089840 | receptor kinase 3 | 23.77 | AT1G21230.1 |
| 560 | HORVU3Hr1G070550 | Transferrin receptor protein 2 | 23.76 | AT5G19740.1 |
| 561 | HORVU4Hr1G009230 | receptor kinase 2 | 23.25 | AT5G10530.1 |
| 562 | HORVU5Hr1G061820 | Protein kinase superfamily protein | 23.24 | AT2G05940.1 |
| 563 | HORVU3Hr1G074940 | Peroxidase superfamily protein | 23.14 | AT4G11290.1 |
| 564 | HORVU7Hr1G040850 | unknown function | 23.04 | AT5G18520.1 |
| 565 | HORVU2Hr1G109330 | WRKY family transcription factor | 22.92 | AT3G56400.1 |
| 566 | HORVU3Hr1G020140 | WRKY DNA-binding protein 50 | 22.86 | AT5G26170.1 |
| 567 | HORVU5Hr1G080300 | Dehydration-responsive element-binding protein 1B | 22.50 | AT4G25470.1 |
| 568 | HORVU1Hr1G090870 | 2-oxoglutarate (2OG) and Fe(II)-dependent oxygenase superfamily protein | 22.49 | AT1G52820.1 |
| 569 | HORVU5Hr1G121520 | unknown function | 22.44 | AT5G18980.1 |
| 570 | HORVU3Hr1G018610 | receptor kinase 2 | 22.37 | AT3G53810.1 |
| 571 | HORVU4Hr1G090140 | germin-like protein 9 | 22.22 | AT5G39150.1 |
| 572 | HORVU1Hr1G082670 | AP2/B3 transcription factor family protein | 22.21 | AT1G68840.2 |
| 573 | HORVU6Hr1G063480 | cysteine synthase D1 | 22.21 | AT2G43750.2 |
| 574 | HORVU6Hr1G022590 | RING/U-box superfamily protein | 22.20 | AT3G05200.1 |
| 575 | HORVU3Hr1G077890 | Chromosome 3B, genomic scaffold, cultivar Chinese Spring | 22.11 | AT4G21920.1 |
| 576 | HORVU5Hr1G094710 | Disease resistance-responsive (dirigent-like protein) family protein | 22.05 | AT1G58170.1 |
| 577 | HORVU3Hr1G064960 | Protein kinase superfamily protein | 22.03 | AT5G60900.1 |
| 578 | HORVU0Hr1G000780 | unknown function | 21.91 | AT5G24275.4 |
| 579 | HORVU2Hr1G100360 | SAUR-like auxin-responsive protein family | 21.89 | AT2G24400.1 |
| 580 | HORVU7Hr1G049610 | Sterile alpha motif (SAM) domain-containing protein | 21.89 | AT1G80520.1 |
| 581 | HORVU5Hr1G054510 | BZIP transcription factor | 21.86 | AT1G75390.1 |
| 582 | HORVU3Hr1G105680 | 0 | 21.84 | #N/A |
| 583 | HORVU1Hr1G093480 | tryptophan synthase alpha chain | 21.82 | AT4G02610.1 |
| 584 | HORVU5Hr1G006460 | unknown function | 21.81 | AT2G38470.1 |
| 585 | HORVU1Hr1G019410 | HXXXD-type acyl-transferase family protein | 21.78 | AT5G48930.1 |
| 586 | HORVU5Hr1G123330 | Cysteine/Histidine-rich C1 domain family protein | 21.72 | AT2G37810.1 |
| 587 | HORVU3Hr1G096890 | Avr9/Cf-9 rapidly elicited protein | 21.62 | AT4G29110.1 |
| 588 | HORVU3Hr1G087690 | ammonium transporter 2 | 21.60 | AT2G38290.1 |
| 589 | HORVU1Hr1G057880 | Eukaryotic peptide chain release factor subunit 1-2 | 21.47 | AT1G12920.1 |
| 590 | HORVU1Hr1G057650 | rRNA N-glycosidase | 21.38 | AT5G33406.1 |
| 591 | HORVU6Hr1G020570 | ATP-dependent RNA helicase DeaD | 21.32 | AT5G60990.1 |
| 592 | HORVU7Hr1G059520 | undescribed protein | 21.30 | AT4G31210.1 |
| 593 | HORVU0Hr1G000760 | undescribed protein | 21.20 | AT4G19140.1 |
| 594 | HORVU1Hr1G016660 | Peroxidase superfamily protein | 21.19 | AT4G33420.1 |

| | | | | |
|---|---|---|---|---|
| 595 | HORVU6Hr1G058820 | phenylalanine ammonia-lyase 2 | 21.18 | AT2G37040.1 |
| 596 | HORVU4Hr1G056500 | 2-oxoglutarate (2OG) and Fe(II)-dependent oxygenase superfamily protein | 21.16 | AT3G11180.1 |
| 597 | HORVU2Hr1G120010 | Chaperone protein DnaJ | 21.11 | AT5G23240.1 |
| 598 | HORVU7Hr1G096870 | SAUR-like auxin-responsive protein family | 21.08 | AT3G20220.1 |
| 599 | HORVU7Hr1G107630 | Homeodomain-like superfamily protein | 20.94 | AT1G14600.1 |
| 600 | HORVU7Hr1G083270 | WRKY DNA-binding protein 70 | 20.91 | AT3G56400.1 |
| 601 | HORVU5Hr1G065620 | 2-oxoglutarate (2OG) and Fe(II)-dependent oxygenase superfamily protein | 20.81 | AT5G20400.1 |
| 602 | HORVU2Hr1G045200 | Glutathione S-transferase family protein | 20.78 | AT1G10370.1 |
| 603 | HORVU6Hr1G088430 | 2-oxoglutarate (2OG) and Fe(II)-dependent oxygenase superfamily protein | 20.72 | AT1G49390.1 |
| 604 | HORVU2Hr1G098700 | undescribed protein | 20.68 | #N/A |
| 605 | HORVU5Hr1G006480 | undescribed protein | 20.62 | AT5G66750.1 |
| 606 | HORVU0Hr1G009460 | germin-like protein 9 | 20.50 | AT5G39110.1 |
| 607 | HORVU4Hr1G089560 | Chromosome 3B, genomic scaffold, cultivar Chinese Spring | 20.49 | AT5G42900.2 |
| 608 | HORVU2Hr1G081510 | unknown function | 20.18 | AT1G55775.1 |
| 609 | HORVU2Hr1G070880 | Protein TIFY 9 | 20.17 | AT5G13220.1 |
| 610 | HORVU7Hr1G033430 | Chaperone DnaJ domain protein | 20.16 | AT3G14200.1 |
| 611 | HORVU7Hr1G010340 | Anthocyanin 5-aromatic acyltransferase-like | 19.96 | AT5G39080.1 |
| 612 | HORVU2Hr1G094330 | GDSL esterase/lipase | 19.96 | AT5G41890.1 |
| 613 | HORVU5Hr1G045850 | Glutathione S-transferase family protein | 19.63 | AT2G29420.1 |
| 614 | HORVU7Hr1G002850 | 26S protease regulatory subunit 6A homolog | 19.56 | AT3G01610.2 |
| 615 | HORVU2Hr1G091190 | sugar transporter 9 | 19.51 | AT2G20780.1 |
| 616 | HORVU2Hr1G017350 | NAC domain containing protein 100 | 19.43 | AT5G61430.1 |
| 617 | HORVU5Hr1G034300 | undescribed protein | 19.40 | AT5G10140.4 |
| 618 | HORVU2Hr1G126100 | Protein NRT1/ PTR FAMILY 5.10 | 19.36 | AT1G72125.1 |
| 619 | HORVU6Hr1G067920 | Protein NRT1/ PTR FAMILY 8.1 | 19.22 | AT1G62200.1 |
| 620 | HORVU7Hr1G028660 | unknown function | 19.15 | AT3G17240.3 |
| 621 | HORVU0Hr1G006150 | receptor kinase 1 | 19.14 | AT5G38240.2 |
| 622 | HORVU2Hr1G070700 | Glycerophosphodiester phosphodiesterase GDE1 | 19.09 | AT5G41080.2 |
| 623 | HORVU5Hr1G060620 | Centromere protein V | 18.94 | AT5G16940.2 |
| 624 | HORVU4Hr1G000700 | Ethylene-responsive transcription factor 1B | 18.85 | AT3G23240.1 |
| 625 | HORVU2Hr1G103930 | NAC domain containing protein 1 | 18.84 | AT3G12977.1 |
| 626 | HORVU5Hr1G045580 | mini zinc finger 2 | 18.81 | AT3G28917.1 |
| 627 | HORVU4Hr1G039300 | Kelch repeat-containing protein 3 | 18.80 | AT1G68050.1 |
| 628 | HORVU5Hr1G057320 | LAG1 longevity assurance homolog 3 | 18.80 | AT1G13580.3 |
| 629 | HORVU5Hr1G094700 | Disease resistance-responsive (dirigent-like protein) family protein | 18.78 | AT1G58170.1 |
| 630 | HORVU5Hr1G093350 | Solute carrier family 22 member 1 | 18.66 | AT3G13050.4 |
| 631 | HORVU1Hr1G092310 | Glucan endo-1,3-beta-glucosidase 13 | 18.57 | AT4G26830.1 |
| 632 | HORVU1Hr1G080300 | WRKY DNA-binding protein 51 | 18.54 | AT5G64810.1 |

| | | | | |
|---|---|---|---|---|
| 633 | HORVU1Hr1G015210 | Leucine-rich repeat receptor-like protein kinase family protein | 18.50 | AT4G08850.1 |
| 634 | HORVU4Hr1G083650 | Protein NRT1/ PTR FAMILY 5.2 | 18.47 | AT5G46050.1 |
| 635 | HORVU7Hr1G076480 | Non-lysosomal glucosylceramidase | 18.42 | AT5G49900.3 |
| 636 | HORVU1Hr1G085240 | Core-2/I-branching beta-1,6-N-acetylglucosaminyltransferase family protein | 18.39 | AT5G11730.1 |
| 637 | HORVU3Hr1G062920 | Chromosome 3B, genomic scaffold, cultivar Chinese Spring | 18.37 | AT3G49210.2 |
| 638 | HORVU2Hr1G082370 | 2-oxoglutarate (2OG) and Fe(II)-dependent oxygenase superfamily protein | 18.34 | AT1G14130.1 |
| 639 | HORVU3Hr1G077950 | Subtilisin-like protease | 18.24 | AT4G10540.1 |
| 640 | HORVU4Hr1G087630 | Cytochrome P450 superfamily protein | 18.21 | AT2G30490.1 |
| 641 | HORVU0Hr1G000340 | cystathionine beta-lyase | 18.11 | AT1G64660.1 |
| 642 | HORVU5Hr1G050970 | unknown protein; BEST Arabidopsis thaliana protein match is: unknown protein . | 18.07 | AT2G24100.1 |
| 643 | HORVU3Hr1G028870 | undescribed protein | 18.01 | AT4G19510.9 |
| 644 | HORVU5Hr1G063430 | glutamate receptor 2.8 | 17.99 | AT2G29120.1 |
| 645 | HORVU5Hr1G104670 | Glutathione S-transferase family protein | 17.91 | AT3G62760.1 |
| 646 | HORVU2Hr1G028430 | ribonuclease 1 | 17.80 | AT2G02990.1 |
| 647 | HORVU6Hr1G011120 | Glutathione S-transferase family protein | 17.75 | AT4G19880.1 |
| 648 | HORVU3Hr1G077110 | receptor kinase 2 | 17.52 | AT4G21390.1 |
| 649 | HORVU3Hr1G013650 | Gamma-glutamyltranspeptidase 3 | 17.36 | AT4G29210.1 |
| 650 | HORVU1Hr1G075640 | ABC transporter ATP-binding protein NatA | 17.36 | AT3G47730.1 |
| 651 | HORVU0Hr1G015930 | Ergosterol biosynthetic protein 28 | 17.31 | AT1G10030.1 |
| 652 | HORVU5Hr1G081500 | Protein STAY-GREEN, chloroplastic | 17.29 | AT4G22920.2 |
| 653 | HORVU3Hr1G057730 | unknown protein; Has 19 Blast hits to 19 proteins in 6 species: Archae - 0; Bacteria - 0; Metazoa - 0; Fungi - 0; Plants - 19; Viruses - 0; Other Eukaryotes - 0 (source: NCBI BLink). | 17.11 | AT2G01400.1 |
| 654 | HORVU4Hr1G078210 | purine permease 3 | 16.98 | AT1G28220.1 |
| 655 | HORVU2Hr1G099640 | VACUOLAR SORTING RECEPTOR 5 | 16.93 | AT2G34940.1 |
| 656 | HORVU6Hr1G014340 | undescribed protein | 16.89 | AT2G40660.1 |
| 657 | HORVU5Hr1G097940 | undescribed protein | 16.88 | AT2G15960.1 |
| 658 | HORVU1Hr1G026200 | Chromosome 3B, genomic scaffold, cultivar Chinese Spring | 16.86 | AT5G04160.1 |
| 659 | HORVU1Hr1G072720 | unknown protein; Has 30 Blast hits to 30 proteins in 10 species: Archae - 0; Bacteria - 0; Metazoa - 0; Fungi - 0; Plants - 30; Viruses - 0; Other Eukaryotes - 0 (source: NCBI BLink). | 16.84 | AT5G02090.1 |
| 660 | HORVU5Hr1G006330 | Glutathione S-transferase family protein | 16.80 | AT3G62760.1 |
| 661 | HORVU3Hr1G089900 | undescribed protein | 16.77 | AT3G32090.1 |
| 662 | HORVU2Hr1G117840 | Protein kinase superfamily protein | 16.69 | AT4G21390.1 |
| 663 | HORVU2Hr1G117620 | undescribed protein | 16.67 | #N/A |
| 664 | HORVU5Hr1G058020 | Major facilitator superfamily protein | 16.66 | AT5G10190.2 |
| 665 | HORVU7Hr1G083550 | Peroxidase superfamily protein | 16.63 | AT5G66390.1 |

| | | | | |
|---|---|---|---|---|
| 666 | HORVU2Hr1G108290 | Solute carrier family 22 member 15-like | 16.60 | AT3G20660.1 |
| 667 | HORVU7Hr1G106760 | unknown function | 16.51 | AT5G16610.2 |
| 668 | HORVU7Hr1G083370 | Alpha/beta hydrolase | 16.50 | AT5G13800.3 |
| 669 | HORVU6Hr1G078110 | UDP-Glycosyltransferase superfamily protein | 16.31 | AT1G22400.1 |
| 670 | HORVU7Hr1G032740 | unknown function | 16.27 | AT3G16670.1 |
| 671 | HORVU4Hr1G012530 | Plant protein of unknown function (DUF863) | 16.23 | AT5G67390.3 |
| 672 | HORVU2Hr1G013170 | 2-oxoglutarate dehydrogenase, E1 component | 16.21 | AT3G55410.2 |
| 673 | HORVU2Hr1G017360 | NAC domain containing protein 100 | 16.20 | AT5G61430.1 |
| 674 | HORVU1Hr1G070220 | alanine:glyoxylate aminotransferase 3 | 16.19 | AT3G08860.1 |
| 675 | HORVU6Hr1G070740 | undescribed protein | 16.19 | AT2G41905.1 |
| 676 | HORVU3Hr1G093550 | receptor kinase 2 | 16.11 | AT1G56130.2 |
| 677 | HORVU6Hr1G077280 | Armadillo repeat-containing protein 6 | 16.08 | AT4G33945.1 |
| 678 | HORVU4Hr1G021040 | exocyst subunit exo70 family protein A2 | 15.99 | AT5G52340.1 |
| 679 | HORVU3Hr1G090170 | tropinone reductase | 15.98 | AT5G06060.1 |
| 680 | HORVU7Hr1G028730 | receptor kinase 2 | 15.89 | AT2G37710.1 |
| 681 | HORVU5Hr1G048810 | Coffea canephora DH200=94 genomic scaffold, scaffold_93 | 15.87 | AT2G17710.1 |
| 682 | HORVU1Hr1G052250 | undescribed protein | 15.84 | AT3G54710.1 |
| 683 | HORVU0Hr1G039620 | Receptor-like protein kinase | 15.73 | AT1G21230.1 |
| 684 | HORVU7Hr1G118560 | Blue copper protein | 15.67 | AT3G17675.1 |
| 685 | HORVU5Hr1G085780 | SAUR-like auxin-responsive protein family | 15.63 | AT1G29450.2 |
| 686 | HORVU3Hr1G065320 | ABC transporter B family member 4 | 15.36 | AT2G47000.7 |
| 687 | HORVU2Hr1G101980 | Ubiquinol oxidase | 15.32 | AT3G22370.1 |
| 688 | HORVU6Hr1G090630 | alpha/beta-Hydrolases superfamily protein | 15.20 | AT5G06570.2 |
| 689 | HORVU1Hr1G075620 | ABC transporter ATP-binding protein NatA | 15.15 | AT3G47780.1 |
| 690 | HORVU1Hr1G002590 | receptor-like protein kinase 4 | 15.09 | AT1G66920.2 |
| 691 | HORVU5Hr1G115780 | Epoxide hydrolase 3 | 15.03 | AT4G02340.1 |
| 692 | HORVU2Hr1G044650 | receptor kinase 1 | 15.02 | AT4G23180.1 |
| 693 | HORVU2Hr1G042520 | Protein kinase superfamily protein | 14.97 | AT4G21390.1 |
| 694 | HORVU3Hr1G051820 | Protein kinase superfamily protein | 14.94 | AT2G05940.1 |
| 695 | HORVU3Hr1G063430 | MATE efflux family protein | 14.76 | AT1G71140.1 |
| 696 | HORVU7Hr1G048570 | Glycosyltransferase family 61 protein | 14.76 | AT3G18180.1 |
| 697 | HORVU3Hr1G064450 | lipid phosphate phosphatase 2 | 14.74 | AT3G02600.9 |
| 698 | HORVU2Hr1G017470 | NAC domain protein, | 14.74 | AT1G69490.1 |
| 699 | HORVU2Hr1G109590 | HXXXD-type acyl-transferase family protein | 14.70 | AT5G48930.1 |
| 700 | HORVU1Hr1G002900 | unknown function | 14.67 | AT3G14470.2 |
| 701 | HORVU3Hr1G066220 | ABC transporter G family member 19 | 14.66 | AT3G55130.1 |
| 702 | HORVU5Hr1G061770 | Cathepsin B-like cysteine proteinase | 14.66 | AT5G60360.1 |
| 703 | HORVU1Hr1G059130 | Protein phosphatase 2C family protein | 14.59 | AT3G05640.3 |
| 704 | HORVU0Hr1G007680 | inositol polyphosphate 5-phosphatase 11 | 14.56 | AT1G47510.1 |
| 705 | HORVU5Hr1G063480 | glutamate receptor 2.8 | 14.51 | AT2G29120.1 |
| 706 | HORVU5Hr1G023990 | unknown function | 14.47 | AT1G67120.1 |

| | | | | |
|---|---|---|---|---|
| 707 | HORVU2Hr1G112580 | endonuclease 4 | 14.43 | AT4G21585.4 |
| 708 | HORVU5Hr1G057740 | unknown function | 14.43 | AT2G26110.1 |
| 709 | HORVU1Hr1G069800 | Non-specific lipid-transfer protein | 14.38 | AT5G01870.1 |
| 710 | HORVU5Hr1G075470 | Peptidyl-prolyl cis-trans isomerase | 14.26 | AT4G14200.1 |
| 711 | HORVU4Hr1G043910 | Protein disulfide-isomerase | 14.10 | AT1G21750.1 |
| 712 | HORVU6Hr1G067380 | myb domain protein 63 | 14.09 | AT1G79180.1 |
| 713 | HORVU3Hr1G090070 | undescribed protein | 14.07 | AT1G20400.1 |
| 714 | HORVU4Hr1G007180 | microsomal glutathione s-transferase, putative | 14.06 | AT1G65820.1 |
| 715 | HORVU6Hr1G090560 | Glutathione S-transferase family protein | 13.95 | AT1G10360.1 |
| 716 | HORVU2Hr1G125600 | unknown function | 13.89 | AT4G35920.5 |
| 717 | HORVU7Hr1G057650 | RING/U-box superfamily protein | 13.81 | AT2G17730.1 |
| 718 | HORVU4Hr1G083210 | Anthranilate phosphoribosyltransferase | 13.79 | AT5G17990.1 |
| 719 | HORVU4Hr1G023390 | unknown protein; BEST Arabidopsis thaliana protein match is: unknown protein . | 13.77 | AT3G48860.2 |
| 720 | HORVU1Hr1G026520 | HXXXD-type acyl-transferase family protein | 13.76 | AT3G62160.1 |
| 721 | HORVU2Hr1G098110 | nudix hydrolase homolog 8 | 13.76 | AT5G47240.1 |
| 722 | HORVU7Hr1G108820 | unknown function | 13.70 | AT2G18680.1 |
| 723 | HORVU6Hr1G009360 | Peroxidase superfamily protein | 13.69 | AT5G05340.1 |
| 724 | HORVU4Hr1G038810 | undescribed protein | 13.59 | AT2G34140.1 |
| 725 | HORVU2Hr1G000120 | undescribed protein | 13.48 | AT1G59960.1 |
| 726 | HORVU1Hr1G049400 | undescribed protein | 13.45 | AT4G25990.3 |
| 727 | HORVU4Hr1G072560 | Blue copper protein | 13.43 | AT2G32300.2 |
| 728 | HORVU5Hr1G106640 | rRNA N-glycosidase | 13.43 | AT5G07280.1 |
| 729 | HORVU2Hr1G075970 | Peroxidase superfamily protein | 13.42 | AT4G08390.3 |
| 730 | HORVU6Hr1G000070 | Rhodanese-like domain-containing protein 17 | 13.35 | AT5G66170.3 |
| 731 | HORVU7Hr1G077470 | GDP-mannose transporter 1 | 13.22 | AT2G13650.3 |
| 732 | HORVU6Hr1G013610 | beta-1,4-N-acetylglucosaminyltransferase family protein | 13.10 | AT2G13290.2 |
| 733 | HORVU5Hr1G019880 | unknown function | 13.09 | AT5G39380.5 |
| 734 | HORVU1Hr1G083700 | Galactosyltransferase family protein | 13.04 | AT5G57500.1 |
| 735 | HORVU6Hr1G005400 | Epoxide hydrolase 4 | 12.96 | AT3G51000.1 |
| 736 | HORVU3Hr1G028860 | cationic amino acid transporter 5 | 12.93 | AT2G34960.1 |
| 737 | HORVU6Hr1G028590 | undescribed protein | 12.78 | AT3G01430.1 |
| 738 | HORVU5Hr1G084740 | phospholipase D delta | 12.61 | AT4G35790.3 |
| 739 | HORVU4Hr1G076970 | Cyanate hydratase | 12.54 | AT3G23490.3 |
| 740 | HORVU7Hr1G079160 | Chromosome 3B, genomic scaffold, cultivar Chinese Spring | 12.47 | AT2G29220.1 |
| 741 | HORVU6Hr1G067740 | Protein NRT1/ PTR FAMILY 8.3 | 12.47 | AT2G02040.1 |
| 742 | HORVU7Hr1G048970 | caffeoyl-CoA 3-O-methyltransferase | 12.47 | AT4G34050.1 |
| 743 | HORVU2Hr1G080140 | Aldose 1-epimerase | 12.34 | AT5G15140.1 |
| 744 | HORVU4Hr1G071020 | sugar transporter 9 | 12.33 | AT3G18830.1 |
| 745 | HORVU0Hr1G007750 | 0 | 12.31 | #N/A |

| | | | | |
|---|---|---|---|---|
| 746 | HORVU2Hr1G064150 | sugar transporter 1 | 12.14 | AT1G11260.1 |
| 747 | HORVU4Hr1G062440 | Phosphoenolpyruvate carboxykinase [ATP] | 12.08 | AT4G37870.1 |
| 748 | HORVU7Hr1G038370 | Beta-1,3-galactosyltransferase 15 | 12.05 | AT1G26810.2 |
| 749 | HORVU7Hr1G073460 | Tryptophan synthase beta chain 1 | 12.05 | AT4G27070.1 |
| 750 | HORVU5Hr1G063270 | GRAS family transcription factor | 12.02 | AT1G07520.1 |
| 751 | HORVU1Hr1G037150 | Protein kinase superfamily protein | 11.95 | AT5G03320.2 |
| 752 | HORVU5Hr1G045640 | NAC domain protein, | 11.94 | AT5G53950.1 |
| 753 | HORVU4Hr1G000070 | Poly(A) RNA polymerase protein 1 | 11.91 | AT3G45750.1 |
| 754 | HORVU7Hr1G046130 | undescribed protein | 11.89 | AT5G42740.2 |
| 755 | HORVU7Hr1G099340 | receptor kinase 1 | 11.80 | AT4G23280.2 |
| 756 | HORVU5Hr1G012120 | receptor kinase 2 | 11.79 | AT5G10530.1 |
| 757 | HORVU7Hr1G110570 | Protein of unknown function (DUF679) | 11.77 | AT4G24310.1 |
| 758 | HORVU1Hr1G089830 | Dehydrogenase/reductase SDR family member 12 | 11.75 | AT1G01800.1 |
| 759 | HORVU5Hr1G079690 | Acyl-coenzyme A thioesterase 9, mitochondrial | 11.70 | AT5G48370.2 |
| 760 | HORVU0Hr1G002930 | unknown function | 11.68 | AT2G15760.1 |
| 761 | HORVU1Hr1G085720 | undescribed protein | 11.57 | AT1G09480.2 |
| 762 | HORVU3Hr1G116350 | Ankyrin repeat family protein | 11.53 | AT2G28840.2 |
| 763 | HORVU5Hr1G046900 | Peroxidase superfamily protein | 11.53 | AT5G05340.1 |
| 764 | HORVU2Hr1G066200 | Core-2/I-branching beta-1,6-N-acetylglucosaminyltransferase family protein | 11.44 | AT1G71070.1 |
| 765 | HORVU2Hr1G081790 | amino acid permease 2 | 11.41 | AT5G09220.1 |
| 766 | HORVU7Hr1G051060 | undescribed protein | 11.40 | AT1G66880.1 |
| 767 | HORVU2Hr1G100090 | Solute carrier family 22 member 6-A | 11.31 | AT3G13050.4 |
| 768 | HORVU2Hr1G117870 | Protein kinase superfamily protein | 11.24 | AT4G21390.1 |
| 769 | HORVU6Hr1G060500 | membrane steroid binding protein 1 | 11.24 | AT3G48890.1 |
| 770 | HORVU5Hr1G024400 | Calmodulin-binding protein | 11.15 | AT2G18750.3 |
| 771 | HORVU1Hr1G092240 | Glucan endo-1,3-beta-glucosidase 13 | 11.13 | AT5G55180.2 |
| 772 | HORVU6Hr1G084080 | MATE efflux family protein | 11.09 | AT3G23560.1 |
| 773 | HORVU1Hr1G057460 | Aldolase-type TIM barrel family protein | 11.08 | AT5G64250.1 |
| 774 | HORVU7Hr1G044160 | Embryogenesis transmembrane protein-like | 10.95 | AT4G11000.1 |
| 775 | HORVU7Hr1G079460 | unknown function | 10.95 | AT5G53710.2 |
| 776 | HORVU6Hr1G066250 | Tetratricopeptide repeat (TPR)-like superfamily protein | 10.94 | AT3G18420.1 |
| 777 | HORVU7Hr1G029370 | undescribed protein | 10.92 | AT5G35360.3 |
| 778 | HORVU1Hr1G037550 | RING finger protein 141 | 10.91 | AT5G01520.1 |
| 779 | HORVU6Hr1G008880 | Heat shock 70 kDa protein C | 10.90 | AT5G42020.1 |
| 780 | HORVU2Hr1G109950 | undescribed protein | 10.89 | AT1G30450.4 |
| 781 | HORVU6Hr1G068990 | Adenine nucleotide alpha hydrolases-like superfamily protein | 10.87 | AT2G21620.1 |
| 782 | HORVU5Hr1G109190 | germin-like protein 4 | 10.84 | AT1G18980.1 |
| 783 | HORVU5Hr1G109290 | Calcium-binding EF-hand family protein | 10.79 | AT2G44310.1 |
| 784 | HORVU5Hr1G117910 | ferredoxin 3 | 10.76 | AT2G27510.1 |

| | | | | |
|---|---|---|---|---|
| 785 | HORVU0Hr1G030830 | WRKY family transcription factor | 10.75 | AT3G56400.1 |
| 786 | HORVU1Hr1G021160 | Glutathione S-transferase family protein | 10.73 | AT3G62760.1 |
| 787 | HORVU0Hr1G015180 | AT-hook motif nuclear-localized protein 1 | 10.67 | AT4G12080.1 |
| 788 | HORVU2Hr1G089760 | undescribed protein | 10.65 | AT5G18700.1 |
| 789 | HORVU5Hr1G087880 | Auxin efflux carrier family protein | 10.65 | AT5G65980.1 |
| 790 | HORVU1Hr1G065820 | Chromosome 3B, genomic scaffold, cultivar Chinese Spring | 10.64 | AT3G14460.1 |
| 791 | HORVU3Hr1G076890 | sphingosine kinase 1 | 10.63 | AT4G21534.2 |
| 792 | HORVU3Hr1G105190 | Glucan endo-1,3-beta-glucosidase GI | 10.60 | AT3G57260.2 |
| 793 | HORVU1Hr1G009130 | unknown function | 10.59 | AT2G18690.1 |
| 794 | HORVU2Hr1G017460 | NAC domain protein, | 10.57 | AT1G69490.1 |
| 795 | HORVU2Hr1G114190 | UDP-Glycosyltransferase superfamily protein | 10.51 | AT2G43820.1 |
| 796 | HORVU2Hr1G004720 | UDP-Glycosyltransferase superfamily protein | 10.49 | AT1G22400.1 |
| 797 | HORVU3Hr1G001060 | Chromosome 3B, genomic scaffold, cultivar Chinese Spring | 10.47 | AT1G49000.1 |
| 798 | HORVU3Hr1G093400 | RNA-binding protein 1 | 10.45 | AT3G23830.2 |
| 799 | HORVU2Hr1G093040 | undescribed protein | 10.43 | AT4G00060.4 |
| 800 | HORVU3Hr1G076400 | undescribed protein | 10.42 | AT2G32210.1 |
| 801 | HORVU2Hr1G107100 | Protein translation factor SUI1 homolog 1 | 10.40 | AT5G54760.4 |
| 802 | HORVU3Hr1G014270 | Phloem-specific lectin | 10.37 | AT2G02340.1 |
| 803 | HORVU2Hr1G064160 | sugar transporter 1 | 10.32 | AT1G11260.1 |
| 804 | HORVU5Hr1G051960 | Pathogenesis-related thaumatin superfamily protein | 10.31 | AT1G77700.1 |
| 805 | HORVU2Hr1G090200 | UDP-Glycosyltransferase superfamily protein | 10.23 | AT3G53150.1 |
| 806 | HORVU6Hr1G030880 | Receptor-like protein kinase 5 | 10.17 | AT2G20850.1 |
| 807 | HORVU7Hr1G096660 | histone H2A 2 | 10.15 | AT4G27230.2 |
| 808 | HORVU3Hr1G062940 | O-acyltransferase (WSD1-like) family protein | 10.14 | AT5G53390.1 |
| 809 | HORVU5Hr1G115230 | non-specific phospholipase C1 | 10.11 | AT1G07230.1 |
| 810 | HORVU5Hr1G005180 | Pathogenesis-related thaumatin superfamily protein | 10.11 | AT4G11650.1 |
| 811 | HORVU7Hr1G050670 | lipoxygenase 2 | 10.11 | AT3G45140.1 |
| 812 | HORVU2Hr1G109370 | Adenosylhomocysteinase | 10.05 | AT4G13940.1 |
| 813 | HORVU4Hr1G000810 | undescribed protein | 10.04 | AT1G63720.1 |
| 814 | HORVU2Hr1G001940 | undescribed protein | 10.04 | AT4G25120.1 |
| 815 | HORVU3Hr1G105870 | Amino-acid permease BAT1 homolog | 10.01 | AT2G01170.1 |
| 816 | HORVU1Hr1G063390 | AAA-ATPase 1 | 10.01 | AT5G40010.1 |
| 817 | HORVU7Hr1G028840 | 2-oxoglutarate (2OG) and Fe(II)-dependent oxygenase superfamily protein | 9.98 | AT5G54000.2 |
| 818 | HORVU2Hr1G017400 | NAC domain protein, | 9.97 | AT5G39610.1 |
| 819 | HORVU4Hr1G011430 | pfkB-like carbohydrate kinase family protein | 9.94 | AT5G58730.1 |
| 820 | HORVU6Hr1G081850 | Subtilisin-like protease | 9.93 | AT3G14067.1 |
| 821 | HORVU4Hr1G084740 | sugar transporter 1 | 9.92 | AT1G34580.1 |
| 822 | HORVU5Hr1G019820 | ATP-dependent zinc metalloprotease FtsH 2 | 9.89 | AT5G40010.1 |

| | | | | |
|---|---|---|---|---|
| 823 | HORVU7Hr1G043690 | UDP-Glycosyltransferase superfamily protein | 9.87 | AT4G01070.1 |
| 824 | HORVU7Hr1G079630 | undescribed protein | 9.86 | AT2G33600.1 |
| 825 | HORVU6Hr1G035420 | cinnamyl alcohol dehydrogenase 5 | 9.83 | AT3G19450.1 |
| 826 | HORVU5Hr1G104580 | UDP-Glycosyltransferase superfamily protein | 9.82 | AT2G15480.2 |
| 827 | HORVU7Hr1G000970 | HXXXD-type acyl-transferase family protein | 9.79 | AT5G42830.1 |
| 828 | HORVU4Hr1G067490 | Myosin-related, putative | 9.75 | AT1G51405.2 |
| 829 | HORVU4Hr1G014970 | Kelch-like protein 23 | 9.69 | AT2G44130.1 |
| 830 | HORVU1Hr1G075950 | unknown function | 9.67 | AT3G16330.1 |
| 831 | HORVU5Hr1G095130 | Beta-glucosidase C | 9.64 | AT5G04885.2 |
| 832 | HORVU4Hr1G069530 | unknown function | 9.64 | AT5G43150.1 |
| 833 | HORVU7Hr1G011250 | E3 ubiquitin-protein ligase XBAT33 | 9.61 | AT5G07270.1 |
| 834 | HORVU1Hr1G002470 | receptor-like protein kinase 4 | 9.57 | AT1G70250.1 |
| 835 | HORVU6Hr1G093740 | Receptor-like protein kinase | 9.56 | AT1G21230.1 |
| 836 | HORVU1Hr1G039720 | Protein NRT1/ PTR FAMILY 5.2 | 9.54 | AT5G46050.1 |
| 837 | HORVU5Hr1G062090 | Aldehyde dehydrogenase family 7 member A1 | 9.54 | AT1G54100.2 |
| 838 | HORVU2Hr1G118080 | Heavy metal transport/detoxification superfamily protein | 9.53 | AT5G52740.1 |
| 839 | HORVU2Hr1G090170 | UDP-Glycosyltransferase superfamily protein | 9.53 | AT2G36790.1 |
| 840 | HORVU1Hr1G070690 | PEBP (phosphatidylethanolamine-binding protein) family protein | 9.49 | AT5G01300.1 |
| 841 | HORVU2Hr1G038830 | Ferredoxin--NADP reductase | 9.48 | AT4G05390.1 |
| 842 | HORVU4Hr1G023680 | jacalin-related lectin 23 | 9.48 | AT1G19715.2 |
| 843 | HORVU3Hr1G027700 | Class I glutamine amidotransferase-like superfamily protein | 9.44 | AT1G53280.1 |
| 844 | HORVU2Hr1G001790 | Vacuolar protein sorting-associated protein 27 | 9.39 | AT5G16880.4 |
| 845 | HORVU4Hr1G065800 | Inactive poly [ADP-ribose] polymerase RCD1 | 9.36 | AT1G32230.3 |
| 846 | HORVU3Hr1G006640 | receptor kinase 2 | 9.35 | AT4G23180.1 |
| 847 | HORVU7Hr1G101590 | Cytochrome P450 superfamily protein | 9.34 | AT3G14690.1 |
| 848 | HORVU7Hr1G090280 | Tetratricopeptide repeat (TPR)-like superfamily protein | 9.33 | AT5G20190.1 |
| 849 | HORVU1Hr1G002600 | receptor-like protein kinase 4 | 9.32 | AT1G70250.1 |
| 850 | HORVU3Hr1G066450 | IAA-amino acid hydrolase ILR1-like 2 | 9.32 | AT1G51760.1 |
| 851 | HORVU6Hr1G085870 | Transmembrane 9 superfamily member 11 | 9.31 | AT5G35160.4 |
| 852 | HORVU6Hr1G032710 | 50S ribosomal protein L7/L12 | 9.29 | AT3G06040.3 |
| 853 | HORVU2Hr1G072210 | Arogenate dehydratase/prephenate dehydratase 6, chloroplastic | 9.25 | AT5G53460.3 |
| 854 | HORVU0Hr1G009390 | germin-like protein 2 | 9.24 | AT5G39110.1 |
| 855 | HORVU2Hr1G014890 | receptor kinase 2 | 9.23 | AT3G53810.1 |
| 856 | HORVU7Hr1G050660 | lipoxygenase 2 | 9.06 | AT3G45140.2 |
| 857 | HORVU3Hr1G070560 | Transferrin receptor protein 2 | 8.98 | AT5G19740.1 |
| 858 | HORVU1Hr1G020690 | Peroxidase superfamily protein | 8.97 | AT5G06730.1 |
| 859 | HORVU5Hr1G050240 | Cyclase family protein | 8.94 | AT4G35220.1 |
| 860 | HORVU5Hr1G098030 | undescribed protein | 8.88 | AT2G15960.1 |

| | | | | |
|---|---|---|---|---|
| 861 | HORVU3Hr1G001790 | Chromosome 3B, genomic scaffold, cultivar Chinese Spring | 8.88 | AT2G29050.2 |
| 862 | HORVU7Hr1G031430 | glutamate receptor 2.9 | 8.80 | AT5G11210.3 |
| 863 | HORVU2Hr1G114590 | Synaptotagmin-5 | 8.78 | AT1G05500.1 |
| 864 | HORVU6Hr1G000180 | 60 kDa chaperonin 2 | 8.78 | AT3G13470.1 |
| 865 | HORVU5Hr1G084900 | Glucan endo-1,3-beta-glucosidase 13 | 8.74 | AT5G55180.1 |
| 866 | HORVU1Hr1G012680 | UDP-Glycosyltransferase superfamily protein | 8.71 | AT2G36970.1 |
| 867 | HORVU6Hr1G090170 | 50S ribosomal protein L14 | 8.71 | AT3G04400.2 |
| 868 | HORVU1Hr1G003720 | Mitochondrial import inner membrane translocase subunit TIM14-2 | 8.69 | AT3G09700.1 |
| 869 | HORVU5Hr1G096930 | Cytochrome P450 superfamily protein | 8.67 | AT4G37330.1 |
| 870 | HORVU1Hr1G093890 | Pathogen-related protein | 8.66 | AT1G78780.5 |
| 871 | HORVU5Hr1G097270 | Peroxidase superfamily protein | 8.65 | AT3G49960.1 |
| 872 | HORVU2Hr1G019180 | UDP-glucuronic acid decarboxylase 1 | 8.64 | AT2G47650.1 |
| 873 | HORVU1Hr1G080680 | Cytochrome P450 superfamily protein | 8.58 | AT3G48520.1 |
| 874 | HORVU1Hr1G054390 | lipoxygenase 1 | 8.57 | AT3G22400.1 |
| 875 | HORVU5Hr1G024160 | Adenine nucleotide alpha hydrolases-like superfamily protein | 8.56 | AT2G47710.1 |
| 876 | HORVU2Hr1G099530 | cationic amino acid transporter 8 | 8.55 | AT1G17120.1 |
| 877 | HORVU2Hr1G027370 | tRNA pseudouridine synthase B | 8.54 | AT3G57150.1 |
| 878 | HORVU6Hr1G057680 | inter-alpha-trypsin inhibitor heavy chain-related | 8.53 | AT5G60710.1 |
| 879 | HORVU2Hr1G110120 | Adenosylhomocysteinase | 8.50 | AT4G13940.1 |
| 880 | HORVU1Hr1G049100 | Glutathione S-transferase family protein | 8.49 | AT1G10360.1 |
| 881 | HORVU4Hr1G076520 | Protein kinase superfamily protein | 8.49 | AT2G05940.1 |
| 882 | HORVU4Hr1G068530 | unknown protein; Has 65 Blast hits to 65 proteins in 11 species: Archae - 0; Bacteria - 0; Metazoa - 0; Fungi - 0; Plants - 65; Viruses - 0; Other Eukaryotes - 0 (source: NCBI BLink). | 8.48 | AT3G57450.1 |
| 883 | HORVU2Hr1G031410 | Transmembrane protein 87A | 8.41 | AT1G72480.1 |
| 884 | HORVU1Hr1G056600 | Protein kinase superfamily protein | 8.41 | AT3G51990.1 |
| 885 | HORVU6Hr1G070130 | undescribed protein | 8.41 | AT2G43420.1 |
| 886 | HORVU2Hr1G078890 | unknown function | 8.40 | AT4G37030.1 |
| 887 | HORVU2Hr1G096510 | 2-oxoglutarate (2OG) and Fe(II)-dependent oxygenase superfamily protein | 8.38 | AT5G20400.1 |
| 888 | HORVU5Hr1G114440 | unknown function | 8.37 | AT4G35920.5 |
| 889 | HORVU5Hr1G070360 | Protein kinase family protein | 8.32 | AT1G21240.1 |
| 890 | HORVU0Hr1G039970 | Leucine-rich receptor-like protein kinase family protein | 8.31 | AT3G47090.1 |
| 891 | HORVU5Hr1G016400 | Peroxisomal membrane 22 kDa (Mpv17/PMP22) family protein | 8.28 | AT4G14305.1 |
| 892 | HORVU4Hr1G085750 | Hydroxymethylglutaryl-CoA synthase | 8.27 | AT4G11820.2 |
| 893 | HORVU1Hr1G074840 | unknown function | 8.25 | AT5G17210.1 |
| 894 | HORVU5Hr1G080310 | Dehydration-responsive element-binding protein 1B | 8.21 | AT4G25470.1 |

| | | | | |
|---|---|---|---|---|
| 895 | HORVU2Hr1G009830 | Cystic fibrosis transmembrane conductance regulator | 8.21 | AT2G47800.1 |
| 896 | HORVU6Hr1G009420 | undescribed protein | 8.18 | AT3G59100.1 |
| 897 | HORVU6Hr1G024850 | Cytochrome P450 superfamily protein | 8.18 | AT3G48310.1 |
| 898 | HORVU3Hr1G065390 | Lipid A export ATP-binding/permease protein MsbA | 8.13 | AT2G47000.6 |
| 899 | HORVU1Hr1G080320 | aldehyde dehydrogenase 12A1 | 8.12 | AT5G62530.1 |
| 900 | HORVU2Hr1G017950 | B12D protein | 8.12 | AT3G48140.1 |
| 901 | HORVU2Hr1G102110 | Bifunctional inhibitor/lipid-transfer protein/seed storage 2S albumin superfamily protein | 8.10 | AT5G48485.1 |
| 902 | HORVU5Hr1G087240 | receptor kinase 3 | 8.10 | AT4G21390.2 |
| 903 | HORVU1Hr1G084620 | CASP-like protein 1U2 | 8.07 | AT4G15630.1 |
| 904 | HORVU4Hr1G090130 | germin-like protein 2 | 8.05 | AT5G39110.1 |
| 905 | HORVU5Hr1G014060 | calmodulin-like 11 | 8.04 | AT3G22930.1 |
| 906 | HORVU7Hr1G050680 | lipoxygenase 2 | 8.04 | AT3G45140.1 |
| 907 | HORVU1Hr1G043920 | Plant basic secretory protein (BSP) family protein | 7.98 | AT2G15220.1 |
| 908 | HORVU3Hr1G117370 | Glutathione S-transferase family protein | 7.97 | AT3G62760.1 |
| 909 | HORVU6Hr1G052060 | unknown function | 7.96 | AT5G41240.1 |
| 910 | HORVU3Hr1G028260 | RING/U-box superfamily protein | 7.96 | AT2G27940.1 |
| 911 | HORVU4Hr1G060280 | AT-hook motif nuclear-localized protein 22 | 7.95 | AT1G76500.1 |
| 912 | HORVU0Hr1G009400 | germin-like protein 9 | 7.90 | AT5G39110.1 |
| 913 | HORVU5Hr1G084860 | Cysteine-rich receptor-like protein kinase 10 | 7.89 | AT3G45860.1 |
| 914 | HORVU2Hr1G114260 | Glycerol kinase | 7.89 | AT1G80460.1 |
| 915 | HORVU2Hr1G022500 | undescribed protein | 7.88 | AT3G03770.3 |
| 916 | HORVU7Hr1G108450 | Leucine-rich repeat receptor-like protein kinase family protein | 7.88 | AT5G20480.2 |
| 917 | HORVU7Hr1G033520 | receptor-like protein kinase 1 | 7.80 | AT5G10530.1 |
| 918 | HORVU2Hr1G042210 | Serine/threonine-protein kinase | 7.79 | AT2G19130.1 |
| 919 | HORVU2Hr1G093020 | undescribed protein | 7.77 | AT5G08780.1 |
| 920 | HORVU1Hr1G051200 | ERD (early-responsive to dehydration stress) family protein | 7.74 | AT1G32090.1 |
| 921 | HORVU7Hr1G118610 | Protein kinase superfamily protein | 7.70 | AT1G68690.3 |
| 922 | HORVU4Hr1G088860 | Major facilitator superfamily protein | 7.66 | AT1G64650.1 |
| 923 | HORVU6Hr1G068160 | undescribed protein | 7.64 | AT5G61120.3 |
| 924 | HORVU2Hr1G023450 | Carboxypeptidase Y homolog A | 7.63 | AT3G63470.1 |
| 925 | HORVU4Hr1G023740 | receptor-like protein kinase 1 | 7.58 | AT1G72300.1 |
| 926 | HORVU6Hr1G005900 | alpha/beta-Hydrolases superfamily protein | 7.58 | AT1G47480.1 |
| 927 | HORVU1Hr1G089310 | B12D protein | 7.58 | AT3G48140.1 |
| 928 | HORVU3Hr1G048610 | Protein transport protein Sec61 subunit beta | 7.58 | AT3G60540.3 |
| 929 | HORVU6Hr1G089510 | germin-like protein 5 | 7.49 | AT1G02335.1 |
| 930 | HORVU2Hr1G039850 | MATE efflux family protein | 7.48 | AT1G15170.2 |
| 931 | HORVU6Hr1G086560 | Protein phosphatase 2C family protein | 7.46 | AT2G25620.2 |

| | | | | |
|---|---|---|---|---|
| 932 | HORVU7Hr1G019240 | sugar transporter 9 | 7.46 | AT5G61520.2 |
| 933 | HORVU5Hr1G062410 | Hypersensitive-induced response protein 1 | 7.45 | AT5G62740.1 |
| 934 | HORVU3Hr1G004850 | UDP-Glycosyltransferase superfamily protein | 7.45 | AT4G01070.1 |
| 935 | HORVU1Hr1G090430 | CTP synthase family protein | 7.43 | AT1G30820.1 |
| 936 | HORVU5Hr1G096950 | Cytochrome P450 superfamily protein | 7.40 | AT5G67310.1 |
| 937 | HORVU2Hr1G105240 | purine permease 11 | 7.39 | AT1G44750.1 |
| 938 | HORVU2Hr1G090260 | Protein RETICULATA, chloroplastic | 7.38 | AT2G37860.4 |
| 939 | HORVU7Hr1G090950 | adenylate kinase 1 | 7.38 | AT5G63400.1 |
| 940 | HORVU7Hr1G017950 | Cytochrome P450 superfamily protein | 7.37 | AT2G45560.1 |
| 941 | HORVU1Hr1G000660 | Leucine-rich receptor-like protein kinase family protein | 7.33 | AT1G35710.1 |
| 942 | HORVU1Hr1G045110 | RING finger protein 141 | 7.29 | AT3G47160.1 |
| 943 | HORVU4Hr1G052010 | WRKY DNA-binding protein 46 | 7.29 | AT2G46400.1 |
| 944 | HORVU1Hr1G021150 | Glutathione S-transferase family protein | 7.29 | AT3G62760.1 |
| 945 | HORVU3Hr1G063840 | alpha/beta-Hydrolases superfamily protein | 7.29 | AT1G77420.1 |
| 946 | HORVU3Hr1G070540 | Transferrin receptor protein 2 | 7.22 | AT5G19740.1 |
| 947 | HORVU5Hr1G072730 | undescribed protein | 7.22 | #N/A |
| 948 | HORVU7Hr1G040730 | Cysteine-rich venom protein | 7.22 | AT5G57625.1 |
| 949 | HORVU4Hr1G078690 | Ras-related protein Rab-6A | 7.21 | AT2G44610.1 |
| 950 | HORVU2Hr1G036800 | Geranylgeranyl diphosphate synthase | 7.19 | AT4G36810.1 |
| 951 | HORVU2Hr1G090280 | YELLOW STRIPE like 7 | 7.19 | AT1G65730.1 |
| 952 | HORVU6Hr1G074690 | receptor-like protein kinase 4 | 7.17 | AT1G34300.1 |
| 953 | HORVU2Hr1G043400 | undescribed protein | 7.16 | #N/A |
| 954 | HORVU5Hr1G047060 | plastid transcriptionally active 6 | 7.16 | AT1G21600.2 |
| 955 | HORVU2Hr1G108460 | unknown function | 7.11 | AT3G16660.2 |
| 956 | HORVU2Hr1G095330 | undescribed protein | 7.11 | AT3G29810.1 |
| 957 | HORVU2Hr1G043890 | Xylanase inhibitor protein 1 | 7.09 | AT5G24090.1 |
| 958 | HORVU4Hr1G000910 | Peptidyl-prolyl cis-trans isomerase | 7.07 | AT3G56070.2 |
| 959 | HORVU7Hr1G048690 | BTB/POZ domain-containing protein | 7.07 | AT1G55760.1 |
| 960 | HORVU4Hr1G078830 | Heat shock 70 kDa protein 17 | 7.07 | AT4G16660.1 |
| 961 | HORVU5Hr1G103770 | alpha/beta-Hydrolases superfamily protein | 7.07 | AT5G06570.2 |
| 962 | HORVU7Hr1G091350 | SAUR-like auxin-responsive protein family | 7.07 | AT2G46690.1 |
| 963 | HORVU2Hr1G032360 | undescribed protein | 7.06 | AT5G61412.1 |
| 964 | HORVU7Hr1G080950 | WRKY DNA-binding protein 27 | 7.03 | AT4G01250.1 |
| 965 | HORVU3Hr1G096050 | Transaldolase 1 | 7.03 | AT5G13420.1 |
| 966 | HORVU4Hr1G084330 | Inter-alpha-trypsin inhibitor heavy chain H4 | 7.01 | AT5G60710.1 |
| 967 | HORVU7Hr1G106960 | Protein LIGHT-DEPENDENT SHORT HYPOCOTYLS 4 | 6.97 | AT1G07090.1 |
| 968 | HORVU7Hr1G037910 | shikimate kinase 1 | 6.95 | AT2G21940.5 |
| 969 | HORVU2Hr1G066920 | Voltage-gated potassium channel subunit beta-3 | 6.89 | AT1G60730.3 |
| 970 | HORVU3Hr1G056100 | Cytochrome P450 superfamily protein | 6.88 | AT3G14680.1 |

| | | | | |
|---|---|---|---|---|
| 971 | HORVU7Hr1G037510 | Protein of unknown function, DUF547 | 6.88 | AT4G08550.2 |
| 972 | HORVU7Hr1G073050 | External alternative NAD(P)H-ubiquinone oxidoreductase B3, mitochondrial | 6.88 | AT4G21490.2 |
| 973 | HORVU4Hr1G066270 | allene oxide synthase | 6.88 | AT5G42650.1 |
| 974 | HORVU2Hr1G044910 | receptor kinase 2 | 6.87 | AT4G23180.3 |
| 975 | HORVU2Hr1G034370 | receptor-like protein kinase 2 | 6.84 | AT1G74360.1 |
| 976 | HORVU4Hr1G021000 | undescribed protein | 6.83 | AT3G10195.1 |
| 977 | HORVU6Hr1G070710 | unknown function | 6.75 | AT3G12650.1 |
| 978 | HORVU5Hr1G112210 | Transmembrane amino acid transporter family protein | 6.74 | AT3G11900.2 |
| 979 | HORVU2Hr1G065100 | unknown protein; Has 978 Blast hits to 254 proteins in 81 species: Archae - 0; Bacteria - 8; Metazoa - 109; Fungi - 53; Plants - 41; Viruses - 0; Other Eukaryotes - 767 (source: NCBI BLink). | 6.74 | AT2G39950.9 |
| 980 | HORVU5Hr1G093160 | ATP-dependent RNA helicase DeaD | 6.72 | AT4G16630.1 |
| 981 | HORVU6Hr1G081000 | allene oxide cyclase 4 | 6.69 | AT3G25780.1 |
| 982 | HORVU7Hr1G019280 | sugar transporter 1 | 6.67 | AT5G61520.1 |
| 983 | HORVU5Hr1G074360 | undescribed protein | 6.66 | AT5G35320.1 |
| 984 | HORVU3Hr1G117910 | PLAC8 family protein | 6.64 | AT5G05350.1 |
| 985 | HORVU4Hr1G019410 | alkaline/neutral invertase | 6.62 | AT4G09510.1 |
| 986 | HORVU4Hr1G076570 | lipoxygenase 3 | 6.62 | AT1G72520.1 |
| 987 | HORVU2Hr1G004230 | 12-oxophytodienoate reductase 2 | 6.62 | AT1G76680.1 |
| 988 | HORVU5Hr1G118750 | Zinc finger protein 668 | 6.61 | AT1G34370.2 |
| 989 | HORVU5Hr1G062510 | Dormancy/auxin associated family protein | 6.59 | AT1G56220.2 |
| 990 | HORVU3Hr1G062430 | Chromosome 3B, genomic scaffold, cultivar Chinese Spring | 6.58 | AT4G21700.1 |
| 991 | HORVU2Hr1G077220 | 12-oxophytodienoate reductase 2 | 6.54 | AT1G76690.1 |
| 992 | HORVU2Hr1G044680 | receptor kinase 1 | 6.54 | AT4G23180.1 |
| 993 | HORVU2Hr1G024160 | chaperone protein dnaJ-related | 6.52 | AT2G38000.1 |
| 994 | HORVU5Hr1G066130 | unknown function | 6.51 | AT3G47510.1 |
| 995 | HORVU4Hr1G064010 | Protein kinase superfamily protein | 6.49 | AT1G77280.2 |
| 996 | HORVU0Hr1G018830 | Cell division protein FtsZ | 6.49 | AT5G55280.1 |
| 997 | HORVU3Hr1G071880 | unknown protein; Has 35333 Blast hits to 34131 proteins in 2444 species: Archae - 798; Bacteria - 22429; Metazoa - 974; Fungi - 991; Plants - 531; Viruses - 0; Other Eukaryotes - 9610 (source: NCBI BLink). | 6.49 | AT1G06475.1 |
| 998 | HORVU2Hr1G009580 | ABC transporter C family member 14 | 6.49 | AT3G62700.1 |
| 999 | HORVU4Hr1G088720 | Palmitoyl-protein thioesterase 1 | 6.47 | AT3G60340.2 |
| 1000 | HORVU2Hr1G022540 | undescribed protein | 6.44 | AT2G03370.2 |
| 1001 | HORVU5Hr1G079180 | 3-beta hydroxysteroid dehydrogenase/isomerase family protein | 6.42 | AT2G43420.1 |
| 1002 | HORVU5Hr1G104050 | purine permease 3 | 6.40 | AT1G28220.1 |
| 1003 | HORVU1Hr1G040940 | Heat shock 70 kDa protein 17 | 6.39 | AT4G16660.1 |
| 1004 | HORVU0Hr1G020420 | 17.6 kDa class II heat shock protein | 6.37 | AT5G37670.1 |

| | | | | |
|---|---|---|---|---|
| 1005 | HORVU7Hr1G116060 | Cytochrome P450 superfamily protein | 6.37 | AT3G48310.1 |
| 1006 | HORVU0Hr1G005420 | Ubiquinol oxidase | 6.36 | AT3G22360.1 |
| 1007 | HORVU7Hr1G048430 | Glycosyltransferase | 6.32 | AT3G18180.1 |
| 1008 | HORVU5Hr1G106550 | histone H1-3 | 6.32 | AT2G30620.2 |
| 1009 | HORVU7Hr1G091100 | Glucan endo-1,3-beta-glucosidase 13 | 6.29 | AT2G05790.1 |
| 1010 | HORVU4Hr1G012510 | unknown function | 6.26 | AT3G10020.1 |
| 1011 | HORVU0Hr1G015140 | undescribed protein | 6.21 | AT5G09850.5 |
| 1012 | HORVU7Hr1G010250 | Peroxisomal membrane protein 11-5 | 6.20 | AT1G01820.1 |
| 1013 | HORVU1Hr1G042370 | receptor kinase 1 | 6.20 | AT1G70520.1 |
| 1014 | HORVU7Hr1G042260 | Cytochrome P450 superfamily protein | 6.18 | AT5G07990.1 |
| 1015 | HORVU1Hr1G073330 | HXXXD-type acyl-transferase family protein | 6.17 | AT2G39980.1 |
| 1016 | HORVU2Hr1G121440 | Cathepsin B-like cysteine proteinase | 6.15 | AT5G43060.1 |
| 1017 | HORVU4Hr1G046610 | Heavy metal transport/detoxification superfamily protein | 6.15 | AT1G56210.1 |
| 1018 | HORVU3Hr1G018650 | pyruvate decarboxylase-2 | 6.13 | AT5G01320.1 |
| 1019 | HORVU0Hr1G005070 | F-box protein | 6.12 | AT1G47056.1 |
| 1020 | HORVU5Hr1G055060 | IAA-amino acid hydrolase ILR1-like 9 | 6.10 | AT3G02875.1 |
| 1021 | HORVU1Hr1G001950 | 12-oxophytodienoate reductase 2 | 6.10 | AT1G76680.1 |
| 1022 | HORVU7Hr1G043810 | Embryogenesis transmembrane protein-like | 6.09 | AT4G11000.1 |
| 1023 | HORVU2Hr1G084200 | undescribed protein | 6.07 | AT3G11402.1 |
| 1024 | HORVU7Hr1G013170 | Flavin-containing monooxygenase family protein | 6.05 | AT1G21430.1 |
| 1025 | HORVU3Hr1G078840 | UDP-Glycosyltransferase superfamily protein | 6.02 | AT2G30150.2 |
| 1026 | HORVU6Hr1G093180 | Electron transfer flavoprotein subunit beta | 6.00 | AT5G43430.1 |
| 1027 | HORVU2Hr1G064530 | glutamate receptor 3.4 | 5.97 | AT1G05200.6 |
| 1028 | HORVU1Hr1G017050 | Glutathione S-transferase family protein | 5.95 | AT3G62760.1 |
| 1029 | HORVU2Hr1G039050 | copper transporter 1 | 5.95 | AT2G26975.1 |
| 1030 | HORVU2Hr1G073320 | 60S ribosomal protein L5-1 | 5.95 | AT5G39740.2 |
| 1031 | HORVU6Hr1G013750 | Protein kinase superfamily protein | 5.95 | AT2G19130.1 |
| 1032 | HORVU2Hr1G010590 | Leucine-rich repeat receptor-like protein kinase family protein | 5.92 | AT3G47570.1 |
| 1033 | HORVU6Hr1G021320 | Lipid A export ATP-binding/permease protein MsbA | 5.91 | AT2G47000.7 |
| 1034 | HORVU1Hr1G047050 | undescribed protein | 5.91 | AT1G02110.1 |
| 1035 | HORVU1Hr1G038680 | 60 kDa chaperonin 1 | 5.90 | AT3G23990.1 |
| 1036 | HORVU7Hr1G091250 | alpha-amylase-like | 5.88 | AT4G25000.1 |
| 1037 | HORVU2Hr1G122290 | unknown function | 5.86 | AT1G48720.1 |
| 1038 | HORVU2Hr1G089160 | 3-hydroxyacyl-[acyl-carrier-protein] dehydratase FabZ | 5.86 | AT2G22230.1 |
| 1039 | HORVU5Hr1G040740 | Long-chain-fatty-acid--CoA ligase 1 | 5.86 | AT5G27600.1 |
| 1040 | HORVU1Hr1G011430 | Pyrimidine-specific ribonucleoside hydrolase RihA | 5.85 | AT2G36310.1 |
| 1041 | HORVU4Hr1G078470 | alcohol dehydrogenase 1 | 5.85 | AT5G42250.1 |
| 1042 | HORVU3Hr1G106720 | Pyrroline-5-carboxylate reductase | 5.81 | AT5G14800.1 |

| | | | | |
|---|---|---|---|---|
| 1043 | HORVU4Hr1G087340 | 60 kDa jasmonate-induced protein | 5.80 | AT2G35330.2 |
| 1044 | HORVU2Hr1G018570 | Peroxidase superfamily protein | 5.78 | AT5G05340.1 |
| 1045 | HORVU7Hr1G079830 | Diacylglycerol kinase | 5.77 | AT2G18730.1 |
| 1046 | HORVU2Hr1G036590 | sugar transporter 9 | 5.77 | AT3G18830.1 |
| 1047 | HORVU5Hr1G060070 | unknown function | 5.77 | AT1G70160.1 |
| 1048 | HORVU6Hr1G085370 | Elongation factor 1-alpha | 5.77 | AT5G60390.3 |
| 1049 | HORVU6Hr1G069400 | formate dehydrogenase | 5.75 | AT5G14780.3 |
| 1050 | HORVU4Hr1G048700 | Zinc finger protein CONSTANS-LIKE 13 | 5.74 | AT2G47890.1 |
| 1051 | HORVU2Hr1G023470 | undescribed protein | 5.74 | AT2G33100.2 |
| 1052 | HORVU1Hr1G053070 | metallo-beta-lactamase family protein | 5.73 | AT4G33540.2 |
| 1053 | HORVU4Hr1G067840 | GDP-mannose transporter GONST3 | 5.73 | AT1G76340.1 |
| 1054 | HORVU3Hr1G117760 | DnaJ protein P58IPK homolog | 5.73 | AT5G03160.1 |
| 1055 | HORVU2Hr1G028330 | EF hand family protein | 5.72 | AT5G04990.1 |
| 1056 | HORVU5Hr1G105480 | alpha/beta-Hydrolases superfamily protein | 5.72 | AT1G47480.1 |
| 1057 | HORVU2Hr1G022350 | UDP-N-acetylglucosamine--dolichyl-phosphate N-acetylglucosaminephosphotransferase | 5.71 | AT3G57220.1 |
| 1058 | HORVU1Hr1G072250 | Chitinase 2 | 5.70 | AT4G20095.1 |
| 1059 | HORVU7Hr1G118450 | receptor kinase 2 | 5.70 | AT4G38830.1 |
| 1060 | HORVU4Hr1G081100 | Glutathione S-transferase family protein | 5.70 | AT3G03190.1 |
| 1061 | HORVU2Hr1G121990 | calreticulin 1b | 5.68 | AT1G56340.2 |
| 1062 | HORVU6Hr1G043980 | Dolichyl-diphosphooligosaccharide--protein glycosyltransferase subunit 4A | 5.67 | AT3G09455.2 |
| 1063 | HORVU2Hr1G101920 | Ubiquinol oxidase | 5.65 | AT3G22370.1 |
| 1064 | HORVU7Hr1G086580 | protein kinase family protein | 5.64 | AT5G57035.1 |
| 1065 | HORVU7Hr1G088770 | undescribed protein | 5.60 | AT2G20420.1 |
| 1066 | HORVU5Hr1G073580 | DNA-binding bromodomain-containing protein | 5.58 | AT2G44430.1 |
| 1067 | HORVU2Hr1G109380 | Carboxypeptidase Y | 5.57 | AT3G10410.1 |
| 1068 | HORVU1Hr1G079070 | Farnesyl pyrophosphate synthase | 5.54 | AT5G47770.1 |
| 1069 | HORVU7Hr1G026720 | early nodulin-related | 5.53 | AT5G25940.1 |
| 1070 | HORVU7Hr1G047080 | MATE efflux family protein | 5.52 | AT5G52450.1 |
| 1071 | HORVU4Hr1G005450 | Protein of unknown function, DUF538 | 5.51 | AT5G19860.1 |
| 1072 | HORVU7Hr1G071020 | RING/U-box superfamily protein | 5.51 | AT3G43430.1 |
| 1073 | HORVU7Hr1G083180 | Protein of unknown function (DUF1218) | 5.50 | AT1G13380.1 |
| 1074 | HORVU4Hr1G013330 | zinc finger protein 7 | 5.49 | AT1G24625.1 |
| 1075 | HORVU5Hr1G077010 | NAC domain containing protein 41 | 5.49 | AT2G33480.2 |
| 1076 | HORVU5Hr1G076250 | purple acid phosphatase 27 | 5.49 | AT5G50400.1 |
| 1077 | HORVU0Hr1G027290 | 1-acyl-sn-glycerol-3-phosphate acyltransferase 2 | 5.44 | AT3G18850.4 |
| 1078 | HORVU6Hr1G018420 | calnexin 1 | 5.44 | AT5G61790.1 |
| 1079 | HORVU4Hr1G084340 | Inter-alpha-trypsin inhibitor heavy chain H1 | 5.44 | AT5G60710.1 |
| 1080 | HORVU1Hr1G001660 | receptor-like protein kinase 4 | 5.42 | AT1G66920.2 |
| 1081 | HORVU3Hr1G095820 | cyclin-dependent kinase C;1 | 5.41 | AT5G50860.1 |

| | | | | |
|---|---|---|---|---|
| 1082 | HORVU7Hr1G078530 | Protein kinase superfamily protein | 5.38 | AT1G63500.1 |
| 1083 | HORVU5Hr1G002120 | ADP-ribosylation factor family protein | 5.36 | AT2G18390.1 |
| 1084 | HORVU4Hr1G018150 | sterol C4-methyl oxidase 1-2 | 5.34 | AT4G22756.1 |
| 1085 | HORVU1Hr1G053440 | Proline dehydrogenase 2, mitochondrial | 5.33 | AT5G38710.1 |
| 1086 | HORVU5Hr1G002150 | RNA-binding protein 1 | 5.33 | AT4G39260.3 |
| 1087 | HORVU2Hr1G077990 | F-box protein SKIP19 | 5.32 | AT4G05460.1 |
| 1088 | HORVU3Hr1G033550 | Carboxypeptidase Y homolog A | 5.32 | AT3G17180.2 |
| 1089 | HORVU3Hr1G028060 | GDSL esterase/lipase | 5.29 | AT5G45910.1 |
| 1090 | HORVU1Hr1G068170 | tubulin beta chain 2 | 5.28 | AT5G12250.1 |
| 1091 | HORVU4Hr1G081940 | 4-coumarate--CoA ligase-like 4 | 5.27 | AT1G20510.1 |
| 1092 | HORVU5Hr1G115250 | unknown function | 5.24 | AT1G19530.2 |
| 1093 | HORVU4Hr1G064790 | glutamate decarboxylase | 5.24 | AT5G17330.1 |
| 1094 | HORVU3Hr1G085930 | Protein phosphatase 2C family protein | 5.24 | AT2G29380.1 |
| 1095 | HORVU1Hr1G092290 | receptor kinase 1 | 5.22 | AT4G23180.1 |
| 1096 | HORVU4Hr1G002270 | 5-methyltetrahydropteroyltriglutamate--homocysteine methyltransferase 2 | 5.22 | AT5G17920.2 |
| 1097 | HORVU2Hr1G121060 | polyamine oxidase 2 | 5.22 | AT1G65840.1 |
| 1098 | HORVU4Hr1G021010 | Two-component response regulator-like APRR5 | 5.20 | AT5G24470.1 |
| 1099 | HORVU5Hr1G063340 | ATP-dependent zinc metalloprotease FtsH 2 | 5.19 | AT5G57480.1 |
| 1100 | HORVU7Hr1G121700 | catalase 2 | 5.18 | AT4G35090.1 |
| 1101 | HORVU2Hr1G111600 | Adenine nucleotide alpha hydrolases-like superfamily protein | 5.17 | AT1G68300.1 |
| 1102 | HORVU3Hr1G093100 | Bifunctional inhibitor/lipid-transfer protein/seed storage 2S albumin superfamily protein | 5.16 | AT5G55410.1 |
| 1103 | HORVU1Hr1G045610 | membrane steroid binding protein 1 | 5.16 | AT5G52240.1 |
| 1104 | HORVU1Hr1G002460 | receptor-like protein kinase 4 | 5.16 | AT1G66910.1 |
| 1105 | HORVU5Hr1G019110 | Amino acid permease family protein | 5.16 | AT3G19553.1 |
| 1106 | HORVU2Hr1G072180 | prephenate dehydratase 1 | 5.13 | AT1G08250.1 |
| 1107 | HORVU5Hr1G087780 | Glutamyl-tRNA(Gln) amidotransferase subunit A | 5.12 | AT4G34880.2 |
| 1108 | HORVU6Hr1G065240 | undescribed protein | 5.12 | AT5G42900.3 |
| 1109 | HORVU5Hr1G063420 | glutamate receptor 2.8 | 5.09 | AT2G29120.1 |
| 1110 | HORVU2Hr1G121050 | polyamine oxidase 4 | 5.09 | AT1G65840.1 |
| 1111 | HORVU4Hr1G074990 | DOF zinc finger protein 1 | 5.08 | AT5G39660.3 |
| 1112 | HORVU3Hr1G004140 | Bowman-Birk type trypsin inhibitor | 5.07 | AT1G23580.1 |
| 1113 | HORVU5Hr1G096940 | Cytochrome P450 superfamily protein | 5.06 | AT4G37330.1 |
| 1114 | HORVU5Hr1G054060 | ferrochelatase 1 | 5.02 | AT5G26030.2 |
| 1115 | HORVU7Hr1G106600 | patatin-like protein 6 | 4.99 | AT3G54950.1 |
| 1116 | HORVU7Hr1G081910 | 60S ribosomal protein L37a | 4.97 | AT3G10950.1 |
| 1117 | HORVU7Hr1G002130 | receptor-like protein kinase 2 | 4.96 | AT3G47570.1 |
| 1118 | HORVU5Hr1G056370 | Domain of unknown function (DUF220) | 4.96 | AT1G70480.1 |
| 1119 | HORVU4Hr1G074450 | Alba DNA/RNA-binding protein | 4.95 | AT3G07030.3 |

| | | | | |
|---|---|---|---|---|
| 1120 | HORVU2Hr1G121070 | Adenylate kinase | 4.94 | AT2G37250.1 |
| 1121 | HORVU6Hr1G061140 | GATA transcription factor 5 | 4.92 | AT5G66320.2 |
| 1122 | HORVU4Hr1G076840 | Protein TIFY 3A | 4.92 | AT1G74950.1 |
| 1123 | HORVU4Hr1G069320 | Divalent metal cation transporter MntH | 4.92 | AT1G47240.1 |
| 1124 | HORVU1Hr1G054560 | DYNAMIN-like 1E | 4.92 | AT3G60190.1 |
| 1125 | HORVU3Hr1G082590 | Abhydrolase domain-containing protein 1 | 4.91 | AT1G34340.1 |
| 1126 | HORVU5Hr1G001920 | Leucine-rich receptor-like protein kinase family protein | 4.91 | AT1G09970.1 |
| 1127 | HORVU1Hr1G045220 | Calcium uniporter protein 2, mitochondrial | 4.89 | AT2G23790.1 |
| 1128 | HORVU3Hr1G088570 | fructose-bisphosphate aldolase 2 | 4.87 | AT2G36460.1 |
| 1129 | HORVU3Hr1G058170 | ADP,ATP carrier protein 1 | 4.84 | AT1G80300.1 |
| 1130 | HORVU1Hr1G049900 | Leucine-rich repeat family protein | 4.83 | AT4G35470.1 |
| 1131 | HORVU7Hr1G075710 | N-(5'-phosphoribosyl)anthranilate isomerase | 4.81 | AT1G07780.9 |
| 1132 | HORVU0Hr1G005210 | Protein of unknown function, DUF584 | 4.81 | AT2G28400.1 |
| 1133 | HORVU2Hr1G113260 | N-terminal nucleophile aminohydrolases (Ntn hydrolases) superfamily protein | 4.81 | AT3G16150.1 |
| 1134 | HORVU5Hr1G073990 | proton pump interactor 1 | 4.80 | AT1G20970.1 |
| 1135 | HORVU3Hr1G099630 | unknown function | 4.80 | AT3G20980.1 |
| 1136 | HORVU7Hr1G090830 | Copine family protein 2 | 4.79 | AT5G14420.4 |
| 1137 | HORVU3Hr1G067910 | Alliinase | 4.77 | AT1G34060.1 |
| 1138 | HORVU6Hr1G008930 | receptor-like protein kinase 2 | 4.76 | AT3G47570.1 |
| 1139 | HORVU6Hr1G070270 | ATP-dependent 6-phosphofructokinase 7 | 4.76 | AT1G76550.1 |
| 1140 | HORVU4Hr1G056610 | protein kinase family protein | 4.72 | AT3G01090.2 |
| 1141 | HORVU2Hr1G020680 | Protein kinase superfamily protein | 4.72 | AT1G54610.3 |
| 1142 | HORVU5Hr1G071720 | undescribed protein | 4.69 | AT5G04690.1 |
| 1143 | HORVU6Hr1G062830 | Ceramide kinase | 4.68 | AT5G51290.1 |
| 1144 | HORVU7Hr1G089960 | phospholipase D P2 | 4.68 | AT3G15730.1 |
| 1145 | HORVU3Hr1G090730 | Chromosome 3B, genomic scaffold, cultivar Chinese Spring | 4.68 | AT1G33780.1 |
| 1146 | HORVU4Hr1G072310 | FAD/NAD(P)-binding oxidoreductase family protein | 4.68 | AT4G38540.1 |
| 1147 | HORVU7Hr1G008050 | Receptor-like protein kinase | 4.67 | AT1G35710.1 |
| 1148 | HORVU7Hr1G034350 | Chalcone-flavonone isomerase family protein | 4.66 | AT2G26310.5 |
| 1149 | HORVU3Hr1G080830 | Cytochrome P450 superfamily protein | 4.66 | AT2G30490.1 |
| 1150 | HORVU2Hr1G053620 | Senescence-associated protein, putative | 4.66 | AT1G27320.1 |
| 1151 | HORVU6Hr1G059940 | GRAM domain-containing protein / ABA-responsive protein-related | 4.66 | AT5G08350.1 |
| 1152 | HORVU5Hr1G077120 | Esterase/lipase/thioesterase family protein | 4.64 | AT1G54570.1 |
| 1153 | HORVU2Hr1G036440 | Heat shock 70 kDa protein 17 | 4.63 | AT4G16660.1 |
| 1154 | HORVU6Hr1G064650 | unknown function | 4.63 | AT1G76240.1 |
| 1155 | HORVU6Hr1G065210 | transcription factor-related | 4.63 | AT2G27230.2 |
| 1156 | HORVU0Hr1G000050 | receptor-like protein kinase 4 | 4.62 | AT5G38240.2 |
| 1157 | HORVU1Hr1G045630 | membrane steroid binding protein 1 | 4.61 | AT3G48890.1 |

| | | | | |
|---|---|---|---|---|
| 1158 | HORVU2Hr1G090730 | GRAM domain-containing protein / ABA-responsive protein-related | 4.61 | AT5G23370.1 |
| 1159 | HORVU1Hr1G092300 | receptor kinase 2 | 4.59 | AT4G23190.2 |
| 1160 | HORVU5Hr1G056790 | arginine methyltransferase 11 | 4.59 | AT4G29510.1 |
| 1161 | HORVU2Hr1G118950 | receptor kinase 1 | 4.59 | AT3G09010.1 |
| 1162 | HORVU0Hr1G009430 | 0 | 4.59 | #N/A |
| 1163 | HORVU4Hr1G074770 | unknown function | 4.57 | AT1G13990.1 |
| 1164 | HORVU2Hr1G123030 | Calcium-dependent lipid-binding (CaLB domain) family protein | 4.57 | AT1G63220.1 |
| 1165 | HORVU3Hr1G090920 | NAC domain protein, | 4.56 | AT1G01720.1 |
| 1166 | HORVU2Hr1G028540 | Lipase/lipooxygenase, putative isoform 3 | 4.56 | AT1G55280.1 |
| 1167 | HORVU6Hr1G076510 | matrix metalloproteinase | 4.54 | AT1G24140.1 |
| 1168 | HORVU3Hr1G094720 | Chromosome 3B, genomic scaffold, cultivar Chinese Spring | 4.53 | AT3G01960.1 |
| 1169 | HORVU3Hr1G081300 | U-box domain-containing protein 16 | 4.52 | AT5G01830.1 |
| 1170 | HORVU5Hr1G081860 | Chaperone protein ClpB | 4.52 | AT5G51070.1 |
| 1171 | HORVU6Hr1G018370 | calnexin 1 | 4.51 | AT5G61790.1 |
| 1172 | HORVU1Hr1G047570 | undescribed protein | 4.51 | AT5G24660.1 |
| 1173 | HORVU7Hr1G055900 | Eukaryotic protein of unknown function (DUF914) | 4.51 | AT3G59330.2 |
| 1174 | HORVU5Hr1G072740 | undescribed protein | 4.50 | #N/A |
| 1175 | HORVU4Hr1G054060 | Argininosuccinate lyase | 4.50 | AT5G10920.1 |
| 1176 | HORVU2Hr1G125110 | Peroxidase superfamily protein | 4.50 | AT1G71695.1 |
| 1177 | HORVU5Hr1G043460 | calmodulin like 23 | 4.49 | AT3G10190.1 |
| 1178 | HORVU3Hr1G054640 | NAD(P)-linked oxidoreductase superfamily protein | 4.49 | AT1G60690.1 |
| 1179 | HORVU7Hr1G045140 | 39S ribosomal protein L46, mitochondrial | 4.48 | AT1G14620.1 |
| 1180 | HORVU4Hr1G079330 | Pheophorbide a oxygenase, chloroplastic | 4.48 | AT3G44880.1 |
| 1181 | HORVU2Hr1G022140 | Nucleolar protein 56 | 4.47 | AT3G12860.1 |
| 1182 | HORVU0Hr1G006610 | 0 | 4.46 | #N/A |
| 1183 | HORVU7Hr1G033390 | Stromal cell-derived factor 2-like protein | 4.46 | AT2G25110.1 |
| 1184 | HORVU6Hr1G076520 | unknown function | 4.46 | AT1G19020.1 |
| 1185 | HORVU2Hr1G091130 | Thioredoxin superfamily protein | 4.46 | AT3G15360.1 |
| 1186 | HORVU2Hr1G049420 | Transmembrane protein 41B | 4.46 | AT4G17790.1 |
| 1187 | HORVU6Hr1G053140 | Senescence-associated protein, putative | 4.46 | AT1G27320.1 |
| 1188 | HORVU4Hr1G007550 | receptor-like kinase 902 | 4.45 | AT1G48480.1 |
| 1189 | HORVU1Hr1G081550 | malate dehydrogenase | 4.44 | AT3G47520.1 |
| 1190 | HORVU2Hr1G046590 | BNR/Asp-box repeat family protein | 4.44 | AT5G57700.3 |
| 1191 | HORVU5Hr1G058070 | Major facilitator superfamily protein | 4.43 | AT1G78130.1 |
| 1192 | HORVU5Hr1G089620 | receptor kinase 2 | 4.41 | AT1G21240.1 |
| 1193 | HORVU3Hr1G100350 | Xylanase inhibitor | 4.39 | AT1G03220.1 |
| 1194 | HORVU3Hr1G081880 | Galactose-binding protein | 4.38 | AT1G22882.1 |
| 1195 | HORVU4Hr1G058340 | Cytochrome P450 superfamily protein | 4.36 | AT3G14690.1 |

| | | | | |
|---|---|---|---|---|
| 1196 | HORVU5Hr1G069090 | alpha/beta-Hydrolases superfamily protein | 4.35 | AT1G47480.1 |
| 1197 | HORVU1Hr1G025890 | Boron transporter 4 | 4.35 | AT1G15460.2 |
| 1198 | HORVU6Hr1G067430 | oligopeptide transporter 4 | 4.35 | AT5G64410.1 |
| 1199 | HORVU3Hr1G023400 | UDP-Glycosyltransferase superfamily protein | 4.34 | AT2G36800.1 |
| 1200 | HORVU7Hr1G119870 | undescribed protein | 4.33 | AT1G80245.7 |
| 1201 | HORVU1Hr1G064860 | Isopentenyl-diphosphate Delta-isomerase I | 4.33 | AT5G16440.1 |
| 1202 | HORVU3Hr1G073150 | Glucan endo-1,3-beta-glucosidase GI | 4.32 | AT3G57260.1 |
| 1203 | HORVU5Hr1G007550 | Ubiquitin carboxyl-terminal hydrolase 8 | 4.31 | AT5G22030.1 |
| 1204 | HORVU2Hr1G091270 | beta-xylosidase 2 | 4.31 | AT5G10560.1 |
| 1205 | HORVU4Hr1G022170 | unknown function | 4.30 | AT3G52710.1 |
| 1206 | HORVU6Hr1G084740 | Cytochrome c oxidase copper chaperone 2 | 4.30 | AT1G53030.1 |
| 1207 | HORVU2Hr1G043860 | histone H2A 2 | 4.30 | AT4G27230.2 |
| 1208 | HORVU6Hr1G089960 | undescribed protein | 4.29 | AT1G47890.1 |
| 1209 | HORVU2Hr1G022620 | Disease resistance protein | 4.29 | AT1G50180.1 |
| 1210 | HORVU5Hr1G013360 | 40S ribosomal protein S20 | 4.28 | AT5G62300.2 |
| 1211 | HORVU6Hr1G075590 | Kelch repeat-containing protein | 4.28 | AT1G14330.1 |
| 1212 | HORVU4Hr1G023260 | BTB/POZ domain-containing protein | 4.27 | AT2G30520.1 |
| 1213 | HORVU3Hr1G057970 | undescribed protein | 4.27 | AT1G66730.1 |
| 1214 | HORVU7Hr1G110630 | Disease resistance protein | 4.26 | AT3G07040.1 |
| 1215 | HORVU1Hr1G027700 | WRKY DNA-binding protein 50 | 4.26 | AT5G26170.1 |
| 1216 | HORVU1Hr1G049210 | Glutathione S-transferase family protein | 4.24 | AT1G10360.1 |
| 1217 | HORVU3Hr1G040710 | PENTATRICOPEPTIDE REPEAT 596 | 4.23 | AT1G80270.5 |
| 1218 | HORVU1Hr1G022810 | Protein disulfide isomerase-like 2-1 | 4.23 | AT2G47470.1 |
| 1219 | HORVU5Hr1G120570 | Membrane protein insertase YidC | 4.22 | AT1G24490.1 |
| 1220 | HORVU5Hr1G112610 | rRNA N-glycosidase | 4.22 | AT3G59490.3 |
| 1221 | HORVU5Hr1G124160 | Chlorophyll a-b binding protein 2, chloroplastic | 4.21 | AT2G34420.1 |
| 1222 | HORVU5Hr1G048890 | Carbamoyl-phosphate synthase small chain | 4.21 | AT3G27740.1 |
| 1223 | HORVU3Hr1G108900 | LORELEI-LIKE-GPI-ANCHORED PROTEIN 1 | 4.19 | AT5G56170.1 |
| 1224 | HORVU7Hr1G007540 | unknown function | 4.18 | AT1G09070.1 |
| 1225 | HORVU1Hr1G022020 | alpha/beta-Hydrolases superfamily protein | 4.18 | AT3G14360.1 |
| 1226 | HORVU3Hr1G096190 | Protein kinase family protein | 4.15 | AT1G21230.1 |
| 1227 | HORVU2Hr1G016650 | Guanine nucleotide-binding protein-like 3 homolog | 4.15 | AT3G07050.1 |
| 1228 | HORVU2Hr1G089810 | undescribed protein | 4.14 | AT1G58190.1 |
| 1229 | HORVU3Hr1G078850 | UDP-Glycosyltransferase superfamily protein | 4.14 | AT2G30150.2 |
| 1230 | HORVU1Hr1G033630 | undescribed protein | 4.14 | AT1G04985.1 |
| 1231 | HORVU3Hr1G076750 | Secretory carrier-associated membrane protein 2 | 4.13 | AT1G61250.1 |
| 1232 | HORVU5Hr1G106960 | Major facilitator superfamily protein | 4.13 | AT2G39210.1 |
| 1233 | HORVU3Hr1G090410 | Chromosome 3B, genomic scaffold, cultivar Chinese Spring | 4.13 | AT3G59880.1 |
| 1234 | HORVU5Hr1G024420 | Senescence-associated protein, putative | 4.13 | AT1G27320.1 |

| | | | | |
|---|---|---|---|---|
| 1235 | HORVU2Hr1G113460 | Cysteine proteinases superfamily protein | 4.12 | AT1G47128.1 |
| 1236 | HORVU5Hr1G026120 | Syntaxin-132 | 4.11 | AT3G03800.2 |
| 1237 | HORVU5Hr1G125130 | 60 kDa chaperonin 2 | 4.09 | AT2G28000.1 |
| 1238 | HORVU5Hr1G098090 | allene oxide synthase | 4.09 | AT5G42650.1 |
| 1239 | HORVU6Hr1G031640 | unknown function | 4.09 | AT4G39720.4 |
| 1240 | HORVU2Hr1G037750 | receptor kinase 3 | 4.07 | AT4G23180.1 |
| 1241 | HORVU1Hr1G094070 | Cytoplasmic membrane protein | 4.06 | AT4G10140.1 |
| 1242 | HORVU4Hr1G052320 | unknown function | 4.06 | AT3G56710.2 |
| 1243 | HORVU2Hr1G072830 | alpha/beta-Hydrolases superfamily protein | 4.05 | AT3G48410.1 |
| 1244 | HORVU3Hr1G006290 | Alanine--tRNA ligase | 4.05 | AT3G16565.6 |
| 1245 | HORVU7Hr1G099390 | receptor kinase 1 | 4.03 | AT4G11530.1 |
| 1246 | HORVU1Hr1G044960 | CBS domain-containing protein CBSX5 | 4.03 | AT4G27460.1 |
| 1247 | HORVU2Hr1G046550 | Protein of unknown function, DUF584 | 4.02 | AT1G61930.1 |
| 1248 | HORVU3Hr1G087740 | TRICHOME BIREFRINGENCE-LIKE 28 | 4.01 | AT3G55990.1 |
| 1249 | HORVU2Hr1G020690 | Nucleolar protein 58 | 4.01 | AT5G27120.1 |
| 1250 | HORVU5Hr1G117900 | actin 7 | 4.00 | AT5G09810.1 |
| 1251 | HORVU1Hr1G043710 | Protein of unknown function (DUF789) | 4.00 | AT1G03610.1 |
| 1252 | HORVU5Hr1G085020 | receptor kinase 1 | 3.99 | AT4G21410.1 |
| 1253 | HORVU6Hr1G068150 | Ubiquinol oxidase | 3.99 | AT3G22370.1 |
| 1254 | HORVU1Hr1G072060 | 30S ribosomal protein S8 | 3.98 | AT5G59850.1 |
| 1255 | HORVU7Hr1G045420 | Protein FATTY ACID EXPORT 1, chloroplastic | 3.97 | AT3G57280.1 |
| 1256 | HORVU3Hr1G080500 | Cytochrome P450 superfamily protein | 3.96 | AT4G13290.1 |
| 1257 | HORVU3Hr1G057550 | Single-stranded DNA-binding protein | 3.96 | AT3G18580.1 |
| 1258 | HORVU2Hr1G092820 | YELLOW STRIPE like 2 | 3.96 | AT5G24380.1 |
| 1259 | HORVU3Hr1G099620 | Chromosome 3B, genomic scaffold, cultivar Chinese Spring | 3.95 | AT3G20980.1 |
| 1260 | HORVU4Hr1G003400 | Coffea canephora DH200=94 genomic scaffold, scaffold_6 | 3.93 | AT2G01820.1 |
| 1261 | HORVU2Hr1G092880 | YELLOW STRIPE like 2 | 3.93 | AT5G24380.1 |
| 1262 | HORVU3Hr1G098400 | methyl esterase 1 | 3.92 | AT2G23620.1 |
| 1263 | HORVU7Hr1G088260 | Inositol oxygenase 1 | 3.92 | AT1G14520.4 |
| 1264 | HORVU1Hr1G021760 | BTB-POZ and MATH domain 2 | 3.91 | AT3G06190.1 |
| 1265 | HORVU7Hr1G117000 | Chaperone protein htpG family protein | 3.91 | AT4G24190.1 |
| 1266 | HORVU6Hr1G002030 | Multiple organellar RNA editing factor 5, mitochondrial | 3.91 | AT1G32580.1 |
| 1267 | HORVU5Hr1G099820 | citrate synthase 3 | 3.91 | AT3G58750.1 |
| 1268 | HORVU4Hr1G010470 | sugar transporter 9 | 3.91 | AT3G18830.1 |
| 1269 | HORVU1Hr1G015770 | receptor kinase 2 | 3.90 | AT5G40380.1 |
| 1270 | HORVU1Hr1G047390 | SAUR-like auxin-responsive protein family | 3.88 | AT2G46690.1 |
| 1271 | HORVU6Hr1G001980 | Ankyrin repeat family protein | 3.88 | AT5G07840.1 |
| 1272 | HORVU5Hr1G104850 | receptor kinase 2 | 3.87 | AT2G37710.1 |
| 1273 | HORVU5Hr1G000800 | Protein NRT1/ PTR FAMILY 2.11 | 3.86 | AT5G62680.1 |

| | | | | |
|---|---|---|---|---|
| 1274 | HORVU6Hr1G014960 | Ubiquitin family protein | 3.86 | AT4G05050.3 |
| 1275 | HORVU7Hr1G046980 | Mitochondrial glycoprotein family protein | 3.86 | AT5G02050.1 |
| 1276 | HORVU6Hr1G055250 | Isochorismatase hydrolase family protein | 3.86 | AT2G22570.1 |
| 1277 | HORVU2Hr1G056750 | Small nuclear ribonucleoprotein-associated protein | 3.85 | AT5G44500.2 |
| 1278 | HORVU5Hr1G077020 | Multiple organellar RNA editing factor 8, chloroplastic/mitochondrial | 3.85 | AT1G53260.1 |
| 1279 | HORVU3Hr1G040350 | Coatomer, beta subunit | 3.84 | AT4G31480.9 |
| 1280 | HORVU2Hr1G048220 | Transmembrane protein 184A | 3.84 | AT4G21570.1 |
| 1281 | HORVU6Hr1G057270 | Subtilisin-like protease SBT3.6 | 3.82 | AT1G71950.1 |
| 1282 | HORVU2Hr1G006140 | undescribed protein | 3.82 | #N/A |
| 1283 | HORVU6Hr1G084590 | Isopentenyl-diphosphate Delta-isomerase | 3.82 | AT1G79690.2 |
| 1284 | HORVU7Hr1G100100 | histone H2A 6 | 3.82 | AT5G02560.1 |
| 1285 | HORVU2Hr1G016720 | Protein kinase superfamily protein | 3.82 | AT1G26970.1 |
| 1286 | HORVU4Hr1G003180 | SBP (S-ribonuclease binding protein) family protein | 3.82 | AT3G12920.1 |
| 1287 | HORVU2Hr1G075010 | nicotinate phosphoribosyltransferase 2 | 3.82 | AT2G23420.1 |
| 1288 | HORVU4Hr1G003060 | Histone-lysine N-methyltransferase | 3.81 | AT4G00570.1 |
| 1289 | HORVU6Hr1G014910 | D.melanogaster polytene | 3.80 | AT1G65349.1 |
| 1290 | HORVU3Hr1G087680 | Phytol kinase 1, chloroplastic | 3.80 | AT5G58560.1 |
| 1291 | HORVU0Hr1G013910 | undescribed protein | 3.77 | AT3G29060.1 |
| 1292 | HORVU2Hr1G010560 | Thioredoxin superfamily protein | 3.77 | AT1G20225.1 |
| 1293 | HORVU1Hr1G068030 | Cytochrome P450 superfamily protein | 3.77 | AT2G40890.1 |
| 1294 | HORVU5Hr1G008300 | 30S ribosomal protein S10 | 3.76 | AT5G62300.2 |
| 1295 | HORVU2Hr1G056490 | undescribed protein | 3.76 | AT2G40030.1 |
| 1296 | HORVU5Hr1G008820 | p21-activated protein kinase-interacting protein 1-like | 3.75 | AT1G65030.1 |
| 1297 | HORVU0Hr1G022290 | receptor kinase 2 | 3.75 | AT5G10530.1 |
| 1298 | HORVU2Hr1G028510 | Ribosomal protein S4 | 3.74 | AT5G15200.1 |
| 1299 | HORVU1Hr1G000140 | Phospholipid-transporting ATPase 10 | 3.74 | AT1G68710.2 |
| 1300 | HORVU5Hr1G071270 | F-box and associated interaction domains-containing protein | 3.74 | AT3G22730.1 |
| 1301 | HORVU6Hr1G063560 | S-adenosylmethionine synthase 2 | 3.73 | AT3G17390.1 |
| 1302 | HORVU3Hr1G024210 | ABC transporter C family member 3 | 3.73 | AT3G13080.5 |
| 1303 | HORVU6Hr1G075400 | Exocyst complex component SEC6 | 3.73 | AT1G71820.1 |
| 1304 | HORVU3Hr1G083830 | 29 kDa ribonucleoprotein B, chloroplastic | 3.73 | AT2G37220.1 |
| 1305 | HORVU3Hr1G116740 | myb domain protein 59 | 3.72 | AT5G59780.3 |
| 1306 | HORVU5Hr1G098190 | Cold acclimation protein WCOR413 family | 3.72 | AT3G50830.1 |
| 1307 | HORVU7Hr1G048810 | Transmembrane 9 superfamily member 12 | 3.70 | AT4G12650.1 |
| 1308 | HORVU1Hr1G037760 | Divalent ion symporter | 3.70 | AT1G02260.2 |
| 1309 | HORVU5Hr1G068230 | bZIP transcription factor 27 | 3.69 | AT3G56850.1 |
| 1310 | HORVU5Hr1G054180 | Phosphoenolpyruvate/phosphate translocator 1, chloroplastic | 3.69 | AT5G33320.1 |
| 1311 | HORVU7Hr1G112980 | DNA-directed RNA polymerase 2A | 3.69 | AT2G24120.1 |

| | | | | |
|---|---|---|---|---|
| 1312 | HORVU1Hr1G062900 | Argininosuccinate synthase | 3.68 | AT4G24830.1 |
| 1313 | HORVU2Hr1G007570 | S-adenosylmethionine-dependent methyltransferase | 3.68 | AT2G41380.1 |
| 1314 | HORVU2Hr1G022490 | ATP-dependent RNA helicase DeaD | 3.67 | AT5G60990.1 |
| 1315 | HORVU3Hr1G096550 | glutamine cyclotransferase | 3.67 | AT4G25720.1 |
| 1316 | HORVU4Hr1G084930 | TVP38/TMEM64 family membrane protein slr0305 | 3.66 | AT2G02370.3 |
| 1317 | HORVU2Hr1G087090 | adenine phosphoribosyltransferase 5 | 3.65 | AT1G80050.1 |
| 1318 | HORVU5Hr1G080170 | ATP-dependent RNA helicase DeaD | 3.64 | AT5G62190.1 |
| 1319 | HORVU6Hr1G071780 | Transmembrane 9 superfamily member 12 | 3.64 | AT4G12650.1 |
| 1320 | HORVU1Hr1G074590 | aspartate aminotransferase | 3.63 | AT2G22250.3 |
| 1321 | HORVU7Hr1G000850 | RING/U-box superfamily protein | 3.62 | AT3G19950.3 |
| 1322 | HORVU3Hr1G002920 | Glycosyltransferase | 3.62 | AT3G18180.1 |
| 1323 | HORVU3Hr1G080940 | unknown function | 3.62 | AT1G47970.1 |
| 1324 | HORVU7Hr1G028290 | Cytochrome P450 superfamily protein | 3.62 | AT3G26300.1 |
| 1325 | HORVU6Hr1G090810 | Acetyl-coenzyme A synthetase | 3.62 | AT5G36880.2 |
| 1326 | HORVU5Hr1G123490 | Protein BCCIP homolog | 3.62 | AT2G44510.1 |
| 1327 | HORVU3Hr1G052980 | Chromosome 3B, genomic scaffold, cultivar Chinese Spring | 3.60 | AT2G01410.1 |
| 1328 | HORVU6Hr1G016150 | ubiquitin 4 | 3.59 | AT4G05050.3 |
| 1329 | HORVU7Hr1G085180 | UDP-galactose transporter 3 | 3.59 | AT1G14360.1 |
| 1330 | HORVU2Hr1G031700 | unknown function | 3.58 | AT4G30140.1 |
| 1331 | HORVU7Hr1G013710 | 3-phosphoshikimate 1-carboxyvinyltransferase 2 | 3.58 | AT2G45300.4 |
| 1332 | HORVU3Hr1G038060 | Glycosyl hydrolase, family 31 | 3.56 | AT2G44970.3 |
| 1333 | HORVU1Hr1G044020 | myosin XI B | 3.56 | AT1G04160.1 |
| 1334 | HORVU2Hr1G085160 | inositol transporter 1 | 3.55 | AT2G43330.1 |
| 1335 | HORVU2Hr1G005520 | Stress responsive A/B Barrel Domain | 3.55 | AT3G17210.2 |
| 1336 | HORVU2Hr1G005990 | Nucleotide-diphospho-sugar transferase family protein | 3.55 | AT4G19975.1 |
| 1337 | HORVU5Hr1G118010 | RNA-binding protein 1 | 3.55 | AT3G26420.1 |
| 1338 | HORVU5Hr1G027590 | Zinc finger CCCH domain-containing protein 67 | 3.54 | AT2G41900.1 |
| 1339 | HORVU3Hr1G066810 | Cytokinin riboside 5'-monophosphate phosphoribohydrolase LOG8 | 3.53 | AT2G35990.1 |
| 1340 | HORVU3Hr1G083560 | Uricase | 3.53 | AT2G26230.1 |
| 1341 | HORVU5Hr1G028030 | sugar transporter 9 | 3.53 | AT3G18830.1 |
| 1342 | HORVU5Hr1G044780 | Glutamine--fructose-6-phosphate aminotransferase [isomerizing] 2 | 3.52 | AT3G24090.1 |
| 1343 | HORVU3Hr1G073220 | aspartate aminotransferase 3 | 3.52 | AT5G11520.1 |
| 1344 | HORVU3Hr1G023780 | HXXXD-type acyl-transferase family protein | 3.52 | AT5G42830.1 |
| 1345 | HORVU5Hr1G113180 | Heat shock 70 kDa protein 3 | 3.52 | AT5G02500.1 |
| 1346 | HORVU3Hr1G092660 | Dolichyl-diphosphooligosaccharide--protein glycosyltransferase subunit 2 | 3.52 | AT4G21150.3 |
| 1347 | HORVU6Hr1G091860 | rRNA/tRNA 2'-O-methyltransferase fibrillarin-like protein 1 | 3.51 | AT5G52470.1 |

| | | | | |
|---|---|---|---|---|
| 1348 | HORVU6Hr1G021700 | polyubiquitin 3 | 3.51 | AT5G20620.1 |
| 1349 | HORVU5Hr1G011460 | ATP-dependent RNA helicase DeaD | 3.51 | AT3G22330.1 |
| 1350 | HORVU4Hr1G047580 | H/ACA ribonucleoprotein complex subunit 1-like protein 2 | 3.51 | AT3G03920.1 |
| 1351 | HORVU4Hr1G085800 | flowering promoting factor 1 | 3.50 | AT5G10625.1 |
| 1352 | HORVU4Hr1G043760 | Leucine-rich repeat receptor-like protein kinase family protein | 3.50 | AT1G34420.1 |
| 1353 | HORVU5Hr1G096480 | RING/U-box superfamily protein | 3.50 | AT1G72310.1 |
| 1354 | HORVU3Hr1G019480 | myb-like transcription factor family protein | 3.50 | AT5G56840.1 |
| 1355 | HORVU6Hr1G016200 | ubiquitin 4 | 3.49 | AT4G05320.7 |
| 1356 | HORVU5Hr1G001090 | BEL1-like homeodomain 6 | 3.49 | AT4G34610.4 |
| 1357 | HORVU3Hr1G006330 | Signal peptidase complex subunit 3A | 3.48 | AT3G05230.1 |
| 1358 | HORVU6Hr1G018460 | unknown function | 3.48 | AT1G48120.1 |
| 1359 | HORVU5Hr1G109910 | 30S ribosomal protein S5 | 3.47 | AT3G57490.1 |
| 1360 | HORVU7Hr1G100230 | Heavy metal transport/detoxification superfamily protein | 3.47 | AT1G66240.1 |
| 1361 | HORVU7Hr1G012700 | Lung seven transmembrane receptor family protein | 3.46 | AT5G42090.1 |
| 1362 | HORVU5Hr1G061140 | Dihydrolipoyllysine-residue acetyltransferase component of pyruvate dehydrogenase complex | 3.46 | AT3G25860.1 |
| 1363 | HORVU4Hr1G003100 | aminopeptidase M1 | 3.46 | AT4G33090.1 |
| 1364 | HORVU7Hr1G051070 | Aminoacylase-1 | 3.46 | AT4G38220.1 |
| 1365 | HORVU1Hr1G050330 | undescribed protein | 3.45 | AT3G01990.8 |
| 1366 | HORVU4Hr1G009560 | Sulfite exporter TauE/SafE family protein | 3.45 | AT2G36630.1 |
| 1367 | HORVU1Hr1G073040 | Phosphatidylinositol:ceramide inositolphosphotransferase | 3.45 | AT3G54020.1 |
| 1368 | HORVU4Hr1G064870 | Protein NRT1/ PTR FAMILY 8.3 | 3.44 | AT2G02040.1 |
| 1369 | HORVU2Hr1G110110 | serine hydroxymethyltransferase 4 | 3.44 | AT4G13930.1 |
| 1370 | HORVU3Hr1G024500 | Glutathione S-transferase family protein | 3.44 | AT1G59700.1 |
| 1371 | HORVU7Hr1G036000 | Dipeptidyl peptidase family member 6 | 3.44 | AT5G36210.1 |
| 1372 | HORVU4Hr1G066310 | Wall-associated receptor kinase-like 14 | 3.44 | AT4G00955.1 |
| 1373 | HORVU7Hr1G010320 | Protein MID1-COMPLEMENTING ACTIVITY 2 | 3.44 | AT2G17780.4 |
| 1374 | HORVU7Hr1G108600 | DUF21 domain-containing protein | 3.43 | AT4G14240.1 |
| 1375 | HORVU6Hr1G058740 | cryptochrome 2 | 3.42 | AT1G04400.2 |
| 1376 | HORVU4Hr1G062740 | 8-amino-7-oxononanoate synthase | 3.42 | AT4G36480.2 |
| 1377 | HORVU7Hr1G077630 | F-box family protein | 3.41 | AT5G39450.1 |
| 1378 | HORVU2Hr1G043900 | Xylanase inhibitor protein 1 | 3.41 | AT5G24090.1 |
| 1379 | HORVU3Hr1G067130 | calcineurin B-like protein 10 | 3.41 | AT4G33000.1 |
| 1380 | HORVU5Hr1G067570 | Protein disulfide isomerase-like 2-3 | 3.41 | AT1G04980.1 |
| 1381 | HORVU1Hr1G088880 | Chlorophyll a-b binding protein, chloroplastic | 3.38 | AT2G34420.1 |
| 1382 | HORVU5Hr1G075490 | Peptidyl-prolyl cis-trans isomerase | 3.38 | AT3G25220.1 |
| 1383 | HORVU2Hr1G120150 | 2-succinylbenzoate--CoA ligase | 3.37 | AT5G16370.1 |
| 1384 | HORVU0Hr1G027680 | Metal-dependent protein hydrolase | 3.37 | AT5G41970.1 |

| | | | | |
|---|---|---|---|---|
| 1385 | HORVU1Hr1G064210 | Cold shock protein 1 | 3.37 | AT4G36020.2 |
| 1386 | HORVU5Hr1G026780 | Auxin-binding protein 4 | 3.37 | AT4G02980.1 |
| 1387 | HORVU5Hr1G120600 | ER lumen protein retaining receptor family protein | 3.35 | AT3G25040.1 |
| 1388 | HORVU1Hr1G008230 | dual specificity protein phosphatase 1 | 3.35 | AT3G55270.1 |
| 1389 | HORVU7Hr1G018850 | unknown function | 3.35 | AT4G00850.2 |
| 1390 | HORVU3Hr1G049610 | Carbamoyl-phosphate synthase large chain | 3.34 | AT1G29900.1 |
| 1391 | HORVU3Hr1G040270 | Coatomer, beta subunit | 3.34 | AT4G31480.6 |
| 1392 | HORVU3Hr1G085640 | Chromosome 3B, genomic scaffold, cultivar Chinese Spring | 3.34 | AT3G54000.3 |
| 1393 | HORVU1Hr1G074110 | histone deacetylase 8 | 3.34 | AT1G08460.1 |
| 1394 | HORVU4Hr1G046230 | Acyl carrier protein | 3.34 | AT1G65290.1 |
| 1395 | HORVU7Hr1G009730 | undescribed protein | 3.34 | AT1G66930.2 |
| 1396 | HORVU7Hr1G031850 | 60S ribosomal protein L35a-3 | 3.32 | AT1G74270.1 |
| 1397 | HORVU7Hr1G035340 | 3-phosphoserine phosphatase | 3.32 | AT1G18640.2 |
| 1398 | HORVU6Hr1G080000 | SAUR-like auxin-responsive protein family | 3.32 | AT2G46690.1 |
| 1399 | HORVU2Hr1G029870 | Phospho-2-dehydro-3-deoxyheptonate aldolase 2, chloroplastic | 3.31 | AT1G22410.1 |
| 1400 | HORVU3Hr1G069200 | BnaA08g17720D protein | 3.31 | AT4G12543.1 |
| 1401 | HORVU3Hr1G045480 | dehydroquinate dehydratase, putative / shikimate dehydrogenase, putative | 3.31 | AT3G06350.1 |
| 1402 | HORVU7Hr1G033360 | 2-oxoglutarate (2OG) and Fe(II)-dependent oxygenase superfamily protein | 3.30 | AT4G10490.1 |
| 1403 | HORVU1Hr1G013820 | isovaleryl-CoA-dehydrogenase | 3.30 | AT3G45300.1 |
| 1404 | HORVU5Hr1G072480 | Protein kinase family protein | 3.30 | AT1G21240.1 |
| 1405 | HORVU4Hr1G075710 | 60S ribosomal protein L4-1 | 3.29 | AT3G09630.1 |
| 1406 | HORVU2Hr1G005140 | Glyoxylate/hydroxypyruvate reductase B | 3.29 | AT2G45630.2 |
| 1407 | HORVU3Hr1G078650 | Nucleotide-sugar transporter family protein | 3.29 | AT4G32390.1 |
| 1408 | HORVU7Hr1G113350 | NADPH HC toxin reductase | 3.28 | AT4G27250.1 |
| 1409 | HORVU6Hr1G011970 | Calcium-binding EF-hand family protein | 3.28 | AT2G46600.1 |
| 1410 | HORVU4Hr1G038960 | Alpha-1,4-glucan-protein synthase [UDP-forming] | 3.27 | AT5G15650.1 |
| 1411 | HORVU5Hr1G067110 | Ribosomal L5P family protein | 3.27 | ATMG00210.1 |
| 1412 | HORVU3Hr1G066720 | Glycylpeptide N-tetradecanoyltransferase 1 | 3.27 | AT5G57020.1 |
| 1413 | HORVU2Hr1G078350 | Peptidyl-prolyl cis-trans isomerase | 3.27 | AT4G25340.1 |
| 1414 | HORVU3Hr1G055650 | loricrin-related | 3.27 | AT1G64140.1 |
| 1415 | HORVU1Hr1G083710 | Guanine nucleotide-binding protein subunit beta-like protein A | 3.27 | AT1G18080.1 |
| 1416 | HORVU1Hr1G080040 | ADP,ATP carrier protein 2, mitochondrial | 3.27 | AT5G56450.1 |
| 1417 | HORVU7Hr1G056980 | 6-phosphogluconolactonase 5 | 3.26 | AT5G24400.1 |
| 1418 | HORVU4Hr1G071360 | Cytochrome b561 and DOMON domain-containing protein | 3.25 | AT3G25290.2 |
| 1419 | HORVU7Hr1G074660 | Transmembrane amino acid transporter family protein | 3.24 | AT5G40780.1 |

| | | | | |
|---|---|---|---|---|
| 1420 | HORVU0Hr1G004790 | Unplaced genomic scaffold supercont1.2, whole genome shotgun sequence | 3.24 | AT1G21370.2 |
| 1421 | HORVU7Hr1G049370 | heat shock protein 81-3 | 3.24 | AT5G56010.1 |
| 1422 | HORVU4Hr1G028310 | myb domain protein 59 | 3.24 | AT5G59780.3 |
| 1423 | HORVU1Hr1G013990 | Exosome complex component RRP41 homolog | 3.24 | AT3G61620.2 |
| 1424 | HORVU2Hr1G060680 | Transcription factor PIF5 | 3.23 | AT2G20180.7 |
| 1425 | HORVU5Hr1G106460 | Diphosphomevalonate decarboxylase | 3.23 | AT3G54250.1 |
| 1426 | HORVU3Hr1G108720 | NADPH:quinone oxidoreductase | 3.23 | AT3G27890.1 |
| 1427 | HORVU2Hr1G086380 | hydroxycinnamoyl-CoA shikimate/quinate hydroxycinnamoyl transferase | 3.21 | AT5G48930.1 |
| 1428 | HORVU1Hr1G048970 | phospholipase D P2 | 3.20 | AT2G42010.1 |
| 1429 | HORVU5Hr1G056670 | Leucine-rich repeat transmembrane protein kinase | 3.20 | AT1G07650.4 |
| 1430 | HORVU7Hr1G106230 | Exonuclease 3'-5' domain-containing protein 2 | 3.20 | AT5G01830.1 |
| 1431 | HORVU2Hr1G031910 | Stress responsive alpha-beta barrel domain protein | 3.20 | AT2G31670.1 |
| 1432 | HORVU1Hr1G030390 | Abscisic acid receptor PYR1 | 3.20 | AT5G53160.1 |
| 1433 | HORVU4Hr1G004460 | Protein NRT1/ PTR FAMILY 2.9 | 3.19 | AT3G47960.1 |
| 1434 | HORVU1Hr1G015660 | bZIP transcription factor 53 | 3.19 | AT3G62420.1 |
| 1435 | HORVU5Hr1G122350 | Elongation factor Tu | 3.19 | AT4G02930.1 |
| 1436 | HORVU1Hr1G023510 | GDSL esterase/lipase | 3.19 | AT1G56670.1 |
| 1437 | HORVU3Hr1G092520 | Chromosome 3B, genomic scaffold, cultivar Chinese Spring | 3.18 | AT4G34900.1 |
| 1438 | HORVU6Hr1G021520 | Peroxidase superfamily protein | 3.18 | AT5G05340.1 |
| 1439 | HORVU1Hr1G083330 | Quinolone resistance protein NorA | 3.17 | AT5G10190.1 |
| 1440 | HORVU5Hr1G066930 | Superoxide dismutase [Cu-Zn] 4A | 3.15 | AT1G08830.2 |
| 1441 | HORVU7Hr1G111600 | Protein transport protein Sec61 subunit gamma | 3.15 | AT5G50460.1 |
| 1442 | HORVU2Hr1G083940 | glucose-6-phosphate dehydrogenase 6 | 3.15 | AT5G40760.2 |
| 1443 | HORVU5Hr1G036790 | O-fucosyltransferase family protein | 3.15 | AT1G35510.1 |
| 1444 | HORVU7Hr1G048330 | Alpha-glucan water dikinase 2 | 3.14 | AT1G10760.3 |
| 1445 | HORVU5Hr1G089780 | Molybdenum cofactor sulfurase family protein | 3.13 | AT1G30910.1 |
| 1446 | HORVU3Hr1G106140 | Glucan endo-1,3-beta-glucosidase GI | 3.12 | AT4G16260.1 |
| 1447 | HORVU4Hr1G008700 | glutamate decarboxylase | 3.12 | AT5G17330.1 |
| 1448 | HORVU7Hr1G033870 | zinc-finger protein 3 | 3.11 | AT1G27730.1 |
| 1449 | HORVU3Hr1G014730 | Translocon-associated protein subunit beta | 3.11 | AT5G14030.4 |
| 1450 | HORVU5Hr1G023530 | Calmodulin-binding protein | 3.11 | AT5G57580.1 |
| 1451 | HORVU3Hr1G077000 | Adenine nucleotide alpha hydrolases-like superfamily protein | 3.11 | AT1G11360.4 |
| 1452 | HORVU2Hr1G006250 | Dihydrolipoyllysine-residue succinyltransferase component of 2-oxoglutarate dehydrogenase complex | 3.10 | AT5G55070.2 |
| 1453 | HORVU4Hr1G058840 | Arginine biosynthesis bifunctional protein ArgJ 1 | 3.10 | AT2G37500.1 |
| 1454 | HORVU6Hr1G006560 | 1-deoxy-D-xylulose 5-phosphate synthase 3 | 3.10 | AT1G55510.1 |

| | | | | |
|---|---|---|---|---|
| 1455 | HORVU5Hr1G110180 | phosphate transporter 1;7 | 3.09 | AT3G54700.2 |
| 1456 | HORVU6Hr1G072910 | Eukaryotic aspartyl protease family protein | 3.09 | AT1G49050.1 |
| 1457 | HORVU1Hr1G003600 | Protein of unknown function (DUF707) | 3.08 | AT1G67850.4 |
| 1458 | HORVU3Hr1G011460 | Mitochondrial glycoprotein family protein | 3.08 | AT5G05990.1 |
| 1459 | HORVU3Hr1G070270 | Amino-acid acetyltransferase | 3.08 | AT4G37670.2 |
| 1460 | HORVU7Hr1G018190 | GDP-D-mannose 4,6-dehydratase 1 | 3.07 | AT3G51160.1 |
| 1461 | HORVU5Hr1G028550 | ureide permease 1 | 3.07 | AT2G03530.5 |
| 1462 | HORVU1Hr1G025670 | Auxin-responsive protein IAA15 | 3.07 | AT2G22670.4 |
| 1463 | HORVU4Hr1G009040 | Histone H2B.1 | 3.06 | AT3G45980.1 |
| 1464 | HORVU1Hr1G028200 | hexokinase 2 | 3.06 | AT4G29130.1 |
| 1465 | HORVU7Hr1G028910 | Pyrophosphate-energized proton pump 1 | 3.05 | AT1G15690.1 |
| 1466 | HORVU2Hr1G068670 | Acyl-[acyl-carrier-protein] 6-desaturase | 3.05 | AT2G43710.1 |
| 1467 | HORVU3Hr1G084170 | Glucan endo-1,3-beta-glucosidase 14 | 3.04 | AT2G27500.2 |
| 1468 | HORVU5Hr1G028870 | GRF1-interacting factor 3 | 3.04 | AT4G00850.1 |
| 1469 | HORVU5Hr1G095140 | Chromosome 3B, genomic scaffold, cultivar Chinese Spring | 3.04 | AT5G56050.1 |
| 1470 | HORVU3Hr1G090500 | DOF zinc finger protein 1 | 3.04 | AT3G55370.1 |
| 1471 | HORVU4Hr1G074270 | 40S ribosomal protein S27 | 3.03 | AT3G61110.1 |
| 1472 | HORVU7Hr1G071240 | 60S ribosomal protein L34 | 3.03 | AT1G26880.1 |
| 1473 | HORVU6Hr1G016180 | undescribed protein | 3.03 | AT1G16360.2 |
| 1474 | HORVU7Hr1G008690 | Protein kinase superfamily protein | 3.03 | AT4G19110.1 |
| 1475 | HORVU6Hr1G002280 | undescribed protein | 3.03 | AT3G62960.1 |
| 1476 | HORVU7Hr1G091740 | post-illumination chlorophyll fluorescence increase | 3.02 | AT3G15840.1 |
| 1477 | HORVU3Hr1G094450 | Mannosyl-oligosaccharide glucosidase | 3.02 | AT1G67490.1 |
| 1478 | HORVU3Hr1G002280 | sucrose phosphate synthase 1F | 3.02 | AT5G20280.1 |
| 1479 | HORVU4Hr1G014130 | 40S ribosomal protein S27 | 3.02 | AT3G61110.1 |
| 1480 | HORVU4Hr1G000850 | Calcium-dependent lipid-binding (CaLB domain) family protein | 3.02 | AT5G23950.2 |
| 1481 | HORVU1Hr1G062940 | peroxisomal adenine nucleotide carrier 1 | 3.01 | AT3G05290.1 |
| 1482 | HORVU5Hr1G043190 | 2-isopropylmalate synthase 1 | 3.01 | AT1G74040.1 |
| 1483 | HORVU4Hr1G052210 | calmodulin 5 | 3.01 | AT3G43810.2 |
| 1484 | HORVU2Hr1G037940 | 40S ribosomal protein S13-1 | 3.01 | AT4G00100.1 |
| 1485 | HORVU2Hr1G089490 | Protein of unknown function (DUF3317) | 3.00 | AT1G06515.2 |
| 1486 | HORVU4Hr1G064710 | Transmembrane 9 superfamily member 3 | 3.00 | AT2G01970.1 |
| 1487 | HORVU2Hr1G005320 | arginase | 3.00 | AT4G08900.1 |
| 1488 | HORVU1Hr1G052040 | polyol/monosaccharide transporter 5 | 3.00 | AT4G35300.8 |
| 1489 | HORVU3Hr1G064080 | zinc finger protein-related | -3.01 | AT3G18290.1 |
| 1490 | HORVU7Hr1G115990 | Disease resistance protein | -3.01 | AT3G07040.1 |
| 1491 | HORVU6Hr1G060720 | cytochrome B561-1 | -3.01 | AT4G25570.2 |
| 1492 | HORVU0Hr1G004760 | 0 | -3.01 | #N/A |
| 1493 | HORVU5Hr1G070160 | Glycogen debranching enzyme | -3.02 | AT4G09020.1 |

| | | | | |
|---|---|---|---|---|
| 1494 | HORVU1Hr1G004430 | NAD(P)-binding Rossmann-fold superfamily protein | -3.02 | AT2G37660.1 |
| 1495 | HORVU2Hr1G118740 | allantoinase | -3.02 | AT4G04955.1 |
| 1496 | HORVU0Hr1G022500 | Ubiquitin system component Cue protein | -3.02 | AT5G32440.3 |
| 1497 | HORVU4Hr1G012240 | Histidine triad (HIT) protein | -3.02 | AT5G48545.1 |
| 1498 | HORVU2Hr1G111840 | Glutathione S-transferase family protein | -3.03 | AT1G10360.1 |
| 1499 | HORVU5Hr1G018840 | unknown protein; LOCATED IN: chloroplast; EXPRESSED IN: 24 plant structures; EXPRESSED DURING: 15 growth stages; BEST Arabidopsis thaliana protein match is: unknown protein . | -3.03 | AT2G14910.1 |
| 1500 | HORVU4Hr1G050230 | Pentatricopeptide repeat-containing protein | -3.03 | AT1G62680.2 |
| 1501 | HORVU5Hr1G079600 | undescribed protein | -3.03 | AT3G57300.2 |
| 1502 | HORVU6Hr1G047770 | chaperone protein dnaJ-related | -3.03 | AT5G06130.2 |
| 1503 | HORVU2Hr1G035220 | fatty acid desaturase 6 | -3.03 | AT4G30950.1 |
| 1504 | HORVU5Hr1G118340 | nicotinate phosphoribosyltransferase 1 | -3.04 | AT4G36940.1 |
| 1505 | HORVU2Hr1G047020 | undescribed protein | -3.04 | #N/A |
| 1506 | HORVU5Hr1G020200 | RNA-binding protein 47 | -3.04 | AT4G00830.5 |
| 1507 | HORVU4Hr1G071050 | alpha/beta-Hydrolases superfamily protein | -3.04 | AT1G29840.1 |
| 1508 | HORVU4Hr1G078610 | Mitochondrial import inner membrane translocase subunit Tim17/Tim22/Tim23 family protein | -3.05 | AT1G34630.1 |
| 1509 | HORVU7Hr1G107330 | SAUR-like auxin-responsive protein family | -3.05 | AT1G56150.1 |
| 1510 | HORVU3Hr1G030210 | Abscisic acid receptor PYR1 | -3.05 | AT2G26040.1 |
| 1511 | HORVU1Hr1G041290 | undescribed protein | -3.05 | AT1G16400.1 |
| 1512 | HORVU4Hr1G077420 | receptor kinase 2 | -3.05 | AT1G53730.1 |
| 1513 | HORVU5Hr1G074200 | serine/threonine protein phosphatase 2A | -3.06 | AT3G58500.1 |
| 1514 | HORVU6Hr1G035260 | 30S ribosomal protein S21 | -3.06 | AT3G27160.2 |
| 1515 | HORVU5Hr1G078960 | CDGSH iron-sulfur domain-containing protein 2 | -3.06 | AT5G51720.1 |
| 1516 | HORVU7Hr1G116600 | unknown function | -3.06 | AT3G19810.2 |
| 1517 | HORVU4Hr1G069390 | Protein kinase superfamily protein | -3.07 | AT5G67380.1 |
| 1518 | HORVU7Hr1G118440 | Disease resistance protein | -3.07 | AT3G07040.1 |
| 1519 | HORVU3Hr1G056580 | Ribosomal protein L31 | -3.07 | AT1G75350.1 |
| 1520 | HORVU6Hr1G014700 | calcium-dependent protein kinase 28 | -3.07 | AT5G66210.2 |
| 1521 | HORVU6Hr1G069190 | monodehydroascorbate reductase 4 | -3.07 | AT3G27820.1 |
| 1522 | HORVU5Hr1G120210 | Ras-related protein Rab-25 | -3.07 | AT1G09630.1 |
| 1523 | HORVU4Hr1G059050 | Sterol 3-beta-glucosyltransferase UGT80A2 | -3.08 | AT3G07020.1 |
| 1524 | HORVU6Hr1G074830 | undescribed protein | -3.08 | AT1G50000.1 |
| 1525 | HORVU6Hr1G034490 | receptor kinase 2 | -3.08 | AT2G33580.1 |
| 1526 | HORVU5Hr1G100140 | Photosystem I reaction center subunit III | -3.08 | AT1G31330.1 |
| 1527 | HORVU6Hr1G067890 | Guanosine-3',5'-bis(diphosphate) 3'-pyrophosphohydrolase | -3.09 | AT1G54130.1 |
| 1528 | HORVU6Hr1G079070 | unknown function | -3.10 | AT4G19100.1 |
| 1529 | HORVU2Hr1G019650 | F-actin-capping protein subunit alpha | -3.10 | AT3G05520.1 |

| | | | | |
|---|---|---|---|---|
| 1530 | HORVU2Hr1G123610 | undescribed protein | -3.10 | AT3G26040.1 |
| 1531 | HORVU7Hr1G013600 | Transketolase | -3.11 | AT3G60750.1 |
| 1532 | HORVU2Hr1G072010 | Mitochondrial import inner membrane translocase subunit Tim17/Tim22/Tim23 family protein | -3.11 | AT5G24650.1 |
| 1533 | HORVU1Hr1G082380 | Hexosyltransferase | -3.11 | AT5G47780.1 |
| 1534 | HORVU3Hr1G096780 | Kelch repeat-containing protein | -3.12 | AT1G67480.3 |
| 1535 | HORVU6Hr1G035560 | unknown protein; Has 83 Blast hits to 82 proteins in 37 species: Archae - 0; Bacteria - 51; Metazoa - 0; Fungi - 0; Plants - 30; Viruses - 0; Other Eukaryotes - 2 (source: NCBI BLink). | -3.12 | AT4G13150.5 |
| 1536 | HORVU5Hr1G040400 | TLD domain-containing protein 1 | -3.12 | AT5G06260.3 |
| 1537 | HORVU2Hr1G079840 | Lipase/lipooxygenase, PLAT/LH2 family protein | -3.12 | AT4G39730.1 |
| 1538 | HORVU6Hr1G013620 | transferases, transferring glycosyl groups | -3.12 | AT4G14100.1 |
| 1539 | HORVU1Hr1G021800 | NAC domain protein, | -3.12 | AT1G69490.1 |
| 1540 | HORVU3Hr1G073480 | Chromosome 3B, genomic scaffold, cultivar Chinese Spring | -3.12 | AT5G48960.1 |
| 1541 | HORVU6Hr1G090380 | Haloacid dehalogenase-like hydrolase (HAD) superfamily protein | -3.13 | AT2G33255.1 |
| 1542 | HORVU3Hr1G024540 | unknown protein; BEST Arabidopsis thaliana protein match is: unknown protein . | -3.14 | AT5G25360.3 |
| 1543 | HORVU6Hr1G095270 | MATE efflux family protein | -3.15 | AT4G38380.5 |
| 1544 | HORVU7Hr1G092460 | Alpha/beta hydrolase domain-containing protein 13 | -3.15 | AT3G30380.2 |
| 1545 | HORVU3Hr1G064730 | Threonine synthase | -3.15 | AT4G29840.1 |
| 1546 | HORVU1Hr1G010820 | Ribonuclease II/R family protein | -3.15 | AT5G02250.1 |
| 1547 | HORVU7Hr1G026580 | BnaC07g25690D protein | -3.15 | AT3G29240.2 |
| 1548 | HORVU3Hr1G066420 | Regulator of Vps4 activity in the MVB pathway protein | -3.15 | AT1G34220.1 |
| 1549 | HORVU7Hr1G046280 | Thioredoxin domain-containing protein 17 | -3.15 | AT5G42850.2 |
| 1550 | HORVU6Hr1G017710 | undescribed protein | -3.16 | AT4G22160.2 |
| 1551 | HORVU3Hr1G006270 | Chromosome 3B, genomic scaffold, cultivar Chinese Spring | -3.16 | AT1G18060.1 |
| 1552 | HORVU3Hr1G004320 | MULTISPECIES: hypothetical protein | -3.16 | AT5G19730.1 |
| 1553 | HORVU3Hr1G115840 | Dual-specificity RNA methyltransferase RlmN | -3.16 | AT2G39670.1 |
| 1554 | HORVU5Hr1G118070 | undescribed protein | -3.17 | AT3G49670.1 |
| 1555 | HORVU3Hr1G029200 | phosphoenolpyruvate carboxylase 3 | -3.17 | AT3G14940.2 |
| 1556 | HORVU6Hr1G043170 | FAR1-related sequence 5 | -3.17 | AT4G38180.1 |
| 1557 | HORVU5Hr1G053950 | Bifunctional aspartokinase/homoserine dehydrogenase 1 | -3.17 | AT4G19710.2 |
| 1558 | HORVU4Hr1G072930 | Photosynthetic NDH subunit of subcomplex B 2, chloroplastic | -3.17 | AT1G64770.1 |
| 1559 | HORVU6Hr1G016240 | Transcription factor HBP-1a | -3.17 | AT2G35530.1 |
| 1560 | HORVU5Hr1G103750 | unknown function | -3.18 | AT3G10650.2 |
| 1561 | HORVU6Hr1G042840 | Monosaccharide-sensing protein 2 | -3.18 | AT4G35300.5 |

| | | | | |
|---|---|---|---|---|
| 1562 | HORVU4Hr1G021190 | Late embryogenesis abundant (LEA) hydroxyproline-rich glycoprotein family | -3.19 | AT3G44220.1 |
| 1563 | HORVU4Hr1G017680 | NAD kinase 2 | -3.19 | AT1G21640.1 |
| 1564 | HORVU3Hr1G023470 | undescribed protein | -3.19 | AT5G15300.1 |
| 1565 | HORVU4Hr1G084410 | 50S ribosomal protein L5 | -3.19 | AT4G01310.1 |
| 1566 | HORVU6Hr1G088370 | GATA transcription factor 9 | -3.19 | AT4G32890.1 |
| 1567 | HORVU2Hr1G037020 | alpha/beta-Hydrolases superfamily protein | -3.19 | AT5G19850.1 |
| 1568 | HORVU7Hr1G040390 | Alanine--tRNA ligase | -3.19 | AT5G22800.1 |
| 1569 | HORVU4Hr1G020030 | Abscisic stress-ripening protein 1 | -3.19 | AT5G37710.1 |
| 1570 | HORVU4Hr1G014070 | UPF0454 protein C12orf49 homolog | -3.19 | AT3G59390.2 |
| 1571 | HORVU4Hr1G056060 | APRATAXIN-like | -3.20 | AT5G01310.1 |
| 1572 | HORVU5Hr1G024250 | nucleoside diphosphate kinase 2 | -3.20 | AT5G63310.1 |
| 1573 | HORVU4Hr1G007420 | Zinc finger protein CONSTANS-LIKE 16 | -3.20 | AT1G25440.1 |
| 1574 | HORVU3Hr1G084660 | Phosphoglucan, water dikinase, chloroplastic | -3.20 | AT5G01260.3 |
| 1575 | HORVU4Hr1G085050 | purple acid phosphatase 27 | -3.21 | AT5G50400.1 |
| 1576 | HORVU6Hr1G004980 | unknown function | -3.21 | AT3G45050.4 |
| 1577 | HORVU2Hr1G091680 | Remorin family protein | -3.21 | AT5G23750.3 |
| 1578 | HORVU5Hr1G112340 | dihydroflavonol 4-reductase-like1 | -3.21 | AT5G58490.1 |
| 1579 | HORVU4Hr1G064580 | microtubule-associated protein 65-7 | -3.21 | AT1G14690.2 |
| 1580 | HORVU1Hr1G066910 | undescribed protein | -3.22 | #N/A |
| 1581 | HORVU3Hr1G050430 | protein kinase family protein | -3.22 | AT5G12000.1 |
| 1582 | HORVU5Hr1G031410 | undescribed protein | -3.22 | AT5G29624.1 |
| 1583 | HORVU4Hr1G068990 | unknown function | -3.22 | AT2G03440.1 |
| 1584 | HORVU3Hr1G097280 | undescribed protein | -3.23 | AT1G10130.1 |
| 1585 | HORVU6Hr1G055960 | Bidirectional sugar transporter N3 | -3.23 | AT4G10850.1 |
| 1586 | HORVU6Hr1G014130 | undescribed protein | -3.23 | AT5G51545.1 |
| 1587 | HORVU4Hr1G057040 | undescribed protein | -3.23 | #N/A |
| 1588 | HORVU7Hr1G095810 | mitogen-activated protein kinase 16 | -3.23 | AT3G18040.4 |
| 1589 | HORVU6Hr1G092960 | Aquaporin-like superfamily protein | -3.24 | AT4G00430.1 |
| 1590 | HORVU5Hr1G062890 | Glycine--tRNA ligase beta subunit | -3.24 | AT3G48110.1 |
| 1591 | HORVU3Hr1G063700 | Chromosome 3B, genomic scaffold, cultivar Chinese Spring | -3.24 | AT5G38570.1 |
| 1592 | HORVU1Hr1G022840 | Peroxisomal membrane protein 2 | -3.25 | AT1G52870.2 |
| 1593 | HORVU5Hr1G070230 | Dicarboxylate transporter 2.1, chloroplastic | -3.25 | AT5G64290.1 |
| 1594 | HORVU6Hr1G028680 | Two-component response regulator ARR12 | -3.25 | AT2G25180.1 |
| 1595 | HORVU5Hr1G014290 | Auxin-responsive protein IAA31 | -3.25 | AT1G04240.1 |
| 1596 | HORVU6Hr1G067390 | Protein kinase superfamily protein | -3.25 | AT5G16810.2 |
| 1597 | HORVU2Hr1G109710 | TLD-domain containing nucleolar protein | -3.26 | AT5G39590.1 |
| 1598 | HORVU5Hr1G050180 | Na+/H+ antiporter 2 | -3.26 | AT3G19490.1 |
| 1599 | HORVU1Hr1G076460 | Non-symbiotic hemoglobin 1 | -3.26 | AT2G16060.1 |
| 1600 | HORVU7Hr1G012560 | arginine decarboxylase 2 | -3.26 | AT4G34710.2 |
| 1601 | HORVU3Hr1G050080 | Protein of unknown function (DUF1230) | -3.27 | AT5G67370.1 |

| | | | | |
|---|---|---|---|---|
| 1602 | HORVU2Hr1G023480 | Uncharacterised protein family (UPF0114) | -3.27 | AT5G13720.1 |
| 1603 | HORVU6Hr1G082060 | 10 kDa chaperonin | -3.27 | AT5G20720.4 |
| 1604 | HORVU5Hr1G026640 | ankyrin repeat family protein | -3.27 | AT3G04710.1 |
| 1605 | HORVU2Hr1G116090 | undescribed protein | -3.27 | AT1G56000.2 |
| 1606 | HORVU3Hr1G032440 | Two-component response regulator-like APRR2 | -3.28 | AT2G20570.1 |
| 1607 | HORVU2Hr1G060390 | Early-responsive to dehydration stress protein (ERD4) | -3.28 | AT1G30360.1 |
| 1608 | HORVU7Hr1G049070 | caffeoyl-CoA 3-O-methyltransferase | -3.29 | AT4G34050.1 |
| 1609 | HORVU6Hr1G039660 | CASP-like protein 6 | -3.29 | AT5G54980.1 |
| 1610 | HORVU5Hr1G062030 | Alcohol dehydrogenase | -3.29 | AT4G37980.1 |
| 1611 | HORVU0Hr1G039910 | Protein kinase superfamily protein | -3.29 | AT3G55950.1 |
| 1612 | HORVU4Hr1G042610 | Rho GTPase-activating protein 7 | -3.29 | AT5G19390.2 |
| 1613 | HORVU7Hr1G061500 | Transposon Ty1-PL Gag-Pol polyprotein | -3.30 | AT4G23160.3 |
| 1614 | HORVU4Hr1G054580 | PHD and RING finger domain-containing protein 1 | -3.30 | AT3G05670.3 |
| 1615 | HORVU2Hr1G105830 | Tetratricopeptide repeat (TPR)-like superfamily protein | -3.31 | AT1G78915.1 |
| 1616 | HORVU5Hr1G009890 | Pentatricopeptide repeat (PPR) superfamily protein | -3.32 | AT4G21190.1 |
| 1617 | HORVU0Hr1G011850 | undescribed protein | -3.32 | AT4G10330.1 |
| 1618 | HORVU1Hr1G079410 | Protein of unknown function (DUF1639) | -3.32 | AT1G55340.1 |
| 1619 | HORVU5Hr1G004630 | Photosystem II protein D1 | -3.33 | ATCG00020.1 |
| 1620 | HORVU2Hr1G076180 | Regulator of chromosome condensation (RCC1) family protein | -3.33 | AT5G63860.1 |
| 1621 | HORVU4Hr1G066880 | Cytochrome P450 superfamily protein | -3.33 | AT3G56630.1 |
| 1622 | HORVU2Hr1G101390 | undescribed protein | -3.34 | AT5G24780.1 |
| 1623 | HORVU6Hr1G041720 | Glycine--tRNA ligase alpha subunit | -3.34 | AT3G48110.1 |
| 1624 | HORVU3Hr1G031400 | Transcription factor bHLH68 | -3.34 | AT4G29100.2 |
| 1625 | HORVU6Hr1G004230 | undescribed protein | -3.34 | AT1G02120.2 |
| 1626 | HORVU7Hr1G113460 | Chaperone protein DnaJ | -3.35 | AT1G80920.1 |
| 1627 | HORVU2Hr1G057700 | Oxygen-evolving enhancer protein 1, chloroplastic | -3.35 | AT3G50820.1 |
| 1628 | HORVU2Hr1G095890 | HMG1/2-like protein | -3.36 | AT1G20693.2 |
| 1629 | HORVU5Hr1G007980 | Glutathione-regulated potassium-efflux system protein | -3.36 | AT4G04850.2 |
| 1630 | HORVU2Hr1G078410 | Demethylmenaquinone methyltransferase | -3.36 | AT1G78140.1 |
| 1631 | HORVU7Hr1G076030 | Beta-carotene isomerase D27, chloroplastic | -3.36 | AT1G64680.1 |
| 1632 | HORVU7Hr1G002350 | Disease resistance protein | -3.36 | AT3G14470.2 |
| 1633 | HORVU7Hr1G025730 | Ribosomal protein L11 methyltransferase | -3.36 | AT4G28830.4 |
| 1634 | HORVU4Hr1G076930 | unknown function | -3.37 | AT5G08390.1 |
| 1635 | HORVU2Hr1G084910 | basic helix-loop-helix (bHLH) DNA-binding superfamily protein | -3.37 | AT1G05710.5 |
| 1636 | HORVU7Hr1G019830 | Acyl-ACP thioesterase | -3.37 | AT1G08510.1 |
| 1637 | HORVU2Hr1G038580 | F-box domain containing protein | -3.37 | AT2G13980.1 |

| | | | | |
|---|---|---|---|---|
| 1638 | HORVU7Hr1G041570 | alpha/beta-Hydrolases superfamily protein | -3.37 | AT5G37710.1 |
| 1639 | HORVU5Hr1G016730 | Protein kinase superfamily protein | -3.37 | AT2G23450.2 |
| 1640 | HORVU6Hr1G085730 | Alpha/beta hydrolase domain-containing protein 13 | -3.38 | AT4G31020.2 |
| 1641 | HORVU4Hr1G003800 | BEL1-like homeodomain 6 | -3.38 | AT4G34610.4 |
| 1642 | HORVU4Hr1G057450 | 50S ribosomal protein L10 | -3.38 | AT5G13510.1 |
| 1643 | HORVU6Hr1G060100 | pH-response regulator protein palA/RIM20 | -3.39 | AT1G15130.1 |
| 1644 | HORVU2Hr1G126540 | translocon at the outer membrane of chloroplasts 64-V | -3.39 | AT1G08980.1 |
| 1645 | HORVU4Hr1G057050 | undescribed protein | -3.39 | AT1G10900.3 |
| 1646 | HORVU1Hr1G053670 | Bifunctional inhibitor/lipid-transfer protein/seed storage 2S albumin superfamily protein | -3.40 | AT4G12470.1 |
| 1647 | HORVU5Hr1G019650 | Heparanase-like protein 1 | -3.40 | AT5G07830.1 |
| 1648 | HORVU5Hr1G045440 | ADP-ribosylation factor 1 | -3.40 | AT2G47170.3 |
| 1649 | HORVU6Hr1G063650 | GRAS family transcription factor | -3.40 | AT4G00150.1 |
| 1650 | HORVU0Hr1G038350 | Histidine protein methyltransferase 1 homolog | -3.41 | AT2G43320.2 |
| 1651 | HORVU2Hr1G073370 | photosystem I subunit O | -3.42 | AT1G08380.1 |
| 1652 | HORVU3Hr1G087500 | IQ-domain 3 | -3.42 | AT5G03040.3 |
| 1653 | HORVU4Hr1G058580 | Lipid A export ATP-binding/permease protein MsbA | -3.42 | AT3G55320.1 |
| 1654 | HORVU4Hr1G073930 | glutathione reductase | -3.43 | AT3G54660.1 |
| 1655 | HORVU4Hr1G025330 | 2,3-bisphosphoglycerate-dependent phosphoglycerate mutase | -3.43 | AT5G22620.6 |
| 1656 | HORVU6Hr1G083930 | Protein CYPRO4 | -3.43 | AT3G19240.1 |
| 1657 | HORVU7Hr1G039810 | WD-40 repeat family protein | -3.43 | AT1G24130.1 |
| 1658 | HORVU0Hr1G038050 | 0 | -3.44 | #N/A |
| 1659 | HORVU2Hr1G104830 | undescribed protein | -3.44 | AT5G40510.1 |
| 1660 | HORVU3Hr1G083680 | Harpin induced gene 1 homolog | -3.44 | AT2G35980.1 |
| 1661 | HORVU4Hr1G072220 | pseudouridine synthase family protein | -3.44 | AT2G39140.1 |
| 1662 | HORVU5Hr1G053270 | Rhodanese-like domain-containing protein 10 | -3.44 | AT2G42220.1 |
| 1663 | HORVU1Hr1G068020 | Aldehyde dehydrogenase family 2 member C4 | -3.44 | AT3G24503.1 |
| 1664 | HORVU1Hr1G064200 | RNA-binding KH domain-containing protein | -3.45 | AT1G09660.3 |
| 1665 | HORVU6Hr1G050440 | senescence-associated family protein | -3.45 | AT1G66330.2 |
| 1666 | HORVU1Hr1G086070 | Auxin-responsive protein IAA19 | -3.45 | AT2G22670.5 |
| 1667 | HORVU5Hr1G052030 | C2 domain-containing protein 5 | -3.45 | AT5G47710.4 |
| 1668 | HORVU2Hr1G119760 | Pentatricopeptide repeat-containing protein | -3.45 | AT1G50270.1 |
| 1669 | HORVU7Hr1G056310 | undescribed protein | -3.45 | AT1G33750.1 |
| 1670 | HORVU3Hr1G114290 | Protein of unknown function (DUF707) | -3.45 | AT1G13000.5 |
| 1671 | HORVU4Hr1G005670 | Nuclear transcription factor Y subunit A-9 | -3.46 | AT5G12840.5 |
| 1672 | HORVU7Hr1G094870 | Eukaryotic aspartyl protease family protein | -3.46 | AT5G10770.1 |
| 1673 | HORVU3Hr1G107290 | UPF0001 protein YBL036C | -3.46 | AT4G26860.1 |
| 1674 | HORVU4Hr1G079440 | exocyst subunit exo70 family protein F1 | -3.46 | AT5G50380.1 |

| | | | | |
|---|---|---|---|---|
| 1675 | HORVU6Hr1G073180 | methyltransferase type 11 | -3.47 | AT3G18000.1 |
| 1676 | HORVU4Hr1G016190 | 60S ribosomal protein L10-1 | -3.47 | AT1G66580.1 |
| 1677 | HORVU0Hr1G016150 | RNA-binding protein 39 | -3.47 | AT4G09040.2 |
| 1678 | HORVU0Hr1G008940 | heat shock factor 3 | -3.48 | AT5G16820.2 |
| 1679 | HORVU3Hr1G075820 | Chromosome 3B, genomic scaffold, cultivar Chinese Spring | -3.48 | AT5G43630.4 |
| 1680 | HORVU3Hr1G016240 | receptor kinase 2 | -3.48 | AT1G60800.2 |
| 1681 | HORVU3Hr1G029210 | SCAR-like protein 2 | -3.49 | AT2G38440.1 |
| 1682 | HORVU5Hr1G018690 | Eukaryotic aspartyl protease family protein | -3.49 | AT2G03200.1 |
| 1683 | HORVU3Hr1G098170 | Mitochondrial substrate carrier family protein | -3.49 | AT5G48970.1 |
| 1684 | HORVU2Hr1G101440 | Structural molecule, putative isoform 5 | -3.49 | AT1G51110.1 |
| 1685 | HORVU4Hr1G048300 | undescribed protein | -3.49 | AT1G67160.1 |
| 1686 | HORVU3Hr1G094240 | 23S rRNA (uracil(747)-C(5))-methyltransferase RlmC | -3.49 | AT3G27180.1 |
| 1687 | HORVU3Hr1G084750 | Auxin transporter-like protein 1 | -3.50 | AT2G38120.1 |
| 1688 | HORVU3Hr1G054220 | undescribed protein | -3.50 | AT1G19270.2 |
| 1689 | HORVU5Hr1G113990 | Pentatricopeptide repeat-containing protein | -3.50 | AT1G59720.1 |
| 1690 | HORVU5Hr1G014300 | Auxin-responsive protein IAA30 | -3.50 | AT3G04730.1 |
| 1691 | HORVU3Hr1G016700 | NADH:ubiquinone oxidoreductase, 17.2kDa subunit | -3.52 | AT4G26965.1 |
| 1692 | HORVU5Hr1G088200 | Carboxyl-terminal-processing protease | -3.52 | AT4G17740.2 |
| 1693 | HORVU6Hr1G063250 | undescribed protein | -3.52 | AT3G27190.1 |
| 1694 | HORVU5Hr1G106120 | Opaque-2-2 protein | -3.52 | AT4G02640.4 |
| 1695 | HORVU6Hr1G049260 | Ribulose bisphosphate carboxylase large chain | -3.53 | ATCG00490.1 |
| 1696 | HORVU3Hr1G092020 | BTB/POZ and TAZ domain-containing protein 2 | -3.53 | AT3G48360.1 |
| 1697 | HORVU5Hr1G060370 | vacuolar iron transporter 1 | -3.53 | AT2G01770.1 |
| 1698 | HORVU5Hr1G025960 | acyl carrier protein 5 | -3.53 | AT4G25050.1 |
| 1699 | HORVU6Hr1G058580 | myb domain protein 15 | -3.53 | AT3G23250.1 |
| 1700 | HORVU5Hr1G071250 | D6 protein kinase like 2 | -3.53 | AT5G47750.1 |
| 1701 | HORVU3Hr1G030250 | unknown function | -3.53 | AT3G48200.1 |
| 1702 | HORVU7Hr1G062930 | Peptidyl-prolyl cis-trans isomerase, chloroplastic | -3.54 | AT3G01480.1 |
| 1703 | HORVU0Hr1G035820 | serine acetyltransferase 1;1 | -3.54 | AT5G56760.1 |
| 1704 | HORVU3Hr1G059340 | undescribed protein | -3.54 | AT1G22380.1 |
| 1705 | HORVU5Hr1G010960 | Glutaredoxin family protein | -3.55 | AT2G47870.1 |
| 1706 | HORVU0Hr1G019710 | alpha/beta-Hydrolases superfamily protein | -3.55 | AT1G47480.1 |
| 1707 | HORVU5Hr1G073310 | BnaCnng49070D protein | -3.55 | AT4G12070.1 |
| 1708 | HORVU1Hr1G016360 | Triose phosphate/phosphate translocator, chloroplastic | -3.55 | AT5G46110.1 |
| 1709 | HORVU3Hr1G027610 | Class I glutamine amidotransferase-like superfamily protein | -3.55 | AT1G53280.1 |
| 1710 | HORVU7Hr1G036380 | RING/U-box superfamily protein | -3.55 | AT5G27420.1 |
| 1711 | HORVU6Hr1G092970 | Aquaporin-like superfamily protein | -3.55 | AT4G00430.1 |
| 1712 | HORVU2Hr1G114580 | Clustered mitochondria protein | -3.56 | AT4G28080.1 |

| | | | | |
|---|---|---|---|---|
| 1713 | HORVU6Hr1G011060 | unknown function | -3.56 | AT2G29100.1 |
| 1714 | HORVU5Hr1G008980 | receptor kinase 3 | -3.56 | AT1G21230.1 |
| 1715 | HORVU6Hr1G017860 | L-aspartate oxidase | -3.57 | AT5G14760.1 |
| 1716 | HORVU4Hr1G074810 | D-ribulose-5-phosphate-3-epimerase | -3.57 | AT5G61410.2 |
| 1717 | HORVU1Hr1G090990 | Protein CHUP1, chloroplastic | -3.57 | AT3G25690.6 |
| 1718 | HORVU1Hr1G034570 | Ribosomal RNA small subunit methyltransferase I | -3.57 | AT1G45110.1 |
| 1719 | HORVU0Hr1G014950 | Protein kinase superfamily protein | -3.57 | AT1G61590.1 |
| 1720 | HORVU3Hr1G034210 | HAD-superfamily hydrolase, subfamily IG, 5'-nucleotidase | -3.58 | AT5G48960.1 |
| 1721 | HORVU6Hr1G025880 | unknown function | -3.58 | AT2G04360.1 |
| 1722 | HORVU5Hr1G124210 | PGR5-LIKE A | -3.58 | AT4G22890.3 |
| 1723 | HORVU6Hr1G014480 | adenylate kinase family protein | -3.58 | AT5G35170.1 |
| 1724 | HORVU5Hr1G096370 | UDP-glucose 6-dehydrogenase family protein | -3.59 | AT5G15490.1 |
| 1725 | HORVU6Hr1G015770 | Photosystem II 5 kD protein | -3.59 | AT1G51400.1 |
| 1726 | HORVU5Hr1G098490 | Cellulose-synthase-like C4 | -3.59 | AT4G07960.2 |
| 1727 | HORVU1Hr1G086110 | Protein of unknown function (DUF760) | -3.60 | AT3G07310.1 |
| 1728 | HORVU2Hr1G018730 | alkyl hydroperoxide reductase | -3.60 | AT1G21350.5 |
| 1729 | HORVU3Hr1G049970 | undescribed protein | -3.60 | AT4G24330.1 |
| 1730 | HORVU7Hr1G054530 | Lipid transfer protein-like | -3.61 | AT1G27950.1 |
| 1731 | HORVU6Hr1G026680 | Beta-1,3-galactosyltransferase 7 | -3.61 | AT1G05170.2 |
| 1732 | HORVU4Hr1G020850 | undescribed protein | -3.61 | AT1G51630.1 |
| 1733 | HORVU4Hr1G083080 | serine/threonine protein kinase 1 | -3.61 | AT1G01140.1 |
| 1734 | HORVU3Hr1G073040 | Phosphatidate cytidylyltransferase family protein | -3.62 | AT1G62430.1 |
| 1735 | HORVU7Hr1G036130 | MADS-box transcription factor 55 | -3.62 | AT2G22540.2 |
| 1736 | HORVU3Hr1G060730 | unknown function | -3.62 | AT2G42975.1 |
| 1737 | HORVU2Hr1G127440 | purple acid phosphatase 22 | -3.62 | AT3G52820.1 |
| 1738 | HORVU2Hr1G086040 | Photosystem II reaction center PsbP family protein | -3.62 | AT1G77090.1 |
| 1739 | HORVU5Hr1G008080 | undescribed protein | -3.63 | AT1G04150.1 |
| 1740 | HORVU4Hr1G057890 | Glutathione S-transferase family protein | -3.64 | AT5G02790.1 |
| 1741 | HORVU4Hr1G048960 | UBA domain-containing protein 7 | -3.64 | AT5G44180.3 |
| 1742 | HORVU7Hr1G021630 | GDSL esterase/lipase | -3.64 | AT3G16370.1 |
| 1743 | HORVU1Hr1G072750 | uracil phosphoribosyltransferase | -3.64 | AT3G53900.2 |
| 1744 | HORVU4Hr1G029350 | protein kinase family protein | -3.65 | AT5G49470.3 |
| 1745 | HORVU2Hr1G123270 | unknown function | -3.65 | AT5G67610.1 |
| 1746 | HORVU0Hr1G019500 | HXXXD-type acyl-transferase family protein | -3.65 | AT1G28680.1 |
| 1747 | HORVU5Hr1G110960 | Transcription initiation factor IIE subunit beta | -3.65 | AT4G20330.1 |
| 1748 | HORVU6Hr1G052230 | undescribed protein | -3.66 | AT5G32470.1 |
| 1749 | HORVU3Hr1G054770 | ATP-dependent zinc metalloprotease FtsH | -3.66 | AT5G53170.1 |
| 1750 | HORVU1Hr1G064290 | Alpha beta-hydrolase-like protein | -3.66 | AT5G38520.1 |
| 1751 | HORVU2Hr1G089340 | phosphoenolpyruvate carboxylase kinase 1 | -3.66 | AT1G08650.1 |

| | | | | |
|---|---|---|---|---|
| 1752 | HORVU4Hr1G078780 | unknown function | -3.66 | AT4G28760.2 |
| 1753 | HORVU2Hr1G116130 | bZIP transcription factor family protein | -3.66 | AT5G65210.7 |
| 1754 | HORVU1Hr1G006860 | Glucose-6-phosphate isomerase | -3.67 | AT5G42740.2 |
| 1755 | HORVU6Hr1G045140 | P0696G06.27 protein | -3.67 | AT1G72550.2 |
| 1756 | HORVU3Hr1G028810 | Heme-binding protein 2 | -3.68 | AT1G17100.1 |
| 1757 | HORVU5Hr1G016970 | Retrovirus-related Pol polyprotein LINE-1 | -3.68 | AT3G50380.4 |
| 1758 | HORVU3Hr1G041820 | Protein kinase superfamily protein | -3.68 | AT1G48210.7 |
| 1759 | HORVU7Hr1G051760 | chaperone protein dnaJ-related | -3.68 | AT2G24395.1 |
| 1760 | HORVU5Hr1G112710 | Leucine-rich repeat-containing protein 49 | -3.69 | AT1G78230.1 |
| 1761 | HORVU4Hr1G021950 | Signal recognition particle 54 kDa protein, chloroplastic | -3.69 | AT5G03940.1 |
| 1762 | HORVU2Hr1G059380 | RNA binding | -3.69 | AT3G52380.1 |
| 1763 | HORVU2Hr1G099510 | terpene synthase 03 | -3.69 | AT1G79460.1 |
| 1764 | HORVU7Hr1G012530 | arginine decarboxylase 2 | -3.70 | AT4G34710.2 |
| 1765 | HORVU2Hr1G093580 | Protein kinase superfamily protein | -3.70 | AT2G44830.3 |
| 1766 | HORVU4Hr1G011250 | Sorghum bicolor protein targeted either to mitochondria or chloroplast proteins T50848 | -3.70 | AT5G05250.1 |
| 1767 | HORVU6Hr1G090250 | unknown function | -3.70 | AT5G37360.1 |
| 1768 | HORVU1Hr1G047820 | Aldo-keto reductase family 4 member C9 | -3.70 | AT5G53580.1 |
| 1769 | HORVU1Hr1G083160 | Non-specific lipid-transfer protein 2 | -3.70 | AT3G18280.2 |
| 1770 | HORVU0Hr1G016560 | unknown function | -3.70 | AT5G62140.1 |
| 1771 | HORVU2Hr1G023540 | Magnesium-chelatase subunit ChlH, chloroplastic | -3.70 | AT5G13630.1 |
| 1772 | HORVU2Hr1G058230 | Protein S-acyltransferase 24 | -3.71 | AT5G20350.1 |
| 1773 | HORVU6Hr1G093570 | Prolyl 4-hydroxylase subunit alpha-1 | -3.71 | AT1G20270.1 |
| 1774 | HORVU7Hr1G049200 | F-box protein At-B | -3.71 | AT1G55590.1 |
| 1775 | HORVU7Hr1G026100 | undescribed protein | -3.71 | AT3G54920.1 |
| 1776 | HORVU2Hr1G035550 | undescribed protein | -3.71 | AT2G23390.2 |
| 1777 | HORVU1Hr1G018080 | ATP-dependent RNA helicase DOB1 | -3.71 | AT1G59760.1 |
| 1778 | HORVU5Hr1G071920 | Photosystem I reaction center subunit V | -3.72 | AT1G55670.1 |
| 1779 | HORVU0Hr1G016250 | RNA binding | -3.72 | AT3G08010.1 |
| 1780 | HORVU7Hr1G019510 | Dormancy/auxin associated family protein | -3.72 | AT1G56220.1 |
| 1781 | HORVU4Hr1G016470 | Protein kinase superfamily protein | -3.72 | AT4G31390.1 |
| 1782 | HORVU1Hr1G092860 | Mitochondrial substrate carrier family protein | -3.73 | AT4G01100.1 |
| 1783 | HORVU4Hr1G072150 | 4-coumarate--CoA ligase-like 1 | -3.73 | AT4G05160.1 |
| 1784 | HORVU4Hr1G050630 | eukaryotic translation initiation factor 2 beta subunit | -3.74 | AT5G20920.3 |
| 1785 | HORVU3Hr1G078830 | unknown function | -3.74 | AT3G17930.1 |
| 1786 | HORVU7Hr1G014840 | Chromosome 3B, genomic scaffold, cultivar Chinese Spring | -3.74 | AT1G32928.1 |
| 1787 | HORVU4Hr1G023230 | undescribed protein | -3.74 | #N/A |
| 1788 | HORVU6Hr1G008590 | 30S ribosomal protein S1 | -3.74 | AT1G12800.1 |
| 1789 | HORVU5Hr1G073380 | Ethylene insensitive 3 family protein | -3.75 | AT1G73730.1 |

| | | | | |
|---|---|---|---|---|
| 1790 | HORVU3Hr1G031460 | Auxin-responsive protein IAA17 | -3.75 | AT4G29080.1 |
| 1791 | HORVU6Hr1G014100 | Protein LOW PSII ACCUMULATION 2, chloroplastic | -3.75 | AT5G51545.1 |
| 1792 | HORVU7Hr1G052400 | TBC1 domain family member 5 homolog A | -3.76 | AT4G29950.1 |
| 1793 | HORVU3Hr1G043870 | Glucose-6-phosphate isomerase | -3.76 | AT5G42740.3 |
| 1794 | HORVU3Hr1G088910 | Mitochondrial intermediate peptidase | -3.76 | AT1G67690.1 |
| 1795 | HORVU5Hr1G076100 | ACT domain-containing protein | -3.76 | AT1G69040.2 |
| 1796 | HORVU3Hr1G057020 | Dual-specificity RNA methyltransferase RlmN | -3.77 | AT1G60230.1 |
| 1797 | HORVU2Hr1G010520 | Eukaryotic aspartyl protease family protein | -3.77 | AT4G35880.1 |
| 1798 | HORVU5Hr1G072120 | unknown function | -3.77 | AT2G33180.1 |
| 1799 | HORVU6Hr1G032540 | Protein kinase superfamily protein | -3.78 | AT1G61590.1 |
| 1800 | HORVU4Hr1G004210 | ferredoxin 3 | -3.78 | AT1G32550.1 |
| 1801 | HORVU6Hr1G046980 | High mobility group B protein 15 | -3.78 | AT1G04880.1 |
| 1802 | HORVU3Hr1G065630 | Protein kinase superfamily protein | -3.79 | AT5G55090.1 |
| 1803 | HORVU6Hr1G060810 | Pentapeptide repeat-containing protein | -3.79 | AT1G12250.2 |
| 1804 | HORVU2Hr1G104920 | glutamate receptor 3.3 | -3.79 | AT1G42540.4 |
| 1805 | HORVU4Hr1G008210 | histidine kinase 3 | -3.79 | AT2G01830.6 |
| 1806 | HORVU7Hr1G060530 | Translation initiation factor IF-2 | -3.80 | AT1G17220.1 |
| 1807 | HORVU6Hr1G095000 | Disease resistance protein | -3.81 | AT3G14470.2 |
| 1808 | HORVU3Hr1G043300 | zinc induced facilitator-like 2 | -3.81 | AT3G43790.4 |
| 1809 | HORVU3Hr1G078200 | Protein CHLOROPLAST ENHANCING STRESS TOLERANCE, chloroplastic | -3.81 | AT5G44650.1 |
| 1810 | HORVU1Hr1G020000 | Protease Do-like 9 | -3.81 | AT2G47940.2 |
| 1811 | HORVU3Hr1G036380 | NAC domain protein, | -3.82 | AT5G17260.1 |
| 1812 | HORVU2Hr1G060880 | Oxygen-evolving enhancer protein 2, chloroplastic | -3.82 | AT1G06680.1 |
| 1813 | HORVU7Hr1G027740 | undescribed protein | -3.82 | AT1G05990.1 |
| 1814 | HORVU7Hr1G086340 | undescribed protein | -3.83 | AT2G32210.1 |
| 1815 | HORVU7Hr1G108650 | undescribed protein | -3.83 | #N/A |
| 1816 | HORVU4Hr1G007380 | undescribed protein | -3.83 | AT1G63660.2 |
| 1817 | HORVU7Hr1G097660 | Chromosome 3B, genomic scaffold, cultivar Chinese Spring | -3.83 | AT5G06570.2 |
| 1818 | HORVU2Hr1G034400 | Eukaryotic translation initiation factor 5A | -3.83 | AT1G69410.1 |
| 1819 | HORVU2Hr1G101040 | calcium-transporting ATPase, putative | -3.83 | AT5G57110.3 |
| 1820 | HORVU3Hr1G005770 | ABC transporter G family member 7 | -3.84 | AT2G01320.7 |
| 1821 | HORVU4Hr1G079600 | undescribed protein | -3.85 | AT4G32700.2 |
| 1822 | HORVU4Hr1G029400 | undescribed protein | -3.87 | AT5G10490.2 |
| 1823 | HORVU0Hr1G001490 | Photosystem I reaction center subunit VI, chloroplastic | -3.88 | AT1G52230.1 |
| 1824 | HORVU7Hr1G021180 | Single-stranded DNA-binding protein WHY1, chloroplastic | -3.88 | AT2G02740.1 |
| 1825 | HORVU6Hr1G005020 | unknown function | -3.88 | AT3G45050.4 |
| 1826 | HORVU6Hr1G009480 | Low PSII Accumulation 3 | -3.88 | AT1G73060.1 |

| | | | | |
|---|---|---|---|---|
| 1827 | HORVU7Hr1G100310 | 2-oxoglutarate (2OG) and Fe(II)-dependent oxygenase superfamily protein | -3.88 | AT3G50210.2 |
| 1828 | HORVU2Hr1G079180 | unknown function | -3.89 | AT2G14095.1 |
| 1829 | HORVU4Hr1G013420 | BTB/POZ domain-containing protein | -3.89 | AT2G14820.2 |
| 1830 | HORVU4Hr1G023580 | Zeaxanthin epoxidase, chloroplastic | -3.89 | AT2G21530.1 |
| 1831 | HORVU3Hr1G064340 | Protein PLASTID MOVEMENT IMPAIRED 2 | -3.89 | AT1G66840.2 |
| 1832 | HORVU2Hr1G008380 | FKBP-like peptidyl-prolyl cis-trans isomerase family protein | -3.89 | AT5G13410.1 |
| 1833 | HORVU0Hr1G008530 | undescribed protein | -3.90 | AT5G41190.1 |
| 1834 | HORVU6Hr1G069390 | histidinol phosphate aminotransferase 1 | -3.90 | AT5G10330.2 |
| 1835 | HORVU5Hr1G119790 | carotene isomerase | -3.90 | AT1G57770.1 |
| 1836 | HORVU7Hr1G020590 | Fe superoxide dismutase 3 | -3.90 | AT5G23310.1 |
| 1837 | HORVU4Hr1G067280 | Homocysteine S-methyltransferase family protein | -3.90 | AT3G25900.1 |
| 1838 | HORVU1Hr1G020240 | NBS-LRR disease resistance protein homologue | -3.91 | AT4G19510.8 |
| 1839 | HORVU7Hr1G012940 | Peptidyl-prolyl cis-trans isomerase-like 3 | -3.92 | AT1G01940.1 |
| 1840 | HORVU2Hr1G018340 | protein kinase family protein | -3.92 | AT2G30360.1 |
| 1841 | HORVU2Hr1G087010 | Calcium uptake protein 1, mitochondrial | -3.93 | AT4G32060.1 |
| 1842 | HORVU4Hr1G014590 | unknown function | -3.93 | AT5G67550.1 |
| 1843 | HORVU5Hr1G043800 | undescribed protein | -3.93 | #N/A |
| 1844 | HORVU2Hr1G106100 | Transmembrane protein 41B | -3.94 | AT4G17790.1 |
| 1845 | HORVU1Hr1G065250 | Potassium channel AKT2 | -3.94 | AT4G22200.1 |
| 1846 | HORVU4Hr1G026670 | Chromosome 3B, genomic scaffold, cultivar Chinese Spring | -3.95 | AT1G15660.1 |
| 1847 | HORVU3Hr1G020830 | metacaspase 1 | -3.96 | AT1G02170.1 |
| 1848 | HORVU0Hr1G004060 | VQ motif family protein | -3.96 | AT3G18690.1 |
| 1849 | HORVU2Hr1G105930 | RNA-binding protein 28 | -3.96 | AT2G35410.1 |
| 1850 | HORVU5Hr1G042310 | ACT domain-containing protein | -3.96 | AT5G04740.1 |
| 1851 | HORVU4Hr1G014910 | Ankyrin repeat protein SKIP35 | -3.96 | AT2G44090.5 |
| 1852 | HORVU3Hr1G057240 | Glutaredoxin family protein | -3.96 | AT1G64500.1 |
| 1853 | HORVU5Hr1G017400 | Mutator-like transposase | -3.97 | AT3G05850.1 |
| 1854 | HORVU1Hr1G088870 | Chlorophyll a-b binding protein, chloroplastic | -3.98 | AT2G34420.1 |
| 1855 | HORVU4Hr1G015260 | Chlorophyll a-b binding protein 2, chloroplastic | -3.98 | AT4G10340.1 |
| 1856 | HORVU7Hr1G099350 | undescribed protein | -3.99 | AT3G06010.1 |
| 1857 | HORVU3Hr1G062700 | Polyketide cyclase/dehydrase and lipid transport superfamily protein | -3.99 | AT1G02475.1 |
| 1858 | HORVU3Hr1G069730 | UDP-Glycosyltransferase superfamily protein | -4.00 | AT3G16520.3 |
| 1859 | HORVU6Hr1G075540 | undescribed protein | -4.01 | AT1G52990.2 |
| 1860 | HORVU4Hr1G040950 | 50S ribosomal protein L6 | -4.01 | AT1G05190.1 |
| 1861 | HORVU4Hr1G010430 | BEL1-type homeodomain protein | -4.01 | AT2G23760.3 |
| 1862 | HORVU3Hr1G089650 | Rhodanese/Cell cycle control phosphatase superfamily protein | -4.01 | AT3G59780.1 |
| 1863 | HORVU7Hr1G021430 | HXXXD-type acyl-transferase family protein | -4.02 | AT1G31490.1 |
| 1864 | HORVU3Hr1G069650 | BURP domain-containing protein | -4.02 | AT5G25610.1 |

| | | | | |
|---|---|---|---|---|
| 1865 | HORVU1Hr1G016410 | undescribed protein | -4.03 | AT1G16460.4 |
| 1866 | HORVU5Hr1G125110 | RNA polymerase sigma-C factor | -4.03 | AT2G36990.1 |
| 1867 | HORVU1Hr1G092670 | Late embryogenesis abundant protein | -4.04 | AT2G46140.1 |
| 1868 | HORVU5Hr1G070950 | undescribed protein | -4.05 | AT4G22970.3 |
| 1869 | HORVU7Hr1G021950 | xyloglucan endotransglucosylase/hydrolase 25 | -4.05 | AT5G57550.1 |
| 1870 | HORVU5Hr1G065330 | cinnamoyl coa reductase 1 | -4.05 | AT1G15950.3 |
| 1871 | HORVU7Hr1G106540 | Alpha-glucosidase | -4.05 | AT5G11720.1 |
| 1872 | HORVU3Hr1G066240 | chloride channel E | -4.06 | AT4G35440.1 |
| 1873 | HORVU5Hr1G013400 | Chromosome 3B, genomic scaffold, cultivar Chinese Spring | -4.06 | AT5G50760.1 |
| 1874 | HORVU7Hr1G029750 | Protein kinase superfamily protein | -4.08 | AT3G46920.1 |
| 1875 | HORVU3Hr1G065720 | undescribed protein | -4.08 | AT1G28240.1 |
| 1876 | HORVU0Hr1G005330 | RING/U-box superfamily protein | -4.09 | AT3G61460.1 |
| 1877 | HORVU7Hr1G092940 | Potassium transporter family protein | -4.09 | AT5G14880.1 |
| 1878 | HORVU2Hr1G086950 | Calcium-binding mitochondrial carrier protein | -4.09 | AT4G32060.2 |
| 1879 | HORVU1Hr1G094480 | ATP-dependent Clp protease proteolytic subunit 2 | -4.09 | AT1G49970.1 |
| 1880 | HORVU7Hr1G042080 | Zinc finger protein CONSTANS-LIKE 16 | -4.09 | AT1G25440.1 |
| 1881 | HORVU3Hr1G108540 | Two-component response regulator ARR8 | -4.10 | AT3G57040.2 |
| 1882 | HORVU7Hr1G099560 | calmodulin 6 | -4.10 | AT1G18210.2 |
| 1883 | HORVU7Hr1G024560 | undescribed protein | -4.10 | AT5G15180.1 |
| 1884 | HORVU1Hr1G079270 | Polyketide cyclase/dehydrase and lipid transport superfamily protein | -4.10 | AT3G13062.1 |
| 1885 | HORVU3Hr1G065640 | Protein kinase superfamily protein | -4.11 | AT4G26890.1 |
| 1886 | HORVU4Hr1G070720 | Auxilin-related protein 2 | -4.11 | AT1G75100.1 |
| 1887 | HORVU2Hr1G023170 | Thioredoxin reductase | -4.12 | AT2G41680.1 |
| 1888 | HORVU3Hr1G090760 | Alpha/beta hydrolase domain-containing protein 11 | -4.12 | AT4G10030.1 |
| 1889 | HORVU2Hr1G084820 | undescribed protein | -4.13 | AT3G06110.3 |
| 1890 | HORVU0Hr1G018970 | myb-like transcription factor family protein | -4.13 | AT1G49010.1 |
| 1891 | HORVU5Hr1G050610 | PLC-like phosphodiesterases superfamily protein | -4.14 | AT5G67130.1 |
| 1892 | HORVU7Hr1G056320 | undescribed protein | -4.14 | AT3G22780.1 |
| 1893 | HORVU5Hr1G033660 | Valine--tRNA ligase | -4.15 | AT1G14610.1 |
| 1894 | HORVU7Hr1G030810 | glutathione peroxidase 1 | -4.15 | AT2G25080.1 |
| 1895 | HORVU5Hr1G082420 | Chlorophyll a-b binding protein, chloroplastic | -4.16 | AT3G27690.2 |
| 1896 | HORVU2Hr1G043170 | Pentatricopeptide repeat-containing protein | -4.16 | AT5G42310.1 |
| 1897 | HORVU7Hr1G101270 | auxin response factor 16 | -4.16 | AT4G30080.1 |
| 1898 | HORVU4Hr1G015820 | Rpr4901.2 | -4.16 | AT5G63510.1 |
| 1899 | HORVU2Hr1G041610 | Cytochrome b6-f complex iron-sulfur subunit | -4.16 | AT4G03280.1 |
| 1900 | HORVU7Hr1G113850 | WRKY family transcription factor | -4.17 | AT1G80840.1 |
| 1901 | HORVU2Hr1G105900 | unknown function | -4.17 | AT5G16660.1 |
| 1902 | HORVU6Hr1G040040 | Protein of unknown function (DUF3511) | -4.17 | AT5G11970.1 |

| | | | | |
|---|---|---|---|---|
| 1903 | HORVU6Hr1G063510 | unknown function | -4.18 | AT5G21222.4 |
| 1904 | HORVU4Hr1G057910 | Glutathione S-transferase family protein | -4.18 | AT5G02790.1 |
| 1905 | HORVU5Hr1G072920 | Zinc finger A20 and AN1 domain-containing stress-associated protein 1 | -4.18 | AT3G12630.1 |
| 1906 | HORVU3Hr1G085380 | alpha-glucan phosphorylase 2 | -4.18 | AT3G46970.1 |
| 1907 | HORVU3Hr1G003050 | glutamate decarboxylase | -4.19 | AT5G17330.1 |
| 1908 | HORVU5Hr1G121250 | Aspartokinase 1, chloroplastic | -4.19 | AT5G13280.1 |
| 1909 | HORVU2Hr1G104790 | unknown function | -4.19 | #N/A |
| 1910 | HORVU5Hr1G066260 | calmodulin like 42 | -4.19 | AT3G29000.1 |
| 1911 | HORVU1Hr1G070740 | UDP-glucose pyrophosphorylase 3 | -4.20 | AT3G56040.1 |
| 1912 | HORVU1Hr1G000040 | 50S ribosomal protein L28 | -4.20 | AT2G33450.1 |
| 1913 | HORVU3Hr1G097200 | auxin response factor 2 | -4.21 | AT5G62000.5 |
| 1914 | HORVU4Hr1G084890 | Endoribonuclease Dicer homolog 1 | -4.21 | AT1G01040.2 |
| 1915 | HORVU7Hr1G025590 | undescribed protein | -4.21 | AT5G15180.1 |
| 1916 | HORVU2Hr1G041320 | 3'(2'),5'-bisphosphate nucleotidase 1 | -4.22 | AT4G05090.1 |
| 1917 | HORVU7Hr1G101810 | 50S ribosomal protein L24 | -4.22 | AT5G54600.1 |
| 1918 | HORVU3Hr1G005830 | ABC transporter G family member 7 | -4.22 | AT2G01320.3 |
| 1919 | HORVU3Hr1G089290 | Late embryogenesis abundant protein Lea5-A | -4.22 | AT4G02380.3 |
| 1920 | HORVU3Hr1G065960 | Farnesyl pyrophosphate synthase | -4.23 | AT5G47770.1 |
| 1921 | HORVU2Hr1G023560 | Magnesium-chelatase subunit H | -4.24 | AT5G13630.1 |
| 1922 | HORVU2Hr1G023530 | undescribed protein | -4.24 | AT1G73875.1 |
| 1923 | HORVU4Hr1G076690 | Protein of unknown function (DUF1677) | -4.24 | AT4G14819.1 |
| 1924 | HORVU2Hr1G121310 | Serine/threonine-protein kinase | -4.24 | AT2G19130.1 |
| 1925 | HORVU2Hr1G108350 | Chromosome 3B, genomic scaffold, cultivar Chinese Spring | -4.25 | AT2G35460.1 |
| 1926 | HORVU3Hr1G013970 | NAD(P)-binding Rossmann-fold superfamily protein | -4.26 | AT2G37540.1 |
| 1927 | HORVU3Hr1G030580 | Replication protein A 32 kDa subunit B | -4.26 | AT2G24490.2 |
| 1928 | HORVU7Hr1G027270 | MATE efflux family protein | -4.26 | AT5G65380.1 |
| 1929 | HORVU0Hr1G022240 | Sec14p-like phosphatidylinositol transfer family protein | -4.26 | AT5G63060.1 |
| 1930 | HORVU5Hr1G038840 | pumilio 7 | -4.27 | AT1G22240.1 |
| 1931 | HORVU6Hr1G042170 | undescribed protein | -4.27 | AT1G61900.2 |
| 1932 | HORVU3Hr1G096910 | Beta-glucosidase 3 | -4.27 | AT3G60140.2 |
| 1933 | HORVU5Hr1G040640 | undescribed protein | -4.28 | AT4G02070.2 |
| 1934 | HORVU2Hr1G035990 | undescribed protein | -4.28 | AT1G15640.1 |
| 1935 | HORVU2Hr1G066470 | undescribed protein | -4.28 | AT4G28760.2 |
| 1936 | HORVU2Hr1G034540 | unknown function | -4.29 | AT3G15110.1 |
| 1937 | HORVU6Hr1G033970 | glycerol dehydrogenase | -4.29 | AT3G21340.2 |
| 1938 | HORVU7Hr1G012690 | magnesium-protoporphyrin IX methyltransferase | -4.29 | AT4G25080.6 |
| 1939 | HORVU3Hr1G116060 | undescribed protein | -4.30 | AT5G21280.2 |
| 1940 | HORVU3Hr1G086210 | GRAS family transcription factor | -4.30 | AT1G07530.1 |

| | | | | |
|---|---|---|---|---|
| 1941 | HORVU0Hr1G040200 | coenzyme F420 hydrogenase family / dehydrogenase, beta subunit family | -4.30 | AT1G04620.1 |
| 1942 | HORVU0Hr1G003890 | Phosphoadenosine phosphosulfate reductase | -4.30 | AT4G21990.1 |
| 1943 | HORVU5Hr1G043200 | strictosidine synthase-like 2 | -4.31 | AT3G57030.1 |
| 1944 | HORVU1Hr1G030540 | basic helix-loop-helix (bHLH) DNA-binding superfamily protein | -4.31 | AT4G29100.2 |
| 1945 | HORVU2Hr1G094160 | Beta-carotene hydroxylase | -4.31 | AT4G25700.1 |
| 1946 | HORVU2Hr1G009850 | Thylakoid lumenal 15.0 kDa protein 2, chloroplastic | -4.31 | AT5G52970.2 |
| 1947 | HORVU3Hr1G086310 | O-fucosyltransferase family protein | -4.32 | AT2G01480.1 |
| 1948 | HORVU3Hr1G053910 | maternal effect embryo arrest 18 | -4.32 | AT2G34090.1 |
| 1949 | HORVU5Hr1G001650 | Protein of unknown function (DUF1218) | -4.33 | AT5G17210.1 |
| 1950 | HORVU4Hr1G041000 | undescribed protein | -4.33 | AT1G72990.2 |
| 1951 | HORVU4Hr1G002020 | cationic amino acid transporter 2 | -4.34 | AT1G58030.1 |
| 1952 | HORVU2Hr1G118410 | Plant cysteine oxidase 3 | -4.35 | AT1G18490.1 |
| 1953 | HORVU4Hr1G010120 | Prostamide/prostaglandin F synthase | -4.35 | AT5G65840.1 |
| 1954 | HORVU6Hr1G017890 | Chromosome 3B, genomic scaffold, cultivar Chinese Spring | -4.35 | AT1G70420.1 |
| 1955 | HORVU5Hr1G064130 | Pentatricopeptide repeat-containing protein | -4.35 | AT4G30825.1 |
| 1956 | HORVU1Hr1G045280 | BnaC08g35780D protein | -4.36 | AT2G21385.1 |
| 1957 | HORVU2Hr1G107990 | Thioredoxin superfamily protein | -4.36 | AT1G11530.1 |
| 1958 | HORVU1Hr1G052780 | pollen development | -4.37 | AT3G10405.1 |
| 1959 | HORVU2Hr1G076100 | Glutathione S-transferase family protein | -4.37 | AT1G77290.3 |
| 1960 | HORVU5Hr1G022760 | Solute carrier family 40 member 1 | -4.38 | AT5G26820.1 |
| 1961 | HORVU2Hr1G038740 | Aquaporin-like superfamily protein | -4.38 | AT3G54820.1 |
| 1962 | HORVU5Hr1G064500 | undescribed protein | -4.39 | AT1G12190.1 |
| 1963 | HORVU1Hr1G063090 | Beta-mannosidase B | -4.40 | AT1G09010.1 |
| 1964 | HORVU2Hr1G111440 | Ribulose bisphosphate carboxylase/oxygenase activase, chloroplastic | -4.40 | AT1G73110.1 |
| 1965 | HORVU1Hr1G090740 | unknown function | -4.40 | AT2G25605.1 |
| 1966 | HORVU3Hr1G110160 | Signal recognition particle receptor FtsY | -4.41 | AT2G45770.1 |
| 1967 | HORVU1Hr1G056510 | 50S ribosomal protein L1 | -4.41 | AT3G63490.1 |
| 1968 | HORVU7Hr1G113210 | Mog1/PsbP/DUF1795-like photosystem II reaction center PsbP family protein | -4.41 | AT5G11450.2 |
| 1969 | HORVU2Hr1G095400 | Iron-sulfur cluster assembly protein HesB | -4.42 | AT5G52190.1 |
| 1970 | HORVU4Hr1G013380 | LOB domain-containing protein 37 | -4.43 | AT4G37540.1 |
| 1971 | HORVU4Hr1G090840 | ATP-dependent RNA helicase DeaD | -4.43 | AT3G06980.1 |
| 1972 | HORVU6Hr1G028250 | Cytochrome P450 superfamily protein | -4.43 | AT3G48280.1 |
| 1973 | HORVU7Hr1G024210 | P-loop containing nucleoside triphosphate hydrolases superfamily protein | -4.44 | AT1G73170.1 |
| 1974 | HORVU1Hr1G084520 | Photosynthetic NDH subunit of subcomplex B 3, chloroplastic | -4.44 | AT4G32590.1 |
| 1975 | HORVU2Hr1G011960 | DnaJ/Hsp40 cysteine-rich domain superfamily protein | -4.44 | AT2G34860.2 |
| 1976 | HORVU3Hr1G049900 | glyoxylate reductase 2 | -4.44 | AT1G17650.1 |

| | | | | |
|---|---|---|---|---|
| 1977 | HORVU2Hr1G003840 | Glutamyl-tRNA(Gln) amidotransferase subunit A | -4.44 | AT4G34880.2 |
| 1978 | HORVU2Hr1G072020 | Chaperone protein ClpB | -4.45 | AT5G51070.1 |
| 1979 | HORVU4Hr1G084830 | 50S ribosomal protein L11 | -4.45 | AT1G32990.1 |
| 1980 | HORVU1Hr1G072520 | undescribed protein | -4.46 | AT3G32040.1 |
| 1981 | HORVU5Hr1G049280 | undescribed protein | -4.46 | AT1G50730.1 |
| 1982 | HORVU0Hr1G012860 | Alcohol dehydrogenase | -4.46 | AT4G37980.1 |
| 1983 | HORVU3Hr1G116470 | Pectin acetylesterase 5 | -4.47 | AT3G09410.1 |
| 1984 | HORVU1Hr1G081370 | unknown protein; BEST Arabidopsis thaliana protein match is: unknown protein . | -4.48 | AT3G14830.4 |
| 1985 | HORVU7Hr1G116780 | Eukaryotic aspartyl protease family protein | -4.48 | AT3G59080.1 |
| 1986 | HORVU1Hr1G000430 | Chalcone--flavonone isomerase 1B-1 | -4.48 | AT1G53520.1 |
| 1987 | HORVU7Hr1G035320 | Tyrosine phosphatase family protein | -4.48 | AT1G05000.1 |
| 1988 | HORVU6Hr1G025110 | Elongation factor 4 | -4.48 | AT5G08650.1 |
| 1989 | HORVU6Hr1G076860 | DEAD/DEAH box helicase, putative | -4.49 | AT1G70070.1 |
| 1990 | HORVU3Hr1G039930 | Magnesium-protoporphyrin IX monomethyl ester [oxidative] cyclase 3 | -4.49 | AT3G56940.1 |
| 1991 | HORVU2Hr1G110900 | Protein kinase superfamily protein | -4.50 | AT3G13530.1 |
| 1992 | HORVU1Hr1G022800 | BEST Arabidopsis thaliana protein match is: 3'-5' exonuclease domain-containing protein / K homology domain-containing protein / KH domain-containing protein . | -4.51 | AT2G25920.1 |
| 1993 | HORVU2Hr1G104580 | At5g37260-like protein | -4.51 | AT5G37260.1 |
| 1994 | HORVU5Hr1G109440 | Lactoylglutathione lyase | -4.52 | AT1G67280.1 |
| 1995 | HORVU6Hr1G043920 | undescribed protein | -4.52 | AT1G06950.1 |
| 1996 | HORVU3Hr1G049730 | Chromosome 3B, genomic scaffold, cultivar Chinese Spring | -4.52 | AT4G14368.1 |
| 1997 | HORVU7Hr1G038770 | dehydroascorbate reductase 1 | -4.52 | AT5G16710.1 |
| 1998 | HORVU2Hr1G036280 | GRAS family transcription factor | -4.52 | AT5G48150.4 |
| 1999 | HORVU1Hr1G095410 | Zinc finger protein CONSTANS-LIKE 4 | -4.53 | AT5G53420.1 |
| 2000 | HORVU5Hr1G001180 | lipoxygenase 2 | -4.53 | AT3G45140.1 |
| 2001 | HORVU6Hr1G049050 | Protein kinase superfamily protein | -4.53 | AT5G64940.2 |
| 2002 | HORVU6Hr1G014140 | undescribed protein | -4.53 | AT4G14500.2 |
| 2003 | HORVU4Hr1G085930 | Protein kinase superfamily protein | -4.53 | AT3G61080.1 |
| 2004 | HORVU3Hr1G085420 | alpha-glucan phosphorylase 2 | -4.54 | AT3G46970.1 |
| 2005 | HORVU2Hr1G032170 | Acidic leucine-rich nuclear phosphoprotein 32-related protein 1 | -4.54 | AT1G09760.1 |
| 2006 | HORVU0Hr1G013890 | Peroxisomal (S)-2-hydroxy-acid oxidase | -4.54 | AT3G14420.2 |
| 2007 | HORVU3Hr1G013600 | unknown function | -4.55 | AT5G15802.1 |
| 2008 | HORVU7Hr1G117020 | Basic-leucine zipper (bZIP) transcription factor family protein | -4.56 | AT4G35040.4 |
| 2009 | HORVU4Hr1G011090 | ATP synthase delta-subunit gene | -4.56 | AT4G09650.1 |
| 2010 | HORVU7Hr1G091440 | CASP-like protein 1 | -4.56 | AT4G15630.1 |
| 2011 | HORVU5Hr1G043550 | Saccharopine dehydrogenase | -4.57 | AT1G50450.1 |

| | | | | |
|---|---|---|---|---|
| 2012 | HORVU3Hr1G025430 | undescribed protein | -4.57 | AT3G50370.1 |
| 2013 | HORVU4Hr1G027180 | proline transporter 1 | -4.57 | AT2G39890.2 |
| 2014 | HORVU6Hr1G025120 | Elongation factor 4 | -4.57 | AT5G08650.1 |
| 2015 | HORVU6Hr1G090510 | O-methyltransferase family protein | -4.57 | AT4G35160.1 |
| 2016 | HORVU5Hr1G020410 | unknown protein; BEST Arabidopsis thaliana protein match is: unknown protein . | -4.58 | AT5G61040.1 |
| 2017 | HORVU6Hr1G011760 | 0 | -4.58 | #N/A |
| 2018 | HORVU3Hr1G116490 | Pectin acetylesterase 5 | -4.59 | AT3G09410.1 |
| 2019 | HORVU7Hr1G082590 | Haloacid dehalogenase-like hydrolase (HAD) superfamily protein | -4.60 | AT3G48420.1 |
| 2020 | HORVU3Hr1G086240 | Calcium-dependent lipid-binding (CaLB domain) family protein | -4.60 | AT3G55470.2 |
| 2021 | HORVU1Hr1G061830 | Beta-propeller domain-containing protein, methanol dehydrogenase | -4.62 | AT1G54780.1 |
| 2022 | HORVU1Hr1G011520 | Glutamyl-tRNA reductase family protein | -4.62 | AT1G58290.1 |
| 2023 | HORVU6Hr1G074820 | Lysine-specific demethylase REF6 | -4.62 | AT3G48430.2 |
| 2024 | HORVU1Hr1G051150 | Antiholin-like protein LrgB | -4.62 | AT1G32080.1 |
| 2025 | HORVU3Hr1G080770 | Chromosome 3B, genomic scaffold, cultivar Chinese Spring | -4.63 | AT5G05190.1 |
| 2026 | HORVU1Hr1G067110 | Ethylene-responsive transcription factor 8 | -4.63 | AT5G44210.1 |
| 2027 | HORVU3Hr1G032490 | Triose phosphate/phosphate translocator, chloroplastic | -4.64 | AT5G46110.4 |
| 2028 | HORVU7Hr1G074890 | Succinate dehydrogenase [ubiquinone] iron-sulfur subunit 1, mitochondrial | -4.64 | AT5G40650.1 |
| 2029 | HORVU7Hr1G038120 | shikimate kinase like 1 | -4.66 | AT3G26900.3 |
| 2030 | HORVU5Hr1G013840 | L-galactose dehydrogenase | -4.66 | AT4G33670.1 |
| 2031 | HORVU7Hr1G114170 | 50S ribosomal protein L35 | -4.67 | AT2G24090.1 |
| 2032 | HORVU6Hr1G081670 | tyrosine aminotransferase 3 | -4.67 | AT5G36160.1 |
| 2033 | HORVU3Hr1G117810 | Zinc finger CCCH domain-containing protein 33 | -4.67 | AT3G55980.1 |
| 2034 | HORVU5Hr1G036160 | CBS domain-containing protein CBSCBSPB3 | -4.67 | AT3G52950.2 |
| 2035 | HORVU4Hr1G067320 | SprT-like domain-containing protein Spartan | -4.67 | AT3G24340.1 |
| 2036 | HORVU3Hr1G093260 | Protein of unknown function (DUF506) | -4.67 | AT3G07350.1 |
| 2037 | HORVU2Hr1G029430 | WRKY family transcription factor | -4.68 | AT4G11070.2 |
| 2038 | HORVU1Hr1G029850 | calmodulin like 23 | -4.68 | AT5G37770.2 |
| 2039 | HORVU5Hr1G056390 | Galactose mutarotase-like superfamily protein | -4.68 | AT5G66530.3 |
| 2040 | HORVU4Hr1G076050 | O-methyltransferase family protein | -4.69 | AT4G35160.1 |
| 2041 | HORVU3Hr1G006150 | unknown protein; INVOLVED IN: biological_process unknown; LOCATED IN: chloroplast; EXPRESSED IN: 22 plant structures; EXPRESSED DURING: 13 growth stages; Has 74 Blast hits to 74 proteins in 29 species: Archae - 0; Bacteria - 19; Metazoa - 0; Fungi - 0; Plants - 49; Viruses - 0; Other Eukaryotes - 6 (source: NCBI BLink). | -4.70 | AT1G18060.1 |
| 2042 | HORVU5Hr1G016350 | DA1 | -4.70 | AT4G36860.3 |
| 2043 | HORVU6Hr1G015120 | undescribed protein | -4.72 | AT1G78080.1 |

| | | | | |
|---|---|---|---|---|
| 2044 | HORVU2Hr1G032400 | alpha/beta-Hydrolases superfamily protein | -4.72 | AT1G13820.1 |
| 2045 | HORVU7Hr1G030800 | Pco114914 | -4.73 | AT2G28630.2 |
| 2046 | HORVU0Hr1G005340 | 0 | -4.73 | #N/A |
| 2047 | HORVU4Hr1G006730 | Plastid-lipid associated protein PAP / fibrillin family protein | -4.73 | AT3G26070.1 |
| 2048 | HORVU4Hr1G079710 | Divalent ion symporter | -4.74 | AT1G02260.2 |
| 2049 | HORVU2Hr1G073760 | Peroxiredoxin-2 | -4.74 | AT5G06290.1 |
| 2050 | HORVU3Hr1G002820 | receptor kinase 1 | -4.74 | AT4G05200.1 |
| 2051 | HORVU1Hr1G010450 | undescribed protein | -4.75 | AT1G31730.1 |
| 2052 | HORVU2Hr1G060480 | Photosystem I reaction center subunit psaK, chloroplastic | -4.75 | AT1G30380.1 |
| 2053 | HORVU4Hr1G053530 | PsbB mRNA maturation factor Mbb1, chloroplastic | -4.76 | AT3G17040.1 |
| 2054 | HORVU7Hr1G122750 | BnaA09g56450D protein | -4.76 | AT1G16080.1 |
| 2055 | HORVU2Hr1G032970 | 6-phosphogluconolactonase 1 | -4.77 | AT1G13700.3 |
| 2056 | HORVU5Hr1G020110 | Protein of unknown function (DUF1336) | -4.78 | AT3G29180.3 |
| 2057 | HORVU6Hr1G050160 | GTP binding Elongation factor Tu family protein | -4.78 | AT5G13650.2 |
| 2058 | HORVU0Hr1G005570 | Ribulose bisphosphate carboxylase/oxygenase activase 2, chloroplastic | -4.80 | AT1G73110.2 |
| 2059 | HORVU7Hr1G071440 | undescribed protein | -4.80 | AT3G01160.1 |
| 2060 | HORVU3Hr1G014890 | Disease resistance protein (CC-NBS-LRR class) family | -4.80 | AT3G07040.1 |
| 2061 | HORVU6Hr1G075850 | Aquaporin-like superfamily protein | -4.81 | AT5G37820.1 |
| 2062 | HORVU4Hr1G011600 | P-loop containing nucleoside triphosphate hydrolases superfamily protein | -4.81 | AT2G30630.1 |
| 2063 | HORVU5Hr1G091810 | RNA binding | -4.82 | AT4G24770.2 |
| 2064 | HORVU5Hr1G062780 | Abhydrolase domain-containing protein 1 | -4.82 | AT2G03140.6 |
| 2065 | HORVU3Hr1G030540 | Replication protein A 32 kDa subunit B | -4.82 | AT2G24490.2 |
| 2066 | HORVU7Hr1G023250 | unknown protein; BEST Arabidopsis thaliana protein match is: unknown protein . | -4.82 | AT3G14830.4 |
| 2067 | HORVU6Hr1G027910 | Testis-expressed sequence 2 protein | -4.83 | AT1G17820.1 |
| 2068 | HORVU0Hr1G007950 | undescribed protein | -4.83 | #N/A |
| 2069 | HORVU4Hr1G069050 | unknown protein; Has 38 Blast hits to 38 proteins in 17 species: Archae - 0; Bacteria - 0; Metazoa - 0; Fungi - 0; Plants - 38; Viruses - 0; Other Eukaryotes - 0 (source: NCBI BLink). | -4.84 | AT2G03420.1 |
| 2070 | HORVU2Hr1G073740 | Protein FAM210B | -4.85 | AT2G27290.1 |
| 2071 | HORVU1Hr1G064750 | EG2771 | -4.85 | AT5G14910.2 |
| 2072 | HORVU3Hr1G065700 | undescribed protein | -4.86 | AT3G28510.1 |
| 2073 | HORVU5Hr1G080690 | RHOMBOID-like protein 3 | -4.86 | AT1G63120.1 |
| 2074 | HORVU5Hr1G028670 | undescribed protein | -4.86 | AT3G03090.1 |
| 2075 | HORVU7Hr1G096430 | myb-like transcription factor family protein | -4.86 | AT3G13040.3 |
| 2076 | HORVU1Hr1G055710 | Ethylene-responsive transcription factor 2 | -4.87 | AT4G17490.1 |
| 2077 | HORVU4Hr1G031320 | methionine S-methyltransferase | -4.87 | AT5G49810.1 |
| 2078 | HORVU4Hr1G009080 | undescribed protein | -4.88 | AT1G16800.2 |

| | | | | |
|---|---|---|---|---|
| 2079 | HORVU6Hr1G020730 | DTW domain-containing protein | -4.88 | AT5G54880.1 |
| 2080 | HORVU7Hr1G080520 | unknown function | -4.88 | AT3G45750.1 |
| 2081 | HORVU2Hr1G003460 | ATP-dependent Clp protease adapter protein ClpS | -4.89 | AT1G68660.1 |
| 2082 | HORVU2Hr1G046740 | Exonuclease 3'-5' domain-containing protein 2 | -4.89 | AT2G32415.1 |
| 2083 | HORVU7Hr1G069930 | P0696G06.27 protein | -4.90 | AT3G59780.1 |
| 2084 | HORVU6Hr1G084390 | UDP-glucuronic acid 4-epimerase | -4.91 | #N/A |
| 2085 | HORVU3Hr1G078820 | Photosystem II reaction center PsbP family protein | -4.91 | AT3G56650.1 |
| 2086 | HORVU3Hr1G059950 | undescribed protein | -4.92 | AT5G47490.2 |
| 2087 | HORVU3Hr1G061690 | Protein DEHYDRATION-INDUCED 19 homolog 3 | -4.92 | AT3G05700.1 |
| 2088 | HORVU1Hr1G027830 | unknown protein; BEST Arabidopsis thaliana protein match is: unknown protein . | -4.92 | AT5G12010.1 |
| 2089 | HORVU2Hr1G043920 | beta-amylase 5 | -4.92 | AT4G15210.1 |
| 2090 | HORVU4Hr1G011120 | unknown function | -4.93 | AT3G28460.1 |
| 2091 | HORVU5Hr1G066350 | N-lysine methyltransferase setd6 | -4.93 | AT1G14030.1 |
| 2092 | HORVU3Hr1G037600 | respiratory burst oxidase homologue D | -4.94 | AT5G47910.1 |
| 2093 | HORVU2Hr1G088180 | triglyceride lipases;triglyceride lipases | -4.95 | AT4G13550.2 |
| 2094 | HORVU3Hr1G084590 | Isoprenyl transferase | -4.96 | AT5G58770.1 |
| 2095 | HORVU2Hr1G028160 | Protein of unknown function (DUF1997) | -4.96 | AT5G39530.1 |
| 2096 | HORVU3Hr1G114960 | undescribed protein | -4.96 | AT2G29525.4 |
| 2097 | HORVU3Hr1G085970 | Protein of unknown function (DUF668) | -4.96 | AT3G23160.1 |
| 2098 | HORVU6Hr1G021460 | Zinc finger protein CONSTANS-LIKE 7 | -4.97 | AT5G57180.2 |
| 2099 | HORVU7Hr1G034290 | Protein kinase superfamily protein | -4.97 | AT2G05940.1 |
| 2100 | HORVU7Hr1G029720 | RING/U-box superfamily protein | -4.97 | AT2G15580.1 |
| 2101 | HORVU1Hr1G070250 | WRKY DNA-binding protein 33 | -4.97 | AT2G38470.1 |
| 2102 | HORVU3Hr1G096830 | Carboxypeptidase Y homolog A | -4.98 | AT4G12910.1 |
| 2103 | HORVU5Hr1G073950 | dihydroflavonol 4-reductase-like1 | -4.98 | AT2G33590.1 |
| 2104 | HORVU4Hr1G086250 | shikimate kinase like 1 | -4.98 | AT3G26900.3 |
| 2105 | HORVU3Hr1G116000 | undescribed protein | -4.99 | AT3G27040.1 |
| 2106 | HORVU7Hr1G101740 | Calcium-binding EF-hand family protein | -5.00 | AT4G27280.1 |
| 2107 | HORVU6Hr1G081330 | Protein kinase superfamily protein | -5.01 | AT2G05940.1 |
| 2108 | HORVU0Hr1G017440 | Homoserine O-acetyltransferase | -5.01 | AT3G15900.1 |
| 2109 | HORVU7Hr1G074600 | unknown function | -5.01 | AT5G47920.1 |
| 2110 | HORVU6Hr1G090160 | unknown function | -5.02 | AT5G40810.2 |
| 2111 | HORVU6Hr1G091260 | Auxin-responsive protein IAA10 | -5.02 | AT2G33310.2 |
| 2112 | HORVU5Hr1G103320 | Accelerated cell death 11 | -5.02 | AT2G34690.1 |
| 2113 | HORVU4Hr1G002700 | Protein of unknown function (DUF581) | -5.02 | AT3G22550.1 |
| 2114 | HORVU1Hr1G051360 | plastid transcriptionally active 18 | -5.03 | AT2G32180.2 |
| 2115 | HORVU2Hr1G095070 | S-adenosyl-L-methionine-dependent methyltransferases superfamily protein | -5.03 | AT4G00750.1 |
| 2116 | HORVU2Hr1G094090 | RING zinc finger protein-like | -5.03 | #N/A |
| 2117 | HORVU7Hr1G113560 | receptor-like protein kinase 4 | -5.04 | AT5G10530.1 |

| | | | | |
|---|---|---|---|---|
| 2118 | HORVU1Hr1G069430 | Acetyltransferase NSI | -5.05 | AT1G32070.1 |
| 2119 | HORVU1Hr1G078140 | Photosystem II reaction center W protein, chloroplastic | -5.06 | AT2G30570.1 |
| 2120 | HORVU1Hr1G060740 | Lipase | -5.06 | AT1G10740.4 |
| 2121 | HORVU2Hr1G041080 | sugar transporter 1 | -5.06 | AT1G11260.1 |
| 2122 | HORVU3Hr1G035230 | FHA domain-containing protein | -5.07 | AT3G02400.1 |
| 2123 | HORVU3Hr1G072260 | vascular plant one zinc finger protein | -5.07 | AT1G28520.5 |
| 2124 | HORVU6Hr1G088200 | Protein kinase superfamily protein | -5.08 | AT2G39190.2 |
| 2125 | HORVU4Hr1G020080 | Protein of unknown function (DUF793) | -5.09 | AT1G18740.1 |
| 2126 | HORVU5Hr1G071680 | 0 | -5.09 | #N/A |
| 2127 | HORVU7Hr1G067990 | unknown function | -5.09 | AT5G38660.1 |
| 2128 | HORVU3Hr1G070380 | Chromosome 3B, genomic scaffold, cultivar Chinese Spring | -5.10 | AT5G56550.1 |
| 2129 | HORVU6Hr1G012320 | Glutathione-specific gamma-glutamylcyclotransferase 1 | -5.10 | AT4G31290.1 |
| 2130 | HORVU4Hr1G078440 | unknown function | -5.10 | AT3G25590.1 |
| 2131 | HORVU0Hr1G005920 | Pentatricopeptide repeat-containing protein | -5.10 | AT5G27270.1 |
| 2132 | HORVU1Hr1G087600 | Glutaredoxin family protein | -5.11 | AT1G28480.1 |
| 2133 | HORVU2Hr1G041620 | undescribed protein | -5.11 | AT4G36440.1 |
| 2134 | HORVU5Hr1G037920 | histone deacetylase 14 | -5.11 | AT4G33470.1 |
| 2135 | HORVU2Hr1G019270 | undescribed protein | -5.11 | AT2G06520.1 |
| 2136 | HORVU3Hr1G086970 | Metal tolerance protein 5 | -5.11 | AT2G39450.1 |
| 2137 | HORVU7Hr1G010790 | aspartate aminotransferase 1 | -5.12 | AT2G30970.2 |
| 2138 | HORVU6Hr1G072740 | calcium sensing receptor | -5.12 | AT5G23060.1 |
| 2139 | HORVU5Hr1G011100 | Protoporphyrinogen oxidase | -5.12 | AT4G01690.1 |
| 2140 | HORVU4Hr1G048440 | RNA-binding protein 1 | -5.12 | AT2G37220.1 |
| 2141 | HORVU1Hr1G093430 | BnaC06g06240D protein | -5.13 | AT1G53250.1 |
| 2142 | HORVU7Hr1G075700 | Reticulon family protein | -5.13 | AT4G11220.1 |
| 2143 | HORVU3Hr1G073740 | unknown protein; LOCATED IN: chloroplast; Has 35333 Blast hits to 34131 proteins in 2444 species: Archae - 798; Bacteria - 22429; Metazoa - 974; Fungi - 991; Plants - 531; Viruses - 0; Other Eukaryotes - 9610 (source: NCBI BLink). | -5.13 | AT5G55570.1 |
| 2144 | HORVU4Hr1G016210 | unknown function | -5.14 | AT1G32520.1 |
| 2145 | HORVU3Hr1G098160 | Calcium-dependent lipid-binding (CaLB domain) family protein | -5.14 | AT1G09070.1 |
| 2146 | HORVU7Hr1G100790 | Remorin family protein | -5.14 | AT3G57540.1 |
| 2147 | HORVU5Hr1G095990 | Lipid A export ATP-binding/permease protein MsbA | -5.15 | AT5G39040.1 |
| 2148 | HORVU5Hr1G082630 | triosephosphate isomerase | -5.15 | AT2G21170.3 |
| 2149 | HORVU3Hr1G094600 | HCP-like superfamily protein with MYND-type zinc finger | -5.16 | AT1G67340.1 |
| 2150 | HORVU6Hr1G000640 | SKP1-like 4 | -5.16 | AT5G42190.1 |
| 2151 | HORVU0Hr1G015530 | receptor kinase 2 | -5.17 | AT1G16670.1 |

| | | | | |
|---|---|---|---|---|
| 2152 | HORVU7Hr1G043360 | undescribed protein | -5.17 | AT5G52630.1 |
| 2153 | HORVU7Hr1G097250 | Peptidyl-prolyl cis-trans isomerase B | -5.18 | AT5G35100.3 |
| 2154 | HORVU3Hr1G014850 | Chromosome 3B, genomic scaffold, cultivar Chinese Spring | -5.20 | AT3G23740.1 |
| 2155 | HORVU7Hr1G000630 | Disease resistance protein RPP13 | -5.20 | AT3G46530.1 |
| 2156 | HORVU3Hr1G112930 | Disease resistance protein | -5.20 | AT3G07040.1 |
| 2157 | HORVU3Hr1G013770 | GTPase Der | -5.21 | AT3G12080.1 |
| 2158 | HORVU1Hr1G010440 | undescribed protein | -5.21 | AT4G23470.1 |
| 2159 | HORVU4Hr1G042550 | undescribed protein | -5.22 | AT2G29640.1 |
| 2160 | HORVU2Hr1G059910 | undescribed protein | -5.23 | AT2G07240.1 |
| 2161 | HORVU1Hr1G088270 | IMP dehydrogenase/GMP reductase | -5.23 | AT5G01930.1 |
| 2162 | HORVU3Hr1G089550 | ribonuclease 2 | -5.24 | AT2G39780.1 |
| 2163 | HORVU1Hr1G019770 | Protein kinase superfamily protein | -5.25 | AT2G05940.1 |
| 2164 | HORVU6Hr1G071930 | DNA-directed RNA polymerase II subunit 4 | -5.25 | AT4G15950.1 |
| 2165 | HORVU3Hr1G062490 | 30S ribosomal protein S20 | -5.25 | AT3G15190.1 |
| 2166 | HORVU5Hr1G039960 | GCN5 family acetyltransferase | -5.26 | AT1G72030.1 |
| 2167 | HORVU1Hr1G082540 | Protein phosphatase 2C family protein | -5.26 | AT4G33500.1 |
| 2168 | HORVU4Hr1G052470 | 30S ribosomal protein S1 homolog A | -5.27 | AT5G30510.1 |
| 2169 | HORVU2Hr1G096360 | Aquaporin-like superfamily protein | -5.27 | AT4G00430.1 |
| 2170 | HORVU0Hr1G008850 | ACT domain repeat 8 | -5.28 | AT1G12420.2 |
| 2171 | HORVU4Hr1G024290 | undescribed protein | -5.28 | AT4G02640.4 |
| 2172 | HORVU1Hr1G075580 | oxidative stress 3 . | -5.28 | AT5G56550.1 |
| 2173 | HORVU3Hr1G107070 | Phycoerythrobilin:ferredoxin oxidoreductase | -5.28 | AT3G09150.3 |
| 2174 | HORVU3Hr1G065770 | Bidirectional sugar transporter N3 | -5.29 | AT3G14770.1 |
| 2175 | HORVU0Hr1G022820 | NAC domain containing protein 94 | -5.31 | AT2G17040.1 |
| 2176 | HORVU6Hr1G061870 | cyclin p4;3 | -5.31 | AT5G07450.1 |
| 2177 | HORVU4Hr1G054910 | xyloglucan xylosyltransferase 5 | -5.33 | AT3G62720.2 |
| 2178 | HORVU3Hr1G094820 | thylakoid rhodanese-like | -5.33 | AT3G25480.1 |
| 2179 | HORVU5Hr1G011960 | Short-chain dehydrogenase/reductase family protein | -5.33 | AT4G20760.1 |
| 2180 | HORVU5Hr1G038630 | ribulose bisphosphate carboxylase small chain 1A | -5.34 | AT1G67090.1 |
| 2181 | HORVU2Hr1G098650 | Protein argonaute 2 | -5.34 | AT1G31280.1 |
| 2182 | HORVU1Hr1G013250 | undescribed protein | -5.34 | AT1G23450.1 |
| 2183 | HORVU6Hr1G009510 | chlororespiratory reduction 6 | -5.34 | AT2G47910.1 |
| 2184 | HORVU7Hr1G090800 | Lipid A export ATP-binding/permease protein MsbA | -5.35 | AT4G25450.3 |
| 2185 | HORVU5Hr1G059570 | unknown function | -5.35 | AT1G62780.1 |
| 2186 | HORVU2Hr1G080260 | Oxygen-evolving enhancer protein 3-1, chloroplastic | -5.36 | AT3G01440.1 |
| 2187 | HORVU2Hr1G079970 | undescribed protein | -5.36 | AT5G57200.1 |
| 2188 | HORVU2Hr1G115690 | Protein kinase superfamily protein | -5.36 | AT1G71810.1 |
| 2189 | HORVU3Hr1G014860 | Protein of unknown function (DUF 3339) | -5.37 | AT5G63500.1 |

| | | | | |
|---|---|---|---|---|
| 2190 | HORVU5Hr1G028460 | Plant calmodulin-binding protein-related | -5.37 | AT5G15430.2 |
| 2191 | HORVU4Hr1G060970 | 50S ribosomal protein L4 | -5.38 | AT1G07320.2 |
| 2192 | HORVU6Hr1G063820 | undescribed protein | -5.40 | AT1G11420.1 |
| 2193 | HORVU6Hr1G062040 | 50S ribosomal protein L19 | -5.41 | AT4G17560.1 |
| 2194 | HORVU4Hr1G082040 | catalase 2 | -5.41 | AT4G35090.1 |
| 2195 | HORVU2Hr1G007350 | flowering promoting factor 1 | -5.41 | AT5G24860.1 |
| 2196 | HORVU1Hr1G004400 | unknown function | -5.41 | AT5G02160.1 |
| 2197 | HORVU0Hr1G000960 | Nuclear transport factor 2 (NTF2) family protein | -5.41 | AT3G04890.1 |
| 2198 | HORVU4Hr1G060130 | RING/U-box superfamily protein | -5.43 | AT5G59550.1 |
| 2199 | HORVU0Hr1G008540 | undescribed protein | -5.46 | AT5G26850.4 |
| 2200 | HORVU3Hr1G014780 | shikimate kinase like 1 | -5.46 | AT3G26900.3 |
| 2201 | HORVU2Hr1G015720 | Hexosyltransferase | -5.46 | AT1G56600.1 |
| 2202 | HORVU5Hr1G069380 | Heavy metal transport/detoxification superfamily protein | -5.46 | AT1G22990.1 |
| 2203 | HORVU7Hr1G000900 | plastocyanin 1 | -5.46 | AT1G76100.1 |
| 2204 | HORVU1Hr1G018010 | unknown function | -5.47 | AT3G62070.1 |
| 2205 | HORVU7Hr1G091570 | Endoglucanase 18 | -5.47 | AT1G19940.2 |
| 2206 | HORVU2Hr1G113790 | undescribed protein | -5.47 | AT3G13100.1 |
| 2207 | HORVU4Hr1G060630 | Fructose-1,6-bisphosphatase class 1 | -5.48 | AT3G54050.2 |
| 2208 | HORVU2Hr1G017420 | Stress responsive protein | -5.48 | AT2G39050.1 |
| 2209 | HORVU5Hr1G098340 | 30S ribosomal protein S9 | -5.48 | AT1G74970.1 |
| 2210 | HORVU0Hr1G023150 | Protein of unknown function (DUF1218) | -5.48 | AT1G31720.6 |
| 2211 | HORVU5Hr1G111310 | magnesium chelatase i2 | -5.49 | AT1G08520.1 |
| 2212 | HORVU0Hr1G021950 | Glucan endo-1,3-beta-glucosidase 14 | -5.49 | AT2G26600.1 |
| 2213 | HORVU1Hr1G061020 | unknown function | -5.49 | AT4G15545.1 |
| 2214 | HORVU6Hr1G085120 | C2 domain-containing protein / GRAM domain-containing protein | -5.51 | AT3G59660.1 |
| 2215 | HORVU2Hr1G037620 | Lactation elevated protein 1 | -5.51 | AT4G15810.1 |
| 2216 | HORVU1Hr1G077160 | unknown protein; Has 640 Blast hits to 638 proteins in 201 species: Archae - 0; Bacteria - 293; Metazoa - 0; Fungi - 71; Plants - 72; Viruses - 0; Other Eukaryotes - 204 (source: NCBI BLink). | -5.51 | AT3G01060.3 |
| 2217 | HORVU5Hr1G087920 | plastid movement impaired1 | -5.52 | AT1G42550.1 |
| 2218 | HORVU3Hr1G114220 | ATP-dependent RNA helicase RhlB | -5.52 | AT5G19210.2 |
| 2219 | HORVU2Hr1G075180 | undescribed protein | -5.52 | AT5G61350.1 |
| 2220 | HORVU4Hr1G086110 | F-box protein PP2-B1 | -5.52 | AT3G61060.1 |
| 2221 | HORVU6Hr1G054430 | methionine aminopeptidase 1B | -5.53 | AT1G13270.1 |
| 2222 | HORVU1Hr1G038620 | plastid transcriptionally active 3 | -5.54 | AT3G04260.1 |
| 2223 | HORVU6Hr1G053680 | Elongation factor Tu | -5.54 | AT4G20360.2 |
| 2224 | HORVU2Hr1G113850 | undescribed protein | -5.54 | AT4G14465.1 |
| 2225 | HORVU5Hr1G122820 | Ankyrin repeat domain-containing protein 2A | -5.54 | AT4G35450.3 |

| | | | | |
|---|---|---|---|---|
| 2226 | HORVU5Hr1G056130 | WRKY family transcription factor | -5.54 | AT4G11070.2 |
| 2227 | HORVU4Hr1G003040 | regulatory protein (NPR1) | -5.55 | AT5G45110.1 |
| 2228 | HORVU5Hr1G073330 | Lysine-specific demethylase 8 | -5.55 | AT3G20810.1 |
| 2229 | HORVU6Hr1G005920 | NADPH:quinone oxidoreductase | -5.56 | AT3G27890.1 |
| 2230 | HORVU3Hr1G063280 | D-glycerate 3-kinase, chloroplastic | -5.57 | AT1G80380.2 |
| 2231 | HORVU2Hr1G120490 | Two-component response regulator ORR6 | -5.59 | AT3G57040.2 |
| 2232 | HORVU5Hr1G024470 | elongation factor Ts family protein | -5.59 | AT4G29060.1 |
| 2233 | HORVU0Hr1G031850 | myb-like transcription factor family protein | -5.59 | AT5G52660.1 |
| 2234 | HORVU1Hr1G041640 | undescribed protein | -5.60 | AT1G37020.1 |
| 2235 | HORVU5Hr1G049980 | Protein translocase subunit SecY | -5.61 | AT2G18710.1 |
| 2236 | HORVU2Hr1G037610 | undescribed protein | -5.61 | #N/A |
| 2237 | HORVU6Hr1G078290 | 50S ribosomal protein L29 | -5.61 | AT5G65220.1 |
| 2238 | HORVU0Hr1G017140 | Methyltransferase-like protein 7A | -5.62 | AT1G69523.1 |
| 2239 | HORVU1Hr1G046550 | undescribed protein | -5.63 | AT2G38130.2 |
| 2240 | HORVU6Hr1G025170 | undescribed protein | -5.64 | #N/A |
| 2241 | HORVU6Hr1G087720 | Thioredoxin-like 2-2, chloroplastic | -5.64 | AT1G76080.1 |
| 2242 | HORVU3Hr1G003410 | tryptophan synthase alpha chain | -5.64 | AT4G02610.1 |
| 2243 | HORVU6Hr1G025730 | Quinone oxidoreductase-like protein 2 homolog | -5.65 | AT1G23740.1 |
| 2244 | HORVU1Hr1G074940 | Tubby-like F-box protein 8 | -5.65 | AT1G25280.1 |
| 2245 | HORVU5Hr1G037660 | amino acid permease 3 | -5.65 | AT5G09220.1 |
| 2246 | HORVU2Hr1G068880 | alpha/beta-Hydrolases superfamily protein | -5.66 | AT5G38360.1 |
| 2247 | HORVU7Hr1G034430 | receptor-like protein kinase 2 | -5.67 | AT2G26330.1 |
| 2248 | HORVU3Hr1G055260 | myb domain protein 66 | -5.68 | AT4G01060.1 |
| 2249 | HORVU7Hr1G109380 | Peptidyl-prolyl cis-trans isomerase | -5.68 | AT5G45680.1 |
| 2250 | HORVU2Hr1G093220 | Pentatricopeptide repeat-containing protein | -5.68 | AT1G02150.1 |
| 2251 | HORVU1Hr1G068500 | unknown function | -5.69 | AT3G51510.1 |
| 2252 | HORVU3Hr1G068300 | Cytosolic Fe-S cluster assembly factor NUBP2 homolog | -5.69 | AT3G24430.1 |
| 2253 | HORVU5Hr1G120250 | 30S ribosomal protein S6 | -5.70 | AT1G64510.1 |
| 2254 | HORVU1Hr1G072180 | Riboflavin biosynthesis protein RibBA | -5.70 | AT5G59750.1 |
| 2255 | HORVU5Hr1G124520 | unknown function | -5.71 | AT1G17850.2 |
| 2256 | HORVU5Hr1G118680 | Mitochondrial transcription termination factor family protein | -5.72 | AT4G09620.2 |
| 2257 | HORVU7Hr1G067110 | rubredoxin family protein | -5.73 | AT5G17170.1 |
| 2258 | HORVU5Hr1G046250 | RING/U-box superfamily protein | -5.73 | AT5G01980.1 |
| 2259 | HORVU4Hr1G054770 | COBRA-like protein 7 | -5.74 | AT4G16120.1 |
| 2260 | HORVU6Hr1G067470 | xyloglucan endotransglucosylase/hydrolase 28 | -5.74 | AT2G01850.1 |
| 2261 | HORVU1Hr1G070190 | Nodulin-like / Major Facilitator Superfamily protein | -5.74 | AT2G30300.1 |
| 2262 | HORVU2Hr1G121200 | nodulin MtN21 /EamA-like transporter family protein | -5.75 | AT5G64700.1 |
| 2263 | HORVU7Hr1G102730 | Protein of unknown function, DUF642 | -5.76 | AT5G25460.1 |

| | | | | |
|---|---|---|---|---|
| 2264 | HORVU2Hr1G091860 | DNA helicase INO80 | -5.77 | AT5G66750.1 |
| 2265 | HORVU2Hr1G112450 | purple acid phosphatase 21 | -5.77 | AT3G52810.1 |
| 2266 | HORVU1Hr1G053510 | GRAS family transcription factor | -5.78 | AT4G00150.1 |
| 2267 | HORVU5Hr1G050120 | undescribed protein | -5.79 | AT3G53210.2 |
| 2268 | HORVU4Hr1G016510 | Chaperone protein DnaJ | -5.80 | AT4G09350.1 |
| 2269 | HORVU5Hr1G068520 | EF hand family protein | -5.80 | AT1G66400.1 |
| 2270 | HORVU5Hr1G081750 | alpha/beta hydrolase | -5.81 | AT4G25290.1 |
| 2271 | HORVU1Hr1G022110 | Ran guanine nucleotide release factor | -5.81 | AT1G69680.1 |
| 2272 | HORVU5Hr1G103440 | zinc-finger protein 1 | -5.82 | AT5G48890.2 |
| 2273 | HORVU0Hr1G001280 | Calmodulin | -5.83 | AT3G43810.1 |
| 2274 | HORVU6Hr1G014080 | StAR-related lipid transfer protein 7, mitochondrial | -5.83 | AT1G64720.1 |
| 2275 | HORVU2Hr1G089030 | Plastid transcriptionally active 5, putative isoform 1 | -5.83 | AT4G13670.1 |
| 2276 | HORVU7Hr1G105500 | Nuclear transcription factor Y subunit B-2 | -5.83 | AT4G14540.1 |
| 2277 | HORVU2Hr1G115920 | Protein kinase superfamily protein | -5.84 | AT3G09830.3 |
| 2278 | HORVU5Hr1G118100 | Transposable element protein, putative | -5.84 | AT3G28674.1 |
| 2279 | HORVU6Hr1G068640 | syntaxin, putative | -5.84 | AT5G46860.1 |
| 2280 | HORVU1Hr1G091520 | Transcription factor RADIALIS | -5.85 | AT1G75250.2 |
| 2281 | HORVU1Hr1G017120 | Protein kinase superfamily protein | -5.85 | AT2G47060.4 |
| 2282 | HORVU3Hr1G018800 | RING/U-box superfamily protein | -5.86 | AT4G34040.1 |
| 2283 | HORVU5Hr1G094210 | Protein transport protein sec23-1 | -5.86 | AT5G43670.1 |
| 2284 | HORVU3Hr1G020640 | Chromosome 3B, genomic scaffold, cultivar Chinese Spring | -5.86 | AT4G01935.1 |
| 2285 | HORVU5Hr1G039210 | Chaperone DnaJ-domain superfamily protein | -5.86 | AT5G21430.1 |
| 2286 | HORVU4Hr1G060140 | Pentatricopeptide repeat-containing protein | -5.86 | AT5G59600.1 |
| 2287 | HORVU2Hr1G106880 | chloroplastic lipocalin | -5.87 | AT3G47860.1 |
| 2288 | HORVU1Hr1G050130 | Mitochondrial substrate carrier family protein | -5.87 | AT5G46800.3 |
| 2289 | HORVU1Hr1G005600 | ACT domain-containing protein | -5.87 | AT2G39570.1 |
| 2290 | HORVU3Hr1G083350 | Fructose-1,6-bisphosphatase class 1 | -5.89 | AT1G43670.1 |
| 2291 | HORVU7Hr1G023600 | Thiocyanate methyltransferase 1 | -5.91 | AT2G43940.1 |
| 2292 | HORVU2Hr1G112330 | purple acid phosphatase 22 | -5.92 | AT3G52820.1 |
| 2293 | HORVU7Hr1G051200 | NAD(P)H-quinone oxidoreductase subunit 3, chloroplastic | -5.93 | ATCG00440.1 |
| 2294 | HORVU3Hr1G018110 | Protein phosphatase PTC7 homolog | -5.95 | AT2G30170.1 |
| 2295 | HORVU1Hr1G022660 | Chlororespiratory reduction31 | -5.96 | AT4G23890.1 |
| 2296 | HORVU1Hr1G061230 | resistance to phytophthora 1 | -5.98 | AT2G48070.2 |
| 2297 | HORVU2Hr1G089940 | Aquaporin-like superfamily protein | -5.98 | AT2G37170.2 |
| 2298 | HORVU6Hr1G013670 | cellulose synthase 1 | -5.98 | AT4G32410.1 |
| 2299 | HORVU6Hr1G020960 | UDP-Glycosyltransferase superfamily protein | -5.98 | AT4G01070.1 |
| 2300 | HORVU1Hr1G049760 | Homeobox protein BEL1 homolog | -5.99 | AT5G41410.1 |
| 2301 | HORVU1Hr1G090150 | undescribed protein | -5.99 | AT3G48110.1 |

| | | | | |
|---|---|---|---|---|
| 2302 | HORVU3Hr1G067760 | S-adenosyl-L-methionine-dependent methyltransferases superfamily protein | -6.02 | AT4G22530.2 |
| 2303 | HORVU7Hr1G036450 | O-fucosyltransferase family protein | -6.02 | AT1G22460.1 |
| 2304 | HORVU1Hr1G001890 | Tubulin-specific chaperone E | -6.03 | AT1G71440.1 |
| 2305 | HORVU5Hr1G010870 | Glutaredoxin family protein | -6.03 | AT2G47870.1 |
| 2306 | HORVU6Hr1G064620 | Dehydrin COR410 | -6.05 | AT1G20440.1 |
| 2307 | HORVU1Hr1G093180 | RNA polymerase sigma-B factor | -6.05 | AT5G24120.3 |
| 2308 | HORVU7Hr1G108930 | F-box family protein | -6.06 | AT5G07610.1 |
| 2309 | HORVU6Hr1G054110 | Protein HHL1, chloroplastic | -6.07 | AT1G67700.4 |
| 2310 | HORVU7Hr1G060130 | Superoxide dismutase [Cu-Zn] 4A | -6.07 | AT2G28190.1 |
| 2311 | HORVU3Hr1G086380 | ENTH/VHS family protein | -6.08 | AT3G46540.3 |
| 2312 | HORVU4Hr1G076000 | O-methyltransferase family protein | -6.08 | AT4G35160.1 |
| 2313 | HORVU2Hr1G079920 | Chlorophyll a-b binding protein, chloroplastic | -6.08 | AT1G15820.1 |
| 2314 | HORVU6Hr1G001690 | RING/U-box superfamily protein | -6.12 | AT5G60580.3 |
| 2315 | HORVU3Hr1G070360 | Chromosome 3B, genomic scaffold, cultivar Chinese Spring | -6.13 | AT4G26288.1 |
| 2316 | HORVU3Hr1G070690 | hexokinase 1 | -6.13 | AT4G29130.1 |
| 2317 | HORVU5Hr1G021340 | Rubredoxin-like superfamily protein | -6.14 | AT1G54500.1 |
| 2318 | HORVU5Hr1G097150 | Subtilisin-like protease | -6.14 | AT5G67360.1 |
| 2319 | HORVU4Hr1G041010 | undescribed protein | -6.15 | AT1G25290.2 |
| 2320 | HORVU2Hr1G118060 | Heavy metal transport/detoxification superfamily protein | -6.15 | AT1G01490.2 |
| 2321 | HORVU4Hr1G052990 | haloacid dehalogenase-like hydrolase family protein | -6.18 | AT1G56500.3 |
| 2322 | HORVU3Hr1G105260 | unknown function | -6.18 | AT4G31560.1 |
| 2323 | HORVU7Hr1G052500 | Oxysterol-binding protein-related protein 4C | -6.18 | AT5G57240.5 |
| 2324 | HORVU1Hr1G041990 | RING/FYVE/PHD zinc finger superfamily protein | -6.19 | AT2G45530.1 |
| 2325 | HORVU6Hr1G049120 | Photosystem II reaction center protein M | -6.20 | ATCG00220.1 |
| 2326 | HORVU5Hr1G087830 | Histone superfamily protein | -6.20 | AT5G59970.2 |
| 2327 | HORVU5Hr1G073870 | dihydroflavonol 4-reductase-like1 | -6.21 | AT1G76470.1 |
| 2328 | HORVU7Hr1G036090 | kinesin 4 | -6.21 | AT5G10470.1 |
| 2329 | HORVU2Hr1G080510 | undescribed protein | -6.22 | AT2G47800.1 |
| 2330 | HORVU1Hr1G004650 | purple acid phosphatase 22 | -6.23 | AT3G52820.1 |
| 2331 | HORVU2Hr1G101720 | Multiple organellar RNA editing factor 5, mitochondrial | -6.23 | AT1G32580.1 |
| 2332 | HORVU6Hr1G071520 | Eukaryotic aspartyl protease family protein | -6.25 | AT5G10770.1 |
| 2333 | HORVU4Hr1G009380 | Lariat debranching enzyme | -6.25 | AT3G22436.1 |
| 2334 | HORVU7Hr1G109580 | tRNA pseudouridine synthase A | -6.26 | AT5G35400.1 |
| 2335 | HORVU2Hr1G123390 | unknown function | -6.26 | AT4G10220.1 |
| 2336 | HORVU5Hr1G117780 | Nodulation signaling protein 1 | -6.26 | AT3G13840.1 |
| 2337 | HORVU2Hr1G104740 | RAB GDP dissociation inhibitor 2 | -6.27 | AT2G44100.2 |
| 2338 | HORVU2Hr1G071850 | Protein phosphatase 2C family protein | -6.27 | AT3G16560.4 |
| 2339 | HORVU4Hr1G014080 | enzyme binding;tetrapyrrole binding | -6.28 | AT3G59400.1 |

| | | | | |
|---|---|---|---|---|
| 2340 | HORVU2Hr1G101340 | undescribed protein | -6.28 | #N/A |
| 2341 | HORVU5Hr1G100990 | Obg-like ATPase 1 | -6.29 | AT1G56050.1 |
| 2342 | HORVU7Hr1G076840 | vesicle-associated membrane protein 725 | -6.30 | AT4G10170.2 |
| 2343 | HORVU2Hr1G069880 | Protein of unknown function (DUF 3339) | -6.31 | AT3G48660.1 |
| 2344 | HORVU4Hr1G000230 | PsbP-like protein 1 | -6.31 | AT3G55330.1 |
| 2345 | HORVU6Hr1G012270 | terpene synthase 14 | -6.32 | AT1G61680.1 |
| 2346 | HORVU3Hr1G062250 | Chlororespiratory reduction 41 | -6.32 | AT1G51100.1 |
| 2347 | HORVU5Hr1G118120 | unknown function | -6.36 | AT1G40087.1 |
| 2348 | HORVU1Hr1G054190 | unknown function | -6.36 | AT5G65750.1 |
| 2349 | HORVU3Hr1G070340 | Protein-tyrosine phosphatase mitochondrial 1-like protein | -6.37 | AT2G35680.1 |
| 2350 | HORVU2Hr1G113050 | sulfate transporter 91 | -6.37 | AT1G23090.1 |
| 2351 | HORVU4Hr1G037290 | Ferredoxin-thioredoxin reductase, catalytic chain | -6.39 | AT2G04700.3 |
| 2352 | HORVU2Hr1G066790 | UDP-Glycosyltransferase superfamily protein | -6.39 | AT1G22360.1 |
| 2353 | HORVU1Hr1G014240 | undescribed protein | -6.40 | AT1G70290.1 |
| 2354 | HORVU6Hr1G085170 | Photosystem II 10 kDa polypeptide | -6.40 | AT1G79040.1 |
| 2355 | HORVU6Hr1G009350 | Acylamino-acid-releasing enzyme | -6.42 | AT4G14570.1 |
| 2356 | HORVU7Hr1G040790 | U-box domain-containing protein 21 | -6.43 | AT5G37490.1 |
| 2357 | HORVU1Hr1G045520 | Thylakoid lumenal 17.4 kDa protein, chloroplastic | -6.43 | AT5G53490.2 |
| 2358 | HORVU5Hr1G084490 | Eukaryotic aspartyl protease family protein | -6.43 | AT4G35880.1 |
| 2359 | HORVU1Hr1G069180 | Protein LURP-one-related 8 | -6.45 | AT2G38640.1 |
| 2360 | HORVU6Hr1G067660 | Phosphoribulokinase / Uridine kinase family | -6.46 | AT1G32060.1 |
| 2361 | HORVU6Hr1G011670 | unknown function | -6.47 | AT2G24960.1 |
| 2362 | HORVU1Hr1G068960 | MATE efflux family protein | -6.48 | AT1G51340.1 |
| 2363 | HORVU1Hr1G046360 | undescribed protein | -6.49 | AT5G49110.4 |
| 2364 | HORVU1Hr1G031930 | undescribed protein | -6.50 | AT1G55640.1 |
| 2365 | HORVU2Hr1G121110 | auxin response factor 6 | -6.50 | AT5G37020.1 |
| 2366 | HORVU5Hr1G098890 | malate dehydrogenase | -6.50 | AT2G22780.1 |
| 2367 | HORVU1Hr1G031940 | undescribed protein | -6.51 | AT4G19650.2 |
| 2368 | HORVU5Hr1G098980 | HAD-superfamily hydrolase, subfamily IG, 5'-nucleotidase | -6.52 | AT1G75210.3 |
| 2369 | HORVU3Hr1G094270 | Chromosome 3B, genomic scaffold, cultivar Chinese Spring | -6.52 | AT2G35450.1 |
| 2370 | HORVU5Hr1G039850 | actin 7 | -6.52 | AT5G09810.1 |
| 2371 | HORVU3Hr1G095700 | metacaspase 1 | -6.52 | AT1G02170.1 |
| 2372 | HORVU3Hr1G112010 | Abscisic stress-ripening protein 1 | -6.53 | #N/A |
| 2373 | HORVU0Hr1G000900 | undescribed protein | -6.53 | AT5G17310.5 |
| 2374 | HORVU2Hr1G077630 | Histone H2B.1 | -6.54 | AT3G45980.1 |
| 2375 | HORVU6Hr1G068760 | Inorganic pyrophosphatase | -6.54 | AT1G01050.2 |
| 2376 | HORVU5Hr1G033540 | ribulose bisphosphate carboxylase small chain 1A | -6.55 | AT1G67090.1 |
| 2377 | HORVU4Hr1G063420 | 3-ketoacyl-CoA synthase 1 | -6.57 | AT1G01120.1 |

| | | | | |
|---|---|---|---|---|
| 2378 | HORVU3Hr1G096650 | NAD(P)H dehydrogenase 18 | -6.58 | AT5G43750.1 |
| 2379 | HORVU3Hr1G010030 | Repressor of RNA polymerase III transcription | -6.58 | AT5G13240.1 |
| 2380 | HORVU4Hr1G006530 | beta glucosidase 43 | -6.59 | AT3G18080.1 |
| 2381 | HORVU5Hr1G009460 | Pentatricopeptide repeat (PPR) superfamily protein | -6.60 | AT5G24060.1 |
| 2382 | HORVU5Hr1G063380 | Pentatricopeptide repeat-containing protein | -6.60 | AT5G10690.2 |
| 2383 | HORVU3Hr1G046900 | UDP-N-acetylglucosamine--N-acetylmuramyl-(pentapeptide) pyrophosphoryl-undecaprenol N-acetylglucosamine transferase | -6.60 | AT1G73740.1 |
| 2384 | HORVU4Hr1G063270 | Filament-like plant protein 4 | -6.61 | AT1G19835.6 |
| 2385 | HORVU4Hr1G066860 | glutamine synthetase 1;4 | -6.62 | AT5G16570.1 |
| 2386 | HORVU2Hr1G039100 | Protein TIC 20-I, chloroplastic | -6.66 | AT1G04940.1 |
| 2387 | HORVU5Hr1G079950 | plastid movement impaired1 | -6.67 | AT1G42550.1 |
| 2388 | HORVU4Hr1G009120 | undescribed protein | -6.72 | AT4G37910.2 |
| 2389 | HORVU0Hr1G001170 | unknown function | -6.73 | AT3G51510.1 |
| 2390 | HORVU7Hr1G078490 | Casparian strip membrane protein 2 | -6.74 | AT2G27370.1 |
| 2391 | HORVU7Hr1G080270 | unknown function | -6.74 | AT5G08560.3 |
| 2392 | HORVU5Hr1G060250 | annexin 5 | -6.74 | AT1G68090.1 |
| 2393 | HORVU5Hr1G082310 | Protein kinase superfamily protein | -6.75 | AT5G47070.1 |
| 2394 | HORVU3Hr1G113920 | undescribed protein | -6.75 | AT5G04895.1 |
| 2395 | HORVU3Hr1G019750 | Auxin-responsive protein IAA2 | -6.76 | AT1G51950.1 |
| 2396 | HORVU3Hr1G047030 | Ferredoxin-thioredoxin reductase, catalytic chain | -6.76 | AT2G04700.3 |
| 2397 | HORVU5Hr1G062880 | Kelch repeat-containing protein | -6.77 | AT1G14330.1 |
| 2398 | HORVU6Hr1G075280 | ATP synthase delta-subunit gene | -6.80 | AT4G09650.1 |
| 2399 | HORVU5Hr1G056270 | undescribed protein | -6.81 | AT1G23050.1 |
| 2400 | HORVU4Hr1G004590 | Inositol hexakisphosphate and diphosphoinositol-pentakisphosphate kinase | -6.81 | AT5G15070.4 |
| 2401 | HORVU1Hr1G063010 | S-adenosyl-L-methionine-dependent methyltransferases superfamily protein | -6.81 | AT3G60910.1 |
| 2402 | HORVU2Hr1G065700 | unknown function | -6.82 | AT5G39380.5 |
| 2403 | HORVU7Hr1G098410 | Inositol polyphosphate multikinase | -6.82 | AT5G61760.2 |
| 2404 | HORVU2Hr1G017450 | unknown function | -6.84 | AT2G39050.1 |
| 2405 | HORVU3Hr1G030010 | GDT1-like protein 1, chloroplastic | -6.87 | AT1G64150.1 |
| 2406 | HORVU3Hr1G109830 | undescribed protein | -6.87 | AT2G16440.1 |
| 2407 | HORVU2Hr1G029900 | Protein kinase superfamily protein | -6.89 | AT1G78290.3 |
| 2408 | HORVU5Hr1G097060 | Protein phosphatase 2C family protein | -6.90 | AT4G33920.1 |
| 2409 | HORVU1Hr1G042490 | aspartate aminotransferase | -6.92 | AT1G77670.1 |
| 2410 | HORVU7Hr1G058030 | 3-beta hydroxysteroid dehydrogenase | -6.92 | AT4G35250.1 |
| 2411 | HORVU7Hr1G095100 | stress enhanced protein 1 | -6.92 | AT4G34190.1 |
| 2412 | HORVU7Hr1G118340 | Heavy metal transport/detoxification superfamily protein | -6.93 | AT1G23000.1 |
| 2413 | HORVU3Hr1G087180 | CCT motif family protein | -6.93 | AT5G41380.1 |

| | | | | |
|---|---|---|---|---|
| 2414 | HORVU4Hr1G086660 | Cyclin-dependent protein kinase inhibitor SMR1 | -6.94 | AT5G02420.1 |
| 2415 | HORVU2Hr1G007080 | Nuclear transport factor 2 (NTF2) family protein | -6.95 | AT1G71480.1 |
| 2416 | HORVU7Hr1G069710 | ACT domain-containing protein | -6.95 | AT1G16880.1 |
| 2417 | HORVU4Hr1G079230 | Aquaporin-like superfamily protein | -6.96 | AT2G36830.1 |
| 2418 | HORVU2Hr1G007840 | 60 kDa chaperonin 2 | -6.96 | AT2G28000.1 |
| 2419 | HORVU4Hr1G075160 | unknown function | -6.96 | AT1G14345.1 |
| 2420 | HORVU4Hr1G007260 | unknown function | -6.97 | AT1G73885.1 |
| 2421 | HORVU2Hr1G090980 | Starch synthase 2, chloroplastic/amyloplastic | -6.97 | AT1G32900.1 |
| 2422 | HORVU3Hr1G075960 | Jacalin-related lectin 3 | -6.98 | AT1G19715.2 |
| 2423 | HORVU7Hr1G096140 | Thylakoid lumenal 16.5 kDa protein, chloroplastic | -6.98 | AT4G02530.1 |
| 2424 | HORVU4Hr1G081310 | serine acetyltransferase 3;2 | -7.03 | AT2G17640.1 |
| 2425 | HORVU2Hr1G029490 | unknown protein; BEST Arabidopsis thaliana protein match is: unknown protein . | -7.07 | AT5G65030.1 |
| 2426 | HORVU0Hr1G014060 | unknown function | -7.08 | AT4G20880.1 |
| 2427 | HORVU6Hr1G032680 | FKBP-like peptidyl-prolyl cis-trans isomerase family protein | -7.12 | AT4G26555.1 |
| 2428 | HORVU0Hr1G039030 | nodulin MtN21 /EamA-like transporter family protein | -7.12 | AT1G25270.1 |
| 2429 | HORVU1Hr1G064780 | receptor-like protein kinase 2 | -7.14 | AT5G48380.1 |
| 2430 | HORVU6Hr1G064910 | Argonaute family protein | -7.15 | AT1G48410.3 |
| 2431 | HORVU5Hr1G067530 | 1-aminocyclopropane-1-carboxylate oxidase 1 | -7.16 | AT1G05010.1 |
| 2432 | HORVU1Hr1G026780 | Protein of unknown function (DUF581) | -7.16 | AT5G11460.1 |
| 2433 | HORVU3Hr1G080640 | Auxin efflux carrier family protein | -7.17 | AT5G01990.1 |
| 2434 | HORVU7Hr1G033500 | Peroxiredoxin Q, chloroplastic | -7.17 | AT4G16410.1 |
| 2435 | HORVU1Hr1G000720 | Protein of unknown function (DUF 3339) | -7.17 | AT5G08391.1 |
| 2436 | HORVU2Hr1G045620 | Protein LHCP TRANSLOCATION DEFECT | -7.18 | AT1G50900.1 |
| 2437 | HORVU7Hr1G114890 | UDP-Glycosyltransferase superfamily protein | -7.19 | AT4G01070.1 |
| 2438 | HORVU4Hr1G050730 | undescribed protein | -7.19 | AT5G15710.2 |
| 2439 | HORVU4Hr1G009980 | unknown function | -7.20 | AT1G44920.1 |
| 2440 | HORVU2Hr1G076010 | PsbP domain-containing protein 2, chloroplastic | -7.20 | AT2G28605.1 |
| 2441 | HORVU2Hr1G039510 | Cytochrome c | -7.21 | AT5G45040.1 |
| 2442 | HORVU2Hr1G101730 | ascorbate peroxidase 4 | -7.21 | AT4G09010.1 |
| 2443 | HORVU1Hr1G055140 | beta-amylase 1 | -7.23 | AT4G17090.1 |
| 2444 | HORVU4Hr1G050700 | Receptor-like protein kinase | -7.23 | AT3G51550.1 |
| 2445 | HORVU4Hr1G076990 | undescribed protein | -7.24 | AT5G26670.1 |
| 2446 | HORVU7Hr1G050520 | undescribed protein | -7.24 | AT3G10180.2 |
| 2447 | HORVU6Hr1G072050 | Ribulose-1,5-bisphosphate carboxylase/oxygenase small subunit N-methyltransferase I | -7.27 | AT3G07670.2 |
| 2448 | HORVU1Hr1G089760 | 50S ribosomal protein L7/L12 | -7.28 | AT3G27850.1 |

| | | | | |
|---|---|---|---|---|
| 2449 | HORVU2Hr1G094190 | Nuclear transport factor 2 (NTF2) family protein | -7.29 | AT2G46100.1 |
| 2450 | HORVU6Hr1G061780 | undescribed protein | -7.30 | AT4G18240.1 |
| 2451 | HORVU7Hr1G079190 | PSRP4 | -7.31 | AT2G38140.1 |
| 2452 | HORVU3Hr1G026500 | B-box zinc finger family protein | -7.32 | AT1G78600.1 |
| 2453 | HORVU2Hr1G104730 | #N/A | -7.33 | #N/A |
| 2454 | HORVU1Hr1G073790 | Zinc finger protein 282 | -7.34 | AT1G08290.1 |
| 2455 | HORVU4Hr1G046120 | undescribed protein | -7.35 | AT5G23010.3 |
| 2456 | HORVU3Hr1G048410 | undescribed protein | -7.36 | AT1G53580.2 |
| 2457 | HORVU3Hr1G071530 | 50S ribosomal protein L13 | -7.36 | AT1G78630.1 |
| 2458 | HORVU3Hr1G084830 | Nematode resistance protein-like HSPRO2 | -7.36 | AT3G55840.1 |
| 2459 | HORVU1Hr1G084090 | oligopeptide transporter 7 | -7.37 | AT4G10770.1 |
| 2460 | HORVU3Hr1G034640 | unknown function | -7.38 | AT4G01935.1 |
| 2461 | HORVU3Hr1G007290 | Calmodulin-binding protein | -7.39 | AT5G57580.1 |
| 2462 | HORVU2Hr1G115620 | Loricrin-like protein | -7.40 | AT5G64550.1 |
| 2463 | HORVU7Hr1G040720 | Endoglucanase 16 | -7.40 | AT2G32990.1 |
| 2464 | HORVU5Hr1G009430 | undescribed protein | -7.41 | AT5G44760.1 |
| 2465 | HORVU5Hr1G086710 | Flavin-containing monooxygenase family protein | -7.43 | AT1G19250.1 |
| 2466 | HORVU0Hr1G012920 | receptor-like kinase 1 | -7.44 | AT1G48480.1 |
| 2467 | HORVU4Hr1G080810 | Late embryogenesis abundant protein D-7 | -7.46 | AT1G52690.2 |
| 2468 | HORVU7Hr1G113020 | Low temperature and salt responsive protein family | -7.46 | AT4G28088.1 |
| 2469 | HORVU3Hr1G077570 | Sulfite exporter TauE/SafE family protein | -7.47 | AT4G21250.1 |
| 2470 | HORVU0Hr1G004850 | Peroxidase superfamily protein | -7.47 | AT4G17690.1 |
| 2471 | HORVU3Hr1G026870 | Protein kinase superfamily protein | -7.48 | AT5G58950.1 |
| 2472 | HORVU3Hr1G007420 | calmodulin like 23 | -7.49 | AT5G37770.2 |
| 2473 | HORVU7Hr1G083670 | Cytochrome P450 superfamily protein | -7.49 | AT5G06900.1 |
| 2474 | HORVU6Hr1G066340 | oxidative stress 3 | -7.50 | AT5G56550.1 |
| 2475 | HORVU7Hr1G041080 | SOUL heme-binding family protein | -7.50 | AT3G10130.1 |
| 2476 | HORVU4Hr1G062920 | Protein disulfide-isomerase LQY1, chloroplastic | -7.53 | AT1G75690.1 |
| 2477 | HORVU4Hr1G032780 | Peptide chain release factor 1 | -7.54 | AT3G62910.1 |
| 2478 | HORVU7Hr1G042480 | aluminum-activated malate transporter 9 | -7.54 | AT3G18440.1 |
| 2479 | HORVU3Hr1G010460 | Peptidase M50 family protein | -7.55 | AT5G05740.2 |
| 2480 | HORVU7Hr1G118020 | Eukaryotic aspartyl protease family protein | -7.55 | AT4G35880.1 |
| 2481 | HORVU3Hr1G093510 | Zinc finger CCCH domain-containing protein 12 | -7.56 | AT5G16540.4 |
| 2482 | HORVU3Hr1G105620 | Glucan endo-1,3-beta-glucosidase | -7.57 | AT4G16260.1 |
| 2483 | HORVU0Hr1G015440 | Ethylene responsive transcription factor 4 | -7.58 | #N/A |
| 2484 | HORVU1Hr1G015000 | unknown function | -7.58 | AT1G50732.1 |
| 2485 | HORVU2Hr1G092280 | NAD(P)H-quinone oxidoreductase subunit M | -7.59 | AT4G37925.1 |
| 2486 | HORVU3Hr1G093220 | undescribed protein | -7.62 | AT1G51630.1 |
| 2487 | HORVU7Hr1G064100 | proton gradient regulation 5 | -7.62 | AT2G05620.2 |

| | | | | |
|---|---|---|---|---|
| 2488 | HORVU6Hr1G021050 | unknown function | -7.64 | AT3G57062.2 |
| 2489 | HORVU5Hr1G110380 | purple acid phosphatase 10 | -7.64 | AT2G16430.2 |
| 2490 | HORVU3Hr1G066960 | Phospholipase A1-II 5 | -7.66 | AT2G42690.1 |
| 2491 | HORVU4Hr1G067930 | Ninja-family protein | -7.66 | AT1G69260.1 |
| 2492 | HORVU7Hr1G041380 | Caleosin-related family protein | -7.70 | AT1G70680.2 |
| 2493 | HORVU3Hr1G068840 | WD-40 repeat family protein | -7.72 | AT3G15354.5 |
| 2494 | HORVU5Hr1G115340 | 50S ribosomal protein L18 | -7.73 | AT1G48350.1 |
| 2495 | HORVU3Hr1G091150 | Protein of unknown function (DUF803) | -7.73 | AT1G71900.1 |
| 2496 | HORVU7Hr1G045640 | Epoxide hydrolase 3 | -7.73 | AT4G02340.1 |
| 2497 | HORVU7Hr1G104490 | glutamate receptor 3.4 | -7.74 | AT1G05200.6 |
| 2498 | HORVU1Hr1G090190 | WRKY family transcription factor | -7.76 | AT4G23810.1 |
| 2499 | HORVU2Hr1G097670 | Plastid-lipid associated protein | -7.76 | AT4G00030.1 |
| 2500 | HORVU7Hr1G049870 | alanine:glyoxylate aminotransferase | -7.78 | AT2G13360.3 |
| 2501 | HORVU6Hr1G052420 | Glycerophosphodiester phosphodiesterase GDPDL4 | -7.79 | AT5G55480.1 |
| 2502 | HORVU1Hr1G024070 | Protein kinase superfamily protein | -7.79 | AT3G46290.1 |
| 2503 | HORVU7Hr1G113890 | unknown function | -7.80 | AT4G31805.1 |
| 2504 | HORVU2Hr1G014710 | Methyltransferase domain protein | -7.82 | AT5G01710.1 |
| 2505 | HORVU2Hr1G080870 | vacuolar iron transporter 1 | -7.82 | AT2G01770.1 |
| 2506 | HORVU2Hr1G045970 | unknown function | -7.85 | AT3G21710.2 |
| 2507 | HORVU1Hr1G063220 | unknown function | -7.85 | AT1G03160.1 |
| 2508 | HORVU1Hr1G090130 | WRKY DNA-binding protein 30 | -7.86 | AT4G11070.2 |
| 2509 | HORVU6Hr1G012750 | undescribed protein | -7.86 | AT3G47460.1 |
| 2510 | HORVU6Hr1G078480 | undescribed protein | -7.88 | AT5G11360.2 |
| 2511 | HORVU2Hr1G086920 | Major facilitator superfamily protein | -7.90 | AT1G80530.1 |
| 2512 | HORVU4Hr1G002350 | N utilization substance protein B homolog | -7.90 | AT4G26370.1 |
| 2513 | HORVU7Hr1G083690 | Two-component response regulator-like APRR2 | -7.92 | AT2G20570.1 |
| 2514 | HORVU2Hr1G031570 | undescribed protein | -7.92 | AT3G23550.1 |
| 2515 | HORVU5Hr1G103590 | Syntaxin-121 | -7.92 | AT1G61290.2 |
| 2516 | HORVU2Hr1G072890 | Glucan endo-1,3-beta-glucosidase 14 | -7.93 | AT2G27500.2 |
| 2517 | HORVU1Hr1G079400 | undescribed protein | -7.93 | AT1G20110.1 |
| 2518 | HORVU6Hr1G059990 | unknown function | -7.94 | AT5G14895.1 |
| 2519 | HORVU5Hr1G093860 | Structural constituent of ribosome protein | -7.94 | AT4G19390.2 |
| 2520 | HORVU7Hr1G105950 | undescribed protein | -7.94 | #N/A |
| 2521 | HORVU3Hr1G037780 | PQ-loop repeat family protein / transmembrane family protein | -7.94 | AT2G41050.3 |
| 2522 | HORVU2Hr1G039070 | calcium-dependent protein kinase 19 | -7.95 | AT5G12480.1 |
| 2523 | HORVU4Hr1G067020 | Caleosin-related family protein | -7.95 | AT2G33380.1 |
| 2524 | HORVU7Hr1G109180 | VACUOLAR SORTING RECEPTOR 6 | -7.95 | AT1G30900.1 |
| 2525 | HORVU1Hr1G054930 | SBP (S-ribonuclease-binding protein) family protein | -7.96 | AT4G17680.1 |
| 2526 | HORVU2Hr1G022780 | Haloacid dehalogenase-like hydrolase domain-containing protein 3 | -8.00 | AT2G41250.1 |

| | | | | |
|---|---|---|---|---|
| 2527 | HORVU0Hr1G007660 | 0 | -8.02 | #N/A |
| 2528 | HORVU1Hr1G023160 | Flavin-containing monooxygenase family protein | -8.06 | AT1G19250.1 |
| 2529 | HORVU7Hr1G113830 | WRKY family transcription factor | -8.07 | AT1G80840.1 |
| 2530 | HORVU5Hr1G103990 | early nodulin-like protein 14 | -8.10 | AT2G26720.1 |
| 2531 | HORVU2Hr1G017080 | alpha/beta-Hydrolases superfamily protein | -8.10 | AT5G17780.2 |
| 2532 | HORVU6Hr1G029160 | Protein phosphatase 2C family protein | -8.10 | AT4G31860.1 |
| 2533 | HORVU5Hr1G063820 | Fatty acid hydroxylase superfamily | -8.12 | AT5G57800.1 |
| 2534 | HORVU2Hr1G037550 | tubulin alpha-4 chain | -8.12 | AT5G19780.1 |
| 2535 | HORVU3Hr1G089510 | formin homology 1 | -8.12 | AT3G25500.1 |
| 2536 | HORVU1Hr1G035870 | undescribed protein | -8.14 | AT4G15760.1 |
| 2537 | HORVU0Hr1G013900 | L-lactate dehydrogenase | -8.14 | AT3G14415.3 |
| 2538 | HORVU6Hr1G033290 | MLO-like protein 1 | -8.16 | AT4G02600.2 |
| 2539 | HORVU5Hr1G110000 | Cellulose synthase family protein | -8.18 | AT5G05170.1 |
| 2540 | HORVU6Hr1G009170 | NAD(P)-binding Rossmann-fold superfamily protein | -8.19 | AT1G72640.1 |
| 2541 | HORVU4Hr1G038570 | 30S ribosomal protein S5 | -8.19 | AT2G33800.1 |
| 2542 | HORVU3Hr1G077530 | Sulfite exporter TauE/SafE family protein | -8.20 | AT1G61740.1 |
| 2543 | HORVU4Hr1G024950 | Double Clp-N motif-containing P-loop nucleoside triphosphate hydrolases superfamily protein | -8.20 | AT2G29970.1 |
| 2544 | HORVU7Hr1G079720 | GDSL esterase/lipase | -8.22 | AT5G45910.1 |
| 2545 | HORVU7Hr1G114700 | unknown function | -8.24 | AT5G11070.1 |
| 2546 | HORVU5Hr1G087210 | Receptor-like serine/threonine-protein kinase SD1-8 | -8.24 | AT4G21390.2 |
| 2547 | HORVU2Hr1G104810 | undescribed protein | -8.28 | ATCG01050.1 |
| 2548 | HORVU4Hr1G004160 | Exostosin-2 | -8.29 | AT5G04500.1 |
| 2549 | HORVU7Hr1G082380 | Class I glutamine amidotransferase-like superfamily protein | -8.29 | AT4G34020.1 |
| 2550 | HORVU0Hr1G021630 | Auxin-responsive protein IAA21 | -8.29 | AT4G29080.1 |
| 2551 | HORVU6Hr1G077440 | Threonine endopeptidase | -8.31 | AT4G38100.1 |
| 2552 | HORVU6Hr1G015430 | alpha/beta-Hydrolases superfamily protein | -8.37 | AT5G17670.1 |
| 2553 | HORVU1Hr1G094050 | Protein kinase superfamily protein | -8.38 | AT3G59110.1 |
| 2554 | HORVU7Hr1G086120 | Leucine-rich repeat receptor-like protein kinase family protein | -8.39 | AT3G47570.1 |
| 2555 | HORVU6Hr1G053550 | beta-1,4-N-acetylglucosaminyltransferase family protein | -8.39 | AT5G14480.1 |
| 2556 | HORVU5Hr1G106790 | GTPase Obg | -8.39 | AT5G18570.1 |
| 2557 | HORVU7Hr1G121210 | At5g37260-like protein | -8.39 | AT5G37260.1 |
| 2558 | HORVU2Hr1G013870 | fatty acid desaturase 8 | -8.40 | AT5G05580.1 |
| 2559 | HORVU3Hr1G097330 | undescribed protein | -8.41 | AT5G19180.1 |
| 2560 | HORVU4Hr1G066230 | allene oxide synthase | -8.44 | AT5G42650.1 |
| 2561 | HORVU5Hr1G102530 | Hfr-2-like protein | -8.44 | AT5G44010.2 |
| 2562 | HORVU5Hr1G087970 | plastid movement impaired1 | -8.44 | AT1G42550.1 |

| | | | | |
|---|---|---|---|---|
| 2563 | HORVU3Hr1G095330 | 50S ribosomal protein L27 | -8.48 | AT5G40950.1 |
| 2564 | HORVU1Hr1G093390 | TPR repeat-containing thioredoxin TTL1 | -8.49 | AT1G53300.1 |
| 2565 | HORVU2Hr1G049560 | FKBP-like peptidyl-prolyl cis-trans isomerase family protein | -8.51 | AT1G20810.2 |
| 2566 | HORVU7Hr1G052920 | undescribed protein | -8.51 | AT2G17510.1 |
| 2567 | HORVU7Hr1G078020 | Pentatricopeptide repeat (PPR) superfamily protein | -8.52 | AT1G64430.2 |
| 2568 | HORVU3Hr1G078780 | Ribosomal protein L34 | -8.53 | AT1G29070.1 |
| 2569 | HORVU1Hr1G066530 | 22.3 kDa class VI heat shock protein | -8.56 | AT5G54660.1 |
| 2570 | HORVU3Hr1G029260 | Peptide-N4-(N-acetyl-beta-glucosaminyl)asparagine amidase A protein | -8.58 | AT3G14920.1 |
| 2571 | HORVU3Hr1G081180 | expansin A1 | -8.58 | AT1G69530.2 |
| 2572 | HORVU3Hr1G117590 | 23 kDa jasmonate-induced protein | -8.61 | AT1G53990.1 |
| 2573 | HORVU0Hr1G005320 | IQ-domain 22 | -8.63 | AT4G23060.1 |
| 2574 | HORVU6Hr1G053730 | Protein GLE1 | -8.66 | AT1G13120.1 |
| 2575 | HORVU7Hr1G114510 | BZIP transcription factor | -8.66 | AT4G36730.2 |
| 2576 | HORVU5Hr1G100870 | undescribed protein | -8.70 | AT5G64640.1 |
| 2577 | HORVU4Hr1G069990 | Diacylglycerol kinase | -8.70 | AT1G48390.3 |
| 2578 | HORVU4Hr1G077310 | Ethylene-responsive transcription factor 1 | -8.71 | AT1G72360.3 |
| 2579 | HORVU7Hr1G002200 | nodulin MtN21 /EamA-like transporter family protein | -8.72 | AT3G30340.1 |
| 2580 | HORVU0Hr1G029600 | undescribed protein | -8.73 | AT3G17240.3 |
| 2581 | HORVU5Hr1G098370 | Cytochrome c biogenesis protein CcsB | -8.73 | AT1G49380.1 |
| 2582 | HORVU7Hr1G085980 | Salt overly sensitive 1 | -8.74 | AT2G01980.1 |
| 2583 | HORVU4Hr1G011160 | Endoglucanase 10 | -8.74 | AT5G49720.1 |
| 2584 | HORVU0Hr1G009860 | PI-PLC X domain-containing protein | -8.75 | AT1G13680.1 |
| 2585 | HORVU5Hr1G092700 | Nuclear transcription factor Y subunit A-10 | -8.78 | AT5G06510.3 |
| 2586 | HORVU4Hr1G017030 | GDP-L-galactose phosphorylase 2 | -8.79 | AT5G55120.1 |
| 2587 | HORVU7Hr1G073860 | 2-succinylbenzoate--CoA ligase | -8.82 | AT1G30520.2 |
| 2588 | HORVU5Hr1G050510 | kinesin 4 | -8.82 | AT5G47820.2 |
| 2589 | HORVU1Hr1G071500 | undescribed protein | -8.83 | AT3G59100.1 |
| 2590 | HORVU4Hr1G035330 | Tryptophan synthase beta chain 2 | -8.84 | AT5G38530.1 |
| 2591 | HORVU7Hr1G042160 | Protein CURVATURE THYLAKOID 1A, chloroplastic | -8.84 | AT4G01150.1 |
| 2592 | HORVU2Hr1G106080 | WEB family protein At5g16730, chloroplastic | -8.84 | AT5G16730.1 |
| 2593 | HORVU4Hr1G023830 | FRAGILE HISTIDINE TRIAD | -8.85 | AT5G58240.2 |
| 2594 | HORVU4Hr1G021140 | Harpin-induced protein 1 containing protein, expressed | -8.85 | AT2G27260.1 |
| 2595 | HORVU4Hr1G016260 | ATP-dependent protease La (LON) domain protein | -8.85 | AT1G35340.1 |
| 2596 | HORVU4Hr1G074320 | ABC-type Co2+ transport system, permease component | -8.86 | AT4G19100.1 |
| 2597 | HORVU2Hr1G111590 | unknown function | -8.86 | AT3G22210.1 |
| 2598 | HORVU2Hr1G072880 | ferredoxin 3 | -8.89 | AT2G27510.1 |

| | | | | |
|---|---|---|---|---|
| 2599 | HORVU2Hr1G021700 | 4-alpha-glucanotransferase | -8.91 | AT2G40840.1 |
| 2600 | HORVU3Hr1G010310 | Disease resistance protein | -8.92 | AT3G07040.1 |
| 2601 | HORVU3Hr1G090270 | thylakoid rhodanese-like | -8.92 | AT3G25480.1 |
| 2602 | HORVU6Hr1G080750 | sulfite reductase | -8.94 | AT2G15620.1 |
| 2603 | HORVU6Hr1G077900 | Non-specific lipid transfer protein-like 1 | -8.97 | AT5G64080.2 |
| 2604 | HORVU3Hr1G067110 | Glycine dehydrogenase (decarboxylating) | -8.97 | AT2G26080.1 |
| 2605 | HORVU3Hr1G077660 | nuclear transport factor 2A | -8.97 | AT1G27310.1 |
| 2606 | HORVU6Hr1G034620 | Peroxiredoxin-2B | -8.98 | AT3G52960.1 |
| 2607 | HORVU5Hr1G034830 | WRKY family transcription factor | -8.98 | AT4G11070.2 |
| 2608 | HORVU4Hr1G075730 | unknown function | -9.00 | AT3G52105.1 |
| 2609 | HORVU4Hr1G056100 | Mitochondrial substrate carrier family protein | -9.02 | AT5G01340.1 |
| 2610 | HORVU3Hr1G016460 | undescribed protein | -9.04 | AT1G07990.2 |
| 2611 | HORVU3Hr1G065570 | undescribed protein | -9.06 | #N/A |
| 2612 | HORVU2Hr1G084750 | 2-phosphoglycolate phosphatase 1 | -9.07 | AT5G36790.3 |
| 2613 | HORVU2Hr1G078980 | Folic acid and derivative biosynthetic process | -9.11 | AT4G24380.4 |
| 2614 | HORVU7Hr1G050530 | basic helix-loop-helix (bHLH) DNA-binding superfamily protein | -9.11 | AT2G42280.1 |
| 2615 | HORVU7Hr1G117370 | Senescence/dehydration-associated protein-related | -9.13 | AT3G51250.1 |
| 2616 | HORVU3Hr1G107090 | calmodulin like 23 | -9.14 | AT1G24620.1 |
| 2617 | HORVU5Hr1G111330 | unknown function | -9.15 | AT4G13500.1 |
| 2618 | HORVU7Hr1G000120 | Protein of unknown function (DUF3754) | -9.24 | AT2G46915.1 |
| 2619 | HORVU2Hr1G001500 | undescribed protein | -9.24 | AT4G14500.2 |
| 2620 | HORVU0Hr1G016780 | Cytochrome P450 superfamily protein | -9.24 | AT5G45340.1 |
| 2621 | HORVU5Hr1G120650 | polyribonucleotide nucleotidyltransferase, putative | -9.25 | AT3G23700.1 |
| 2622 | HORVU3Hr1G044430 | undescribed protein | -9.26 | AT1G57770.1 |
| 2623 | HORVU2Hr1G025720 | C-repeat binding factor 3-like protein | -9.30 | AT4G25490.1 |
| 2624 | HORVU1Hr1G013970 | 3-oxo-5-alpha-steroid 4-dehydrogenase family protein | -9.33 | AT5G16010.1 |
| 2625 | HORVU2Hr1G016440 | Pentatricopeptide repeat-containing protein | -9.34 | AT1G13040.2 |
| 2626 | HORVU2Hr1G111570 | alpha/beta-Hydrolases superfamily protein | -9.36 | AT5G67050.1 |
| 2627 | HORVU6Hr1G028220 | Disease resistance protein | -9.36 | AT3G07040.1 |
| 2628 | HORVU2Hr1G063740 | alanine aminotransferase 2 | -9.38 | AT1G70580.4 |
| 2629 | HORVU2Hr1G072530 | Calcineurin-like metallo-phosphoesterase superfamily protein | -9.39 | AT3G09970.1 |
| 2630 | HORVU7Hr1G109280 | Fructose-1,6-bisphosphatase class 1 | -9.41 | AT5G64380.1 |
| 2631 | HORVU3Hr1G093230 | undescribed protein | -9.43 | AT5G16750.1 |
| 2632 | HORVU3Hr1G017000 | Tetratricopeptide repeat 6 | -9.44 | AT1G62340.1 |
| 2633 | HORVU6Hr1G075520 | undescribed protein | -9.44 | AT3G12940.1 |
| 2634 | HORVU5Hr1G114990 | GDP-mannose transporter | -9.44 | AT2G13650.3 |
| 2635 | HORVU5Hr1G115460 | ATP-dependent RNA helicase DeaD | -9.44 | AT5G26742.2 |
| 2636 | HORVU4Hr1G068870 | Pentatricopeptide repeat-containing protein | -9.44 | AT3G29290.5 |

| | | | | |
|---|---|---|---|---|
| 2637 | HORVU2Hr1G077970 | 3-oxoacyl-[acyl-carrier-protein] synthase 2 | -9.45 | AT5G46290.1 |
| 2638 | HORVU5Hr1G031210 | undescribed protein | -9.50 | AT3G47600.1 |
| 2639 | HORVU4Hr1G083230 | S-adenosyl-L-methionine-dependent methyltransferases superfamily protein | -9.52 | AT5G44600.1 |
| 2640 | HORVU7Hr1G095160 | KAR-UP F-box 1 | -9.53 | AT1G31350.1 |
| 2641 | HORVU2Hr1G093210 | cyclin p4;1 | -9.53 | AT2G44740.1 |
| 2642 | HORVU3Hr1G019820 | 2-oxoglutarate (2OG) and Fe(II)-dependent oxygenase superfamily protein | -9.57 | AT1G35190.1 |
| 2643 | HORVU6Hr1G058780 | Protein kinase superfamily protein | -9.58 | AT1G08650.1 |
| 2644 | HORVU5Hr1G104230 | Glucan endo-1,3-beta-glucosidase 8 | -9.62 | AT2G19440.2 |
| 2645 | HORVU3Hr1G050800 | Chromosome 3B, genomic scaffold, cultivar Chinese Spring | -9.67 | AT1G74730.1 |
| 2646 | HORVU6Hr1G003300 | nitrate reductase 1 | -9.68 | AT1G77760.1 |
| 2647 | HORVU6Hr1G005050 | flowering promoting factor 1 | -9.72 | AT5G24860.1 |
| 2648 | HORVU2Hr1G007070 | Cyclopropane-fatty-acyl-phospholipid synthase | -9.73 | AT3G23510.1 |
| 2649 | HORVU3Hr1G028410 | RING/U-box superfamily protein | -9.75 | AT2G42350.1 |
| 2650 | HORVU6Hr1G031870 | Retrotransposon protein, putative, Ty1-copia subclass | -9.75 | AT3G16785.3 |
| 2651 | HORVU5Hr1G101020 | ribose-5-phosphate isomerase 2 | -9.76 | AT5G44520.2 |
| 2652 | HORVU0Hr1G014040 | PI-PLC X domain-containing protein | -9.77 | AT1G13680.1 |
| 2653 | HORVU5Hr1G095540 | Outward rectifying potassium channel protein | -9.80 | AT5G55630.2 |
| 2654 | HORVU4Hr1G076180 | unknown function | -9.82 | AT2G26340.1 |
| 2655 | HORVU2Hr1G091360 | receptor kinase 2 | -9.83 | AT2G37710.1 |
| 2656 | HORVU7Hr1G050540 | DNA binding protein | -9.83 | AT1G51140.1 |
| 2657 | HORVU7Hr1G055890 | 50S ribosomal protein 6, chloroplastic | -9.85 | AT5G17870.1 |
| 2658 | HORVU4Hr1G021820 | DUF3049 family protein | -9.85 | AT5G22390.1 |
| 2659 | HORVU5Hr1G115000 | Thylakoid soluble phosphoprotein TSP9 protein | -9.85 | AT3G47070.1 |
| 2660 | HORVU4Hr1G065660 | MATE efflux family protein | -9.86 | AT1G58340.1 |
| 2661 | HORVU4Hr1G015250 | Zinc-finger homeodomain protein 4 | -9.87 | AT1G14440.2 |
| 2662 | HORVU5Hr1G082460 | Endoglucanase 24 | -9.89 | AT1G75680.1 |
| 2663 | HORVU3Hr1G095300 | stress enhanced protein 1 | -9.91 | AT4G34190.1 |
| 2664 | HORVU3Hr1G089190 | Pentatricopeptide repeat (PPR) superfamily protein | -9.97 | AT3G59300.1 |
| 2665 | HORVU5Hr1G093640 | Auxin-responsive protein IAA12 | -10.01 | AT1G04240.1 |
| 2666 | HORVU6Hr1G067620 | Pentatricopeptide repeat-containing protein | -10.01 | AT1G31920.1 |
| 2667 | HORVU2Hr1G125280 | unknown function | -10.02 | AT1G45688.1 |
| 2668 | HORVU3Hr1G088020 | dihydroflavonol 4-reductase-like1 | -10.03 | AT5G58490.1 |
| 2669 | HORVU2Hr1G118050 | undescribed protein | -10.05 | AT1G01490.2 |
| 2670 | HORVU1Hr1G038750 | Double-stranded RNA-binding protein 5 | -10.07 | AT5G37180.3 |
| 2671 | HORVU3Hr1G055940 | Kinesin-related protein 6 | -10.10 | AT3G16060.1 |
| 2672 | HORVU2Hr1G007000 | Cyclopropane-fatty-acyl-phospholipid synthase | -10.15 | AT3G23510.1 |
| 2673 | HORVU6Hr1G068690 | Cytochrome P450 superfamily protein | -10.25 | AT5G45340.1 |
| 2674 | HORVU7Hr1G015160 | unknown function | -10.25 | AT1G32928.1 |

| | | | | |
|---|---|---|---|---|
| 2675 | HORVU6Hr1G068910 | alpha/beta hydrolase | -10.25 | AT4G36530.2 |
| 2676 | HORVU2Hr1G115710 | unknown protein; LOCATED IN: chloroplast; EXPRESSED IN: 22 plant structures; EXPRESSED DURING: 13 growth stages; BEST Arabidopsis thaliana protein match is: unknown protein . | -10.26 | AT2G21960.1 |
| 2677 | HORVU7Hr1G056240 | Peptidyl-prolyl cis-trans isomerase | -10.26 | AT2G43560.1 |
| 2678 | HORVU5Hr1G093850 | Phototropic-responsive NPH3 family protein | -10.28 | AT5G66560.1 |
| 2679 | HORVU3Hr1G082280 | Glutamine--fructose-6-phosphate aminotransferase [isomerizing] | -10.28 | AT2G16570.1 |
| 2680 | HORVU1Hr1G076350 | trehalose-6-phosphate synthase | -10.28 | AT1G78580.4 |
| 2681 | HORVU5Hr1G016270 | Sulfurtransferase | -10.32 | AT1G27320.1 |
| 2682 | HORVU6Hr1G018830 | 50S ribosomal protein L3 | -10.34 | AT2G43030.1 |
| 2683 | HORVU4Hr1G019570 | fructose-bisphosphate aldolase 2 | -10.35 | AT4G38970.1 |
| 2684 | HORVU6Hr1G010050 | Protein kinase superfamily protein | -10.37 | AT2G17220.1 |
| 2685 | HORVU2Hr1G036110 | NADP-dependent glyceraldehyde-3-phosphate dehydrogenase | -10.43 | AT2G24270.4 |
| 2686 | HORVU6Hr1G000650 | Glyoxylate/hydroxypyruvate reductase B | -10.45 | AT1G68010.1 |
| 2687 | HORVU7Hr1G081590 | Protein kinase superfamily protein | -10.46 | AT3G46290.1 |
| 2688 | HORVU2Hr1G108070 | Glycosyltransferase family 61 protein | -10.47 | AT3G18170.1 |
| 2689 | HORVU5Hr1G109680 | Protein of Unknown Function (DUF239) | -10.47 | AT1G55360.1 |
| 2690 | HORVU7Hr1G092550 | Transmembrane amino acid transporter family protein | -10.47 | AT3G30390.3 |
| 2691 | HORVU4Hr1G060440 | Digalactosyldiacylglycerol synthase 2, chloroplastic | -10.47 | AT4G00550.2 |
| 2692 | HORVU6Hr1G082710 | UDP-Glycosyltransferase superfamily protein | -10.47 | AT1G22360.1 |
| 2693 | HORVU2Hr1G099890 | Ethylene-responsive transcription factor 4 | -10.49 | AT3G15210.1 |
| 2694 | HORVU5Hr1G053230 | RNA-binding protein 1 | -10.49 | AT3G52150.2 |
| 2695 | HORVU7Hr1G043030 | HD1 | -10.50 | AT3G02380.1 |
| 2696 | HORVU4Hr1G015330 | Rieske (2Fe-2S) domain-containing protein | -10.51 | AT1G71500.1 |
| 2697 | HORVU2Hr1G077460 | cytochrome P450, family 87, subfamily A, polypeptide 6 | -10.52 | AT1G64940.1 |
| 2698 | HORVU2Hr1G092180 | Elongation factor G | -10.52 | AT1G62750.1 |
| 2699 | HORVU2Hr1G022920 | glutamate synthase 1 | -10.53 | AT5G04140.1 |
| 2700 | HORVU0Hr1G017160 | Glutaredoxin family protein | -10.54 | AT5G39865.1 |
| 2701 | HORVU5Hr1G048300 | Double Clp-N motif-containing P-loop nucleoside triphosphate hydrolases superfamily protein | -10.55 | AT2G29970.1 |
| 2702 | HORVU1Hr1G024400 | 6-phosphofructo-2-kinase/fructose-2,6-bisphosphatase | -10.58 | AT1G07110.1 |
| 2703 | HORVU5Hr1G042160 | purple acid phosphatase 22 | -10.58 | AT3G52820.1 |
| 2704 | HORVU3Hr1G039790 | 50S ribosomal protein L40 | -10.58 | AT3G56910.1 |
| 2705 | HORVU2Hr1G111540 | Farnesyl pyrophosphate synthase 2 | -10.60 | AT5G47770.1 |
| 2706 | HORVU1Hr1G073300 | myb-like transcription factor family protein | -10.64 | AT2G38090.1 |
| 2707 | HORVU6Hr1G022680 | Late embryogenesis abundant (LEA) hydroxyproline-rich glycoprotein family | -10.64 | AT5G06320.1 |

| | | | | |
|---|---|---|---|---|
| 2708 | HORVU2Hr1G010830 | alcohol dehydrogenase 1 | -10.66 | AT4G39330.1 |
| 2709 | HORVU3Hr1G091140 | NAD(P)H-quinone oxidoreductase subunit N | -10.66 | AT5G58260.1 |
| 2710 | HORVU4Hr1G042560 | undescribed protein | -10.66 | AT1G32960.1 |
| 2711 | HORVU3Hr1G019830 | undescribed protein | -10.68 | AT3G49180.1 |
| 2712 | HORVU1Hr1G064970 | amino acid permease 2 | -10.69 | AT5G09220.1 |
| 2713 | HORVU3Hr1G001080 | Chromosome 3B, genomic scaffold, cultivar Chinese Spring | -10.71 | AT2G35350.1 |
| 2714 | HORVU2Hr1G015470 | Protein phosphatase 2C family protein | -10.74 | AT1G09160.2 |
| 2715 | HORVU7Hr1G115040 | 30S ribosomal protein S17 | -10.75 | AT1G79850.1 |
| 2716 | HORVU6Hr1G009500 | Peroxidase superfamily protein | -10.75 | AT5G05340.1 |
| 2717 | HORVU4Hr1G051820 | Retrotransposon protein, putative, Ty1-copia subclass | -10.75 | AT4G23160.3 |
| 2718 | HORVU2Hr1G071100 | Myb/SANT-like DNA-binding domain protein | -10.76 | AT2G33550.1 |
| 2719 | HORVU1Hr1G068910 | unknown function | -10.77 | AT3G47510.1 |
| 2720 | HORVU4Hr1G062330 | unknown function | -10.77 | AT4G11000.1 |
| 2721 | HORVU1Hr1G025440 | Embryo-specific protein 3, (ATS3) | -10.80 | AT2G41475.1 |
| 2722 | HORVU4Hr1G056080 | 2-oxoglutarate (2OG) and Fe(II)-dependent oxygenase superfamily protein | -10.80 | AT1G06640.1 |
| 2723 | HORVU7Hr1G098510 | Peroxidase superfamily protein | -10.82 | AT2G18980.1 |
| 2724 | HORVU3Hr1G082730 | Photosystem II 22 kDa protein, chloroplastic | -10.82 | AT1G44575.3 |
| 2725 | HORVU5Hr1G001730 | Leucine-rich receptor-like protein kinase family protein | -10.85 | AT5G49660.1 |
| 2726 | HORVU2Hr1G066020 | undescribed protein | -10.87 | AT3G55380.2 |
| 2727 | HORVU6Hr1G000620 | Ferredoxin--NADP reductase | -10.88 | AT5G66190.1 |
| 2728 | HORVU3Hr1G010370 | DNA damage-binding protein 2 | -10.92 | AT5G58760.1 |
| 2729 | HORVU2Hr1G061010 | undescribed protein | -10.93 | AT5G07380.4 |
| 2730 | HORVU7Hr1G112280 | undescribed protein | -10.94 | #N/A |
| 2731 | HORVU6Hr1G031370 | Calcineurin-like metallo-phosphoesterase superfamily protein | -11.03 | AT1G07010.1 |
| 2732 | HORVU5Hr1G022750 | PsbP domain-containing protein 1, chloroplastic | -11.03 | AT4G15510.5 |
| 2733 | HORVU5Hr1G018220 | Two-component response regulator ARR18 | -11.03 | AT2G03500.1 |
| 2734 | HORVU1Hr1G061970 | 'putative endo-1,3;1,4-beta-D-glucanase' | -11.03 | AT3G23600.1 |
| 2735 | HORVU6Hr1G090240 | undescribed protein | -11.05 | #N/A |
| 2736 | HORVU3Hr1G022540 | Auxin-responsive protein IAA1 | -11.06 | AT1G04250.1 |
| 2737 | HORVU5Hr1G072980 | serine carboxypeptidase-like 18 | -11.07 | AT1G33540.1 |
| 2738 | HORVU5Hr1G050990 | ribulose bisphosphate carboxylase small chain 1A | -11.20 | AT1G67090.1 |
| 2739 | HORVU3Hr1G115610 | Vacuolar-processing enzyme | -11.25 | AT4G32940.1 |
| 2740 | HORVU2Hr1G076880 | Protein NRT1/ PTR FAMILY 4.3 | -11.31 | AT1G59740.1 |
| 2741 | HORVU3Hr1G082740 | Photosystem II 22 kDa protein, chloroplastic | -11.35 | AT1G44575.3 |
| 2742 | HORVU2Hr1G039000 | unknown function | -11.42 | AT2G01870.1 |
| 2743 | HORVU7Hr1G084420 | RAS-related GTP-binding nuclear protein 2 | -11.45 | AT5G55190.1 |

| | | | | |
|---|---|---|---|---|
| 2744 | HORVU2Hr1G010630 | ribulose bisphosphate carboxylase small chain 1A | -11.49 | AT1G67090.1 |
| 2745 | HORVU0Hr1G022410 | undescribed protein | -11.52 | AT1G05860.3 |
| 2746 | HORVU4Hr1G011500 | serine hydroxymethyltransferase 2 | -11.53 | AT4G37930.1 |
| 2747 | HORVU7Hr1G078690 | cytochrome B561-1 | -11.57 | AT4G25570.1 |
| 2748 | HORVU2Hr1G089970 | Aquaporin-like superfamily protein | -11.58 | AT2G16850.1 |
| 2749 | HORVU6Hr1G066050 | RING/U-box superfamily protein | -11.59 | AT3G61460.1 |
| 2750 | HORVU1Hr1G075520 | Jacalin-related lectin 3 | -11.61 | AT1G19715.1 |
| 2751 | HORVU1Hr1G049420 | Heavy metal transport/detoxification superfamily protein | -11.62 | AT1G01490.2 |
| 2752 | HORVU6Hr1G086210 | RING-H2 zinc finger protein-like | -11.62 | AT2G15580.1 |
| 2753 | HORVU6Hr1G073100 | Protein CURVATURE THYLAKOID 1A, chloroplastic | -11.62 | AT4G01150.1 |
| 2754 | HORVU4Hr1G075400 | undescribed protein | -11.63 | AT5G02400.1 |
| 2755 | HORVU1Hr1G001900 | undescribed protein | -11.66 | AT1G21640.1 |
| 2756 | HORVU4Hr1G048560 | Tubby-like F-box protein 6 | -11.72 | AT2G47900.3 |
| 2757 | HORVU4Hr1G027260 | Ribulose bisphosphate carboxylase/oxygenase activase A, chloroplastic | -11.73 | AT2G39730.1 |
| 2758 | HORVU6Hr1G063310 | trehalose-6-phosphate phosphatase | -11.75 | AT5G51460.4 |
| 2759 | HORVU5Hr1G018790 | Cytochrome P450 superfamily protein | -11.76 | AT1G66540.2 |
| 2760 | HORVU5Hr1G074000 | NAD(P)-binding Rossmann-fold superfamily protein | -11.77 | AT3G18890.1 |
| 2761 | HORVU1Hr1G065810 | Secretory carrier-associated membrane protein 3 | -11.79 | AT1G61250.1 |
| 2762 | HORVU7Hr1G114710 | unknown function | -11.82 | AT5G11070.1 |
| 2763 | HORVU6Hr1G075900 | Geranylgeranyl diphosphate reductase | -11.82 | AT1G74470.1 |
| 2764 | HORVU3Hr1G024970 | GDSL esterase/lipase | -11.84 | AT1G28650.1 |
| 2765 | HORVU6Hr1G072540 | signal peptide peptidase | -11.85 | AT1G73990.2 |
| 2766 | HORVU3Hr1G117770 | Zinc finger CCCH domain-containing protein 33 | -11.85 | AT5G58620.1 |
| 2767 | HORVU3Hr1G036600 | TRAM, LAG1 and CLN8 (TLC) lipid-sensing domain containing protein | -11.86 | AT1G35180.1 |
| 2768 | HORVU4Hr1G002050 | 2-methyl-6-phytyl-1,4-hydroquinone methyltransferase 2, chloroplastic | -11.89 | AT3G63410.1 |
| 2769 | HORVU1Hr1G026650 | Leucine-rich repeat receptor-like protein kinase family protein | -11.92 | AT3G24480.1 |
| 2770 | HORVU1Hr1G007810 | UDP-Glycosyltransferase superfamily protein | -12.00 | AT1G22360.1 |
| 2771 | HORVU1Hr1G005120 | outer plastid envelope protein 16-1 | -12.02 | AT2G28900.1 |
| 2772 | HORVU6Hr1G088610 | terpene synthase 14 | -12.05 | AT1G61680.1 |
| 2773 | HORVU0Hr1G023540 | trehalose-6-phosphate phosphatase | -12.06 | AT5G51460.4 |
| 2774 | HORVU6Hr1G055660 | Dehydrogenase/reductase SDR family member 7B | -12.08 | AT3G59710.3 |
| 2775 | HORVU4Hr1G009440 | basic helix-loop-helix (bHLH) DNA-binding superfamily protein | -12.12 | AT4G37850.2 |
| 2776 | HORVU5Hr1G020500 | Protein of unknown function (DUF1118) | -12.15 | AT5G08050.1 |
| 2777 | HORVU2Hr1G060010 | L-lactate dehydrogenase | -12.15 | AT3G14420.2 |
| 2778 | HORVU2Hr1G108090 | undescribed protein | -12.18 | AT5G45100.2 |

| | | | | |
|---|---|---|---|---|
| 2779 | HORVU6Hr1G067120 | unknown protein; Has 149 Blast hits to 140 proteins in 44 species: Archae - 0; Bacteria - 6; Metazoa - 40; Fungi - 6; Plants - 39; Viruses - 0; Other Eukaryotes - 58 (source: NCBI BLink). | -12.28 | AT2G35200.1 |
| 2780 | HORVU4Hr1G002900 | Protein of unknown function (DUF1677) | -12.32 | AT3G22540.1 |
| 2781 | HORVU2Hr1G084420 | BnaA09g28020D protein | -12.35 | AT1G27300.1 |
| 2782 | HORVU7Hr1G071000 | undescribed protein | -12.36 | #N/A |
| 2783 | HORVU5Hr1G111640 | 50S ribosomal protein L17 | -12.42 | AT3G54210.1 |
| 2784 | HORVU0Hr1G020750 | beta glucosidase 17 | -12.42 | AT2G44480.1 |
| 2785 | HORVU1Hr1G013760 | undescribed protein | -12.45 | AT3G08530.1 |
| 2786 | HORVU7Hr1G077750 | ferredoxin 1 | -12.46 | AT1G60950.1 |
| 2787 | HORVU7Hr1G120660 | PHYTOENE SYNTHASE | -12.49 | AT5G17230.4 |
| 2788 | HORVU5Hr1G022570 | Stress responsive alpha-beta barrel domain protein | -12.54 | AT2G32500.2 |
| 2789 | HORVU7Hr1G112290 | Chromosome 3B, genomic scaffold, cultivar Chinese Spring | -12.56 | AT2G24710.3 |
| 2790 | HORVU7Hr1G112240 | undescribed protein | -12.60 | AT5G40250.1 |
| 2791 | HORVU7Hr1G044780 | undescribed protein | -12.64 | AT1G21450.1 |
| 2792 | HORVU1Hr1G060610 | 1-deoxy-D-xylulose 5-phosphate synthase 1 | -12.64 | AT4G15560.1 |
| 2793 | HORVU4Hr1G004800 | Tetratricopeptide repeat (TPR)-like superfamily protein | -12.65 | AT1G27150.1 |
| 2794 | HORVU5Hr1G025610 | Glutaredoxin family protein | -12.66 | AT2G47880.1 |
| 2795 | HORVU7Hr1G047210 | Phosphate transporter PHO1-3 | -12.69 | AT1G68740.1 |
| 2796 | HORVU6Hr1G080420 | sugar transporter 2 | -12.70 | AT2G48020.3 |
| 2797 | HORVU7Hr1G057100 | Cytochrome P450 superfamily protein | -12.73 | AT3G61880.2 |
| 2798 | HORVU2Hr1G085300 | haloacid dehalogenase-like hydrolase family protein | -12.73 | AT4G39970.1 |
| 2799 | HORVU2Hr1G071940 | Photosystem II stability/assembly factor HCF136 | -12.74 | AT5G23120.1 |
| 2800 | HORVU7Hr1G038300 | Transmembrane amino acid transporter family protein | -12.91 | AT3G28960.5 |
| 2801 | HORVU4Hr1G085960 | Peroxisomal membrane protein 11-1 | -12.92 | AT1G01820.1 |
| 2802 | HORVU4Hr1G057020 | Low temperature and salt responsive protein family | -12.92 | AT2G38905.1 |
| 2803 | HORVU3Hr1G056830 | glycerol-3-phosphate acyltransferase 1 | -13.02 | AT1G06520.1 |
| 2804 | HORVU7Hr1G036720 | germin-like protein 1 | -13.05 | AT1G72610.1 |
| 2805 | HORVU4Hr1G025010 | undescribed protein | -13.13 | #N/A |
| 2806 | HORVU7Hr1G121250 | Aquaporin-like superfamily protein | -13.14 | AT4G18910.1 |
| 2807 | HORVU2Hr1G098820 | Trypsin family protein | -13.20 | AT5G45030.2 |
| 2808 | HORVU3Hr1G078860 | UDP-Glycosyltransferase superfamily protein | -13.25 | AT2G30140.1 |
| 2809 | HORVU3Hr1G018050 | undescribed protein | -13.25 | AT4G28706.4 |
| 2810 | HORVU6Hr1G002550 | Nitric oxide-associated protein 1 | -13.27 | AT3G47450.2 |
| 2811 | HORVU1Hr1G022490 | unknown function | -13.27 | AT5G60360.3 |
| 2812 | HORVU4Hr1G008160 | PHD finger protein | -13.29 | AT1G66170.1 |
| 2813 | HORVU7Hr1G119100 | Serine/threonine-protein kinase | -13.30 | AT1G34300.1 |

| | | | | |
|---|---|---|---|---|
| 2814 | HORVU5Hr1G024410 | rapid alkalinization factor 23 | -13.34 | AT4G15800.1 |
| 2815 | HORVU5Hr1G025820 | Retrotransposon protein, putative, unclassified | -13.36 | AT3G62300.1 |
| 2816 | HORVU3Hr1G040880 | F-box family protein | -13.38 | AT2G32560.1 |
| 2817 | HORVU2Hr1G074770 | Abscisic stress-ripening protein 3 | -13.39 | AT2G03140.9 |
| 2818 | HORVU5Hr1G070140 | undescribed protein | -13.40 | AT3G61180.1 |
| 2819 | HORVU7Hr1G082350 | UDP-Glycosyltransferase superfamily protein | -13.41 | AT3G16520.3 |
| 2820 | HORVU1Hr1G047100 | Aquaporin-like superfamily protein | -13.42 | AT4G10380.1 |
| 2821 | HORVU2Hr1G038570 | Ribosome-recycling factor | -13.42 | AT3G63190.1 |
| 2822 | HORVU6Hr1G045650 | undescribed protein | -13.45 | AT2G41720.1 |
| 2823 | HORVU3Hr1G024980 | undescribed protein | -13.45 | AT3G28223.1 |
| 2824 | HORVU5Hr1G051010 | ribulose bisphosphate carboxylase small chain 1A | -13.51 | AT1G67090.1 |
| 2825 | HORVU1Hr1G069830 | expansin A1 | -13.52 | AT1G69530.2 |
| 2826 | HORVU7Hr1G088920 | Heat stress transcription factor C-2b | -13.58 | AT3G24520.1 |
| 2827 | HORVU3Hr1G058230 | undescribed protein | -13.59 | AT4G21200.3 |
| 2828 | HORVU2Hr1G096270 | calcium-dependent protein kinase 29 | -13.61 | AT1G76040.2 |
| 2829 | HORVU7Hr1G091560 | C-repeat binding factor 3-like protein | -13.62 | AT4G25470.1 |
| 2830 | HORVU5Hr1G077140 | Esterase/lipase/thioesterase family protein | -13.66 | AT5G41130.1 |
| 2831 | HORVU6Hr1G093030 | 50S ribosomal protein L9 | -13.71 | AT3G44890.1 |
| 2832 | HORVU7Hr1G050500 | unknown protein; INVOLVED IN: biological_process unknown; LOCATED IN: chloroplast; EXPRESSED IN: 21 plant structures; EXPRESSED DURING: 12 growth stages; BEST Arabidopsis thaliana protein match is: unknown protein . | -13.74 | AT5G12900.1 |
| 2833 | HORVU3Hr1G057400 | BnaA07g15680D protein | -13.76 | AT5G42070.1 |
| 2834 | HORVU3Hr1G019320 | unknown protein; BEST Arabidopsis thaliana protein match is: unknown protein . | -13.77 | AT3G11760.1 |
| 2835 | HORVU2Hr1G111530 | alpha/beta-Hydrolases superfamily protein | -13.83 | AT3G14360.1 |
| 2836 | HORVU4Hr1G013190 | Protein of unknown function (DUF620) | -13.84 | AT3G19540.1 |
| 2837 | HORVU1Hr1G076910 | serine/threonine protein kinase 1 | -13.87 | AT4G18700.1 |
| 2838 | HORVU4Hr1G072060 | Heavy metal transport/detoxification superfamily protein | -13.87 | AT5G03380.1 |
| 2839 | HORVU7Hr1G089850 | Plant protein of unknown function (DUF247) | -13.93 | AT3G50120.1 |
| 2840 | HORVU6Hr1G077610 | RING/U-box superfamily protein | -13.95 | AT3G05200.1 |
| 2841 | HORVU2Hr1G009980 | Eukaryotic aspartyl protease family protein | -14.07 | AT2G03200.1 |
| 2842 | HORVU2Hr1G089960 | undescribed protein | -14.08 | AT5G63950.2 |
| 2843 | HORVU7Hr1G065970 | undescribed protein | -14.11 | AT3G48430.1 |
| 2844 | HORVU1Hr1G070290 | proline-rich family protein | -14.13 | AT5G07020.1 |
| 2845 | HORVU1Hr1G052790 | undescribed protein | -14.13 | AT1G78000.7 |
| 2846 | HORVU7Hr1G039140 | Glutathione S-transferase family protein | -14.18 | AT3G62760.1 |
| 2847 | HORVU7Hr1G009800 | LEA | -14.19 | AT1G32560.1 |
| 2848 | HORVU7Hr1G077740 | ferredoxin 1 | -14.24 | AT1G60950.1 |
| 2849 | HORVU2Hr1G076970 | Pentatricopeptide repeat-containing protein | -14.26 | AT5G08490.8 |

| | | | | |
|---|---|---|---|---|
| 2850 | HORVU2Hr1G096850 | Haloacid dehalogenase-like hydrolase (HAD) superfamily protein | -14.29 | AT4G11570.2 |
| 2851 | HORVU3Hr1G085270 | Fructose-1,6-bisphosphatase class 1 | -14.32 | AT3G55800.1 |
| 2852 | HORVU7Hr1G038530 | Pentatricopeptide repeat-containing protein | -14.37 | AT5G06540.1 |
| 2853 | HORVU6Hr1G010940 | undescribed protein | -14.43 | AT4G16120.1 |
| 2854 | HORVU4Hr1G032020 | undescribed protein | -14.53 | AT5G62630.1 |
| 2855 | HORVU0Hr1G031970 | Protein DEHYDRATION-INDUCED 19 homolog 5 | -14.57 | AT3G05700.1 |
| 2856 | HORVU6Hr1G010900 | undescribed protein | -14.64 | AT2G39330.2 |
| 2857 | HORVU2Hr1G094280 | Protein FATTY ACID EXPORT 4, chloroplastic | -14.73 | AT1G33265.1 |
| 2858 | HORVU2Hr1G068610 | violaxanthin de-epoxidase-related | -14.75 | AT1G08550.3 |
| 2859 | HORVU1Hr1G017310 | undescribed protein | -14.82 | AT5G17980.1 |
| 2860 | HORVU7Hr1G058120 | Chlorophyll a-b binding protein 1B, chloroplastic | -14.85 | AT2G34430.1 |
| 2861 | HORVU0Hr1G017560 | Cytochrome P450 superfamily protein | -14.96 | AT1G31800.1 |
| 2862 | HORVU4Hr1G014010 | NAD(P)H dehydrogenase subunit 48 | -14.98 | AT1G15980.1 |
| 2863 | HORVU0Hr1G021130 | undescribed protein | -15.05 | AT4G29900.2 |
| 2864 | HORVU7Hr1G076790 | Core-2/I-branching beta-1,6-N-acetylglucosaminyltransferase family protein | -15.06 | AT1G10280.1 |
| 2865 | HORVU1Hr1G084390 | GDSL esterase/lipase | -15.09 | AT5G45910.1 |
| 2866 | HORVU0Hr1G025440 | unknown protein; BEST Arabidopsis thaliana protein match is: unknown protein . | -15.09 | AT3G15630.1 |
| 2867 | HORVU4Hr1G087350 | UPF0187 protein YneE | -15.13 | AT3G61320.1 |
| 2868 | HORVU3Hr1G063630 | N-terminal protein myristoylation | -15.14 | AT3G03280.1 |
| 2869 | HORVU5Hr1G009200 | NAD-dependent epimerase/dehydratase | -15.24 | AT1G09340.1 |
| 2870 | HORVU0Hr1G004830 | glyceraldehyde-3-phosphate dehydrogenase C2 | -15.28 | AT1G12900.5 |
| 2871 | HORVU7Hr1G098400 | Chlororespiratory reduction 42 | -15.31 | AT5G20935.1 |
| 2872 | HORVU7Hr1G022050 | undescribed protein | -15.32 | AT1G10170.4 |
| 2873 | HORVU2Hr1G019560 | Adenine nucleotide alpha hydrolases-like superfamily protein | -15.33 | AT2G47710.1 |
| 2874 | HORVU2Hr1G029460 | WRKY family transcription factor | -15.33 | AT4G11070.2 |
| 2875 | HORVU1Hr1G090110 | WRKY family transcription factor | -15.45 | AT5G24110.1 |
| 2876 | HORVU7Hr1G120190 | Ycf48-like protein | -15.53 | AT5G23120.2 |
| 2877 | HORVU7Hr1G041230 | Protein TIFY 3A | -15.57 | AT1G19180.2 |
| 2878 | HORVU3Hr1G085180 | myb-like transcription factor family protein | -15.65 | AT2G38090.1 |
| 2879 | HORVU3Hr1G031130 | Pentatricopeptide repeat-containing protein | -15.71 | AT5G25630.1 |
| 2880 | HORVU6Hr1G011790 | serine/threonine protein kinase 2 | -15.72 | AT3G04810.3 |
| 2881 | HORVU4Hr1G008180 | Protein kinase superfamily protein | -15.73 | AT1G66150.1 |
| 2882 | HORVU2Hr1G080480 | 30S ribosomal protein 3, chloroplastic | -15.80 | AT1G68590.2 |
| 2883 | HORVU3Hr1G052490 | 5'-methylthioadenosine/S-adenosylhomocysteine nucleosidase | -15.87 | AT4G28940.1 |
| 2884 | HORVU5Hr1G053480 | Protein FLUORESCENT IN BLUE LIGHT, chloroplastic | -15.87 | AT3G14110.2 |
| 2885 | HORVU3Hr1G005420 | Lecithin-cholesterol acyltransferase-like 1 | -15.92 | AT1G27480.1 |
| 2886 | HORVU3Hr1G051330 | One-helix protein 2 | -15.93 | AT1G34000.1 |

| | | | | |
|---|---|---|---|---|
| 2887 | HORVU5Hr1G046480 | Chalcone--flavonone isomerase | -15.96 | AT5G05270.2 |
| 2888 | HORVU1Hr1G067300 | phosphoglycerate kinase 1 | -15.98 | AT3G12780.1 |
| 2889 | HORVU7Hr1G111190 | D-3-phosphoglycerate dehydrogenase | -16.01 | AT4G34200.1 |
| 2890 | HORVU6Hr1G051650 | Oxygen-evolving enhancer protein 3-1, chloroplastic | -16.01 | AT1G14150.1 |
| 2891 | HORVU7Hr1G057460 | beta-galactosidase 11 | -16.03 | AT4G38590.1 |
| 2892 | HORVU5Hr1G040430 | undescribed protein | -16.04 | AT5G48970.2 |
| 2893 | HORVU5Hr1G075660 | UDP-glucose 4-epimerase | -16.06 | AT3G23820.1 |
| 2894 | HORVU5Hr1G119830 | UDP-Glycosyltransferase superfamily protein | -16.06 | AT2G36800.1 |
| 2895 | HORVU7Hr1G115600 | heptahelical transmembrane protein1 | -16.09 | AT5G20270.1 |
| 2896 | HORVU1Hr1G043900 | Chloroplast inner envelope protein, putative, expressed | -16.17 | AT1G06950.1 |
| 2897 | HORVU2Hr1G079020 | Ovarian cancer-associated gene 2 protein homolog | -16.33 | AT4G24380.6 |
| 2898 | HORVU4Hr1G082800 | unknown function | -16.43 | AT4G38225.1 |
| 2899 | HORVU5Hr1G088550 | undescribed protein | -16.46 | AT3G43190.2 |
| 2900 | HORVU2Hr1G111580 | undescribed protein | -16.47 | AT1G22990.1 |
| 2901 | HORVU5Hr1G075570 | FASCICLIN-like arabinogalactan 1 | -16.49 | AT4G12730.1 |
| 2902 | HORVU2Hr1G090400 | Protein of unknown function (DUF177) | -16.49 | AT3G19800.2 |
| 2903 | HORVU1Hr1G087570 | FASCICLIN-like arabinogalactan 6 | -16.52 | AT2G20520.1 |
| 2904 | HORVU2Hr1G092710 | Homeobox-leucine zipper protein family | -16.56 | AT2G46680.1 |
| 2905 | HORVU3Hr1G050590 | WRKY DNA-binding protein 50 | -16.57 | AT5G26170.1 |
| 2906 | HORVU6Hr1G070290 | unknown function | -16.59 | AT4G35320.1 |
| 2907 | HORVU2Hr1G039880 | Cyclophilin-like peptidyl-prolyl cis-trans isomerase family protein | -16.60 | AT3G15520.3 |
| 2908 | HORVU1Hr1G029180 | Late embryogenesis abundant (LEA) hydroxyproline-rich glycoprotein family | -16.62 | AT1G17620.1 |
| 2909 | HORVU5Hr1G112670 | Chalcone--flavonone isomerase | -16.83 | AT3G55120.1 |
| 2910 | HORVU3Hr1G071770 | cellulose synthase family protein | -16.88 | AT4G18780.1 |
| 2911 | HORVU2Hr1G039020 | unknown function | -16.96 | AT2G29750.1 |
| 2912 | HORVU6Hr1G084650 | Clustered mitochondria protein | -17.04 | AT4G28080.1 |
| 2913 | HORVU4Hr1G063870 | Subtilisin-like protease | -17.09 | AT4G34980.1 |
| 2914 | HORVU3Hr1G029750 | unknown protein; BEST Arabidopsis thaliana protein match is: unknown protein . | -17.15 | AT5G41761.1 |
| 2915 | HORVU5Hr1G001280 | #N/A | -17.22 | #N/A |
| 2916 | HORVU4Hr1G083020 | hydroxyproline-rich glycoprotein family protein | -17.29 | AT5G51680.1 |
| 2917 | HORVU7Hr1G120060 | Cathepsin B-like cysteine proteinase 1 | -17.32 | AT2G34080.1 |
| 2918 | HORVU7Hr1G036410 | unknown function | -17.36 | AT1G35210.1 |
| 2919 | HORVU1Hr1G024860 | arabinogalactan protein 16 | -17.41 | AT2G46330.1 |
| 2920 | HORVU1Hr1G048610 | RHOMBOID-like 2 | -17.45 | AT1G63120.1 |
| 2921 | HORVU4Hr1G078510 | 1,4-dihydroxy-2-naphthoate phytyltransferase | -17.49 | AT1G60600.2 |
| 2922 | HORVU3Hr1G108550 | undescribed protein | -17.50 | AT3G43920.5 |
| 2923 | HORVU6Hr1G050760 | undescribed protein | -17.50 | AT5G02390.3 |
| 2924 | HORVU3Hr1G059810 | 50S ribosomal protein L7/L12 | -17.51 | AT3G27850.1 |

| | | | | |
|---|---|---|---|---|
| 2925 | HORVU1Hr1G035720 | ribulose bisphosphate carboxylase small chain 1A | -17.56 | AT1G67090.1 |
| 2926 | HORVU7Hr1G006880 | undescribed protein | -17.56 | AT3G60380.1 |
| 2927 | HORVU4Hr1G075090 | unknown protein; Has 39 Blast hits to 39 proteins in 10 species: Archae - 0; Bacteria - 0; Metazoa - 0; Fungi - 0; Plants - 39; Viruses - 0; Other Eukaryotes - 0 (source: NCBI BLink). | -17.58 | AT2G35290.1 |
| 2928 | HORVU1Hr1G026080 | undescribed protein | -17.70 | AT4G33500.1 |
| 2929 | HORVU1Hr1G050830 | dihydroflavonol 4-reductase-like1 | -17.71 | AT2G23910.1 |
| 2930 | HORVU4Hr1G080820 | Group 3 late embryogenesis abundant protein | -17.75 | AT1G52690.2 |
| 2931 | HORVU6Hr1G080870 | zinc finger protein 8 | -17.75 | AT2G41940.1 |
| 2932 | HORVU2Hr1G002870 | undescribed protein | -17.78 | AT4G21120.1 |
| 2933 | HORVU7Hr1G021550 | Myosin heavy chain-related-like | -17.79 | AT3G49055.1 |
| 2934 | HORVU2Hr1G074280 | unknown function | -17.81 | AT5G09820.2 |
| 2935 | HORVU5Hr1G112100 | unknown function | -18.05 | AT3G11740.1 |
| 2936 | HORVU6Hr1G004860 | 3-beta hydroxysteroid dehydrogenase | -18.10 | AT3G63140.1 |
| 2937 | HORVU3Hr1G109500 | Plant protein of unknown function (DUF247) | -18.14 | AT3G50150.1 |
| 2938 | HORVU5Hr1G110940 | UDP-Glycosyltransferase superfamily protein | -18.15 | AT5G49690.1 |
| 2939 | HORVU6Hr1G054890 | undescribed protein | -18.20 | AT5G06760.1 |
| 2940 | HORVU4Hr1G073650 | Heat stress transcription factor A-2d | -18.28 | AT3G22830.1 |
| 2941 | HORVU3Hr1G082210 | myb domain protein 4 | -18.39 | AT4G38620.1 |
| 2942 | HORVU4Hr1G010770 | unknown protein; Has 65 Blast hits to 65 proteins in 11 species: Archae - 0; Bacteria - 0; Metazoa - 0; Fungi - 0; Plants - 65; Viruses - 0; Other Eukaryotes - 0 (source: NCBI BLink). | -18.54 | AT3G57450.1 |
| 2943 | HORVU1Hr1G063450 | alpha/beta-Hydrolases superfamily protein | -18.58 | AT5G16080.1 |
| 2944 | HORVU6Hr1G077790 | Phosphate-responsive 1 family protein | -18.84 | AT4G08950.1 |
| 2945 | HORVU1Hr1G066510 | Domain of unknown function (DUF1995) | -18.85 | AT3G20680.1 |
| 2946 | HORVU2Hr1G039960 | Receptor-like protein kinase 5 | -18.90 | AT2G20850.1 |
| 2947 | HORVU3Hr1G094000 | Auxin efflux carrier family protein | -18.93 | AT5G16530.2 |
| 2948 | HORVU2Hr1G075110 | PLC-like phosphodiesterases superfamily protein | -18.94 | AT1G49740.1 |
| 2949 | HORVU1Hr1G019680 | undescribed protein | -18.99 | AT5G01130.1 |
| 2950 | HORVU5Hr1G067010 | Homeobox-leucine zipper protein family | -19.00 | AT5G06710.1 |
| 2951 | HORVU3Hr1G000900 | Peptidyl-prolyl cis-trans isomerase B | -19.10 | AT1G74070.1 |
| 2952 | HORVU7Hr1G022310 | NAC domain protein, | -19.12 | AT5G63790.1 |
| 2953 | HORVU1Hr1G054260 | basic helix-loop-helix (bHLH) DNA-binding superfamily protein | -19.23 | AT4G00050.1 |
| 2954 | HORVU5Hr1G096260 | UDP-Glycosyltransferase superfamily protein | -19.24 | AT3G02100.1 |
| 2955 | HORVU1Hr1G014140 | unknown protein; BEST Arabidopsis thaliana protein match is: unknown protein . | -19.36 | AT2G15020.1 |
| 2956 | HORVU3Hr1G011030 | Plant protein of unknown function (DUF247) | -19.39 | AT5G22540.1 |
| 2957 | HORVU3Hr1G093310 | Transcription factor bHLH62 | -19.40 | AT3G07340.1 |
| 2958 | HORVU7Hr1G058940 | RNA-binding protein 1 | -19.40 | AT4G24770.2 |

| | | | | |
|---|---|---|---|---|
| 2959 | HORVU5Hr1G094770 | photosystem II 11 kDa protein-related | -19.58 | AT1G05385.2 |
| 2960 | HORVU3Hr1G093990 | Haloalkane dehalogenase 2 | -19.69 | AT4G12830.1 |
| 2961 | HORVU5Hr1G018970 | 0 | -19.84 | #N/A |
| 2962 | HORVU5Hr1G062240 | Chlorophyll a-b binding protein 7, chloroplastic | -19.86 | AT1G19150.1 |
| 2963 | HORVU5Hr1G099020 | Leucine carboxyl methyltransferase | -19.90 | AT5G42760.2 |
| 2964 | HORVU7Hr1G119930 | Cathepsin B-like cysteine proteinase 1 | -19.93 | AT2G34080.1 |
| 2965 | HORVU5Hr1G037620 | undescribed protein | -19.94 | #N/A |
| 2966 | HORVU2Hr1G002010 | Peroxidase superfamily protein | -19.94 | AT1G77490.1 |
| 2967 | HORVU6Hr1G010890 | undescribed protein | -20.10 | AT2G14540.1 |
| 2968 | HORVU4Hr1G059150 | Cysteine-rich receptor-like protein kinase 25 | -20.12 | AT5G48540.1 |
| 2969 | HORVU2Hr1G103130 | Aminomethyltransferase | -20.12 | AT1G11860.2 |
| 2970 | HORVU1Hr1G012110 | xylulose kinase-1 | -20.26 | AT2G21370.1 |
| 2971 | HORVU4Hr1G077350 | unknown function | -20.34 | AT5G20700.1 |
| 2972 | HORVU6Hr1G078500 | fucosyltransferase 1 | -20.34 | AT2G03220.1 |
| 2973 | HORVU3Hr1G025820 | Mitochondrial substrate carrier family protein | -20.55 | AT5G19760.1 |
| 2974 | HORVU5Hr1G098920 | Pectin acetylesterase 9 | -20.63 | AT5G23870.4 |
| 2975 | HORVU3Hr1G068200 | Heavy metal transport/detoxification superfamily protein | -20.64 | AT3G24450.1 |
| 2976 | HORVU2Hr1G084960 | undescribed protein | -20.76 | ATMG00870.1 |
| 2977 | HORVU4Hr1G015570 | Sucrose-phosphate synthase family protein | -20.79 | AT4G10120.3 |
| 2978 | HORVU7Hr1G078480 | unknown function | -20.84 | AT1G26220.1 |
| 2979 | HORVU4Hr1G036300 | undescribed protein | -20.86 | AT2G48060.3 |
| 2980 | HORVU3Hr1G089910 | Trigger factor | -21.27 | AT2G30695.3 |
| 2981 | HORVU7Hr1G090340 | phosphatidic acid phosphatase-related / PAP2-related | -21.32 | AT3G15820.1 |
| 2982 | HORVU2Hr1G109040 | myb-like transcription factor family protein | -21.36 | AT2G01060.1 |
| 2983 | HORVU4Hr1G071730 | Inositol-3-phosphate synthase | -21.37 | AT2G22240.2 |
| 2984 | HORVU2Hr1G126040 | NAC domain containing protein 76 | -21.43 | AT1G71930.2 |
| 2985 | HORVU4Hr1G060200 | RNA polymerase sigma-C factor | -21.43 | AT1G08540.1 |
| 2986 | HORVU2Hr1G111300 | Bifunctional lysine-specific demethylase and histidyl-hydroxylase NO66 | -21.55 | AT3G11560.5 |
| 2987 | HORVU4Hr1G089990 | Protein WEAK CHLOROPLAST MOVEMENT UNDER BLUE LIGHT 1 | -21.59 | AT2G26570.2 |
| 2988 | HORVU3Hr1G000150 | Chromosome 3B, genomic scaffold, cultivar Chinese Spring | -21.64 | AT1G10585.1 |
| 2989 | HORVU3Hr1G061130 | FASCICLIN-like arabinogalactan-protein 11 | -21.66 | AT5G03170.1 |
| 2990 | HORVU4Hr1G066870 | Cytochrome P450 superfamily protein | -21.87 | AT3G56630.1 |
| 2991 | HORVU2Hr1G099470 | Bifunctional inhibitor/lipid-transfer protein/seed storage 2S albumin superfamily protein | -21.92 | AT3G22142.1 |
| 2992 | HORVU0Hr1G038040 | thionin 2.1 | -22.03 | AT1G66100.1 |
| 2993 | HORVU5Hr1G064230 | Cellulose synthase family protein | -22.03 | AT5G17420.1 |
| 2994 | HORVU3Hr1G095080 | MADS-box transcription factor family protein | -22.10 | AT4G22950.2 |
| 2995 | HORVU2Hr1G029450 | Transcription factor WRKY | -22.12 | AT4G11070.2 |

| | | | | |
|---|---|---|---|---|
| 2996 | HORVU3Hr1G013180 | Serine/threonine-protein kinase | -22.22 | AT2G19130.1 |
| 2997 | HORVU7Hr1G117710 | O-methyltransferase family protein | -22.24 | AT4G35160.1 |
| 2998 | HORVU4Hr1G028720 | xyloglucan endotransglucosylase/hydrolase 5 | -22.36 | AT5G13870.3 |
| 2999 | HORVU1Hr1G051700 | Protein CURVATURE THYLAKOID 1B, chloroplastic | -22.37 | AT2G46820.2 |
| 3000 | HORVU7Hr1G096250 | high chlorophyll fluorescence phenotype 173 | -22.37 | AT1G16720.1 |
| 3001 | HORVU6Hr1G094040 | undescribed protein | -22.37 | AT3G62360.2 |
| 3002 | HORVU2Hr1G063460 | Peroxidase superfamily protein | -22.46 | AT4G37530.1 |
| 3003 | HORVU2Hr1G065450 | flowering promoting factor 1 | -22.55 | AT5G24860.1 |
| 3004 | HORVU5Hr1G120390 | Thiopurine S-methyltransferase | -22.67 | AT2G43940.1 |
| 3005 | HORVU3Hr1G066390 | Basic helix-loop-helix protein A | -22.80 | AT2G46510.1 |
| 3006 | HORVU7Hr1G080510 | phosphoenolpyruvate carboxylase 3 | -22.83 | AT3G14940.2 |
| 3007 | HORVU5Hr1G067490 | 1-aminocyclopropane-1-carboxylate oxidase 1 | -23.01 | AT1G05010.1 |
| 3008 | HORVU7Hr1G096370 | unknown protein; Has 30201 Blast hits to 17322 proteins in 780 species: Archae - 12; Bacteria - 1396; Metazoa - 17338; Fungi - 3422; Plants - 5037; Viruses - 0; Other Eukaryotes - 2996 (source: NCBI BLink). | -23.04 | AT1G78995.1 |
| 3009 | HORVU6Hr1G094410 | unknown function | -23.23 | AT3G19900.1 |
| 3010 | HORVU7Hr1G006980 | NBS-LRR disease resistance protein | -23.28 | AT3G07040.1 |
| 3011 | HORVU2Hr1G021270 | Dehydrogenase/reductase SDR family member 4 | -23.30 | AT4G03140.1 |
| 3012 | HORVU4Hr1G016450 | undescribed protein | -23.40 | AT5G09860.1 |
| 3013 | HORVU2Hr1G029950 | undescribed protein | -23.40 | AT2G27660.1 |
| 3014 | HORVU3Hr1G095000 | undescribed protein | -23.49 | AT5G09950.2 |
| 3015 | HORVU6Hr1G069030 | unknown function | -23.51 | AT3G09510.1 |
| 3016 | HORVU5Hr1G096420 | WRKY DNA-binding protein 57 | -23.57 | AT1G69310.4 |
| 3017 | HORVU3Hr1G058210 | undescribed protein | -23.62 | AT1G59820.1 |
| 3018 | HORVU6Hr1G035470 | myb domain protein r1 | -23.71 | AT5G67300.1 |
| 3019 | HORVU6Hr1G061270 | Aldehyde dehydrogenase family 3 member F1 | -23.75 | AT4G36250.1 |
| 3020 | HORVU2Hr1G043780 | GRAS family transcription factor | -23.81 | AT5G48150.4 |
| 3021 | HORVU1Hr1G057640 | peptide chain release factor, putative | -23.83 | AT3G57190.1 |
| 3022 | HORVU3Hr1G056960 | Siroheme synthase | -24.03 | AT5G40850.1 |
| 3023 | HORVU4Hr1G082700 | glyceraldehyde-3-phosphate dehydrogenase C2 | -24.09 | AT1G42970.1 |
| 3024 | HORVU2Hr1G095080 | Cytochrome P450 superfamily protein | -24.14 | AT1G12740.1 |
| 3025 | HORVU3Hr1G086190 | Photosystem II reaction center Psb28 protein | -24.38 | AT1G13270.1 |
| 3026 | HORVU7Hr1G036990 | undescribed protein | -24.40 | AT3G06530.5 |
| 3027 | HORVU5Hr1G032980 | fructose-bisphosphate aldolase 2 | -24.41 | AT4G38970.1 |
| 3028 | HORVU0Hr1G017530 | BnaC01g00720D protein | -24.46 | AT2G46020.6 |
| 3029 | HORVU4Hr1G057000 | Low temperature and salt responsive protein family | -24.48 | AT2G38905.1 |
| 3030 | HORVU2Hr1G009970 | Eukaryotic aspartyl protease family protein | -24.81 | AT2G03200.1 |
| 3031 | HORVU6Hr1G077770 | EXORDIUM like 2 | -24.96 | AT4G08950.1 |

| | | | | |
|---|---|---|---|---|
| 3032 | HORVU2Hr1G021870 | Tryptophan/tyrosine permease | -25.24 | AT5G19500.1 |
| 3033 | HORVU1Hr1G071410 | undescribed protein | -25.25 | AT3G09530.1 |
| 3034 | HORVU3Hr1G087420 | DNA-3-methyladenine glycosylase 1 | -25.27 | AT1G13635.2 |
| 3035 | HORVU2Hr1G008090 | Transmembrane amino acid transporter family protein | -25.33 | AT2G39130.2 |
| 3036 | HORVU5Hr1G035300 | P-loop containing nucleoside triphosphate hydrolases superfamily protein | -25.40 | AT2G46620.1 |
| 3037 | HORVU5Hr1G077390 | Transcription factor bHLH137 | -25.50 | AT5G50915.4 |
| 3038 | HORVU7Hr1G047690 | Chromosome 3B, genomic scaffold, cultivar Chinese Spring | -25.73 | AT4G20970.1 |
| 3039 | HORVU5Hr1G081060 | Cytochrome P450 superfamily protein | -25.76 | AT2G46660.1 |
| 3040 | HORVU1Hr1G013740 | Dihydrolipoyl dehydrogenase | -26.12 | AT1G48030.5 |
| 3041 | HORVU7Hr1G037600 | sodium/calcium exchanger family protein / calcium-binding EF hand family protein | -26.14 | AT1G53210.1 |
| 3042 | HORVU6Hr1G093940 | Peroxidase superfamily protein | -26.19 | AT1G77490.1 |
| 3043 | HORVU4Hr1G040100 | undescribed protein | -26.28 | AT1G28520.3 |
| 3044 | HORVU5Hr1G069480 | Protein of unknown function, DUF547 | -26.33 | AT4G37080.3 |
| 3045 | HORVU2Hr1G001320 | Protein NRT1/ PTR FAMILY 8.5 | -26.47 | AT1G62200.1 |
| 3046 | HORVU1Hr1G091530 | DnaJ homolog subfamily C member 2 | -26.59 | AT1G75250.2 |
| 3047 | HORVU3Hr1G025020 | Chromosome 3B, genomic scaffold, cultivar Chinese Spring | -26.74 | AT5G19875.1 |
| 3048 | HORVU1Hr1G069560 | Protein kinase superfamily protein | -26.78 | AT5G01920.2 |
| 3049 | HORVU6Hr1G071950 | Zinc finger protein CONSTANS-LIKE 10 | -26.91 | AT5G48250.1 |
| 3050 | HORVU5Hr1G099170 | Protein NRT1/ PTR FAMILY 6.4 | -27.11 | AT3G21670.1 |
| 3051 | HORVU3Hr1G024920 | undescribed protein | -27.27 | AT5G17650.1 |
| 3052 | HORVU4Hr1G043630 | undescribed protein | -27.28 | AT2G46930.1 |
| 3053 | HORVU5Hr1G112880 | zinc-finger protein 1 | -27.36 | AT3G53600.1 |
| 3054 | HORVU1Hr1G050730 | undescribed protein | -27.39 | AT2G35470.1 |
| 3055 | HORVU2Hr1G095380 | undescribed protein | -27.54 | AT1G71120.1 |
| 3056 | HORVU3Hr1G040050 | undescribed protein | -27.73 | AT4G15720.1 |
| 3057 | HORVU7Hr1G097670 | Macrophage migration inhibitory factor homolog | -27.77 | AT5G57170.1 |
| 3058 | HORVU7Hr1G098370 | Xyloglucan endotransglucosylase/hydrolase family protein | -27.81 | AT5G57560.1 |
| 3059 | HORVU3Hr1G076570 | undescribed protein | -27.87 | AT5G56540.1 |
| 3060 | HORVU1Hr1G062450 | unknown protein; BEST Arabidopsis thaliana protein match is: unknown protein . | -28.09 | AT3G06070.1 |
| 3061 | HORVU2Hr1G005570 | 2-oxoglutarate (2OG) and Fe(II)-dependent oxygenase superfamily protein | -28.20 | AT1G06620.1 |
| 3062 | HORVU2Hr1G116660 | BnaC07g36720D protein | -28.25 | AT4G21445.1 |
| 3063 | HORVU4Hr1G082430 | undescribed protein | -28.25 | AT1G27090.1 |
| 3064 | HORVU3Hr1G110660 | NADH dehydrogenase-like complex O | -28.46 | AT1G74880.1 |
| 3065 | HORVU7Hr1G087630 | undescribed protein | -28.50 | AT5G57580.1 |
| 3066 | HORVU6Hr1G095380 | multidrug MFS transporter | -28.92 | AT2G21280.2 |
| 3067 | HORVU6Hr1G048770 | undescribed protein | -28.95 | AT2G38360.1 |

| | | | | |
|---|---|---|---|---|
| 3068 | HORVU3Hr1G092820 | myb domain protein 112 | -28.96 | AT1G48000.1 |
| 3069 | HORVU4Hr1G086020 | 4-hydroxy-tetrahydrodipicolinate reductase | -29.14 | AT5G52100.1 |
| 3070 | HORVU3Hr1G039310 | undescribed protein | -29.17 | AT1G29720.1 |
| 3071 | HORVU5Hr1G077520 | unknown protein; BEST Arabidopsis thaliana protein match is: unknown protein . | -29.34 | AT4G22758.2 |
| 3072 | HORVU1Hr1G082310 | Ethylene-responsive transcription factor 4 | -29.39 | AT4G34410.1 |
| 3073 | HORVU7Hr1G119060 | GDSL esterase/lipase | -29.77 | AT5G33370.1 |
| 3074 | HORVU5Hr1G096490 | Blue copper protein | -29.89 | AT2G27035.1 |
| 3075 | HORVU2Hr1G065420 | undescribed protein | -30.08 | AT4G22390.2 |
| 3076 | HORVU3Hr1G114250 | unknown function | -30.09 | AT3G10550.1 |
| 3077 | HORVU5Hr1G068110 | basic helix-loop-helix (bHLH) DNA-binding superfamily protein | -30.14 | AT5G67060.2 |
| 3078 | HORVU6Hr1G093240 | High-light-induced protein, chloroplastic | -30.37 | AT5G02120.1 |
| 3079 | HORVU7Hr1G037020 | undescribed protein | -30.40 | #N/A |
| 3080 | HORVU3Hr1G039670 | undescribed protein | -30.86 | #N/A |
| 3081 | HORVU4Hr1G043650 | Rubredoxin-like superfamily protein | -30.99 | AT5G51010.1 |
| 3082 | HORVU3Hr1G058830 | TRICHOME BIREFRINGENCE-LIKE 19 | -31.03 | AT5G15900.1 |
| 3083 | HORVU5Hr1G118060 | Harpin-induced protein 1 containing protein | -31.04 | AT2G35460.1 |
| 3084 | HORVU6Hr1G048660 | E3 SUMO-protein ligase SIZ2 | -31.19 | AT3G16800.5 |
| 3085 | HORVU7Hr1G037010 | undescribed protein | -31.25 | AT1G33410.2 |
| 3086 | HORVU5Hr1G049590 | undescribed protein | -31.25 | AT1G66210.1 |
| 3087 | HORVU5Hr1G062720 | SAUR-like auxin-responsive protein family | -31.27 | AT5G20820.1 |
| 3088 | HORVU6Hr1G026370 | BEST Arabidopsis thaliana protein match is: Galact s in 8 species: Archae - 0; Bacteria - 0; Metazoa - 0; Fungi - 0; Plants - 30; Viruses - 0; Other Eukaryotes - 0 (source: NCBI BLink). | -31.47 | AT1G30260.1 |
| 3089 | HORVU1Hr1G044080 | Late embryogenesis abundant (LEA) hydroxyproline-rich glycoprotein family | -31.64 | AT1G65690.1 |
| 3090 | HORVU1Hr1G026070 | 0 | -31.65 | #N/A |
| 3091 | HORVU5Hr1G072770 | undescribed protein | -31.95 | AT3G21570.1 |
| 3092 | HORVU6Hr1G056210 | Retrotransposon protein, putative, unclassified | -32.03 | AT1G43760.1 |
| 3093 | HORVU6Hr1G010520 | unknown protein; Has 52 Blast hits to 46 proteins in 20 species: Archae - 0; Bacteria - 0; Metazoa - 0; Fungi - 0; Plants - 45; Viruses - 0; Other Eukaryotes - 7 (source: NCBI BLink). | -32.19 | AT3G50685.1 |
| 3094 | HORVU2Hr1G075950 | zinc finger homeodomain 1 | -32.36 | AT2G02540.2 |
| 3095 | HORVU6Hr1G059850 | 3'(2'),5'-bisphosphate nucleotidase 2 | -32.54 | AT4G05090.1 |
| 3096 | HORVU5Hr1G028180 | LIM domain-containing protein 2 | -32.85 | AT1G10200.1 |
| 3097 | HORVU2Hr1G035020 | undescribed protein | -32.96 | AT4G25070.2 |
| 3098 | HORVU6Hr1G011990 | unknown function | -33.25 | AT3G61870.2 |
| 3099 | HORVU4Hr1G040110 | undescribed protein | -33.39 | AT3G52330.1 |
| 3100 | HORVU7Hr1G049860 | FASCICLIN-like arabinogalactan-protein 12 | -34.46 | AT2G04780.2 |
| 3101 | HORVU3Hr1G083520 | Glutathione S-transferase family protein | -34.60 | AT1G17170.1 |
| 3102 | HORVU5Hr1G112060 | thionin 2.2 | -34.81 | AT1G66100.1 |

| | | | | |
|---|---|---|---|---|
| 3103 | HORVU6Hr1G066450 | At5g37260-like protein | -34.91 | AT1G18330.1 |
| 3104 | HORVU4Hr1G088160 | Eukaryotic aspartyl protease family protein | -35.41 | AT2G03200.1 |
| 3105 | HORVU6Hr1G076040 | Photosynthetic NDH subcomplex B 4 | -35.88 | AT1G18730.5 |
| 3106 | HORVU3Hr1G078150 | Ethylene-responsive transcription factor 4 | -36.31 | AT3G15210.1 |
| 3107 | HORVU3Hr1G057090 | carbonic anhydrase 2 | -36.39 | AT5G14740.9 |
| 3108 | HORVU3Hr1G087210 | respiratory burst oxidase protein F | -36.56 | AT5G60010.1 |
| 3109 | HORVU4Hr1G056210 | Protein phosphatase 2C family protein | -36.60 | AT2G30020.1 |
| 3110 | HORVU0Hr1G002410 | unknown function | -36.75 | AT5G05190.1 |
| 3111 | HORVU3Hr1G026920 | metallothionein 3 | -36.79 | AT3G15353.1 |
| 3112 | HORVU4Hr1G088110 | Eukaryotic aspartyl protease family protein | -36.93 | AT2G03200.1 |
| 3113 | HORVU5Hr1G043090 | Two-component response regulator ARR8 | -37.00 | AT2G41310.1 |
| 3114 | HORVU5Hr1G023160 | Transmembrane amino acid transporter family protein | -37.12 | AT1G47670.1 |
| 3115 | HORVU3Hr1G105360 | cytokinin oxidase/dehydrogenase 1 | -37.17 | AT2G41510.2 |
| 3116 | HORVU6Hr1G090190 | RNA-binding protein 1 | -37.57 | AT1G60000.1 |
| 3117 | HORVU2Hr1G104030 | Receptor-like protein kinase 2 | -37.67 | AT5G46330.2 |
| 3118 | HORVU7Hr1G023140 | Photosynthetic NDH subunit of subcomplex B 3, chloroplastic | -38.17 | AT3G16250.1 |
| 3119 | HORVU5Hr1G097870 | Cytochrome b6-f complex subunit 7 | -38.34 | AT2G26500.2 |
| 3120 | HORVU0Hr1G009120 | Hydrolase, alpha/beta fold family protein, expressed | -38.59 | AT5G21950.1 |
| 3121 | HORVU3Hr1G083790 | unknown function | -38.92 | AT2G36145.1 |
| 3122 | HORVU0Hr1G012330 | undescribed protein | -39.11 | AT5G42390.1 |
| 3123 | HORVU6Hr1G035460 | undescribed protein | -39.30 | AT1G65640.2 |
| 3124 | HORVU1Hr1G029840 | Retroelement pol polyprotein-like | -39.36 | AT2G05120.2 |
| 3125 | HORVU4Hr1G081040 | ABC transporter G family member 45 | -39.61 | AT2G36380.1 |
| 3126 | HORVU7Hr1G030210 | Jacalin-like lectin domain containing protein | -39.62 | AT1G73040.1 |
| 3127 | HORVU3Hr1G040180 | undescribed protein | -40.08 | AT2G39435.6 |
| 3128 | HORVU5Hr1G010950 | Glutaredoxin family protein | -40.11 | AT5G18600.1 |
| 3129 | HORVU1Hr1G000170 | undescribed protein | -41.32 | AT1G48490.3 |
| 3130 | HORVU4Hr1G009250 | undescribed protein | -41.51 | AT5G44800.1 |
| 3131 | HORVU5Hr1G059210 | undescribed protein | -41.97 | AT3G02230.1 |
| 3132 | HORVU0Hr1G002390 | unknown function | -42.10 | AT4G05060.1 |
| 3133 | HORVU0Hr1G022690 | Senescence-associated protein DIN1 | -43.18 | AT4G35770.1 |
| 3134 | HORVU7Hr1G095870 | 0 | -43.46 | #N/A |
| 3135 | HORVU1Hr1G024180 | FASCICLIN-like arabinogalactan-protein 11 | -43.46 | AT5G03170.1 |
| 3136 | HORVU5Hr1G030220 | undescribed protein | -44.66 | AT1G80490.3 |
| 3137 | HORVU1Hr1G078350 | Phosphoethanolamine N-methyltransferase 1 | -46.14 | AT3G18000.1 |
| 3138 | HORVU2Hr1G040790 | organic cation/carnitine transporter 2 | -46.41 | AT1G79360.1 |
| 3139 | HORVU5Hr1G073800 | dihydroflavonol 4-reductase-like1 | -46.41 | AT2G33590.1 |
| 3140 | HORVU1Hr1G033190 | undescribed protein | -46.69 | AT3G15095.2 |
| 3141 | HORVU2Hr1G063190 | undescribed protein | -47.45 | AT1G24485.3 |

| | | | | |
|---|---|---|---|---|
| 3142 | HORVU7Hr1G041500 | nudix hydrolase homolog 21 | -48.12 | AT1G73540.1 |
| 3143 | HORVU3Hr1G044870 | undescribed protein | -48.22 | AT2G29440.1 |
| 3144 | HORVU7Hr1G007800 | undescribed protein | -48.65 | AT1G32560.1 |
| 3145 | HORVU5Hr1G101990 | undescribed protein | -48.69 | #N/A |
| 3146 | HORVU6Hr1G091840 | MATE efflux family protein | -48.81 | AT4G00350.1 |
| 3147 | HORVU2Hr1G116390 | Chalcone synthase 1 | -50.01 | AT5G13930.1 |
| 3148 | HORVU7Hr1G085070 | undescribed protein | -50.10 | AT5G07680.2 |
| 3149 | HORVU7Hr1G027860 | Glycogen debranching enzyme | -50.22 | AT5G04360.2 |
| 3150 | HORVU3Hr1G050380 | unknown protein; BEST Arabidopsis thaliana protein match is: unknown protein . | -51.31 | AT5G12010.1 |
| 3151 | HORVU7Hr1G052750 | Cytochrome b561 and DOMON domain-containing protein | -51.89 | AT3G25290.2 |
| 3152 | HORVU1Hr1G091540 | Transcription factor RADIALIS | -52.15 | AT4G39250.1 |
| 3153 | HORVU6Hr1G048760 | undescribed protein | -52.34 | AT2G27980.1 |
| 3154 | HORVU1Hr1G065700 | undescribed protein | -53.25 | AT3G09660.2 |
| 3155 | HORVU0Hr1G016540 | Ethylene-responsive transcription factor 4 | -53.95 | AT4G25470.1 |
| 3156 | HORVU6Hr1G025050 | undescribed protein | -54.29 | AT2G20930.1 |
| 3157 | HORVU7Hr1G002210 | Ferredoxin--NADP reductase | -55.08 | AT1G20020.3 |
| 3158 | HORVU2Hr1G040800 | Solute carrier family 22 member 3 | -56.38 | AT1G79360.1 |
| 3159 | HORVU4Hr1G068930 | unknown function | -56.38 | AT4G30845.1 |
| 3160 | HORVU2Hr1G012460 | Adenosine diphosphate glucose pyrophosphatase | -57.90 | AT2G24630.2 |
| 3161 | HORVU2Hr1G076630 | undescribed protein | -59.26 | #N/A |
| 3162 | HORVU5Hr1G102620 | Amaranthin-like lectin | -59.38 | AT3G23800.2 |
| 3163 | HORVU5Hr1G053060 | BnaA04g27590D protein | -59.54 | AT3G57990.1 |
| 3164 | HORVU1Hr1G032950 | undescribed protein | -59.55 | AT4G12680.1 |
| 3165 | HORVU0Hr1G003480 | Organic cation/carnitine transporter 3 | -60.40 | AT1G79360.1 |
| 3166 | HORVU2Hr1G101230 | undescribed protein | -62.88 | AT5G20960.2 |
| 3167 | HORVU3Hr1G073860 | chlororespiratory reduction 7 | -63.47 | AT5G39210.1 |
| 3168 | HORVU4Hr1G076650 | undescribed protein | -64.59 | AT1G09650.1 |
| 3169 | HORVU6Hr1G076880 | Glycine cleavage system H protein | -65.34 | AT1G32470.1 |
| 3170 | HORVU2Hr1G002720 | PLANT CADMIUM RESISTANCE 1 | -66.83 | AT1G14880.1 |
| 3171 | HORVU3Hr1G078170 | unknown function | -66.86 | AT4G23160.3 |
| 3172 | HORVU7Hr1G090850 | fructose-bisphosphate aldolase 2 | -67.88 | AT4G26530.3 |
| 3173 | HORVU1Hr1G076730 | gibberellin 2-oxidase | -68.30 | AT1G78440.1 |
| 3174 | HORVU2Hr1G001620 | Pyruvate dehydrogenase E1 component subunit alpha | -68.73 | AT1G32920.1 |
| 3175 | HORVU5Hr1G080450 | C-repeat-binding factor 4 | -68.88 | AT4G25470.1 |
| 3176 | HORVU3Hr1G012490 | undescribed protein | -69.26 | AT5G51920.1 |
| 3177 | HORVU3Hr1G108680 | Transcription factor ORG2 | -70.09 | AT3G56970.1 |
| 3178 | HORVU4Hr1G076660 | unknown function | -70.95 | AT1G78410.2 |
| 3179 | HORVU7Hr1G081510 | DnaJ/Hsp40 cysteine-rich domain superfamily protein | -71.08 | AT3G47650.1 |

| | | | | |
|---|---|---|---|---|
| 3180 | HORVU3Hr1G085520 | laccase 7 | -74.04 | AT3G09220.1 |
| 3181 | HORVU1Hr1G007630 | TRICHOME BIREFRINGENCE-LIKE 11 | -75.34 | AT5G19160.1 |
| 3182 | HORVU2Hr1G094870 | MATE efflux family protein | -77.01 | AT4G23030.1 |
| 3183 | HORVU7Hr1G095170 | Pre-mRNA-splicing factor syf2, putative | -77.89 | AT2G16860.2 |
| 3184 | HORVU0Hr1G000620 | Chaperone protein dnaJ 3 | -78.46 | AT1G11040.1 |
| 3185 | HORVU6Hr1G075080 | FKBP-like peptidyl-prolyl cis-trans isomerase family protein | -80.20 | AT4G39710.2 |
| 3186 | HORVU6Hr1G068660 | ROP guanine nucleotide exchange factor 5 | -80.43 | AT1G31650.1 |
| 3187 | HORVU4Hr1G009240 | undescribed protein | -80.83 | AT4G24110.1 |
| 3188 | HORVU4Hr1G087760 | Bifunctional inhibitor/lipid-transfer protein/seed storage 2S albumin superfamily protein | -83.71 | AT2G45180.1 |
| 3189 | HORVU2Hr1G066100 | basic helix-loop-helix (bHLH) DNA-binding superfamily protein | -87.50 | AT5G57150.1 |
| 3190 | HORVU2Hr1G073030 | Carbonic anhydrase | -87.70 | AT3G52720.4 |
| 3191 | HORVU1Hr1G093660 | ribonuclease 3 | -97.51 | AT1G26820.1 |
| 3192 | HORVU2Hr1G077650 | ferric reduction oxidase 7 | -97.84 | AT5G49740.3 |
| 3193 | HORVU1Hr1G089380 | Subtilisin-like protease | -102.20 | AT4G10550.1 |
| 3194 | HORVU7Hr1G084850 | undescribed protein | -104.00 | AT4G11450.1 |
| 3195 | HORVU3Hr1G013900 | Cytochrome P450 superfamily protein | -104.60 | AT3G26330.1 |
| 3196 | HORVU2Hr1G010670 | ribulose bisphosphate carboxylase small chain 1A | -105.64 | AT1G67090.1 |
| 3197 | HORVU6Hr1G041610 | ROP guanine nucleotide exchange factor 5 | -107.73 | AT1G31650.1 |
| 3198 | HORVU4Hr1G090860 | metacaspase 1 | -113.07 | AT1G02170.1 |
| 3199 | HORVU0Hr1G012340 | undescribed protein | -116.07 | AT2G46130.1 |
| 3200 | HORVU5Hr1G089230 | Fatty acyl-CoA reductase 1 | -116.71 | AT5G22500.1 |
| 3201 | HORVU7Hr1G092880 | undescribed protein | -119.41 | AT3G03750.2 |
| 3202 | HORVU0Hr1G019740 | unknown function | -124.87 | AT2G26110.1 |
| 3203 | HORVU2Hr1G010690 | ribulose bisphosphate carboxylase small chain 1A | -126.62 | AT1G67090.1 |
| 3204 | HORVU6Hr1G084860 | purple acid phosphatase 10 | -129.29 | AT2G16430.2 |
| 3205 | HORVU3Hr1G027970 | TRICHOME BIREFRINGENCE-LIKE 38 | -129.93 | AT1G29050.1 |
| 3206 | HORVU7Hr1G084860 | undescribed protein | -130.36 | AT3G03305.1 |
| 3207 | HORVU4Hr1G069100 | beta-xylosidase 3 | -145.10 | AT1G78060.1 |
| 3208 | HORVU7Hr1G095860 | undescribed protein | -145.27 | AT5G24165.2 |
| 3209 | HORVU5Hr1G004700 | terpene synthase 21 | -153.43 | AT5G23960.1 |
| 3210 | HORVU5Hr1G005740 | PHYTOENE SYNTHASE | -161.87 | AT5G17230.4 |
| 3211 | HORVU7Hr1G010690 | Acid phosphatase 1 | -193.16 | AT4G29260.1 |
| 3212 | HORVU1Hr1G093570 | Ribonuclease T2 family protein | -199.97 | AT1G14220.1 |
| 3213 | HORVU2Hr1G114680 | 36.4 kDa proline-rich protein | -204.32 | AT3G22142.1 |
| 3214 | HORVU7Hr1G111010 | unknown function | -208.75 | AT3G06130.1 |
| 3215 | HORVU7Hr1G098330 | xyloglucan endotransglucosylase/hydrolase 25 | -219.22 | AT5G57550.1 |
| 3216 | HORVU4Hr1G058590 | unknown function | -263.29 | AT1G80830.2 |

| | | | | |
|---|---|---|---|---|
| 3217 | HORVU7Hr1G098320 | xyloglucan endotransglucosylase/hydrolase 25 | -293.65 | AT5G57550.1 |
| 3218 | HORVU7Hr1G098260 | xyloglucan endotransglucosylase/hydrolase 25 | -319.28 | AT4G25810.1 |
| 3219 | HORVU3Hr1G083360 | Fructose-1,6-bisphosphatase class 1 | -469.73 | AT1G43670.1 |
| 3220 | HORVU7Hr1G098280 | xyloglucan endotransglucosylase/hydrolase 25 | -508.39 | AT4G25810.1 |
| 3221 | HORVU1Hr1G088840 | Phospholipase A1-II 7 | -3505.51 | AT4G18550.1 |

SUPPLIMENTARY TABLE 3

| S No | Feature ID | Barley Annotation | EDGE test: BWT_0Hr vs BWT_72Hr_Bs, tagwise dispersions - Fold change | Ara-Port |
|---|---|---|---|---|
| 1 | HORVU2Hr1G004640 | Cytochrome P450 superfamily protein | 177.37 | AT3G26300.1 |
| 2 | HORVU2Hr1G004600 | Cytochrome P450 superfamily protein | 139.47 | AT3G26310.1 |
| 3 | HORVU2Hr1G004610 | Cytochrome P450 superfamily protein | 117.65 | AT2G45550.1 |
| 4 | HORVU1Hr1G089700 | O-methyltransferase 1 | 108.10 | AT5G54160.1 |
| 5 | HORVU6Hr1G012640 | Chymotrypsin inhibitor | 102.14 | AT5G43580.1 |
| 6 | HORVU2Hr1G098710 | undescribed protein | 100.57 | AT3G49845.2 |
| 7 | HORVU2Hr1G004530 | Cytochrome P450 superfamily protein | 99.84 | AT5G25120.1 |
| 8 | HORVU7Hr1G012240 | undescribed protein | 97.83 | AT5G63840.2 |
| 9 | HORVU7Hr1G011930 | undescribed protein | 94.91 | AT1G45474.2 |
| 10 | HORVU2Hr1G004620 | Copalyl diphosphate synthase 2 | 94.14 | AT4G02780.1 |
| 11 | HORVU6Hr1G001850 | undescribed protein | 90.36 | AT2G29780.1 |
| 12 | HORVU1Hr1G089620 | O-methyltransferase 1 | 86.52 | AT5G54160.1 |
| 13 | HORVU5Hr1G041660 | protein kinase family protein | 86.28 | AT1G51890.1 |
| 14 | HORVU7Hr1G011950 | undescribed protein | 81.79 | AT5G35620.3 |
| 15 | HORVU3Hr1G112050 | undescribed protein | 79.11 | AT3G55640.3 |
| 16 | HORVU7Hr1G033530 | Cysteine-rich venom protein | 77.60 | AT2G14610.1 |
| 17 | HORVU6Hr1G014600 | Bowman-Birk type trypsin inhibitor | 76.65 | AT1G61420.1 |
| 18 | HORVU1Hr1G089520 | O-methyltransferase family protein | 76.38 | AT4G35160.1 |
| 19 | HORVU6Hr1G020540 | Cytochrome P450 superfamily protein | 75.86 | AT3G26230.1 |
| 20 | HORVU1Hr1G065670 | undescribed protein | 74.39 | AT5G47910.1 |
| 21 | HORVU1Hr1G089540 | Chalcone synthase 2 | 73.93 | AT5G13930.1 |
| 22 | HORVU6Hr1G005720 | high affinity nitrate transporter 2.6 | 73.73 | AT1G08090.1 |

| | | | | |
|---|---|---|---|---|
| 23 | HORVU6Hr1G013690 | undescribed protein | 73.02 | AT3G51350.1 |
| 24 | HORVU7Hr1G011940 | undescribed protein | 72.83 | AT5G60540.1 |
| 25 | HORVU2Hr1G028660 | Chymotrypsin inhibitor | 71.47 | AT2G38900.2 |
| 26 | HORVU5Hr1G096850 | Cytochrome P450 superfamily protein | 71.24 | AT5G57260.1 |
| 27 | HORVU2Hr1G004480 | Cytochrome P450 superfamily protein | 71.08 | AT3G26300.1 |
| 28 | HORVU2Hr1G126150 | 1-deoxy-D-xylulose 5-phosphate synthase 1 | 71.06 | AT4G15560.1 |
| 29 | HORVU2Hr1G122570 | early nodulin-like protein 12 | 68.19 | AT5G26330.1 |
| 30 | HORVU3Hr1G099760 | L-gulonolactone oxidase 2 | 67.54 | AT2G46750.1 |
| 31 | HORVU2Hr1G079440 | UDP-Glycosyltransferase superfamily protein | 65.85 | AT1G22360.1 |
| 32 | HORVU3Hr1G074960 | Peroxidase superfamily protein | 65.55 | AT4G11290.1 |
| 33 | HORVU1Hr1G005510 | Protein kinase family protein | 64.16 | AT1G21240.1 |
| 34 | HORVU0Hr1G011730 | unknown function | 63.98 | AT5G65440.2 |
| 35 | HORVU2Hr1G028720 | Chymotrypsin inhibitor | 63.46 | AT5G43580.1 |
| 36 | HORVU2Hr1G122580 | Blue copper protein | 61.79 | AT2G32300.1 |
| 37 | HORVU0Hr1G011720 | Major pollen allergen Bet v 1-B | 58.38 | AT1G35260.1 |
| 38 | HORVU4Hr1G001250 | Caffeic acid 3-O-methyltransferase | 55.18 | AT5G54160.1 |
| 39 | HORVU5Hr1G124650 | ABC transporter G family member 48 | 55.00 | AT1G66950.1 |
| 40 | HORVU7Hr1G017640 | Enolase | 54.89 | AT2G36530.1 |
| 41 | HORVU2Hr1G028670 | Chymotrypsin inhibitor | 54.87 | AT5G43580.1 |
| 42 | HORVU5Hr1G055950 | Cysteine-rich venom protein | 54.75 | AT4G33720.1 |
| 43 | HORVU5Hr1G056040 | Cysteine-rich venom protein | 54.57 | AT4G33720.1 |
| 44 | HORVU5Hr1G114700 | UDP-Glycosyltransferase superfamily protein | 54.30 | AT3G11340.1 |
| 45 | HORVU7Hr1G076120 | Peroxidase superfamily protein | 54.23 | AT4G33420.1 |
| 46 | HORVU4Hr1G071370 | Membrane protein | 54.20 | AT5G47530.1 |
| 47 | HORVU7Hr1G019380 | Cytochrome P450 superfamily protein | 52.24 | AT3G26300.1 |
| 48 | HORVU2Hr1G104240 | germin-like protein 2 | 51.93 | AT5G39160.1 |
| 49 | HORVU4Hr1G010740 | undescribed protein | 50.60 | AT1G72060.1 |
| 50 | HORVU3Hr1G032370 | Isoflavone reductase-like protein | 50.25 | AT4G39230.1 |
| 51 | HORVU3Hr1G097860 | Laccase-19 | 49.59 | AT3G09220.1 |
| 52 | HORVU5Hr1G023720 | Pathogenesis-related protein STH-2 | 49.23 | AT1G35260.1 |
| 53 | HORVU4Hr1G078370 | undescribed protein | 48.33 | AT1G78880.1 |
| 54 | HORVU6Hr1G053710 | High-affinity nitrate transporter 3.1 | 46.89 | AT5G50200.3 |
| 55 | HORVU7Hr1G009760 | undescribed protein | 46.51 | AT4G26440.1 |
| 56 | HORVU1Hr1G084900 | 1-deoxy-D-xylulose 5-phosphate synthase 1 | 46.40 | AT4G15560.1 |
| 57 | HORVU7Hr1G010110 | undescribed protein | 46.22 | AT2G18640.1 |
| 58 | HORVU2Hr1G076320 | undescribed protein | 45.63 | AT4G34600.1 |
| 59 | HORVU5Hr1G051970 | Pathogenesis-related thaumatin superfamily protein | 45.40 | AT4G11650.1 |

| | | | | |
|---|---|---|---|---|
| 60 | HORVU0Hr1G008830 | MLO-like protein 1 | 45.28 | AT4G02600.2 |
| 61 | HORVU2Hr1G124300 | Glutathione S-transferase family protein | 44.92 | AT1G78380.1 |
| 62 | HORVU5Hr1G056420 | Flavin-containing monooxygenase family protein | 44.82 | AT1G19250.1 |
| 63 | HORVU5Hr1G075540 | Oxidoreductase, zinc-binding dehydrogenase family protein | 44.60 | AT4G13010.1 |
| 64 | HORVU4Hr1G002640 | Pathogenesis-related thaumatin superfamily protein | 44.50 | AT4G11650.1 |
| 65 | HORVU4Hr1G064790 | glutamate decarboxylase | 44.30 | AT5G17330.1 |
| 66 | HORVU5Hr1G099390 | NAC domain containing protein 42 | 43.84 | AT2G43000.1 |
| 67 | HORVU3Hr1G109350 | receptor kinase 1 | 43.37 | AT4G03230.3 |
| 68 | HORVU1Hr1G009490 | Bifunctional inhibitor/lipid-transfer protein/seed storage 2S albumin superfamily protein | 42.92 | AT3G22600.1 |
| 69 | HORVU2Hr1G071570 | Potassium transporter family protein | 42.42 | AT4G13420.1 |
| 70 | HORVU2Hr1G033520 | dihydroflavonol 4-reductase | 41.65 | AT4G27250.1 |
| 71 | HORVU2Hr1G095460 | Glutathione S-transferase family protein | 40.75 | AT3G62760.1 |
| 72 | HORVU3Hr1G077790 | Polyphenol oxidase, chloroplastic | 40.48 | AT3G02210.1 |
| 73 | HORVU5Hr1G009840 | RAB GDP dissociation inhibitor 2 | 40.47 | AT3G59920.1 |
| 74 | HORVU5Hr1G072520 | FAD-binding Berberine family protein | 40.43 | AT4G20820.1 |
| 75 | HORVU2Hr1G122270 | Glutathione S-transferase family protein | 40.19 | AT1G10360.1 |
| 76 | HORVU2Hr1G122620 | Blue copper protein | 40.15 | AT2G31050.1 |
| 77 | HORVU4Hr1G084330 | Inter-alpha-trypsin inhibitor heavy chain H4 | 40.09 | AT5G60710.1 |
| 78 | HORVU7Hr1G008830 | Glutathione S-transferase family protein | 39.87 | AT1G10360.1 |
| 79 | HORVU3Hr1G001840 | undescribed protein | 39.55 | #N/A |
| 80 | HORVU5Hr1G005230 | 0 | 39.54 | AT4G11650.1 |
| 81 | HORVU1Hr1G020720 | Peroxidase superfamily protein | 38.41 | AT5G19890.1 |
| 82 | HORVU5Hr1G123550 | receptor kinase 2 | 38.36 | AT4G21390.1 |
| 83 | HORVU2Hr1G001780 | WRKY DNA-binding protein 75 | 38.21 | AT5G13080.1 |
| 84 | HORVU7Hr1G019400 | receptor kinase 1 | 38.18 | AT2G37710.1 |
| 85 | HORVU1Hr1G022430 | Leucine aminopeptidase 1 | 37.98 | AT2G24200.3 |
| 86 | HORVU4Hr1G002650 | Pathogenesis-related thaumatin superfamily protein | 37.25 | AT4G11650.1 |
| 87 | HORVU1Hr1G062380 | CASP-like protein 1 | 36.42 | AT3G06390.1 |
| 88 | HORVU5Hr1G106010 | Cysteine-rich secretory protein 3 | 36.18 | AT3G19690.1 |
| 89 | HORVU7Hr1G010990 | L-tyrosine decarboxylase | 36.12 | AT2G20340.1 |
| 90 | HORVU3Hr1G094340 | Chromosome 3B, genomic scaffold, cultivar Chinese Spring | 36.05 | AT1G14590.1 |
| 91 | HORVU1Hr1G088550 | undescribed protein | 36.03 | AT4G10930.2 |
| 92 | HORVU3Hr1G112060 | undescribed protein | 35.91 | AT2G42270.1 |
| 93 | HORVU5Hr1G023690 | Pathogenesis-related protein STH-21 | 35.89 | AT5G51770.1 |

| | | | | |
|---|---|---|---|---|
| 94 | HORVU6Hr1G088400 | Leucine-rich receptor-like protein kinase family protein | 35.49 | AT3G47570.1 |
| 95 | HORVU6Hr1G077940 | unknown function | 35.42 | AT1G66170.1 |
| 96 | HORVU6Hr1G077960 | UDP-Glycosyltransferase superfamily protein | 35.06 | AT1G22360.2 |
| 97 | HORVU3Hr1G105400 | Glucan endo-1,3-beta-glucosidase GV | 34.82 | AT3G57270.1 |
| 98 | HORVU2Hr1G092440 | undescribed protein | 34.60 | AT2G32240.1 |
| 99 | HORVU3Hr1G112080 | undescribed protein | 33.83 | AT2G14680.4 |
| 100 | HORVU4Hr1G011760 | UDP-Glycosyltransferase superfamily protein | 33.70 | AT3G11340.1 |
| 101 | HORVU3Hr1G027910 | Cathepsin B-like cysteine proteinase 2 | 33.64 | AT5G45890.1 |
| 102 | HORVU3Hr1G112030 | Abscisic stress-ripening protein 2 | 33.56 | AT3G01410.2 |
| 103 | HORVU5Hr1G099460 | NAC domain protein, | 33.53 | AT2G43000.1 |
| 104 | HORVU3Hr1G098620 | Chromosome 3B, genomic scaffold, cultivar Chinese Spring | 33.41 | AT4G08350.1 |
| 105 | HORVU3Hr1G111600 | Pathogenesis-related protein | 33.30 | AT1G64790.1 |
| 106 | HORVU2Hr1G109640 | Cytochrome P450 superfamily protein | 32.85 | AT2G45550.1 |
| 107 | HORVU3Hr1G112140 | Abscisic stress-ripening protein 1 | 32.61 | #N/A |
| 108 | HORVU6Hr1G021870 | Homeobox-leucine zipper protein family | 32.41 | AT5G06710.1 |
| 109 | HORVU3Hr1G094250 | Chromosome 3B, genomic scaffold, cultivar Chinese Spring | 31.97 | AT4G19975.1 |
| 110 | HORVU7Hr1G097350 | YELLOW STRIPE like 3 | 31.89 | AT5G53550.6 |
| 111 | HORVU1Hr1G090930 | unknown function | 31.82 | AT2G31900.4 |
| 112 | HORVU1Hr1G049070 | Glutathione S-transferase family protein | 31.66 | AT1G10370.1 |
| 113 | HORVU5Hr1G023730 | Pathogenesis-related protein STH-21 | 31.62 | AT1G35260.1 |
| 114 | HORVU3Hr1G096860 | Potassium transporter family protein | 31.54 | AT4G13420.1 |
| 115 | HORVU3Hr1G109430 | receptor kinase 2 | 31.25 | AT4G03230.3 |
| 116 | HORVU7Hr1G010120 | unknown function | 31.20 | AT5G01130.1 |
| 117 | HORVU7Hr1G107670 | Two-component response regulator ORR21 | 31.13 | AT1G14600.1 |
| 118 | HORVU2Hr1G119500 | Late embryogenesis abundant (LEA) hydroxyproline-rich glycoprotein family | 30.76 | AT5G36970.1 |
| 119 | HORVU6Hr1G061420 | Cytochrome b5 | 30.64 | AT5G53560.1 |
| 120 | HORVU2Hr1G088640 | Caleosin-related family protein | 30.55 | AT2G33380.1 |
| 121 | HORVU3Hr1G105630 | Glucan endo-1,3-beta-glucosidase GI | 30.28 | AT4G16260.1 |
| 122 | HORVU2Hr1G079420 | UDP-Glycosyltransferase superfamily protein | 30.17 | AT1G22360.1 |
| 123 | HORVU3Hr1G018980 | basic helix-loop-helix (bHLH) DNA-binding family protein | 30.04 | AT5G56960.1 |
| 124 | HORVU5Hr1G066460 | unknown protein; BEST Arabidopsis thaliana protein match is: unknown protein . | 29.92 | AT1G23110.7 |
| 125 | HORVU5Hr1G080090 | serine carboxypeptidase-like 17 | 29.75 | AT1G33540.2 |
| 126 | HORVU2Hr1G004540 | KS protein | 29.75 | AT1G79460.1 |

| | | | | |
|---|---|---|---|---|
| 127 | HORVU2Hr1G114390 | L-tyrosine decarboxylase | 29.52 | AT2G20340.1 |
| 128 | HORVU2Hr1G098700 | undescribed protein | 29.50 | #N/A |
| 129 | HORVU3Hr1G113120 | Wound-induced protein | 29.50 | AT3G04720.1 |
| 130 | HORVU2Hr1G004510 | Cytochrome P450 superfamily protein | 29.39 | AT5G25120.1 |
| 131 | HORVU3Hr1G100190 | Jasmonate O-methyltransferase | 29.38 | AT5G04370.1 |
| 132 | HORVU3Hr1G020210 | WRKY DNA-binding protein 72 | 28.82 | AT5G15130.1 |
| 133 | HORVU4Hr1G009230 | receptor kinase 2 | 28.80 | AT5G10530.1 |
| 134 | HORVU2Hr1G124970 | Peroxidase superfamily protein | 28.58 | AT1G71695.1 |
| 135 | HORVU7Hr1G110570 | Protein of unknown function (DUF679) | 28.13 | AT4G24310.1 |
| 136 | HORVU1Hr1G014500 | undescribed protein | 27.72 | AT1G66730.1 |
| 137 | HORVU2Hr1G102720 | Protein NRT1/ PTR FAMILY 8.3 | 27.71 | AT2G02040.1 |
| 138 | HORVU3Hr1G069320 | WRKY DNA-binding protein 65 | 27.49 | AT1G29280.1 |
| 139 | HORVU2Hr1G003210 | Cytochrome P450 superfamily protein | 27.44 | AT1G11680.1 |
| 140 | HORVU7Hr1G082180 | undescribed protein | 27.29 | AT1G16030.1 |
| 141 | HORVU2Hr1G114440 | L-tyrosine decarboxylase | 27.07 | AT2G20340.1 |
| 142 | HORVU6Hr1G088440 | 2-oxoglutarate (2OG) and Fe(II)-dependent oxygenase superfamily protein | 26.98 | AT5G20400.1 |
| 143 | HORVU2Hr1G071270 | Ethylene-responsive transcription factor 1 | 26.96 | AT5G13330.1 |
| 144 | HORVU3Hr1G109170 | calmodulin like 37 | 26.47 | AT5G42380.1 |
| 145 | HORVU2Hr1G112120 | Anthranilate N-benzoyltransferase protein 1 | 26.41 | AT5G48930.1 |
| 146 | HORVU1Hr1G063390 | AAA-ATPase 1 | 26.33 | AT5G40010.1 |
| 147 | HORVU2Hr1G012140 | Leucine-rich repeat receptor-like protein kinase family protein | 26.30 | AT1G74190.1 |
| 148 | HORVU1Hr1G092650 | UDP-Glycosyltransferase superfamily protein | 26.23 | AT3G16520.3 |
| 149 | HORVU6Hr1G077930 | UDP-Glycosyltransferase superfamily protein | 26.08 | AT1G22380.1 |
| 150 | HORVU5Hr1G106850 | ABC transporter G family member 48 | 26.04 | AT1G66950.1 |
| 151 | HORVU7Hr1G056250 | undescribed protein | 25.94 | AT4G28460.1 |
| 152 | HORVU3Hr1G006450 | Xylanase inhibitor | 25.82 | AT1G03230.1 |
| 153 | HORVU2Hr1G102710 | Protein NRT1/ PTR FAMILY 8.3 | 25.72 | AT2G02040.1 |
| 154 | HORVU1Hr1G001220 | receptor kinase 3 | 25.62 | AT4G23180.3 |
| 155 | HORVU1Hr1G075620 | ABC transporter ATP-binding protein NatA | 25.45 | AT3G47780.1 |
| 156 | HORVU2Hr1G098770 | undescribed protein | 25.44 | AT5G03130.1 |
| 157 | HORVU1Hr1G010120 | L-tyrosine decarboxylase | 25.25 | AT2G20340.1 |
| 158 | HORVU5Hr1G089840 | receptor kinase 3 | 25.18 | AT1G21230.1 |
| 159 | HORVU6Hr1G069710 | receptor kinase 2 | 25.15 | AT4G21390.1 |
| 160 | HORVU1Hr1G089590 | undescribed protein | 25.11 | AT1G67120.1 |
| 161 | HORVU2Hr1G122610 | Blue copper protein | 25.02 | AT2G31050.1 |

| | | | | |
|---|---|---|---|---|
| 162 | HORVU7Hr1G035730 | EF hand family protein | 24.98 | AT3G50360.1 |
| 163 | HORVU3Hr1G095730 | Glutathione S-transferase family protein | 24.94 | AT1G10360.1 |
| 164 | HORVU1Hr1G072690 | Aldose reductase A | 24.67 | AT2G37790.1 |
| 165 | HORVU2Hr1G004550 | Cytochrome P450 superfamily protein | 24.52 | AT1G64940.1 |
| 166 | HORVU4Hr1G087630 | Cytochrome P450 superfamily protein | 24.52 | AT2G30490.1 |
| 167 | HORVU7Hr1G033870 | zinc-finger protein 3 | 24.38 | AT1G27730.1 |
| 168 | HORVU7Hr1G009680 | Leucine-rich receptor-like protein kinase family protein | 24.35 | AT3G47570.1 |
| 169 | HORVU1Hr1G061140 | Pyrophosphate-energized proton pump 1 | 24.22 | AT1G15690.1 |
| 170 | HORVU7Hr1G043530 | serpin 2 | 24.11 | AT1G47710.1 |
| 171 | HORVU5Hr1G057800 | L-ascorbate oxidase | 23.85 | AT4G39830.4 |
| 172 | HORVU1Hr1G014450 | undescribed protein | 23.83 | AT5G23430.1 |
| 173 | HORVU2Hr1G114450 | L-tyrosine decarboxylase | 23.78 | AT2G20340.1 |
| 174 | HORVU2Hr1G112060 | undescribed protein | 23.71 | AT1G49560.1 |
| 175 | HORVU6Hr1G089590 | U-box domain-containing protein | 23.68 | AT2G35930.1 |
| 176 | HORVU3Hr1G018740 | sugar transporter 1 | 23.59 | AT5G61520.1 |
| 177 | HORVU3Hr1G087020 | alpha/beta-Hydrolases superfamily protein | 23.57 | AT2G39420.1 |
| 178 | HORVU6Hr1G073510 | Potassium transporter family protein | 23.45 | AT4G13420.1 |
| 179 | HORVU5Hr1G083340 | purple acid phosphatase 15 | 23.41 | AT3G07130.1 |
| 180 | HORVU6Hr1G001840 | ubiquitin-protein ligase 4 | 23.28 | AT4G38600.3 |
| 181 | HORVU2Hr1G040130 | Internal alternative NAD(P)H-ubiquinone oxidoreductase A1, mitochondrial | 23.26 | AT1G07180.1 |
| 182 | HORVU4Hr1G064070 | U-box domain-containing protein 21 | 23.25 | AT5G37490.1 |
| 183 | HORVU7Hr1G110840 | Nodulin-like / Major Facilitator Superfamily protein | 23.25 | AT2G39210.1 |
| 184 | HORVU1Hr1G015090 | tyrosine aminotransferase 3 | 23.16 | AT5G53970.1 |
| 185 | HORVU7Hr1G107360 | SAUR-like auxin-responsive protein family | 23.07 | AT1G56150.1 |
| 186 | HORVU7Hr1G107640 | 0 | 23.05 | AT1G14600.1 |
| 187 | HORVU7Hr1G118770 | Disease resistance-responsive (dirigent-like protein) family protein | 22.98 | AT5G42500.1 |
| 188 | HORVU2Hr1G067710 | undescribed protein | 22.82 | AT4G34480.1 |
| 189 | HORVU6Hr1G070130 | undescribed protein | 22.80 | AT2G43420.1 |
| 190 | HORVU7Hr1G073340 | early nodulin-like protein 15 | 22.73 | AT2G32300.2 |
| 191 | HORVU5Hr1G078630 | respiratory burst oxidase homologue D | 22.50 | AT5G47910.1 |
| 192 | HORVU7Hr1G091860 | L-tyrosine decarboxylase | 22.47 | AT2G20340.1 |
| 193 | HORVU6Hr1G038470 | unknown function | 22.42 | AT2G25735.1 |
| 194 | HORVU2Hr1G111780 | receptor-like protein kinase 4 | 22.25 | AT4G32300.1 |
| 195 | HORVU3Hr1G086840 | C2H2-like zinc finger protein | 22.11 | AT2G28710.1 |
| 196 | HORVU5Hr1G061580 | Secretory carrier-associated membrane protein 4 | 22.01 | AT2G20840.1 |

| | | | | |
|---|---|---|---|---|
| 197 | HORVU3Hr1G094370 | Chromosome 3B, genomic scaffold, cultivar Chinese Spring | 21.93 | AT4G19975.1 |
| 198 | HORVU2Hr1G101150 | xyloglucan endotransglucosylase/hydrolase 13 | 21.88 | AT5G57540.1 |
| 199 | HORVU2Hr1G044340 | Peroxidase superfamily protein | 21.87 | AT5G05340.1 |
| 200 | HORVU1Hr1G088300 | WRKY DNA-binding protein 71 | 21.83 | AT1G29860.1 |
| 201 | HORVU3Hr1G011870 | Cytochrome P450 superfamily protein | 21.73 | AT5G57260.1 |
| 202 | HORVU3Hr1G020140 | WRKY DNA-binding protein 50 | 21.63 | AT5G26170.1 |
| 203 | HORVU2Hr1G088660 | Caleosin-related family protein | 21.62 | AT4G26740.1 |
| 204 | HORVU2Hr1G109440 | Cytochrome P450 superfamily protein | 21.55 | AT4G36220.1 |
| 205 | HORVU3Hr1G006130 | GDSL esterase/lipase | 21.45 | AT3G26430.1 |
| 206 | HORVU4Hr1G079620 | Cytochrome P450 superfamily protein | 21.44 | AT2G26170.1 |
| 207 | HORVU1Hr1G002590 | receptor-like protein kinase 4 | 21.23 | AT1G66920.2 |
| 208 | HORVU5Hr1G047920 | Two-component response regulator ORR21 | 21.22 | AT2G38300.1 |
| 209 | HORVU7Hr1G081700 | lipid phosphate phosphatase 3 | 21.00 | AT3G02600.9 |
| 210 | HORVU5Hr1G111040 | UDP-Glycosyltransferase superfamily protein | 20.73 | AT5G49690.1 |
| 211 | HORVU4Hr1G078320 | undescribed protein | 20.67 | AT1G51850.2 |
| 212 | HORVU7Hr1G040740 | Cysteine-rich venom protein | 20.66 | AT5G57625.1 |
| 213 | HORVU3Hr1G111620 | undescribed protein | 20.66 | AT5G65090.3 |
| 214 | HORVU6Hr1G079700 | nitrate reductase 1 | 20.62 | AT1G77760.1 |
| 215 | HORVU7Hr1G010850 | L-tyrosine decarboxylase | 20.60 | AT2G20340.1 |
| 216 | HORVU7Hr1G107370 | SAUR-like auxin-responsive protein family | 20.56 | AT1G56150.1 |
| 217 | HORVU5Hr1G115750 | PLANT CADMIUM RESISTANCE 2 | 20.54 | AT1G14870.1 |
| 218 | HORVU7Hr1G042300 | Basic blue protein | 20.48 | AT2G02850.1 |
| 219 | HORVU5Hr1G077920 | beta glucosidase 11 | 20.38 | AT1G02850.2 |
| 220 | HORVU3Hr1G099550 | Eukaryotic aspartyl protease family protein | 20.27 | AT1G03230.1 |
| 221 | HORVU2Hr1G003230 | Chromosome 3B, genomic scaffold, cultivar Chinese Spring | 20.09 | AT3G14470.2 |
| 222 | HORVU4Hr1G053150 | Plant protein of unknown function (DUF247) | 19.84 | AT3G50170.1 |
| 223 | HORVU2Hr1G044360 | Peroxidase superfamily protein | 19.82 | AT5G05340.1 |
| 224 | HORVU5Hr1G094710 | Disease resistance-responsive (dirigent-like protein) family protein | 19.78 | AT1G58170.1 |
| 225 | HORVU3Hr1G002900 | receptor kinase 2 | 19.69 | AT1G70520.1 |
| 226 | HORVU4Hr1G081590 | receptor-like protein kinase 4 | 19.53 | AT5G10530.1 |
| 227 | HORVU4Hr1G010720 | undescribed protein | 19.48 | AT1G72060.1 |
| 228 | HORVU7Hr1G019390 | receptor kinase 2 | 19.34 | AT2G37710.1 |
| 229 | HORVU1Hr1G065650 | undescribed protein | 19.28 | AT2G05635.2 |
| 230 | HORVU1Hr1G065150 | undescribed protein | 19.06 | AT2G43550.1 |

| | | | | |
|---|---|---|---|---|
| 231 | HORVU1Hr1G058520 | beta glucosidase 11 | 18.91 | AT1G02850.5 |
| 232 | HORVU3Hr1G050810 | Chromosome 3B, genomic scaffold, cultivar Chinese Spring | 18.90 | AT3G21305.1 |
| 233 | HORVU3Hr1G077130 | receptor kinase 2 | 18.80 | AT4G21390.1 |
| 234 | HORVU0Hr1G019300 | Glutathione S-transferase family protein | 18.72 | AT3G09270.1 |
| 235 | HORVU0Hr1G021480 | Arabidopsis thaliana protein of unknown function (DUF821) | 18.70 | AT5G23850.1 |
| 236 | HORVU4Hr1G074080 | undescribed protein | 18.67 | AT2G30620.2 |
| 237 | HORVU4Hr1G087840 | Protein of unknown function (DUF1218) | 18.64 | AT5G17210.1 |
| 238 | HORVU3Hr1G006440 | Xylanase inhibitor | 18.55 | AT1G03230.1 |
| 239 | HORVU1Hr1G049260 | Glutathione S-transferase family protein | 18.30 | AT1G10370.1 |
| 240 | HORVU2Hr1G067700 | undescribed protein | 18.26 | AT2G19820.1 |
| 241 | HORVU0Hr1G000840 | undescribed protein | 18.13 | AT5G47910.1 |
| 242 | HORVU5Hr1G099470 | NAC domain protein, | 17.89 | AT2G43000.1 |
| 243 | HORVU5Hr1G000910 | Serine/threonine-protein kinase | 17.79 | AT4G32300.1 |
| 244 | HORVU3Hr1G096280 | Sec14p-like phosphatidylinositol transfer family protein | 17.77 | AT1G01630.1 |
| 245 | HORVU3Hr1G099530 | UDP-Glycosyltransferase superfamily protein | 17.75 | AT3G11340.1 |
| 246 | HORVU5Hr1G109190 | germin-like protein 4 | 17.72 | AT1G18980.1 |
| 247 | HORVU2Hr1G113120 | Protein kinase superfamily protein | 17.71 | AT5G59700.1 |
| 248 | HORVU4Hr1G068530 | unknown protein; Has 65 Blast hits to 65 proteins in 11 species: Archae - 0; Bacteria - 0; Metazoa - 0; Fungi - 0; Plants - 65; Viruses - 0; Other Eukaryotes - 0 (source: NCBI BLink). | 17.67 | AT3G57450.1 |
| 249 | HORVU6Hr1G067980 | Protein NRT1/ PTR FAMILY 8.3 | 17.58 | AT2G02040.1 |
| 250 | HORVU3Hr1G026330 | Subtilisin-like protease | 17.50 | AT4G10540.1 |
| 251 | HORVU6Hr1G094790 | receptor-like protein kinase 1 | 17.41 | AT3G21630.1 |
| 252 | HORVU3Hr1G006140 | GDSL esterase/lipase | 17.38 | AT3G26430.1 |
| 253 | HORVU7Hr1G092510 | ABC transporter G family member 44 | 17.01 | AT1G15520.1 |
| 254 | HORVU6Hr1G073990 | early nodulin-like protein 19 | 16.99 | AT2G32300.2 |
| 255 | HORVU7Hr1G057650 | RING/U-box superfamily protein | 16.93 | AT2G17730.1 |
| 256 | HORVU7Hr1G046180 | undescribed protein | 16.87 | AT2G29580.1 |
| 257 | HORVU6Hr1G060500 | membrane steroid binding protein 1 | 16.82 | AT3G48890.1 |
| 258 | HORVU3Hr1G023230 | UDP-Glycosyltransferase superfamily protein | 16.78 | AT3G53150.1 |
| 259 | HORVU7Hr1G106760 | unknown function | 16.73 | AT5G16610.2 |
| 260 | HORVU1Hr1G075640 | ABC transporter ATP-binding protein NatA | 16.58 | AT3G47730.1 |
| 261 | HORVU1Hr1G090860 | unknown function | 16.56 | AT3G62650.2 |
| 262 | HORVU2Hr1G034670 | Auxin-responsive GH3 family protein | 16.52 | AT2G14960.1 |
| 263 | HORVU1Hr1G008360 | laccase 7 | 16.49 | AT3G09220.1 |

| | | | | |
|---|---|---|---|---|
| 264 | HORVU2Hr1G079580 | unknown function | 16.45 | AT2G14095.1 |
| 265 | HORVU2Hr1G111760 | receptor-like protein kinase 4 | 16.42 | AT4G32300.1 |
| 266 | HORVU2Hr1G096380 | Branched-chain-amino-acid aminotransferase 6 | 16.41 | AT5G65780.1 |
| 267 | HORVU1Hr1G062030 | Chitinase family protein | 16.38 | AT3G12500.1 |
| 268 | HORVU5Hr1G112610 | rRNA N-glycosidase | 16.24 | AT3G59490.3 |
| 269 | HORVU5Hr1G045640 | NAC domain protein, | 16.10 | AT5G53950.1 |
| 270 | HORVU7Hr1G086340 | undescribed protein | 16.04 | AT2G32210.1 |
| 271 | HORVU7Hr1G021260 | WRKY DNA-binding protein 61 | 15.77 | AT5G15130.1 |
| 272 | HORVU4Hr1G084810 | 2-oxoglutarate (2OG) and Fe(II)-dependent oxygenase superfamily protein | 15.76 | AT5G24530.1 |
| 273 | HORVU6Hr1G074370 | undescribed protein | 15.72 | AT1G44970.1 |
| 274 | HORVU5Hr1G034830 | WRKY family transcription factor | 15.69 | AT4G11070.2 |
| 275 | HORVU1Hr1G090460 | undescribed protein | 15.64 | AT1G78410.2 |
| 276 | HORVU3Hr1G065320 | ABC transporter B family member 4 | 15.59 | AT2G47000.7 |
| 277 | HORVU2Hr1G018430 | Peroxidase superfamily protein | 15.45 | AT5G05340.1 |
| 278 | HORVU4Hr1G056890 | LOB domain-containing protein 1 | 15.42 | AT1G07900.1 |
| 279 | HORVU7Hr1G108150 | Protein kinase superfamily protein | 15.41 | AT2G05940.1 |
| 280 | HORVU2Hr1G012980 | Cytochrome P450 superfamily protein | 15.39 | AT1G13110.1 |
| 281 | HORVU4Hr1G020230 | rRNA N-glycosidase | 15.37 | AT3G56810.1 |
| 282 | HORVU3Hr1G021500 | Transposable element protein, putative | 15.35 | AT1G62270.1 |
| 283 | HORVU3Hr1G083680 | Harpin induced gene 1 homolog | 15.34 | AT2G35980.1 |
| 284 | HORVU7Hr1G032740 | unknown function | 15.31 | AT3G16670.1 |
| 285 | HORVU2Hr1G063690 | MATE efflux family protein | 15.28 | AT3G23550.1 |
| 286 | HORVU1Hr1G085240 | Core-2/I-branching beta-1,6-N-acetylglucosaminyltransferase family protein | 15.27 | AT5G11730.1 |
| 287 | HORVU3Hr1G001950 | undescribed protein | 15.08 | AT4G39970.1 |
| 288 | HORVU3Hr1G002910 | 0 | 15.07 | AT1G56120.3 |
| 289 | HORVU3Hr1G024820 | Chromosome 3B, genomic scaffold, cultivar Chinese Spring | 15.05 | AT5G06570.4 |
| 290 | HORVU1Hr1G080680 | Cytochrome P450 superfamily protein | 15.04 | AT3G48520.1 |
| 291 | HORVU4Hr1G051450 | Phospholipid-transporting ATPase 1 | 14.96 | AT5G04930.1 |
| 292 | HORVU3Hr1G064470 | Chitinase family protein | 14.93 | AT3G12500.1 |
| 293 | HORVU2Hr1G013760 | MLO-like protein 1 | 14.91 | AT2G44110.2 |
| 294 | HORVU5Hr1G124810 | receptor kinase 1 | 14.87 | AT4G23230.1 |
| 295 | HORVU2Hr1G091190 | sugar transporter 9 | 14.75 | AT2G20780.1 |
| 296 | HORVU4Hr1G051290 | NAC domain protein, | 14.75 | AT1G69490.1 |
| 297 | HORVU3Hr1G090170 | tropinone reductase | 14.69 | AT5G06060.1 |
| 298 | HORVU3Hr1G086980 | alpha/beta-Hydrolases superfamily protein | 14.68 | AT2G39420.1 |
| 299 | HORVU3Hr1G011990 | germin-like protein 2 | 14.65 | AT5G39100.1 |

| | | | | |
|---|---|---|---|---|
| 300 | HORVU1Hr1G004980 | Glycosyltransferase family 61 protein | 14.47 | AT2G41640.1 |
| 301 | HORVU2Hr1G013170 | 2-oxoglutarate dehydrogenase, E1 component | 14.47 | AT3G55410.2 |
| 302 | HORVU4Hr1G056500 | 2-oxoglutarate (2OG) and Fe(II)-dependent oxygenase superfamily protein | 14.45 | AT3G11180.1 |
| 303 | HORVU3Hr1G057730 | unknown protein; Has 19 Blast hits to 19 proteins in 6 species: Archae - 0; Bacteria - 0; Metazoa - 0; Fungi - 0; Plants - 19; Viruses - 0; Other Eukaryotes - 0 (source: NCBI BLink). | 14.44 | AT2G01400.1 |
| 304 | HORVU3Hr1G014140 | NAC domain protein, | 14.43 | AT1G61110.1 |
| 305 | HORVU7Hr1G038330 | amino acid permease 1 | 14.29 | AT1G58360.1 |
| 306 | HORVU2Hr1G112580 | endonuclease 4 | 14.20 | AT4G21585.4 |
| 307 | HORVU3Hr1G022780 | Protein kinase family protein | 14.05 | AT5G02070.1 |
| 308 | HORVU3Hr1G024660 | Acetyl esterase | 13.93 | AT5G06570.4 |
| 309 | HORVU7Hr1G092800 | Tryptophan synthase beta chain 2 | 13.92 | AT5G38530.1 |
| 310 | HORVU3Hr1G028020 | GDSL esterase/lipase | 13.88 | AT1G28570.2 |
| 311 | HORVU2Hr1G103180 | L-lactate dehydrogenase | 13.71 | AT4G18360.4 |
| 312 | HORVU1Hr1G014460 | undescribed protein | 13.65 | #N/A |
| 313 | HORVU5Hr1G063270 | GRAS family transcription factor | 13.61 | AT1G07520.1 |
| 314 | HORVU7Hr1G080950 | WRKY DNA-binding protein 27 | 13.55 | AT4G01250.1 |
| 315 | HORVU5Hr1G046730 | undescribed protein | 13.51 | AT3G17700.2 |
| 316 | HORVU2Hr1G018510 | Peroxidase superfamily protein | 13.51 | AT5G05340.1 |
| 317 | HORVU4Hr1G074780 | Protein of unknown function (DUF1997) | 13.48 | AT5G39530.1 |
| 318 | HORVU6Hr1G069980 | receptor kinase 2 | 13.47 | AT5G06740.1 |
| 319 | HORVU1Hr1G037270 | Chitinase 2 | 13.42 | AT4G19810.2 |
| 320 | HORVU3Hr1G089900 | undescribed protein | 13.41 | AT3G32090.1 |
| 321 | HORVU7Hr1G084920 | Chitinase 1 | 13.40 | AT1G14430.1 |
| 322 | HORVU5Hr1G063430 | glutamate receptor 2.8 | 13.38 | AT2G29120.1 |
| 323 | HORVU2Hr1G067340 | exocyst subunit exo70 family protein H6 | 13.29 | AT1G07725.1 |
| 324 | HORVU1Hr1G067840 | Pectin lyase-like superfamily protein | 13.29 | AT1G02460.1 |
| 325 | HORVU3Hr1G018610 | receptor kinase 2 | 13.23 | AT3G53810.1 |
| 326 | HORVU4Hr1G005440 | germin-like protein 4 | 13.23 | AT1G18980.1 |
| 327 | HORVU7Hr1G075760 | L-gulonolactone oxidase 2 | 13.19 | AT2G46750.1 |
| 328 | HORVU0Hr1G016280 | undescribed protein | 13.07 | AT5G20360.6 |
| 329 | HORVU0Hr1G016310 | Endochitinase A | 13.05 | AT3G54420.1 |
| 330 | HORVU5Hr1G066250 | Vacuolar-processing enzyme | 13.05 | AT2G25940.1 |
| 331 | HORVU3Hr1G098340 | methyl esterase 1 | 13.03 | AT2G23620.1 |
| 332 | HORVU3Hr1G056100 | Cytochrome P450 superfamily protein | 13.01 | AT3G14680.1 |
| 333 | HORVU7Hr1G084610 | 3-ketoacyl-CoA synthase 11 | 12.95 | AT2G26640.1 |
| 334 | HORVU7Hr1G077970 | PLC-like phosphodiesterases superfamily protein | 12.90 | AT4G36945.1 |

| | | | | |
|---|---|---|---|---|
| 335 | HORVU3Hr1G113620 | Wound-induced protein | 12.89 | AT3G04720.1 |
| 336 | HORVU4Hr1G084930 | TVP38/TMEM64 family membrane protein slr0305 | 12.84 | AT2G02370.3 |
| 337 | HORVU2Hr1G075970 | Peroxidase superfamily protein | 12.79 | AT4G08390.3 |
| 338 | HORVU7Hr1G114660 | Indole-3-glycerol phosphate synthase | 12.75 | AT2G04400.1 |
| 339 | HORVU3Hr1G018920 | Chromosome 3B, genomic scaffold, cultivar Chinese Spring | 12.65 | AT1G71010.1 |
| 340 | HORVU1Hr1G019500 | sucrose-phosphatase 1 | 12.62 | AT2G35840.4 |
| 341 | HORVU2Hr1G101980 | Ubiquinol oxidase | 12.61 | AT3G22370.1 |
| 342 | HORVU3Hr1G053350 | ABC transporter G family member 36 | 12.60 | AT1G15520.1 |
| 343 | HORVU7Hr1G101310 | Calcium-dependent lipid-binding (CaLB domain) family protein | 12.53 | AT5G04220.2 |
| 344 | HORVU5Hr1G043460 | calmodulin like 23 | 12.52 | AT3G10190.1 |
| 345 | HORVU3Hr1G080100 | NAC domain containing protein 102 | 12.52 | AT5G63790.1 |
| 346 | HORVU7Hr1G024350 | Long-chain-fatty-acid--CoA ligase 1 | 12.51 | AT4G23850.1 |
| 347 | HORVU1Hr1G009130 | unknown function | 12.46 | AT2G18690.1 |
| 348 | HORVU5Hr1G052150 | Indole-3-glycerol phosphate synthase | 12.45 | AT5G48220.1 |
| 349 | HORVU5Hr1G000940 | receptor kinase 2 | 12.43 | AT5G10530.1 |
| 350 | HORVU2Hr1G085260 | Protein NRT1/ PTR FAMILY 4.3 | 12.42 | AT1G59740.1 |
| 351 | HORVU2Hr1G028780 | Chymotrypsin inhibitor | 12.37 | AT2G38900.2 |
| 352 | HORVU5Hr1G041590 | Leucine-rich repeat protein kinase family protein | 12.37 | AT4G29180.1 |
| 353 | HORVU1Hr1G065660 | undescribed protein | 12.28 | AT4G19140.1 |
| 354 | HORVU3Hr1G104990 | undescribed protein | 12.24 | #N/A |
| 355 | HORVU1Hr1G051580 | PLAC8 family protein | 12.22 | AT1G63830.3 |
| 356 | HORVU4Hr1G076750 | methyl esterase 1 | 12.21 | AT2G23620.1 |
| 357 | HORVU1Hr1G019770 | Protein kinase superfamily protein | 12.16 | AT2G05940.1 |
| 358 | HORVU1Hr1G015210 | Leucine-rich repeat receptor-like protein kinase family protein | 12.15 | AT4G08850.1 |
| 359 | HORVU6Hr1G027640 | SAUR25-auxin-responsive SAUR family member | 12.12 | AT1G56150.1 |
| 360 | HORVU4Hr1G054770 | COBRA-like protein 7 | 12.08 | AT4G16120.1 |
| 361 | HORVU1Hr1G019750 | Laccase-19 | 12.08 | AT3G09220.1 |
| 362 | HORVU2Hr1G090170 | UDP-Glycosyltransferase superfamily protein | 11.88 | AT2G36790.1 |
| 363 | HORVU0Hr1G005420 | Ubiquinol oxidase | 11.88 | AT3G22360.1 |
| 364 | HORVU4Hr1G028780 | myb domain protein 112 | 11.81 | AT5G49620.1 |
| 365 | HORVU3Hr1G068700 | Ribose-phosphate pyrophosphokinase 4 | 11.79 | AT1G10700.1 |
| 366 | HORVU3Hr1G065390 | Lipid A export ATP-binding/permease protein MsbA | 11.79 | AT2G47000.6 |
| 367 | HORVU2Hr1G018480 | Peroxidase superfamily protein | 11.78 | AT5G05340.1 |
| 368 | HORVU2Hr1G033610 | dihydroflavonol 4-reductase | 11.76 | AT4G27250.1 |

| | | | | |
|---|---|---|---|---|
| 369 | HORVU5Hr1G062090 | Aldehyde dehydrogenase family 7 member A1 | 11.75 | AT1G54100.2 |
| 370 | HORVU5Hr1G087240 | receptor kinase 3 | 11.73 | AT4G21390.2 |
| 371 | HORVU5Hr1G060620 | Centromere protein V | 11.71 | AT5G16940.2 |
| 372 | HORVU7Hr1G013170 | Flavin-containing monooxygenase family protein | 11.69 | AT1G21430.1 |
| 373 | HORVU5Hr1G095130 | Beta-glucosidase C | 11.67 | AT5G04885.2 |
| 374 | HORVU3Hr1G028620 | Cytochrome P450 61 | 11.65 | AT2G34500.1 |
| 375 | HORVU4Hr1G081430 | NAC domain containing protein 36 | 11.62 | AT2G17040.1 |
| 376 | HORVU6Hr1G053700 | undescribed protein | 11.61 | AT3G27473.1 |
| 377 | HORVU1Hr1G038780 | unknown protein; BEST Arabidopsis thaliana protein match is: unknown protein . | 11.56 | AT1G07040.1 |
| 378 | HORVU4Hr1G063660 | UDP-Glycosyltransferase superfamily protein | 11.56 | AT3G02100.1 |
| 379 | HORVU3Hr1G006760 | Chromosome 3B, genomic scaffold, cultivar Chinese Spring | 11.53 | AT3G14460.1 |
| 380 | HORVU7Hr1G038370 | Beta-1,3-galactosyltransferase 15 | 11.52 | AT1G26810.2 |
| 381 | HORVU4Hr1G004000 | Major facilitator superfamily protein | 11.50 | AT2G39210.1 |
| 382 | HORVU6Hr1G067920 | Protein NRT1/ PTR FAMILY 8.1 | 11.46 | AT1G62200.1 |
| 383 | HORVU3Hr1G117910 | PLAC8 family protein | 11.44 | AT5G05350.1 |
| 384 | HORVU1Hr1G081410 | Chaperone protein DnaJ | 11.42 | AT2G42750.1 |
| 385 | HORVU5Hr1G094700 | Disease resistance-responsive (dirigent-like protein) family protein | 11.39 | AT1G58170.1 |
| 386 | HORVU7Hr1G107330 | SAUR-like auxin-responsive protein family | 11.32 | AT1G56150.1 |
| 387 | HORVU7Hr1G106600 | patatin-like protein 6 | 11.17 | AT3G54950.1 |
| 388 | HORVU3Hr1G077950 | Subtilisin-like protease | 11.12 | AT4G10540.1 |
| 389 | HORVU4Hr1G085820 | UDP-galactose transporter 2 | 11.06 | AT4G23010.1 |
| 390 | HORVU2Hr1G125250 | receptor kinase 2 | 11.06 | AT4G23180.3 |
| 391 | HORVU2Hr1G007570 | S-adenosylmethionine-dependent methyltransferase | 11.06 | AT2G41380.1 |
| 392 | HORVU6Hr1G074690 | receptor-like protein kinase 4 | 11.05 | AT1G34300.1 |
| 393 | HORVU3Hr1G002000 | germin-like protein 9 | 10.99 | AT5G38960.1 |
| 394 | HORVU0Hr1G030830 | WRKY family transcription factor | 10.97 | AT3G56400.1 |
| 395 | HORVU4Hr1G005380 | germin-like protein 4 | 10.95 | AT1G18980.1 |
| 396 | HORVU5Hr1G068320 | Heat stress transcription factor B-1 | 10.87 | AT4G36990.1 |
| 397 | HORVU2Hr1G100360 | SAUR-like auxin-responsive protein family | 10.85 | AT2G24400.1 |
| 398 | HORVU5Hr1G115870 | PLANT CADMIUM RESISTANCE 2 | 10.84 | AT1G14870.1 |
| 399 | HORVU1Hr1G073820 | Glucan endo-1,3-beta-glucosidase 14 | 10.77 | AT2G27500.2 |
| 400 | HORVU2Hr1G114590 | Synaptotagmin-5 | 10.75 | AT1G05500.1 |
| 401 | HORVU2Hr1G101920 | Ubiquinol oxidase | 10.70 | AT3G22370.1 |

| | | | | |
|---|---|---|---|---|
| 402 | HORVU5Hr1G104580 | UDP-Glycosyltransferase superfamily protein | 10.69 | AT2G15480.2 |
| 403 | HORVU6Hr1G031640 | unknown function | 10.66 | AT4G39720.4 |
| 404 | HORVU2Hr1G008220 | G-type lectin S-receptor-like serine/threonine-protein kinase | 10.65 | AT4G21380.2 |
| 405 | HORVU2Hr1G006100 | receptor kinase 2 | 10.64 | AT3G53810.1 |
| 406 | HORVU2Hr1G110230 | Protein kinase superfamily protein | 10.63 | AT1G78290.3 |
| 407 | HORVU5Hr1G118750 | Zinc finger protein 668 | 10.62 | AT1G34370.2 |
| 408 | HORVU4Hr1G017390 | Eukaryotic aspartyl protease family protein | 10.57 | AT4G33490.2 |
| 409 | HORVU7Hr1G086250 | undescribed protein | 10.50 | AT3G24460.2 |
| 410 | HORVU5Hr1G095580 | Cathepsin B-like cysteine proteinase | 10.49 | AT5G45890.1 |
| 411 | HORVU4Hr1G071330 | Cytochrome b561 and DOMON domain-containing protein | 10.48 | AT3G59070.1 |
| 412 | HORVU2Hr1G120660 | receptor kinase 2 | 10.45 | AT3G53810.1 |
| 413 | HORVU7Hr1G029460 | undescribed protein | 10.44 | AT2G17780.4 |
| 414 | HORVU1Hr1G067870 | Cysteine proteinase inhibitor | 10.36 | AT3G12490.1 |
| 415 | HORVU1Hr1G027700 | WRKY DNA-binding protein 50 | 10.32 | AT5G26170.1 |
| 416 | HORVU4Hr1G001040 | Leucine-rich repeat receptor-like protein kinase family protein | 10.30 | AT2G34930.1 |
| 417 | HORVU3Hr1G113100 | Beta-fructofuranosidase, insoluble isoenzyme 4 | 10.26 | AT3G13790.2 |
| 418 | HORVU3Hr1G067510 | WRKY DNA-binding protein 50 | 10.26 | AT5G26170.1 |
| 419 | HORVU3Hr1G072210 | undescribed protein | 10.26 | AT3G63150.1 |
| 420 | HORVU2Hr1G100090 | Solute carrier family 22 member 6-A | 10.23 | AT3G13050.4 |
| 421 | HORVU7Hr1G003630 | receptor kinase 2 | 10.19 | AT4G21390.1 |
| 422 | HORVU3Hr1G111550 | undescribed protein | 10.18 | AT5G43210.1 |
| 423 | HORVU1Hr1G089310 | B12D protein | 10.17 | AT3G48140.1 |
| 424 | HORVU0Hr1G005300 | 2-oxoglutarate (2OG) and Fe(II)-dependent oxygenase superfamily protein | 10.12 | AT5G24530.1 |
| 425 | HORVU3Hr1G117470 | 2-oxoglutarate (2OG) and Fe(II)-dependent oxygenase superfamily protein | 10.08 | AT1G35190.2 |
| 426 | HORVU2Hr1G093020 | undescribed protein | 9.99 | AT5G08780.1 |
| 427 | HORVU5Hr1G072740 | undescribed protein | 9.96 | #N/A |
| 428 | HORVU5Hr1G096210 | UDP-Glycosyltransferase superfamily protein | 9.94 | AT3G02100.1 |
| 429 | HORVU7Hr1G028710 | 2-oxoglutarate (2OG) and Fe(II)-dependent oxygenase superfamily protein | 9.94 | AT5G20400.1 |
| 430 | HORVU6Hr1G008640 | catalase 1 | 9.94 | AT1G20630.1 |
| 431 | HORVU2Hr1G028330 | EF hand family protein | 9.91 | AT5G04990.1 |

| | | | | |
|---|---|---|---|---|
| 432 | HORVU6Hr1G062380 | Ethylene-responsive transcription factor 1 | 9.83 | AT4G17500.1 |
| 433 | HORVU5Hr1G052760 | Heavy metal transport/detoxification superfamily protein | 9.82 | AT3G20180.3 |
| 434 | HORVU2Hr1G016720 | Protein kinase superfamily protein | 9.81 | AT1G26970.1 |
| 435 | HORVU4Hr1G061120 | Anthranilate synthase component 1 | 9.79 | AT5G05730.1 |
| 436 | HORVU0Hr1G006150 | receptor kinase 1 | 9.77 | AT5G38240.2 |
| 437 | HORVU6Hr1G083050 | lysine-ketoglutarate reductase/saccharopine dehydrogenase bifunctional enzyme | 9.69 | AT4G33150.3 |
| 438 | HORVU1Hr1G041720 | Nucleotide/sugar transporter family protein | 9.64 | AT2G28315.1 |
| 439 | HORVU7Hr1G089160 | FAD-binding Berberine family protein | 9.61 | AT5G44400.1 |
| 440 | HORVU1Hr1G052430 | Chitinase family protein | 9.58 | AT3G12500.1 |
| 441 | HORVU1Hr1G039720 | Protein NRT1/ PTR FAMILY 5.2 | 9.56 | AT5G46050.1 |
| 442 | HORVU3Hr1G013380 | Leucine-rich repeat protein kinase family protein | 9.56 | AT4G29990.2 |
| 443 | HORVU2Hr1G010560 | Thioredoxin superfamily protein | 9.56 | AT1G20225.1 |
| 444 | HORVU7Hr1G116650 | Glutathione S-transferase family protein | 9.56 | AT3G62760.1 |
| 445 | HORVU2Hr1G076470 | U-box domain-containing protein | 9.55 | AT5G64660.1 |
| 446 | HORVU3Hr1G117480 | 2-oxoglutarate (2OG) and Fe(II)-dependent oxygenase superfamily protein | 9.54 | AT4G16765.6 |
| 447 | HORVU3Hr1G108360 | Orotidine 5'-phosphate decarboxylase | 9.49 | AT3G54470.1 |
| 448 | HORVU2Hr1G028430 | ribonuclease 1 | 9.42 | AT2G02990.1 |
| 449 | HORVU1Hr1G074310 | receptor kinase 1 | 9.34 | AT4G23180.1 |
| 450 | HORVU4Hr1G076520 | Protein kinase superfamily protein | 9.34 | AT2G05940.1 |
| 451 | HORVU7Hr1G101740 | Calcium-binding EF-hand family protein | 9.31 | AT4G27280.1 |
| 452 | HORVU3Hr1G018690 | receptor kinase 2 | 9.28 | AT2G37710.1 |
| 453 | HORVU1Hr1G081300 | Heat stress transcription factor A-4d | 9.21 | AT4G18880.1 |
| 454 | HORVU1Hr1G054290 | Subtilisin-like protease SBT3.17 | 9.21 | AT1G71950.1 |
| 455 | HORVU2Hr1G004720 | UDP-Glycosyltransferase superfamily protein | 9.18 | AT1G22400.1 |
| 456 | HORVU5Hr1G063480 | glutamate receptor 2.8 | 9.16 | AT2G29120.1 |
| 457 | HORVU5Hr1G122410 | undescribed protein | 9.14 | AT1G69850.1 |
| 458 | HORVU3Hr1G093100 | Bifunctional inhibitor/lipid-transfer protein/seed storage 2S albumin superfamily protein | 9.11 | AT5G55410.1 |
| 459 | HORVU2Hr1G089020 | Disease resistance protein | 9.09 | AT4G26090.1 |
| 460 | HORVU7Hr1G089530 | RING/U-box superfamily protein | 9.08 | AT1G53820.1 |
| 461 | HORVU3Hr1G110580 | undescribed protein | 9.03 | AT4G04750.3 |
| 462 | HORVU2Hr1G094640 | unknown function | 9.01 | AT5G54080.2 |
| 463 | HORVU3Hr1G071470 | ABC transporter G family member 32 | 8.95 | AT1G15520.1 |
| 464 | HORVU3Hr1G105560 | Glucan endo-1,3-beta-glucosidase GI | 8.94 | AT4G16260.1 |

| | | | | |
|---|---|---|---|---|
| 465 | HORVU6Hr1G084080 | MATE efflux family protein | 8.93 | AT3G23560.1 |
| 466 | HORVU5Hr1G104620 | Type IV inositol polyphosphate 5-phosphatase 7 | 8.92 | AT2G32010.4 |
| 467 | HORVU7Hr1G007520 | Leucine-rich repeat receptor-like protein kinase family protein | 8.92 | AT5G44700.1 |
| 468 | HORVU4Hr1G052320 | unknown function | 8.92 | AT3G56710.2 |
| 469 | HORVU5Hr1G045850 | Glutathione S-transferase family protein | 8.91 | AT2G29420.1 |
| 470 | HORVU6Hr1G077120 | U-box domain-containing protein 21 | 8.90 | AT5G37490.1 |
| 471 | HORVU2Hr1G045200 | Glutathione S-transferase family protein | 8.79 | AT1G10370.1 |
| 472 | HORVU4Hr1G067450 | sugar transporter protein 7 | 8.75 | AT5G26340.1 |
| 473 | HORVU2Hr1G124930 | Peroxidase superfamily protein | 8.74 | AT1G71695.1 |
| 474 | HORVU7Hr1G121290 | flavin-dependent monooxygenase 1 | 8.74 | AT1G19250.1 |
| 475 | HORVU5Hr1G102900 | Cysteine-rich receptor-like protein kinase 25 | 8.74 | AT5G48540.1 |
| 476 | HORVU7Hr1G002740 | laccase 7 | 8.73 | AT3G09220.1 |
| 477 | HORVU2Hr1G123480 | CCR4-NOT transcription complex subunit 8 | 8.66 | AT5G22250.1 |
| 478 | HORVU3Hr1G070700 | Leucine-rich receptor-like protein kinase family protein | 8.63 | AT5G25930.1 |
| 479 | HORVU7Hr1G091250 | alpha-amylase-like | 8.55 | AT4G25000.1 |
| 480 | HORVU2Hr1G091410 | unknown function | 8.52 | AT3G14460.1 |
| 481 | HORVU1Hr1G063740 | NAC domain protein, | 8.46 | AT1G01720.1 |
| 482 | HORVU2Hr1G075220 | unknown function | 8.39 | AT1G45688.2 |
| 483 | HORVU1Hr1G070250 | WRKY DNA-binding protein 33 | 8.39 | AT2G38470.1 |
| 484 | HORVU2Hr1G085280 | Chitinase | 8.39 | AT3G54420.1 |
| 485 | HORVU5Hr1G010130 | undescribed protein | 8.35 | AT4G16120.1 |
| 486 | HORVU3Hr1G007290 | Calmodulin-binding protein | 8.31 | AT5G57580.1 |
| 487 | HORVU5Hr1G112210 | Transmembrane amino acid transporter family protein | 8.31 | AT3G11900.2 |
| 488 | HORVU7Hr1G028370 | Pheophorbide a oxygenase, chloroplastic | 8.30 | AT4G25650.1 |
| 489 | HORVU6Hr1G068990 | Adenine nucleotide alpha hydrolases-like superfamily protein | 8.26 | AT2G21620.1 |
| 490 | HORVU1Hr1G021150 | Glutathione S-transferase family protein | 8.26 | AT3G62760.1 |
| 491 | HORVU4Hr1G084340 | Inter-alpha-trypsin inhibitor heavy chain H1 | 8.22 | AT5G60710.1 |
| 492 | HORVU6Hr1G005900 | alpha/beta-Hydrolases superfamily protein | 8.22 | AT1G47480.1 |
| 493 | HORVU7Hr1G086260 | undescribed protein | 8.21 | AT3G46990.1 |
| 494 | HORVU3Hr1G001740 | Chromosome 3B, genomic scaffold, cultivar Chinese Spring | 8.14 | AT5G22050.1 |
| 495 | HORVU3Hr1G085890 | ABC transporter C family member 3 | 8.04 | AT3G13080.4 |
| 496 | HORVU4Hr1G053130 | calmodulin 3 | 8.03 | AT5G49480.1 |
| 497 | HORVU1Hr1G071380 | unknown function | 7.98 | AT1G74540.1 |

| | | | | |
|---|---|---|---|---|
| 498 | HORVU4Hr1G071360 | Cytochrome b561 and DOMON domain-containing protein | 7.89 | AT3G25290.2 |
| 499 | HORVU5Hr1G072730 | undescribed protein | 7.88 | #N/A |
| 500 | HORVU1Hr1G071670 | 2-oxoglutarate (2OG) and Fe(II)-dependent oxygenase superfamily protein | 7.84 | AT5G24530.1 |
| 501 | HORVU5Hr1G024400 | Calmodulin-binding protein | 7.84 | AT2G18750.3 |
| 502 | HORVU5Hr1G114440 | unknown function | 7.83 | AT4G35920.5 |
| 503 | HORVU1Hr1G021140 | Glutathione S-transferase family protein | 7.81 | AT3G62760.1 |
| 504 | HORVU4Hr1G072480 | Blue copper protein | 7.77 | AT2G32300.1 |
| 505 | HORVU7Hr1G042120 | Protein NRT1/ PTR FAMILY 3.1 | 7.75 | AT1G68570.1 |
| 506 | HORVU3Hr1G028260 | RING/U-box superfamily protein | 7.75 | AT2G27940.1 |
| 507 | HORVU2Hr1G044640 | receptor kinase 1 | 7.69 | AT4G23160.1 |
| 508 | HORVU3Hr1G063430 | MATE efflux family protein | 7.67 | AT1G71140.1 |
| 509 | HORVU3Hr1G092680 | Aldehyde dehydrogenase family 2 member C4 | 7.67 | AT3G24503.1 |
| 510 | HORVU2Hr1G083570 | Cytochrome P450 superfamily protein | 7.65 | AT3G26330.1 |
| 511 | HORVU7Hr1G031360 | glutamate receptor 3.3 | 7.64 | AT5G11210.1<br>1 |
| 512 | HORVU7Hr1G101590 | Cytochrome P450 superfamily protein | 7.60 | AT3G14690.1 |
| 513 | HORVU5Hr1G097270 | Peroxidase superfamily protein | 7.60 | AT3G49960.1 |
| 514 | HORVU7Hr1G083550 | Peroxidase superfamily protein | 7.56 | AT5G66390.1 |
| 515 | HORVU3Hr1G033740 | WRKY family transcription factor | 7.52 | AT1G62300.1 |
| 516 | HORVU1Hr1G037150 | Protein kinase superfamily protein | 7.52 | AT5G03320.2 |
| 517 | HORVU2Hr1G120420 | undescribed protein | 7.50 | AT5G22010.2 |
| 518 | HORVU7Hr1G104350 | early nodulin-like protein 9 | 7.44 | AT3G20570.1 |
| 519 | HORVU4Hr1G009210 | Sec14p-like phosphatidylinositol transfer family protein | 7.43 | AT1G75170.2 |
| 520 | HORVU4Hr1G009190 | Sec14p-like phosphatidylinositol transfer family protein | 7.41 | AT1G75170.2 |
| 521 | HORVU4Hr1G055280 | Protein of unknown function (DUF1262) | 7.38 | AT1G13520.1 |
| 522 | HORVU1Hr1G054390 | lipoxygenase 1 | 7.37 | AT3G22400.1 |
| 523 | HORVU7Hr1G075710 | N-(5'-phosphoribosyl)anthranilate isomerase | 7.34 | AT1G07780.9 |
| 524 | HORVU6Hr1G066250 | Tetratricopeptide repeat (TPR)-like superfamily protein | 7.34 | AT3G18420.1 |
| 525 | HORVU0Hr1G021610 | F-box protein PP2-B7 | 7.33 | AT2G02360.1 |
| 526 | HORVU2Hr1G077220 | 12-oxophytodienoate reductase 2 | 7.32 | AT1G76690.1 |
| 527 | HORVU4Hr1G083210 | Anthranilate phosphoribosyltransferase | 7.32 | AT5G17990.1 |
| 528 | HORVU1Hr1G093950 | receptor-like protein kinase 1 | 7.32 | AT5G10530.1 |
| 529 | HORVU4Hr1G076840 | Protein TIFY 3A | 7.31 | AT1G74950.1 |
| 530 | HORVU6Hr1G018070 | Zinc transporter 2 | 7.30 | AT1G55910.1 |
| 531 | HORVU2Hr1G032780 | methyl esterase 13 | 7.27 | AT1G26360.1 |

| | | | | |
|---|---|---|---|---|
| 532 | HORVU2Hr1G036570 | sugar transporter 1 | 7.26 | AT2G16120.1 |
| 533 | HORVU7Hr1G028730 | receptor kinase 2 | 7.24 | AT2G37710.1 |
| 534 | HORVU5Hr1G118590 | ammonium transporter 2 | 7.24 | AT2G38290.1 |
| 535 | HORVU2Hr1G038830 | Ferredoxin--NADP reductase | 7.23 | AT4G05390.1 |
| 536 | HORVU7Hr1G036380 | RING/U-box superfamily protein | 7.22 | AT5G27420.1 |
| 537 | HORVU2Hr1G008130 | Serine/threonine-protein kinase BRI1-like 1 | 7.21 | AT4G21380.2 |
| 538 | HORVU4Hr1G059170 | NAD(P)H dehydrogenase (quinone) | 7.19 | AT4G27270.1 |
| 539 | HORVU1Hr1G093480 | tryptophan synthase alpha chain | 7.19 | AT4G02610.1 |
| 540 | HORVU3Hr1G077580 | Peroxidase superfamily protein | 7.17 | AT5G15180.1 |
| 541 | HORVU7Hr1G101240 | Indole-3-glycerol phosphate lyase, chloroplastic | 7.16 | AT1G33320.3 |
| 542 | HORVU7Hr1G077710 | phosphatidylinositol-4-phosphate 5-kinase family protein | 7.13 | AT1G34260.3 |
| 543 | HORVU5Hr1G058170 | unknown function | 7.12 | AT4G39720.4 |
| 544 | HORVU2Hr1G017950 | B12D protein | 7.06 | AT3G48140.1 |
| 545 | HORVU7Hr1G074660 | Transmembrane amino acid transporter family protein | 7.03 | AT5G40780.1 |
| 546 | HORVU4Hr1G055270 | Protein of unknown function, DUF538 | 7.00 | AT5G37070.1 |
| 547 | HORVU0Hr1G039970 | Leucine-rich receptor-like protein kinase family protein | 6.91 | AT3G47090.1 |
| 548 | HORVU2Hr1G003830 | Amidase family protein, expressed | 6.89 | AT4G34880.1 |
| 549 | HORVU4Hr1G086660 | Cyclin-dependent protein kinase inhibitor SMR1 | 6.88 | AT5G02420.1 |
| 550 | HORVU2Hr1G044440 | glucan endo-1,3-beta-glucosidase-like protein 3 | 6.87 | AT2G39640.1 |
| 551 | HORVU7Hr1G008140 | receptor-like protein kinase 4 | 6.87 | AT5G10530.1 |
| 552 | HORVU2Hr1G109330 | WRKY family transcription factor | 6.83 | AT3G56400.1 |
| 553 | HORVU7Hr1G121260 | Flavin-containing monooxygenase family protein | 6.82 | AT1G19250.1 |
| 554 | HORVU3Hr1G022640 | alpha/beta-Hydrolases superfamily protein | 6.80 | AT1G47480.1 |
| 555 | HORVU6Hr1G057270 | Subtilisin-like protease SBT3.6 | 6.78 | AT1G71950.1 |
| 556 | HORVU3Hr1G096360 | HXXXD-type acyl-transferase family protein | 6.77 | AT3G62160.1 |
| 557 | HORVU4Hr1G076820 | ZIM motif family protein | 6.75 | AT2G17390.1 |
| 558 | HORVU6Hr1G074440 | annexin 7 | 6.73 | AT5G10230.1 |
| 559 | HORVU6Hr1G066920 | Acetylornithine deacetylase | 6.73 | AT4G17830.1 |
| 560 | HORVU2Hr1G123390 | unknown function | 6.70 | AT4G10220.1 |
| 561 | HORVU2Hr1G080890 | Anthranilate synthase | 6.68 | AT1G25220.1 |
| 562 | HORVU3Hr1G023400 | UDP-Glycosyltransferase superfamily protein | 6.67 | AT2G36800.1 |
| 563 | HORVU5Hr1G062410 | Hypersensitive-induced response protein 1 | 6.65 | AT5G62740.1 |

| | | | | |
|---|---|---|---|---|
| 564 | HORVU5Hr1G105930 | CASP-like protein 1U3 | 6.65 | AT4G15630.1 |
| 565 | HORVU5Hr1G014060 | calmodulin-like 11 | 6.64 | AT3G22930.1 |
| 566 | HORVU3Hr1G092250 | E3 ubiquitin-protein ligase RGLG2 | 6.64 | AT5G14420.4 |
| 567 | HORVU1Hr1G017700 | Hydrophobic protein LTI6B | 6.64 | AT3G05880.1 |
| 568 | HORVU5Hr1G098090 | allene oxide synthase | 6.63 | AT5G42650.1 |
| 569 | HORVU2Hr1G017350 | NAC domain containing protein 100 | 6.62 | AT5G61430.1 |
| 570 | HORVU5Hr1G010120 | Protein WIR1A | 6.60 | AT2G23450.2 |
| 571 | HORVU1Hr1G001570 | O-methyltransferase family protein | 6.59 | AT4G35160.1 |
| 572 | HORVU1Hr1G081550 | malate dehydrogenase | 6.56 | AT3G47520.1 |
| 573 | HORVU2Hr1G118110 | Heavy metal transport/detoxification superfamily protein | 6.56 | AT1G01490.2 |
| 574 | HORVU3Hr1G085810 | Aldose reductase | 6.55 | AT2G37790.1 |
| 575 | HORVU0Hr1G002930 | unknown function | 6.52 | AT2G15760.1 |
| 576 | HORVU2Hr1G044450 | Glucan endo-1,3-beta-glucosidase 3 | 6.51 | AT5G55180.2 |
| 577 | HORVU5Hr1G062290 | Protein TIFY 3B | 6.50 | AT1G74950.1 |
| 578 | HORVU7Hr1G028840 | 2-oxoglutarate (2OG) and Fe(II)-dependent oxygenase superfamily protein | 6.48 | AT5G54000.2 |
| 579 | HORVU4Hr1G071020 | sugar transporter 9 | 6.46 | AT3G18830.1 |
| 580 | HORVU7Hr1G030570 | undescribed protein | 6.41 | AT2G24040.1 |
| 581 | HORVU4Hr1G083070 | undescribed protein | 6.40 | #N/A |
| 582 | HORVU2Hr1G017470 | NAC domain protein, | 6.38 | AT1G69490.1 |
| 583 | HORVU2Hr1G022080 | Cytochrome P450 superfamily protein | 6.38 | AT2G46950.1 |
| 584 | HORVU4Hr1G090140 | germin-like protein 9 | 6.38 | AT5G39150.1 |
| 585 | HORVU7Hr1G047070 | MATE efflux family protein | 6.37 | AT5G52450.1 |
| 586 | HORVU7Hr1G116310 | Cytochrome P450 superfamily protein | 6.36 | AT3G14660.1 |
| 587 | HORVU4Hr1G067840 | GDP-mannose transporter GONST3 | 6.36 | AT1G76340.1 |
| 588 | HORVU3Hr1G012100 | Protein kinase superfamily protein | 6.34 | AT4G32300.1 |
| 589 | HORVU2Hr1G036590 | sugar transporter 9 | 6.33 | AT3G18830.1 |
| 590 | HORVU5Hr1G066260 | calmodulin like 42 | 6.32 | AT3G29000.1 |
| 591 | HORVU1Hr1G070580 | receptor-like kinase 902 | 6.32 | AT4G18640.1 |
| 592 | HORVU4Hr1G078210 | purine permease 3 | 6.31 | AT1G28220.1 |
| 593 | HORVU3Hr1G016860 | Chymotrypsin inhibitor | 6.29 | AT2G38870.1 |
| 594 | HORVU6Hr1G057680 | inter-alpha-trypsin inhibitor heavy chain-related | 6.28 | AT5G60710.1 |
| 595 | HORVU3Hr1G014270 | Phloem-specific lectin | 6.27 | AT2G02340.1 |
| 596 | HORVU7Hr1G108460 | Leucine-rich repeat receptor-like protein kinase family protein | 6.27 | AT5G20480.2 |
| 597 | HORVU7Hr1G039900 | Family of unknown function (DUF716) | 6.26 | AT3G01360.2 |
| 598 | HORVU2Hr1G045900 | glucose-6-phosphate/phosphate translocator 2 | 6.20 | AT1G61800.1 |
| 599 | HORVU2Hr1G042210 | Serine/threonine-protein kinase | 6.19 | AT2G19130.1 |
| 600 | HORVU2Hr1G121060 | polyamine oxidase 2 | 6.18 | AT1G65840.1 |

| | | | | |
|---|---|---|---|---|
| 601 | HORVU5Hr1G119270 | Protein kinase family protein | 6.15 | AT1G21230.1 |
| 602 | HORVU4Hr1G084740 | sugar transporter 1 | 6.13 | AT1G34580.1 |
| 603 | HORVU3Hr1G087050 | Ankyrin repeat family protein | 6.12 | AT3G12360.1 |
| 604 | HORVU2Hr1G044270 | receptor kinase 1 | 6.11 | AT4G05200.2 |
| 605 | HORVU0Hr1G009460 | germin-like protein 9 | 6.10 | AT5G39110.1 |
| 606 | HORVU6Hr1G004770 | unknown function | 6.07 | AT1G21230.1 |
| 607 | HORVU2Hr1G114100 | Leucine-rich repeat receptor-like protein kinase family protein | 6.07 | AT1G74170.2 |
| 608 | HORVU1Hr1G091600 | Glucose-1-phosphate adenylyltransferase family protein | 6.06 | AT1G27680.1 |
| 609 | HORVU4Hr1G072560 | Blue copper protein | 6.06 | AT2G32300.2 |
| 610 | HORVU1Hr1G020450 | 1-aminocyclopropane-1-carboxylate oxidase | 6.05 | AT1G77330.1 |
| 611 | HORVU5Hr1G110180 | phosphate transporter 1;7 | 6.03 | AT3G54700.2 |
| 612 | HORVU3Hr1G079620 | Protein RETICULATA-RELATED 3, chloroplastic | 6.02 | AT3G08640.1 |
| 613 | HORVU0Hr1G007620 | Leucine-rich receptor-like protein kinase family protein | 6.00 | AT4G08850.1 |
| 614 | HORVU7Hr1G019280 | sugar transporter 1 | 5.99 | AT5G61520.1 |
| 615 | HORVU1Hr1G049280 | Glutathione S-transferase family protein | 5.97 | AT1G10370.1 |
| 616 | HORVU6Hr1G023280 | 0 | 5.95 | AT1G17240.1 |
| 617 | HORVU7Hr1G113350 | NADPH HC toxin reductase | 5.91 | AT4G27250.1 |
| 618 | HORVU3Hr1G024650 | Serine/threonine-protein kinase | 5.91 | AT2G19130.1 |
| 619 | HORVU5Hr1G110220 | phosphate transporter 1;7 | 5.90 | AT3G54700.2 |
| 620 | HORVU5Hr1G058020 | Major facilitator superfamily protein | 5.90 | AT5G10190.2 |
| 621 | HORVU3Hr1G059480 | Acidic endochitinase | 5.89 | AT5G24090.1 |
| 622 | HORVU6Hr1G074310 | undescribed protein | 5.89 | AT3G57020.2 |
| 623 | HORVU4Hr1G007180 | microsomal glutathione s-transferase, putative | 5.87 | AT1G65820.1 |
| 624 | HORVU1Hr1G052250 | undescribed protein | 5.87 | AT3G54710.1 |
| 625 | HORVU2Hr1G125600 | unknown function | 5.84 | AT4G35920.5 |
| 626 | HORVU2Hr1G080140 | Aldose 1-epimerase | 5.84 | AT5G15140.1 |
| 627 | HORVU2Hr1G008140 | receptor kinase 3 | 5.80 | AT4G27300.1 |
| 628 | HORVU2Hr1G009830 | Cystic fibrosis transmembrane conductance regulator | 5.79 | AT2G47800.1 |
| 629 | HORVU5Hr1G047060 | plastid transcriptionally active 6 | 5.75 | AT1G21600.2 |
| 630 | HORVU6Hr1G089960 | undescribed protein | 5.73 | AT1G47890.1 |
| 631 | HORVU5Hr1G081860 | Chaperone protein ClpB | 5.73 | AT5G51070.1 |
| 632 | HORVU3Hr1G077890 | Chromosome 3B, genomic scaffold, cultivar Chinese Spring | 5.72 | AT4G21920.1 |
| 633 | HORVU5Hr1G107290 | Actin cross-linking protein | 5.69 | AT1G59710.2 |
| 634 | HORVU4Hr1G003400 | Coffea canephora DH200=94 genomic scaffold, scaffold_6 | 5.68 | AT2G01820.1 |

| | | | | |
|---|---|---|---|---|
| 635 | HORVU7Hr1G079460 | unknown function | 5.67 | AT5G53710.2 |
| 636 | HORVU2Hr1G118060 | Heavy metal transport/detoxification superfamily protein | 5.67 | AT1G01490.2 |
| 637 | HORVU5Hr1G011540 | 2-oxoglutarate (2OG) and Fe(II)-dependent oxygenase superfamily protein | 5.67 | AT2G36690.2 |
| 638 | HORVU4Hr1G072310 | FAD/NAD(P)-binding oxidoreductase family protein | 5.66 | AT4G38540.1 |
| 639 | HORVU1Hr1G080320 | aldehyde dehydrogenase 12A1 | 5.66 | AT5G62530.1 |
| 640 | HORVU1Hr1G074660 | Subtilisin-like protease SBT3.6 | 5.65 | AT1G71950.1 |
| 641 | HORVU6Hr1G031530 | Cell cycle control protein 50A | 5.61 | AT1G54320.1 |
| 642 | HORVU6Hr1G023530 | Protein kinase domain containing protein, expressed | 5.59 | AT1G21230.1 |
| 643 | HORVU3Hr1G090220 | Protein kinase superfamily protein | 5.59 | AT2G19130.1 |
| 644 | HORVU6Hr1G024850 | Cytochrome P450 superfamily protein | 5.59 | AT3G48310.1 |
| 645 | HORVU5Hr1G063420 | glutamate receptor 2.8 | 5.58 | AT2G29120.1 |
| 646 | HORVU6Hr1G034380 | N-terminal protein myristoylation | 5.57 | AT1G76600.1 |
| 647 | HORVU6Hr1G034570 | receptor kinase 2 | 5.57 | AT2G33580.1 |
| 648 | HORVU5Hr1G012040 | receptor kinase 2 | 5.56 | AT5G65600.1 |
| 649 | HORVU5Hr1G065620 | 2-oxoglutarate (2OG) and Fe(II)-dependent oxygenase superfamily protein | 5.56 | AT5G20400.1 |
| 650 | HORVU2Hr1G109590 | HXXXD-type acyl-transferase family protein | 5.55 | AT5G48930.1 |
| 651 | HORVU7Hr1G019240 | sugar transporter 9 | 5.55 | AT5G61520.2 |
| 652 | HORVU3Hr1G099630 | unknown function | 5.55 | AT3G20980.1 |
| 653 | HORVU7Hr1G118560 | Blue copper protein | 5.54 | AT3G17675.1 |
| 654 | HORVU4Hr1G000850 | Calcium-dependent lipid-binding (CaLB domain) family protein | 5.54 | AT5G23950.2 |
| 655 | HORVU6Hr1G093180 | Electron transfer flavoprotein subunit beta | 5.53 | AT5G43430.1 |
| 656 | HORVU2Hr1G027480 | Cytochrome P450 superfamily protein | 5.53 | AT2G46950.1 |
| 657 | HORVU5Hr1G024160 | Adenine nucleotide alpha hydrolases-like superfamily protein | 5.51 | AT2G47710.1 |
| 658 | HORVU6Hr1G021320 | Lipid A export ATP-binding/permease protein MsbA | 5.49 | AT2G47000.7 |
| 659 | HORVU1Hr1G016840 | Peroxidase superfamily protein | 5.49 | AT1G05260.1 |
| 660 | HORVU2Hr1G044870 | receptor kinase 2 | 5.44 | AT4G23180.1 |
| 661 | HORVU1Hr1G028200 | hexokinase 2 | 5.41 | AT4G29130.1 |
| 662 | HORVU5Hr1G071140 | cellulose synthase like E1 | 5.39 | AT1G55850.1 |
| 663 | HORVU1Hr1G000140 | Phospholipid-transporting ATPase 10 | 5.39 | AT1G68710.2 |
| 664 | HORVU5Hr1G071720 | undescribed protein | 5.38 | AT5G04690.1 |
| 665 | HORVU7Hr1G091100 | Glucan endo-1,3-beta-glucosidase 13 | 5.38 | AT2G05790.1 |
| 666 | HORVU4Hr1G083650 | Protein NRT1/ PTR FAMILY 5.2 | 5.38 | AT5G46050.1 |

| | | | | |
|---|---|---|---|---|
| 667 | HORVU1Hr1G089830 | Dehydrogenase/reductase SDR family member 12 | 5.36 | AT1G01800.1 |
| 668 | HORVU5Hr1G063340 | ATP-dependent zinc metalloprotease FtsH 2 | 5.35 | AT5G57480.1 |
| 669 | HORVU3Hr1G086220 | Calcium-dependent lipid-binding (CaLB domain) family protein | 5.35 | AT3G55470.2 |
| 670 | HORVU2Hr1G098390 | receptor kinase 2 | 5.35 | AT1G56120.3 |
| 671 | HORVU4Hr1G007340 | Homologous-pairing protein 2 homolog | 5.33 | AT1G13330.1 |
| 672 | HORVU1Hr1G074330 | Fructose-1,6-bisphosphatase class 1 | 5.32 | AT1G43670.1 |
| 673 | HORVU2Hr1G121440 | Cathepsin B-like cysteine proteinase | 5.32 | AT5G43060.1 |
| 674 | HORVU3Hr1G007420 | calmodulin like 23 | 5.31 | AT5G37770.2 |
| 675 | HORVU2Hr1G062730 | Protein NRT1/ PTR FAMILY 4.5 | 5.30 | AT1G27040.2 |
| 676 | HORVU7Hr1G024930 | Subtilisin-like protease | 5.29 | AT4G10540.1 |
| 677 | HORVU4Hr1G084600 | unknown function | 5.28 | AT2G40095.3 |
| 678 | HORVU3Hr1G079000 | Dihydropyrimidinase | 5.28 | AT5G12200.1 |
| 679 | HORVU0Hr1G005160 | L-allo-threonine aldolase | 5.27 | AT1G08630.6 |
| 680 | HORVU1Hr1G088680 | Receptor-like protein kinase | 5.27 | AT3G51550.1 |
| 681 | HORVU3Hr1G028280 | RING/U-box superfamily protein | 5.26 | AT2G42360.1 |
| 682 | HORVU1Hr1G066120 | Kinesin-like protein | 5.25 | AT1G03280.3 |
| 683 | HORVU6Hr1G005400 | Epoxide hydrolase 4 | 5.24 | AT3G51000.1 |
| 684 | HORVU5Hr1G117910 | ferredoxin 3 | 5.22 | AT2G27510.1 |
| 685 | HORVU7Hr1G079630 | undescribed protein | 5.22 | AT2G33600.1 |
| 686 | HORVU5Hr1G120570 | Membrane protein insertase YidC | 5.21 | AT1G24490.1 |
| 687 | HORVU4Hr1G078690 | Ras-related protein Rab-6A | 5.20 | AT2G44610.1 |
| 688 | HORVU7Hr1G090830 | Copine family protein 2 | 5.16 | AT5G14420.4 |
| 689 | HORVU5Hr1G089780 | Molybdenum cofactor sulfurase family protein | 5.15 | AT1G30910.1 |
| 690 | HORVU4Hr1G016770 | alcohol dehydrogenase 1 | 5.12 | AT1G77120.1 |
| 691 | HORVU1Hr1G020410 | 1-aminocyclopropane-1-carboxylate oxidase | 5.11 | AT1G77330.1 |
| 692 | HORVU5Hr1G001920 | Leucine-rich receptor-like protein kinase family protein | 5.10 | AT1G09970.1 |
| 693 | HORVU2Hr1G007760 | ABC transporter family protein | 5.08 | AT5G60740.1 |
| 694 | HORVU7Hr1G059050 | Pyrimidine-specific ribonucleoside hydrolase RihA | 5.06 | AT2G36310.1 |
| 695 | HORVU7Hr1G079610 | undescribed protein | 5.06 | #N/A |
| 696 | HORVU0Hr1G026470 | undescribed protein | 5.06 | AT4G34215.2 |
| 697 | HORVU2Hr1G117840 | Protein kinase superfamily protein | 5.04 | AT4G21390.1 |
| 698 | HORVU1Hr1G080300 | WRKY DNA-binding protein 51 | 5.04 | AT5G64810.1 |
| 699 | HORVU7Hr1G118020 | Eukaryotic aspartyl protease family protein | 5.03 | AT4G35880.1 |
| 700 | HORVU4Hr1G010030 | Protein kinase superfamily protein | 5.02 | AT1G34300.1 |

| | | | | |
|---|---|---|---|---|
| 701 | HORVU7Hr1G028750 | F-box domain containing protein, expressed | 5.00 | AT3G06240.1 |
| 702 | HORVU3Hr1G028060 | GDSL esterase/lipase | 4.99 | AT5G45910.1 |
| 703 | HORVU5Hr1G012120 | receptor kinase 2 | 4.94 | AT5G10530.1 |
| 704 | HORVU5Hr1G064020 | Tetraspanin family protein | 4.94 | AT2G23810.1 |
| 705 | HORVU7Hr1G037910 | shikimate kinase 1 | 4.93 | AT2G21940.5 |
| 706 | HORVU6Hr1G011120 | Glutathione S-transferase family protein | 4.92 | AT4G19880.1 |
| 707 | HORVU4Hr1G023390 | unknown protein; BEST Arabidopsis thaliana protein match is: unknown protein . | 4.92 | AT3G48860.2 |
| 708 | HORVU7Hr1G030480 | cinnamoyl coa reductase | 4.91 | AT1G80820.1 |
| 709 | HORVU7Hr1G074640 | lysine histidine transporter 1 | 4.88 | AT5G40780.1 |
| 710 | HORVU2Hr1G120260 | Chromatin structure-remodeling complex subunit snf21 | 4.85 | AT2G28290.1 |
| 711 | HORVU5Hr1G096930 | Cytochrome P450 superfamily protein | 4.84 | AT4G37330.1 |
| 712 | HORVU5Hr1G087880 | Auxin efflux carrier family protein | 4.84 | AT5G65980.1 |
| 713 | HORVU4Hr1G009340 | Protein kinase superfamily protein | 4.83 | AT2G16250.1 |
| 714 | HORVU5Hr1G107410 | AAA-ATPase 1 | 4.81 | AT3G28510.1 |
| 715 | HORVU1Hr1G016660 | Peroxidase superfamily protein | 4.77 | AT4G33420.1 |
| 716 | HORVU7Hr1G113850 | WRKY family transcription factor | 4.77 | AT1G80840.1 |
| 717 | HORVU0Hr1G009400 | germin-like protein 9 | 4.74 | AT5G39110.1 |
| 718 | HORVU2Hr1G004230 | 12-oxophytodienoate reductase 2 | 4.73 | AT1G76680.1 |
| 719 | HORVU2Hr1G064160 | sugar transporter 1 | 4.72 | AT1G11260.1 |
| 720 | HORVU5Hr1G077010 | NAC domain containing protein 41 | 4.72 | AT2G33480.2 |
| 721 | HORVU7Hr1G073460 | Tryptophan synthase beta chain 1 | 4.71 | AT4G27070.1 |
| 722 | HORVU3Hr1G076060 | purple acid phosphatase 10 | 4.71 | AT2G16430.2 |
| 723 | HORVU2Hr1G096510 | 2-oxoglutarate (2OG) and Fe(II)-dependent oxygenase superfamily protein | 4.71 | AT5G20400.1 |
| 724 | HORVU1Hr1G002890 | HXXXD-type acyl-transferase family protein | 4.70 | AT1G03940.1 |
| 725 | HORVU1Hr1G049210 | Glutathione S-transferase family protein | 4.70 | AT1G10360.1 |
| 726 | HORVU2Hr1G123590 | Protein kinase family protein | 4.67 | AT1G21210.1 |
| 727 | HORVU7Hr1G079600 | unknown function | 4.66 | AT1G17147.1 |
| 728 | HORVU3Hr1G093550 | receptor kinase 2 | 4.65 | AT1G56130.2 |
| 729 | HORVU3Hr1G077110 | receptor kinase 2 | 4.63 | AT4G21390.1 |
| 730 | HORVU1Hr1G062250 | UDP-Glycosyltransferase superfamily protein | 4.63 | AT3G16520.3 |
| 731 | HORVU3Hr1G003920 | receptor kinase 2 | 4.63 | AT4G00970.1 |
| 732 | HORVU0Hr1G009470 | germin-like protein 2 | 4.59 | AT5G39110.1 |
| 733 | HORVU1Hr1G007310 | Calcium-transporting ATPase 1 | 4.55 | AT1G07670.2 |
| 734 | HORVU1Hr1G057460 | Aldolase-type TIM barrel family protein | 4.55 | AT5G64250.1 |
| 735 | HORVU2Hr1G036800 | Geranylgeranyl diphosphate synthase | 4.53 | AT4G36810.1 |

| | | | | |
|---|---|---|---|---|
| 736 | HORVU0Hr1G007680 | inositol polyphosphate 5-phosphatase 11 | 4.53 | AT1G47510.1 |
| 737 | HORVU2Hr1G014890 | receptor kinase 2 | 4.50 | AT3G53810.1 |
| 738 | HORVU0Hr1G012940 | Alcohol dehydrogenase | 4.49 | AT4G37980.2 |
| 739 | HORVU2Hr1G103890 | receptor kinase 2 | 4.49 | AT1G16670.1 |
| 740 | HORVU3Hr1G049640 | calmodulin-binding family protein | 4.49 | AT4G33050.3 |
| 741 | HORVU2Hr1G087090 | adenine phosphoribosyltransferase 5 | 4.47 | AT1G80050.1 |
| 742 | HORVU7Hr1G073050 | External alternative NAD(P)H-ubiquinone oxidoreductase B3, mitochondrial | 4.45 | AT4G21490.2 |
| 743 | HORVU4Hr1G058550 | Peptidase M50B-like protein | 4.45 | AT1G67060.1 |
| 744 | HORVU3Hr1G088200 | WRKY DNA-binding protein 33 | 4.44 | AT2G38470.1 |
| 745 | HORVU3Hr1G002350 | receptor-like protein kinase 4 | 4.42 | AT5G38280.1 |
| 746 | HORVU2Hr1G116910 | diacylglycerol kinase 5 | 4.41 | AT2G20900.3 |
| 747 | HORVU7Hr1G038160 | sphingoid base hydroxylase 2 | 4.40 | AT1G14290.1 |
| 748 | HORVU7Hr1G022050 | undescribed protein | 4.40 | AT1G10170.4 |
| 749 | HORVU3Hr1G076750 | Secretory carrier-associated membrane protein 2 | 4.39 | AT1G61250.1 |
| 750 | HORVU1Hr1G019410 | HXXXD-type acyl-transferase family protein | 4.38 | AT5G48930.1 |
| 751 | HORVU2Hr1G100450 | HKT14-1 | 4.38 | AT4G10310.1 |
| 752 | HORVU1Hr1G074350 | actin 7 | 4.36 | AT5G09810.1 |
| 753 | HORVU7Hr1G104190 | undescribed protein | 4.36 | AT3G47100.1 |
| 754 | HORVU2Hr1G078510 | lipoxygenase 3 | 4.35 | AT1G17420.1 |
| 755 | HORVU5Hr1G040740 | Long-chain-fatty-acid--CoA ligase 1 | 4.34 | AT5G27600.1 |
| 756 | HORVU2Hr1G107550 | Aldehyde dehydrogenase family 9 member A1-B | 4.33 | AT2G14170.1 |
| 757 | HORVU1Hr1G045610 | membrane steroid binding protein 1 | 4.33 | AT5G52240.1 |
| 758 | HORVU5Hr1G115230 | non-specific phospholipase C1 | 4.32 | AT1G07230.1 |
| 759 | HORVU2Hr1G085270 | Chitinase | 4.32 | AT3G54420.1 |
| 760 | HORVU2Hr1G117940 | Heavy metal transport/detoxification superfamily protein | 4.29 | AT1G01490.2 |
| 761 | HORVU3Hr1G081100 | receptor kinase 1 | 4.28 | AT3G08760.3 |
| 762 | HORVU1Hr1G049100 | Glutathione S-transferase family protein | 4.26 | AT1G10360.1 |
| 763 | HORVU3Hr1G087740 | TRICHOME BIREFRINGENCE-LIKE 28 | 4.26 | AT3G55990.1 |
| 764 | HORVU7Hr1G122130 | undescribed protein | 4.24 | AT2G25800.1 |
| 765 | HORVU5Hr1G014980 | N-carbamoyl-L-amino acid hydrolase | 4.20 | AT5G43600.1 |
| 766 | HORVU4Hr1G005450 | Protein of unknown function, DUF538 | 4.19 | AT5G19860.1 |
| 767 | HORVU6Hr1G033980 | Citrate synthase family protein | 4.19 | AT3G60100.3 |
| 768 | HORVU6Hr1G090600 | Glutathione S-transferase family protein | 4.17 | AT1G10370.1 |
| 769 | HORVU3Hr1G111360 | undescribed protein | 4.16 | AT5G01950.8 |
| 770 | HORVU2Hr1G078890 | unknown function | 4.14 | AT4G37030.1 |
| 771 | HORVU6Hr1G028790 | WRKY family transcription factor | 4.14 | AT1G80840.1 |

| | | | | |
|---|---|---|---|---|
| 772 | HORVU3Hr1G108900 | LORELEI-LIKE-GPI-ANCHORED PROTEIN 1 | 4.13 | AT5G56170.1 |
| 773 | HORVU2Hr1G084210 | Leucine-rich repeat receptor-like protein kinase family protein | 4.13 | AT3G28450.1 |
| 774 | HORVU4Hr1G087340 | 60 kDa jasmonate-induced protein | 4.10 | AT2G35330.2 |
| 775 | HORVU3Hr1G031020 | Cytochrome P450 superfamily protein | 4.09 | AT3G48320.1 |
| 776 | HORVU0Hr1G001430 | WRKY DNA-binding protein 51 | 4.09 | AT5G64810.1 |
| 777 | HORVU3Hr1G106950 | Calcium-binding EF-hand family protein | 4.08 | AT1G24620.1 |
| 778 | HORVU6Hr1G084740 | Cytochrome c oxidase copper chaperone 2 | 4.07 | AT1G53030.1 |
| 779 | HORVU3Hr1G002920 | Glycosyltransferase | 4.06 | AT3G18180.1 |
| 780 | HORVU7Hr1G010250 | Peroxisomal membrane protein 11-5 | 4.03 | AT1G01820.1 |
| 781 | HORVU5Hr1G089620 | receptor kinase 2 | 4.02 | AT1G21240.1 |
| 782 | HORVU1Hr1G038680 | 60 kDa chaperonin 1 | 4.01 | AT3G23990.1 |
| 783 | HORVU7Hr1G121950 | Retrotransposon protein, putative, unclassified | 4.01 | AT4G29090.1 |
| 784 | HORVU1Hr1G072720 | unknown protein; Has 30 Blast hits to 30 proteins in 10 species: Archae - 0; Bacteria - 0; Metazoa - 0; Fungi - 0; Plants - 30; Viruses - 0; Other Eukaryotes - 0 (source: NCBI BLink). | 4.00 | AT5G02090.1 |
| 785 | HORVU7Hr1G037510 | Protein of unknown function, DUF547 | 4.00 | AT4G08550.2 |
| 786 | HORVU3Hr1G061700 | callose synthase 5 | 3.99 | AT4G03550.1 |
| 787 | HORVU6Hr1G077890 | Non-specific lipid transfer protein-like 1 | 3.99 | AT5G64080.2 |
| 788 | HORVU2Hr1G048220 | Transmembrane protein 184A | 3.97 | AT4G21570.1 |
| 789 | HORVU5Hr1G068060 | Mitochondrial phosphate carrier protein 2, mitochondrial | 3.97 | AT5G14040.1 |
| 790 | HORVU1Hr1G086170 | undescribed protein | 3.95 | AT1G53708.1 |
| 791 | HORVU5Hr1G009930 | undescribed protein | 3.93 | AT5G37660.1 |
| 792 | HORVU6Hr1G084590 | Isopentenyl-diphosphate Delta-isomerase | 3.93 | AT1G79690.2 |
| 793 | HORVU7Hr1G008760 | soluble N-ethylmaleimide-sensitive factor adaptor protein 33 | 3.92 | AT5G61210.2 |
| 794 | HORVU5Hr1G096940 | Cytochrome P450 superfamily protein | 3.92 | AT4G37330.1 |
| 795 | HORVU6Hr1G061450 | YELLOW STRIPE like 3 | 3.92 | AT5G53550.6 |
| 796 | HORVU7Hr1G108820 | unknown function | 3.92 | AT2G18680.1 |
| 797 | HORVU6Hr1G018460 | unknown function | 3.92 | AT1G48120.1 |
| 798 | HORVU7Hr1G050670 | lipoxygenase 2 | 3.92 | AT3G45140.1 |
| 799 | HORVU7Hr1G116880 | Protein of unknown function (DUF668) | 3.91 | AT1G30755.6 |
| 800 | HORVU1Hr1G073280 | Glycosyl hydrolase family protein | 3.90 | AT3G47000.1 |
| 801 | HORVU2Hr1G113180 | D-3-phosphoglycerate dehydrogenase | 3.90 | AT4G34200.1 |
| 802 | HORVU3Hr1G105290 | Glucan endo-1,3-beta-glucosidase GI | 3.89 | AT3G57240.1 |
| 803 | HORVU5Hr1G023990 | unknown function | 3.86 | AT1G67120.1 |
| 804 | HORVU0Hr1G017540 | glutathione reductase | 3.86 | AT3G24170.3 |

| | | | | |
|---|---|---|---|---|
| 805 | HORVU2Hr1G117490 | Serine/threonine-protein kinase | 3.85 | AT1G65790.3 |
| 806 | HORVU4Hr1G072880 | phosphoserine aminotransferase | 3.85 | AT4G35630.1 |
| 807 | HORVU1Hr1G084720 | unknown function | 3.81 | AT3G20630.1 |
| 808 | HORVU6Hr1G041840 | unknown function | 3.81 | ATMG00860.1 |
| 809 | HORVU6Hr1G066930 | Acetylornithine deacetylase | 3.81 | AT4G17830.1 |
| 810 | HORVU2Hr1G040900 | NADPH--cytochrome P450 reductase 2 | 3.80 | AT4G30210.3 |
| 811 | HORVU6Hr1G093740 | Receptor-like protein kinase | 3.80 | AT1G21230.1 |
| 812 | HORVU2Hr1G064150 | sugar transporter 1 | 3.80 | AT1G11260.1 |
| 813 | HORVU7Hr1G051070 | Aminoacylase-1 | 3.79 | AT4G38220.1 |
| 814 | HORVU2Hr1G016000 | lon protease 1 | 3.77 | AT5G26860.1 |
| 815 | HORVU5Hr1G099670 | Class E vacuolar protein-sorting machinery protein hse1 | 3.75 | AT4G32760.1 |
| 816 | HORVU6Hr1G078650 | fucosyltransferase 1 | 3.74 | AT2G03220.1 |
| 817 | HORVU6Hr1G077010 | Endoglucanase 7 | 3.74 | AT2G32990.1 |
| 818 | HORVU1Hr1G039600 | Leucine-rich repeat receptor-like protein kinase family protein | 3.74 | AT4G04220.1 |
| 819 | HORVU6Hr1G024100 | Fatty acid oxidation complex subunit alpha | 3.73 | AT4G29010.1 |
| 820 | HORVU7Hr1G099340 | receptor kinase 1 | 3.73 | AT4G23280.2 |
| 821 | HORVU2Hr1G031460 | Adipocyte plasma membrane-associated protein | 3.72 | AT3G51430.2 |
| 822 | HORVU5Hr1G050940 | cinnamoyl coa reductase | 3.72 | AT1G15950.3 |
| 823 | HORVU5Hr1G089790 | receptor kinase 3 | 3.72 | AT1G21240.1 |
| 824 | HORVU5Hr1G099820 | citrate synthase 3 | 3.71 | AT3G58750.1 |
| 825 | HORVU1Hr1G053260 | U-box domain-containing protein | 3.71 | AT2G45910.2 |
| 826 | HORVU4Hr1G054060 | Argininosuccinate lyase | 3.70 | AT5G10920.1 |
| 827 | HORVU1Hr1G045580 | undescribed protein | 3.70 | AT1G55020.1 |
| 828 | HORVU4Hr1G076570 | lipoxygenase 3 | 3.69 | AT1G72520.1 |
| 829 | HORVU4Hr1G043530 | Phytoalexin deficient 4 | 3.69 | AT3G52430.1 |
| 830 | HORVU5Hr1G093860 | Structural constituent of ribosome protein | 3.68 | AT4G19390.2 |
| 831 | HORVU1Hr1G058810 | UDP-glucuronic acid decarboxylase 1 | 3.67 | AT3G62830.2 |
| 832 | HORVU6Hr1G059420 | YELLOW STRIPE like 7 | 3.66 | AT1G65730.1 |
| 833 | HORVU7Hr1G005270 | cellulose synthase-like D3 | 3.65 | AT3G03050.1 |
| 834 | HORVU3Hr1G024210 | ABC transporter C family member 3 | 3.64 | AT3G13080.5 |
| 835 | HORVU4Hr1G081940 | 4-coumarate--CoA ligase-like 4 | 3.63 | AT1G20510.1 |
| 836 | HORVU5Hr1G097940 | undescribed protein | 3.61 | AT2G15960.1 |
| 837 | HORVU1Hr1G069800 | Non-specific lipid-transfer protein | 3.61 | AT5G01870.1 |
| 838 | HORVU1Hr1G070640 | RING/U-box superfamily protein | 3.60 | AT3G11110.1 |
| 839 | HORVU2Hr1G078860 | Protein phosphatase 2C family protein | 3.59 | AT1G47380.1 |
| 840 | HORVU5Hr1G085310 | UDP-Glycosyltransferase superfamily protein | 3.59 | AT1G07250.1 |

| | | | | |
|---|---|---|---|---|
| 841 | HORVU2Hr1G086380 | hydroxycinnamoyl-CoA shikimate/quinate hydroxycinnamoyl transferase | 3.58 | AT5G48930.1 |
| 842 | HORVU1Hr1G063410 | Protein O-linked-mannose beta-1,4-N-acetylglucosaminyltransferase 2 | 3.58 | AT3G18170.1 |
| 843 | HORVU2Hr1G104450 | D-2-hydroxyglutarate dehydrogenase, mitochondrial | 3.57 | AT4G36400.2 |
| 844 | HORVU7Hr1G053300 | RPM1 interacting protein 4 transcript 2 | 3.57 | AT3G25070.2 |
| 845 | HORVU3Hr1G087680 | Phytol kinase 1, chloroplastic | 3.55 | AT5G58560.1 |
| 846 | HORVU4Hr1G061130 | receptor kinase 1 | 3.53 | AT3G09010.1 |
| 847 | HORVU1Hr1G073840 | ferredoxin 3 | 3.53 | AT2G27510.1 |
| 848 | HORVU5Hr1G097500 | Zinc finger protein 1 | 3.53 | AT1G27730.1 |
| 849 | HORVU3Hr1G071890 | Protein TIC 22-like, chloroplastic | 3.51 | AT3G23710.1 |
| 850 | HORVU2Hr1G017400 | NAC domain protein, | 3.51 | AT5G39610.1 |
| 851 | HORVU5Hr1G103180 | Ferredoxin--NADP reductase | 3.51 | AT4G05390.1 |
| 852 | HORVU6Hr1G065210 | transcription factor-related | 3.50 | AT2G27230.2 |
| 853 | HORVU3Hr1G053380 | ABC transporter G family member 37 | 3.49 | AT1G15520.1 |
| 854 | HORVU2Hr1G015410 | Wound-induced protein 1 | 3.49 | AT5G01740.1 |
| 855 | HORVU1Hr1G073040 | Phosphatidylinositol:ceramide inositolphosphotransferase | 3.46 | AT3G54020.1 |
| 856 | HORVU2Hr1G039850 | MATE efflux family protein | 3.46 | AT1G15170.2 |
| 857 | HORVU5Hr1G076250 | purple acid phosphatase 27 | 3.46 | AT5G50400.1 |
| 858 | HORVU2Hr1G013200 | LITAF-domain-containing protein | 3.45 | AT5G13190.1 |
| 859 | HORVU1Hr1G011430 | Pyrimidine-specific ribonucleoside hydrolase RihA | 3.44 | AT2G36310.1 |
| 860 | HORVU3Hr1G073220 | aspartate aminotransferase 3 | 3.44 | AT5G11520.1 |
| 861 | HORVU7Hr1G106420 | Chromosome 3B, genomic scaffold, cultivar Chinese Spring | 3.43 | AT2G38880.8 |
| 862 | HORVU5Hr1G106550 | histone H1-3 | 3.41 | AT2G30620.2 |
| 863 | HORVU1Hr1G020430 | 1-aminocyclopropane-1-carboxylate oxidase | 3.39 | AT1G77330.1 |
| 864 | HORVU3Hr1G070550 | Transferrin receptor protein 2 | 3.39 | AT5G19740.1 |
| 865 | HORVU3Hr1G096050 | Transaldolase 1 | 3.37 | AT5G13420.1 |
| 866 | HORVU7Hr1G017950 | Cytochrome P450 superfamily protein | 3.36 | AT2G45560.1 |
| 867 | HORVU6Hr1G013750 | Protein kinase superfamily protein | 3.36 | AT2G19130.1 |
| 868 | HORVU7Hr1G020580 | Cadmium tolerant 1 | 3.34 | #N/A |
| 869 | HORVU3Hr1G086040 | Major facilitator superfamily protein | 3.34 | AT2G39210.1 |
| 870 | HORVU7Hr1G040030 | unknown function | 3.34 | AT1G19020.1 |
| 871 | HORVU5Hr1G006900 | cationic amino acid transporter 2 | 3.34 | AT1G58030.1 |
| 872 | HORVU2Hr1G015360 | Wound-induced protein 1 | 3.33 | AT5G01740.1 |
| 873 | HORVU4Hr1G027150 | Mitochondrial uncoupling protein 1 | 3.33 | AT3G54110.1 |
| 874 | HORVU4Hr1G064720 | Deoxymugineic acid synthase1 | 3.33 | AT1G59960.1 |
| 875 | HORVU7Hr1G007540 | unknown function | 3.32 | AT1G09070.1 |

| | | | | |
|---|---|---|---|---|
| 876 | HORVU1Hr1G038330 | 3-ketoacyl-CoA thiolase | 3.32 | AT2G33150.1 |
| 877 | HORVU4Hr1G027660 | metallothionein 2A | 3.31 | AT3G09390.2 |
| 878 | HORVU5Hr1G092960 | Ornithine aminotransferase | 3.31 | AT5G46180.1 |
| 879 | HORVU5Hr1G115100 | GRAM domain-containing protein / ABA-responsive protein-related | 3.28 | AT5G13200.1 |
| 880 | HORVU6Hr1G071780 | Transmembrane 9 superfamily member 12 | 3.28 | AT4G12650.1 |
| 881 | HORVU2Hr1G104110 | UDP-glucose 4-epimerase 4 | 3.27 | AT1G30620.6 |
| 882 | HORVU2Hr1G075470 | Protein kinase superfamily protein | 3.27 | AT5G63650.1 |
| 883 | HORVU3Hr1G007680 | receptor-like protein kinase 4 | 3.26 | AT5G38260.1 |
| 884 | HORVU3Hr1G099620 | Chromosome 3B, genomic scaffold, cultivar Chinese Spring | 3.26 | AT3G20980.1 |
| 885 | HORVU5Hr1G079690 | Acyl-coenzyme A thioesterase 9, mitochondrial | 3.26 | AT5G48370.2 |
| 886 | HORVU7Hr1G022970 | Proteasome subunit beta type-5-A | 3.25 | AT1G13060.1 |
| 887 | HORVU4Hr1G002180 | Ankyrin repeat domain-containing protein 50 | 3.24 | AT5G02620.4 |
| 888 | HORVU6Hr1G090560 | Glutathione S-transferase family protein | 3.24 | AT1G10360.1 |
| 889 | HORVU2Hr1G114260 | Glycerol kinase | 3.23 | AT1G80460.1 |
| 890 | HORVU2Hr1G011610 | SNARE-interacting protein KEULE | 3.22 | AT1G12360.1 |
| 891 | HORVU2Hr1G022620 | Disease resistance protein | 3.22 | AT1G50180.1 |
| 892 | HORVU3Hr1G085460 | SOUL heme-binding family protein | 3.22 | AT2G37970.1 |
| 893 | HORVU3Hr1G083560 | Uricase | 3.22 | AT2G26230.1 |
| 894 | HORVU1Hr1G092300 | receptor kinase 2 | 3.21 | AT4G23190.2 |
| 895 | HORVU5Hr1G118460 | Protein kinase superfamily protein | 3.21 | AT1G34300.1 |
| 896 | HORVU5Hr1G012890 | Epoxide hydrolase 4 | 3.20 | AT3G51000.1 |
| 897 | HORVU4Hr1G081670 | respiratory burst oxidase homologue D | 3.17 | AT5G47910.1 |
| 898 | HORVU4Hr1G078130 | GRAM domain family protein | 3.16 | AT1G28200.2 |
| 899 | HORVU4Hr1G007050 | Phytanoyl-CoA dioxygenase domain-containing protein 1 | 3.16 | AT2G01490.2 |
| 900 | HORVU2Hr1G091270 | beta-xylosidase 2 | 3.15 | AT5G10560.1 |
| 901 | HORVU4Hr1G063890 | unknown function | 3.15 | AT5G53670.1 |
| 902 | HORVU3Hr1G018650 | pyruvate decarboxylase-2 | 3.15 | AT5G01320.1 |
| 903 | HORVU1Hr1G076870 | proline iminopeptidase | 3.14 | AT2G14260.3 |
| 904 | HORVU3Hr1G013880 | Fatty acid oxidation complex subunit alpha | 3.14 | AT4G29010.1 |
| 905 | HORVU6Hr1G006560 | 1-deoxy-D-xylulose 5-phosphate synthase 3 | 3.13 | AT1G55510.1 |
| 906 | HORVU3Hr1G096190 | Protein kinase family protein | 3.13 | AT1G21230.1 |
| 907 | HORVU4Hr1G059120 | UDP-glucose 4-epimerase | 3.12 | AT1G78570.1 |
| 908 | HORVU5Hr1G050350 | Glyoxylate/hydroxypyruvate reductase A | 3.12 | AT1G72190.1 |
| 909 | HORVU6Hr1G066470 | Calcium-dependent protein kinase family protein | 3.12 | AT4G35310.2 |

| | | | | |
|---|---|---|---|---|
| 910 | HORVU3Hr1G090070 | undescribed protein | 3.11 | AT1G20400.1 |
| 911 | HORVU2Hr1G080070 | Aldose 1-epimerase | 3.10 | AT5G15140.1 |
| 912 | HORVU1Hr1G084800 | 2-oxoglutarate (2OG) and Fe(II)-dependent oxygenase superfamily protein | 3.10 | AT4G13400.1 |
| 913 | HORVU6Hr1G081000 | allene oxide cyclase 4 | 3.10 | AT3G25780.1 |
| 914 | HORVU2Hr1G028540 | Lipase/lipooxygenase, putative isoform 3 | 3.10 | AT1G55280.1 |
| 915 | HORVU2Hr1G092820 | YELLOW STRIPE like 2 | 3.09 | AT5G24380.1 |
| 916 | HORVU2Hr1G037750 | receptor kinase 3 | 3.08 | AT4G23180.1 |
| 917 | HORVU1Hr1G002470 | receptor-like protein kinase 4 | 3.06 | AT1G70250.1 |
| 918 | HORVU3Hr1G015150 | Sphingosine-1-phosphate lyase | 3.06 | AT1G27980.1 |
| 919 | HORVU7Hr1G109810 | Core-2/I-branching beta-1,6-N-acetylglucosaminyltransferase family protein | 3.05 | AT4G31350.4 |
| 920 | HORVU2Hr1G065750 | WRKY family transcription factor | 3.05 | AT4G31550.1 |
| 921 | HORVU4Hr1G084510 | protein kinase family protein | 3.03 | AT5G24010.1 |
| 922 | HORVU6Hr1G091130 | 3-ketoacyl-CoA thiolase | 3.03 | AT2G33150.1 |
| 923 | HORVU3Hr1G093400 | RNA-binding protein 1 | 3.03 | AT3G23830.2 |
| 924 | HORVU4Hr1G075160 | unknown function | -3.00 | AT1G14345.1 |
| 925 | HORVU5Hr1G056390 | Galactose mutarotase-like superfamily protein | -3.00 | AT5G66530.3 |
| 926 | HORVU3Hr1G030540 | Replication protein A 32 kDa subunit B | -3.00 | AT2G24490.2 |
| 927 | HORVU3Hr1G096650 | NAD(P)H dehydrogenase 18 | -3.01 | AT5G43750.1 |
| 928 | HORVU4Hr1G072240 | unknown function | -3.01 | AT2G36895.2 |
| 929 | HORVU5Hr1G046250 | RING/U-box superfamily protein | -3.01 | AT5G01980.1 |
| 930 | HORVU6Hr1G010520 | unknown protein; Has 52 Blast hits to 46 proteins in 20 species: Archae - 0; Bacteria - 0; Metazoa - 0; Fungi - 0; Plants - 45; Viruses - 0; Other Eukaryotes - 7 (source: NCBI BLink). | -3.02 | AT3G50685.1 |
| 931 | HORVU2Hr1G013870 | fatty acid desaturase 8 | -3.03 | AT5G05580.1 |
| 932 | HORVU5Hr1G112670 | Chalcone--flavonone isomerase | -3.03 | AT3G55120.1 |
| 933 | HORVU0Hr1G031970 | Protein DEHYDRATION-INDUCED 19 homolog 5 | -3.03 | AT3G05700.1 |
| 934 | HORVU2Hr1G092280 | NAD(P)H-quinone oxidoreductase subunit M | -3.03 | AT4G37925.1 |
| 935 | HORVU1Hr1G073170 | LSD1 zinc finger family protein | -3.03 | AT1G32540.1 |
| 936 | HORVU7Hr1G106280 | auxin response factor 6 | -3.03 | AT5G37020.1 |
| 937 | HORVU7Hr1G105250 | DUF4408 domain protein | -3.03 | AT5G57510.1 |
| 938 | HORVU0Hr1G004830 | glyceraldehyde-3-phosphate dehydrogenase C2 | -3.03 | AT1G12900.5 |
| 939 | HORVU0Hr1G032640 | RNAase | -3.03 | #N/A |
| 940 | HORVU7Hr1G120190 | Ycf48-like protein | -3.03 | AT5G23120.2 |

| | | | | |
|---|---|---|---|---|
| 941 | HORVU7Hr1G064100 | proton gradient regulation 5 | -3.03 | AT2G05620.2 |
| 942 | HORVU3Hr1G047030 | Ferredoxin-thioredoxin reductase, catalytic chain | -3.04 | AT2G04700.3 |
| 943 | HORVU3Hr1G062490 | 30S ribosomal protein S20 | -3.04 | AT3G15190.1 |
| 944 | HORVU3Hr1G029560 | undescribed protein | -3.05 | AT1G61230.1 |
| 945 | HORVU1Hr1G070190 | Nodulin-like / Major Facilitator Superfamily protein | -3.06 | AT2G30300.1 |
| 946 | HORVU4Hr1G090780 | Disease resistance-responsive (dirigent-like protein) family protein | -3.07 | AT1G22900.1 |
| 947 | HORVU4Hr1G060970 | 50S ribosomal protein L4 | -3.07 | AT1G07320.2 |
| 948 | HORVU5Hr1G046480 | Chalcone--flavonone isomerase | -3.07 | AT5G05270.2 |
| 949 | HORVU2Hr1G101040 | calcium-transporting ATPase, putative | -3.07 | AT5G57110.3 |
| 950 | HORVU6Hr1G003160 | O-methyltransferase family protein | -3.07 | AT4G35160.1 |
| 951 | HORVU5Hr1G027740 | undescribed protein | -3.07 | AT5G55060.1 |
| 952 | HORVU6Hr1G093030 | 50S ribosomal protein L9 | -3.08 | AT3G44890.1 |
| 953 | HORVU7Hr1G041080 | SOUL heme-binding family protein | -3.08 | AT3G10130.1 |
| 954 | HORVU7Hr1G058940 | RNA-binding protein 1 | -3.08 | AT4G24770.2 |
| 955 | HORVU1Hr1G000430 | Chalcone--flavonone isomerase 1B-1 | -3.09 | AT1G53520.1 |
| 956 | HORVU5Hr1G050120 | undescribed protein | -3.09 | AT3G53210.2 |
| 957 | HORVU5Hr1G120250 | 30S ribosomal protein S6 | -3.09 | AT1G64510.1 |
| 958 | HORVU4Hr1G006200 | Phosphatidylinositol-4-phosphate 5-kinase family protein | -3.09 | AT1G60890.2 |
| 959 | HORVU1Hr1G028680 | Thioredoxin superfamily protein | -3.09 | AT4G29670.1 |
| 960 | HORVU6Hr1G068720 | carotenoid cleavage dioxygenase 1 | -3.09 | AT4G19170.1 |
| 961 | HORVU3Hr1G088580 | Retrotransposon protein, putative, unclassified | -3.09 | AT3G24255.3 |
| 962 | HORVU1Hr1G088270 | IMP dehydrogenase/GMP reductase | -3.10 | AT5G01930.1 |
| 963 | HORVU4Hr1G016510 | Chaperone protein DnaJ | -3.10 | AT4G09350.1 |
| 964 | HORVU4Hr1G025010 | undescribed protein | -3.11 | #N/A |
| 965 | HORVU6Hr1G049050 | Protein kinase superfamily protein | -3.11 | AT5G64940.2 |
| 966 | HORVU7Hr1G068610 | undescribed protein | -3.11 | AT5G41150.1 |
| 967 | HORVU2Hr1G091680 | Remorin family protein | -3.12 | AT5G23750.3 |
| 968 | HORVU6Hr1G065710 | SPX domain-containing membrane protein | -3.12 | AT4G11810.3 |
| 969 | HORVU6Hr1G085170 | Photosystem II 10 kDa polypeptide | -3.12 | AT1G79040.1 |
| 970 | HORVU6Hr1G075900 | Geranylgeranyl diphosphate reductase | -3.12 | AT1G74470.1 |
| 971 | HORVU7Hr1G038300 | Transmembrane amino acid transporter family protein | -3.12 | AT3G28960.5 |
| 972 | HORVU2Hr1G112450 | purple acid phosphatase 21 | -3.12 | AT3G52810.1 |
| 973 | HORVU3Hr1G007480 | unknown protein; BEST Arabidopsis thaliana protein match is: unknown protein . | -3.12 | AT1G16840.4 |
| 974 | HORVU2Hr1G115710 | unknown protein; LOCATED IN: chloroplast; EXPRESSED IN: 22 plant | -3.13 | AT2G21960.1 |

| | | | | |
|---|---|---|---|---|
| | | structures; EXPRESSED DURING: 13 growth stages; BEST Arabidopsis thaliana protein match is: unknown protein . | | |
| 975 | HORVU3Hr1G084660 | Phosphoglucan, water dikinase, chloroplastic | -3.14 | AT5G01260.3 |
| 976 | HORVU0Hr1G019860 | hypothetical protein chrM:11918-12241 FORWARD | -3.14 | ATMG00030.1 |
| 977 | HORVU4Hr1G074810 | D-ribulose-5-phosphate-3-epimerase | -3.14 | AT5G61410.2 |
| 978 | HORVU2Hr1G053620 | Senescence-associated protein, putative | -3.15 | AT1G27320.1 |
| 979 | HORVU2Hr1G119760 | Pentatricopeptide repeat-containing protein | -3.15 | AT1G50270.1 |
| 980 | HORVU6Hr1G015430 | alpha/beta-Hydrolases superfamily protein | -3.15 | AT5G17670.1 |
| 981 | HORVU1Hr1G061830 | Beta-propeller domain-containing protein, methanol dehydrogenase | -3.15 | AT1G54780.1 |
| 982 | HORVU5Hr1G100990 | Obg-like ATPase 1 | -3.15 | AT1G56050.1 |
| 983 | HORVU7Hr1G115040 | 30S ribosomal protein S17 | -3.15 | AT1G79850.1 |
| 984 | HORVU7Hr1G079190 | PSRP4 | -3.16 | AT2G38140.1 |
| 985 | HORVU7Hr1G086810 | GATA transcription factor 23 | -3.16 | AT5G56860.1 |
| 986 | HORVU3Hr1G089390 | Protein of unknown function (DUF1685) | -3.16 | AT2G31560.3 |
| 987 | HORVU6Hr1G070420 | fatty acid desaturase 2 | -3.16 | AT3G12120.2 |
| 988 | HORVU3Hr1G010030 | Repressor of RNA polymerase III transcription | -3.16 | AT5G13240.1 |
| 989 | HORVU7Hr1G114170 | 50S ribosomal protein L35 | -3.16 | AT2G24090.1 |
| 990 | HORVU6Hr1G033970 | glycerol dehydrogenase | -3.16 | AT3G21340.2 |
| 991 | HORVU2Hr1G005570 | 2-oxoglutarate (2OG) and Fe(II)-dependent oxygenase superfamily protein | -3.17 | AT1G06620.1 |
| 992 | HORVU5Hr1G001650 | Protein of unknown function (DUF1218) | -3.17 | AT5G17210.1 |
| 993 | HORVU4Hr1G024050 | NAD(P)-binding Rossmann-fold superfamily protein | -3.18 | AT2G39080.1 |
| 994 | HORVU7Hr1G002210 | Ferredoxin--NADP reductase | -3.18 | AT1G20020.3 |
| 995 | HORVU2Hr1G057700 | Oxygen-evolving enhancer protein 1, chloroplastic | -3.18 | AT3G50820.1 |
| 996 | HORVU4Hr1G060440 | Digalactosyldiacylglycerol synthase 2, chloroplastic | -3.18 | AT4G00550.2 |
| 997 | HORVU5Hr1G112580 | rRNA N-glycosidase | -3.18 | AT3G59490.3 |
| 998 | HORVU2Hr1G116160 | undescribed protein | -3.18 | #N/A |
| 999 | HORVU3Hr1G083470 | Chromosome 3B, genomic scaffold, cultivar Chinese Spring | -3.18 | AT2G41430.5 |
| 1000 | HORVU7Hr1G097670 | Macrophage migration inhibitory factor homolog | -3.19 | AT5G57170.1 |
| 1001 | HORVU0Hr1G031330 | RNAase | -3.20 | #N/A |
| 1002 | HORVU6Hr1G055960 | Bidirectional sugar transporter N3 | -3.20 | AT4G10850.1 |

| | | | | |
|---|---|---|---|---|
| 1003 | HORVU1Hr1G077020 | Magnesium transporter NIPA2 | -3.20 | AT1G34470.1 |
| 1004 | HORVU4Hr1G012530 | Plant protein of unknown function (DUF863) | -3.21 | AT5G67390.3 |
| 1005 | HORVU7Hr1G060130 | Superoxide dismutase [Cu-Zn] 4A | -3.22 | AT2G28190.1 |
| 1006 | HORVU4Hr1G071730 | Inositol-3-phosphate synthase | -3.22 | AT2G22240.2 |
| 1007 | HORVU6Hr1G054110 | Protein HHL1, chloroplastic | -3.23 | AT1G67700.4 |
| 1008 | HORVU2Hr1G060880 | Oxygen-evolving enhancer protein 2, chloroplastic | -3.23 | AT1G06680.1 |
| 1009 | HORVU1Hr1G050130 | Mitochondrial substrate carrier family protein | -3.24 | AT5G46800.3 |
| 1010 | HORVU1Hr1G057680 | beta-1,3-glucanase_putative | -3.24 | AT4G16260.1 |
| 1011 | HORVU6Hr1G072200 | DOF zinc finger protein 1 | -3.24 | AT5G60200.1 |
| 1012 | HORVU5Hr1G065330 | cinnamoyl coa reductase 1 | -3.24 | AT1G15950.3 |
| 1013 | HORVU5Hr1G114990 | GDP-mannose transporter | -3.24 | AT2G13650.3 |
| 1014 | HORVU4Hr1G010430 | BEL1-type homeodomain protein | -3.25 | AT2G23760.3 |
| 1015 | HORVU6Hr1G091360 | OTU domain-containing protein 5-B | -3.25 | AT5G04250.2 |
| 1016 | HORVU3Hr1G018110 | Protein phosphatase PTC7 homolog | -3.25 | AT2G30170.1 |
| 1017 | HORVU5Hr1G080980 | 5-keto-4-deoxy-D-glucarate aldolase | -3.25 | AT4G10750.1 |
| 1018 | HORVU3Hr1G032490 | Triose phosphate/phosphate translocator, chloroplastic | -3.26 | AT5G46110.4 |
| 1019 | HORVU5Hr1G053270 | Rhodanese-like domain-containing protein 10 | -3.27 | AT2G42220.1 |
| 1020 | HORVU3Hr1G049900 | glyoxylate reductase 2 | -3.27 | AT1G17650.1 |
| 1021 | HORVU7Hr1G088160 | AP-3 complex subunit mu-1 | -3.27 | AT1G58230.3 |
| 1022 | HORVU0Hr1G025070 | Retrotransposon protein | -3.27 | AT2G07706.1 |
| 1023 | HORVU4Hr1G074990 | DOF zinc finger protein 1 | -3.27 | AT5G39660.3 |
| 1024 | HORVU6Hr1G018830 | 50S ribosomal protein L3 | -3.28 | AT2G43030.1 |
| 1025 | HORVU1Hr1G013250 | undescribed protein | -3.29 | AT1G23450.1 |
| 1026 | HORVU6Hr1G091260 | Auxin-responsive protein IAA10 | -3.29 | AT2G33310.2 |
| 1027 | HORVU3Hr1G006150 | unknown protein; INVOLVED IN: biological_process unknown; LOCATED IN: chloroplast; EXPRESSED IN: 22 plant structures; EXPRESSED DURING: 13 growth stages; Has 74 Blast hits to 74 proteins in 29 species: Archae - 0; Bacteria - 19; Metazoa - 0; Fungi - 0; Plants - 49; Viruses - 0; Other Eukaryotes - 6 (source: NCBI BLink). | -3.30 | AT1G18060.1 |
| 1028 | HORVU5Hr1G009200 | NAD-dependent epimerase/dehydratase | -3.30 | AT1G09340.1 |
| 1029 | HORVU4Hr1G085050 | purple acid phosphatase 27 | -3.30 | AT5G50400.1 |
| 1030 | HORVU7Hr1G121630 | UPF0392 protein | -3.30 | AT4G37420.1 |
| 1031 | HORVU4Hr1G062920 | Protein disulfide-isomerase LQY1, chloroplastic | -3.30 | AT1G75690.1 |
| 1032 | HORVU4Hr1G004210 | ferredoxin 3 | -3.30 | AT1G32550.1 |

| | | | | |
|---|---|---|---|---|
| 1033 | HORVU6Hr1G016070 | Sodium Bile acid symporter family | -3.31 | AT3G25410.1 |
| 1034 | HORVU4Hr1G089990 | Protein WEAK CHLOROPLAST MOVEMENT UNDER BLUE LIGHT 1 | -3.32 | AT2G26570.2 |
| 1035 | HORVU4Hr1G004590 | Inositol hexakisphosphate and diphosphoinositol-pentakisphosphate kinase | -3.32 | AT5G15070.4 |
| 1036 | HORVU7Hr1G062930 | Peptidyl-prolyl cis-trans isomerase, chloroplastic | -3.33 | AT3G01480.1 |
| 1037 | HORVU1Hr1G012110 | xylulose kinase-1 | -3.33 | AT2G21370.1 |
| 1038 | HORVU3Hr1G082730 | Photosystem II 22 kDa protein, chloroplastic | -3.34 | AT1G44575.3 |
| 1039 | HORVU4Hr1G084830 | 50S ribosomal protein L11 | -3.34 | AT1G32990.1 |
| 1040 | HORVU2Hr1G115080 | Retrotransposon protein, putative, unclassified | -3.34 | AT4G20520.1 |
| 1041 | HORVU5Hr1G025960 | acyl carrier protein 5 | -3.35 | AT4G25050.1 |
| 1042 | HORVU2Hr1G105930 | RNA-binding protein 28 | -3.35 | AT2G35410.1 |
| 1043 | HORVU6Hr1G078500 | fucosyltransferase 1 | -3.36 | AT2G03220.1 |
| 1044 | HORVU4Hr1G062890 | Nuclear transcription factor Y subunit C-4 | -3.36 | AT5G63470.2 |
| 1045 | HORVU6Hr1G090160 | unknown function | -3.37 | AT5G40810.2 |
| 1046 | HORVU3Hr1G085270 | Fructose-1,6-bisphosphatase class 1 | -3.37 | AT3G55800.1 |
| 1047 | HORVU3Hr1G098290 | methyl esterase 3 | -3.37 | AT2G23610.1 |
| 1048 | HORVU1Hr1G046400 | Non-specific lipid-transfer protein 2 | -3.38 | AT5G38170.1 |
| 1049 | HORVU0Hr1G010770 | undescribed protein | -3.38 | AT3G26790.1 |
| 1050 | HORVU3Hr1G011620 | Pentapeptide repeat-containing protein | -3.39 | AT2G44920.2 |
| 1051 | HORVU1Hr1G091850 | unknown function | -3.40 | AT2G42900.1 |
| 1052 | HORVU6Hr1G073180 | methyltransferase type 11 | -3.40 | AT3G18000.1 |
| 1053 | HORVU0Hr1G024890 | RNAase | -3.40 | #N/A |
| 1054 | HORVU2Hr1G062850 | unknown function | -3.40 | AT4G33625.3 |
| 1055 | HORVU6Hr1G075280 | ATP synthase delta-subunit gene | -3.41 | AT4G09650.1 |
| 1056 | HORVU4Hr1G087350 | UPF0187 protein YneE | -3.41 | AT3G61320.1 |
| 1057 | HORVU4Hr1G006730 | Plastid-lipid associated protein PAP / fibrillin family protein | -3.42 | AT3G26070.1 |
| 1058 | HORVU5Hr1G111640 | 50S ribosomal protein L17 | -3.42 | AT3G54210.1 |
| 1059 | HORVU7Hr1G055890 | 50S ribosomal protein 6, chloroplastic | -3.42 | AT5G17870.1 |
| 1060 | HORVU4Hr1G085930 | Protein kinase superfamily protein | -3.42 | AT3G61080.1 |
| 1061 | HORVU7Hr1G026100 | undescribed protein | -3.42 | AT3G54920.1 |
| 1062 | HORVU3Hr1G083790 | unknown function | -3.43 | AT2G36145.1 |
| 1063 | HORVU7Hr1G101810 | 50S ribosomal protein L24 | -3.43 | AT5G54600.1 |
| 1064 | HORVU5Hr1G048240 | fatty acid desaturase 8 | -3.44 | AT5G05580.1 |
| 1065 | HORVU5Hr1G124160 | Chlorophyll a-b binding protein 2, chloroplastic | -3.44 | AT2G34420.1 |
| 1066 | HORVU2Hr1G009850 | Thylakoid lumenal 15.0 kDa protein 2, chloroplastic | -3.45 | AT5G52970.2 |

| | | | | |
|---|---|---|---|---|
| 1067 | HORVU3Hr1G082280 | Glutamine--fructose-6-phosphate aminotransferase [isomerizing] | -3.45 | AT2G16570.1 |
| 1068 | HORVU2Hr1G103130 | Aminomethyltransferase | -3.46 | AT1G11860.2 |
| 1069 | HORVU2Hr1G041610 | Cytochrome b6-f complex iron-sulfur subunit | -3.47 | AT4G03280.1 |
| 1070 | HORVU3Hr1G093990 | Haloalkane dehalogenase 2 | -3.47 | AT4G12830.1 |
| 1071 | HORVU7Hr1G037600 | sodium/calcium exchanger family protein / calcium-binding EF hand family protein | -3.47 | AT1G53210.1 |
| 1072 | HORVU1Hr1G072180 | Riboflavin biosynthesis protein RibBA | -3.47 | AT5G59750.1 |
| 1073 | HORVU5Hr1G022570 | Stress responsive alpha-beta barrel domain protein | -3.51 | AT2G32500.2 |
| 1074 | HORVU6Hr1G004860 | 3-beta hydroxysteroid dehydrogenase | -3.51 | AT3G63140.1 |
| 1075 | HORVU2Hr1G001600 | Pyruvate dehydrogenase E1 component subunit alpha | -3.51 | AT1G01090.1 |
| 1076 | HORVU1Hr1G087570 | FASCICLIN-like arabinogalactan 6 | -3.51 | AT2G20520.1 |
| 1077 | HORVU7Hr1G122750 | BnaA09g56450D protein | -3.51 | AT1G16080.1 |
| 1078 | HORVU7Hr1G046320 | Chlorophyll a-b binding protein 1B-21, chloroplastic | -3.51 | AT3G54890.1 |
| 1079 | HORVU6Hr1G009170 | NAD(P)-binding Rossmann-fold superfamily protein | -3.51 | AT1G72640.1 |
| 1080 | HORVU5Hr1G058300 | trehalose-6-phosphate phosphatase | -3.52 | AT5G65140.1 |
| 1081 | HORVU3Hr1G095700 | metacaspase 1 | -3.52 | AT1G02170.1 |
| 1082 | HORVU3Hr1G039790 | 50S ribosomal protein L40 | -3.52 | AT3G56910.1 |
| 1083 | HORVU3Hr1G025020 | Chromosome 3B, genomic scaffold, cultivar Chinese Spring | -3.52 | AT5G19875.1 |
| 1084 | HORVU2Hr1G090400 | Protein of unknown function (DUF177) | -3.53 | AT3G19800.2 |
| 1085 | HORVU3Hr1G050430 | protein kinase family protein | -3.53 | AT5G12000.1 |
| 1086 | HORVU2Hr1G042370 | cellulose-synthase like D2 | -3.54 | AT5G16910.1 |
| 1087 | HORVU7Hr1G042480 | aluminum-activated malate transporter 9 | -3.54 | AT3G18440.1 |
| 1088 | HORVU1Hr1G060610 | 1-deoxy-D-xylulose 5-phosphate synthase 1 | -3.54 | AT4G15560.1 |
| 1089 | HORVU5Hr1G118680 | Mitochondrial transcription termination factor family protein | -3.55 | AT4G09620.2 |
| 1090 | HORVU7Hr1G000900 | plastocyanin 1 | -3.56 | AT1G76100.1 |
| 1091 | HORVU7Hr1G039140 | Glutathione S-transferase family protein | -3.56 | AT3G62760.1 |
| 1092 | HORVU3Hr1G036600 | TRAM, LAG1 and CLN8 (TLC) lipid-sensing domain containing protein | -3.58 | AT1G35180.1 |
| 1093 | HORVU5Hr1G043090 | Two-component response regulator ARR8 | -3.58 | AT2G41310.1 |
| 1094 | HORVU4Hr1G087610 | Transcription factor bHLH35 | -3.61 | AT5G57150.5 |
| 1095 | HORVU6Hr1G000090 | sulfurtransferase protein 16 | -3.61 | AT2G17850.1 |
| 1096 | HORVU1Hr1G051700 | Protein CURVATURE THYLAKOID 1B, chloroplastic | -3.62 | AT2G46820.2 |

| | | | | |
|---|---|---|---|---|
| 1097 | HORVU0Hr1G001170 | unknown function | -3.62 | AT3G51510.1 |
| 1098 | HORVU4Hr1G087230 | Ectonucleoside triphosphate diphosphohydrolase 5 | -3.63 | AT5G18280.1 |
| 1099 | HORVU7Hr1G077740 | ferredoxin 1 | -3.64 | AT1G60950.1 |
| 1100 | HORVU2Hr1G113050 | sulfate transporter 91 | -3.64 | AT1G23090.1 |
| 1101 | HORVU2Hr1G037550 | tubulin alpha-4 chain | -3.65 | AT5G19780.1 |
| 1102 | HORVU5Hr1G019000 | FAR1-related sequence 5 | -3.65 | AT4G38180.1 |
| 1103 | HORVU4Hr1G031320 | methionine S-methyltransferase | -3.65 | AT5G49810.1 |
| 1104 | HORVU0Hr1G003270 | Oxygen-evolving enhancer protein 3-1, chloroplastic | -3.66 | AT4G21280.1 |
| 1105 | HORVU6Hr1G076040 | Photosynthetic NDH subcomplex B 4 | -3.66 | AT1G18730.5 |
| 1106 | HORVU2Hr1G070440 | ACT domain repeat 4 | -3.66 | AT1G69040.2 |
| 1107 | HORVU5Hr1G111330 | unknown function | -3.66 | AT4G13500.1 |
| 1108 | HORVU2Hr1G113260 | N-terminal nucleophile aminohydrolases (Ntn hydrolases) superfamily protein | -3.66 | AT3G16150.1 |
| 1109 | HORVU6Hr1G033160 | chlorophyll A/B binding protein 3 | -3.67 | AT1G61520.3 |
| 1110 | HORVU2Hr1G065110 | D-inositol 3-phosphate glycosyltransferase | -3.67 | AT5G01220.1 |
| 1111 | HORVU2Hr1G039020 | unknown function | -3.67 | AT2G29750.1 |
| 1112 | HORVU1Hr1G013740 | Dihydrolipoyl dehydrogenase | -3.67 | AT1G48030.5 |
| 1113 | HORVU4Hr1G067320 | SprT-like domain-containing protein Spartan | -3.67 | AT3G24340.1 |
| 1114 | HORVU7Hr1G073860 | 2-succinylbenzoate--CoA ligase | -3.68 | AT1G30520.2 |
| 1115 | HORVU5Hr1G087250 | Chlorophyll a-b binding protein 2, chloroplastic | -3.69 | AT2G34420.1 |
| 1116 | HORVU7Hr1G085980 | Salt overly sensitive 1 | -3.69 | AT2G01980.1 |
| 1117 | HORVU3Hr1G073740 | unknown protein; LOCATED IN: chloroplast; Has 35333 Blast hits to 34131 proteins in 2444 species: Archae - 798; Bacteria - 22429; Metazoa - 974; Fungi - 991; Plants - 531; Viruses - 0; Other Eukaryotes - 9610 (source: NCBI BLink). | -3.70 | AT5G55570.1 |
| 1118 | HORVU1Hr1G064750 | EG2771 | -3.70 | AT5G14910.2 |
| 1119 | HORVU6Hr1G011990 | unknown function | -3.70 | AT3G61870.2 |
| 1120 | HORVU3Hr1G071530 | 50S ribosomal protein L13 | -3.71 | AT1G78630.1 |
| 1121 | HORVU6Hr1G025170 | undescribed protein | -3.71 | #N/A |
| 1122 | HORVU7Hr1G018800 | undescribed protein | -3.71 | AT5G64440.1 |
| 1123 | HORVU2Hr1G123750 | glutathione peroxidase 6 | -3.72 | AT4G11600.1 |
| 1124 | HORVU7Hr1G038770 | dehydroascorbate reductase 1 | -3.73 | AT5G16710.1 |
| 1125 | HORVU2Hr1G038570 | Ribosome-recycling factor | -3.73 | AT3G63190.1 |
| 1126 | HORVU7Hr1G083690 | Two-component response regulator-like APRR2 | -3.73 | AT2G20570.1 |

| | | | | |
|---|---|---|---|---|
| 1127 | HORVU6Hr1G032680 | FKBP-like peptidyl-prolyl cis-trans isomerase family protein | -3.74 | AT4G26555.1 |
| 1128 | HORVU6Hr1G095380 | multidrug MFS transporter | -3.74 | AT2G21280.2 |
| 1129 | HORVU0Hr1G024880 | Conserved protein | -3.74 | AT2G26790.2 |
| 1130 | HORVU3Hr1G009210 | Photosystem I reaction center subunit XI | -3.74 | AT4G12800.2 |
| 1131 | HORVU2Hr1G070500 | Unknow protein | -3.74 | AT1G33040.1 |
| 1132 | HORVU3Hr1G065570 | undescribed protein | -3.75 | #N/A |
| 1133 | HORVU2Hr1G066800 | undescribed protein | -3.77 | AT5G24160.1 |
| 1134 | HORVU7Hr1G111190 | D-3-phosphoglycerate dehydrogenase | -3.77 | AT4G34200.1 |
| 1135 | HORVU5Hr1G051010 | ribulose bisphosphate carboxylase small chain 1A | -3.77 | AT1G67090.1 |
| 1136 | HORVU5Hr1G053060 | BnaA04g27590D protein | -3.77 | AT3G57990.1 |
| 1137 | HORVU6Hr1G076880 | Glycine cleavage system H protein | -3.78 | AT1G32470.1 |
| 1138 | HORVU4Hr1G042560 | undescribed protein | -3.78 | AT1G32960.1 |
| 1139 | HORVU5Hr1G050510 | kinesin 4 | -3.79 | AT5G47820.2 |
| 1140 | HORVU5Hr1G101020 | ribose-5-phosphate isomerase 2 | -3.79 | AT5G44520.2 |
| 1141 | HORVU6Hr1G063110 | undescribed protein | -3.79 | AT5G64740.1 |
| 1142 | HORVU3Hr1G078780 | Ribosomal protein L34 | -3.79 | AT1G29070.1 |
| 1143 | HORVU3Hr1G105790 | Glucan endo-1,3-beta-glucosidase GI | -3.80 | AT3G57270.1 |
| 1144 | HORVU5Hr1G099020 | Leucine carboxyl methyltransferase | -3.80 | AT5G42760.2 |
| 1145 | HORVU7Hr1G069410 | undescribed protein | -3.81 | AT1G15640.2 |
| 1146 | HORVU0Hr1G024560 | Ribulose bisphosphate carboxylase large chain | -3.82 | ATCG00490.1 |
| 1147 | HORVU0Hr1G017890 | undescribed protein | -3.82 | AT5G63630.2 |
| 1148 | HORVU4Hr1G071070 | Dehydrodolichyl diphosphate syntase complex subunit NUS1 | -3.82 | AT1G11755.1 |
| 1149 | HORVU5Hr1G035300 | P-loop containing nucleoside triphosphate hydrolases superfamily protein | -3.82 | AT2G46620.1 |
| 1150 | HORVU1Hr1G054240 | expansin B2 | -3.83 | AT1G65680.1 |
| 1151 | HORVU3Hr1G087420 | DNA-3-methyladenine glycosylase 1 | -3.83 | AT1G13635.2 |
| 1152 | HORVU6Hr1G021460 | Zinc finger protein CONSTANS-LIKE 7 | -3.84 | AT5G57180.2 |
| 1153 | HORVU1Hr1G047710 | Plant invertase/pectin methylesterase inhibitor superfamily protein | -3.85 | AT5G62360.1 |
| 1154 | HORVU4Hr1G059950 | ROTUNDIFOLIA like 7 | -3.85 | AT2G39705.1 |
| 1155 | HORVU5Hr1G044410 | Hydroxymethylglutaryl-CoA lyase | -3.87 | AT2G26800.1 |
| 1156 | HORVU1Hr1G045520 | Thylakoid lumenal 17.4 kDa protein, chloroplastic | -3.87 | AT5G53490.2 |
| 1157 | HORVU0Hr1G034360 | undescribed protein | -3.88 | AT3G20460.1 |
| 1158 | HORVU4Hr1G009980 | unknown function | -3.90 | AT1G44920.1 |
| 1159 | HORVU5Hr1G115000 | Thylakoid soluble phosphoprotein TSP9 protein | -3.91 | AT3G47070.1 |
| 1160 | HORVU2Hr1G111300 | Bifunctional lysine-specific demethylase and histidyl-hydroxylase NO66 | -3.92 | AT3G11560.5 |

| | | | | |
|---|---|---|---|---|
| 1161 | HORVU1Hr1G035870 | undescribed protein | -3.93 | AT4G15760.1 |
| 1162 | HORVU5Hr1G098340 | 30S ribosomal protein S9 | -3.93 | AT1G74970.1 |
| 1163 | HORVU0Hr1G012860 | Alcohol dehydrogenase | -3.96 | AT4G37980.1 |
| 1164 | HORVU4Hr1G018070 | Auxilin-related protein 2 | -3.96 | AT1G04160.2 |
| 1165 | HORVU5Hr1G066280 | Chlorophyll a-b binding protein 1B-20, chloroplastic | -3.96 | AT3G47470.1 |
| 1166 | HORVU2Hr1G041280 | Chlorophyll a-b binding protein CP29 | -3.97 | AT3G08940.1 |
| 1167 | HORVU7Hr1G121210 | At5g37260-like protein | -3.98 | AT5G37260.1 |
| 1168 | HORVU3Hr1G094270 | Chromosome 3B, genomic scaffold, cultivar Chinese Spring | -3.99 | AT2G35450.1 |
| 1169 | HORVU7Hr1G115540 | Pyrophosphate-energized vacuolar membrane proton pump | -4.00 | AT1G15690.1 |
| 1170 | HORVU3Hr1G113920 | undescribed protein | -4.01 | AT5G04895.1 |
| 1171 | HORVU3Hr1G089370 | Protein of unknown function (DUF1685) | -4.01 | AT1G05870.9 |
| 1172 | HORVU3Hr1G081180 | expansin A1 | -4.02 | AT1G69530.2 |
| 1173 | HORVU1Hr1G043820 | alpha-galactosidase 2 | -4.02 | AT5G08370.1 |
| 1174 | HORVU7Hr1G085350 | F-box/kelch-repeat protein | -4.03 | AT1G80440.1 |
| 1175 | HORVU5Hr1G062240 | Chlorophyll a-b binding protein 7, chloroplastic | -4.04 | AT1G19150.1 |
| 1176 | HORVU3Hr1G013970 | NAD(P)-binding Rossmann-fold superfamily protein | -4.05 | AT2G37540.1 |
| 1177 | HORVU2Hr1G073760 | Peroxiredoxin-2 | -4.07 | AT5G06290.1 |
| 1178 | HORVU7Hr1G097250 | Peptidyl-prolyl cis-trans isomerase B | -4.07 | AT5G35100.3 |
| 1179 | HORVU2Hr1G062290 | ORF58e | -4.07 | AT1G17250.1 |
| 1180 | HORVU5Hr1G096260 | UDP-Glycosyltransferase superfamily protein | -4.08 | AT3G02100.1 |
| 1181 | HORVU7Hr1G030800 | Pco114914 | -4.09 | AT2G28630.2 |
| 1182 | HORVU2Hr1G001500 | undescribed protein | -4.10 | AT4G14500.2 |
| 1183 | HORVU4Hr1G048990 | undescribed protein | -4.10 | AT5G20220.3 |
| 1184 | HORVU3Hr1G089650 | Rhodanese/Cell cycle control phosphatase superfamily protein | -4.10 | AT3G59780.1 |
| 1185 | HORVU1Hr1G067300 | phosphoglycerate kinase 1 | -4.10 | AT3G12780.1 |
| 1186 | HORVU6Hr1G092750 | C2H2-like zinc finger protein | -4.12 | AT5G04390.1 |
| 1187 | HORVU1Hr1G001900 | undescribed protein | -4.12 | AT1G21640.1 |
| 1188 | HORVU2Hr1G060480 | Photosystem I reaction center subunit psaK, chloroplastic | -4.12 | AT1G30380.1 |
| 1189 | HORVU1Hr1G054190 | unknown function | -4.13 | AT5G65750.1 |
| 1190 | HORVU5Hr1G109440 | Lactoylglutathione lyase | -4.13 | AT1G67280.1 |
| 1191 | HORVU3Hr1G029260 | Peptide-N4-(N-acetyl-beta-glucosaminyl)asparagine amidase A protein | -4.14 | AT3G14920.1 |
| 1192 | HORVU2Hr1G071360 | undescribed protein | -4.14 | AT2G21880.1 |
| 1193 | HORVU7Hr1G042160 | Protein CURVATURE THYLAKOID 1A, chloroplastic | -4.14 | AT4G01150.1 |

| | | | | |
|---|---|---|---|---|
| 1194 | HORVU1Hr1G022840 | Peroxisomal membrane protein 2 | -4.14 | AT1G52870.2 |
| 1195 | HORVU3Hr1G095330 | 50S ribosomal protein L27 | -4.15 | AT5G40950.1 |
| 1196 | HORVU7Hr1G056240 | Peptidyl-prolyl cis-trans isomerase | -4.16 | AT2G43560.1 |
| 1197 | HORVU2Hr1G044790 | undescribed protein | -4.17 | AT1G52820.1 |
| 1198 | HORVU2Hr1G080480 | 30S ribosomal protein 3, chloroplastic | -4.17 | AT1G68590.2 |
| 1199 | HORVU4Hr1G009520 | tubulin alpha-4 chain | -4.18 | AT4G14960.2 |
| 1200 | HORVU3Hr1G093310 | Transcription factor bHLH62 | -4.19 | AT3G07340.1 |
| 1201 | HORVU7Hr1G096140 | Thylakoid lumenal 16.5 kDa protein, chloroplastic | -4.20 | AT4G02530.1 |
| 1202 | HORVU5Hr1G059140 | unknown protein; BEST Arabidopsis thaliana protein match is: unknown protein . | -4.20 | AT4G38060.1 |
| 1203 | HORVU5Hr1G100140 | Photosystem I reaction center subunit III | -4.22 | AT1G31330.1 |
| 1204 | HORVU3Hr1G091140 | NAD(P)H-quinone oxidoreductase subunit N | -4.22 | AT5G58260.1 |
| 1205 | HORVU1Hr1G045590 | Glutamyl-tRNA reductase family protein | -4.22 | AT1G58290.1 |
| 1206 | HORVU2Hr1G102620 | unknown function | -4.23 | AT5G17165.1 |
| 1207 | HORVU3Hr1G075870 | Photosystem II reaction center W protein, chloroplastic | -4.23 | AT2G30570.1 |
| 1208 | HORVU1Hr1G011520 | Glutamyl-tRNA reductase family protein | -4.23 | AT1G58290.1 |
| 1209 | HORVU4Hr1G083230 | S-adenosyl-L-methionine-dependent methyltransferases superfamily protein | -4.24 | AT5G44600.1 |
| 1210 | HORVU2Hr1G089030 | Plastid transcriptionally active 5, putative isoform 1 | -4.25 | AT4G13670.1 |
| 1211 | HORVU4Hr1G090720 | Jacalin-related lectin 34 | -4.26 | AT3G16460.2 |
| 1212 | HORVU0Hr1G032630 | undescribed protein | -4.26 | AT5G56400.1 |
| 1213 | HORVU6Hr1G010940 | undescribed protein | -4.27 | AT4G16120.1 |
| 1214 | HORVU5Hr1G005740 | PHYTOENE SYNTHASE | -4.28 | AT5G17230.4 |
| 1215 | HORVU4Hr1G073730 | Protein of unknown function (DUF506) | -4.31 | AT2G38820.2 |
| 1216 | HORVU2Hr1G106880 | chloroplastic lipocalin | -4.31 | AT3G47860.1 |
| 1217 | HORVU7Hr1G009800 | LEA | -4.31 | AT1G32560.1 |
| 1218 | HORVU1Hr1G023160 | Flavin-containing monooxygenase family protein | -4.32 | AT1G19250.1 |
| 1219 | HORVU3Hr1G088020 | dihydroflavonol 4-reductase-like1 | -4.32 | AT5G58490.1 |
| 1220 | HORVU2Hr1G113270 | Isoaspartyl peptidase/L-asparaginase | -4.32 | AT3G16150.1 |
| 1221 | HORVU5Hr1G124380 | Light-repressed protein A | -4.32 | AT5G24490.1 |
| 1222 | HORVU5Hr1G014300 | Auxin-responsive protein IAA30 | -4.34 | AT3G04730.1 |
| 1223 | HORVU1Hr1G050830 | dihydroflavonol 4-reductase-like1 | -4.36 | AT2G23910.1 |
| 1224 | HORVU5Hr1G060250 | annexin 5 | -4.36 | AT1G68090.1 |
| 1225 | HORVU6Hr1G053730 | Protein GLE1 | -4.37 | AT1G13120.1 |
| 1226 | HORVU2Hr1G116390 | Chalcone synthase 1 | -4.37 | AT5G13930.1 |
| 1227 | HORVU5Hr1G098980 | HAD-superfamily hydrolase, subfamily IG, 5'-nucleotidase | -4.38 | AT1G75210.3 |

| | | | | |
|---|---|---|---|---|
| 1228 | HORVU4Hr1G073650 | Heat stress transcription factor A-2d | -4.39 | AT3G22830.1 |
| 1229 | HORVU2Hr1G073370 | photosystem I subunit O | -4.40 | AT1G08380.1 |
| 1230 | HORVU7Hr1G117710 | O-methyltransferase family protein | -4.40 | AT4G35160.1 |
| 1231 | HORVU7Hr1G006880 | undescribed protein | -4.41 | AT3G60380.1 |
| 1232 | HORVU3Hr1G078940 | tubulin beta chain 2 | -4.41 | AT5G12250.1 |
| 1233 | HORVU0Hr1G021630 | Auxin-responsive protein IAA21 | -4.43 | AT4G29080.1 |
| 1234 | HORVU1Hr1G075580 | oxidative stress 3 . | -4.44 | AT5G56550.1 |
| 1235 | HORVU1Hr1G045080 | PPR containing protein, putative | -4.46 | AT1G28100.6 |
| 1236 | HORVU0Hr1G007360 | Chlorophyllide a oxygenase, chloroplastic | -4.46 | AT1G44446.1 |
| 1237 | HORVU6Hr1G056000 | CONSTANS-like 3 | -4.47 | AT5G24930.1 |
| 1238 | HORVU3Hr1G000900 | Peptidyl-prolyl cis-trans isomerase B | -4.48 | AT1G74070.1 |
| 1239 | HORVU3Hr1G059330 | undescribed protein | -4.48 | #N/A |
| 1240 | HORVU2Hr1G023560 | Magnesium-chelatase subunit H | -4.49 | AT5G13630.1 |
| 1241 | HORVU0Hr1G001490 | Photosystem I reaction center subunit VI, chloroplastic | -4.50 | AT1G52230.1 |
| 1242 | HORVU5Hr1G050990 | ribulose bisphosphate carboxylase small chain 1A | -4.50 | AT1G67090.1 |
| 1243 | HORVU1Hr1G004400 | unknown function | -4.51 | AT5G02160.1 |
| 1244 | HORVU7Hr1G069520 | RNA polymerase sigma-C factor | -4.54 | AT1G64860.1 |
| 1245 | HORVU1Hr1G076120 | Protein of unknown function (DUF506) | -4.56 | AT4G32480.1 |
| 1246 | HORVU2Hr1G085300 | haloacid dehalogenase-like hydrolase family protein | -4.56 | AT4G39970.1 |
| 1247 | HORVU5Hr1G032350 | Chromosome 3B, genomic scaffold, cultivar Chinese Spring | -4.57 | AT3G55270.1 |
| 1248 | HORVU1Hr1G005120 | outer plastid envelope protein 16-1 | -4.57 | AT2G28900.1 |
| 1249 | HORVU5Hr1G109710 | Photosystem I reaction center subunit II | -4.58 | AT1G03130.1 |
| 1250 | HORVU3Hr1G089380 | Protein of unknown function (DUF1685) | -4.58 | AT1G05870.9 |
| 1251 | HORVU2Hr1G079630 | Protein OBERON 3 | -4.59 | AT1G14740.1 |
| 1252 | HORVU4Hr1G076990 | undescribed protein | -4.61 | AT5G26670.1 |
| 1253 | HORVU3Hr1G039930 | Magnesium-protoporphyrin IX monomethyl ester [oxidative] cyclase 3 | -4.61 | AT3G56940.1 |
| 1254 | HORVU3Hr1G094820 | thylakoid rhodanese-like | -4.63 | AT3G25480.1 |
| 1255 | HORVU2Hr1G074280 | unknown function | -4.63 | AT5G09820.2 |
| 1256 | HORVU1Hr1G073010 | unknown function | -4.64 | AT3G54000.3 |
| 1257 | HORVU2Hr1G049560 | FKBP-like peptidyl-prolyl cis-trans isomerase family protein | -4.64 | AT1G20810.2 |
| 1258 | HORVU6Hr1G049120 | Photosystem II reaction center protein M | -4.65 | ATCG00220.1 |
| 1259 | HORVU6Hr1G071520 | Eukaryotic aspartyl protease family protein | -4.69 | AT5G10770.1 |
| 1260 | HORVU2Hr1G077650 | ferric reduction oxidase 7 | -4.70 | AT5G49740.3 |
| 1261 | HORVU2Hr1G036960 | Chlorophyll a-b binding protein 7, chloroplastic | -4.70 | AT3G61470.1 |

| | | | | |
|---|---|---|---|---|
| 1262 | HORVU2Hr1G121640 | Chromosome 3B, genomic scaffold, cultivar Chinese Spring | -4.71 | AT5G26040.1 |
| 1263 | HORVU2Hr1G095080 | Cytochrome P450 superfamily protein | -4.71 | AT1G12740.1 |
| 1264 | HORVU2Hr1G042030 | undescribed protein | -4.75 | #N/A |
| 1265 | HORVU5Hr1G103440 | zinc-finger protein 1 | -4.76 | AT5G48890.2 |
| 1266 | HORVU4Hr1G013420 | BTB/POZ domain-containing protein | -4.77 | AT2G14820.2 |
| 1267 | HORVU3Hr1G025430 | undescribed protein | -4.78 | AT3G50370.1 |
| 1268 | HORVU7Hr1G118130 | Cytokinin dehydrogenase 11 | -4.80 | AT5G21482.1 |
| 1269 | HORVU5Hr1G113900 | Photosystem I reaction center subunit IV | -4.81 | AT2G20260.1 |
| 1270 | HORVU3Hr1G059950 | undescribed protein | -4.82 | AT5G47490.2 |
| 1271 | HORVU5Hr1G018680 | undescribed protein | -4.83 | AT5G56220.2 |
| 1272 | HORVU2Hr1G066020 | undescribed protein | -4.83 | AT3G55380.2 |
| 1273 | HORVU4Hr1G049000 | undescribed protein | -4.86 | AT5G13700.1 |
| 1274 | HORVU1Hr1G091540 | Transcription factor RADIALIS | -4.87 | AT4G39250.1 |
| 1275 | HORVU1Hr1G057640 | peptide chain release factor, putative | -4.88 | AT3G57190.1 |
| 1276 | HORVU5Hr1G043200 | strictosidine synthase-like 2 | -4.90 | AT3G57030.1 |
| 1277 | HORVU7Hr1G052750 | Cytochrome b561 and DOMON domain-containing protein | -4.91 | AT3G25290.2 |
| 1278 | HORVU6Hr1G075080 | FKBP-like peptidyl-prolyl cis-trans isomerase family protein | -4.92 | AT4G39710.2 |
| 1279 | HORVU3Hr1G057090 | carbonic anhydrase 2 | -4.92 | AT5G14740.9 |
| 1280 | HORVU1Hr1G087320 | xyloglucan endotransglucosylase/hydrolase 25 | -4.96 | AT5G57550.1 |
| 1281 | HORVU7Hr1G023140 | Photosynthetic NDH subunit of subcomplex B 3, chloroplastic | -4.97 | AT3G16250.1 |
| 1282 | HORVU2Hr1G108430 | undescribed protein | -4.98 | #N/A |
| 1283 | HORVU4Hr1G045210 | Mitochondrial transcription termination factor family protein | -4.99 | AT2G36000.1 |
| 1284 | HORVU1Hr1G035720 | ribulose bisphosphate carboxylase small chain 1A | -5.00 | AT1G67090.1 |
| 1285 | HORVU3Hr1G005500 | Transcription factor bHLH35 | -5.00 | AT5G57150.5 |
| 1286 | HORVU6Hr1G015120 | undescribed protein | -5.01 | AT1G78080.1 |
| 1287 | HORVU2Hr1G040780 | Chlorophyll a-b binding protein 13, chloroplastic | -5.02 | AT5G54270.1 |
| 1288 | HORVU4Hr1G002050 | 2-methyl-6-phytyl-1,4-hydroquinone methyltransferase 2, chloroplastic | -5.04 | AT3G63410.1 |
| 1289 | HORVU3Hr1G083360 | Fructose-1,6-bisphosphatase class 1 | -5.06 | AT1G43670.1 |
| 1290 | HORVU4Hr1G014080 | enzyme binding;tetrapyrrole binding | -5.06 | AT3G59400.1 |
| 1291 | HORVU7Hr1G069590 | undescribed protein | -5.09 | AT5G49360.1 |
| 1292 | HORVU3Hr1G021750 | Serine incorporator 1 | -5.11 | AT3G24460.2 |
| 1293 | HORVU3Hr1G077230 | Protein A | -5.12 | AT1G79190.2 |
| 1294 | HORVU7Hr1G092880 | undescribed protein | -5.14 | AT3G03750.2 |
| 1295 | HORVU7Hr1G105500 | Nuclear transcription factor Y subunit B-2 | -5.14 | AT4G14540.1 |

| | | | | |
|---|---|---|---|---|
| 1296 | HORVU2Hr1G038580 | F-box domain containing protein | -5.17 | AT2G13980.1 |
| 1297 | HORVU4Hr1G011160 | Endoglucanase 10 | -5.19 | AT5G49720.1 |
| 1298 | HORVU4Hr1G032020 | undescribed protein | -5.20 | AT5G62630.1 |
| 1299 | HORVU3Hr1G055260 | myb domain protein 66 | -5.20 | AT4G01060.1 |
| 1300 | HORVU5Hr1G071920 | Photosystem I reaction center subunit V | -5.26 | AT1G55670.1 |
| 1301 | HORVU7Hr1G098660 | Fe/S biogenesis protein NfuA | -5.27 | AT4G25910.1 |
| 1302 | HORVU3Hr1G009490 | Non-specific lipid-transfer protein 4.2 | -5.28 | AT5G59310.1 |
| 1303 | HORVU2Hr1G001320 | Protein NRT1/ PTR FAMILY 8.5 | -5.30 | AT1G62200.1 |
| 1304 | HORVU5Hr1G028180 | LIM domain-containing protein 2 | -5.31 | AT1G10200.1 |
| 1305 | HORVU1Hr1G043900 | Chloroplast inner envelope protein, putative, expressed | -5.32 | AT1G06950.1 |
| 1306 | HORVU5Hr1G074000 | NAD(P)-binding Rossmann-fold superfamily protein | -5.33 | AT3G18890.1 |
| 1307 | HORVU2Hr1G065420 | undescribed protein | -5.33 | AT4G22390.2 |
| 1308 | HORVU5Hr1G119230 | oligopeptide transporter 4 | -5.35 | AT5G64410.1 |
| 1309 | HORVU3Hr1G040880 | F-box family protein | -5.37 | AT2G32560.1 |
| 1310 | HORVU5Hr1G095480 | unknown function | -5.37 | AT4G16380.1 |
| 1311 | HORVU4Hr1G067020 | Caleosin-related family protein | -5.38 | AT2G33380.1 |
| 1312 | HORVU7Hr1G107840 | KY49.ctg7180000000017_quiver, whole genome shotgun sequence | -5.39 | AT1G14740.1 |
| 1313 | HORVU6Hr1G055350 | undescribed protein | -5.39 | AT1G60760.1 |
| 1314 | HORVU2Hr1G017480 | undescribed protein | -5.41 | AT4G32430.2 |
| 1315 | HORVU2Hr1G019820 | Photosystem I reaction center subunit N, chloroplastic | -5.41 | AT5G64040.1 |
| 1316 | HORVU5Hr1G124120 | gibberellin 20 oxidase 2 | -5.41 | AT4G25420.1 |
| 1317 | HORVU7Hr1G112290 | Chromosome 3B, genomic scaffold, cultivar Chinese Spring | -5.43 | AT2G24710.3 |
| 1318 | HORVU5Hr1G100870 | undescribed protein | -5.45 | AT5G64640.1 |
| 1319 | HORVU2Hr1G019270 | undescribed protein | -5.46 | AT2G06520.1 |
| 1320 | HORVU3Hr1G025820 | Mitochondrial substrate carrier family protein | -5.48 | AT5G19760.1 |
| 1321 | HORVU5Hr1G001750 | undescribed protein | -5.50 | AT5G17210.1 |
| 1322 | HORVU6Hr1G090250 | unknown function | -5.53 | AT5G37360.1 |
| 1323 | HORVU2Hr1G079920 | Chlorophyll a-b binding protein, chloroplastic | -5.56 | AT1G15820.1 |
| 1324 | HORVU3Hr1G080430 | Protein LURP-one-related 8 | -5.56 | AT2G38640.1 |
| 1325 | HORVU1Hr1G007630 | TRICHOME BIREFRINGENCE-LIKE 11 | -5.57 | AT5G19160.1 |
| 1326 | HORVU3Hr1G014790 | thylakoid lumenal 17.9 kDa protein, chloroplast | -5.59 | AT4G24930.1 |
| 1327 | HORVU5Hr1G016270 | Sulfurtransferase | -5.62 | AT1G27320.1 |
| 1328 | HORVU2Hr1G101730 | ascorbate peroxidase 4 | -5.63 | AT4G09010.1 |
| 1329 | HORVU3Hr1G109830 | undescribed protein | -5.65 | AT2G16440.1 |
| 1330 | HORVU5Hr1G049590 | undescribed protein | -5.69 | AT1G66210.1 |

| | | | | |
|---|---|---|---|---|
| 1331 | HORVU5Hr1G114230 | Bowman-Birk type trypsin inhibitor | -5.70 | AT4G21550.3 |
| 1332 | HORVU5Hr1G102530 | Hfr-2-like protein | -5.70 | AT5G44010.2 |
| 1333 | HORVU3Hr1G039670 | undescribed protein | -5.70 | #N/A |
| 1334 | HORVU1Hr1G061970 | 'putative endo-1,3;1,4-beta-D-glucanase' | -5.73 | AT3G23600.1 |
| 1335 | HORVU5Hr1G109260 | Chlorophyll a-b binding protein, chloroplastic | -5.75 | AT2G34420.1 |
| 1336 | HORVU2Hr1G090980 | Starch synthase 2, chloroplastic/amyloplastic | -5.79 | AT1G32900.1 |
| 1337 | HORVU0Hr1G028250 | Retrotransposon protein | -5.87 | AT2G07706.1 |
| 1338 | HORVU6Hr1G015770 | Photosystem II 5 kD protein | -5.88 | AT1G51400.1 |
| 1339 | HORVU2Hr1G113790 | undescribed protein | -5.89 | AT3G13100.1 |
| 1340 | HORVU2Hr1G079180 | unknown function | -5.89 | AT2G14095.1 |
| 1341 | HORVU3Hr1G052710 | myb-like transcription factor family protein | -5.90 | AT5G56840.1 |
| 1342 | HORVU3Hr1G011630 | undescribed protein | -5.90 | AT2G23380.2 |
| 1343 | HORVU5Hr1G064800 | Cell wall-associated hydrolase | -5.95 | AT4G22180.1 |
| 1344 | HORVU3Hr1G065090 | undescribed protein | -6.02 | AT1G15140.1 |
| 1345 | HORVU3Hr1G049730 | Chromosome 3B, genomic scaffold, cultivar Chinese Spring | -6.04 | AT4G14368.1 |
| 1346 | HORVU5Hr1G094110 | undescribed protein | -6.06 | AT5G62600.1 |
| 1347 | HORVU4Hr1G054240 | cAMP-regulated phosphoprotein 19-related protein | -6.11 | AT4G16146.1 |
| 1348 | HORVU6Hr1G048120 | undescribed protein | -6.12 | AT5G26110.3 |
| 1349 | HORVU3Hr1G087210 | respiratory burst oxidase protein F | -6.14 | AT5G60010.1 |
| 1350 | HORVU5Hr1G079600 | undescribed protein | -6.16 | AT3G57300.2 |
| 1351 | HORVU5Hr1G082420 | Chlorophyll a-b binding protein, chloroplastic | -6.18 | AT3G27690.2 |
| 1352 | HORVU3Hr1G014590 | serine/threonine protein phosphatase 2A | -6.19 | AT1G64040.1 |
| 1353 | HORVU1Hr1G056630 | UDP-glucose 4-epimerase | -6.20 | AT4G33030.1 |
| 1354 | HORVU4Hr1G079600 | undescribed protein | -6.20 | AT4G32700.2 |
| 1355 | HORVU6Hr1G016890 | Chlorophyll a-b binding protein, chloroplastic | -6.22 | AT2G34420.1 |
| 1356 | HORVU7Hr1G036780 | EXORDIUM like 2 | -6.22 | AT5G64260.1 |
| 1357 | HORVU3Hr1G026920 | metallothionein 3 | -6.26 | AT3G15353.1 |
| 1358 | HORVU4Hr1G090750 | Retrotransposon protein, putative, unclassified | -6.30 | AT1G55250.3 |
| 1359 | HORVU7Hr1G081510 | DnaJ/Hsp40 cysteine-rich domain superfamily protein | -6.38 | AT3G47650.1 |
| 1360 | HORVU1Hr1G078380 | Chlorophyll a-b binding protein, chloroplastic | -6.39 | AT2G34420.1 |
| 1361 | HORVU5Hr1G110610 | undescribed protein | -6.45 | AT4G39590.1 |

| | | | | |
|---|---|---|---|---|
| 1362 | HORVU2Hr1G070140 | Heavy metal transport/detoxification superfamily protein | -6.48 | AT5G50740.5 |
| 1363 | HORVU7Hr1G056700 | BAG family molecular chaperone regulator 3 | -6.48 | AT5G07220.1 |
| 1364 | HORVU6Hr1G028970 | undescribed protein | -6.49 | AT1G65870.1 |
| 1365 | HORVU3Hr1G069280 | Chromosome 3B, genomic scaffold, cultivar Chinese Spring | -6.53 | AT2G42760.1 |
| 1366 | HORVU5Hr1G102620 | Amaranthin-like lectin | -6.54 | AT3G23800.2 |
| 1367 | HORVU4Hr1G087580 | Chromosome 3B, genomic scaffold, cultivar Chinese Spring | -6.56 | AT1G10586.1 |
| 1368 | HORVU4Hr1G005320 | Protein of unknown function, DUF538 | -6.57 | AT1G55265.1 |
| 1369 | HORVU5Hr1G053480 | Protein FLUORESCENT IN BLUE LIGHT, chloroplastic | -6.58 | AT3G14110.2 |
| 1370 | HORVU6Hr1G035460 | undescribed protein | -6.60 | AT1G65640.2 |
| 1371 | HORVU2Hr1G010630 | ribulose bisphosphate carboxylase small chain 1A | -6.62 | AT1G67090.1 |
| 1372 | HORVU5Hr1G113910 | undescribed protein | -6.64 | AT1G26090.1 |
| 1373 | HORVU0Hr1G034230 | Retrotransposon protein | -6.75 | AT2G07706.1 |
| 1374 | HORVU2Hr1G121770 | Chromosome 3B, genomic scaffold, cultivar Chinese Spring | -6.77 | AT1G71140.1 |
| 1375 | HORVU7Hr1G119060 | GDSL esterase/lipase | -6.80 | AT5G33370.1 |
| 1376 | HORVU6Hr1G073100 | Protein CURVATURE THYLAKOID 1A, chloroplastic | -6.83 | AT4G01150.1 |
| 1377 | HORVU0Hr1G013070 | cinnamoyl coa reductase | -6.86 | AT1G15950.7 |
| 1378 | HORVU3Hr1G068200 | Heavy metal transport/detoxification superfamily protein | -6.92 | AT3G24450.1 |
| 1379 | HORVU2Hr1G089340 | phosphoenolpyruvate carboxylase kinase 1 | -6.98 | AT1G08650.1 |
| 1380 | HORVU3Hr1G027220 | Polyketide cyclase/dehydrase and lipid transport superfamily protein | -7.04 | AT2G25770.2 |
| 1381 | HORVU3Hr1G110660 | NADH dehydrogenase-like complex O | -7.06 | AT1G74880.1 |
| 1382 | HORVU1Hr1G007810 | UDP-Glycosyltransferase superfamily protein | -7.06 | AT1G22360.1 |
| 1383 | HORVU5Hr1G064230 | Cellulose synthase family protein | -7.07 | AT5G17420.1 |
| 1384 | HORVU3Hr1G024920 | undescribed protein | -7.08 | AT5G17650.1 |
| 1385 | HORVU3Hr1G110330 | Phospholipase A1-II 6 | -7.17 | AT1G06800.1 |
| 1386 | HORVU6Hr1G093940 | Peroxidase superfamily protein | -7.17 | AT1G77490.1 |
| 1387 | HORVU2Hr1G010690 | ribulose bisphosphate carboxylase small chain 1A | -7.20 | AT1G67090.1 |
| 1388 | HORVU5Hr1G113060 | actin depolymerizing factor 4 | -7.23 | AT4G00680.1 |
| 1389 | HORVU3Hr1G085520 | laccase 7 | -7.25 | AT3G09220.1 |
| 1390 | HORVU5Hr1G109610 | unknown function | -7.28 | AT5G43150.1 |
| 1391 | HORVU5Hr1G010960 | Glutaredoxin family protein | -7.30 | AT2G47870.1 |
| 1392 | HORVU2Hr1G074730 | unknown function | -7.35 | AT3G55480.1 |

| | | | | |
|---|---|---|---|---|
| 1393 | HORVU6Hr1G016940 | Chlorophyll a-b binding protein, chloroplastic | -7.35 | AT2G34420.1 |
| 1394 | HORVU5Hr1G022750 | PsbP domain-containing protein 1, chloroplastic | -7.38 | AT4G15510.5 |
| 1395 | HORVU3Hr1G071770 | cellulose synthase family protein | -7.51 | AT4G18780.1 |
| 1396 | HORVU3Hr1G004080 | undescribed protein | -7.58 | AT2G35110.1 |
| 1397 | HORVU6Hr1G049260 | Ribulose bisphosphate carboxylase large chain | -7.58 | ATCG00490.1 |
| 1398 | HORVU2Hr1G077630 | Histone H2B.1 | -7.58 | AT3G45980.1 |
| 1399 | HORVU5Hr1G097870 | Cytochrome b6-f complex subunit 7 | -7.58 | AT2G26500.2 |
| 1400 | HORVU0Hr1G020630 | Benzoate O-methyltransferase | -7.58 | AT5G04370.3 |
| 1401 | HORVU4Hr1G069690 | undescribed protein | -7.60 | AT1G24430.1 |
| 1402 | HORVU2Hr1G045970 | unknown function | -7.61 | AT3G21710.2 |
| 1403 | HORVU2Hr1G000420 | HXXXD-type acyl-transferase family protein | -7.62 | AT3G62160.1 |
| 1404 | HORVU1Hr1G039250 | cellulose synthase A4 | -7.64 | AT5G44030.3 |
| 1405 | HORVU1Hr1G007690 | General transcription factor 2-related zinc finger protein | -7.69 | AT2G06541.2 |
| 1406 | HORVU4Hr1G009380 | Lariat debranching enzyme | -7.72 | AT3G22436.1 |
| 1407 | HORVU0Hr1G034200 | undescribed protein | -7.74 | AT5G56400.1 |
| 1408 | HORVU6Hr1G091650 | Chlorophyll a-b binding protein, chloroplastic | -7.84 | AT2G34420.1 |
| 1409 | HORVU1Hr1G088920 | Chlorophyll a-b binding protein, chloroplastic | -7.91 | AT2G34420.1 |
| 1410 | HORVU2Hr1G041270 | Chlorophyll a-b binding protein 6, chloroplastic | -7.94 | AT5G01530.1 |
| 1411 | HORVU5Hr1G109250 | Chlorophyll a-b binding protein, chloroplastic | -8.06 | AT1G29930.1 |
| 1412 | HORVU5Hr1G086710 | Flavin-containing monooxygenase family protein | -8.07 | AT1G19250.1 |
| 1413 | HORVU1Hr1G089240 | Cytochrome c oxidase subunit 1 | -8.09 | ATMG01360.1 |
| 1414 | HORVU6Hr1G051650 | Oxygen-evolving enhancer protein 3-1, chloroplastic | -8.10 | AT1G14150.1 |
| 1415 | HORVU7Hr1G065970 | undescribed protein | -8.13 | AT3G48430.1 |
| 1416 | HORVU3Hr1G039310 | undescribed protein | -8.13 | AT1G29720.1 |
| 1417 | HORVU4Hr1G086020 | 4-hydroxy-tetrahydrodipicolinate reductase | -8.18 | AT5G52100.1 |
| 1418 | HORVU1Hr1G024890 | NAD(P)H dehydrogenase 18 | -8.30 | AT5G43750.1 |
| 1419 | HORVU6Hr1G004000 | Acid beta-fructofuranosidase | -8.41 | AT1G62660.2 |
| 1420 | HORVU0Hr1G038990 | Protein of unknown function (DUF3511) | -8.56 | AT4G09890.1 |
| 1421 | HORVU3Hr1G096910 | Beta-glucosidase 3 | -8.62 | AT3G60140.2 |
| 1422 | HORVU5Hr1G001180 | lipoxygenase 2 | -8.66 | AT3G45140.1 |
| 1423 | HORVU5Hr1G018970 | 0 | -8.69 | AT1G22540.1 |

| | | | | |
|---|---|---|---|---|
| 1424 | HORVU5Hr1G019480 | Disease resistance-responsive (dirigent-like protein) family protein | -8.71 | AT3G58090.1 |
| 1425 | HORVU1Hr1G075160 | Isoflavone reductase-like protein | -8.76 | AT1G75290.1 |
| 1426 | HORVU5Hr1G067530 | 1-aminocyclopropane-1-carboxylate oxidase 1 | -8.79 | AT1G05010.1 |
| 1427 | HORVU1Hr1G089180 | Chlorophyll a-b binding protein, chloroplastic | -8.80 | AT2G34420.1 |
| 1428 | HORVU6Hr1G016860 | undescribed protein | -8.93 | AT5G18890.1 |
| 1429 | HORVU4Hr1G088080 | expansin B2 | -9.20 | AT1G65680.1 |
| 1430 | HORVU7Hr1G047690 | Chromosome 3B, genomic scaffold, cultivar Chinese Spring | -9.32 | AT4G20970.1 |
| 1431 | HORVU3Hr1G089910 | Trigger factor | -9.38 | AT2G30695.3 |
| 1432 | HORVU0Hr1G012330 | undescribed protein | -9.44 | AT5G42390.1 |
| 1433 | HORVU2Hr1G102110 | Bifunctional inhibitor/lipid-transfer protein/seed storage 2S albumin superfamily protein | -9.45 | AT5G48485.1 |
| 1434 | HORVU0Hr1G012340 | undescribed protein | -9.47 | AT2G46130.1 |
| 1435 | HORVU2Hr1G075170 | undescribed protein | -9.51 | AT1G67910.2 |
| 1436 | HORVU2Hr1G102100 | Bifunctional inhibitor/lipid-transfer protein/seed storage 2S albumin superfamily protein | -9.56 | AT5G48485.1 |
| 1437 | HORVU7Hr1G086120 | Leucine-rich repeat receptor-like protein kinase family protein | -9.60 | AT3G47570.1 |
| 1438 | HORVU7Hr1G120660 | PHYTOENE SYNTHASE | -9.63 | AT5G17230.4 |
| 1439 | HORVU7Hr1G040380 | Chlorophyll a-b binding protein, chloroplastic | -9.69 | AT2G34420.1 |
| 1440 | HORVU2Hr1G073030 | Carbonic anhydrase | -9.72 | AT3G52720.4 |
| 1441 | HORVU7Hr1G112130 | undescribed protein | -9.74 | AT5G28080.2 |
| 1442 | HORVU6Hr1G016880 | Chlorophyll a-b binding protein, chloroplastic | -9.78 | AT2G34420.1 |
| 1443 | HORVU7Hr1G010690 | Acid phosphatase 1 | -9.78 | AT4G29260.1 |
| 1444 | HORVU6Hr1G050760 | undescribed protein | -9.80 | AT5G02390.3 |
| 1445 | HORVU1Hr1G071910 | Plant protein 1589 of unknown function | -9.83 | AT5G02580.1 |
| 1446 | HORVU2Hr1G010670 | ribulose bisphosphate carboxylase small chain 1A | -9.91 | AT1G67090.1 |
| 1447 | HORVU5Hr1G023160 | Transmembrane amino acid transporter family protein | -9.94 | AT1G47670.1 |
| 1448 | HORVU5Hr1G093640 | Auxin-responsive protein IAA12 | -9.99 | AT1G04240.1 |
| 1449 | HORVU4Hr1G018150 | sterol C4-methyl oxidase 1-2 | -10.06 | AT4G22756.1 |
| 1450 | HORVU7Hr1G058120 | Chlorophyll a-b binding protein 1B, chloroplastic | -10.09 | AT2G34430.1 |
| 1451 | HORVU4Hr1G002900 | Protein of unknown function (DUF1677) | -10.22 | AT3G22540.1 |
| 1452 | HORVU4Hr1G076050 | O-methyltransferase family protein | -10.40 | AT4G35160.1 |
| 1453 | HORVU2Hr1G065450 | flowering promoting factor 1 | -10.58 | AT5G24860.1 |

| | | | | |
|---|---|---|---|---|
| 1454 | HORVU7Hr1G098370 | Xyloglucan endotransglucosylase/hydrolase family protein | -10.61 | AT5G57560.1 |
| 1455 | HORVU5Hr1G004700 | terpene synthase 21 | -10.83 | AT5G23960.1 |
| 1456 | HORVU6Hr1G012270 | terpene synthase 14 | -10.86 | AT1G61680.1 |
| 1457 | HORVU4Hr1G088160 | Eukaryotic aspartyl protease family protein | -10.87 | AT2G03200.1 |
| 1458 | HORVU2Hr1G094280 | Protein FATTY ACID EXPORT 4, chloroplastic | -10.96 | AT1G33265.1 |
| 1459 | HORVU4Hr1G018180 | sterol C4-methyl oxidase 1-2 | -11.03 | AT4G22756.1 |
| 1460 | HORVU0Hr1G022410 | undescribed protein | -11.15 | AT1G05860.3 |
| 1461 | HORVU5Hr1G067490 | 1-aminocyclopropane-1-carboxylate oxidase 1 | -11.20 | AT1G05010.1 |
| 1462 | HORVU2Hr1G038040 | unknown function | -11.38 | AT5G43150.1 |
| 1463 | HORVU1Hr1G093660 | ribonuclease 3 | -11.39 | AT1G26820.1 |
| 1464 | HORVU1Hr1G073630 | undescribed protein | -11.40 | AT2G06541.2 |
| 1465 | HORVU2Hr1G064730 | Protein of unknown function, DUF584 | -11.52 | AT1G11700.1 |
| 1466 | HORVU2Hr1G094390 | histone H1-3 | -11.52 | AT2G18050.1 |
| 1467 | HORVU2Hr1G051050 | undescribed protein | -11.54 | AT4G24560.1 |
| 1468 | HORVU2Hr1G063190 | undescribed protein | -11.54 | AT1G24485.3 |
| 1469 | HORVU5Hr1G075660 | UDP-glucose 4-epimerase | -11.79 | AT3G23820.1 |
| 1470 | HORVU7Hr1G122470 | sulfotransferase 4A | -11.93 | AT3G45070.3 |
| 1471 | HORVU7Hr1G040370 | Chlorophyll a-b binding protein, chloroplastic | -11.93 | AT2G34420.1 |
| 1472 | HORVU6Hr1G061270 | Aldehyde dehydrogenase family 3 member F1 | -12.04 | AT4G36250.1 |
| 1473 | HORVU2Hr1G074770 | Abscisic stress-ripening protein 3 | -12.80 | AT2G03140.9 |
| 1474 | HORVU3Hr1G108550 | undescribed protein | -12.83 | AT3G43920.5 |
| 1475 | HORVU4Hr1G088110 | Eukaryotic aspartyl protease family protein | -12.93 | AT2G03200.1 |
| 1476 | HORVU2Hr1G051040 | undescribed protein | -13.01 | AT3G11670.1 |
| 1477 | HORVU0Hr1G005360 | serine carboxypeptidase-like 19 | -13.23 | AT1G33540.1 |
| 1478 | HORVU7Hr1G002620 | undescribed protein | -13.47 | AT1G72540.1 |
| 1479 | HORVU7Hr1G090410 | polyamine oxidase 1 | -13.54 | AT5G13700.1 |
| 1480 | HORVU4Hr1G090860 | metacaspase 1 | -13.76 | AT1G02170.1 |
| 1481 | HORVU3Hr1G001680 | TPX2 (targeting protein for Xklp2) protein family | -13.77 | AT3G23090.5 |
| 1482 | HORVU2Hr1G009970 | Eukaryotic aspartyl protease family protein | -14.13 | AT2G03200.1 |
| 1483 | HORVU1Hr1G069830 | expansin A1 | -14.25 | AT1G69530.2 |
| 1484 | HORVU2Hr1G002720 | PLANT CADMIUM RESISTANCE 1 | -14.43 | AT1G14880.1 |
| 1485 | HORVU5Hr1G063630 | Early light-induced protein 1, chloroplastic | -14.72 | AT3G22840.1 |
| 1486 | HORVU1Hr1G057860 | cytokinin oxidase/dehydrogenase 1 | -14.86 | AT2G41510.2 |

| | | | | |
|---|---|---|---|---|
| 1487 | HORVU2Hr1G032710 | unknown function | -15.36 | AT1G74160.1 |
| 1488 | HORVU6Hr1G016850 | Chlorophyll a-b binding protein, chloroplastic | -15.74 | AT2G34420.1 |
| 1489 | HORVU3Hr1G025300 | undescribed protein | -16.16 | AT2G42320.4 |
| 1490 | HORVU1Hr1G083580 | undescribed protein | -16.24 | AT5G25240.1 |
| 1491 | HORVU4Hr1G081040 | ABC transporter G family member 45 | -16.38 | AT2G36380.1 |
| 1492 | HORVU5Hr1G067200 | RING/U-box superfamily protein | -16.60 | AT3G61460.1 |
| 1493 | HORVU6Hr1G025050 | undescribed protein | -16.79 | AT2G20930.1 |
| 1494 | HORVU3Hr1G094580 | undescribed protein | -17.56 | AT3G24050.1 |
| 1495 | HORVU0Hr1G022690 | Senescence-associated protein DIN1 | -18.04 | AT4G35770.1 |
| 1496 | HORVU4Hr1G080820 | Group 3 late embryogenesis abundant protein | -18.32 | AT1G52690.2 |
| 1497 | HORVU6Hr1G012110 | terpene synthase 14 | -18.35 | AT1G61680.1 |
| 1498 | HORVU1Hr1G088900 | Chlorophyll a-b binding protein, chloroplastic | -18.39 | AT1G29930.1 |
| 1499 | HORVU4Hr1G058590 | unknown function | -18.45 | AT1G80830.2 |
| 1500 | HORVU3Hr1G073860 | chlororespiratory reduction 7 | -19.09 | AT5G39210.1 |
| 1501 | HORVU6Hr1G079190 | Peroxidase superfamily protein | -19.25 | AT5G15180.1 |
| 1502 | HORVU3Hr1G069650 | BURP domain-containing protein | -19.36 | AT5G25610.1 |
| 1503 | HORVU4Hr1G066230 | allene oxide synthase | -19.68 | AT5G42650.1 |
| 1504 | HORVU6Hr1G009360 | Peroxidase superfamily protein | -19.70 | AT5G05340.1 |
| 1505 | HORVU4Hr1G028720 | xyloglucan endotransglucosylase/hydrolase 5 | -20.53 | AT5G13870.3 |
| 1506 | HORVU2Hr1G090670 | sulfotransferase 4B | -20.80 | AT1G13420.1 |
| 1507 | HORVU5Hr1G082160 | O-acyltransferase (WSD1-like) family protein | -23.08 | AT5G53390.1 |
| 1508 | HORVU4Hr1G057000 | Low temperature and salt responsive protein family | -23.17 | AT2G38905.1 |
| 1509 | HORVU4Hr1G040110 | undescribed protein | -23.53 | AT3G52330.1 |
| 1510 | HORVU7Hr1G036720 | germin-like protein 1 | -23.80 | AT1G72610.1 |
| 1511 | HORVU5Hr1G077390 | Transcription factor bHLH137 | -24.17 | AT5G50915.4 |
| 1512 | HORVU1Hr1G089380 | Subtilisin-like protease | -24.59 | AT4G10550.1 |
| 1513 | HORVU4Hr1G076000 | O-methyltransferase family protein | -25.54 | AT4G35160.1 |
| 1514 | HORVU5Hr1G010950 | Glutaredoxin family protein | -25.92 | AT5G18600.1 |
| 1515 | HORVU1Hr1G093570 | Ribonuclease T2 family protein | -27.75 | AT1G14220.1 |
| 1516 | HORVU7Hr1G007800 | undescribed protein | -30.08 | AT1G32560.1 |
| 1517 | HORVU2Hr1G003470 | Cytochrome P450 superfamily protein | -30.14 | AT2G22330.2 |
| 1518 | HORVU3Hr1G027970 | TRICHOME BIREFRINGENCE-LIKE 38 | -32.28 | AT1G29050.1 |
| 1519 | HORVU6Hr1G005050 | flowering promoting factor 1 | -33.02 | AT5G24860.1 |
| 1520 | HORVU5Hr1G084230 | Aquaporin-like superfamily protein | -37.05 | AT2G37170.2 |
| 1521 | HORVU5Hr1G101880 | Protein phosphatase 2C 35 | -40.08 | AT1G60650.2 |
| 1522 | HORVU1Hr1G088840 | Phospholipase A1-II 7 | -41.32 | AT4G18550.1 |
| 1523 | HORVU2Hr1G103160 | undescribed protein | -42.01 | AT5G39690.1 |

| | | | | |
|---|---|---|---|---|
| 1524 | HORVU1Hr1G016410 | undescribed protein | -43.17 | AT1G16460.4 |
| 1525 | HORVU7Hr1G120060 | Cathepsin B-like cysteine proteinase 1 | -43.50 | AT2G34080.1 |
| 1526 | HORVU7Hr1G119930 | Cathepsin B-like cysteine proteinase 1 | -43.76 | AT2G34080.1 |
| 1527 | HORVU5Hr1G089230 | Fatty acyl-CoA reductase 1 | -61.94 | AT5G22500.1 |
| 1528 | HORVU7Hr1G098320 | xyloglucan endotransglucosylase/hydrolase 25 | -71.19 | AT5G57550.1 |
| 1529 | HORVU7Hr1G098330 | xyloglucan endotransglucosylase/hydrolase 25 | -97.06 | AT5G57550.1 |
| 1530 | HORVU7Hr1G111010 | unknown function | -108.99 | AT3G06130.1 |
| 1531 | HORVU7Hr1G098280 | xyloglucan endotransglucosylase/hydrolase 25 | -133.06 | AT4G25810.1 |
| 1532 | HORVU2Hr1G043930 | undescribed protein | -208.98 | AT4G12650.1 |

SUPPLEMENTARY TABLE 4

| S No | 845 exclusively in "Harrington_Up": | |
|---|---|---|
| 1 | HORVU7Hr1G122120 | AT4G11650.1 |
| 2 | HORVU3Hr1G003980 | ATCG00180.1 |
| 3 | HORVU5Hr1G067760 | #N/A |
| 4 | HORVU2Hr1G118020 | AT1G09580.1 |
| 5 | HORVU2Hr1G118010 | AT3G47570.1 |
| 6 | HORVU5Hr1G122520 | AT3G25727.1 |
| 7 | HORVU7Hr1G085450 | AT5G54260.1 |
| 8 | HORVU6Hr1G066380 | AT1G72610.1 |
| 9 | HORVU3Hr1G105600 | AT4G16260.1 |
| 10 | HORVU3Hr1G105480 | AT4G16260.2 |
| 11 | HORVU5Hr1G023660 | AT5G51770.1 |
| 12 | HORVU5Hr1G009940 | AT2G33150.1 |
| 13 | HORVU3Hr1G105420 | AT3G57240.1 |
| 14 | HORVU3Hr1G091780 | AT2G18140.1 |
| 15 | HORVU3Hr1G104940 | AT1G71830.1 |
| 16 | HORVU5Hr1G118800 | AT3G62760.1 |
| 17 | HORVU3Hr1G091740 | AT5G06730.1 |
| 18 | HORVU5Hr1G106020 | AT3G19690.1 |
| 19 | HORVU1Hr1G043910 | AT2G15130.1 |
| 20 | HORVU3Hr1G091750 | AT5G06730.1 |
| 21 | HORVU7Hr1G085420 | AT3G29830.2 |
| 22 | HORVU1Hr1G009800 | AT3G22600.1 |
| 23 | HORVU5Hr1G119640 | AT1G33600.2 |
| 24 | HORVU3Hr1G077880 | AT3G20340.1 |
| 25 | HORVU5Hr1G081620 | AT2G46790.3 |

| | | |
|---|---|---|
| 26 | HORVU4Hr1G077300 | AT1G18197.1 |
| 27 | HORVU7Hr1G010840 | AT4G28680.4 |
| 28 | HORVU4Hr1G071300 | AT4G33680.1 |
| 29 | HORVU0Hr1G008950 | AT3G49680.1 |
| 30 | HORVU3Hr1G084210 | AT2G27510.1 |
| 31 | HORVU2Hr1G122630 | AT5G26330.1 |
| 32 | HORVU7Hr1G017810 | AT5G60900.1 |
| 33 | HORVU2Hr1G039670 | AT1G52900.1 |
| 34 | HORVU7Hr1G080190 | AT3G62560.1 |
| 35 | HORVU7Hr1G007480 | AT3G47570.1 |
| 36 | HORVU1Hr1G091960 | AT1G06475.1 |
| 37 | HORVU2Hr1G124960 | AT1G71695.1 |
| 38 | HORVU7Hr1G009310 | AT5G20400.1 |
| 39 | HORVU1Hr1G089530 | AT1G19450.1 |
| 40 | HORVU7Hr1G043540 | AT3G48280.1 |
| 41 | HORVU6Hr1G089460 | AT1G15950.3 |
| 42 | HORVU6Hr1G013710 | AT2G15220.1 |
| 43 | HORVU7Hr1G108490 | AT3G47570.1 |
| 44 | HORVU6Hr1G033060 | AT2G36290.1 |
| 45 | HORVU2Hr1G118030 | AT1G48980.3 |
| 46 | HORVU7Hr1G002820 | AT3G01420.1 |
| 47 | HORVU1Hr1G044000 | AT2G15220.1 |
| 48 | HORVU1Hr1G049230 | AT1G10360.1 |
| 49 | HORVU3Hr1G091730 | AT5G06730.1 |
| 50 | HORVU2Hr1G002750 | AT5G10530.1 |
| 51 | HORVU4Hr1G018100 | AT1G77400.1 |
| 52 | HORVU1Hr1G049290 | AT5G63420.1 |
| 53 | HORVU6Hr1G001520 | AT4G29990.1 |
| 54 | HORVU2Hr1G119250 | AT2G22170.1 |
| 55 | HORVU7Hr1G122110 | AT5G02140.2 |
| 56 | HORVU6Hr1G002530 | AT1G21230.1 |
| 57 | HORVU3Hr1G111640 | AT1G10360.1 |
| 58 | HORVU7Hr1G040780 | AT4G06634.2 |
| 59 | HORVU2Hr1G124320 | AT1G78380.1 |
| 60 | HORVU3Hr1G098610 | AT5G06860.1 |
| 61 | HORVU6Hr1G066420 | AT5G48930.1 |
| 62 | HORVU7Hr1G034270 | AT2G40330.1 |
| 63 | HORVU4Hr1G007190 | AT2G02850.1 |
| 64 | HORVU2Hr1G112040 | #N/A |
| 65 | HORVU5Hr1G079680 | AT2G43840.1 |
| 66 | HORVU7Hr1G011860 | AT3G04960.2 |

| | | |
|---|---|---|
| 67 | HORVU4Hr1G065000 | AT5G06730.1 |
| 68 | HORVU7Hr1G024360 | AT5G59310.1 |
| 69 | HORVU3Hr1G006620 | AT4G23160.1 |
| 70 | HORVU6Hr1G077180 | AT3G17100.2 |
| 71 | HORVU5Hr1G010880 | AT5G39100.1 |
| 72 | HORVU4Hr1G014670 | AT1G18480.1 |
| 73 | HORVU3Hr1G026270 | AT4G10520.1 |
| 74 | HORVU6Hr1G055000 | AT5G13930.1 |
| 75 | HORVU3Hr1G004050 | AT1G68320.1 |
| 76 | HORVU6Hr1G094480 | AT1G12160.1 |
| 77 | HORVU2Hr1G017440 | AT2G39050.1 |
| 78 | HORVU4Hr1G054870 | AT2G26040.1 |
| 79 | HORVU6Hr1G000130 | AT5G66170.3 |
| 80 | HORVU4Hr1G066240 | AT2G16060.1 |
| 81 | HORVU7Hr1G091880 | AT4G37290.1 |
| 82 | HORVU3Hr1G002830 | AT1G10360.1 |
| 83 | HORVU7Hr1G025930 | #N/A |
| 84 | HORVU3Hr1G112040 | AT1G71695.1 |
| 85 | HORVU6Hr1G001530 | AT1G49100.1 |
| 86 | HORVU5Hr1G123540 | AT4G23180.1 |
| 87 | HORVU4Hr1G005540 | AT1G18970.1 |
| 88 | HORVU7Hr1G085480 | #N/A |
| 89 | HORVU5Hr1G014170 | AT5G28770.2 |
| 90 | HORVU7Hr1G031650 | AT4G36670.1 |
| 91 | HORVU7Hr1G091800 | AT5G39150.1 |
| 92 | HORVU5Hr1G111240 | AT4G21410.3 |
| 93 | HORVU3Hr1G074230 | AT4G37390.1 |
| 94 | HORVU5Hr1G115900 | AT1G14870.1 |
| 95 | HORVU7Hr1G035980 | AT5G22870.1 |
| 96 | HORVU4Hr1G012750 | AT5G50300.2 |
| 97 | HORVU7Hr1G049580 | AT1G21230.1 |
| 98 | HORVU2Hr1G076490 | AT3G19380.1 |
| 99 | HORVU1Hr1G003210 | AT5G20710.1 |
| 100 | HORVU2Hr1G121170 | AT3G61040.2 |
| 101 | HORVU4Hr1G057550 | AT5G02810.1 |
| 102 | HORVU3Hr1G086680 | AT2G28710.1 |
| 103 | HORVU3Hr1G021140 | AT1G22770.1 |
| 104 | HORVU6Hr1G073160 | AT2G18690.1 |
| 105 | HORVU6Hr1G004300 | AT1G21240.2 |
| 106 | HORVU1Hr1G010210 | AT3G48740.1 |
| 107 | HORVU5Hr1G074030 | AT5G66580.1 |

| | | |
|---|---|---|
| 108 | HORVU2Hr1G113080 | AT1G16260.2 |
| 109 | HORVU3Hr1G025810 | AT5G45810.1 |
| 110 | HORVU3Hr1G006730 | AT4G23180.1 |
| 111 | HORVU3Hr1G035680 | AT5G24860.1 |
| 112 | HORVU5Hr1G122510 | AT3G47570.1 |
| 113 | HORVU6Hr1G088420 | AT1G49390.1 |
| 114 | HORVU3Hr1G001990 | AT3G05950.1 |
| 115 | HORVU7Hr1G100810 | AT1G06620.1 |
| 116 | HORVU7Hr1G006940 | AT1G56130.2 |
| 117 | HORVU1Hr1G090820 | AT2G25625.2 |
| 118 | HORVU2Hr1G083020 | AT3G48270.1 |
| 119 | HORVU2Hr1G030660 | AT5G42900.2 |
| 120 | HORVU2Hr1G004500 | AT2G30770.1 |
| 121 | HORVU6Hr1G088960 | AT3G14690.1 |
| 122 | HORVU3Hr1G006800 | AT4G23280.1 |
| 123 | HORVU6Hr1G001420 | AT1G21230.1 |
| 124 | HORVU1Hr1G025530 | AT1G15380.2 |
| 125 | HORVU5Hr1G093090 | AT4G21120.1 |
| 126 | HORVU4Hr1G043960 | AT2G03200.1 |
| 127 | HORVU5Hr1G066720 | AT1G55230.1 |
| 128 | HORVU7Hr1G035190 | AT4G18930.1 |
| 129 | HORVU7Hr1G040170 | AT3G21950.1 |
| 130 | HORVU6Hr1G074060 | AT5G13930.1 |
| 131 | HORVU4Hr1G063590 | AT2G17880.1 |
| 132 | HORVU5Hr1G047730 | AT2G40300.1 |
| 133 | HORVU5Hr1G106000 | AT3G05200.1 |
| 134 | HORVU4Hr1G064480 | AT5G19790.1 |
| 135 | HORVU3Hr1G021150 | AT1G22770.1 |
| 136 | HORVU3Hr1G056110 | AT3G14690.1 |
| 137 | HORVU3Hr1G087570 | AT5G05600.2 |
| 138 | HORVU3Hr1G026050 | AT4G14440.1 |
| 139 | HORVU3Hr1G011850 | AT3G48270.1 |
| 140 | HORVU2Hr1G045350 | AT3G03800.2 |
| 141 | HORVU7Hr1G101300 | AT5G04220.2 |
| 142 | HORVU1Hr1G016870 | AT1G05260.1 |
| 143 | HORVU3Hr1G109510 | AT4G20020.2 |
| 144 | HORVU7Hr1G113290 | AT2G38465.1 |
| 145 | HORVU1Hr1G092680 | AT5G44390.1 |
| 146 | HORVU1Hr1G090730 | AT2G26640.1 |
| 147 | HORVU2Hr1G018260 | AT5G58380.1 |
| 148 | HORVU7Hr1G024940 | AT4G10520.1 |

| | | |
|---|---|---|
| 149 | HORVU5Hr1G086670 | AT1G19250.1 |
| 150 | HORVU3Hr1G064680 | AT2G34250.3 |
| 151 | HORVU3Hr1G108100 | AT3G63520.1 |
| 152 | HORVU2Hr1G044890 | AT4G05200.1 |
| 153 | HORVU4Hr1G056240 | AT5G65010.1 |
| 154 | HORVU2Hr1G084130 | AT5G03860.2 |
| 155 | HORVU6Hr1G079950 | #N/A |
| 156 | HORVU5Hr1G122060 | AT2G02850.1 |
| 157 | HORVU5Hr1G014130 | AT1G47480.1 |
| 158 | HORVU4Hr1G066200 | AT2G16060.1 |
| 159 | HORVU7Hr1G098810 | AT5G42020.1 |
| 160 | HORVU7Hr1G106160 | AT2G40460.1 |
| 161 | HORVU3Hr1G007590 | AT1G18390.2 |
| 162 | HORVU7Hr1G114790 | AT3G09510.1 |
| 163 | HORVU0Hr1G023570 | AT5G48930.1 |
| 164 | HORVU2Hr1G042550 | AT4G23180.2 |
| 165 | HORVU1Hr1G065630 | AT5G50930.1 |
| 166 | HORVU4Hr1G022270 | AT5G05340.1 |
| 167 | HORVU2Hr1G104090 | AT1G16630.1 |
| 168 | HORVU1Hr1G013480 | AT1G34580.1 |
| 169 | HORVU3Hr1G109590 | AT2G41640.1 |
| 170 | HORVU2Hr1G126120 | AT1G22570.2 |
| 171 | HORVU2Hr1G019790 | AT2G47780.1 |
| 172 | HORVU5Hr1G115310 | AT5G66460.1 |
| 173 | HORVU4Hr1G089030 | AT3G15040.1 |
| 174 | HORVU2Hr1G093660 | AT3G23240.1 |
| 175 | HORVU5Hr1G061810 | AT2G05940.1 |
| 176 | HORVU1Hr1G091030 | AT1G70940.1 |
| 177 | HORVU6Hr1G008730 | AT1G20620.4 |
| 178 | HORVU2Hr1G083010 | AT4G12120.1 |
| 179 | HORVU6Hr1G089560 | AT2G35930.1 |
| 180 | HORVU4Hr1G088930 | AT2G46620.1 |
| 181 | HORVU6Hr1G092020 | AT5G20480.2 |
| 182 | HORVU7Hr1G001520 | AT4G00300.1 |
| 183 | HORVU1Hr1G071390 | AT1G74540.1 |
| 184 | HORVU6Hr1G067910 | AT2G02040.1 |
| 185 | HORVU4Hr1G003450 | AT1G22680.1 |
| 186 | HORVU2Hr1G042540 | AT1G32660.1 |
| 187 | HORVU7Hr1G117850 | AT4G35160.1 |
| 188 | HORVU3Hr1G105920 | AT2G01170.1 |
| 189 | HORVU3Hr1G057180 | AT5G62770.1 |

| | | |
|---|---|---|
| 190 | HORVU2Hr1G046370 | AT4G21870.1 |
| 191 | HORVU6Hr1G001370 | #N/A |
| 192 | HORVU6Hr1G004310 | AT1G12120.1 |
| 193 | HORVU7Hr1G026640 | AT4G38180.1 |
| 194 | HORVU5Hr1G070400 | AT1G17840.1 |
| 195 | HORVU2Hr1G061670 | AT2G26170.1 |
| 196 | HORVU7Hr1G107150 | AT2G29690.2 |
| 197 | HORVU3Hr1G091120 | AT5G09220.1 |
| 198 | HORVU1Hr1G077420 | AT5G16760.1 |
| 199 | HORVU6Hr1G057310 | AT1G24062.1 |
| 200 | HORVU5Hr1G061820 | AT2G05940.1 |
| 201 | HORVU3Hr1G074940 | AT4G11290.1 |
| 202 | HORVU7Hr1G040850 | AT5G18520.1 |
| 203 | HORVU5Hr1G080300 | AT4G25470.1 |
| 204 | HORVU1Hr1G090870 | AT1G52820.1 |
| 205 | HORVU5Hr1G121520 | AT5G18980.1 |
| 206 | HORVU1Hr1G082670 | AT1G68840.2 |
| 207 | HORVU6Hr1G063480 | AT2G43750.2 |
| 208 | HORVU6Hr1G022590 | AT3G05200.1 |
| 209 | HORVU3Hr1G064960 | AT5G60900.1 |
| 210 | HORVU0Hr1G000780 | AT5G24275.4 |
| 211 | HORVU7Hr1G049610 | AT1G80520.1 |
| 212 | HORVU5Hr1G054510 | AT1G75390.1 |
| 213 | HORVU3Hr1G105680 | AT4G16260.1 |
| 214 | HORVU5Hr1G006460 | AT2G38470.1 |
| 215 | HORVU5Hr1G123330 | AT2G37810.1 |
| 216 | HORVU3Hr1G096890 | AT4G29110.1 |
| 217 | HORVU3Hr1G087690 | AT2G38290.1 |
| 218 | HORVU1Hr1G057880 | AT1G12920.1 |
| 219 | HORVU1Hr1G057650 | AT5G33406.1 |
| 220 | HORVU6Hr1G020570 | AT5G60990.1 |
| 221 | HORVU7Hr1G059520 | AT4G31210.1 |
| 222 | HORVU0Hr1G000760 | AT4G19140.1 |
| 223 | HORVU6Hr1G058820 | AT2G37040.1 |
| 224 | HORVU2Hr1G120010 | AT5G23240.1 |
| 225 | HORVU7Hr1G096870 | AT3G20220.1 |
| 226 | HORVU7Hr1G107630 | AT1G14600.1 |
| 227 | HORVU7Hr1G083270 | AT3G56400.1 |
| 228 | HORVU6Hr1G088430 | AT1G49390.1 |
| 229 | HORVU5Hr1G006480 | AT5G66750.1 |
| 230 | HORVU4Hr1G089560 | AT5G42900.2 |

| | | |
|---|---|---|
| 231 | HORVU2Hr1G081510 | AT1G55775.1 |
| 232 | HORVU2Hr1G070880 | AT5G13220.1 |
| 233 | HORVU7Hr1G033430 | AT3G14200.1 |
| 234 | HORVU7Hr1G010340 | AT5G39080.1 |
| 235 | HORVU2Hr1G094330 | AT5G41890.1 |
| 236 | HORVU7Hr1G002850 | AT3G01610.2 |
| 237 | HORVU5Hr1G034300 | AT5G10140.4 |
| 238 | HORVU2Hr1G126100 | AT1G72125.1 |
| 239 | HORVU7Hr1G028660 | AT3G17240.3 |
| 240 | HORVU2Hr1G070700 | AT5G41080.2 |
| 241 | HORVU4Hr1G000700 | AT3G23240.1 |
| 242 | HORVU2Hr1G103930 | AT3G12977.1 |
| 243 | HORVU5Hr1G045580 | AT3G28917.1 |
| 244 | HORVU4Hr1G039300 | AT1G68050.1 |
| 245 | HORVU5Hr1G057320 | AT1G13580.3 |
| 246 | HORVU5Hr1G093350 | AT3G13050.4 |
| 247 | HORVU1Hr1G092310 | AT4G26830.1 |
| 248 | HORVU7Hr1G076480 | AT5G49900.3 |
| 249 | HORVU3Hr1G062920 | AT3G49210.2 |
| 250 | HORVU2Hr1G082370 | AT1G14130.1 |
| 251 | HORVU0Hr1G000340 | AT1G64660.1 |
| 252 | HORVU5Hr1G050970 | AT2G24100.1 |
| 253 | HORVU3Hr1G028870 | AT4G19510.9 |
| 254 | HORVU5Hr1G104670 | AT3G62760.1 |
| 255 | HORVU3Hr1G013650 | AT4G29210.1 |
| 256 | HORVU0Hr1G015930 | AT1G10030.1 |
| 257 | HORVU5Hr1G081500 | AT4G22920.2 |
| 258 | HORVU2Hr1G099640 | AT2G34940.1 |
| 259 | HORVU6Hr1G014340 | AT2G40660.1 |
| 260 | HORVU1Hr1G026200 | AT5G04160.1 |
| 261 | HORVU5Hr1G006330 | AT3G62760.1 |
| 262 | HORVU2Hr1G117620 | #N/A |
| 263 | HORVU2Hr1G108290 | AT3G20660.1 |
| 264 | HORVU7Hr1G083370 | AT5G13800.3 |
| 265 | HORVU6Hr1G078110 | AT1G22400.1 |
| 266 | HORVU2Hr1G017360 | AT5G61430.1 |
| 267 | HORVU1Hr1G070220 | AT3G08860.1 |
| 268 | HORVU6Hr1G070740 | AT2G41905.1 |
| 269 | HORVU6Hr1G077280 | AT4G33945.1 |
| 270 | HORVU4Hr1G021040 | AT5G52340.1 |
| 271 | HORVU5Hr1G048810 | AT2G17710.1 |

| | | |
|---|---|---|
| 272 | HORVU0Hr1G039620 | AT1G21230.1 |
| 273 | HORVU5Hr1G085780 | AT1G29450.2 |
| 274 | HORVU6Hr1G090630 | AT5G06570.2 |
| 275 | HORVU5Hr1G115780 | AT4G02340.1 |
| 276 | HORVU2Hr1G044650 | AT4G23180.1 |
| 277 | HORVU2Hr1G042520 | AT4G21390.1 |
| 278 | HORVU3Hr1G051820 | AT2G05940.1 |
| 279 | HORVU7Hr1G048570 | AT3G18180.1 |
| 280 | HORVU3Hr1G064450 | AT3G02600.9 |
| 281 | HORVU1Hr1G002900 | AT3G14470.2 |
| 282 | HORVU3Hr1G066220 | AT3G55130.1 |
| 283 | HORVU5Hr1G061770 | AT5G60360.1 |
| 284 | HORVU1Hr1G059130 | AT3G05640.3 |
| 285 | HORVU5Hr1G057740 | AT2G26110.1 |
| 286 | HORVU5Hr1G075470 | AT4G14200.1 |
| 287 | HORVU4Hr1G043910 | AT1G21750.1 |
| 288 | HORVU6Hr1G067380 | AT1G79180.1 |
| 289 | HORVU1Hr1G026520 | AT3G62160.1 |
| 290 | HORVU2Hr1G098110 | AT5G47240.1 |
| 291 | HORVU4Hr1G038810 | AT2G34140.1 |
| 292 | HORVU2Hr1G000120 | AT1G59960.1 |
| 293 | HORVU1Hr1G049400 | AT4G25990.3 |
| 294 | HORVU5Hr1G106640 | AT5G07280.1 |
| 295 | HORVU6Hr1G000070 | AT5G66170.3 |
| 296 | HORVU7Hr1G077470 | AT2G13650.3 |
| 297 | HORVU6Hr1G013610 | AT2G13290.2 |
| 298 | HORVU5Hr1G019880 | AT5G39380.5 |
| 299 | HORVU1Hr1G083700 | AT5G57500.1 |
| 300 | HORVU3Hr1G028860 | AT2G34960.1 |
| 301 | HORVU6Hr1G028590 | AT3G01430.1 |
| 302 | HORVU5Hr1G084740 | AT4G35790.3 |
| 303 | HORVU4Hr1G076970 | AT3G23490.3 |
| 304 | HORVU7Hr1G079160 | AT2G29220.1 |
| 305 | HORVU6Hr1G067740 | AT2G02040.1 |
| 306 | HORVU7Hr1G048970 | AT4G34050.1 |
| 307 | HORVU0Hr1G007750 | AT2G44130.1 |
| 308 | HORVU4Hr1G062440 | AT4G37870.1 |
| 309 | HORVU4Hr1G000070 | AT3G45750.1 |
| 310 | HORVU7Hr1G046130 | AT5G42740.2 |
| 311 | HORVU1Hr1G085720 | AT1G09480.2 |
| 312 | HORVU3Hr1G116350 | AT2G28840.2 |

| | | |
|---|---|---|
| 313 | HORVU5Hr1G046900 | AT5G05340.1 |
| 314 | HORVU2Hr1G066200 | AT1G71070.1 |
| 315 | HORVU2Hr1G081790 | AT5G09220.1 |
| 316 | HORVU7Hr1G051060 | AT1G66880.1 |
| 317 | HORVU2Hr1G117870 | AT4G21390.1 |
| 318 | HORVU1Hr1G092240 | AT5G55180.2 |
| 319 | HORVU7Hr1G044160 | AT4G11000.1 |
| 320 | HORVU7Hr1G029370 | AT5G35360.3 |
| 321 | HORVU1Hr1G037550 | AT5G01520.1 |
| 322 | HORVU6Hr1G008880 | AT5G42020.1 |
| 323 | HORVU2Hr1G109950 | AT1G30450.4 |
| 324 | HORVU5Hr1G109290 | AT2G44310.1 |
| 325 | HORVU1Hr1G021160 | AT3G62760.1 |
| 326 | HORVU0Hr1G015180 | AT4G12080.1 |
| 327 | HORVU2Hr1G089760 | AT5G18700.1 |
| 328 | HORVU1Hr1G065820 | AT3G14460.1 |
| 329 | HORVU3Hr1G076890 | AT4G21534.2 |
| 330 | HORVU3Hr1G105190 | AT3G57260.2 |
| 331 | HORVU2Hr1G017460 | AT1G69490.1 |
| 332 | HORVU2Hr1G114190 | AT2G43820.1 |
| 333 | HORVU3Hr1G001060 | AT1G49000.1 |
| 334 | HORVU2Hr1G093040 | AT4G00060.4 |
| 335 | HORVU3Hr1G076400 | AT2G32210.1 |
| 336 | HORVU2Hr1G107100 | AT5G54760.4 |
| 337 | HORVU5Hr1G051960 | AT1G77700.1 |
| 338 | HORVU2Hr1G090200 | AT3G53150.1 |
| 339 | HORVU6Hr1G030880 | AT2G20850.1 |
| 340 | HORVU7Hr1G096660 | AT4G27230.2 |
| 341 | HORVU3Hr1G062940 | AT5G53390.1 |
| 342 | HORVU5Hr1G005180 | AT4G11650.1 |
| 343 | HORVU2Hr1G109370 | AT4G13940.1 |
| 344 | HORVU4Hr1G000810 | AT1G63720.1 |
| 345 | HORVU2Hr1G001940 | AT4G25120.1 |
| 346 | HORVU3Hr1G105870 | AT2G01170.1 |
| 347 | HORVU4Hr1G011430 | AT5G58730.1 |
| 348 | HORVU6Hr1G081850 | AT3G14067.1 |
| 349 | HORVU5Hr1G019820 | AT5G40010.1 |
| 350 | HORVU7Hr1G043690 | AT4G01070.1 |
| 351 | HORVU6Hr1G035420 | AT3G19450.1 |
| 352 | HORVU7Hr1G000970 | AT5G42830.1 |
| 353 | HORVU4Hr1G067490 | AT1G51405.2 |

| | | |
|---|---|---|
| 354 | HORVU4Hr1G014970 | AT2G44130.1 |
| 355 | HORVU1Hr1G075950 | AT3G16330.1 |
| 356 | HORVU4Hr1G069530 | AT5G43150.1 |
| 357 | HORVU7Hr1G011250 | AT5G07270.1 |
| 358 | HORVU2Hr1G118080 | AT5G52740.1 |
| 359 | HORVU1Hr1G070690 | AT5G01300.1 |
| 360 | HORVU4Hr1G023680 | AT1G19715.2 |
| 361 | HORVU3Hr1G027700 | AT1G53280.1 |
| 362 | HORVU2Hr1G001790 | AT5G16880.4 |
| 363 | HORVU4Hr1G065800 | AT1G32230.3 |
| 364 | HORVU3Hr1G006640 | AT4G23180.1 |
| 365 | HORVU7Hr1G090280 | AT5G20190.1 |
| 366 | HORVU1Hr1G002600 | AT1G70250.1 |
| 367 | HORVU3Hr1G066450 | AT1G51760.1 |
| 368 | HORVU6Hr1G085870 | AT5G35160.4 |
| 369 | HORVU6Hr1G032710 | AT3G06040.3 |
| 370 | HORVU2Hr1G072210 | AT5G53460.3 |
| 371 | HORVU0Hr1G009390 | AT5G39110.1 |
| 372 | HORVU7Hr1G050660 | AT3G45140.2 |
| 373 | HORVU3Hr1G070560 | AT5G19740.1 |
| 374 | HORVU1Hr1G020690 | AT5G06730.1 |
| 375 | HORVU5Hr1G050240 | AT4G35220.1 |
| 376 | HORVU5Hr1G098030 | AT2G15960.1 |
| 377 | HORVU3Hr1G001790 | AT2G29050.2 |
| 378 | HORVU7Hr1G031430 | AT5G11210.3 |
| 379 | HORVU6Hr1G000180 | AT3G13470.1 |
| 380 | HORVU5Hr1G084900 | AT5G55180.1 |
| 381 | HORVU1Hr1G012680 | AT2G36970.1 |
| 382 | HORVU6Hr1G090170 | AT3G04400.2 |
| 383 | HORVU1Hr1G003720 | AT3G09700.1 |
| 384 | HORVU1Hr1G093890 | AT1G78780.5 |
| 385 | HORVU2Hr1G019180 | AT2G47650.1 |
| 386 | HORVU2Hr1G099530 | AT1G17120.1 |
| 387 | HORVU2Hr1G027370 | AT3G57150.1 |
| 388 | HORVU2Hr1G110120 | AT4G13940.1 |
| 389 | HORVU2Hr1G031410 | AT1G72480.1 |
| 390 | HORVU1Hr1G056600 | AT3G51990.1 |
| 391 | HORVU5Hr1G070360 | AT1G21240.1 |
| 392 | HORVU5Hr1G016400 | AT4G14305.1 |
| 393 | HORVU4Hr1G085750 | AT4G11820.2 |
| 394 | HORVU1Hr1G074840 | AT5G17210.1 |

| | | |
|---|---|---|
| 395 | HORVU5Hr1G080310 | AT4G25470.1 |
| 396 | HORVU6Hr1G009420 | AT3G59100.1 |
| 397 | HORVU1Hr1G084620 | AT4G15630.1 |
| 398 | HORVU4Hr1G090130 | AT5G39110.1 |
| 399 | HORVU7Hr1G050680 | AT3G45140.1 |
| 400 | HORVU1Hr1G043920 | AT2G15220.1 |
| 401 | HORVU3Hr1G117370 | AT3G62760.1 |
| 402 | HORVU6Hr1G052060 | AT5G41240.1 |
| 403 | HORVU4Hr1G060280 | AT1G76500.1 |
| 404 | HORVU5Hr1G084860 | AT3G45860.1 |
| 405 | HORVU2Hr1G022500 | AT3G03770.3 |
| 406 | HORVU7Hr1G108450 | AT5G20480.2 |
| 407 | HORVU7Hr1G033520 | AT5G10530.1 |
| 408 | HORVU1Hr1G051200 | AT1G32090.1 |
| 409 | HORVU7Hr1G118610 | AT1G68690.3 |
| 410 | HORVU4Hr1G088860 | AT1G64650.1 |
| 411 | HORVU6Hr1G068160 | AT5G61120.3 |
| 412 | HORVU2Hr1G023450 | AT3G63470.1 |
| 413 | HORVU4Hr1G023740 | AT1G72300.1 |
| 414 | HORVU3Hr1G048610 | AT3G60540.3 |
| 415 | HORVU6Hr1G089510 | AT1G02335.1 |
| 416 | HORVU6Hr1G086560 | AT2G25620.2 |
| 417 | HORVU3Hr1G004850 | AT4G01070.1 |
| 418 | HORVU1Hr1G090430 | AT1G30820.1 |
| 419 | HORVU5Hr1G096950 | AT5G67310.1 |
| 420 | HORVU2Hr1G105240 | AT1G44750.1 |
| 421 | HORVU2Hr1G090260 | AT2G37860.4 |
| 422 | HORVU7Hr1G090950 | AT5G63400.1 |
| 423 | HORVU1Hr1G000660 | AT1G35710.1 |
| 424 | HORVU1Hr1G045110 | AT3G47160.1 |
| 425 | HORVU4Hr1G052010 | AT2G46400.1 |
| 426 | HORVU3Hr1G063840 | AT1G77420.1 |
| 427 | HORVU3Hr1G070540 | AT5G19740.1 |
| 428 | HORVU7Hr1G040730 | AT5G57625.1 |
| 429 | HORVU2Hr1G090280 | AT1G65730.1 |
| 430 | HORVU2Hr1G043400 | #N/A |
| 431 | HORVU2Hr1G108460 | AT3G16660.2 |
| 432 | HORVU2Hr1G095330 | AT3G29810.1 |
| 433 | HORVU2Hr1G043890 | AT5G24090.1 |
| 434 | HORVU4Hr1G000910 | AT3G56070.2 |
| 435 | HORVU7Hr1G048690 | AT1G55760.1 |

| | | |
|---|---|---|
| 436 | HORVU4Hr1G078830 | AT4G16660.1 |
| 437 | HORVU5Hr1G103770 | AT5G06570.2 |
| 438 | HORVU7Hr1G091350 | AT2G46690.1 |
| 439 | HORVU2Hr1G032360 | AT5G61412.1 |
| 440 | HORVU7Hr1G106960 | AT1G07090.1 |
| 441 | HORVU2Hr1G066920 | AT1G60730.3 |
| 442 | HORVU4Hr1G066270 | AT5G42650.1 |
| 443 | HORVU2Hr1G044910 | AT4G23180.3 |
| 444 | HORVU2Hr1G034370 | AT1G74360.1 |
| 445 | HORVU4Hr1G021000 | AT3G10195.1 |
| 446 | HORVU6Hr1G070710 | AT3G12650.1 |
| 447 | HORVU2Hr1G065100 | AT2G39950.9 |
| 448 | HORVU5Hr1G093160 | AT4G16630.1 |
| 449 | HORVU5Hr1G074360 | AT5G35320.1 |
| 450 | HORVU4Hr1G019410 | AT4G09510.1 |
| 451 | HORVU5Hr1G062510 | AT1G56220.2 |
| 452 | HORVU3Hr1G062430 | AT4G21700.1 |
| 453 | HORVU2Hr1G044680 | AT4G23180.1 |
| 454 | HORVU2Hr1G024160 | AT2G38000.1 |
| 455 | HORVU5Hr1G066130 | AT3G47510.1 |
| 456 | HORVU4Hr1G064010 | AT1G77280.2 |
| 457 | HORVU0Hr1G018830 | AT5G55280.1 |
| 458 | HORVU3Hr1G071880 | AT1G06475.1 |
| 459 | HORVU2Hr1G009580 | AT3G62700.1 |
| 460 | HORVU4Hr1G088720 | AT3G60340.2 |
| 461 | HORVU2Hr1G022540 | AT2G03370.2 |
| 462 | HORVU5Hr1G079180 | AT2G43420.1 |
| 463 | HORVU5Hr1G104050 | AT1G28220.1 |
| 464 | HORVU1Hr1G040940 | AT4G16660.1 |
| 465 | HORVU0Hr1G020420 | AT5G37670.1 |
| 466 | HORVU7Hr1G116060 | AT3G48310.1 |
| 467 | HORVU7Hr1G048430 | AT3G18180.1 |
| 468 | HORVU4Hr1G012510 | AT3G10020.1 |
| 469 | HORVU0Hr1G015140 | AT5G09850.5 |
| 470 | HORVU1Hr1G042370 | AT1G70520.1 |
| 471 | HORVU7Hr1G042260 | AT5G07990.1 |
| 472 | HORVU1Hr1G073330 | AT2G39980.1 |
| 473 | HORVU4Hr1G046610 | AT1G56210.1 |
| 474 | HORVU0Hr1G005070 | AT1G47056.1 |
| 475 | HORVU5Hr1G055060 | AT3G02875.1 |
| 476 | HORVU1Hr1G001950 | AT1G76680.1 |

| | | |
|---|---|---|
| 477 | HORVU7Hr1G043810 | AT4G11000.1 |
| 478 | HORVU2Hr1G084200 | AT3G11402.1 |
| 479 | HORVU3Hr1G078840 | AT2G30150.2 |
| 480 | HORVU2Hr1G064530 | AT1G05200.6 |
| 481 | HORVU1Hr1G017050 | AT3G62760.1 |
| 482 | HORVU2Hr1G039050 | AT2G26975.1 |
| 483 | HORVU2Hr1G073320 | AT5G39740.2 |
| 484 | HORVU2Hr1G010590 | AT3G47570.1 |
| 485 | HORVU1Hr1G047050 | AT1G02110.1 |
| 486 | HORVU2Hr1G122290 | AT1G48720.1 |
| 487 | HORVU2Hr1G089160 | AT2G22230.1 |
| 488 | HORVU4Hr1G078470 | AT5G42250.1 |
| 489 | HORVU3Hr1G106720 | AT5G14800.1 |
| 490 | HORVU2Hr1G018570 | AT5G05340.1 |
| 491 | HORVU7Hr1G079830 | AT2G18730.1 |
| 492 | HORVU5Hr1G060070 | AT1G70160.1 |
| 493 | HORVU6Hr1G085370 | AT5G60390.3 |
| 494 | HORVU6Hr1G069400 | AT5G14780.3 |
| 495 | HORVU4Hr1G048700 | AT2G47890.1 |
| 496 | HORVU2Hr1G023470 | AT2G33100.2 |
| 497 | HORVU1Hr1G053070 | AT4G33540.2 |
| 498 | HORVU3Hr1G117760 | AT5G03160.1 |
| 499 | HORVU5Hr1G105480 | AT1G47480.1 |
| 500 | HORVU2Hr1G022350 | AT3G57220.1 |
| 501 | HORVU1Hr1G072250 | AT4G20095.1 |
| 502 | HORVU7Hr1G118450 | AT4G38830.1 |
| 503 | HORVU4Hr1G081100 | AT3G03190.1 |
| 504 | HORVU2Hr1G121990 | AT1G56340.2 |
| 505 | HORVU6Hr1G043980 | AT3G09455.2 |
| 506 | HORVU7Hr1G086580 | AT5G57035.1 |
| 507 | HORVU7Hr1G088770 | AT2G20420.1 |
| 508 | HORVU5Hr1G073580 | AT2G44430.1 |
| 509 | HORVU2Hr1G109380 | AT3G10410.1 |
| 510 | HORVU1Hr1G079070 | AT5G47770.1 |
| 511 | HORVU7Hr1G026720 | AT5G25940.1 |
| 512 | HORVU7Hr1G047080 | AT5G52450.1 |
| 513 | HORVU7Hr1G071020 | AT3G43430.1 |
| 514 | HORVU7Hr1G083180 | AT1G13380.1 |
| 515 | HORVU4Hr1G013330 | AT1G24625.1 |
| 516 | HORVU0Hr1G027290 | AT3G18850.4 |
| 517 | HORVU6Hr1G018420 | AT5G61790.1 |

| | | |
|---|---|---|
| 518 | HORVU1Hr1G001660 | AT1G66920.2 |
| 519 | HORVU3Hr1G095820 | AT5G50860.1 |
| 520 | HORVU7Hr1G078530 | AT1G63500.1 |
| 521 | HORVU5Hr1G002120 | AT2G18390.1 |
| 522 | HORVU1Hr1G053440 | AT5G38710.1 |
| 523 | HORVU5Hr1G002150 | AT4G39260.3 |
| 524 | HORVU2Hr1G077990 | AT4G05460.1 |
| 525 | HORVU3Hr1G033550 | AT3G17180.2 |
| 526 | HORVU1Hr1G068170 | AT5G12250.1 |
| 527 | HORVU5Hr1G115250 | AT1G19530.2 |
| 528 | HORVU3Hr1G085930 | AT2G29380.1 |
| 529 | HORVU1Hr1G092290 | AT4G23180.1 |
| 530 | HORVU4Hr1G002270 | AT5G17920.2 |
| 531 | HORVU4Hr1G021010 | AT5G24470.1 |
| 532 | HORVU7Hr1G121700 | AT4G35090.1 |
| 533 | HORVU2Hr1G111600 | AT1G68300.1 |
| 534 | HORVU1Hr1G002460 | AT1G66910.1 |
| 535 | HORVU5Hr1G019110 | AT3G19553.1 |
| 536 | HORVU2Hr1G072180 | AT1G08250.1 |
| 537 | HORVU5Hr1G087780 | AT4G34880.2 |
| 538 | HORVU6Hr1G065240 | AT5G42900.3 |
| 539 | HORVU2Hr1G121050 | AT1G65840.1 |
| 540 | HORVU3Hr1G004140 | AT1G23580.1 |
| 541 | HORVU5Hr1G054060 | AT5G26030.2 |
| 542 | HORVU7Hr1G081910 | AT3G10950.1 |
| 543 | HORVU7Hr1G002130 | AT3G47570.1 |
| 544 | HORVU5Hr1G056370 | AT1G70480.1 |
| 545 | HORVU4Hr1G074450 | AT3G07030.3 |
| 546 | HORVU2Hr1G121070 | AT2G37250.1 |
| 547 | HORVU6Hr1G061140 | AT5G66320.2 |
| 548 | HORVU4Hr1G069320 | AT1G47240.1 |
| 549 | HORVU1Hr1G054560 | AT3G60190.1 |
| 550 | HORVU3Hr1G082590 | AT1G34340.1 |
| 551 | HORVU1Hr1G045220 | AT2G23790.1 |
| 552 | HORVU3Hr1G088570 | AT2G36460.1 |
| 553 | HORVU3Hr1G058170 | AT1G80300.1 |
| 554 | HORVU1Hr1G049900 | AT4G35470.1 |
| 555 | HORVU0Hr1G005210 | AT2G28400.1 |
| 556 | HORVU5Hr1G073990 | AT1G20970.1 |
| 557 | HORVU3Hr1G067910 | AT1G34060.1 |
| 558 | HORVU6Hr1G008930 | AT3G47570.1 |

| | | |
|---|---|---|
| 559 | HORVU6Hr1G070270 | AT1G76550.1 |
| 560 | HORVU4Hr1G056610 | AT3G01090.2 |
| 561 | HORVU2Hr1G020680 | AT1G54610.3 |
| 562 | HORVU6Hr1G062830 | AT5G51290.1 |
| 563 | HORVU7Hr1G089960 | AT3G15730.1 |
| 564 | HORVU3Hr1G090730 | AT1G33780.1 |
| 565 | HORVU7Hr1G008050 | AT1G35710.1 |
| 566 | HORVU7Hr1G034350 | AT2G26310.5 |
| 567 | HORVU3Hr1G080830 | AT2G30490.1 |
| 568 | HORVU6Hr1G059940 | AT5G08350.1 |
| 569 | HORVU5Hr1G077120 | AT1G54570.1 |
| 570 | HORVU2Hr1G036440 | AT4G16660.1 |
| 571 | HORVU6Hr1G064650 | AT1G76240.1 |
| 572 | HORVU0Hr1G000050 | AT5G38240.2 |
| 573 | HORVU1Hr1G045630 | AT3G48890.1 |
| 574 | HORVU2Hr1G090730 | AT5G23370.1 |
| 575 | HORVU5Hr1G056790 | AT4G29510.1 |
| 576 | HORVU2Hr1G118950 | AT3G09010.1 |
| 577 | HORVU0Hr1G009430 | AT3G05950.1 |
| 578 | HORVU4Hr1G074770 | AT1G13990.1 |
| 579 | HORVU2Hr1G123030 | AT1G63220.1 |
| 580 | HORVU3Hr1G090920 | AT1G01720.1 |
| 581 | HORVU6Hr1G076510 | AT1G24140.1 |
| 582 | HORVU3Hr1G094720 | AT3G01960.1 |
| 583 | HORVU3Hr1G081300 | AT5G01830.1 |
| 584 | HORVU6Hr1G018370 | AT5G61790.1 |
| 585 | HORVU1Hr1G047570 | AT5G24660.1 |
| 586 | HORVU7Hr1G055900 | AT3G59330.2 |
| 587 | HORVU2Hr1G125110 | AT1G71695.1 |
| 588 | HORVU3Hr1G054640 | AT1G60690.1 |
| 589 | HORVU7Hr1G045140 | AT1G14620.1 |
| 590 | HORVU4Hr1G079330 | AT3G44880.1 |
| 591 | HORVU2Hr1G022140 | AT3G12860.1 |
| 592 | HORVU0Hr1G006610 | AT1G80070.1 |
| 593 | HORVU7Hr1G033390 | AT2G25110.1 |
| 594 | HORVU6Hr1G076520 | AT1G19020.1 |
| 595 | HORVU2Hr1G091130 | AT3G15360.1 |
| 596 | HORVU2Hr1G049420 | AT4G17790.1 |
| 597 | HORVU6Hr1G053140 | AT1G27320.1 |
| 598 | HORVU4Hr1G007550 | AT1G48480.1 |
| 599 | HORVU2Hr1G046590 | AT5G57700.3 |

| | | |
|---|---|---|
| 600 | HORVU5Hr1G058070 | AT1G78130.1 |
| 601 | HORVU3Hr1G100350 | AT1G03220.1 |
| 602 | HORVU3Hr1G081880 | AT1G22882.1 |
| 603 | HORVU4Hr1G058340 | AT3G14690.1 |
| 604 | HORVU5Hr1G069090 | AT1G47480.1 |
| 605 | HORVU1Hr1G025890 | AT1G15460.2 |
| 606 | HORVU6Hr1G067430 | AT5G64410.1 |
| 607 | HORVU7Hr1G119870 | AT1G80245.7 |
| 608 | HORVU1Hr1G064860 | AT5G16440.1 |
| 609 | HORVU3Hr1G073150 | AT3G57260.1 |
| 610 | HORVU5Hr1G007550 | AT5G22030.1 |
| 611 | HORVU4Hr1G022170 | AT3G52710.1 |
| 612 | HORVU2Hr1G043860 | AT4G27230.2 |
| 613 | HORVU5Hr1G013360 | AT5G62300.2 |
| 614 | HORVU6Hr1G075590 | AT1G14330.1 |
| 615 | HORVU4Hr1G023260 | AT2G30520.1 |
| 616 | HORVU3Hr1G057970 | AT1G66730.1 |
| 617 | HORVU7Hr1G110630 | AT3G07040.1 |
| 618 | HORVU3Hr1G040710 | AT1G80270.5 |
| 619 | HORVU1Hr1G022810 | AT2G47470.1 |
| 620 | HORVU5Hr1G048890 | AT3G27740.1 |
| 621 | HORVU1Hr1G022020 | AT3G14360.1 |
| 622 | HORVU2Hr1G016650 | AT3G07050.1 |
| 623 | HORVU2Hr1G089810 | AT1G58190.1 |
| 624 | HORVU3Hr1G078850 | AT2G30150.2 |
| 625 | HORVU1Hr1G033630 | AT1G04985.1 |
| 626 | HORVU5Hr1G106960 | AT2G39210.1 |
| 627 | HORVU3Hr1G090410 | AT3G59880.1 |
| 628 | HORVU5Hr1G024420 | AT1G27320.1 |
| 629 | HORVU2Hr1G113460 | AT1G47128.1 |
| 630 | HORVU5Hr1G026120 | AT3G03800.2 |
| 631 | HORVU5Hr1G125130 | AT2G28000.1 |
| 632 | HORVU1Hr1G094070 | AT4G10140.1 |
| 633 | HORVU2Hr1G072830 | AT3G48410.1 |
| 634 | HORVU3Hr1G006290 | AT3G16565.6 |
| 635 | HORVU7Hr1G099390 | AT4G11530.1 |
| 636 | HORVU1Hr1G044960 | AT4G27460.1 |
| 637 | HORVU2Hr1G046550 | AT1G61930.1 |
| 638 | HORVU2Hr1G020690 | AT5G27120.1 |
| 639 | HORVU5Hr1G117900 | AT5G09810.1 |
| 640 | HORVU1Hr1G043710 | AT1G03610.1 |

| | | |
|---|---|---|
| 641 | HORVU5Hr1G085020 | AT4G21410.1 |
| 642 | HORVU6Hr1G068150 | AT3G22370.1 |
| 643 | HORVU1Hr1G072060 | AT5G59850.1 |
| 644 | HORVU7Hr1G045420 | AT3G57280.1 |
| 645 | HORVU3Hr1G080500 | AT4G13290.1 |
| 646 | HORVU3Hr1G057550 | AT3G18580.1 |
| 647 | HORVU2Hr1G092880 | AT5G24380.1 |
| 648 | HORVU3Hr1G098400 | AT2G23620.1 |
| 649 | HORVU7Hr1G088260 | AT1G14520.4 |
| 650 | HORVU1Hr1G021760 | AT3G06190.1 |
| 651 | HORVU7Hr1G117000 | AT4G24190.1 |
| 652 | HORVU6Hr1G002030 | AT1G32580.1 |
| 653 | HORVU4Hr1G010470 | AT3G18830.1 |
| 654 | HORVU1Hr1G015770 | AT5G40380.1 |
| 655 | HORVU1Hr1G047390 | AT2G46690.1 |
| 656 | HORVU6Hr1G001980 | AT5G07840.1 |
| 657 | HORVU5Hr1G104850 | AT2G37710.1 |
| 658 | HORVU5Hr1G000800 | AT5G62680.1 |
| 659 | HORVU6Hr1G014960 | AT4G05050.3 |
| 660 | HORVU7Hr1G046980 | AT5G02050.1 |
| 661 | HORVU6Hr1G055250 | AT2G22570.1 |
| 662 | HORVU2Hr1G056750 | AT5G44500.2 |
| 663 | HORVU5Hr1G077020 | AT1G53260.1 |
| 664 | HORVU3Hr1G040350 | AT4G31480.9 |
| 665 | HORVU2Hr1G006140 | #N/A |
| 666 | HORVU7Hr1G100100 | AT5G02560.1 |
| 667 | HORVU4Hr1G003180 | AT3G12920.1 |
| 668 | HORVU2Hr1G075010 | AT2G23420.1 |
| 669 | HORVU4Hr1G003060 | AT4G00570.1 |
| 670 | HORVU6Hr1G014910 | AT1G65349.1 |
| 671 | HORVU0Hr1G013910 | AT3G29060.1 |
| 672 | HORVU1Hr1G068030 | AT2G40890.1 |
| 673 | HORVU5Hr1G008300 | AT5G62300.2 |
| 674 | HORVU2Hr1G056490 | AT2G40030.1 |
| 675 | HORVU5Hr1G008820 | AT1G65030.1 |
| 676 | HORVU0Hr1G022290 | AT5G10530.1 |
| 677 | HORVU2Hr1G028510 | AT5G15200.1 |
| 678 | HORVU5Hr1G071270 | AT3G22730.1 |
| 679 | HORVU6Hr1G063560 | AT3G17390.1 |
| 680 | HORVU6Hr1G075400 | AT1G71820.1 |
| 681 | HORVU3Hr1G083830 | AT2G37220.1 |

| | | |
|---|---|---|
| 682 | HORVU3Hr1G116740 | AT5G59780.3 |
| 683 | HORVU5Hr1G098190 | AT3G50830.1 |
| 684 | HORVU7Hr1G048810 | AT4G12650.1 |
| 685 | HORVU1Hr1G037760 | AT1G02260.2 |
| 686 | HORVU5Hr1G068230 | AT3G56850.1 |
| 687 | HORVU5Hr1G054180 | AT5G33320.1 |
| 688 | HORVU7Hr1G112980 | AT2G24120.1 |
| 689 | HORVU1Hr1G062900 | AT4G24830.1 |
| 690 | HORVU2Hr1G022490 | AT5G60990.1 |
| 691 | HORVU3Hr1G096550 | AT4G25720.1 |
| 692 | HORVU5Hr1G080170 | AT5G62190.1 |
| 693 | HORVU1Hr1G074590 | AT2G22250.3 |
| 694 | HORVU7Hr1G000850 | AT3G19950.3 |
| 695 | HORVU3Hr1G080940 | AT1G47970.1 |
| 696 | HORVU7Hr1G028290 | AT3G26300.1 |
| 697 | HORVU6Hr1G090810 | AT5G36880.2 |
| 698 | HORVU5Hr1G123490 | AT2G44510.1 |
| 699 | HORVU3Hr1G052980 | AT2G01410.1 |
| 700 | HORVU6Hr1G016150 | AT4G05050.3 |
| 701 | HORVU7Hr1G085180 | AT1G14360.1 |
| 702 | HORVU2Hr1G031700 | AT4G30140.1 |
| 703 | HORVU7Hr1G013710 | AT2G45300.4 |
| 704 | HORVU3Hr1G038060 | AT2G44970.3 |
| 705 | HORVU1Hr1G044020 | AT1G04160.1 |
| 706 | HORVU2Hr1G085160 | AT2G43330.1 |
| 707 | HORVU2Hr1G005520 | AT3G17210.2 |
| 708 | HORVU2Hr1G005990 | AT4G19975.1 |
| 709 | HORVU5Hr1G118010 | AT3G26420.1 |
| 710 | HORVU5Hr1G027590 | AT2G41900.1 |
| 711 | HORVU3Hr1G066810 | AT2G35990.1 |
| 712 | HORVU5Hr1G028030 | AT3G18830.1 |
| 713 | HORVU5Hr1G044780 | AT3G24090.1 |
| 714 | HORVU3Hr1G023780 | AT5G42830.1 |
| 715 | HORVU5Hr1G113180 | AT5G02500.1 |
| 716 | HORVU3Hr1G092660 | AT4G21150.3 |
| 717 | HORVU6Hr1G091860 | AT5G52470.1 |
| 718 | HORVU6Hr1G021700 | AT5G20620.1 |
| 719 | HORVU5Hr1G011460 | AT3G22330.1 |
| 720 | HORVU4Hr1G047580 | AT3G03920.1 |
| 721 | HORVU4Hr1G085800 | AT5G10625.1 |
| 722 | HORVU4Hr1G043760 | AT1G34420.1 |

| | | |
|---|---|---|
| 723 | HORVU5Hr1G096480 | AT1G72310.1 |
| 724 | HORVU3Hr1G019480 | AT5G56840.1 |
| 725 | HORVU6Hr1G016200 | AT4G05320.7 |
| 726 | HORVU5Hr1G001090 | AT4G34610.4 |
| 727 | HORVU3Hr1G006330 | AT3G05230.1 |
| 728 | HORVU5Hr1G109910 | AT3G57490.1 |
| 729 | HORVU7Hr1G100230 | AT1G66240.1 |
| 730 | HORVU7Hr1G012700 | AT5G42090.1 |
| 731 | HORVU5Hr1G061140 | AT3G25860.1 |
| 732 | HORVU4Hr1G003100 | AT4G33090.1 |
| 733 | HORVU1Hr1G050330 | AT3G01990.8 |
| 734 | HORVU4Hr1G009560 | AT2G36630.1 |
| 735 | HORVU4Hr1G064870 | AT2G02040.1 |
| 736 | HORVU2Hr1G110110 | AT4G13930.1 |
| 737 | HORVU3Hr1G024500 | AT1G59700.1 |
| 738 | HORVU7Hr1G036000 | AT5G36210.1 |
| 739 | HORVU4Hr1G066310 | AT4G00955.1 |
| 740 | HORVU7Hr1G010320 | AT2G17780.4 |
| 741 | HORVU7Hr1G108600 | AT4G14240.1 |
| 742 | HORVU6Hr1G058740 | AT1G04400.2 |
| 743 | HORVU4Hr1G062740 | AT4G36480.2 |
| 744 | HORVU7Hr1G077630 | AT5G39450.1 |
| 745 | HORVU2Hr1G043900 | AT5G24090.1 |
| 746 | HORVU3Hr1G067130 | AT4G33000.1 |
| 747 | HORVU5Hr1G067570 | AT1G04980.1 |
| 748 | HORVU1Hr1G088880 | AT2G34420.1 |
| 749 | HORVU5Hr1G075490 | AT3G25220.1 |
| 750 | HORVU2Hr1G120150 | AT5G16370.1 |
| 751 | HORVU0Hr1G027680 | AT5G41970.1 |
| 752 | HORVU1Hr1G064210 | AT4G36020.2 |
| 753 | HORVU5Hr1G026780 | AT4G02980.1 |
| 754 | HORVU5Hr1G120600 | AT3G25040.1 |
| 755 | HORVU1Hr1G008230 | AT3G55270.1 |
| 756 | HORVU7Hr1G018850 | AT4G00850.2 |
| 757 | HORVU3Hr1G049610 | AT1G29900.1 |
| 758 | HORVU3Hr1G040270 | AT4G31480.6 |
| 759 | HORVU3Hr1G085640 | AT3G54000.3 |
| 760 | HORVU1Hr1G074110 | AT1G08460.1 |
| 761 | HORVU4Hr1G046230 | AT1G65290.1 |
| 762 | HORVU7Hr1G009730 | AT1G66930.2 |
| 763 | HORVU7Hr1G031850 | AT1G74270.1 |

| | | |
|---|---|---|
| 764 | HORVU7Hr1G035340 | AT1G18640.2 |
| 765 | HORVU6Hr1G080000 | AT2G46690.1 |
| 766 | HORVU2Hr1G029870 | AT1G22410.1 |
| 767 | HORVU3Hr1G069200 | AT4G12543.1 |
| 768 | HORVU3Hr1G045480 | AT3G06350.1 |
| 769 | HORVU7Hr1G033360 | AT4G10490.1 |
| 770 | HORVU1Hr1G013820 | AT3G45300.1 |
| 771 | HORVU5Hr1G072480 | AT1G21240.1 |
| 772 | HORVU4Hr1G075710 | AT3G09630.1 |
| 773 | HORVU2Hr1G005140 | AT2G45630.2 |
| 774 | HORVU3Hr1G078650 | AT4G32390.1 |
| 775 | HORVU6Hr1G011970 | AT2G46600.1 |
| 776 | HORVU4Hr1G038960 | AT5G15650.1 |
| 777 | HORVU5Hr1G067110 | ATMG00210.1 |
| 778 | HORVU3Hr1G066720 | AT5G57020.1 |
| 779 | HORVU2Hr1G078350 | AT4G25340.1 |
| 780 | HORVU3Hr1G055650 | AT1G64140.1 |
| 781 | HORVU1Hr1G083710 | AT1G18080.1 |
| 782 | HORVU1Hr1G080040 | AT5G56450.1 |
| 783 | HORVU7Hr1G056980 | AT5G24400.1 |
| 784 | HORVU0Hr1G004790 | AT1G21370.2 |
| 785 | HORVU7Hr1G049370 | AT5G56010.1 |
| 786 | HORVU4Hr1G028310 | AT5G59780.3 |
| 787 | HORVU1Hr1G013990 | AT3G61620.2 |
| 788 | HORVU2Hr1G060680 | AT2G20180.7 |
| 789 | HORVU5Hr1G106460 | AT3G54250.1 |
| 790 | HORVU3Hr1G108720 | AT3G27890.1 |
| 791 | HORVU1Hr1G048970 | AT2G42010.1 |
| 792 | HORVU5Hr1G056670 | AT1G07650.4 |
| 793 | HORVU7Hr1G106230 | AT5G01830.1 |
| 794 | HORVU2Hr1G031910 | AT2G31670.1 |
| 795 | HORVU1Hr1G030390 | AT5G53160.1 |
| 796 | HORVU4Hr1G004460 | AT3G47960.1 |
| 797 | HORVU1Hr1G015660 | AT3G62420.1 |
| 798 | HORVU5Hr1G122350 | AT4G02930.1 |
| 799 | HORVU1Hr1G023510 | AT1G56670.1 |
| 800 | HORVU3Hr1G092520 | AT4G34900.1 |
| 801 | HORVU6Hr1G021520 | AT5G05340.1 |
| 802 | HORVU1Hr1G083330 | AT5G10190.1 |
| 803 | HORVU5Hr1G066930 | AT1G08830.2 |
| 804 | HORVU7Hr1G111600 | AT5G50460.1 |

| | | |
|---|---|---|
| 805 | HORVU2Hr1G083940 | AT5G40760.2 |
| 806 | HORVU5Hr1G036790 | AT1G35510.1 |
| 807 | HORVU7Hr1G048330 | AT1G10760.3 |
| 808 | HORVU3Hr1G106140 | AT4G16260.1 |
| 809 | HORVU4Hr1G008700 | AT5G17330.1 |
| 810 | HORVU3Hr1G014730 | AT5G14030.4 |
| 811 | HORVU5Hr1G023530 | AT5G57580.1 |
| 812 | HORVU3Hr1G077000 | AT1G11360.4 |
| 813 | HORVU2Hr1G006250 | AT5G55070.2 |
| 814 | HORVU4Hr1G058840 | AT2G37500.1 |
| 815 | HORVU6Hr1G072910 | AT1G49050.1 |
| 816 | HORVU1Hr1G003600 | AT1G67850.4 |
| 817 | HORVU3Hr1G011460 | AT5G05990.1 |
| 818 | HORVU3Hr1G070270 | AT4G37670.2 |
| 819 | HORVU7Hr1G018190 | AT3G51160.1 |
| 820 | HORVU5Hr1G028550 | AT2G03530.5 |
| 821 | HORVU1Hr1G025670 | AT2G22670.4 |
| 822 | HORVU4Hr1G009040 | AT3G45980.1 |
| 823 | HORVU7Hr1G028910 | AT1G15690.1 |
| 824 | HORVU2Hr1G068670 | AT2G43710.1 |
| 825 | HORVU3Hr1G084170 | AT2G27500.2 |
| 826 | HORVU5Hr1G028870 | AT4G00850.1 |
| 827 | HORVU5Hr1G095140 | AT5G56050.1 |
| 828 | HORVU3Hr1G090500 | AT3G55370.1 |
| 829 | HORVU4Hr1G074270 | AT3G61110.1 |
| 830 | HORVU7Hr1G071240 | AT1G26880.1 |
| 831 | HORVU6Hr1G016180 | AT1G16360.2 |
| 832 | HORVU7Hr1G008690 | AT4G19110.1 |
| 833 | HORVU6Hr1G002280 | AT3G62960.1 |
| 834 | HORVU7Hr1G091740 | AT3G15840.1 |
| 835 | HORVU3Hr1G094450 | AT1G67490.1 |
| 836 | HORVU3Hr1G002280 | AT5G20280.1 |
| 837 | HORVU4Hr1G014130 | AT3G61110.1 |
| 838 | HORVU1Hr1G062940 | AT3G05290.1 |
| 839 | HORVU5Hr1G043190 | AT1G74040.1 |
| 840 | HORVU4Hr1G052210 | AT3G43810.2 |
| 841 | HORVU2Hr1G037940 | AT4G00100.1 |
| 842 | HORVU2Hr1G089490 | AT1G06515.2 |
| 843 | HORVU4Hr1G064710 | AT2G01970.1 |
| 844 | HORVU2Hr1G005320 | AT4G08900.1 |
| 845 | HORVU1Hr1G052040 | AT4G35300.8 |

SUPPLEMENTARY TABLE 5

| S No | 1338 exclusively in "Harrington_Down": | |
|---|---|---|
| 1 | HORVU3Hr1G064080 | AT3G18290.1 |
| 2 | HORVU7Hr1G115990 | AT3G07040.1 |
| 3 | HORVU6Hr1G060720 | AT4G25570.2 |
| 4 | HORVU0Hr1G004760 | AT5G39830.2 |
| 5 | HORVU5Hr1G070160 | AT4G09020.1 |
| 6 | HORVU1Hr1G004430 | AT2G37660.1 |
| 7 | HORVU2Hr1G118740 | AT4G04955.1 |
| 8 | HORVU0Hr1G022500 | AT5G32440.3 |
| 9 | HORVU4Hr1G012240 | AT5G48545.1 |
| 10 | HORVU2Hr1G111840 | AT1G10360.1 |
| 11 | HORVU5Hr1G018840 | AT2G14910.1 |
| 12 | HORVU4Hr1G050230 | AT1G62680.2 |
| 13 | HORVU6Hr1G047770 | AT5G06130.2 |
| 14 | HORVU2Hr1G035220 | AT4G30950.1 |
| 15 | HORVU5Hr1G118340 | AT4G36940.1 |
| 16 | HORVU2Hr1G047020 | #N/A |
| 17 | HORVU5Hr1G020200 | AT4G00830.5 |
| 18 | HORVU4Hr1G071050 | AT1G29840.1 |
| 19 | HORVU4Hr1G078610 | AT1G34630.1 |
| 20 | HORVU3Hr1G030210 | AT2G26040.1 |
| 21 | HORVU1Hr1G041290 | AT1G16400.1 |
| 22 | HORVU4Hr1G077420 | AT1G53730.1 |
| 23 | HORVU5Hr1G074200 | AT3G58500.1 |
| 24 | HORVU6Hr1G035260 | AT3G27160.2 |
| 25 | HORVU5Hr1G078960 | AT5G51720.1 |
| 26 | HORVU7Hr1G116600 | AT3G19810.2 |
| 27 | HORVU4Hr1G069390 | AT5G67380.1 |
| 28 | HORVU7Hr1G118440 | AT3G07040.1 |
| 29 | HORVU3Hr1G056580 | AT1G75350.1 |
| 30 | HORVU6Hr1G014700 | AT5G66210.2 |
| 31 | HORVU6Hr1G069190 | AT3G27820.1 |
| 32 | HORVU5Hr1G120210 | AT1G09630.1 |
| 33 | HORVU4Hr1G059050 | AT3G07020.1 |
| 34 | HORVU6Hr1G074830 | AT1G50000.1 |
| 35 | HORVU6Hr1G034490 | AT2G33580.1 |
| 36 | HORVU6Hr1G067890 | AT1G54130.1 |
| 37 | HORVU6Hr1G079070 | AT4G19100.1 |

| | | |
|---|---|---|
| 38 | HORVU2Hr1G019650 | AT3G05520.1 |
| 39 | HORVU2Hr1G123610 | AT3G26040.1 |
| 40 | HORVU7Hr1G013600 | AT3G60750.1 |
| 41 | HORVU2Hr1G072010 | AT5G24650.1 |
| 42 | HORVU1Hr1G082380 | AT5G47780.1 |
| 43 | HORVU3Hr1G096780 | AT1G67480.3 |
| 44 | HORVU6Hr1G035560 | AT4G13150.5 |
| 45 | HORVU5Hr1G040400 | AT5G06260.3 |
| 46 | HORVU2Hr1G079840 | AT4G39730.1 |
| 47 | HORVU6Hr1G013620 | AT4G14100.1 |
| 48 | HORVU1Hr1G021800 | AT1G69490.1 |
| 49 | HORVU3Hr1G073480 | AT5G48960.1 |
| 50 | HORVU6Hr1G090380 | AT2G33255.1 |
| 51 | HORVU3Hr1G024540 | AT5G25360.3 |
| 52 | HORVU6Hr1G095270 | AT4G38380.5 |
| 53 | HORVU7Hr1G092460 | AT3G30380.2 |
| 54 | HORVU3Hr1G064730 | AT4G29840.1 |
| 55 | HORVU1Hr1G010820 | AT5G02250.1 |
| 56 | HORVU7Hr1G026580 | AT3G29240.2 |
| 57 | HORVU3Hr1G066420 | AT1G34220.1 |
| 58 | HORVU7Hr1G046280 | AT5G42850.2 |
| 59 | HORVU6Hr1G017710 | AT4G22160.2 |
| 60 | HORVU3Hr1G006270 | AT1G18060.1 |
| 61 | HORVU3Hr1G004320 | AT5G19730.1 |
| 62 | HORVU3Hr1G115840 | AT2G39670.1 |
| 63 | HORVU5Hr1G118070 | AT3G49670.1 |
| 64 | HORVU3Hr1G029200 | AT3G14940.2 |
| 65 | HORVU6Hr1G043170 | AT4G38180.1 |
| 66 | HORVU5Hr1G053950 | AT4G19710.2 |
| 67 | HORVU4Hr1G072930 | AT1G64770.1 |
| 68 | HORVU6Hr1G016240 | AT2G35530.1 |
| 69 | HORVU5Hr1G103750 | AT3G10650.2 |
| 70 | HORVU6Hr1G042840 | AT4G35300.5 |
| 71 | HORVU4Hr1G021190 | AT3G44220.1 |
| 72 | HORVU4Hr1G017680 | AT1G21640.1 |
| 73 | HORVU3Hr1G023470 | AT5G15300.1 |
| 74 | HORVU4Hr1G084410 | AT4G01310.1 |
| 75 | HORVU6Hr1G088370 | AT4G32890.1 |
| 76 | HORVU2Hr1G037020 | AT5G19850.1 |
| 77 | HORVU7Hr1G040390 | AT5G22800.1 |
| 78 | HORVU4Hr1G020030 | AT5G37710.1 |

| | | |
|---|---|---|
| 79 | HORVU4Hr1G014070 | AT3G59390.2 |
| 80 | HORVU4Hr1G056060 | AT5G01310.1 |
| 81 | HORVU5Hr1G024250 | AT5G63310.1 |
| 82 | HORVU4Hr1G007420 | AT1G25440.1 |
| 83 | HORVU6Hr1G004980 | AT3G45050.4 |
| 84 | HORVU5Hr1G112340 | AT5G58490.1 |
| 85 | HORVU4Hr1G064580 | AT1G14690.2 |
| 86 | HORVU1Hr1G066910 | #N/A |
| 87 | HORVU5Hr1G031410 | AT5G29624.1 |
| 88 | HORVU4Hr1G068990 | AT2G03440.1 |
| 89 | HORVU3Hr1G097280 | AT1G10130.1 |
| 90 | HORVU6Hr1G014130 | AT5G51545.1 |
| 91 | HORVU4Hr1G057040 | #N/A |
| 92 | HORVU7Hr1G095810 | AT3G18040.4 |
| 93 | HORVU6Hr1G092960 | AT4G00430.1 |
| 94 | HORVU5Hr1G062890 | AT3G48110.1 |
| 95 | HORVU3Hr1G063700 | AT5G38570.1 |
| 96 | HORVU5Hr1G070230 | AT5G64290.1 |
| 97 | HORVU6Hr1G028680 | AT2G25180.1 |
| 98 | HORVU5Hr1G014290 | AT1G04240.1 |
| 99 | HORVU6Hr1G067390 | AT5G16810.2 |
| 100 | HORVU2Hr1G109710 | AT5G39590.1 |
| 101 | HORVU5Hr1G050180 | AT3G19490.1 |
| 102 | HORVU1Hr1G076460 | AT2G16060.1 |
| 103 | HORVU7Hr1G012560 | AT4G34710.2 |
| 104 | HORVU3Hr1G050080 | AT5G67370.1 |
| 105 | HORVU2Hr1G023480 | AT5G13720.1 |
| 106 | HORVU6Hr1G082060 | AT5G20720.4 |
| 107 | HORVU5Hr1G026640 | AT3G04710.1 |
| 108 | HORVU2Hr1G116090 | AT1G56000.2 |
| 109 | HORVU3Hr1G032440 | AT2G20570.1 |
| 110 | HORVU2Hr1G060390 | AT1G30360.1 |
| 111 | HORVU7Hr1G049070 | AT4G34050.1 |
| 112 | HORVU6Hr1G039660 | AT5G54980.1 |
| 113 | HORVU5Hr1G062030 | AT4G37980.1 |
| 114 | HORVU0Hr1G039910 | AT3G55950.1 |
| 115 | HORVU4Hr1G042610 | AT5G19390.2 |
| 116 | HORVU7Hr1G061500 | AT4G23160.3 |
| 117 | HORVU4Hr1G054580 | AT3G05670.3 |
| 118 | HORVU2Hr1G105830 | AT1G78915.1 |
| 119 | HORVU5Hr1G009890 | AT4G21190.1 |

| | | |
|---|---|---|
| 120 | HORVU0Hr1G011850 | AT4G10330.1 |
| 121 | HORVU1Hr1G079410 | AT1G55340.1 |
| 122 | HORVU5Hr1G004630 | ATCG00020.1 |
| 123 | HORVU2Hr1G076180 | AT5G63860.1 |
| 124 | HORVU4Hr1G066880 | AT3G56630.1 |
| 125 | HORVU2Hr1G101390 | AT5G24780.1 |
| 126 | HORVU6Hr1G041720 | AT3G48110.1 |
| 127 | HORVU3Hr1G031400 | AT4G29100.2 |
| 128 | HORVU6Hr1G004230 | AT1G02120.2 |
| 129 | HORVU7Hr1G113460 | AT1G80920.1 |
| 130 | HORVU2Hr1G095890 | AT1G20693.2 |
| 131 | HORVU5Hr1G007980 | AT4G04850.2 |
| 132 | HORVU2Hr1G078410 | AT1G78140.1 |
| 133 | HORVU7Hr1G076030 | AT1G64680.1 |
| 134 | HORVU7Hr1G002350 | AT3G14470.2 |
| 135 | HORVU7Hr1G025730 | AT4G28830.4 |
| 136 | HORVU4Hr1G076930 | AT5G08390.1 |
| 137 | HORVU2Hr1G084910 | AT1G05710.5 |
| 138 | HORVU7Hr1G019830 | AT1G08510.1 |
| 139 | HORVU7Hr1G041570 | AT5G37710.1 |
| 140 | HORVU5Hr1G016730 | AT2G23450.2 |
| 141 | HORVU6Hr1G085730 | AT4G31020.2 |
| 142 | HORVU4Hr1G003800 | AT4G34610.4 |
| 143 | HORVU4Hr1G057450 | AT5G13510.1 |
| 144 | HORVU6Hr1G060100 | AT1G15130.1 |
| 145 | HORVU2Hr1G126540 | AT1G08980.1 |
| 146 | HORVU4Hr1G057050 | AT1G10900.3 |
| 147 | HORVU1Hr1G053670 | AT4G12470.1 |
| 148 | HORVU5Hr1G019650 | AT5G07830.1 |
| 149 | HORVU5Hr1G045440 | AT2G47170.3 |
| 150 | HORVU6Hr1G063650 | AT4G00150.1 |
| 151 | HORVU0Hr1G038350 | AT2G43320.2 |
| 152 | HORVU3Hr1G087500 | AT5G03040.3 |
| 153 | HORVU4Hr1G058580 | AT3G55320.1 |
| 154 | HORVU4Hr1G073930 | AT3G54660.1 |
| 155 | HORVU4Hr1G025330 | AT5G22620.6 |
| 156 | HORVU6Hr1G083930 | AT3G19240.1 |
| 157 | HORVU7Hr1G039810 | AT1G24130.1 |
| 158 | HORVU0Hr1G038050 | AT5G04950.1 |
| 159 | HORVU2Hr1G104830 | AT5G40510.1 |
| 160 | HORVU4Hr1G072220 | AT2G39140.1 |

| | | |
|---|---|---|
| 161 | HORVU1Hr1G068020 | AT3G24503.1 |
| 162 | HORVU1Hr1G064200 | AT1G09660.3 |
| 163 | HORVU6Hr1G050440 | AT1G66330.2 |
| 164 | HORVU1Hr1G086070 | AT2G22670.5 |
| 165 | HORVU5Hr1G052030 | AT5G47710.4 |
| 166 | HORVU7Hr1G056310 | AT1G33750.1 |
| 167 | HORVU3Hr1G114290 | AT1G13000.5 |
| 168 | HORVU4Hr1G005670 | AT5G12840.5 |
| 169 | HORVU7Hr1G094870 | AT5G10770.1 |
| 170 | HORVU3Hr1G107290 | AT4G26860.1 |
| 171 | HORVU4Hr1G079440 | AT5G50380.1 |
| 172 | HORVU4Hr1G016190 | AT1G66580.1 |
| 173 | HORVU0Hr1G016150 | AT4G09040.2 |
| 174 | HORVU0Hr1G008940 | AT5G16820.2 |
| 175 | HORVU3Hr1G075820 | AT5G43630.4 |
| 176 | HORVU3Hr1G016240 | AT1G60800.2 |
| 177 | HORVU3Hr1G029210 | AT2G38440.1 |
| 178 | HORVU5Hr1G018690 | AT2G03200.1 |
| 179 | HORVU3Hr1G098170 | AT5G48970.1 |
| 180 | HORVU2Hr1G101440 | AT1G51110.1 |
| 181 | HORVU4Hr1G048300 | AT1G67160.1 |
| 182 | HORVU3Hr1G094240 | AT3G27180.1 |
| 183 | HORVU3Hr1G084750 | AT2G38120.1 |
| 184 | HORVU3Hr1G054220 | AT1G19270.2 |
| 185 | HORVU5Hr1G113990 | AT1G59720.1 |
| 186 | HORVU3Hr1G016700 | AT4G26965.1 |
| 187 | HORVU5Hr1G088200 | AT4G17740.2 |
| 188 | HORVU6Hr1G063250 | AT3G27190.1 |
| 189 | HORVU5Hr1G106120 | AT4G02640.4 |
| 190 | HORVU3Hr1G092020 | AT3G48360.1 |
| 191 | HORVU5Hr1G060370 | AT2G01770.1 |
| 192 | HORVU6Hr1G058580 | AT3G23250.1 |
| 193 | HORVU5Hr1G071250 | AT5G47750.1 |
| 194 | HORVU3Hr1G030250 | AT3G48200.1 |
| 195 | HORVU0Hr1G035820 | AT5G56760.1 |
| 196 | HORVU3Hr1G059340 | AT1G22380.1 |
| 197 | HORVU0Hr1G019710 | AT1G47480.1 |
| 198 | HORVU5Hr1G073310 | AT4G12070.1 |
| 199 | HORVU1Hr1G016360 | AT5G46110.1 |
| 200 | HORVU3Hr1G027610 | AT1G53280.1 |
| 201 | HORVU6Hr1G092970 | AT4G00430.1 |

| | | |
|---|---|---|
| 202 | HORVU2Hr1G114580 | AT4G28080.1 |
| 203 | HORVU6Hr1G011060 | AT2G29100.1 |
| 204 | HORVU5Hr1G008980 | AT1G21230.1 |
| 205 | HORVU6Hr1G017860 | AT5G14760.1 |
| 206 | HORVU1Hr1G090990 | AT3G25690.6 |
| 207 | HORVU1Hr1G034570 | AT1G45110.1 |
| 208 | HORVU0Hr1G014950 | AT1G61590.1 |
| 209 | HORVU3Hr1G034210 | AT5G48960.1 |
| 210 | HORVU6Hr1G025880 | AT2G04360.1 |
| 211 | HORVU5Hr1G124210 | AT4G22890.3 |
| 212 | HORVU6Hr1G014480 | AT5G35170.1 |
| 213 | HORVU5Hr1G096370 | AT5G15490.1 |
| 214 | HORVU5Hr1G098490 | AT4G07960.2 |
| 215 | HORVU1Hr1G086110 | AT3G07310.1 |
| 216 | HORVU2Hr1G018730 | AT1G21350.5 |
| 217 | HORVU3Hr1G049970 | AT4G24330.1 |
| 218 | HORVU7Hr1G054530 | AT1G27950.1 |
| 219 | HORVU6Hr1G026680 | AT1G05170.2 |
| 220 | HORVU4Hr1G020850 | AT1G51630.1 |
| 221 | HORVU4Hr1G083080 | AT1G01140.1 |
| 222 | HORVU3Hr1G073040 | AT1G62430.1 |
| 223 | HORVU7Hr1G036130 | AT2G22540.2 |
| 224 | HORVU3Hr1G060730 | AT2G42975.1 |
| 225 | HORVU2Hr1G127440 | AT3G52820.1 |
| 226 | HORVU2Hr1G086040 | AT1G77090.1 |
| 227 | HORVU5Hr1G008080 | AT1G04150.1 |
| 228 | HORVU4Hr1G057890 | AT5G02790.1 |
| 229 | HORVU4Hr1G048960 | AT5G44180.3 |
| 230 | HORVU7Hr1G021630 | AT3G16370.1 |
| 231 | HORVU1Hr1G072750 | AT3G53900.2 |
| 232 | HORVU4Hr1G029350 | AT5G49470.3 |
| 233 | HORVU2Hr1G123270 | AT5G67610.1 |
| 234 | HORVU0Hr1G019500 | AT1G28680.1 |
| 235 | HORVU5Hr1G110960 | AT4G20330.1 |
| 236 | HORVU6Hr1G052230 | AT5G32470.1 |
| 237 | HORVU3Hr1G054770 | AT5G53170.1 |
| 238 | HORVU1Hr1G064290 | AT5G38520.1 |
| 239 | HORVU4Hr1G078780 | AT4G28760.2 |
| 240 | HORVU2Hr1G116130 | AT5G65210.7 |
| 241 | HORVU1Hr1G006860 | AT5G42740.2 |
| 242 | HORVU6Hr1G045140 | AT1G72550.2 |

| | | |
|---|---|---|
| 243 | HORVU3Hr1G028810 | AT1G17100.1 |
| 244 | HORVU5Hr1G016970 | AT3G50380.4 |
| 245 | HORVU3Hr1G041820 | AT1G48210.7 |
| 246 | HORVU7Hr1G051760 | AT2G24395.1 |
| 247 | HORVU5Hr1G112710 | AT1G78230.1 |
| 248 | HORVU4Hr1G021950 | AT5G03940.1 |
| 249 | HORVU2Hr1G059380 | AT3G52380.1 |
| 250 | HORVU2Hr1G099510 | AT1G79460.1 |
| 251 | HORVU7Hr1G012530 | AT4G34710.2 |
| 252 | HORVU2Hr1G093580 | AT2G44830.3 |
| 253 | HORVU4Hr1G011250 | AT5G05250.1 |
| 254 | HORVU1Hr1G047820 | AT5G53580.1 |
| 255 | HORVU1Hr1G083160 | AT3G18280.2 |
| 256 | HORVU0Hr1G016560 | AT5G62140.1 |
| 257 | HORVU2Hr1G023540 | AT5G13630.1 |
| 258 | HORVU2Hr1G058230 | AT5G20350.1 |
| 259 | HORVU6Hr1G093570 | AT1G20270.1 |
| 260 | HORVU7Hr1G049200 | AT1G55590.1 |
| 261 | HORVU2Hr1G035550 | AT2G23390.2 |
| 262 | HORVU1Hr1G018080 | AT1G59760.1 |
| 263 | HORVU0Hr1G016250 | AT3G08010.1 |
| 264 | HORVU7Hr1G019510 | AT1G56220.1 |
| 265 | HORVU4Hr1G016470 | AT4G31390.1 |
| 266 | HORVU1Hr1G092860 | AT4G01100.1 |
| 267 | HORVU4Hr1G072150 | AT4G05160.1 |
| 268 | HORVU4Hr1G050630 | AT5G20920.3 |
| 269 | HORVU3Hr1G078830 | AT3G17930.1 |
| 270 | HORVU7Hr1G014840 | AT1G32928.1 |
| 271 | HORVU4Hr1G023230 | #N/A |
| 272 | HORVU6Hr1G008590 | AT1G12800.1 |
| 273 | HORVU5Hr1G073380 | AT1G73730.1 |
| 274 | HORVU3Hr1G031460 | AT4G29080.1 |
| 275 | HORVU6Hr1G014100 | AT5G51545.1 |
| 276 | HORVU7Hr1G052400 | AT4G29950.1 |
| 277 | HORVU3Hr1G043870 | AT5G42740.3 |
| 278 | HORVU3Hr1G088910 | AT1G67690.1 |
| 279 | HORVU5Hr1G076100 | AT1G69040.2 |
| 280 | HORVU3Hr1G057020 | AT1G60230.1 |
| 281 | HORVU2Hr1G010520 | AT4G35880.1 |
| 282 | HORVU5Hr1G072120 | AT2G33180.1 |
| 283 | HORVU6Hr1G032540 | AT1G61590.1 |

| | | |
|---|---|---|
| 284 | HORVU6Hr1G046980 | AT1G04880.1 |
| 285 | HORVU3Hr1G065630 | AT5G55090.1 |
| 286 | HORVU6Hr1G060810 | AT1G12250.2 |
| 287 | HORVU2Hr1G104920 | AT1G42540.4 |
| 288 | HORVU4Hr1G008210 | AT2G01830.6 |
| 289 | HORVU7Hr1G060530 | AT1G17220.1 |
| 290 | HORVU6Hr1G095000 | AT3G14470.2 |
| 291 | HORVU3Hr1G043300 | AT3G43790.4 |
| 292 | HORVU3Hr1G078200 | AT5G44650.1 |
| 293 | HORVU1Hr1G020000 | AT2G47940.2 |
| 294 | HORVU3Hr1G036380 | AT5G17260.1 |
| 295 | HORVU7Hr1G027740 | AT1G05990.1 |
| 296 | HORVU7Hr1G108650 | #N/A |
| 297 | HORVU4Hr1G007380 | AT1G63660.2 |
| 298 | HORVU7Hr1G097660 | AT5G06570.2 |
| 299 | HORVU2Hr1G034400 | AT1G69410.1 |
| 300 | HORVU3Hr1G005770 | AT2G01320.7 |
| 301 | HORVU4Hr1G029400 | AT5G10490.2 |
| 302 | HORVU7Hr1G021180 | AT2G02740.1 |
| 303 | HORVU6Hr1G005020 | AT3G45050.4 |
| 304 | HORVU6Hr1G009480 | AT1G73060.1 |
| 305 | HORVU7Hr1G100310 | AT3G50210.2 |
| 306 | HORVU4Hr1G023580 | AT2G21530.1 |
| 307 | HORVU3Hr1G064340 | AT1G66840.2 |
| 308 | HORVU2Hr1G008380 | AT5G13410.1 |
| 309 | HORVU0Hr1G008530 | AT5G41190.1 |
| 310 | HORVU6Hr1G069390 | AT5G10330.2 |
| 311 | HORVU5Hr1G119790 | AT1G57770.1 |
| 312 | HORVU7Hr1G020590 | AT5G23310.1 |
| 313 | HORVU4Hr1G067280 | AT3G25900.1 |
| 314 | HORVU1Hr1G020240 | AT4G19510.8 |
| 315 | HORVU7Hr1G012940 | AT1G01940.1 |
| 316 | HORVU2Hr1G018340 | AT2G30360.1 |
| 317 | HORVU2Hr1G087010 | AT4G32060.1 |
| 318 | HORVU4Hr1G014590 | AT5G67550.1 |
| 319 | HORVU5Hr1G043800 | #N/A |
| 320 | HORVU2Hr1G106100 | AT4G17790.1 |
| 321 | HORVU1Hr1G065250 | AT4G22200.1 |
| 322 | HORVU4Hr1G026670 | AT1G15660.1 |
| 323 | HORVU3Hr1G020830 | AT1G02170.1 |
| 324 | HORVU0Hr1G004060 | AT3G18690.1 |

| | | |
|---|---|---|
| 325 | HORVU5Hr1G042310 | AT5G04740.1 |
| 326 | HORVU4Hr1G014910 | AT2G44090.5 |
| 327 | HORVU3Hr1G057240 | AT1G64500.1 |
| 328 | HORVU5Hr1G017400 | AT3G05850.1 |
| 329 | HORVU1Hr1G088870 | AT2G34420.1 |
| 330 | HORVU4Hr1G015260 | AT4G10340.1 |
| 331 | HORVU7Hr1G099350 | AT3G06010.1 |
| 332 | HORVU3Hr1G062700 | AT1G02475.1 |
| 333 | HORVU3Hr1G069730 | AT3G16520.3 |
| 334 | HORVU6Hr1G075540 | AT1G52990.2 |
| 335 | HORVU4Hr1G040950 | AT1G05190.1 |
| 336 | HORVU7Hr1G021430 | AT1G31490.1 |
| 337 | HORVU5Hr1G125110 | AT2G36990.1 |
| 338 | HORVU1Hr1G092670 | AT2G46140.1 |
| 339 | HORVU5Hr1G070950 | AT4G22970.3 |
| 340 | HORVU7Hr1G021950 | AT5G57550.1 |
| 341 | HORVU7Hr1G106540 | AT5G11720.1 |
| 342 | HORVU3Hr1G066240 | AT4G35440.1 |
| 343 | HORVU5Hr1G013400 | AT5G50760.1 |
| 344 | HORVU7Hr1G029750 | AT3G46920.1 |
| 345 | HORVU3Hr1G065720 | AT1G28240.1 |
| 346 | HORVU0Hr1G005330 | AT3G61460.1 |
| 347 | HORVU7Hr1G092940 | AT5G14880.1 |
| 348 | HORVU2Hr1G086950 | AT4G32060.2 |
| 349 | HORVU1Hr1G094480 | AT1G49970.1 |
| 350 | HORVU7Hr1G042080 | AT1G25440.1 |
| 351 | HORVU3Hr1G108540 | AT3G57040.2 |
| 352 | HORVU7Hr1G099560 | AT1G18210.2 |
| 353 | HORVU7Hr1G024560 | AT5G15180.1 |
| 354 | HORVU1Hr1G079270 | AT3G13062.1 |
| 355 | HORVU3Hr1G065640 | AT4G26890.1 |
| 356 | HORVU4Hr1G070720 | AT1G75100.1 |
| 357 | HORVU2Hr1G023170 | AT2G41680.1 |
| 358 | HORVU3Hr1G090760 | AT4G10030.1 |
| 359 | HORVU2Hr1G084820 | AT3G06110.3 |
| 360 | HORVU0Hr1G018970 | AT1G49010.1 |
| 361 | HORVU5Hr1G050610 | AT5G67130.1 |
| 362 | HORVU7Hr1G056320 | AT3G22780.1 |
| 363 | HORVU5Hr1G033660 | AT1G14610.1 |
| 364 | HORVU7Hr1G030810 | AT2G25080.1 |
| 365 | HORVU2Hr1G043170 | AT5G42310.1 |

| | | |
|---|---|---|
| 366 | HORVU7Hr1G101270 | AT4G30080.1 |
| 367 | HORVU4Hr1G015820 | AT5G63510.1 |
| 368 | HORVU2Hr1G105900 | AT5G16660.1 |
| 369 | HORVU6Hr1G040040 | AT5G11970.1 |
| 370 | HORVU6Hr1G063510 | AT5G21222.4 |
| 371 | HORVU4Hr1G057910 | AT5G02790.1 |
| 372 | HORVU5Hr1G072920 | AT3G12630.1 |
| 373 | HORVU3Hr1G085380 | AT3G46970.1 |
| 374 | HORVU3Hr1G003050 | AT5G17330.1 |
| 375 | HORVU5Hr1G121250 | AT5G13280.1 |
| 376 | HORVU2Hr1G104790 | #N/A |
| 377 | HORVU1Hr1G070740 | AT3G56040.1 |
| 378 | HORVU1Hr1G000040 | AT2G33450.1 |
| 379 | HORVU3Hr1G097200 | AT5G62000.5 |
| 380 | HORVU4Hr1G084890 | AT1G01040.2 |
| 381 | HORVU7Hr1G025590 | AT5G15180.1 |
| 382 | HORVU2Hr1G041320 | AT4G05090.1 |
| 383 | HORVU3Hr1G005830 | AT2G01320.3 |
| 384 | HORVU3Hr1G089290 | AT4G02380.3 |
| 385 | HORVU3Hr1G065960 | AT5G47770.1 |
| 386 | HORVU2Hr1G023530 | AT1G73875.1 |
| 387 | HORVU4Hr1G076690 | AT4G14819.1 |
| 388 | HORVU2Hr1G121310 | AT2G19130.1 |
| 389 | HORVU2Hr1G108350 | AT2G35460.1 |
| 390 | HORVU3Hr1G030580 | AT2G24490.2 |
| 391 | HORVU7Hr1G027270 | AT5G65380.1 |
| 392 | HORVU0Hr1G022240 | AT5G63060.1 |
| 393 | HORVU5Hr1G038840 | AT1G22240.1 |
| 394 | HORVU6Hr1G042170 | AT1G61900.2 |
| 395 | HORVU5Hr1G040640 | AT4G02070.2 |
| 396 | HORVU2Hr1G035990 | AT1G15640.1 |
| 397 | HORVU2Hr1G066470 | AT4G28760.2 |
| 398 | HORVU2Hr1G034540 | AT3G15110.1 |
| 399 | HORVU7Hr1G012690 | AT4G25080.6 |
| 400 | HORVU3Hr1G116060 | AT5G21280.2 |
| 401 | HORVU3Hr1G086210 | AT1G07530.1 |
| 402 | HORVU0Hr1G040200 | AT1G04620.1 |
| 403 | HORVU0Hr1G003890 | AT4G21990.1 |
| 404 | HORVU1Hr1G030540 | AT4G29100.2 |
| 405 | HORVU2Hr1G094160 | AT4G25700.1 |
| 406 | HORVU3Hr1G086310 | AT2G01480.1 |

| | | |
|---|---|---|
| 407 | HORVU3Hr1G053910 | AT2G34090.1 |
| 408 | HORVU4Hr1G041000 | AT1G72990.2 |
| 409 | HORVU4Hr1G002020 | AT1G58030.1 |
| 410 | HORVU2Hr1G118410 | AT1G18490.1 |
| 411 | HORVU4Hr1G010120 | AT5G65840.1 |
| 412 | HORVU6Hr1G017890 | AT1G70420.1 |
| 413 | HORVU5Hr1G064130 | AT4G30825.1 |
| 414 | HORVU1Hr1G045280 | AT2G21385.1 |
| 415 | HORVU2Hr1G107990 | AT1G11530.1 |
| 416 | HORVU1Hr1G052780 | AT3G10405.1 |
| 417 | HORVU2Hr1G076100 | AT1G77290.3 |
| 418 | HORVU5Hr1G022760 | AT5G26820.1 |
| 419 | HORVU2Hr1G038740 | AT3G54820.1 |
| 420 | HORVU5Hr1G064500 | AT1G12190.1 |
| 421 | HORVU1Hr1G063090 | AT1G09010.1 |
| 422 | HORVU2Hr1G111440 | AT1G73110.1 |
| 423 | HORVU1Hr1G090740 | AT2G25605.1 |
| 424 | HORVU3Hr1G110160 | AT2G45770.1 |
| 425 | HORVU1Hr1G056510 | AT3G63490.1 |
| 426 | HORVU7Hr1G113210 | AT5G11450.2 |
| 427 | HORVU2Hr1G095400 | AT5G52190.1 |
| 428 | HORVU4Hr1G013380 | AT4G37540.1 |
| 429 | HORVU4Hr1G090840 | AT3G06980.1 |
| 430 | HORVU6Hr1G028250 | AT3G48280.1 |
| 431 | HORVU7Hr1G024210 | AT1G73170.1 |
| 432 | HORVU1Hr1G084520 | AT4G32590.1 |
| 433 | HORVU2Hr1G011960 | AT2G34860.2 |
| 434 | HORVU2Hr1G003840 | AT4G34880.2 |
| 435 | HORVU2Hr1G072020 | AT5G51070.1 |
| 436 | HORVU1Hr1G072520 | AT3G32040.1 |
| 437 | HORVU5Hr1G049280 | AT1G50730.1 |
| 438 | HORVU3Hr1G116470 | AT3G09410.1 |
| 439 | HORVU1Hr1G081370 | AT3G14830.4 |
| 440 | HORVU7Hr1G116780 | AT3G59080.1 |
| 441 | HORVU7Hr1G035320 | AT1G05000.1 |
| 442 | HORVU6Hr1G025110 | AT5G08650.1 |
| 443 | HORVU6Hr1G076860 | AT1G70070.1 |
| 444 | HORVU2Hr1G110900 | AT3G13530.1 |
| 445 | HORVU1Hr1G022800 | AT2G25920.1 |
| 446 | HORVU2Hr1G104580 | AT5G37260.1 |
| 447 | HORVU6Hr1G043920 | AT1G06950.1 |

| | | |
|---|---|---|
| 448 | HORVU2Hr1G036280 | AT5G48150.4 |
| 449 | HORVU1Hr1G095410 | AT5G53420.1 |
| 450 | HORVU6Hr1G014140 | AT4G14500.2 |
| 451 | HORVU3Hr1G085420 | AT3G46970.1 |
| 452 | HORVU2Hr1G032170 | AT1G09760.1 |
| 453 | HORVU0Hr1G013890 | AT3G14420.2 |
| 454 | HORVU3Hr1G013600 | AT5G15802.1 |
| 455 | HORVU7Hr1G117020 | AT4G35040.4 |
| 456 | HORVU4Hr1G011090 | AT4G09650.1 |
| 457 | HORVU7Hr1G091440 | AT4G15630.1 |
| 458 | HORVU5Hr1G043550 | AT1G50450.1 |
| 459 | HORVU4Hr1G027180 | AT2G39890.2 |
| 460 | HORVU6Hr1G025120 | AT5G08650.1 |
| 461 | HORVU6Hr1G090510 | AT4G35160.1 |
| 462 | HORVU5Hr1G020410 | AT5G61040.1 |
| 463 | HORVU6Hr1G011760 | AT3G04790.1 |
| 464 | HORVU3Hr1G116490 | AT3G09410.1 |
| 465 | HORVU7Hr1G082590 | AT3G48420.1 |
| 466 | HORVU3Hr1G086240 | AT3G55470.2 |
| 467 | HORVU6Hr1G074820 | AT3G48430.2 |
| 468 | HORVU1Hr1G051150 | AT1G32080.1 |
| 469 | HORVU3Hr1G080770 | AT5G05190.1 |
| 470 | HORVU1Hr1G067110 | AT5G44210.1 |
| 471 | HORVU7Hr1G074890 | AT5G40650.1 |
| 472 | HORVU7Hr1G038120 | AT3G26900.3 |
| 473 | HORVU5Hr1G013840 | AT4G33670.1 |
| 474 | HORVU6Hr1G081670 | AT5G36160.1 |
| 475 | HORVU3Hr1G117810 | AT3G55980.1 |
| 476 | HORVU5Hr1G036160 | AT3G52950.2 |
| 477 | HORVU3Hr1G093260 | AT3G07350.1 |
| 478 | HORVU2Hr1G029430 | AT4G11070.2 |
| 479 | HORVU1Hr1G029850 | AT5G37770.2 |
| 480 | HORVU5Hr1G016350 | AT4G36860.3 |
| 481 | HORVU2Hr1G032400 | AT1G13820.1 |
| 482 | HORVU0Hr1G005340 | AT5G52190.1 |
| 483 | HORVU4Hr1G079710 | AT1G02260.2 |
| 484 | HORVU3Hr1G002820 | AT4G05200.1 |
| 485 | HORVU1Hr1G010450 | AT1G31730.1 |
| 486 | HORVU4Hr1G053530 | AT3G17040.1 |
| 487 | HORVU2Hr1G032970 | AT1G13700.3 |
| 488 | HORVU5Hr1G020110 | AT3G29180.3 |

| | | |
|---|---|---|
| 489 | HORVU6Hr1G050160 | AT5G13650.2 |
| 490 | HORVU0Hr1G005570 | AT1G73110.2 |
| 491 | HORVU7Hr1G071440 | AT3G01160.1 |
| 492 | HORVU3Hr1G014890 | AT3G07040.1 |
| 493 | HORVU6Hr1G075850 | AT5G37820.1 |
| 494 | HORVU4Hr1G011600 | AT2G30630.1 |
| 495 | HORVU5Hr1G091810 | AT4G24770.2 |
| 496 | HORVU5Hr1G062780 | AT2G03140.6 |
| 497 | HORVU7Hr1G023250 | AT3G14830.4 |
| 498 | HORVU6Hr1G027910 | AT1G17820.1 |
| 499 | HORVU0Hr1G007950 | #N/A |
| 500 | HORVU4Hr1G069050 | AT2G03420.1 |
| 501 | HORVU2Hr1G073740 | AT2G27290.1 |
| 502 | HORVU3Hr1G065700 | AT3G28510.1 |
| 503 | HORVU5Hr1G080690 | AT1G63120.1 |
| 504 | HORVU5Hr1G028670 | AT3G03090.1 |
| 505 | HORVU7Hr1G096430 | AT3G13040.3 |
| 506 | HORVU1Hr1G055710 | AT4G17490.1 |
| 507 | HORVU4Hr1G009080 | AT1G16800.2 |
| 508 | HORVU6Hr1G020730 | AT5G54880.1 |
| 509 | HORVU7Hr1G080520 | AT3G45750.1 |
| 510 | HORVU2Hr1G003460 | AT1G68660.1 |
| 511 | HORVU2Hr1G046740 | AT2G32415.1 |
| 512 | HORVU7Hr1G069930 | AT3G59780.1 |
| 513 | HORVU6Hr1G084390 | #N/A |
| 514 | HORVU3Hr1G078820 | AT3G56650.1 |
| 515 | HORVU3Hr1G061690 | AT3G05700.1 |
| 516 | HORVU1Hr1G027830 | AT5G12010.1 |
| 517 | HORVU2Hr1G043920 | AT4G15210.1 |
| 518 | HORVU4Hr1G011120 | AT3G28460.1 |
| 519 | HORVU5Hr1G066350 | AT1G14030.1 |
| 520 | HORVU3Hr1G037600 | AT5G47910.1 |
| 521 | HORVU2Hr1G088180 | AT4G13550.2 |
| 522 | HORVU3Hr1G084590 | AT5G58770.1 |
| 523 | HORVU2Hr1G028160 | AT5G39530.1 |
| 524 | HORVU3Hr1G114960 | AT2G29525.4 |
| 525 | HORVU3Hr1G085970 | AT3G23160.1 |
| 526 | HORVU7Hr1G034290 | AT2G05940.1 |
| 527 | HORVU7Hr1G029720 | AT2G15580.1 |
| 528 | HORVU3Hr1G096830 | AT4G12910.1 |
| 529 | HORVU5Hr1G073950 | AT2G33590.1 |

| | | |
|---|---|---|
| 530 | HORVU4Hr1G086250 | AT3G26900.3 |
| 531 | HORVU3Hr1G116000 | AT3G27040.1 |
| 532 | HORVU6Hr1G081330 | AT2G05940.1 |
| 533 | HORVU0Hr1G017440 | AT3G15900.1 |
| 534 | HORVU7Hr1G074600 | AT5G47920.1 |
| 535 | HORVU5Hr1G103320 | AT2G34690.1 |
| 536 | HORVU4Hr1G002700 | AT3G22550.1 |
| 537 | HORVU1Hr1G051360 | AT2G32180.2 |
| 538 | HORVU2Hr1G095070 | AT4G00750.1 |
| 539 | HORVU2Hr1G094090 | #N/A |
| 540 | HORVU7Hr1G113560 | AT5G10530.1 |
| 541 | HORVU1Hr1G069430 | AT1G32070.1 |
| 542 | HORVU1Hr1G078140 | AT2G30570.1 |
| 543 | HORVU1Hr1G060740 | AT1G10740.4 |
| 544 | HORVU2Hr1G041080 | AT1G11260.1 |
| 545 | HORVU3Hr1G035230 | AT3G02400.1 |
| 546 | HORVU3Hr1G072260 | AT1G28520.5 |
| 547 | HORVU6Hr1G088200 | AT2G39190.2 |
| 548 | HORVU4Hr1G020080 | AT1G18740.1 |
| 549 | HORVU5Hr1G071680 | AT5G47780.1 |
| 550 | HORVU7Hr1G067990 | AT5G38660.1 |
| 551 | HORVU3Hr1G070380 | AT5G56550.1 |
| 552 | HORVU6Hr1G012320 | AT4G31290.1 |
| 553 | HORVU4Hr1G078440 | AT3G25590.1 |
| 554 | HORVU0Hr1G005920 | AT5G27270.1 |
| 555 | HORVU1Hr1G087600 | AT1G28480.1 |
| 556 | HORVU2Hr1G041620 | AT4G36440.1 |
| 557 | HORVU5Hr1G037920 | AT4G33470.1 |
| 558 | HORVU3Hr1G086970 | AT2G39450.1 |
| 559 | HORVU7Hr1G010790 | AT2G30970.2 |
| 560 | HORVU6Hr1G072740 | AT5G23060.1 |
| 561 | HORVU5Hr1G011100 | AT4G01690.1 |
| 562 | HORVU4Hr1G048440 | AT2G37220.1 |
| 563 | HORVU1Hr1G093430 | AT1G53250.1 |
| 564 | HORVU7Hr1G075700 | AT4G11220.1 |
| 565 | HORVU4Hr1G016210 | AT1G32520.1 |
| 566 | HORVU3Hr1G098160 | AT1G09070.1 |
| 567 | HORVU7Hr1G100790 | AT3G57540.1 |
| 568 | HORVU5Hr1G095990 | AT5G39040.1 |
| 569 | HORVU5Hr1G082630 | AT2G21170.3 |
| 570 | HORVU3Hr1G094600 | AT1G67340.1 |

| | | |
|---|---|---|
| 571 | HORVU6Hr1G000640 | AT5G42190.1 |
| 572 | HORVU0Hr1G015530 | AT1G16670.1 |
| 573 | HORVU7Hr1G043360 | AT5G52630.1 |
| 574 | HORVU3Hr1G014850 | AT3G23740.1 |
| 575 | HORVU7Hr1G000630 | AT3G46530.1 |
| 576 | HORVU3Hr1G112930 | AT3G07040.1 |
| 577 | HORVU3Hr1G013770 | AT3G12080.1 |
| 578 | HORVU1Hr1G010440 | AT4G23470.1 |
| 579 | HORVU4Hr1G042550 | AT2G29640.1 |
| 580 | HORVU2Hr1G059910 | AT2G07240.1 |
| 581 | HORVU3Hr1G089550 | AT2G39780.1 |
| 582 | HORVU6Hr1G071930 | AT4G15950.1 |
| 583 | HORVU5Hr1G039960 | AT1G72030.1 |
| 584 | HORVU1Hr1G082540 | AT4G33500.1 |
| 585 | HORVU4Hr1G052470 | AT5G30510.1 |
| 586 | HORVU2Hr1G096360 | AT4G00430.1 |
| 587 | HORVU0Hr1G008850 | AT1G12420.2 |
| 588 | HORVU4Hr1G024290 | AT4G02640.4 |
| 589 | HORVU3Hr1G107070 | AT3G09150.3 |
| 590 | HORVU3Hr1G065770 | AT3G14770.1 |
| 591 | HORVU0Hr1G022820 | AT2G17040.1 |
| 592 | HORVU6Hr1G061870 | AT5G07450.1 |
| 593 | HORVU4Hr1G054910 | AT3G62720.2 |
| 594 | HORVU5Hr1G011960 | AT4G20760.1 |
| 595 | HORVU5Hr1G038630 | AT1G67090.1 |
| 596 | HORVU2Hr1G098650 | AT1G31280.1 |
| 597 | HORVU6Hr1G009510 | AT2G47910.1 |
| 598 | HORVU7Hr1G090800 | AT4G25450.3 |
| 599 | HORVU5Hr1G059570 | AT1G62780.1 |
| 600 | HORVU2Hr1G080260 | AT3G01440.1 |
| 601 | HORVU2Hr1G079970 | AT5G57200.1 |
| 602 | HORVU2Hr1G115690 | AT1G71810.1 |
| 603 | HORVU3Hr1G014860 | AT5G63500.1 |
| 604 | HORVU5Hr1G028460 | AT5G15430.2 |
| 605 | HORVU6Hr1G063820 | AT1G11420.1 |
| 606 | HORVU6Hr1G062040 | AT4G17560.1 |
| 607 | HORVU4Hr1G082040 | AT4G35090.1 |
| 608 | HORVU2Hr1G007350 | AT5G24860.1 |
| 609 | HORVU0Hr1G000960 | AT3G04890.1 |
| 610 | HORVU4Hr1G060130 | AT5G59550.1 |
| 611 | HORVU0Hr1G008540 | AT5G26850.4 |

| | | |
|---|---|---|
| 612 | HORVU3Hr1G014780 | AT3G26900.3 |
| 613 | HORVU2Hr1G015720 | AT1G56600.1 |
| 614 | HORVU5Hr1G069380 | AT1G22990.1 |
| 615 | HORVU1Hr1G018010 | AT3G62070.1 |
| 616 | HORVU7Hr1G091570 | AT1G19940.2 |
| 617 | HORVU4Hr1G060630 | AT3G54050.2 |
| 618 | HORVU2Hr1G017420 | AT2G39050.1 |
| 619 | HORVU0Hr1G023150 | AT1G31720.6 |
| 620 | HORVU5Hr1G111310 | AT1G08520.1 |
| 621 | HORVU0Hr1G021950 | AT2G26600.1 |
| 622 | HORVU1Hr1G061020 | AT4G15545.1 |
| 623 | HORVU6Hr1G085120 | AT3G59660.1 |
| 624 | HORVU2Hr1G037620 | AT4G15810.1 |
| 625 | HORVU1Hr1G077160 | AT3G01060.3 |
| 626 | HORVU5Hr1G087920 | AT1G42550.1 |
| 627 | HORVU3Hr1G114220 | AT5G19210.2 |
| 628 | HORVU2Hr1G075180 | AT5G61350.1 |
| 629 | HORVU4Hr1G086110 | AT3G61060.1 |
| 630 | HORVU6Hr1G054430 | AT1G13270.1 |
| 631 | HORVU1Hr1G038620 | AT3G04260.1 |
| 632 | HORVU6Hr1G053680 | AT4G20360.2 |
| 633 | HORVU2Hr1G113850 | AT4G14465.1 |
| 634 | HORVU5Hr1G122820 | AT4G35450.3 |
| 635 | HORVU5Hr1G056130 | AT4G11070.2 |
| 636 | HORVU4Hr1G003040 | AT5G45110.1 |
| 637 | HORVU5Hr1G073330 | AT3G20810.1 |
| 638 | HORVU6Hr1G005920 | AT3G27890.1 |
| 639 | HORVU3Hr1G063280 | AT1G80380.2 |
| 640 | HORVU2Hr1G120490 | AT3G57040.2 |
| 641 | HORVU5Hr1G024470 | AT4G29060.1 |
| 642 | HORVU0Hr1G031850 | AT5G52660.1 |
| 643 | HORVU1Hr1G041640 | AT1G37020.1 |
| 644 | HORVU5Hr1G049980 | AT2G18710.1 |
| 645 | HORVU2Hr1G037610 | #N/A |
| 646 | HORVU6Hr1G078290 | AT5G65220.1 |
| 647 | HORVU0Hr1G017140 | AT1G69523.1 |
| 648 | HORVU1Hr1G046550 | AT2G38130.2 |
| 649 | HORVU6Hr1G087720 | AT1G76080.1 |
| 650 | HORVU3Hr1G003410 | AT4G02610.1 |
| 651 | HORVU6Hr1G025730 | AT1G23740.1 |
| 652 | HORVU1Hr1G074940 | AT1G25280.1 |

| | | |
|---|---|---|
| 653 | HORVU5Hr1G037660 | AT5G09220.1 |
| 654 | HORVU2Hr1G068880 | AT5G38360.1 |
| 655 | HORVU7Hr1G034430 | AT2G26330.1 |
| 656 | HORVU7Hr1G109380 | AT5G45680.1 |
| 657 | HORVU2Hr1G093220 | AT1G02150.1 |
| 658 | HORVU1Hr1G068500 | AT3G51510.1 |
| 659 | HORVU3Hr1G068300 | AT3G24430.1 |
| 660 | HORVU5Hr1G124520 | AT1G17850.2 |
| 661 | HORVU7Hr1G067110 | AT5G17170.1 |
| 662 | HORVU6Hr1G067470 | AT2G01850.1 |
| 663 | HORVU2Hr1G121200 | AT5G64700.1 |
| 664 | HORVU7Hr1G102730 | AT5G25460.1 |
| 665 | HORVU2Hr1G091860 | AT5G66750.1 |
| 666 | HORVU1Hr1G053510 | AT4G00150.1 |
| 667 | HORVU5Hr1G068520 | AT1G66400.1 |
| 668 | HORVU5Hr1G081750 | AT4G25290.1 |
| 669 | HORVU1Hr1G022110 | AT1G69680.1 |
| 670 | HORVU0Hr1G001280 | AT3G43810.1 |
| 671 | HORVU6Hr1G014080 | AT1G64720.1 |
| 672 | HORVU2Hr1G115920 | AT3G09830.3 |
| 673 | HORVU5Hr1G118100 | AT3G28674.1 |
| 674 | HORVU6Hr1G068640 | AT5G46860.1 |
| 675 | HORVU1Hr1G091520 | AT1G75250.2 |
| 676 | HORVU1Hr1G017120 | AT2G47060.4 |
| 677 | HORVU3Hr1G018800 | AT4G34040.1 |
| 678 | HORVU5Hr1G094210 | AT5G43670.1 |
| 679 | HORVU3Hr1G020640 | AT4G01935.1 |
| 680 | HORVU5Hr1G039210 | AT5G21430.1 |
| 681 | HORVU4Hr1G060140 | AT5G59600.1 |
| 682 | HORVU1Hr1G005600 | AT2G39570.1 |
| 683 | HORVU3Hr1G083350 | AT1G43670.1 |
| 684 | HORVU7Hr1G023600 | AT2G43940.1 |
| 685 | HORVU2Hr1G112330 | AT3G52820.1 |
| 686 | HORVU7Hr1G051200 | ATCG00440.1 |
| 687 | HORVU1Hr1G022660 | AT4G23890.1 |
| 688 | HORVU1Hr1G061230 | AT2G48070.2 |
| 689 | HORVU2Hr1G089940 | AT2G37170.2 |
| 690 | HORVU6Hr1G013670 | AT4G32410.1 |
| 691 | HORVU6Hr1G020960 | AT4G01070.1 |
| 692 | HORVU1Hr1G049760 | AT5G41410.1 |
| 693 | HORVU1Hr1G090150 | AT3G48110.1 |

| | | |
|---|---|---|
| 694 | HORVU3Hr1G067760 | AT4G22530.2 |
| 695 | HORVU7Hr1G036450 | AT1G22460.1 |
| 696 | HORVU1Hr1G001890 | AT1G71440.1 |
| 697 | HORVU5Hr1G010870 | AT2G47870.1 |
| 698 | HORVU6Hr1G064620 | AT1G20440.1 |
| 699 | HORVU1Hr1G093180 | AT5G24120.3 |
| 700 | HORVU7Hr1G108930 | AT5G07610.1 |
| 701 | HORVU3Hr1G086380 | AT3G46540.3 |
| 702 | HORVU6Hr1G001690 | AT5G60580.3 |
| 703 | HORVU3Hr1G070360 | AT4G26288.1 |
| 704 | HORVU3Hr1G070690 | AT4G29130.1 |
| 705 | HORVU5Hr1G021340 | AT1G54500.1 |
| 706 | HORVU5Hr1G097150 | AT5G67360.1 |
| 707 | HORVU4Hr1G041010 | AT1G25290.2 |
| 708 | HORVU4Hr1G052990 | AT1G56500.3 |
| 709 | HORVU3Hr1G105260 | AT4G31560.1 |
| 710 | HORVU7Hr1G052500 | AT5G57240.5 |
| 711 | HORVU1Hr1G041990 | AT2G45530.1 |
| 712 | HORVU5Hr1G087830 | AT5G59970.2 |
| 713 | HORVU5Hr1G073870 | AT1G76470.1 |
| 714 | HORVU7Hr1G036090 | AT5G10470.1 |
| 715 | HORVU2Hr1G080510 | AT2G47800.1 |
| 716 | HORVU1Hr1G004650 | AT3G52820.1 |
| 717 | HORVU2Hr1G101720 | AT1G32580.1 |
| 718 | HORVU7Hr1G109580 | AT5G35400.1 |
| 719 | HORVU5Hr1G117780 | AT3G13840.1 |
| 720 | HORVU2Hr1G104740 | AT2G44100.2 |
| 721 | HORVU2Hr1G071850 | AT3G16560.4 |
| 722 | HORVU2Hr1G101340 | #N/A |
| 723 | HORVU7Hr1G076840 | AT4G10170.2 |
| 724 | HORVU2Hr1G069880 | AT3G48660.1 |
| 725 | HORVU4Hr1G000230 | AT3G55330.1 |
| 726 | HORVU3Hr1G062250 | AT1G51100.1 |
| 727 | HORVU5Hr1G118120 | AT1G40087.1 |
| 728 | HORVU3Hr1G070340 | AT2G35680.1 |
| 729 | HORVU4Hr1G037290 | AT2G04700.3 |
| 730 | HORVU2Hr1G066790 | AT1G22360.1 |
| 731 | HORVU1Hr1G014240 | AT1G70290.1 |
| 732 | HORVU6Hr1G009350 | AT4G14570.1 |
| 733 | HORVU7Hr1G040790 | AT5G37490.1 |
| 734 | HORVU5Hr1G084490 | AT4G35880.1 |

| | | |
|---|---|---|
| 735 | HORVU1Hr1G069180 | AT2G38640.1 |
| 736 | HORVU6Hr1G067660 | AT1G32060.1 |
| 737 | HORVU6Hr1G011670 | AT2G24960.1 |
| 738 | HORVU1Hr1G068960 | AT1G51340.1 |
| 739 | HORVU1Hr1G046360 | AT5G49110.4 |
| 740 | HORVU1Hr1G031930 | AT1G55640.1 |
| 741 | HORVU2Hr1G121110 | AT5G37020.1 |
| 742 | HORVU5Hr1G098890 | AT2G22780.1 |
| 743 | HORVU1Hr1G031940 | AT4G19650.2 |
| 744 | HORVU5Hr1G039850 | AT5G09810.1 |
| 745 | HORVU3Hr1G112010 | #N/A |
| 746 | HORVU0Hr1G000900 | AT5G17310.5 |
| 747 | HORVU6Hr1G068760 | AT1G01050.2 |
| 748 | HORVU5Hr1G033540 | AT1G67090.1 |
| 749 | HORVU4Hr1G063420 | AT1G01120.1 |
| 750 | HORVU4Hr1G006530 | AT3G18080.1 |
| 751 | HORVU5Hr1G009460 | AT5G24060.1 |
| 752 | HORVU5Hr1G063380 | AT5G10690.2 |
| 753 | HORVU3Hr1G046900 | AT1G73740.1 |
| 754 | HORVU4Hr1G063270 | AT1G19835.6 |
| 755 | HORVU4Hr1G066860 | AT5G16570.1 |
| 756 | HORVU2Hr1G039100 | AT1G04940.1 |
| 757 | HORVU5Hr1G079950 | AT1G42550.1 |
| 758 | HORVU4Hr1G009120 | AT4G37910.2 |
| 759 | HORVU7Hr1G078490 | AT2G27370.1 |
| 760 | HORVU7Hr1G080270 | AT5G08560.3 |
| 761 | HORVU5Hr1G082310 | AT5G47070.1 |
| 762 | HORVU3Hr1G019750 | AT1G51950.1 |
| 763 | HORVU5Hr1G062880 | AT1G14330.1 |
| 764 | HORVU5Hr1G056270 | AT1G23050.1 |
| 765 | HORVU1Hr1G063010 | AT3G60910.1 |
| 766 | HORVU2Hr1G065700 | AT5G39380.5 |
| 767 | HORVU7Hr1G098410 | AT5G61760.2 |
| 768 | HORVU2Hr1G017450 | AT2G39050.1 |
| 769 | HORVU3Hr1G030010 | AT1G64150.1 |
| 770 | HORVU2Hr1G029900 | AT1G78290.3 |
| 771 | HORVU5Hr1G097060 | AT4G33920.1 |
| 772 | HORVU1Hr1G042490 | AT1G77670.1 |
| 773 | HORVU7Hr1G058030 | AT4G35250.1 |
| 774 | HORVU7Hr1G095100 | AT4G34190.1 |
| 775 | HORVU7Hr1G118340 | AT1G23000.1 |

| | | |
|---|---|---|
| 776 | HORVU3Hr1G087180 | AT5G41380.1 |
| 777 | HORVU2Hr1G007080 | AT1G71480.1 |
| 778 | HORVU7Hr1G069710 | AT1G16880.1 |
| 779 | HORVU4Hr1G079230 | AT2G36830.1 |
| 780 | HORVU2Hr1G007840 | AT2G28000.1 |
| 781 | HORVU4Hr1G007260 | AT1G73885.1 |
| 782 | HORVU3Hr1G075960 | AT1G19715.2 |
| 783 | HORVU4Hr1G081310 | AT2G17640.1 |
| 784 | HORVU2Hr1G029490 | AT5G65030.1 |
| 785 | HORVU0Hr1G014060 | AT4G20880.1 |
| 786 | HORVU0Hr1G039030 | AT1G25270.1 |
| 787 | HORVU1Hr1G064780 | AT5G48380.1 |
| 788 | HORVU6Hr1G064910 | AT1G48410.3 |
| 789 | HORVU1Hr1G026780 | AT5G11460.1 |
| 790 | HORVU3Hr1G080640 | AT5G01990.1 |
| 791 | HORVU7Hr1G033500 | AT4G16410.1 |
| 792 | HORVU1Hr1G000720 | AT5G08391.1 |
| 793 | HORVU2Hr1G045620 | AT1G50900.1 |
| 794 | HORVU7Hr1G114890 | AT4G01070.1 |
| 795 | HORVU4Hr1G050730 | AT5G15710.2 |
| 796 | HORVU2Hr1G076010 | AT2G28605.1 |
| 797 | HORVU2Hr1G039510 | AT5G45040.1 |
| 798 | HORVU1Hr1G055140 | AT4G17090.1 |
| 799 | HORVU4Hr1G050700 | AT3G51550.1 |
| 800 | HORVU7Hr1G050520 | AT3G10180.2 |
| 801 | HORVU6Hr1G072050 | AT3G07670.2 |
| 802 | HORVU1Hr1G089760 | AT3G27850.1 |
| 803 | HORVU2Hr1G094190 | AT2G46100.1 |
| 804 | HORVU6Hr1G061780 | AT4G18240.1 |
| 805 | HORVU3Hr1G026500 | AT1G78600.1 |
| 806 | HORVU2Hr1G104730 | #N/A |
| 807 | HORVU1Hr1G073790 | AT1G08290.1 |
| 808 | HORVU4Hr1G046120 | AT5G23010.3 |
| 809 | HORVU3Hr1G048410 | AT1G53580.2 |
| 810 | HORVU3Hr1G084830 | AT3G55840.1 |
| 811 | HORVU1Hr1G084090 | AT4G10770.1 |
| 812 | HORVU3Hr1G034640 | AT4G01935.1 |
| 813 | HORVU2Hr1G115620 | AT5G64550.1 |
| 814 | HORVU7Hr1G040720 | AT2G32990.1 |
| 815 | HORVU5Hr1G009430 | AT5G44760.1 |
| 816 | HORVU0Hr1G012920 | AT1G48480.1 |

| | | |
|---|---|---|
| 817 | HORVU4Hr1G080810 | AT1G52690.2 |
| 818 | HORVU7Hr1G113020 | AT4G28088.1 |
| 819 | HORVU3Hr1G077570 | AT4G21250.1 |
| 820 | HORVU0Hr1G004850 | AT4G17690.1 |
| 821 | HORVU3Hr1G026870 | AT5G58950.1 |
| 822 | HORVU7Hr1G083670 | AT5G06900.1 |
| 823 | HORVU6Hr1G066340 | AT5G56550.1 |
| 824 | HORVU4Hr1G032780 | AT3G62910.1 |
| 825 | HORVU3Hr1G010460 | AT5G05740.2 |
| 826 | HORVU3Hr1G093510 | AT5G16540.4 |
| 827 | HORVU3Hr1G105620 | AT4G16260.1 |
| 828 | HORVU0Hr1G015440 | #N/A |
| 829 | HORVU1Hr1G015000 | AT1G50732.1 |
| 830 | HORVU3Hr1G093220 | AT1G51630.1 |
| 831 | HORVU6Hr1G021050 | AT3G57062.2 |
| 832 | HORVU5Hr1G110380 | AT2G16430.2 |
| 833 | HORVU3Hr1G066960 | AT2G42690.1 |
| 834 | HORVU4Hr1G067930 | AT1G69260.1 |
| 835 | HORVU7Hr1G041380 | AT1G70680.2 |
| 836 | HORVU3Hr1G068840 | AT3G15354.5 |
| 837 | HORVU5Hr1G115340 | AT1G48350.1 |
| 838 | HORVU3Hr1G091150 | AT1G71900.1 |
| 839 | HORVU7Hr1G045640 | AT4G02340.1 |
| 840 | HORVU7Hr1G104490 | AT1G05200.6 |
| 841 | HORVU1Hr1G090190 | AT4G23810.1 |
| 842 | HORVU2Hr1G097670 | AT4G00030.1 |
| 843 | HORVU7Hr1G049870 | AT2G13360.3 |
| 844 | HORVU6Hr1G052420 | AT5G55480.1 |
| 845 | HORVU1Hr1G024070 | AT3G46290.1 |
| 846 | HORVU7Hr1G113890 | AT4G31805.1 |
| 847 | HORVU2Hr1G014710 | AT5G01710.1 |
| 848 | HORVU2Hr1G080870 | AT2G01770.1 |
| 849 | HORVU1Hr1G063220 | AT1G03160.1 |
| 850 | HORVU1Hr1G090130 | AT4G11070.2 |
| 851 | HORVU6Hr1G012750 | AT3G47460.1 |
| 852 | HORVU6Hr1G078480 | AT5G11360.2 |
| 853 | HORVU2Hr1G086920 | AT1G80530.1 |
| 854 | HORVU4Hr1G002350 | AT4G26370.1 |
| 855 | HORVU2Hr1G031570 | AT3G23550.1 |
| 856 | HORVU5Hr1G103590 | AT1G61290.2 |
| 857 | HORVU2Hr1G072890 | AT2G27500.2 |

| | | |
|---|---|---|
| 858 | HORVU1Hr1G079400 | AT1G20110.1 |
| 859 | HORVU6Hr1G059990 | AT5G14895.1 |
| 860 | HORVU7Hr1G105950 | #N/A |
| 861 | HORVU3Hr1G037780 | AT2G41050.3 |
| 862 | HORVU2Hr1G039070 | AT5G12480.1 |
| 863 | HORVU7Hr1G109180 | AT1G30900.1 |
| 864 | HORVU1Hr1G054930 | AT4G17680.1 |
| 865 | HORVU2Hr1G022780 | AT2G41250.1 |
| 866 | HORVU0Hr1G007660 | AT3G58940.2 |
| 867 | HORVU7Hr1G113830 | AT1G80840.1 |
| 868 | HORVU5Hr1G103990 | AT2G26720.1 |
| 869 | HORVU2Hr1G017080 | AT5G17780.2 |
| 870 | HORVU6Hr1G029160 | AT4G31860.1 |
| 871 | HORVU5Hr1G063820 | AT5G57800.1 |
| 872 | HORVU3Hr1G089510 | AT3G25500.1 |
| 873 | HORVU0Hr1G013900 | AT3G14415.3 |
| 874 | HORVU6Hr1G033290 | AT4G02600.2 |
| 875 | HORVU5Hr1G110000 | AT5G05170.1 |
| 876 | HORVU4Hr1G038570 | AT2G33800.1 |
| 877 | HORVU3Hr1G077530 | AT1G61740.1 |
| 878 | HORVU4Hr1G024950 | AT2G29970.1 |
| 879 | HORVU7Hr1G079720 | AT5G45910.1 |
| 880 | HORVU7Hr1G114700 | AT5G11070.1 |
| 881 | HORVU5Hr1G087210 | AT4G21390.2 |
| 882 | HORVU2Hr1G104810 | ATCG01050.1 |
| 883 | HORVU4Hr1G004160 | AT5G04500.1 |
| 884 | HORVU7Hr1G082380 | AT4G34020.1 |
| 885 | HORVU6Hr1G077440 | AT4G38100.1 |
| 886 | HORVU1Hr1G094050 | AT3G59110.1 |
| 887 | HORVU6Hr1G053550 | AT5G14480.1 |
| 888 | HORVU5Hr1G106790 | AT5G18570.1 |
| 889 | HORVU3Hr1G097330 | AT5G19180.1 |
| 890 | HORVU5Hr1G087970 | AT1G42550.1 |
| 891 | HORVU1Hr1G093390 | AT1G53300.1 |
| 892 | HORVU7Hr1G052920 | AT2G17510.1 |
| 893 | HORVU7Hr1G078020 | AT1G64430.2 |
| 894 | HORVU1Hr1G066530 | AT5G54660.1 |
| 895 | HORVU3Hr1G117590 | AT1G53990.1 |
| 896 | HORVU0Hr1G005320 | AT4G23060.1 |
| 897 | HORVU7Hr1G114510 | AT4G36730.2 |
| 898 | HORVU4Hr1G069990 | AT1G48390.3 |

| | | |
|---|---|---|
| 899 | HORVU4Hr1G077310 | AT1G72360.3 |
| 900 | HORVU7Hr1G002200 | AT3G30340.1 |
| 901 | HORVU0Hr1G029600 | AT3G17240.3 |
| 902 | HORVU5Hr1G098370 | AT1G49380.1 |
| 903 | HORVU0Hr1G009860 | AT1G13680.1 |
| 904 | HORVU5Hr1G092700 | AT5G06510.3 |
| 905 | HORVU4Hr1G017030 | AT5G55120.1 |
| 906 | HORVU1Hr1G071500 | AT3G59100.1 |
| 907 | HORVU4Hr1G035330 | AT5G38530.1 |
| 908 | HORVU2Hr1G106080 | AT5G16730.1 |
| 909 | HORVU4Hr1G023830 | AT5G58240.2 |
| 910 | HORVU4Hr1G021140 | AT2G27260.1 |
| 911 | HORVU4Hr1G016260 | AT1G35340.1 |
| 912 | HORVU4Hr1G074320 | AT4G19100.1 |
| 913 | HORVU2Hr1G111590 | AT3G22210.1 |
| 914 | HORVU2Hr1G072880 | AT2G27510.1 |
| 915 | HORVU2Hr1G021700 | AT2G40840.1 |
| 916 | HORVU3Hr1G010310 | AT3G07040.1 |
| 917 | HORVU3Hr1G090270 | AT3G25480.1 |
| 918 | HORVU6Hr1G080750 | AT2G15620.1 |
| 919 | HORVU6Hr1G077900 | AT5G64080.2 |
| 920 | HORVU3Hr1G067110 | AT2G26080.1 |
| 921 | HORVU3Hr1G077660 | AT1G27310.1 |
| 922 | HORVU6Hr1G034620 | AT3G52960.1 |
| 923 | HORVU4Hr1G075730 | AT3G52105.1 |
| 924 | HORVU4Hr1G056100 | AT5G01340.1 |
| 925 | HORVU3Hr1G016460 | AT1G07990.2 |
| 926 | HORVU2Hr1G084750 | AT5G36790.3 |
| 927 | HORVU2Hr1G078980 | AT4G24380.4 |
| 928 | HORVU7Hr1G050530 | AT2G42280.1 |
| 929 | HORVU7Hr1G117370 | AT3G51250.1 |
| 930 | HORVU3Hr1G107090 | AT1G24620.1 |
| 931 | HORVU7Hr1G000120 | AT2G46915.1 |
| 932 | HORVU0Hr1G016780 | AT5G45340.1 |
| 933 | HORVU5Hr1G120650 | AT3G23700.1 |
| 934 | HORVU3Hr1G044430 | AT1G57770.1 |
| 935 | HORVU2Hr1G025720 | AT4G25490.1 |
| 936 | HORVU1Hr1G013970 | AT5G16010.1 |
| 937 | HORVU2Hr1G016440 | AT1G13040.2 |
| 938 | HORVU2Hr1G111570 | AT5G67050.1 |
| 939 | HORVU6Hr1G028220 | AT3G07040.1 |

| | | |
|---|---|---|
| 940 | HORVU2Hr1G063740 | AT1G70580.4 |
| 941 | HORVU2Hr1G072530 | AT3G09970.1 |
| 942 | HORVU7Hr1G109280 | AT5G64380.1 |
| 943 | HORVU3Hr1G093230 | AT5G16750.1 |
| 944 | HORVU3Hr1G017000 | AT1G62340.1 |
| 945 | HORVU6Hr1G075520 | AT3G12940.1 |
| 946 | HORVU5Hr1G115460 | AT5G26742.2 |
| 947 | HORVU4Hr1G068870 | AT3G29290.5 |
| 948 | HORVU2Hr1G077970 | AT5G46290.1 |
| 949 | HORVU5Hr1G031210 | AT3G47600.1 |
| 950 | HORVU7Hr1G095160 | AT1G31350.1 |
| 951 | HORVU2Hr1G093210 | AT2G44740.1 |
| 952 | HORVU3Hr1G019820 | AT1G35190.1 |
| 953 | HORVU6Hr1G058780 | AT1G08650.1 |
| 954 | HORVU5Hr1G104230 | AT2G19440.2 |
| 955 | HORVU3Hr1G050800 | AT1G74730.1 |
| 956 | HORVU6Hr1G003300 | AT1G77760.1 |
| 957 | HORVU2Hr1G007070 | AT3G23510.1 |
| 958 | HORVU3Hr1G028410 | AT2G42350.1 |
| 959 | HORVU6Hr1G031870 | AT3G16785.3 |
| 960 | HORVU0Hr1G014040 | AT1G13680.1 |
| 961 | HORVU5Hr1G095540 | AT5G55630.2 |
| 962 | HORVU4Hr1G076180 | AT2G26340.1 |
| 963 | HORVU2Hr1G091360 | AT2G37710.1 |
| 964 | HORVU7Hr1G050540 | AT1G51140.1 |
| 965 | HORVU4Hr1G021820 | AT5G22390.1 |
| 966 | HORVU4Hr1G065660 | AT1G58340.1 |
| 967 | HORVU4Hr1G015250 | AT1G14440.2 |
| 968 | HORVU5Hr1G082460 | AT1G75680.1 |
| 969 | HORVU3Hr1G095300 | AT4G34190.1 |
| 970 | HORVU3Hr1G089190 | AT3G59300.1 |
| 971 | HORVU6Hr1G067620 | AT1G31920.1 |
| 972 | HORVU2Hr1G125280 | AT1G45688.1 |
| 973 | HORVU2Hr1G118050 | AT1G01490.2 |
| 974 | HORVU1Hr1G038750 | AT5G37180.3 |
| 975 | HORVU3Hr1G055940 | AT3G16060.1 |
| 976 | HORVU2Hr1G007000 | AT3G23510.1 |
| 977 | HORVU6Hr1G068690 | AT5G45340.1 |
| 978 | HORVU7Hr1G015160 | AT1G32928.1 |
| 979 | HORVU6Hr1G068910 | AT4G36530.2 |
| 980 | HORVU5Hr1G093850 | AT5G66560.1 |

| | | |
|---|---|---|
| 981 | HORVU1Hr1G076350 | AT1G78580.4 |
| 982 | HORVU4Hr1G019570 | AT4G38970.1 |
| 983 | HORVU6Hr1G010050 | AT2G17220.1 |
| 984 | HORVU2Hr1G036110 | AT2G24270.4 |
| 985 | HORVU6Hr1G000650 | AT1G68010.1 |
| 986 | HORVU7Hr1G081590 | AT3G46290.1 |
| 987 | HORVU2Hr1G108070 | AT3G18170.1 |
| 988 | HORVU5Hr1G109680 | AT1G55360.1 |
| 989 | HORVU7Hr1G092550 | AT3G30390.3 |
| 990 | HORVU6Hr1G082710 | AT1G22360.1 |
| 991 | HORVU2Hr1G099890 | AT3G15210.1 |
| 992 | HORVU5Hr1G053230 | AT3G52150.2 |
| 993 | HORVU7Hr1G043030 | AT3G02380.1 |
| 994 | HORVU4Hr1G015330 | AT1G71500.1 |
| 995 | HORVU2Hr1G077460 | AT1G64940.1 |
| 996 | HORVU2Hr1G092180 | AT1G62750.1 |
| 997 | HORVU2Hr1G022920 | AT5G04140.1 |
| 998 | HORVU0Hr1G017160 | AT5G39865.1 |
| 999 | HORVU5Hr1G048300 | AT2G29970.1 |
| 1000 | HORVU1Hr1G024400 | AT1G07110.1 |
| 1001 | HORVU5Hr1G042160 | AT3G52820.1 |
| 1002 | HORVU2Hr1G111540 | AT5G47770.1 |
| 1003 | HORVU1Hr1G073300 | AT2G38090.1 |
| 1004 | HORVU6Hr1G022680 | AT5G06320.1 |
| 1005 | HORVU2Hr1G010830 | AT4G39330.1 |
| 1006 | HORVU3Hr1G019830 | AT3G49180.1 |
| 1007 | HORVU1Hr1G064970 | AT5G09220.1 |
| 1008 | HORVU3Hr1G001080 | AT2G35350.1 |
| 1009 | HORVU2Hr1G015470 | AT1G09160.2 |
| 1010 | HORVU6Hr1G009500 | AT5G05340.1 |
| 1011 | HORVU4Hr1G051820 | AT4G23160.3 |
| 1012 | HORVU2Hr1G071100 | AT2G33550.1 |
| 1013 | HORVU1Hr1G068910 | AT3G47510.1 |
| 1014 | HORVU4Hr1G062330 | AT4G11000.1 |
| 1015 | HORVU1Hr1G025440 | AT2G41475.1 |
| 1016 | HORVU4Hr1G056080 | AT1G06640.1 |
| 1017 | HORVU7Hr1G098510 | AT2G18980.1 |
| 1018 | HORVU5Hr1G001730 | AT5G49660.1 |
| 1019 | HORVU6Hr1G000620 | AT5G66190.1 |
| 1020 | HORVU3Hr1G010370 | AT5G58760.1 |
| 1021 | HORVU2Hr1G061010 | AT5G07380.4 |

| | | |
|---|---|---|
| 1022 | HORVU7Hr1G112280 | #N/A |
| 1023 | HORVU6Hr1G031370 | AT1G07010.1 |
| 1024 | HORVU5Hr1G018220 | AT2G03500.1 |
| 1025 | HORVU6Hr1G090240 | #N/A |
| 1026 | HORVU3Hr1G022540 | AT1G04250.1 |
| 1027 | HORVU5Hr1G072980 | AT1G33540.1 |
| 1028 | HORVU3Hr1G115610 | AT4G32940.1 |
| 1029 | HORVU2Hr1G076880 | AT1G59740.1 |
| 1030 | HORVU3Hr1G082740 | AT1G44575.3 |
| 1031 | HORVU2Hr1G039000 | AT2G01870.1 |
| 1032 | HORVU7Hr1G084420 | AT5G55190.1 |
| 1033 | HORVU4Hr1G011500 | AT4G37930.1 |
| 1034 | HORVU7Hr1G078690 | AT4G25570.1 |
| 1035 | HORVU2Hr1G089970 | AT2G16850.1 |
| 1036 | HORVU6Hr1G066050 | AT3G61460.1 |
| 1037 | HORVU1Hr1G075520 | AT1G19715.1 |
| 1038 | HORVU1Hr1G049420 | AT1G01490.2 |
| 1039 | HORVU6Hr1G086210 | AT2G15580.1 |
| 1040 | HORVU4Hr1G075400 | AT5G02400.1 |
| 1041 | HORVU4Hr1G048560 | AT2G47900.3 |
| 1042 | HORVU4Hr1G027260 | AT2G39730.1 |
| 1043 | HORVU6Hr1G063310 | AT5G51460.4 |
| 1044 | HORVU5Hr1G018790 | AT1G66540.2 |
| 1045 | HORVU1Hr1G065810 | AT1G61250.1 |
| 1046 | HORVU7Hr1G114710 | AT5G11070.1 |
| 1047 | HORVU3Hr1G024970 | AT1G28650.1 |
| 1048 | HORVU6Hr1G072540 | AT1G73990.2 |
| 1049 | HORVU3Hr1G117770 | AT5G58620.1 |
| 1050 | HORVU1Hr1G026650 | AT3G24480.1 |
| 1051 | HORVU6Hr1G088610 | AT1G61680.1 |
| 1052 | HORVU0Hr1G023540 | AT5G51460.4 |
| 1053 | HORVU6Hr1G055660 | AT3G59710.3 |
| 1054 | HORVU4Hr1G009440 | AT4G37850.2 |
| 1055 | HORVU5Hr1G020500 | AT5G08050.1 |
| 1056 | HORVU2Hr1G060010 | AT3G14420.2 |
| 1057 | HORVU2Hr1G108090 | AT5G45100.2 |
| 1058 | HORVU6Hr1G067120 | AT2G35200.1 |
| 1059 | HORVU2Hr1G084420 | AT1G27300.1 |
| 1060 | HORVU7Hr1G071000 | #N/A |
| 1061 | HORVU0Hr1G020750 | AT2G44480.1 |
| 1062 | HORVU1Hr1G013760 | AT3G08530.1 |

| | | |
|---|---|---|
| 1063 | HORVU7Hr1G077750 | AT1G60950.1 |
| 1064 | HORVU7Hr1G112240 | AT5G40250.1 |
| 1065 | HORVU7Hr1G044780 | AT1G21450.1 |
| 1066 | HORVU4Hr1G004800 | AT1G27150.1 |
| 1067 | HORVU5Hr1G025610 | AT2G47880.1 |
| 1068 | HORVU7Hr1G047210 | AT1G68740.1 |
| 1069 | HORVU6Hr1G080420 | AT2G48020.3 |
| 1070 | HORVU7Hr1G057100 | AT3G61880.2 |
| 1071 | HORVU2Hr1G071940 | AT5G23120.1 |
| 1072 | HORVU4Hr1G085960 | AT1G01820.1 |
| 1073 | HORVU4Hr1G057020 | AT2G38905.1 |
| 1074 | HORVU3Hr1G056830 | AT1G06520.1 |
| 1075 | HORVU7Hr1G121250 | AT4G18910.1 |
| 1076 | HORVU2Hr1G098820 | AT5G45030.2 |
| 1077 | HORVU3Hr1G078860 | AT2G30140.1 |
| 1078 | HORVU3Hr1G018050 | AT4G28706.4 |
| 1079 | HORVU6Hr1G002550 | AT3G47450.2 |
| 1080 | HORVU1Hr1G022490 | AT5G60360.3 |
| 1081 | HORVU4Hr1G008160 | AT1G66170.1 |
| 1082 | HORVU7Hr1G119100 | AT1G34300.1 |
| 1083 | HORVU5Hr1G024410 | AT4G15800.1 |
| 1084 | HORVU5Hr1G025820 | AT3G62300.1 |
| 1085 | HORVU5Hr1G070140 | AT3G61180.1 |
| 1086 | HORVU7Hr1G082350 | AT3G16520.3 |
| 1087 | HORVU1Hr1G047100 | AT4G10380.1 |
| 1088 | HORVU6Hr1G045650 | AT2G41720.1 |
| 1089 | HORVU3Hr1G024980 | AT3G28223.1 |
| 1090 | HORVU7Hr1G088920 | AT3G24520.1 |
| 1091 | HORVU3Hr1G058230 | AT4G21200.3 |
| 1092 | HORVU2Hr1G096270 | AT1G76040.2 |
| 1093 | HORVU7Hr1G091560 | AT4G25470.1 |
| 1094 | HORVU5Hr1G077140 | AT5G41130.1 |
| 1095 | HORVU7Hr1G050500 | AT5G12900.1 |
| 1096 | HORVU3Hr1G057400 | AT5G42070.1 |
| 1097 | HORVU3Hr1G019320 | AT3G11760.1 |
| 1098 | HORVU2Hr1G111530 | AT3G14360.1 |
| 1099 | HORVU4Hr1G013190 | AT3G19540.1 |
| 1100 | HORVU1Hr1G076910 | AT4G18700.1 |
| 1101 | HORVU4Hr1G072060 | AT5G03380.1 |
| 1102 | HORVU7Hr1G089850 | AT3G50120.1 |
| 1103 | HORVU6Hr1G077610 | AT3G05200.1 |

| | | |
|---|---|---|
| 1104 | HORVU2Hr1G009980 | AT2G03200.1 |
| 1105 | HORVU2Hr1G089960 | AT5G63950.2 |
| 1106 | HORVU1Hr1G070290 | AT5G07020.1 |
| 1107 | HORVU1Hr1G052790 | AT1G78000.7 |
| 1108 | HORVU2Hr1G076970 | AT5G08490.8 |
| 1109 | HORVU2Hr1G096850 | AT4G11570.2 |
| 1110 | HORVU7Hr1G038530 | AT5G06540.1 |
| 1111 | HORVU6Hr1G010900 | AT2G39330.2 |
| 1112 | HORVU2Hr1G068610 | AT1G08550.3 |
| 1113 | HORVU1Hr1G017310 | AT5G17980.1 |
| 1114 | HORVU0Hr1G017560 | AT1G31800.1 |
| 1115 | HORVU4Hr1G014010 | AT1G15980.1 |
| 1116 | HORVU0Hr1G021130 | AT4G29900.2 |
| 1117 | HORVU7Hr1G076790 | AT1G10280.1 |
| 1118 | HORVU1Hr1G084390 | AT5G45910.1 |
| 1119 | HORVU0Hr1G025440 | AT3G15630.1 |
| 1120 | HORVU3Hr1G063630 | AT3G03280.1 |
| 1121 | HORVU7Hr1G098400 | AT5G20935.1 |
| 1122 | HORVU2Hr1G019560 | AT2G47710.1 |
| 1123 | HORVU2Hr1G029460 | AT4G11070.2 |
| 1124 | HORVU1Hr1G090110 | AT5G24110.1 |
| 1125 | HORVU7Hr1G041230 | AT1G19180.2 |
| 1126 | HORVU3Hr1G085180 | AT2G38090.1 |
| 1127 | HORVU3Hr1G031130 | AT5G25630.1 |
| 1128 | HORVU6Hr1G011790 | AT3G04810.3 |
| 1129 | HORVU4Hr1G008180 | AT1G66150.1 |
| 1130 | HORVU3Hr1G052490 | AT4G28940.1 |
| 1131 | HORVU3Hr1G005420 | AT1G27480.1 |
| 1132 | HORVU3Hr1G051330 | AT1G34000.1 |
| 1133 | HORVU7Hr1G057460 | AT4G38590.1 |
| 1134 | HORVU5Hr1G040430 | AT5G48970.2 |
| 1135 | HORVU5Hr1G119830 | AT2G36800.1 |
| 1136 | HORVU7Hr1G115600 | AT5G20270.1 |
| 1137 | HORVU2Hr1G079020 | AT4G24380.6 |
| 1138 | HORVU4Hr1G082800 | AT4G38225.1 |
| 1139 | HORVU5Hr1G088550 | AT3G43190.2 |
| 1140 | HORVU2Hr1G111580 | AT1G22990.1 |
| 1141 | HORVU5Hr1G075570 | AT4G12730.1 |
| 1142 | HORVU2Hr1G092710 | AT2G46680.1 |
| 1143 | HORVU3Hr1G050590 | AT5G26170.1 |
| 1144 | HORVU6Hr1G070290 | AT4G35320.1 |

| | | |
|---|---|---|
| 1145 | HORVU2Hr1G039880 | AT3G15520.3 |
| 1146 | HORVU1Hr1G029180 | AT1G17620.1 |
| 1147 | HORVU6Hr1G084650 | AT4G28080.1 |
| 1148 | HORVU4Hr1G063870 | AT4G34980.1 |
| 1149 | HORVU3Hr1G029750 | AT5G41761.1 |
| 1150 | HORVU5Hr1G001280 | #N/A |
| 1151 | HORVU4Hr1G083020 | AT5G51680.1 |
| 1152 | HORVU7Hr1G036410 | AT1G35210.1 |
| 1153 | HORVU1Hr1G024860 | AT2G46330.1 |
| 1154 | HORVU1Hr1G048610 | AT1G63120.1 |
| 1155 | HORVU4Hr1G078510 | AT1G60600.2 |
| 1156 | HORVU3Hr1G059810 | AT3G27850.1 |
| 1157 | HORVU4Hr1G075090 | AT2G35290.1 |
| 1158 | HORVU1Hr1G026080 | AT4G33500.1 |
| 1159 | HORVU6Hr1G080870 | AT2G41940.1 |
| 1160 | HORVU2Hr1G002870 | AT4G21120.1 |
| 1161 | HORVU7Hr1G021550 | AT3G49055.1 |
| 1162 | HORVU5Hr1G112100 | AT3G11740.1 |
| 1163 | HORVU3Hr1G109500 | AT3G50150.1 |
| 1164 | HORVU5Hr1G110940 | AT5G49690.1 |
| 1165 | HORVU6Hr1G054890 | AT5G06760.1 |
| 1166 | HORVU3Hr1G082210 | AT4G38620.1 |
| 1167 | HORVU4Hr1G010770 | AT3G57450.1 |
| 1168 | HORVU1Hr1G063450 | AT5G16080.1 |
| 1169 | HORVU6Hr1G077790 | AT4G08950.1 |
| 1170 | HORVU1Hr1G066510 | AT3G20680.1 |
| 1171 | HORVU2Hr1G039960 | AT2G20850.1 |
| 1172 | HORVU3Hr1G094000 | AT5G16530.2 |
| 1173 | HORVU2Hr1G075110 | AT1G49740.1 |
| 1174 | HORVU1Hr1G019680 | AT5G01130.1 |
| 1175 | HORVU5Hr1G067010 | AT5G06710.1 |
| 1176 | HORVU7Hr1G022310 | AT5G63790.1 |
| 1177 | HORVU1Hr1G054260 | AT4G00050.1 |
| 1178 | HORVU1Hr1G014140 | AT2G15020.1 |
| 1179 | HORVU3Hr1G011030 | AT5G22540.1 |
| 1180 | HORVU5Hr1G094770 | AT1G05385.2 |
| 1181 | HORVU5Hr1G037620 | #N/A |
| 1182 | HORVU2Hr1G002010 | AT1G77490.1 |
| 1183 | HORVU6Hr1G010890 | AT2G14540.1 |
| 1184 | HORVU4Hr1G059150 | AT5G48540.1 |
| 1185 | HORVU4Hr1G077350 | AT5G20700.1 |

| | | |
|---|---|---|
| 1186 | HORVU5Hr1G098920 | AT5G23870.4 |
| 1187 | HORVU2Hr1G084960 | ATMG00870.1 |
| 1188 | HORVU4Hr1G015570 | AT4G10120.3 |
| 1189 | HORVU7Hr1G078480 | AT1G26220.1 |
| 1190 | HORVU4Hr1G036300 | AT2G48060.3 |
| 1191 | HORVU7Hr1G090340 | AT3G15820.1 |
| 1192 | HORVU2Hr1G109040 | AT2G01060.1 |
| 1193 | HORVU2Hr1G126040 | AT1G71930.2 |
| 1194 | HORVU4Hr1G060200 | AT1G08540.1 |
| 1195 | HORVU3Hr1G000150 | AT1G10585.1 |
| 1196 | HORVU3Hr1G061130 | AT5G03170.1 |
| 1197 | HORVU4Hr1G066870 | AT3G56630.1 |
| 1198 | HORVU2Hr1G099470 | AT3G22142.1 |
| 1199 | HORVU0Hr1G038040 | AT1G66100.1 |
| 1200 | HORVU3Hr1G095080 | AT4G22950.2 |
| 1201 | HORVU2Hr1G029450 | AT4G11070.2 |
| 1202 | HORVU3Hr1G013180 | AT2G19130.1 |
| 1203 | HORVU7Hr1G096250 | AT1G16720.1 |
| 1204 | HORVU6Hr1G094040 | AT3G62360.2 |
| 1205 | HORVU2Hr1G063460 | AT4G37530.1 |
| 1206 | HORVU5Hr1G120390 | AT2G43940.1 |
| 1207 | HORVU3Hr1G066390 | AT2G46510.1 |
| 1208 | HORVU7Hr1G080510 | AT3G14940.2 |
| 1209 | HORVU7Hr1G096370 | AT1G78995.1 |
| 1210 | HORVU6Hr1G094410 | AT3G19900.1 |
| 1211 | HORVU7Hr1G006980 | AT3G07040.1 |
| 1212 | HORVU2Hr1G021270 | AT4G03140.1 |
| 1213 | HORVU4Hr1G016450 | AT5G09860.1 |
| 1214 | HORVU2Hr1G029950 | AT2G27660.1 |
| 1215 | HORVU3Hr1G095000 | AT5G09950.2 |
| 1216 | HORVU6Hr1G069030 | AT3G09510.1 |
| 1217 | HORVU5Hr1G096420 | AT1G69310.4 |
| 1218 | HORVU3Hr1G058210 | AT1G59820.1 |
| 1219 | HORVU6Hr1G035470 | AT5G67300.1 |
| 1220 | HORVU2Hr1G043780 | AT5G48150.4 |
| 1221 | HORVU3Hr1G056960 | AT5G40850.1 |
| 1222 | HORVU4Hr1G082700 | AT1G42970.1 |
| 1223 | HORVU3Hr1G086190 | AT1G13270.1 |
| 1224 | HORVU7Hr1G036990 | AT3G06530.5 |
| 1225 | HORVU5Hr1G032980 | AT4G38970.1 |
| 1226 | HORVU0Hr1G017530 | AT2G46020.6 |

| | | |
|---|---|---|
| 1227 | HORVU6Hr1G077770 | AT4G08950.1 |
| 1228 | HORVU2Hr1G021870 | AT5G19500.1 |
| 1229 | HORVU1Hr1G071410 | AT3G09530.1 |
| 1230 | HORVU2Hr1G008090 | AT2G39130.2 |
| 1231 | HORVU5Hr1G081060 | AT2G46660.1 |
| 1232 | HORVU4Hr1G040100 | AT1G28520.3 |
| 1233 | HORVU5Hr1G069480 | AT4G37080.3 |
| 1234 | HORVU1Hr1G091530 | AT1G75250.2 |
| 1235 | HORVU1Hr1G069560 | AT5G01920.2 |
| 1236 | HORVU6Hr1G071950 | AT5G48250.1 |
| 1237 | HORVU5Hr1G099170 | AT3G21670.1 |
| 1238 | HORVU4Hr1G043630 | AT2G46930.1 |
| 1239 | HORVU5Hr1G112880 | AT3G53600.1 |
| 1240 | HORVU1Hr1G050730 | AT2G35470.1 |
| 1241 | HORVU2Hr1G095380 | AT1G71120.1 |
| 1242 | HORVU3Hr1G040050 | AT4G15720.1 |
| 1243 | HORVU3Hr1G076570 | AT5G56540.1 |
| 1244 | HORVU1Hr1G062450 | AT3G06070.1 |
| 1245 | HORVU2Hr1G116660 | AT4G21445.1 |
| 1246 | HORVU4Hr1G082430 | AT1G27090.1 |
| 1247 | HORVU7Hr1G087630 | AT5G57580.1 |
| 1248 | HORVU6Hr1G048770 | AT2G38360.1 |
| 1249 | HORVU3Hr1G092820 | AT1G48000.1 |
| 1250 | HORVU5Hr1G077520 | AT4G22758.2 |
| 1251 | HORVU1Hr1G082310 | AT4G34410.1 |
| 1252 | HORVU5Hr1G096490 | AT2G27035.1 |
| 1253 | HORVU3Hr1G114250 | AT3G10550.1 |
| 1254 | HORVU5Hr1G068110 | AT5G67060.2 |
| 1255 | HORVU6Hr1G093240 | AT5G02120.1 |
| 1256 | HORVU7Hr1G037020 | #N/A |
| 1257 | HORVU4Hr1G043650 | AT5G51010.1 |
| 1258 | HORVU3Hr1G058830 | AT5G15900.1 |
| 1259 | HORVU5Hr1G118060 | AT2G35460.1 |
| 1260 | HORVU6Hr1G048660 | AT3G16800.5 |
| 1261 | HORVU7Hr1G037010 | AT1G33410.2 |
| 1262 | HORVU5Hr1G062720 | AT5G20820.1 |
| 1263 | HORVU6Hr1G026370 | AT1G30260.1 |
| 1264 | HORVU1Hr1G044080 | AT1G65690.1 |
| 1265 | HORVU1Hr1G026070 | AT3G15630.1 |
| 1266 | HORVU5Hr1G072770 | AT3G21570.1 |
| 1267 | HORVU6Hr1G056210 | AT1G43760.1 |

| | | |
|---|---|---|
| 1268 | HORVU2Hr1G075950 | AT2G02540.2 |
| 1269 | HORVU6Hr1G059850 | AT4G05090.1 |
| 1270 | HORVU2Hr1G035020 | AT4G25070.2 |
| 1271 | HORVU7Hr1G049860 | AT2G04780.2 |
| 1272 | HORVU3Hr1G083520 | AT1G17170.1 |
| 1273 | HORVU5Hr1G112060 | AT1G66100.1 |
| 1274 | HORVU6Hr1G066450 | AT1G18330.1 |
| 1275 | HORVU3Hr1G078150 | AT3G15210.1 |
| 1276 | HORVU4Hr1G056210 | AT2G30020.1 |
| 1277 | HORVU0Hr1G002410 | AT5G05190.1 |
| 1278 | HORVU3Hr1G105360 | AT2G41510.2 |
| 1279 | HORVU6Hr1G090190 | AT1G60000.1 |
| 1280 | HORVU2Hr1G104030 | AT5G46330.2 |
| 1281 | HORVU0Hr1G009120 | AT5G21950.1 |
| 1282 | HORVU1Hr1G029840 | AT2G05120.2 |
| 1283 | HORVU7Hr1G030210 | AT1G73040.1 |
| 1284 | HORVU3Hr1G040180 | AT2G39435.6 |
| 1285 | HORVU1Hr1G000170 | AT1G48490.3 |
| 1286 | HORVU4Hr1G009250 | AT5G44800.1 |
| 1287 | HORVU5Hr1G059210 | AT3G02230.1 |
| 1288 | HORVU0Hr1G002390 | AT4G05060.1 |
| 1289 | HORVU7Hr1G095870 | AT3G18180.1 |
| 1290 | HORVU1Hr1G024180 | AT5G03170.1 |
| 1291 | HORVU5Hr1G030220 | AT1G80490.3 |
| 1292 | HORVU1Hr1G078350 | AT3G18000.1 |
| 1293 | HORVU2Hr1G040790 | AT1G79360.1 |
| 1294 | HORVU5Hr1G073800 | AT2G33590.1 |
| 1295 | HORVU1Hr1G033190 | AT3G15095.2 |
| 1296 | HORVU7Hr1G041500 | AT1G73540.1 |
| 1297 | HORVU3Hr1G044870 | AT2G29440.1 |
| 1298 | HORVU5Hr1G101990 | #N/A |
| 1299 | HORVU6Hr1G091840 | AT4G00350.1 |
| 1300 | HORVU7Hr1G085070 | AT5G07680.2 |
| 1301 | HORVU7Hr1G027860 | AT5G04360.2 |
| 1302 | HORVU3Hr1G050380 | AT5G12010.1 |
| 1303 | HORVU6Hr1G048760 | AT2G27980.1 |
| 1304 | HORVU1Hr1G065700 | AT3G09660.2 |
| 1305 | HORVU0Hr1G016540 | AT4G25470.1 |
| 1306 | HORVU2Hr1G040800 | AT1G79360.1 |
| 1307 | HORVU4Hr1G068930 | AT4G30845.1 |
| 1308 | HORVU2Hr1G012460 | AT2G24630.2 |

| | | |
|---|---|---|
| 1309 | HORVU2Hr1G076630 | #N/A |
| 1310 | HORVU1Hr1G032950 | AT4G12680.1 |
| 1311 | HORVU0Hr1G003480 | AT1G79360.1 |
| 1312 | HORVU2Hr1G101230 | AT5G20960.2 |
| 1313 | HORVU4Hr1G076650 | AT1G09650.1 |
| 1314 | HORVU3Hr1G078170 | AT4G23160.3 |
| 1315 | HORVU7Hr1G090850 | AT4G26530.3 |
| 1316 | HORVU1Hr1G076730 | AT1G78440.1 |
| 1317 | HORVU2Hr1G001620 | AT1G32920.1 |
| 1318 | HORVU5Hr1G080450 | AT4G25470.1 |
| 1319 | HORVU3Hr1G012490 | AT5G51920.1 |
| 1320 | HORVU3Hr1G108680 | AT3G56970.1 |
| 1321 | HORVU4Hr1G076660 | AT1G78410.2 |
| 1322 | HORVU2Hr1G094870 | AT4G23030.1 |
| 1323 | HORVU7Hr1G095170 | AT2G16860.2 |
| 1324 | HORVU0Hr1G000620 | AT1G11040.1 |
| 1325 | HORVU6Hr1G068660 | AT1G31650.1 |
| 1326 | HORVU4Hr1G009240 | AT4G24110.1 |
| 1327 | HORVU4Hr1G087760 | AT2G45180.1 |
| 1328 | HORVU2Hr1G066100 | AT5G57150.1 |
| 1329 | HORVU7Hr1G084850 | AT4G11450.1 |
| 1330 | HORVU3Hr1G013900 | AT3G26330.1 |
| 1331 | HORVU6Hr1G041610 | AT1G31650.1 |
| 1332 | HORVU0Hr1G019740 | AT2G26110.1 |
| 1333 | HORVU6Hr1G084860 | AT2G16430.2 |
| 1334 | HORVU7Hr1G084860 | AT3G03305.1 |
| 1335 | HORVU4Hr1G069100 | AT1G78060.1 |
| 1336 | HORVU7Hr1G095860 | AT5G24165.2 |
| 1337 | HORVU2Hr1G114680 | AT3G22142.1 |
| 1338 | HORVU7Hr1G098260 | AT4G25810.1 |

SUPPLEMENTARY TABLE 6

| S No | 270 exclusively in "Bowman_Up": | |
|---|---|---|
| 1 | HORVU2Hr1G004610 | AT2G45550.1 |
| 2 | HORVU2Hr1G122570 | AT5G26330.1 |
| 3 | HORVU2Hr1G079440 | AT1G22360.1 |
| 4 | HORVU1Hr1G005510 | AT1G21240.1 |
| 5 | HORVU7Hr1G010110 | AT2G18640.1 |
| 6 | HORVU2Hr1G033520 | AT4G27250.1 |
| 7 | HORVU5Hr1G023690 | AT5G51770.1 |

| | | |
|---|---|---|
| 8 | HORVU3Hr1G027910 | AT5G45890.1 |
| 9 | HORVU3Hr1G112030 | AT3G01410.2 |
| 10 | HORVU7Hr1G097350 | AT5G53550.6 |
| 11 | HORVU7Hr1G010120 | AT5G01130.1 |
| 12 | HORVU2Hr1G088640 | AT2G33380.1 |
| 13 | HORVU2Hr1G079420 | AT1G22360.1 |
| 14 | HORVU2Hr1G124970 | AT1G71695.1 |
| 15 | HORVU3Hr1G109170 | AT5G42380.1 |
| 16 | HORVU2Hr1G098770 | AT5G03130.1 |
| 17 | HORVU1Hr1G089590 | AT1G67120.1 |
| 18 | HORVU7Hr1G035730 | AT3G50360.1 |
| 19 | HORVU7Hr1G009680 | AT3G47570.1 |
| 20 | HORVU7Hr1G043530 | AT1G47710.1 |
| 21 | HORVU6Hr1G001840 | AT4G38600.3 |
| 22 | HORVU2Hr1G040130 | AT1G07180.1 |
| 23 | HORVU5Hr1G078630 | AT5G47910.1 |
| 24 | HORVU1Hr1G088300 | AT1G29860.1 |
| 25 | HORVU5Hr1G047920 | AT2G38300.1 |
| 26 | HORVU5Hr1G111040 | AT5G49690.1 |
| 27 | HORVU4Hr1G078320 | AT1G51850.2 |
| 28 | HORVU6Hr1G079700 | AT1G77760.1 |
| 29 | HORVU7Hr1G107370 | AT1G56150.1 |
| 30 | HORVU7Hr1G042300 | AT2G02850.1 |
| 31 | HORVU4Hr1G081590 | AT5G10530.1 |
| 32 | HORVU4Hr1G010720 | AT1G72060.1 |
| 33 | HORVU3Hr1G050810 | AT3G21305.1 |
| 34 | HORVU3Hr1G077130 | AT4G21390.1 |
| 35 | HORVU1Hr1G049260 | AT1G10370.1 |
| 36 | HORVU2Hr1G113120 | AT5G59700.1 |
| 37 | HORVU7Hr1G046180 | AT2G29580.1 |
| 38 | HORVU2Hr1G111760 | AT4G32300.1 |
| 39 | HORVU4Hr1G084810 | AT5G24530.1 |
| 40 | HORVU7Hr1G108150 | AT2G05940.1 |
| 41 | HORVU2Hr1G012980 | AT1G13110.1 |
| 42 | HORVU3Hr1G021500 | AT1G62270.1 |
| 43 | HORVU2Hr1G013760 | AT2G44110.2 |
| 44 | HORVU3Hr1G086980 | AT2G39420.1 |
| 45 | HORVU1Hr1G004980 | AT2G41640.1 |
| 46 | HORVU3Hr1G014140 | AT1G61110.1 |
| 47 | HORVU7Hr1G038330 | AT1G58360.1 |
| 48 | HORVU1Hr1G014460 | #N/A |

| | | |
|---|---|---|
| 49 | HORVU5Hr1G046730 | AT3G17700.2 |
| 50 | HORVU1Hr1G037270 | AT4G19810.2 |
| 51 | HORVU2Hr1G067340 | AT1G07725.1 |
| 52 | HORVU1Hr1G067840 | AT1G02460.1 |
| 53 | HORVU0Hr1G016280 | AT5G20360.6 |
| 54 | HORVU0Hr1G016310 | AT3G54420.1 |
| 55 | HORVU7Hr1G077970 | AT4G36945.1 |
| 56 | HORVU3Hr1G018920 | AT1G71010.1 |
| 57 | HORVU5Hr1G041590 | AT4G29180.1 |
| 58 | HORVU4Hr1G076750 | AT2G23620.1 |
| 59 | HORVU6Hr1G027640 | AT1G56150.1 |
| 60 | HORVU4Hr1G081430 | AT2G17040.1 |
| 61 | HORVU4Hr1G063660 | AT3G02100.1 |
| 62 | HORVU1Hr1G081410 | AT2G42750.1 |
| 63 | HORVU5Hr1G068320 | AT4G36990.1 |
| 64 | HORVU1Hr1G073820 | AT2G27500.2 |
| 65 | HORVU4Hr1G017390 | AT4G33490.2 |
| 66 | HORVU7Hr1G086250 | AT3G24460.2 |
| 67 | HORVU5Hr1G095580 | AT5G45890.1 |
| 68 | HORVU2Hr1G120660 | AT3G53810.1 |
| 69 | HORVU1Hr1G067870 | AT3G12490.1 |
| 70 | HORVU3Hr1G113100 | AT3G13790.2 |
| 71 | HORVU3Hr1G117470 | AT1G35190.2 |
| 72 | HORVU6Hr1G062380 | AT4G17500.1 |
| 73 | HORVU7Hr1G089160 | AT5G44400.1 |
| 74 | HORVU1Hr1G074310 | AT4G23180.1 |
| 75 | HORVU1Hr1G081300 | AT4G18880.1 |
| 76 | HORVU1Hr1G054290 | AT1G71950.1 |
| 77 | HORVU2Hr1G089020 | AT4G26090.1 |
| 78 | HORVU7Hr1G089530 | AT1G53820.1 |
| 79 | HORVU3Hr1G110580 | AT4G04750.3 |
| 80 | HORVU2Hr1G094640 | AT5G54080.2 |
| 81 | HORVU5Hr1G104620 | AT2G32010.4 |
| 82 | HORVU2Hr1G123480 | AT5G22250.1 |
| 83 | HORVU3Hr1G070700 | AT5G25930.1 |
| 84 | HORVU2Hr1G091410 | AT3G14460.1 |
| 85 | HORVU2Hr1G075220 | AT1G45688.2 |
| 86 | HORVU7Hr1G086260 | AT3G46990.1 |
| 87 | HORVU4Hr1G053130 | AT5G49480.1 |
| 88 | HORVU1Hr1G071670 | AT5G24530.1 |
| 89 | HORVU4Hr1G072480 | AT2G32300.1 |

| | | |
|---|---|---|
| 90 | HORVU2Hr1G044640 | AT4G23160.1 |
| 91 | HORVU3Hr1G092680 | AT3G24503.1 |
| 92 | HORVU7Hr1G031360 | AT5G11210.11 |
| 93 | HORVU3Hr1G033740 | AT1G62300.1 |
| 94 | HORVU2Hr1G120420 | AT5G22010.2 |
| 95 | HORVU7Hr1G104350 | AT3G20570.1 |
| 96 | HORVU4Hr1G009210 | AT1G75170.2 |
| 97 | HORVU4Hr1G055280 | AT1G13520.1 |
| 98 | HORVU1Hr1G093950 | AT5G10530.1 |
| 99 | HORVU6Hr1G018070 | AT1G55910.1 |
| 100 | HORVU2Hr1G032780 | AT1G26360.1 |
| 101 | HORVU2Hr1G036570 | AT2G16120.1 |
| 102 | HORVU4Hr1G059170 | AT4G27270.1 |
| 103 | HORVU3Hr1G077580 | AT5G15180.1 |
| 104 | HORVU7Hr1G101240 | AT1G33320.3 |
| 105 | HORVU7Hr1G077710 | AT1G34260.3 |
| 106 | HORVU5Hr1G058170 | AT4G39720.4 |
| 107 | HORVU4Hr1G055270 | AT5G37070.1 |
| 108 | HORVU2Hr1G003830 | AT4G34880.1 |
| 109 | HORVU3Hr1G022640 | AT1G47480.1 |
| 110 | HORVU4Hr1G076820 | AT2G17390.1 |
| 111 | HORVU6Hr1G074440 | AT5G10230.1 |
| 112 | HORVU6Hr1G066920 | AT4G17830.1 |
| 113 | HORVU5Hr1G105930 | AT4G15630.1 |
| 114 | HORVU3Hr1G092250 | AT5G14420.4 |
| 115 | HORVU1Hr1G017700 | AT3G05880.1 |
| 116 | HORVU2Hr1G118110 | AT1G01490.2 |
| 117 | HORVU3Hr1G085810 | AT2G37790.1 |
| 118 | HORVU5Hr1G062290 | AT1G74950.1 |
| 119 | HORVU7Hr1G030570 | AT2G24040.1 |
| 120 | HORVU4Hr1G083070 | #N/A |
| 121 | HORVU2Hr1G022080 | AT2G46950.1 |
| 122 | HORVU7Hr1G047070 | AT5G52450.1 |
| 123 | HORVU3Hr1G012100 | AT4G32300.1 |
| 124 | HORVU1Hr1G070580 | AT4G18640.1 |
| 125 | HORVU7Hr1G108460 | AT5G20480.2 |
| 126 | HORVU7Hr1G039900 | AT3G01360.2 |
| 127 | HORVU2Hr1G045900 | AT1G61800.1 |
| 128 | HORVU5Hr1G119270 | AT1G21230.1 |
| 129 | HORVU3Hr1G087050 | AT3G12360.1 |
| 130 | HORVU2Hr1G114100 | AT1G74170.2 |

| | | |
|---|---|---|
| 131 | HORVU1Hr1G091600 | AT1G27680.1 |
| 132 | HORVU1Hr1G020450 | AT1G77330.1 |
| 133 | HORVU0Hr1G007620 | AT4G08850.1 |
| 134 | HORVU6Hr1G023280 | AT1G17240.1 |
| 135 | HORVU3Hr1G024650 | AT2G19130.1 |
| 136 | HORVU5Hr1G110220 | AT3G54700.2 |
| 137 | HORVU5Hr1G107290 | AT1G59710.2 |
| 138 | HORVU1Hr1G074660 | AT1G71950.1 |
| 139 | HORVU6Hr1G031530 | AT1G54320.1 |
| 140 | HORVU6Hr1G023530 | AT1G21230.1 |
| 141 | HORVU6Hr1G034380 | AT1G76600.1 |
| 142 | HORVU6Hr1G034570 | AT2G33580.1 |
| 143 | HORVU2Hr1G044870 | AT4G23180.1 |
| 144 | HORVU3Hr1G086220 | AT3G55470.2 |
| 145 | HORVU2Hr1G098390 | AT1G56120.3 |
| 146 | HORVU4Hr1G007340 | AT1G13330.1 |
| 147 | HORVU1Hr1G074330 | AT1G43670.1 |
| 148 | HORVU4Hr1G084600 | AT2G40095.3 |
| 149 | HORVU3Hr1G079000 | AT5G12200.1 |
| 150 | HORVU1Hr1G088680 | AT3G51550.1 |
| 151 | HORVU3Hr1G028280 | AT2G42360.1 |
| 152 | HORVU1Hr1G066120 | AT1G03280.3 |
| 153 | HORVU4Hr1G016770 | AT1G77120.1 |
| 154 | HORVU1Hr1G020410 | AT1G77330.1 |
| 155 | HORVU2Hr1G007760 | AT5G60740.1 |
| 156 | HORVU7Hr1G059050 | AT2G36310.1 |
| 157 | HORVU7Hr1G079610 | #N/A |
| 158 | HORVU0Hr1G026470 | AT4G34215.2 |
| 159 | HORVU4Hr1G010030 | AT1G34300.1 |
| 160 | HORVU5Hr1G064020 | AT2G23810.1 |
| 161 | HORVU7Hr1G030480 | AT1G80820.1 |
| 162 | HORVU2Hr1G120260 | AT2G28290.1 |
| 163 | HORVU4Hr1G009340 | AT2G16250.1 |
| 164 | HORVU5Hr1G107410 | AT3G28510.1 |
| 165 | HORVU3Hr1G076060 | AT2G16430.2 |
| 166 | HORVU1Hr1G002890 | AT1G03940.1 |
| 167 | HORVU7Hr1G079600 | AT1G17147.1 |
| 168 | HORVU1Hr1G062250 | AT3G16520.3 |
| 169 | HORVU3Hr1G003920 | AT4G00970.1 |
| 170 | HORVU1Hr1G007310 | AT1G07670.2 |
| 171 | HORVU0Hr1G012940 | AT4G37980.2 |

| | | |
|---|---|---|
| 172 | HORVU2Hr1G103890 | AT1G16670.1 |
| 173 | HORVU3Hr1G049640 | AT4G33050.3 |
| 174 | HORVU4Hr1G058550 | AT1G67060.1 |
| 175 | HORVU3Hr1G088200 | AT2G38470.1 |
| 176 | HORVU3Hr1G002350 | AT5G38280.1 |
| 177 | HORVU2Hr1G116910 | AT2G20900.3 |
| 178 | HORVU7Hr1G038160 | AT1G14290.1 |
| 179 | HORVU2Hr1G100450 | AT4G10310.1 |
| 180 | HORVU1Hr1G074350 | AT5G09810.1 |
| 181 | HORVU7Hr1G104190 | AT3G47100.1 |
| 182 | HORVU2Hr1G078510 | AT1G17420.1 |
| 183 | HORVU2Hr1G107550 | AT2G14170.1 |
| 184 | HORVU3Hr1G081100 | AT3G08760.3 |
| 185 | HORVU7Hr1G122130 | AT2G25800.1 |
| 186 | HORVU5Hr1G014980 | AT5G43600.1 |
| 187 | HORVU6Hr1G033980 | AT3G60100.3 |
| 188 | HORVU3Hr1G111360 | AT5G01950.8 |
| 189 | HORVU6Hr1G028790 | AT1G80840.1 |
| 190 | HORVU2Hr1G084210 | AT3G28450.1 |
| 191 | HORVU0Hr1G001430 | AT5G64810.1 |
| 192 | HORVU3Hr1G106950 | AT1G24620.1 |
| 193 | HORVU3Hr1G061700 | AT4G03550.1 |
| 194 | HORVU6Hr1G077890 | AT5G64080.2 |
| 195 | HORVU5Hr1G068060 | AT5G14040.1 |
| 196 | HORVU1Hr1G086170 | AT1G53708.1 |
| 197 | HORVU7Hr1G008760 | AT5G61210.2 |
| 198 | HORVU6Hr1G061450 | AT5G53550.6 |
| 199 | HORVU7Hr1G116880 | AT1G30755.6 |
| 200 | HORVU1Hr1G073280 | AT3G47000.1 |
| 201 | HORVU0Hr1G017540 | AT3G24170.3 |
| 202 | HORVU2Hr1G117490 | AT1G65790.3 |
| 203 | HORVU6Hr1G041840 | ATMG00860.1 |
| 204 | HORVU6Hr1G066930 | AT4G17830.1 |
| 205 | HORVU2Hr1G040900 | AT4G30210.3 |
| 206 | HORVU2Hr1G016000 | AT5G26860.1 |
| 207 | HORVU5Hr1G099670 | AT4G32760.1 |
| 208 | HORVU6Hr1G078650 | AT2G03220.1 |
| 209 | HORVU6Hr1G077010 | AT2G32990.1 |
| 210 | HORVU1Hr1G039600 | AT4G04220.1 |
| 211 | HORVU6Hr1G024100 | AT4G29010.1 |
| 212 | HORVU2Hr1G031460 | AT3G51430.2 |

| | | |
|---|---|---|
| 213 | HORVU5Hr1G089790 | AT1G21240.1 |
| 214 | HORVU1Hr1G053260 | AT2G45910.2 |
| 215 | HORVU4Hr1G043530 | AT3G52430.1 |
| 216 | HORVU1Hr1G058810 | AT3G62830.2 |
| 217 | HORVU6Hr1G059420 | AT1G65730.1 |
| 218 | HORVU7Hr1G005270 | AT3G03050.1 |
| 219 | HORVU1Hr1G070640 | AT3G11110.1 |
| 220 | HORVU2Hr1G078860 | AT1G47380.1 |
| 221 | HORVU5Hr1G085310 | AT1G07250.1 |
| 222 | HORVU1Hr1G063410 | AT3G18170.1 |
| 223 | HORVU2Hr1G104450 | AT4G36400.2 |
| 224 | HORVU7Hr1G053300 | AT3G25070.2 |
| 225 | HORVU4Hr1G061130 | AT3G09010.1 |
| 226 | HORVU1Hr1G073840 | AT2G27510.1 |
| 227 | HORVU5Hr1G097500 | AT1G27730.1 |
| 228 | HORVU3Hr1G071890 | AT3G23710.1 |
| 229 | HORVU5Hr1G103180 | AT4G05390.1 |
| 230 | HORVU3Hr1G053380 | AT1G15520.1 |
| 231 | HORVU2Hr1G015410 | AT5G01740.1 |
| 232 | HORVU2Hr1G013200 | AT5G13190.1 |
| 233 | HORVU7Hr1G106420 | AT2G38880.8 |
| 234 | HORVU1Hr1G020430 | AT1G77330.1 |
| 235 | HORVU7Hr1G020580 | #N/A |
| 236 | HORVU3Hr1G086040 | AT2G39210.1 |
| 237 | HORVU7Hr1G040030 | AT1G19020.1 |
| 238 | HORVU5Hr1G006900 | AT1G58030.1 |
| 239 | HORVU2Hr1G015360 | AT5G01740.1 |
| 240 | HORVU4Hr1G027150 | AT3G54110.1 |
| 241 | HORVU4Hr1G064720 | AT1G59960.1 |
| 242 | HORVU1Hr1G038330 | AT2G33150.1 |
| 243 | HORVU4Hr1G027660 | AT3G09390.2 |
| 244 | HORVU5Hr1G092960 | AT5G46180.1 |
| 245 | HORVU5Hr1G115100 | AT5G13200.1 |
| 246 | HORVU2Hr1G104110 | AT1G30620.6 |
| 247 | HORVU2Hr1G075470 | AT5G63650.1 |
| 248 | HORVU3Hr1G007680 | AT5G38260.1 |
| 249 | HORVU7Hr1G022970 | AT1G13060.1 |
| 250 | HORVU4Hr1G002180 | AT5G02620.4 |
| 251 | HORVU2Hr1G011610 | AT1G12360.1 |
| 252 | HORVU3Hr1G085460 | AT2G37970.1 |
| 253 | HORVU5Hr1G118460 | AT1G34300.1 |

| | | |
|---|---|---|
| 254 | HORVU5Hr1G012890 | AT3G51000.1 |
| 255 | HORVU4Hr1G081670 | AT5G47910.1 |
| 256 | HORVU4Hr1G078130 | AT1G28200.2 |
| 257 | HORVU4Hr1G007050 | AT2G01490.2 |
| 258 | HORVU4Hr1G063890 | AT5G53670.1 |
| 259 | HORVU1Hr1G076870 | AT2G14260.3 |
| 260 | HORVU3Hr1G013880 | AT4G29010.1 |
| 261 | HORVU4Hr1G059120 | AT1G78570.1 |
| 262 | HORVU5Hr1G050350 | AT1G72190.1 |
| 263 | HORVU6Hr1G066470 | AT4G35310.2 |
| 264 | HORVU2Hr1G080070 | AT5G15140.1 |
| 265 | HORVU1Hr1G084800 | AT4G13400.1 |
| 266 | HORVU3Hr1G015150 | AT1G27980.1 |
| 267 | HORVU7Hr1G109810 | AT4G31350.4 |
| 268 | HORVU2Hr1G065750 | AT4G31550.1 |
| 269 | HORVU4Hr1G084510 | AT5G24010.1 |
| 270 | HORVU6Hr1G091130 | AT2G33150.1 |

SUPPLEMENTARY TABLE 7

| S No | 224exclusively in "Bowman_Down": | |
|---|---|---|
| 1 | HORVU4Hr1G072240 | AT2G36895.2 |
| 2 | HORVU1Hr1G073170 | AT1G32540.1 |
| 3 | HORVU7Hr1G106280 | AT5G37020.1 |
| 4 | HORVU7Hr1G105250 | AT5G57510.1 |
| 5 | HORVU0Hr1G032640 | #N/A |
| 6 | HORVU3Hr1G029560 | AT1G61230.1 |
| 7 | HORVU4Hr1G090780 | AT1G22900.1 |
| 8 | HORVU6Hr1G003160 | AT4G35160.1 |
| 9 | HORVU5Hr1G027740 | AT5G55060.1 |
| 10 | HORVU4Hr1G006200 | AT1G60890.2 |
| 11 | HORVU1Hr1G028680 | AT4G29670.1 |
| 12 | HORVU6Hr1G068720 | AT4G19170.1 |
| 13 | HORVU3Hr1G088580 | AT3G24255.3 |
| 14 | HORVU7Hr1G068610 | AT5G41150.1 |
| 15 | HORVU6Hr1G065710 | AT4G11810.3 |
| 16 | HORVU3Hr1G007480 | AT1G16840.4 |
| 17 | HORVU0Hr1G019860 | ATMG00030.1 |
| 18 | HORVU7Hr1G086810 | AT5G56860.1 |
| 19 | HORVU3Hr1G089390 | AT2G31560.3 |
| 20 | HORVU6Hr1G070420 | AT3G12120.2 |

| | | |
|---|---|---|
| 21 | HORVU4Hr1G024050 | AT2G39080.1 |
| 22 | HORVU5Hr1G112580 | AT3G59490.3 |
| 23 | HORVU2Hr1G116160 | #N/A |
| 24 | HORVU3Hr1G083470 | AT2G41430.5 |
| 25 | HORVU0Hr1G031330 | #N/A |
| 26 | HORVU1Hr1G077020 | AT1G34470.1 |
| 27 | HORVU1Hr1G057680 | AT4G16260.1 |
| 28 | HORVU6Hr1G072200 | AT5G60200.1 |
| 29 | HORVU6Hr1G091360 | AT5G04250.2 |
| 30 | HORVU5Hr1G080980 | AT4G10750.1 |
| 31 | HORVU7Hr1G088160 | AT1G58230.3 |
| 32 | HORVU0Hr1G025070 | AT2G07706.1 |
| 33 | HORVU7Hr1G121630 | AT4G37420.1 |
| 34 | HORVU6Hr1G016070 | AT3G25410.1 |
| 35 | HORVU2Hr1G115080 | AT4G20520.1 |
| 36 | HORVU4Hr1G062890 | AT5G63470.2 |
| 37 | HORVU3Hr1G098290 | AT2G23610.1 |
| 38 | HORVU1Hr1G046400 | AT5G38170.1 |
| 39 | HORVU0Hr1G010770 | AT3G26790.1 |
| 40 | HORVU3Hr1G011620 | AT2G44920.2 |
| 41 | HORVU1Hr1G091850 | AT2G42900.1 |
| 42 | HORVU0Hr1G024890 | #N/A |
| 43 | HORVU2Hr1G062850 | AT4G33625.3 |
| 44 | HORVU5Hr1G048240 | AT5G05580.1 |
| 45 | HORVU2Hr1G001600 | AT1G01090.1 |
| 46 | HORVU7Hr1G046320 | AT3G54890.1 |
| 47 | HORVU5Hr1G058300 | AT5G65140.1 |
| 48 | HORVU2Hr1G042370 | AT5G16910.1 |
| 49 | HORVU4Hr1G087610 | AT5G57150.5 |
| 50 | HORVU6Hr1G000090 | AT2G17850.1 |
| 51 | HORVU4Hr1G087230 | AT5G18280.1 |
| 52 | HORVU5Hr1G019000 | AT4G38180.1 |
| 53 | HORVU0Hr1G003270 | AT4G21280.1 |
| 54 | HORVU2Hr1G070440 | AT1G69040.2 |
| 55 | HORVU6Hr1G033160 | AT1G61520.3 |
| 56 | HORVU2Hr1G065110 | AT5G01220.1 |
| 57 | HORVU5Hr1G087250 | AT2G34420.1 |
| 58 | HORVU7Hr1G018800 | AT5G64440.1 |
| 59 | HORVU2Hr1G123750 | AT4G11600.1 |
| 60 | HORVU0Hr1G024880 | AT2G26790.2 |
| 61 | HORVU3Hr1G009210 | AT4G12800.2 |

| | | |
|---|---|---|
| 62 | HORVU2Hr1G070500 | AT1G33040.1 |
| 63 | HORVU2Hr1G066800 | AT5G24160.1 |
| 64 | HORVU6Hr1G063110 | AT5G64740.1 |
| 65 | HORVU3Hr1G105790 | AT3G57270.1 |
| 66 | HORVU7Hr1G069410 | AT1G15640.2 |
| 67 | HORVU0Hr1G024560 | ATCG00490.1 |
| 68 | HORVU0Hr1G017890 | AT5G63630.2 |
| 69 | HORVU4Hr1G071070 | AT1G11755.1 |
| 70 | HORVU1Hr1G054240 | AT1G65680.1 |
| 71 | HORVU1Hr1G047710 | AT5G62360.1 |
| 72 | HORVU4Hr1G059950 | AT2G39705.1 |
| 73 | HORVU5Hr1G044410 | AT2G26800.1 |
| 74 | HORVU0Hr1G034360 | AT3G20460.1 |
| 75 | HORVU4Hr1G018070 | AT1G04160.2 |
| 76 | HORVU5Hr1G066280 | AT3G47470.1 |
| 77 | HORVU2Hr1G041280 | AT3G08940.1 |
| 78 | HORVU7Hr1G115540 | AT1G15690.1 |
| 79 | HORVU3Hr1G089370 | AT1G05870.9 |
| 80 | HORVU1Hr1G043820 | AT5G08370.1 |
| 81 | HORVU7Hr1G085350 | AT1G80440.1 |
| 82 | HORVU2Hr1G062290 | AT1G17250.1 |
| 83 | HORVU4Hr1G048990 | AT5G20220.3 |
| 84 | HORVU6Hr1G092750 | AT5G04390.1 |
| 85 | HORVU2Hr1G071360 | AT2G21880.1 |
| 86 | HORVU2Hr1G044790 | AT1G52820.1 |
| 87 | HORVU4Hr1G009520 | AT4G14960.2 |
| 88 | HORVU5Hr1G059140 | AT4G38060.1 |
| 89 | HORVU1Hr1G045590 | AT1G58290.1 |
| 90 | HORVU2Hr1G102620 | AT5G17165.1 |
| 91 | HORVU3Hr1G075870 | AT2G30570.1 |
| 92 | HORVU4Hr1G090720 | AT3G16460.2 |
| 93 | HORVU0Hr1G032630 | AT5G56400.1 |
| 94 | HORVU4Hr1G073730 | AT2G38820.2 |
| 95 | HORVU2Hr1G113270 | AT3G16150.1 |
| 96 | HORVU5Hr1G124380 | AT5G24490.1 |
| 97 | HORVU3Hr1G078940 | AT5G12250.1 |
| 98 | HORVU1Hr1G045080 | AT1G28100.6 |
| 99 | HORVU0Hr1G007360 | AT1G44446.1 |
| 100 | HORVU6Hr1G056000 | AT5G24930.1 |
| 101 | HORVU3Hr1G059330 | #N/A |
| 102 | HORVU7Hr1G069520 | AT1G64860.1 |

| | | |
|---|---|---|
| 103 | HORVU1Hr1G076120 | AT4G32480.1 |
| 104 | HORVU5Hr1G032350 | AT3G55270.1 |
| 105 | HORVU5Hr1G109710 | AT1G03130.1 |
| 106 | HORVU3Hr1G089380 | AT1G05870.9 |
| 107 | HORVU2Hr1G079630 | AT1G14740.1 |
| 108 | HORVU1Hr1G073010 | AT3G54000.3 |
| 109 | HORVU2Hr1G036960 | AT3G61470.1 |
| 110 | HORVU2Hr1G121640 | AT5G26040.1 |
| 111 | HORVU2Hr1G042030 | #N/A |
| 112 | HORVU7Hr1G118130 | AT5G21482.1 |
| 113 | HORVU5Hr1G113900 | AT2G20260.1 |
| 114 | HORVU5Hr1G018680 | AT5G56220.2 |
| 115 | HORVU4Hr1G049000 | AT5G13700.1 |
| 116 | HORVU1Hr1G087320 | AT5G57550.1 |
| 117 | HORVU2Hr1G108430 | #N/A |
| 118 | HORVU4Hr1G045210 | AT2G36000.1 |
| 119 | HORVU3Hr1G005500 | AT5G57150.5 |
| 120 | HORVU2Hr1G040780 | AT5G54270.1 |
| 121 | HORVU7Hr1G069590 | AT5G49360.1 |
| 122 | HORVU3Hr1G021750 | AT3G24460.2 |
| 123 | HORVU3Hr1G077230 | AT1G79190.2 |
| 124 | HORVU7Hr1G098660 | AT4G25910.1 |
| 125 | HORVU3Hr1G009490 | AT5G59310.1 |
| 126 | HORVU5Hr1G119230 | AT5G64410.1 |
| 127 | HORVU5Hr1G095480 | AT4G16380.1 |
| 128 | HORVU7Hr1G107840 | AT1G14740.1 |
| 129 | HORVU6Hr1G055350 | AT1G60760.1 |
| 130 | HORVU2Hr1G017480 | AT4G32430.2 |
| 131 | HORVU2Hr1G019820 | AT5G64040.1 |
| 132 | HORVU5Hr1G001750 | AT5G17210.1 |
| 133 | HORVU3Hr1G080430 | AT2G38640.1 |
| 134 | HORVU3Hr1G014790 | AT4G24930.1 |
| 135 | HORVU5Hr1G114230 | AT4G21550.3 |
| 136 | HORVU5Hr1G109260 | AT2G34420.1 |
| 137 | HORVU0Hr1G028250 | AT2G07706.1 |
| 138 | HORVU3Hr1G052710 | AT5G56840.1 |
| 139 | HORVU3Hr1G011630 | AT2G23380.2 |
| 140 | HORVU5Hr1G064800 | AT4G22180.1 |
| 141 | HORVU3Hr1G065090 | AT1G15140.1 |
| 142 | HORVU5Hr1G094110 | AT5G62600.1 |
| 143 | HORVU4Hr1G054240 | AT4G16146.1 |

| | | |
|---|---|---|
| 144 | HORVU6Hr1G048120 | AT5G26110.3 |
| 145 | HORVU3Hr1G014590 | AT1G64040.1 |
| 146 | HORVU1Hr1G056630 | AT4G33030.1 |
| 147 | HORVU6Hr1G016890 | AT2G34420.1 |
| 148 | HORVU7Hr1G036780 | AT5G64260.1 |
| 149 | HORVU4Hr1G090750 | AT1G55250.3 |
| 150 | HORVU1Hr1G078380 | AT2G34420.1 |
| 151 | HORVU5Hr1G110610 | AT4G39590.1 |
| 152 | HORVU2Hr1G070140 | AT5G50740.5 |
| 153 | HORVU7Hr1G056700 | AT5G07220.1 |
| 154 | HORVU6Hr1G028970 | AT1G65870.1 |
| 155 | HORVU3Hr1G069280 | AT2G42760.1 |
| 156 | HORVU4Hr1G087580 | AT1G10586.1 |
| 157 | HORVU4Hr1G005320 | AT1G55265.1 |
| 158 | HORVU5Hr1G113910 | AT1G26090.1 |
| 159 | HORVU0Hr1G034230 | AT2G07706.1 |
| 160 | HORVU2Hr1G121770 | AT1G71140.1 |
| 161 | HORVU0Hr1G013070 | AT1G15950.7 |
| 162 | HORVU3Hr1G027220 | AT2G25770.2 |
| 163 | HORVU3Hr1G110330 | AT1G06800.1 |
| 164 | HORVU5Hr1G113060 | AT4G00680.1 |
| 165 | HORVU5Hr1G109610 | AT5G43150.1 |
| 166 | HORVU2Hr1G074730 | AT3G55480.1 |
| 167 | HORVU6Hr1G016940 | AT2G34420.1 |
| 168 | HORVU3Hr1G004080 | AT2G35110.1 |
| 169 | HORVU0Hr1G020630 | AT5G04370.3 |
| 170 | HORVU4Hr1G069690 | AT1G24430.1 |
| 171 | HORVU2Hr1G000420 | AT3G62160.1 |
| 172 | HORVU1Hr1G039250 | AT5G44030.3 |
| 173 | HORVU1Hr1G007690 | AT2G06541.2 |
| 174 | HORVU0Hr1G034200 | AT5G56400.1 |
| 175 | HORVU6Hr1G091650 | AT2G34420.1 |
| 176 | HORVU1Hr1G088920 | AT2G34420.1 |
| 177 | HORVU2Hr1G041270 | AT5G01530.1 |
| 178 | HORVU5Hr1G109250 | AT1G29930.1 |
| 179 | HORVU1Hr1G089240 | ATMG01360.1 |
| 180 | HORVU1Hr1G024890 | AT5G43750.1 |
| 181 | HORVU6Hr1G004000 | AT1G62660.2 |
| 182 | HORVU0Hr1G038990 | AT4G09890.1 |
| 183 | HORVU5Hr1G019480 | AT3G58090.1 |
| 184 | HORVU1Hr1G075160 | AT1G75290.1 |

| | | |
|---|---|---|
| 185 | HORVU1Hr1G089180 | AT2G34420.1 |
| 186 | HORVU6Hr1G016860 | AT5G18890.1 |
| 187 | HORVU4Hr1G088080 | AT1G65680.1 |
| 188 | HORVU2Hr1G075170 | AT1G67910.2 |
| 189 | HORVU2Hr1G102100 | AT5G48485.1 |
| 190 | HORVU7Hr1G040380 | AT2G34420.1 |
| 191 | HORVU7Hr1G112130 | AT5G28080.2 |
| 192 | HORVU6Hr1G016880 | AT2G34420.1 |
| 193 | HORVU1Hr1G071910 | AT5G02580.1 |
| 194 | HORVU4Hr1G018180 | AT4G22756.1 |
| 195 | HORVU2Hr1G038040 | AT5G43150.1 |
| 196 | HORVU1Hr1G073630 | AT2G06541.2 |
| 197 | HORVU2Hr1G064730 | AT1G11700.1 |
| 198 | HORVU2Hr1G094390 | AT2G18050.1 |
| 199 | HORVU2Hr1G051050 | AT4G24560.1 |
| 200 | HORVU7Hr1G122470 | AT3G45070.3 |
| 201 | HORVU7Hr1G040370 | AT2G34420.1 |
| 202 | HORVU2Hr1G051040 | AT3G11670.1 |
| 203 | HORVU0Hr1G005360 | AT1G33540.1 |
| 204 | HORVU7Hr1G002620 | AT1G72540.1 |
| 205 | HORVU7Hr1G090410 | AT5G13700.1 |
| 206 | HORVU3Hr1G001680 | AT3G23090.5 |
| 207 | HORVU5Hr1G063630 | AT3G22840.1 |
| 208 | HORVU1Hr1G057860 | AT2G41510.2 |
| 209 | HORVU2Hr1G032710 | AT1G74160.1 |
| 210 | HORVU6Hr1G016850 | AT2G34420.1 |
| 211 | HORVU3Hr1G025300 | AT2G42320.4 |
| 212 | HORVU1Hr1G083580 | AT5G25240.1 |
| 213 | HORVU5Hr1G067200 | AT3G61460.1 |
| 214 | HORVU3Hr1G094580 | AT3G24050.1 |
| 215 | HORVU6Hr1G012110 | AT1G61680.1 |
| 216 | HORVU1Hr1G088900 | AT1G29930.1 |
| 217 | HORVU6Hr1G079190 | AT5G15180.1 |
| 218 | HORVU2Hr1G090670 | AT1G13420.1 |
| 219 | HORVU5Hr1G082160 | AT5G53390.1 |
| 220 | HORVU2Hr1G003470 | AT2G22330.2 |
| 221 | HORVU5Hr1G084230 | AT2G37170.2 |
| 222 | HORVU5Hr1G101880 | AT1G60650.2 |
| 223 | HORVU2Hr1G103160 | AT5G39690.1 |
| 224 | HORVU2Hr1G043930 | AT4G12650.1 |

SUPPLEMENTARY TABLE 8

| S No | 634 in "Bowman_Up" and "Harrington_Up": | |
|---|---|---|
| 1 | HORVU2Hr1G004640 | AT3G26300.1 |
| 2 | HORVU2Hr1G004600 | AT3G26310.1 |
| 3 | HORVU1Hr1G089700 | AT5G54160.1 |
| 4 | HORVU6Hr1G012640 | AT5G43580.1 |
| 5 | HORVU2Hr1G098710 | AT3G49845.2 |
| 6 | HORVU2Hr1G004530 | AT5G25120.1 |
| 7 | HORVU7Hr1G012240 | AT5G63840.2 |
| 8 | HORVU7Hr1G011930 | AT1G45474.2 |
| 9 | HORVU2Hr1G004620 | AT4G02780.1 |
| 10 | HORVU6Hr1G001850 | AT2G29780.1 |
| 11 | HORVU1Hr1G089620 | AT5G54160.1 |
| 12 | HORVU5Hr1G041660 | AT1G51890.1 |
| 13 | HORVU7Hr1G011950 | AT5G35620.3 |
| 14 | HORVU3Hr1G112050 | AT3G55640.3 |
| 15 | HORVU7Hr1G033530 | AT2G14610.1 |
| 16 | HORVU6Hr1G014600 | AT1G61420.1 |
| 17 | HORVU1Hr1G089520 | AT4G35160.1 |
| 18 | HORVU6Hr1G020540 | AT3G26230.1 |
| 19 | HORVU1Hr1G065670 | AT5G47910.1 |
| 20 | HORVU1Hr1G089540 | AT5G13930.1 |
| 21 | HORVU6Hr1G005720 | AT1G08090.1 |
| 22 | HORVU6Hr1G013690 | AT3G51350.1 |
| 23 | HORVU7Hr1G011940 | AT5G60540.1 |
| 24 | HORVU2Hr1G028660 | AT2G38900.2 |
| 25 | HORVU5Hr1G096850 | AT5G57260.1 |
| 26 | HORVU2Hr1G004480 | AT3G26300.1 |
| 27 | HORVU2Hr1G126150 | AT4G15560.1 |
| 28 | HORVU3Hr1G099760 | AT2G46750.1 |
| 29 | HORVU3Hr1G074960 | AT4G11290.1 |
| 30 | HORVU0Hr1G011730 | AT5G65440.2 |
| 31 | HORVU2Hr1G028720 | AT5G43580.1 |
| 32 | HORVU2Hr1G122580 | AT2G32300.1 |
| 33 | HORVU0Hr1G011720 | AT1G35260.1 |
| 34 | HORVU4Hr1G001250 | AT5G54160.1 |
| 35 | HORVU5Hr1G124650 | AT1G66950.1 |
| 36 | HORVU7Hr1G017640 | AT2G36530.1 |
| 37 | HORVU2Hr1G028670 | AT5G43580.1 |
| 38 | HORVU5Hr1G055950 | AT4G33720.1 |

| | | |
|---|---|---|
| 39 | HORVU5Hr1G056040 | AT4G33720.1 |
| 40 | HORVU5Hr1G114700 | AT3G11340.1 |
| 41 | HORVU7Hr1G076120 | AT4G33420.1 |
| 42 | HORVU4Hr1G071370 | AT5G47530.1 |
| 43 | HORVU7Hr1G019380 | AT3G26300.1 |
| 44 | HORVU2Hr1G104240 | AT5G39160.1 |
| 45 | HORVU4Hr1G010740 | AT1G72060.1 |
| 46 | HORVU3Hr1G032370 | AT4G39230.1 |
| 47 | HORVU3Hr1G097860 | AT3G09220.1 |
| 48 | HORVU5Hr1G023720 | AT1G35260.1 |
| 49 | HORVU4Hr1G078370 | AT1G78880.1 |
| 50 | HORVU6Hr1G053710 | AT5G50200.3 |
| 51 | HORVU7Hr1G009760 | AT4G26440.1 |
| 52 | HORVU1Hr1G084900 | AT4G15560.1 |
| 53 | HORVU2Hr1G076320 | AT4G34600.1 |
| 54 | HORVU5Hr1G051970 | AT4G11650.1 |
| 55 | HORVU0Hr1G008830 | AT4G02600.2 |
| 56 | HORVU2Hr1G124300 | AT1G78380.1 |
| 57 | HORVU5Hr1G056420 | AT1G19250.1 |
| 58 | HORVU5Hr1G075540 | AT4G13010.1 |
| 59 | HORVU4Hr1G002640 | AT4G11650.1 |
| 60 | HORVU4Hr1G064790 | AT5G17330.1 |
| 61 | HORVU5Hr1G099390 | AT2G43000.1 |
| 62 | HORVU3Hr1G109350 | AT4G03230.3 |
| 63 | HORVU1Hr1G009490 | AT3G22600.1 |
| 64 | HORVU2Hr1G071570 | AT4G13420.1 |
| 65 | HORVU2Hr1G095460 | AT3G62760.1 |
| 66 | HORVU3Hr1G077790 | AT3G02210.1 |
| 67 | HORVU5Hr1G009840 | AT3G59920.1 |
| 68 | HORVU5Hr1G072520 | AT4G20820.1 |
| 69 | HORVU2Hr1G122270 | AT1G10360.1 |
| 70 | HORVU2Hr1G122620 | AT2G31050.1 |
| 71 | HORVU4Hr1G084330 | AT5G60710.1 |
| 72 | HORVU7Hr1G008830 | AT1G10360.1 |
| 73 | HORVU3Hr1G001840 | #N/A |
| 74 | HORVU5Hr1G005230 | AT4G11650.1 |
| 75 | HORVU1Hr1G020720 | AT5G19890.1 |
| 76 | HORVU5Hr1G123550 | AT4G21390.1 |
| 77 | HORVU2Hr1G001780 | AT5G13080.1 |
| 78 | HORVU7Hr1G019400 | AT2G37710.1 |
| 79 | HORVU1Hr1G022430 | AT2G24200.3 |

| | | |
|---|---|---|
| 80 | HORVU4Hr1G002650 | AT4G11650.1 |
| 81 | HORVU1Hr1G062380 | AT3G06390.1 |
| 82 | HORVU5Hr1G106010 | AT3G19690.1 |
| 83 | HORVU7Hr1G010990 | AT2G20340.1 |
| 84 | HORVU3Hr1G094340 | AT1G14590.1 |
| 85 | HORVU1Hr1G088550 | AT4G10930.2 |
| 86 | HORVU3Hr1G112060 | AT2G42270.1 |
| 87 | HORVU6Hr1G088400 | AT3G47570.1 |
| 88 | HORVU6Hr1G077940 | AT1G66170.1 |
| 89 | HORVU6Hr1G077960 | AT1G22360.2 |
| 90 | HORVU3Hr1G105400 | AT3G57270.1 |
| 91 | HORVU2Hr1G092440 | AT2G32240.1 |
| 92 | HORVU3Hr1G112080 | AT2G14680.4 |
| 93 | HORVU4Hr1G011760 | AT3G11340.1 |
| 94 | HORVU5Hr1G099460 | AT2G43000.1 |
| 95 | HORVU3Hr1G098620 | AT4G08350.1 |
| 96 | HORVU3Hr1G111600 | AT1G64790.1 |
| 97 | HORVU2Hr1G109640 | AT2G45550.1 |
| 98 | HORVU3Hr1G112140 | #N/A |
| 99 | HORVU6Hr1G021870 | AT5G06710.1 |
| 100 | HORVU3Hr1G094250 | AT4G19975.1 |
| 101 | HORVU1Hr1G090930 | AT2G31900.4 |
| 102 | HORVU1Hr1G049070 | AT1G10370.1 |
| 103 | HORVU5Hr1G023730 | AT1G35260.1 |
| 104 | HORVU3Hr1G096860 | AT4G13420.1 |
| 105 | HORVU3Hr1G109430 | AT4G03230.3 |
| 106 | HORVU7Hr1G107670 | AT1G14600.1 |
| 107 | HORVU2Hr1G119500 | AT5G36970.1 |
| 108 | HORVU6Hr1G061420 | AT5G53560.1 |
| 109 | HORVU3Hr1G105630 | AT4G16260.1 |
| 110 | HORVU3Hr1G018980 | AT5G56960.1 |
| 111 | HORVU5Hr1G066460 | AT1G23110.7 |
| 112 | HORVU5Hr1G080090 | AT1G33540.2 |
| 113 | HORVU2Hr1G004540 | AT1G79460.1 |
| 114 | HORVU2Hr1G114390 | AT2G20340.1 |
| 115 | HORVU2Hr1G098700 | #N/A |
| 116 | HORVU3Hr1G113120 | AT3G04720.1 |
| 117 | HORVU2Hr1G004510 | AT5G25120.1 |
| 118 | HORVU3Hr1G100190 | AT5G04370.1 |
| 119 | HORVU3Hr1G020210 | AT5G15130.1 |
| 120 | HORVU4Hr1G009230 | AT5G10530.1 |

| | | |
|---|---|---|
| 121 | HORVU7Hr1G110570 | AT4G24310.1 |
| 122 | HORVU1Hr1G014500 | AT1G66730.1 |
| 123 | HORVU2Hr1G102720 | AT2G02040.1 |
| 124 | HORVU3Hr1G069320 | AT1G29280.1 |
| 125 | HORVU2Hr1G003210 | AT1G11680.1 |
| 126 | HORVU7Hr1G082180 | AT1G16030.1 |
| 127 | HORVU2Hr1G114440 | AT2G20340.1 |
| 128 | HORVU6Hr1G088440 | AT5G20400.1 |
| 129 | HORVU2Hr1G071270 | AT5G13330.1 |
| 130 | HORVU2Hr1G112120 | AT5G48930.1 |
| 131 | HORVU1Hr1G063390 | AT5G40010.1 |
| 132 | HORVU2Hr1G012140 | AT1G74190.1 |
| 133 | HORVU1Hr1G092650 | AT3G16520.3 |
| 134 | HORVU6Hr1G077930 | AT1G22380.1 |
| 135 | HORVU5Hr1G106850 | AT1G66950.1 |
| 136 | HORVU7Hr1G056250 | AT4G28460.1 |
| 137 | HORVU3Hr1G006450 | AT1G03230.1 |
| 138 | HORVU2Hr1G102710 | AT2G02040.1 |
| 139 | HORVU1Hr1G001220 | AT4G23180.3 |
| 140 | HORVU1Hr1G075620 | AT3G47780.1 |
| 141 | HORVU1Hr1G010120 | AT2G20340.1 |
| 142 | HORVU5Hr1G089840 | AT1G21230.1 |
| 143 | HORVU6Hr1G069710 | AT4G21390.1 |
| 144 | HORVU2Hr1G122610 | AT2G31050.1 |
| 145 | HORVU3Hr1G095730 | AT1G10360.1 |
| 146 | HORVU1Hr1G072690 | AT2G37790.1 |
| 147 | HORVU2Hr1G004550 | AT1G64940.1 |
| 148 | HORVU4Hr1G087630 | AT2G30490.1 |
| 149 | HORVU7Hr1G033870 | AT1G27730.1 |
| 150 | HORVU1Hr1G061140 | AT1G15690.1 |
| 151 | HORVU5Hr1G057800 | AT4G39830.4 |
| 152 | HORVU1Hr1G014450 | AT5G23430.1 |
| 153 | HORVU2Hr1G114450 | AT2G20340.1 |
| 154 | HORVU2Hr1G112060 | AT1G49560.1 |
| 155 | HORVU6Hr1G089590 | AT2G35930.1 |
| 156 | HORVU3Hr1G018740 | AT5G61520.1 |
| 157 | HORVU3Hr1G087020 | AT2G39420.1 |
| 158 | HORVU6Hr1G073510 | AT4G13420.1 |
| 159 | HORVU5Hr1G083340 | AT3G07130.1 |
| 160 | HORVU4Hr1G064070 | AT5G37490.1 |
| 161 | HORVU7Hr1G110840 | AT2G39210.1 |

| | | |
|---|---|---|
| 162 | HORVU1Hr1G015090 | AT5G53970.1 |
| 163 | HORVU7Hr1G107360 | AT1G56150.1 |
| 164 | HORVU7Hr1G107640 | AT1G14600.1 |
| 165 | HORVU7Hr1G118770 | AT5G42500.1 |
| 166 | HORVU2Hr1G067710 | AT4G34480.1 |
| 167 | HORVU6Hr1G070130 | AT2G43420.1 |
| 168 | HORVU7Hr1G073340 | AT2G32300.2 |
| 169 | HORVU7Hr1G091860 | AT2G20340.1 |
| 170 | HORVU6Hr1G038470 | AT2G25735.1 |
| 171 | HORVU2Hr1G111780 | AT4G32300.1 |
| 172 | HORVU3Hr1G086840 | AT2G28710.1 |
| 173 | HORVU5Hr1G061580 | AT2G20840.1 |
| 174 | HORVU3Hr1G094370 | AT4G19975.1 |
| 175 | HORVU2Hr1G101150 | AT5G57540.1 |
| 176 | HORVU2Hr1G044340 | AT5G05340.1 |
| 177 | HORVU3Hr1G011870 | AT5G57260.1 |
| 178 | HORVU3Hr1G020140 | AT5G26170.1 |
| 179 | HORVU2Hr1G088660 | AT4G26740.1 |
| 180 | HORVU2Hr1G109440 | AT4G36220.1 |
| 181 | HORVU3Hr1G006130 | AT3G26430.1 |
| 182 | HORVU4Hr1G079620 | AT2G26170.1 |
| 183 | HORVU1Hr1G002590 | AT1G66920.2 |
| 184 | HORVU7Hr1G081700 | AT3G02600.9 |
| 185 | HORVU7Hr1G040740 | AT5G57625.1 |
| 186 | HORVU3Hr1G111620 | AT5G65090.3 |
| 187 | HORVU7Hr1G010850 | AT2G20340.1 |
| 188 | HORVU5Hr1G115750 | AT1G14870.1 |
| 189 | HORVU5Hr1G077920 | AT1G02850.2 |
| 190 | HORVU3Hr1G099550 | AT1G03230.1 |
| 191 | HORVU2Hr1G003230 | AT3G14470.2 |
| 192 | HORVU4Hr1G053150 | AT3G50170.1 |
| 193 | HORVU2Hr1G044360 | AT5G05340.1 |
| 194 | HORVU5Hr1G094710 | AT1G58170.1 |
| 195 | HORVU3Hr1G002900 | AT1G70520.1 |
| 196 | HORVU7Hr1G019390 | AT2G37710.1 |
| 197 | HORVU1Hr1G065650 | AT2G05635.2 |
| 198 | HORVU1Hr1G065150 | AT2G43550.1 |
| 199 | HORVU1Hr1G058520 | AT1G02850.5 |
| 200 | HORVU0Hr1G019300 | AT3G09270.1 |
| 201 | HORVU0Hr1G021480 | AT5G23850.1 |
| 202 | HORVU4Hr1G074080 | AT2G30620.2 |

| | | |
|---|---|---|
| 203 | HORVU4Hr1G087840 | AT5G17210.1 |
| 204 | HORVU3Hr1G006440 | AT1G03230.1 |
| 205 | HORVU2Hr1G067700 | AT2G19820.1 |
| 206 | HORVU0Hr1G000840 | AT5G47910.1 |
| 207 | HORVU5Hr1G099470 | AT2G43000.1 |
| 208 | HORVU5Hr1G000910 | AT4G32300.1 |
| 209 | HORVU3Hr1G096280 | AT1G01630.1 |
| 210 | HORVU3Hr1G099530 | AT3G11340.1 |
| 211 | HORVU5Hr1G109190 | AT1G18980.1 |
| 212 | HORVU4Hr1G068530 | AT3G57450.1 |
| 213 | HORVU6Hr1G067980 | AT2G02040.1 |
| 214 | HORVU3Hr1G026330 | AT4G10540.1 |
| 215 | HORVU6Hr1G094790 | AT3G21630.1 |
| 216 | HORVU3Hr1G006140 | AT3G26430.1 |
| 217 | HORVU7Hr1G092510 | AT1G15520.1 |
| 218 | HORVU6Hr1G073990 | AT2G32300.2 |
| 219 | HORVU7Hr1G057650 | AT2G17730.1 |
| 220 | HORVU6Hr1G060500 | AT3G48890.1 |
| 221 | HORVU3Hr1G023230 | AT3G53150.1 |
| 222 | HORVU7Hr1G106760 | AT5G16610.2 |
| 223 | HORVU1Hr1G075640 | AT3G47730.1 |
| 224 | HORVU1Hr1G090860 | AT3G62650.2 |
| 225 | HORVU2Hr1G034670 | AT2G14960.1 |
| 226 | HORVU1Hr1G008360 | AT3G09220.1 |
| 227 | HORVU2Hr1G079580 | AT2G14095.1 |
| 228 | HORVU2Hr1G096380 | AT5G65780.1 |
| 229 | HORVU1Hr1G062030 | AT3G12500.1 |
| 230 | HORVU5Hr1G112610 | AT3G59490.3 |
| 231 | HORVU5Hr1G045640 | AT5G53950.1 |
| 232 | HORVU7Hr1G021260 | AT5G15130.1 |
| 233 | HORVU6Hr1G074370 | AT1G44970.1 |
| 234 | HORVU1Hr1G090460 | AT1G78410.2 |
| 235 | HORVU3Hr1G065320 | AT2G47000.7 |
| 236 | HORVU2Hr1G018430 | AT5G05340.1 |
| 237 | HORVU4Hr1G056890 | AT1G07900.1 |
| 238 | HORVU4Hr1G020230 | AT3G56810.1 |
| 239 | HORVU7Hr1G032740 | AT3G16670.1 |
| 240 | HORVU2Hr1G063690 | AT3G23550.1 |
| 241 | HORVU1Hr1G085240 | AT5G11730.1 |
| 242 | HORVU3Hr1G001950 | AT4G39970.1 |
| 243 | HORVU3Hr1G002910 | AT1G56120.3 |

| | | |
|---|---|---|
| 244 | HORVU3Hr1G024820 | AT5G06570.4 |
| 245 | HORVU1Hr1G080680 | AT3G48520.1 |
| 246 | HORVU4Hr1G051450 | AT5G04930.1 |
| 247 | HORVU3Hr1G064470 | AT3G12500.1 |
| 248 | HORVU5Hr1G124810 | AT4G23230.1 |
| 249 | HORVU2Hr1G091190 | AT2G20780.1 |
| 250 | HORVU4Hr1G051290 | AT1G69490.1 |
| 251 | HORVU3Hr1G090170 | AT5G06060.1 |
| 252 | HORVU3Hr1G011990 | AT5G39100.1 |
| 253 | HORVU2Hr1G013170 | AT3G55410.2 |
| 254 | HORVU4Hr1G056500 | AT3G11180.1 |
| 255 | HORVU3Hr1G057730 | AT2G01400.1 |
| 256 | HORVU2Hr1G112580 | AT4G21585.4 |
| 257 | HORVU3Hr1G022780 | AT5G02070.1 |
| 258 | HORVU3Hr1G024660 | AT5G06570.4 |
| 259 | HORVU7Hr1G092800 | AT5G38530.1 |
| 260 | HORVU3Hr1G028020 | AT1G28570.2 |
| 261 | HORVU2Hr1G103180 | AT4G18360.4 |
| 262 | HORVU5Hr1G063270 | AT1G07520.1 |
| 263 | HORVU7Hr1G080950 | AT4G01250.1 |
| 264 | HORVU2Hr1G018510 | AT5G05340.1 |
| 265 | HORVU4Hr1G074780 | AT5G39530.1 |
| 266 | HORVU6Hr1G069980 | AT5G06740.1 |
| 267 | HORVU3Hr1G089900 | AT3G32090.1 |
| 268 | HORVU7Hr1G084920 | AT1G14430.1 |
| 269 | HORVU5Hr1G063430 | AT2G29120.1 |
| 270 | HORVU3Hr1G018610 | AT3G53810.1 |
| 271 | HORVU4Hr1G005440 | AT1G18980.1 |
| 272 | HORVU7Hr1G075760 | AT2G46750.1 |
| 273 | HORVU5Hr1G066250 | AT2G25940.1 |
| 274 | HORVU3Hr1G098340 | AT2G23620.1 |
| 275 | HORVU3Hr1G056100 | AT3G14680.1 |
| 276 | HORVU7Hr1G084610 | AT2G26640.1 |
| 277 | HORVU3Hr1G113620 | AT3G04720.1 |
| 278 | HORVU4Hr1G084930 | AT2G02370.3 |
| 279 | HORVU2Hr1G075970 | AT4G08390.3 |
| 280 | HORVU7Hr1G114660 | AT2G04400.1 |
| 281 | HORVU1Hr1G019500 | AT2G35840.4 |
| 282 | HORVU2Hr1G101980 | AT3G22370.1 |
| 283 | HORVU3Hr1G053350 | AT1G15520.1 |
| 284 | HORVU7Hr1G101310 | AT5G04220.2 |

| | | |
|---|---|---|
| 285 | HORVU5Hr1G043460 | AT3G10190.1 |
| 286 | HORVU3Hr1G080100 | AT5G63790.1 |
| 287 | HORVU7Hr1G024350 | AT4G23850.1 |
| 288 | HORVU1Hr1G009130 | AT2G18690.1 |
| 289 | HORVU5Hr1G052150 | AT5G48220.1 |
| 290 | HORVU5Hr1G000940 | AT5G10530.1 |
| 291 | HORVU2Hr1G085260 | AT1G59740.1 |
| 292 | HORVU2Hr1G028780 | AT2G38900.2 |
| 293 | HORVU1Hr1G065660 | AT4G19140.1 |
| 294 | HORVU3Hr1G104990 | #N/A |
| 295 | HORVU1Hr1G051580 | AT1G63830.3 |
| 296 | HORVU1Hr1G015210 | AT4G08850.1 |
| 297 | HORVU1Hr1G019750 | AT3G09220.1 |
| 298 | HORVU2Hr1G090170 | AT2G36790.1 |
| 299 | HORVU0Hr1G005420 | AT3G22360.1 |
| 300 | HORVU4Hr1G028780 | AT5G49620.1 |
| 301 | HORVU3Hr1G068700 | AT1G10700.1 |
| 302 | HORVU3Hr1G065390 | AT2G47000.6 |
| 303 | HORVU2Hr1G018480 | AT5G05340.1 |
| 304 | HORVU2Hr1G033610 | AT4G27250.1 |
| 305 | HORVU5Hr1G062090 | AT1G54100.2 |
| 306 | HORVU5Hr1G087240 | AT4G21390.2 |
| 307 | HORVU5Hr1G060620 | AT5G16940.2 |
| 308 | HORVU7Hr1G013170 | AT1G21430.1 |
| 309 | HORVU5Hr1G095130 | AT5G04885.2 |
| 310 | HORVU3Hr1G028620 | AT2G34500.1 |
| 311 | HORVU6Hr1G053700 | AT3G27473.1 |
| 312 | HORVU1Hr1G038780 | AT1G07040.1 |
| 313 | HORVU3Hr1G006760 | AT3G14460.1 |
| 314 | HORVU7Hr1G038370 | AT1G26810.2 |
| 315 | HORVU4Hr1G004000 | AT2G39210.1 |
| 316 | HORVU6Hr1G067920 | AT1G62200.1 |
| 317 | HORVU3Hr1G117910 | AT5G05350.1 |
| 318 | HORVU5Hr1G094700 | AT1G58170.1 |
| 319 | HORVU7Hr1G106600 | AT3G54950.1 |
| 320 | HORVU3Hr1G077950 | AT4G10540.1 |
| 321 | HORVU4Hr1G085820 | AT4G23010.1 |
| 322 | HORVU2Hr1G125250 | AT4G23180.3 |
| 323 | HORVU2Hr1G007570 | AT2G41380.1 |
| 324 | HORVU6Hr1G074690 | AT1G34300.1 |
| 325 | HORVU3Hr1G002000 | AT5G38960.1 |

| | | |
|---|---|---|
| 326 | HORVU0Hr1G030830 | AT3G56400.1 |
| 327 | HORVU4Hr1G005380 | AT1G18980.1 |
| 328 | HORVU2Hr1G100360 | AT2G24400.1 |
| 329 | HORVU5Hr1G115870 | AT1G14870.1 |
| 330 | HORVU2Hr1G114590 | AT1G05500.1 |
| 331 | HORVU2Hr1G101920 | AT3G22370.1 |
| 332 | HORVU5Hr1G104580 | AT2G15480.2 |
| 333 | HORVU6Hr1G031640 | AT4G39720.4 |
| 334 | HORVU2Hr1G008220 | AT4G21380.2 |
| 335 | HORVU2Hr1G006100 | AT3G53810.1 |
| 336 | HORVU2Hr1G110230 | AT1G78290.3 |
| 337 | HORVU5Hr1G118750 | AT1G34370.2 |
| 338 | HORVU4Hr1G071330 | AT3G59070.1 |
| 339 | HORVU7Hr1G029460 | AT2G17780.4 |
| 340 | HORVU1Hr1G027700 | AT5G26170.1 |
| 341 | HORVU4Hr1G001040 | AT2G34930.1 |
| 342 | HORVU3Hr1G067510 | AT5G26170.1 |
| 343 | HORVU3Hr1G072210 | AT3G63150.1 |
| 344 | HORVU2Hr1G100090 | AT3G13050.4 |
| 345 | HORVU7Hr1G003630 | AT4G21390.1 |
| 346 | HORVU3Hr1G111550 | AT5G43210.1 |
| 347 | HORVU1Hr1G089310 | AT3G48140.1 |
| 348 | HORVU0Hr1G005300 | AT5G24530.1 |
| 349 | HORVU2Hr1G093020 | AT5G08780.1 |
| 350 | HORVU5Hr1G072740 | #N/A |
| 351 | HORVU5Hr1G096210 | AT3G02100.1 |
| 352 | HORVU7Hr1G028710 | AT5G20400.1 |
| 353 | HORVU6Hr1G008640 | AT1G20630.1 |
| 354 | HORVU2Hr1G028330 | AT5G04990.1 |
| 355 | HORVU5Hr1G052760 | AT3G20180.3 |
| 356 | HORVU2Hr1G016720 | AT1G26970.1 |
| 357 | HORVU4Hr1G061120 | AT5G05730.1 |
| 358 | HORVU0Hr1G006150 | AT5G38240.2 |
| 359 | HORVU6Hr1G083050 | AT4G33150.3 |
| 360 | HORVU1Hr1G041720 | AT2G28315.1 |
| 361 | HORVU1Hr1G052430 | AT3G12500.1 |
| 362 | HORVU1Hr1G039720 | AT5G46050.1 |
| 363 | HORVU3Hr1G013380 | AT4G29990.2 |
| 364 | HORVU2Hr1G010560 | AT1G20225.1 |
| 365 | HORVU7Hr1G116650 | AT3G62760.1 |
| 366 | HORVU2Hr1G076470 | AT5G64660.1 |

| | | |
|---|---|---|
| 367 | HORVU3Hr1G117480 | AT4G16765.6 |
| 368 | HORVU3Hr1G108360 | AT3G54470.1 |
| 369 | HORVU2Hr1G028430 | AT2G02990.1 |
| 370 | HORVU4Hr1G076520 | AT2G05940.1 |
| 371 | HORVU3Hr1G018690 | AT2G37710.1 |
| 372 | HORVU2Hr1G004720 | AT1G22400.1 |
| 373 | HORVU5Hr1G063480 | AT2G29120.1 |
| 374 | HORVU5Hr1G122410 | AT1G69850.1 |
| 375 | HORVU3Hr1G093100 | AT5G55410.1 |
| 376 | HORVU3Hr1G071470 | AT1G15520.1 |
| 377 | HORVU3Hr1G105560 | AT4G16260.1 |
| 378 | HORVU6Hr1G084080 | AT3G23560.1 |
| 379 | HORVU7Hr1G007520 | AT5G44700.1 |
| 380 | HORVU4Hr1G052320 | AT3G56710.2 |
| 381 | HORVU5Hr1G045850 | AT2G29420.1 |
| 382 | HORVU6Hr1G077120 | AT5G37490.1 |
| 383 | HORVU2Hr1G045200 | AT1G10370.1 |
| 384 | HORVU4Hr1G067450 | AT5G26340.1 |
| 385 | HORVU2Hr1G124930 | AT1G71695.1 |
| 386 | HORVU7Hr1G121290 | AT1G19250.1 |
| 387 | HORVU5Hr1G102900 | AT5G48540.1 |
| 388 | HORVU7Hr1G002740 | AT3G09220.1 |
| 389 | HORVU7Hr1G091250 | AT4G25000.1 |
| 390 | HORVU1Hr1G063740 | AT1G01720.1 |
| 391 | HORVU2Hr1G085280 | AT3G54420.1 |
| 392 | HORVU5Hr1G010130 | AT4G16120.1 |
| 393 | HORVU5Hr1G112210 | AT3G11900.2 |
| 394 | HORVU7Hr1G028370 | AT4G25650.1 |
| 395 | HORVU6Hr1G068990 | AT2G21620.1 |
| 396 | HORVU1Hr1G021150 | AT3G62760.1 |
| 397 | HORVU4Hr1G084340 | AT5G60710.1 |
| 398 | HORVU6Hr1G005900 | AT1G47480.1 |
| 399 | HORVU3Hr1G001740 | AT5G22050.1 |
| 400 | HORVU3Hr1G085890 | AT3G13080.4 |
| 401 | HORVU1Hr1G071380 | AT1G74540.1 |
| 402 | HORVU4Hr1G071360 | AT3G25290.2 |
| 403 | HORVU5Hr1G072730 | #N/A |
| 404 | HORVU5Hr1G024400 | AT2G18750.3 |
| 405 | HORVU5Hr1G114440 | AT4G35920.5 |
| 406 | HORVU1Hr1G021140 | AT3G62760.1 |
| 407 | HORVU7Hr1G042120 | AT1G68570.1 |

| | | |
|---|---|---|
| 408 | HORVU3Hr1G028260 | AT2G27940.1 |
| 409 | HORVU3Hr1G063430 | AT1G71140.1 |
| 410 | HORVU2Hr1G083570 | AT3G26330.1 |
| 411 | HORVU7Hr1G101590 | AT3G14690.1 |
| 412 | HORVU5Hr1G097270 | AT3G49960.1 |
| 413 | HORVU7Hr1G083550 | AT5G66390.1 |
| 414 | HORVU1Hr1G037150 | AT5G03320.2 |
| 415 | HORVU4Hr1G009190 | AT1G75170.2 |
| 416 | HORVU1Hr1G054390 | AT3G22400.1 |
| 417 | HORVU7Hr1G075710 | AT1G07780.9 |
| 418 | HORVU6Hr1G066250 | AT3G18420.1 |
| 419 | HORVU0Hr1G021610 | AT2G02360.1 |
| 420 | HORVU2Hr1G077220 | AT1G76690.1 |
| 421 | HORVU4Hr1G083210 | AT5G17990.1 |
| 422 | HORVU4Hr1G076840 | AT1G74950.1 |
| 423 | HORVU7Hr1G028730 | AT2G37710.1 |
| 424 | HORVU5Hr1G118590 | AT2G38290.1 |
| 425 | HORVU2Hr1G038830 | AT4G05390.1 |
| 426 | HORVU2Hr1G008130 | AT4G21380.2 |
| 427 | HORVU1Hr1G093480 | AT4G02610.1 |
| 428 | HORVU2Hr1G017950 | AT3G48140.1 |
| 429 | HORVU7Hr1G074660 | AT5G40780.1 |
| 430 | HORVU0Hr1G039970 | AT3G47090.1 |
| 431 | HORVU2Hr1G044440 | AT2G39640.1 |
| 432 | HORVU7Hr1G008140 | AT5G10530.1 |
| 433 | HORVU2Hr1G109330 | AT3G56400.1 |
| 434 | HORVU7Hr1G121260 | AT1G19250.1 |
| 435 | HORVU6Hr1G057270 | AT1G71950.1 |
| 436 | HORVU3Hr1G096360 | AT3G62160.1 |
| 437 | HORVU2Hr1G080890 | AT1G25220.1 |
| 438 | HORVU3Hr1G023400 | AT2G36800.1 |
| 439 | HORVU5Hr1G062410 | AT5G62740.1 |
| 440 | HORVU5Hr1G014060 | AT3G22930.1 |
| 441 | HORVU5Hr1G098090 | AT5G42650.1 |
| 442 | HORVU2Hr1G017350 | AT5G61430.1 |
| 443 | HORVU5Hr1G010120 | AT2G23450.2 |
| 444 | HORVU1Hr1G001570 | AT4G35160.1 |
| 445 | HORVU1Hr1G081550 | AT3G47520.1 |
| 446 | HORVU0Hr1G002930 | AT2G15760.1 |
| 447 | HORVU2Hr1G044450 | AT5G55180.2 |
| 448 | HORVU7Hr1G028840 | AT5G54000.2 |

| | | |
|---|---|---|
| 449 | HORVU4Hr1G071020 | AT3G18830.1 |
| 450 | HORVU2Hr1G017470 | AT1G69490.1 |
| 451 | HORVU4Hr1G090140 | AT5G39150.1 |
| 452 | HORVU7Hr1G116310 | AT3G14660.1 |
| 453 | HORVU4Hr1G067840 | AT1G76340.1 |
| 454 | HORVU2Hr1G036590 | AT3G18830.1 |
| 455 | HORVU4Hr1G078210 | AT1G28220.1 |
| 456 | HORVU3Hr1G016860 | AT2G38870.1 |
| 457 | HORVU6Hr1G057680 | AT5G60710.1 |
| 458 | HORVU3Hr1G014270 | AT2G02340.1 |
| 459 | HORVU2Hr1G042210 | AT2G19130.1 |
| 460 | HORVU2Hr1G121060 | AT1G65840.1 |
| 461 | HORVU4Hr1G084740 | AT1G34580.1 |
| 462 | HORVU2Hr1G044270 | AT4G05200.2 |
| 463 | HORVU0Hr1G009460 | AT5G39110.1 |
| 464 | HORVU6Hr1G004770 | AT1G21230.1 |
| 465 | HORVU4Hr1G072560 | AT2G32300.2 |
| 466 | HORVU5Hr1G110180 | AT3G54700.2 |
| 467 | HORVU3Hr1G079620 | AT3G08640.1 |
| 468 | HORVU7Hr1G019280 | AT5G61520.1 |
| 469 | HORVU1Hr1G049280 | AT1G10370.1 |
| 470 | HORVU7Hr1G113350 | AT4G27250.1 |
| 471 | HORVU5Hr1G058020 | AT5G10190.2 |
| 472 | HORVU3Hr1G059480 | AT5G24090.1 |
| 473 | HORVU6Hr1G074310 | AT3G57020.2 |
| 474 | HORVU4Hr1G007180 | AT1G65820.1 |
| 475 | HORVU1Hr1G052250 | AT3G54710.1 |
| 476 | HORVU2Hr1G125600 | AT4G35920.5 |
| 477 | HORVU2Hr1G080140 | AT5G15140.1 |
| 478 | HORVU2Hr1G008140 | AT4G27300.1 |
| 479 | HORVU2Hr1G009830 | AT2G47800.1 |
| 480 | HORVU5Hr1G047060 | AT1G21600.2 |
| 481 | HORVU6Hr1G089960 | AT1G47890.1 |
| 482 | HORVU5Hr1G081860 | AT5G51070.1 |
| 483 | HORVU3Hr1G077890 | AT4G21920.1 |
| 484 | HORVU4Hr1G003400 | AT2G01820.1 |
| 485 | HORVU7Hr1G079460 | AT5G53710.2 |
| 486 | HORVU5Hr1G011540 | AT2G36690.2 |
| 487 | HORVU4Hr1G072310 | AT4G38540.1 |
| 488 | HORVU1Hr1G080320 | AT5G62530.1 |
| 489 | HORVU3Hr1G090220 | AT2G19130.1 |

| | | |
|---|---|---|
| 490 | HORVU6Hr1G024850 | AT3G48310.1 |
| 491 | HORVU5Hr1G063420 | AT2G29120.1 |
| 492 | HORVU5Hr1G012040 | AT5G65600.1 |
| 493 | HORVU5Hr1G065620 | AT5G20400.1 |
| 494 | HORVU2Hr1G109590 | AT5G48930.1 |
| 495 | HORVU7Hr1G019240 | AT5G61520.2 |
| 496 | HORVU3Hr1G099630 | AT3G20980.1 |
| 497 | HORVU7Hr1G118560 | AT3G17675.1 |
| 498 | HORVU4Hr1G000850 | AT5G23950.2 |
| 499 | HORVU6Hr1G093180 | AT5G43430.1 |
| 500 | HORVU2Hr1G027480 | AT2G46950.1 |
| 501 | HORVU5Hr1G024160 | AT2G47710.1 |
| 502 | HORVU6Hr1G021320 | AT2G47000.7 |
| 503 | HORVU1Hr1G016840 | AT1G05260.1 |
| 504 | HORVU1Hr1G028200 | AT4G29130.1 |
| 505 | HORVU5Hr1G071140 | AT1G55850.1 |
| 506 | HORVU1Hr1G000140 | AT1G68710.2 |
| 507 | HORVU5Hr1G071720 | AT5G04690.1 |
| 508 | HORVU7Hr1G091100 | AT2G05790.1 |
| 509 | HORVU4Hr1G083650 | AT5G46050.1 |
| 510 | HORVU1Hr1G089830 | AT1G01800.1 |
| 511 | HORVU5Hr1G063340 | AT5G57480.1 |
| 512 | HORVU2Hr1G121440 | AT5G43060.1 |
| 513 | HORVU2Hr1G062730 | AT1G27040.2 |
| 514 | HORVU7Hr1G024930 | AT4G10540.1 |
| 515 | HORVU0Hr1G005160 | AT1G08630.6 |
| 516 | HORVU6Hr1G005400 | AT3G51000.1 |
| 517 | HORVU5Hr1G117910 | AT2G27510.1 |
| 518 | HORVU7Hr1G079630 | AT2G33600.1 |
| 519 | HORVU5Hr1G120570 | AT1G24490.1 |
| 520 | HORVU4Hr1G078690 | AT2G44610.1 |
| 521 | HORVU7Hr1G090830 | AT5G14420.4 |
| 522 | HORVU5Hr1G089780 | AT1G30910.1 |
| 523 | HORVU5Hr1G001920 | AT1G09970.1 |
| 524 | HORVU2Hr1G117840 | AT4G21390.1 |
| 525 | HORVU1Hr1G080300 | AT5G64810.1 |
| 526 | HORVU7Hr1G028750 | AT3G06240.1 |
| 527 | HORVU3Hr1G028060 | AT5G45910.1 |
| 528 | HORVU5Hr1G012120 | AT5G10530.1 |
| 529 | HORVU7Hr1G037910 | AT2G21940.5 |
| 530 | HORVU6Hr1G011120 | AT4G19880.1 |

| | | |
|---|---|---|
| 531 | HORVU4Hr1G023390 | AT3G48860.2 |
| 532 | HORVU7Hr1G074640 | AT5G40780.1 |
| 533 | HORVU5Hr1G096930 | AT4G37330.1 |
| 534 | HORVU5Hr1G087880 | AT5G65980.1 |
| 535 | HORVU1Hr1G016660 | AT4G33420.1 |
| 536 | HORVU0Hr1G009400 | AT5G39110.1 |
| 537 | HORVU2Hr1G004230 | AT1G76680.1 |
| 538 | HORVU2Hr1G064160 | AT1G11260.1 |
| 539 | HORVU5Hr1G077010 | AT2G33480.2 |
| 540 | HORVU7Hr1G073460 | AT4G27070.1 |
| 541 | HORVU2Hr1G096510 | AT5G20400.1 |
| 542 | HORVU1Hr1G049210 | AT1G10360.1 |
| 543 | HORVU2Hr1G123590 | AT1G21210.1 |
| 544 | HORVU3Hr1G093550 | AT1G56130.2 |
| 545 | HORVU3Hr1G077110 | AT4G21390.1 |
| 546 | HORVU0Hr1G009470 | AT5G39110.1 |
| 547 | HORVU1Hr1G057460 | AT5G64250.1 |
| 548 | HORVU2Hr1G036800 | AT4G36810.1 |
| 549 | HORVU0Hr1G007680 | AT1G47510.1 |
| 550 | HORVU2Hr1G014890 | AT3G53810.1 |
| 551 | HORVU2Hr1G087090 | AT1G80050.1 |
| 552 | HORVU7Hr1G073050 | AT4G21490.2 |
| 553 | HORVU3Hr1G076750 | AT1G61250.1 |
| 554 | HORVU1Hr1G019410 | AT5G48930.1 |
| 555 | HORVU5Hr1G040740 | AT5G27600.1 |
| 556 | HORVU1Hr1G045610 | AT5G52240.1 |
| 557 | HORVU5Hr1G115230 | AT1G07230.1 |
| 558 | HORVU2Hr1G085270 | AT3G54420.1 |
| 559 | HORVU2Hr1G117940 | AT1G01490.2 |
| 560 | HORVU1Hr1G049100 | AT1G10360.1 |
| 561 | HORVU3Hr1G087740 | AT3G55990.1 |
| 562 | HORVU4Hr1G005450 | AT5G19860.1 |
| 563 | HORVU6Hr1G090600 | AT1G10370.1 |
| 564 | HORVU2Hr1G078890 | AT4G37030.1 |
| 565 | HORVU3Hr1G108900 | AT5G56170.1 |
| 566 | HORVU4Hr1G087340 | AT2G35330.2 |
| 567 | HORVU3Hr1G031020 | AT3G48320.1 |
| 568 | HORVU6Hr1G084740 | AT1G53030.1 |
| 569 | HORVU3Hr1G002920 | AT3G18180.1 |
| 570 | HORVU7Hr1G010250 | AT1G01820.1 |
| 571 | HORVU5Hr1G089620 | AT1G21240.1 |

| | | |
|---|---|---|
| 572 | HORVU1Hr1G038680 | AT3G23990.1 |
| 573 | HORVU7Hr1G121950 | AT4G29090.1 |
| 574 | HORVU1Hr1G072720 | AT5G02090.1 |
| 575 | HORVU7Hr1G037510 | AT4G08550.2 |
| 576 | HORVU2Hr1G048220 | AT4G21570.1 |
| 577 | HORVU5Hr1G009930 | AT5G37660.1 |
| 578 | HORVU6Hr1G084590 | AT1G79690.2 |
| 579 | HORVU5Hr1G096940 | AT4G37330.1 |
| 580 | HORVU7Hr1G108820 | AT2G18680.1 |
| 581 | HORVU6Hr1G018460 | AT1G48120.1 |
| 582 | HORVU7Hr1G050670 | AT3G45140.1 |
| 583 | HORVU2Hr1G113180 | AT4G34200.1 |
| 584 | HORVU3Hr1G105290 | AT3G57240.1 |
| 585 | HORVU5Hr1G023990 | AT1G67120.1 |
| 586 | HORVU4Hr1G072880 | AT4G35630.1 |
| 587 | HORVU1Hr1G084720 | AT3G20630.1 |
| 588 | HORVU6Hr1G093740 | AT1G21230.1 |
| 589 | HORVU2Hr1G064150 | AT1G11260.1 |
| 590 | HORVU7Hr1G051070 | AT4G38220.1 |
| 591 | HORVU7Hr1G099340 | AT4G23280.2 |
| 592 | HORVU5Hr1G050940 | AT1G15950.3 |
| 593 | HORVU5Hr1G099820 | AT3G58750.1 |
| 594 | HORVU4Hr1G054060 | AT5G10920.1 |
| 595 | HORVU1Hr1G045580 | AT1G55020.1 |
| 596 | HORVU4Hr1G076570 | AT1G72520.1 |
| 597 | HORVU3Hr1G024210 | AT3G13080.5 |
| 598 | HORVU4Hr1G081940 | AT1G20510.1 |
| 599 | HORVU5Hr1G097940 | AT2G15960.1 |
| 600 | HORVU1Hr1G069800 | AT5G01870.1 |
| 601 | HORVU2Hr1G086380 | AT5G48930.1 |
| 602 | HORVU3Hr1G087680 | AT5G58560.1 |
| 603 | HORVU2Hr1G017400 | AT5G39610.1 |
| 604 | HORVU6Hr1G065210 | AT2G27230.2 |
| 605 | HORVU1Hr1G073040 | AT3G54020.1 |
| 606 | HORVU2Hr1G039850 | AT1G15170.2 |
| 607 | HORVU5Hr1G076250 | AT5G50400.1 |
| 608 | HORVU1Hr1G011430 | AT2G36310.1 |
| 609 | HORVU3Hr1G073220 | AT5G11520.1 |
| 610 | HORVU5Hr1G106550 | AT2G30620.2 |
| 611 | HORVU3Hr1G070550 | AT5G19740.1 |
| 612 | HORVU3Hr1G096050 | AT5G13420.1 |

| | | |
|---|---|---|
| 613 | HORVU7Hr1G017950 | AT2G45560.1 |
| 614 | HORVU6Hr1G013750 | AT2G19130.1 |
| 615 | HORVU7Hr1G007540 | AT1G09070.1 |
| 616 | HORVU6Hr1G071780 | AT4G12650.1 |
| 617 | HORVU3Hr1G099620 | AT3G20980.1 |
| 618 | HORVU5Hr1G079690 | AT5G48370.2 |
| 619 | HORVU6Hr1G090560 | AT1G10360.1 |
| 620 | HORVU2Hr1G114260 | AT1G80460.1 |
| 621 | HORVU2Hr1G022620 | AT1G50180.1 |
| 622 | HORVU3Hr1G083560 | AT2G26230.1 |
| 623 | HORVU1Hr1G092300 | AT4G23190.2 |
| 624 | HORVU2Hr1G091270 | AT5G10560.1 |
| 625 | HORVU3Hr1G018650 | AT5G01320.1 |
| 626 | HORVU6Hr1G006560 | AT1G55510.1 |
| 627 | HORVU3Hr1G096190 | AT1G21230.1 |
| 628 | HORVU3Hr1G090070 | AT1G20400.1 |
| 629 | HORVU6Hr1G081000 | AT3G25780.1 |
| 630 | HORVU2Hr1G028540 | AT1G55280.1 |
| 631 | HORVU2Hr1G092820 | AT5G24380.1 |
| 632 | HORVU2Hr1G037750 | AT4G23180.1 |
| 633 | HORVU1Hr1G002470 | AT1G70250.1 |
| 634 | HORVU3Hr1G093400 | AT3G23830.2 |

SUPPLEMENTARY TABLE 9

| S No | 376 "Bowman_Down" and "Harrington_Down": | |
|---|---|---|
| 1 | HORVU4Hr1G075160 | AT1G14345.1 |
| 2 | HORVU5Hr1G056390 | AT5G66530.3 |
| 3 | HORVU3Hr1G030540 | AT2G24490.2 |
| 4 | HORVU3Hr1G096650 | AT5G43750.1 |
| 5 | HORVU5Hr1G046250 | AT5G01980.1 |
| 6 | HORVU6Hr1G010520 | AT3G50685.1 |
| 7 | HORVU2Hr1G013870 | AT5G05580.1 |
| 8 | HORVU5Hr1G112670 | AT3G55120.1 |
| 9 | HORVU0Hr1G031970 | AT3G05700.1 |
| 10 | HORVU2Hr1G092280 | AT4G37925.1 |
| 11 | HORVU0Hr1G004830 | AT1G12900.5 |
| 12 | HORVU7Hr1G120190 | AT5G23120.2 |
| 13 | HORVU7Hr1G064100 | AT2G05620.2 |
| 14 | HORVU3Hr1G047030 | AT2G04700.3 |
| 15 | HORVU3Hr1G062490 | AT3G15190.1 |

| | | |
|---|---|---|
| 16 | HORVU1Hr1G070190 | AT2G30300.1 |
| 17 | HORVU4Hr1G060970 | AT1G07320.2 |
| 18 | HORVU5Hr1G046480 | AT5G05270.2 |
| 19 | HORVU2Hr1G101040 | AT5G57110.3 |
| 20 | HORVU6Hr1G093030 | AT3G44890.1 |
| 21 | HORVU7Hr1G041080 | AT3G10130.1 |
| 22 | HORVU7Hr1G058940 | AT4G24770.2 |
| 23 | HORVU1Hr1G000430 | AT1G53520.1 |
| 24 | HORVU5Hr1G050120 | AT3G53210.2 |
| 25 | HORVU5Hr1G120250 | AT1G64510.1 |
| 26 | HORVU1Hr1G088270 | AT5G01930.1 |
| 27 | HORVU4Hr1G016510 | AT4G09350.1 |
| 28 | HORVU4Hr1G025010 | #N/A |
| 29 | HORVU6Hr1G049050 | AT5G64940.2 |
| 30 | HORVU2Hr1G091680 | AT5G23750.3 |
| 31 | HORVU6Hr1G085170 | AT1G79040.1 |
| 32 | HORVU6Hr1G075900 | AT1G74470.1 |
| 33 | HORVU7Hr1G038300 | AT3G28960.5 |
| 34 | HORVU2Hr1G112450 | AT3G52810.1 |
| 35 | HORVU2Hr1G115710 | AT2G21960.1 |
| 36 | HORVU3Hr1G084660 | AT5G01260.3 |
| 37 | HORVU4Hr1G074810 | AT5G61410.2 |
| 38 | HORVU2Hr1G119760 | AT1G50270.1 |
| 39 | HORVU6Hr1G015430 | AT5G17670.1 |
| 40 | HORVU1Hr1G061830 | AT1G54780.1 |
| 41 | HORVU5Hr1G100990 | AT1G56050.1 |
| 42 | HORVU7Hr1G115040 | AT1G79850.1 |
| 43 | HORVU7Hr1G079190 | AT2G38140.1 |
| 44 | HORVU3Hr1G010030 | AT5G13240.1 |
| 45 | HORVU7Hr1G114170 | AT2G24090.1 |
| 46 | HORVU6Hr1G033970 | AT3G21340.2 |
| 47 | HORVU2Hr1G005570 | AT1G06620.1 |
| 48 | HORVU5Hr1G001650 | AT5G17210.1 |
| 49 | HORVU7Hr1G002210 | AT1G20020.3 |
| 50 | HORVU2Hr1G057700 | AT3G50820.1 |
| 51 | HORVU4Hr1G060440 | AT4G00550.2 |
| 52 | HORVU7Hr1G097670 | AT5G57170.1 |
| 53 | HORVU6Hr1G055960 | AT4G10850.1 |
| 54 | HORVU7Hr1G060130 | AT2G28190.1 |
| 55 | HORVU4Hr1G071730 | AT2G22240.2 |
| 56 | HORVU6Hr1G054110 | AT1G67700.4 |

| | | |
|---|---|---|
| 57 | HORVU2Hr1G060880 | AT1G06680.1 |
| 58 | HORVU1Hr1G050130 | AT5G46800.3 |
| 59 | HORVU5Hr1G065330 | AT1G15950.3 |
| 60 | HORVU5Hr1G114990 | AT2G13650.3 |
| 61 | HORVU4Hr1G010430 | AT2G23760.3 |
| 62 | HORVU3Hr1G018110 | AT2G30170.1 |
| 63 | HORVU3Hr1G032490 | AT5G46110.4 |
| 64 | HORVU5Hr1G053270 | AT2G42220.1 |
| 65 | HORVU3Hr1G049900 | AT1G17650.1 |
| 66 | HORVU6Hr1G018830 | AT2G43030.1 |
| 67 | HORVU1Hr1G013250 | AT1G23450.1 |
| 68 | HORVU6Hr1G091260 | AT2G33310.2 |
| 69 | HORVU3Hr1G006150 | AT1G18060.1 |
| 70 | HORVU5Hr1G009200 | AT1G09340.1 |
| 71 | HORVU4Hr1G085050 | AT5G50400.1 |
| 72 | HORVU4Hr1G062920 | AT1G75690.1 |
| 73 | HORVU4Hr1G004210 | AT1G32550.1 |
| 74 | HORVU4Hr1G089990 | AT2G26570.2 |
| 75 | HORVU4Hr1G004590 | AT5G15070.4 |
| 76 | HORVU7Hr1G062930 | AT3G01480.1 |
| 77 | HORVU1Hr1G012110 | AT2G21370.1 |
| 78 | HORVU3Hr1G082730 | AT1G44575.3 |
| 79 | HORVU4Hr1G084830 | AT1G32990.1 |
| 80 | HORVU5Hr1G025960 | AT4G25050.1 |
| 81 | HORVU2Hr1G105930 | AT2G35410.1 |
| 82 | HORVU6Hr1G078500 | AT2G03220.1 |
| 83 | HORVU6Hr1G090160 | AT5G40810.2 |
| 84 | HORVU3Hr1G085270 | AT3G55800.1 |
| 85 | HORVU6Hr1G073180 | AT3G18000.1 |
| 86 | HORVU6Hr1G075280 | AT4G09650.1 |
| 87 | HORVU4Hr1G087350 | AT3G61320.1 |
| 88 | HORVU4Hr1G006730 | AT3G26070.1 |
| 89 | HORVU5Hr1G111640 | AT3G54210.1 |
| 90 | HORVU7Hr1G055890 | AT5G17870.1 |
| 91 | HORVU4Hr1G085930 | AT3G61080.1 |
| 92 | HORVU7Hr1G026100 | AT3G54920.1 |
| 93 | HORVU3Hr1G083790 | AT2G36145.1 |
| 94 | HORVU7Hr1G101810 | AT5G54600.1 |
| 95 | HORVU2Hr1G009850 | AT5G52970.2 |
| 96 | HORVU3Hr1G082280 | AT2G16570.1 |
| 97 | HORVU2Hr1G103130 | AT1G11860.2 |

| | | |
|---|---|---|
| 98 | HORVU2Hr1G041610 | AT4G03280.1 |
| 99 | HORVU3Hr1G093990 | AT4G12830.1 |
| 100 | HORVU7Hr1G037600 | AT1G53210.1 |
| 101 | HORVU1Hr1G072180 | AT5G59750.1 |
| 102 | HORVU5Hr1G022570 | AT2G32500.2 |
| 103 | HORVU6Hr1G004860 | AT3G63140.1 |
| 104 | HORVU1Hr1G087570 | AT2G20520.1 |
| 105 | HORVU7Hr1G122750 | AT1G16080.1 |
| 106 | HORVU6Hr1G009170 | AT1G72640.1 |
| 107 | HORVU3Hr1G095700 | AT1G02170.1 |
| 108 | HORVU3Hr1G039790 | AT3G56910.1 |
| 109 | HORVU3Hr1G025020 | AT5G19875.1 |
| 110 | HORVU2Hr1G090400 | AT3G19800.2 |
| 111 | HORVU3Hr1G050430 | AT5G12000.1 |
| 112 | HORVU7Hr1G042480 | AT3G18440.1 |
| 113 | HORVU1Hr1G060610 | AT4G15560.1 |
| 114 | HORVU5Hr1G118680 | AT4G09620.2 |
| 115 | HORVU7Hr1G000900 | AT1G76100.1 |
| 116 | HORVU7Hr1G039140 | AT3G62760.1 |
| 117 | HORVU3Hr1G036600 | AT1G35180.1 |
| 118 | HORVU5Hr1G043090 | AT2G41310.1 |
| 119 | HORVU1Hr1G051700 | AT2G46820.2 |
| 120 | HORVU0Hr1G001170 | AT3G51510.1 |
| 121 | HORVU7Hr1G077740 | AT1G60950.1 |
| 122 | HORVU2Hr1G113050 | AT1G23090.1 |
| 123 | HORVU2Hr1G037550 | AT5G19780.1 |
| 124 | HORVU4Hr1G031320 | AT5G49810.1 |
| 125 | HORVU6Hr1G076040 | AT1G18730.5 |
| 126 | HORVU5Hr1G111330 | AT4G13500.1 |
| 127 | HORVU2Hr1G039020 | AT2G29750.1 |
| 128 | HORVU1Hr1G013740 | AT1G48030.5 |
| 129 | HORVU4Hr1G067320 | AT3G24340.1 |
| 130 | HORVU7Hr1G073860 | AT1G30520.2 |
| 131 | HORVU7Hr1G085980 | AT2G01980.1 |
| 132 | HORVU3Hr1G073740 | AT5G55570.1 |
| 133 | HORVU1Hr1G064750 | AT5G14910.2 |
| 134 | HORVU6Hr1G011990 | AT3G61870.2 |
| 135 | HORVU3Hr1G071530 | AT1G78630.1 |
| 136 | HORVU6Hr1G025170 | #N/A |
| 137 | HORVU7Hr1G038770 | AT5G16710.1 |
| 138 | HORVU2Hr1G038570 | AT3G63190.1 |

| | | |
|---|---|---|
| 139 | HORVU7Hr1G083690 | AT2G20570.1 |
| 140 | HORVU6Hr1G032680 | AT4G26555.1 |
| 141 | HORVU6Hr1G095380 | AT2G21280.2 |
| 142 | HORVU3Hr1G065570 | #N/A |
| 143 | HORVU7Hr1G111190 | AT4G34200.1 |
| 144 | HORVU5Hr1G051010 | AT1G67090.1 |
| 145 | HORVU5Hr1G053060 | AT3G57990.1 |
| 146 | HORVU6Hr1G076880 | AT1G32470.1 |
| 147 | HORVU4Hr1G042560 | AT1G32960.1 |
| 148 | HORVU5Hr1G050510 | AT5G47820.2 |
| 149 | HORVU5Hr1G101020 | AT5G44520.2 |
| 150 | HORVU3Hr1G078780 | AT1G29070.1 |
| 151 | HORVU5Hr1G099020 | AT5G42760.2 |
| 152 | HORVU5Hr1G035300 | AT2G46620.1 |
| 153 | HORVU3Hr1G087420 | AT1G13635.2 |
| 154 | HORVU6Hr1G021460 | AT5G57180.2 |
| 155 | HORVU1Hr1G045520 | AT5G53490.2 |
| 156 | HORVU4Hr1G009980 | AT1G44920.1 |
| 157 | HORVU5Hr1G115000 | AT3G47070.1 |
| 158 | HORVU2Hr1G111300 | AT3G11560.5 |
| 159 | HORVU1Hr1G035870 | AT4G15760.1 |
| 160 | HORVU5Hr1G098340 | AT1G74970.1 |
| 161 | HORVU0Hr1G012860 | AT4G37980.1 |
| 162 | HORVU7Hr1G121210 | AT5G37260.1 |
| 163 | HORVU3Hr1G094270 | AT2G35450.1 |
| 164 | HORVU3Hr1G113920 | AT5G04895.1 |
| 165 | HORVU3Hr1G081180 | AT1G69530.2 |
| 166 | HORVU5Hr1G062240 | AT1G19150.1 |
| 167 | HORVU3Hr1G013970 | AT2G37540.1 |
| 168 | HORVU2Hr1G073760 | AT5G06290.1 |
| 169 | HORVU7Hr1G097250 | AT5G35100.3 |
| 170 | HORVU5Hr1G096260 | AT3G02100.1 |
| 171 | HORVU7Hr1G030800 | AT2G28630.2 |
| 172 | HORVU2Hr1G001500 | AT4G14500.2 |
| 173 | HORVU3Hr1G089650 | AT3G59780.1 |
| 174 | HORVU1Hr1G067300 | AT3G12780.1 |
| 175 | HORVU1Hr1G001900 | AT1G21640.1 |
| 176 | HORVU2Hr1G060480 | AT1G30380.1 |
| 177 | HORVU1Hr1G054190 | AT5G65750.1 |
| 178 | HORVU5Hr1G109440 | AT1G67280.1 |
| 179 | HORVU3Hr1G029260 | AT3G14920.1 |

| | | |
|---|---|---|
| 180 | HORVU7Hr1G042160 | AT4G01150.1 |
| 181 | HORVU1Hr1G022840 | AT1G52870.2 |
| 182 | HORVU3Hr1G095330 | AT5G40950.1 |
| 183 | HORVU7Hr1G056240 | AT2G43560.1 |
| 184 | HORVU2Hr1G080480 | AT1G68590.2 |
| 185 | HORVU3Hr1G093310 | AT3G07340.1 |
| 186 | HORVU7Hr1G096140 | AT4G02530.1 |
| 187 | HORVU5Hr1G100140 | AT1G31330.1 |
| 188 | HORVU3Hr1G091140 | AT5G58260.1 |
| 189 | HORVU1Hr1G011520 | AT1G58290.1 |
| 190 | HORVU4Hr1G083230 | AT5G44600.1 |
| 191 | HORVU2Hr1G089030 | AT4G13670.1 |
| 192 | HORVU6Hr1G010940 | AT4G16120.1 |
| 193 | HORVU5Hr1G005740 | AT5G17230.4 |
| 194 | HORVU2Hr1G106880 | AT3G47860.1 |
| 195 | HORVU7Hr1G009800 | AT1G32560.1 |
| 196 | HORVU1Hr1G023160 | AT1G19250.1 |
| 197 | HORVU3Hr1G088020 | AT5G58490.1 |
| 198 | HORVU5Hr1G014300 | AT3G04730.1 |
| 199 | HORVU1Hr1G050830 | AT2G23910.1 |
| 200 | HORVU5Hr1G060250 | AT1G68090.1 |
| 201 | HORVU6Hr1G053730 | AT1G13120.1 |
| 202 | HORVU2Hr1G116390 | AT5G13930.1 |
| 203 | HORVU5Hr1G098980 | AT1G75210.3 |
| 204 | HORVU4Hr1G073650 | AT3G22830.1 |
| 205 | HORVU2Hr1G073370 | AT1G08380.1 |
| 206 | HORVU7Hr1G117710 | AT4G35160.1 |
| 207 | HORVU7Hr1G006880 | AT3G60380.1 |
| 208 | HORVU0Hr1G021630 | AT4G29080.1 |
| 209 | HORVU1Hr1G075580 | AT5G56550.1 |
| 210 | HORVU3Hr1G000900 | AT1G74070.1 |
| 211 | HORVU2Hr1G023560 | AT5G13630.1 |
| 212 | HORVU0Hr1G001490 | AT1G52230.1 |
| 213 | HORVU5Hr1G050990 | AT1G67090.1 |
| 214 | HORVU1Hr1G004400 | AT5G02160.1 |
| 215 | HORVU2Hr1G085300 | AT4G39970.1 |
| 216 | HORVU1Hr1G005120 | AT2G28900.1 |
| 217 | HORVU4Hr1G076990 | AT5G26670.1 |
| 218 | HORVU3Hr1G039930 | AT3G56940.1 |
| 219 | HORVU3Hr1G094820 | AT3G25480.1 |
| 220 | HORVU2Hr1G074280 | AT5G09820.2 |

| | | |
|---|---|---|
| 221 | HORVU2Hr1G049560 | AT1G20810.2 |
| 222 | HORVU6Hr1G049120 | ATCG00220.1 |
| 223 | HORVU6Hr1G071520 | AT5G10770.1 |
| 224 | HORVU2Hr1G077650 | AT5G49740.3 |
| 225 | HORVU2Hr1G095080 | AT1G12740.1 |
| 226 | HORVU5Hr1G103440 | AT5G48890.2 |
| 227 | HORVU4Hr1G013420 | AT2G14820.2 |
| 228 | HORVU3Hr1G025430 | AT3G50370.1 |
| 229 | HORVU3Hr1G059950 | AT5G47490.2 |
| 230 | HORVU2Hr1G066020 | AT3G55380.2 |
| 231 | HORVU1Hr1G091540 | AT4G39250.1 |
| 232 | HORVU1Hr1G057640 | AT3G57190.1 |
| 233 | HORVU5Hr1G043200 | AT3G57030.1 |
| 234 | HORVU7Hr1G052750 | AT3G25290.2 |
| 235 | HORVU6Hr1G075080 | AT4G39710.2 |
| 236 | HORVU3Hr1G057090 | AT5G14740.9 |
| 237 | HORVU7Hr1G023140 | AT3G16250.1 |
| 238 | HORVU1Hr1G035720 | AT1G67090.1 |
| 239 | HORVU6Hr1G015120 | AT1G78080.1 |
| 240 | HORVU4Hr1G002050 | AT3G63410.1 |
| 241 | HORVU3Hr1G083360 | AT1G43670.1 |
| 242 | HORVU4Hr1G014080 | AT3G59400.1 |
| 243 | HORVU7Hr1G092880 | AT3G03750.2 |
| 244 | HORVU7Hr1G105500 | AT4G14540.1 |
| 245 | HORVU2Hr1G038580 | AT2G13980.1 |
| 246 | HORVU4Hr1G011160 | AT5G49720.1 |
| 247 | HORVU4Hr1G032020 | AT5G62630.1 |
| 248 | HORVU3Hr1G055260 | AT4G01060.1 |
| 249 | HORVU5Hr1G071920 | AT1G55670.1 |
| 250 | HORVU2Hr1G001320 | AT1G62200.1 |
| 251 | HORVU5Hr1G028180 | AT1G10200.1 |
| 252 | HORVU1Hr1G043900 | AT1G06950.1 |
| 253 | HORVU5Hr1G074000 | AT3G18890.1 |
| 254 | HORVU2Hr1G065420 | AT4G22390.2 |
| 255 | HORVU3Hr1G040880 | AT2G32560.1 |
| 256 | HORVU4Hr1G067020 | AT2G33380.1 |
| 257 | HORVU7Hr1G112290 | AT2G24710.3 |
| 258 | HORVU5Hr1G100870 | AT5G64640.1 |
| 259 | HORVU2Hr1G019270 | AT2G06520.1 |
| 260 | HORVU3Hr1G025820 | AT5G19760.1 |
| 261 | HORVU6Hr1G090250 | AT5G37360.1 |

| | | |
|---|---|---|
| 262 | HORVU2Hr1G079920 | AT1G15820.1 |
| 263 | HORVU1Hr1G007630 | AT5G19160.1 |
| 264 | HORVU5Hr1G016270 | AT1G27320.1 |
| 265 | HORVU2Hr1G101730 | AT4G09010.1 |
| 266 | HORVU3Hr1G109830 | AT2G16440.1 |
| 267 | HORVU5Hr1G049590 | AT1G66210.1 |
| 268 | HORVU5Hr1G102530 | AT5G44010.2 |
| 269 | HORVU3Hr1G039670 | #N/A |
| 270 | HORVU1Hr1G061970 | AT3G23600.1 |
| 271 | HORVU2Hr1G090980 | AT1G32900.1 |
| 272 | HORVU6Hr1G015770 | AT1G51400.1 |
| 273 | HORVU2Hr1G113790 | AT3G13100.1 |
| 274 | HORVU2Hr1G079180 | AT2G14095.1 |
| 275 | HORVU3Hr1G049730 | AT4G14368.1 |
| 276 | HORVU3Hr1G087210 | AT5G60010.1 |
| 277 | HORVU5Hr1G079600 | AT3G57300.2 |
| 278 | HORVU5Hr1G082420 | AT3G27690.2 |
| 279 | HORVU4Hr1G079600 | AT4G32700.2 |
| 280 | HORVU3Hr1G026920 | AT3G15353.1 |
| 281 | HORVU7Hr1G081510 | AT3G47650.1 |
| 282 | HORVU5Hr1G102620 | AT3G23800.2 |
| 283 | HORVU5Hr1G053480 | AT3G14110.2 |
| 284 | HORVU6Hr1G035460 | AT1G65640.2 |
| 285 | HORVU2Hr1G010630 | AT1G67090.1 |
| 286 | HORVU7Hr1G119060 | AT5G33370.1 |
| 287 | HORVU6Hr1G073100 | AT4G01150.1 |
| 288 | HORVU3Hr1G068200 | AT3G24450.1 |
| 289 | HORVU2Hr1G089340 | AT1G08650.1 |
| 290 | HORVU3Hr1G110660 | AT1G74880.1 |
| 291 | HORVU1Hr1G007810 | AT1G22360.1 |
| 292 | HORVU5Hr1G064230 | AT5G17420.1 |
| 293 | HORVU3Hr1G024920 | AT5G17650.1 |
| 294 | HORVU6Hr1G093940 | AT1G77490.1 |
| 295 | HORVU2Hr1G010690 | AT1G67090.1 |
| 296 | HORVU3Hr1G085520 | AT3G09220.1 |
| 297 | HORVU5Hr1G010960 | AT2G47870.1 |
| 298 | HORVU5Hr1G022750 | AT4G15510.5 |
| 299 | HORVU3Hr1G071770 | AT4G18780.1 |
| 300 | HORVU6Hr1G049260 | ATCG00490.1 |
| 301 | HORVU2Hr1G077630 | AT3G45980.1 |
| 302 | HORVU5Hr1G097870 | AT2G26500.2 |

| | | |
|---|---|---|
| 303 | HORVU2Hr1G045970 | AT3G21710.2 |
| 304 | HORVU4Hr1G009380 | AT3G22436.1 |
| 305 | HORVU5Hr1G086710 | AT1G19250.1 |
| 306 | HORVU6Hr1G051650 | AT1G14150.1 |
| 307 | HORVU7Hr1G065970 | AT3G48430.1 |
| 308 | HORVU3Hr1G039310 | AT1G29720.1 |
| 309 | HORVU4Hr1G086020 | AT5G52100.1 |
| 310 | HORVU3Hr1G096910 | AT3G60140.2 |
| 311 | HORVU5Hr1G001180 | AT3G45140.1 |
| 312 | HORVU5Hr1G018970 | AT1G22540.1 |
| 313 | HORVU5Hr1G067530 | AT1G05010.1 |
| 314 | HORVU7Hr1G047690 | AT4G20970.1 |
| 315 | HORVU3Hr1G089910 | AT2G30695.3 |
| 316 | HORVU0Hr1G012330 | AT5G42390.1 |
| 317 | HORVU0Hr1G012340 | AT2G46130.1 |
| 318 | HORVU7Hr1G086120 | AT3G47570.1 |
| 319 | HORVU7Hr1G120660 | AT5G17230.4 |
| 320 | HORVU2Hr1G073030 | AT3G52720.4 |
| 321 | HORVU7Hr1G010690 | AT4G29260.1 |
| 322 | HORVU6Hr1G050760 | AT5G02390.3 |
| 323 | HORVU2Hr1G010670 | AT1G67090.1 |
| 324 | HORVU5Hr1G023160 | AT1G47670.1 |
| 325 | HORVU5Hr1G093640 | AT1G04240.1 |
| 326 | HORVU7Hr1G058120 | AT2G34430.1 |
| 327 | HORVU4Hr1G002900 | AT3G22540.1 |
| 328 | HORVU4Hr1G076050 | AT4G35160.1 |
| 329 | HORVU2Hr1G065450 | AT5G24860.1 |
| 330 | HORVU7Hr1G098370 | AT5G57560.1 |
| 331 | HORVU5Hr1G004700 | AT5G23960.1 |
| 332 | HORVU6Hr1G012270 | AT1G61680.1 |
| 333 | HORVU4Hr1G088160 | AT2G03200.1 |
| 334 | HORVU2Hr1G094280 | AT1G33265.1 |
| 335 | HORVU0Hr1G022410 | AT1G05860.3 |
| 336 | HORVU5Hr1G067490 | AT1G05010.1 |
| 337 | HORVU1Hr1G093660 | AT1G26820.1 |
| 338 | HORVU2Hr1G063190 | AT1G24485.3 |
| 339 | HORVU5Hr1G075660 | AT3G23820.1 |
| 340 | HORVU6Hr1G061270 | AT4G36250.1 |
| 341 | HORVU2Hr1G074770 | AT2G03140.9 |
| 342 | HORVU3Hr1G108550 | AT3G43920.5 |
| 343 | HORVU4Hr1G088110 | AT2G03200.1 |

| | | |
|---|---|---|
| 344 | HORVU4Hr1G090860 | AT1G02170.1 |
| 345 | HORVU2Hr1G009970 | AT2G03200.1 |
| 346 | HORVU1Hr1G069830 | AT1G69530.2 |
| 347 | HORVU2Hr1G002720 | AT1G14880.1 |
| 348 | HORVU4Hr1G081040 | AT2G36380.1 |
| 349 | HORVU6Hr1G025050 | AT2G20930.1 |
| 350 | HORVU0Hr1G022690 | AT4G35770.1 |
| 351 | HORVU4Hr1G080820 | AT1G52690.2 |
| 352 | HORVU4Hr1G058590 | AT1G80830.2 |
| 353 | HORVU3Hr1G073860 | AT5G39210.1 |
| 354 | HORVU3Hr1G069650 | AT5G25610.1 |
| 355 | HORVU4Hr1G066230 | AT5G42650.1 |
| 356 | HORVU4Hr1G028720 | AT5G13870.3 |
| 357 | HORVU4Hr1G057000 | AT2G38905.1 |
| 358 | HORVU4Hr1G040110 | AT3G52330.1 |
| 359 | HORVU7Hr1G036720 | AT1G72610.1 |
| 360 | HORVU5Hr1G077390 | AT5G50915.4 |
| 361 | HORVU1Hr1G089380 | AT4G10550.1 |
| 362 | HORVU4Hr1G076000 | AT4G35160.1 |
| 363 | HORVU5Hr1G010950 | AT5G18600.1 |
| 364 | HORVU1Hr1G093570 | AT1G14220.1 |
| 365 | HORVU7Hr1G007800 | AT1G32560.1 |
| 366 | HORVU3Hr1G027970 | AT1G29050.1 |
| 367 | HORVU6Hr1G005050 | AT5G24860.1 |
| 368 | HORVU1Hr1G088840 | AT4G18550.1 |
| 369 | HORVU1Hr1G016410 | AT1G16460.4 |
| 370 | HORVU7Hr1G120060 | AT2G34080.1 |
| 371 | HORVU7Hr1G119930 | AT2G34080.1 |
| 372 | HORVU5Hr1G089230 | AT5G22500.1 |
| 373 | HORVU7Hr1G098320 | AT5G57550.1 |
| 374 | HORVU7Hr1G098330 | AT5G57550.1 |
| 375 | HORVU7Hr1G111010 | AT3G06130.1 |
| 376 | HORVU7Hr1G098280 | AT4G25810.1 |

SUPPLEMENTARY TABLE 10

| S. No. | Feature ID | Barley Annotation | EDGE test: Noninoc_Harr vs Harr72Hpi_Fold change | EDGE test: Noninoc_Bwt vs _Bwt72Hpi_Fold change | Ara. Homology |
|---|---|---|---|---|---|
| 1 | HORVU2Hr1G018510 | Peroxidase superfamily protein | 8262.48 | 13.51 | AT5G05340.1 |
| 2 | HORVU2Hr1G044340 | Peroxidase superfamily protein | 3094.16 | 21.87 | AT5G05340.1 |
| 3 | HORVU2Hr1G018430 | Peroxidase superfamily protein | 729.65 | 15.45 | AT5G05340.1 |
| 4 | HORVU3Hr1G091740 | Peroxidase superfamily protein | 316.99 | NS | AT5G06730.1 |
| 5 | HORVU2Hr1G018480 | Peroxidase superfamily protein | 316.07 | 11.78 | AT5G05340.1 |
| 6 | HORVU3Hr1G091750 | Peroxidase superfamily protein | 226.99 | NS | AT5G06730.1 |
| 7 | HORVU3Hr1G074960 | Peroxidase superfamily protein | 126.01 | 65.55 | AT4G11290.1 |
| 8 | HORVU2Hr1G124960 | Peroxidase superfamily protein | 115.12 | NS | AT1G71695.1 |
| 9 | HORVU1Hr1G020720 | Peroxidase superfamily protein | 98.88 | 38.41 | AT5G19890.1 |
| 10 | HORVU3Hr1G091730 | Peroxidase superfamily protein | 94.04 | NS | AT5G06730.1 |
| 11 | HORVU4Hr1G065000 | Peroxidase superfamily protein | 69.60 | NS | AT5G06730.1 |
| 12 | HORVU3Hr1G112040 | Peroxidase superfamily protein | 55.48 | NS | AT1G71695.1 |
| 13 | HORVU2Hr1G044360 | Peroxidase superfamily protein | 51.03 | 19.82 | AT5G05340.1 |
| 14 | HORVU7Hr1G076120 | Peroxidase superfamily protein | 44.9 | 54.23 | AT4G33420.1 |
| 15 | HORVU1Hr1G016870 | Peroxidase superfamily protein | 30.94 | NS | AT1G05260.1 |
| 16 | HORVU1Hr1G016840 | Peroxidase superfamily protein | 30.09 | 5.49 | AT1G05260.1 |
| 17 | HORVU4Hr1G022270 | Peroxidase superfamily protein | 27.80 | NS | AT5G05340.1 |
| 18 | HORVU2Hr1G124930 | Peroxidase superfamily protein | 24.81 | 8.74 | AT1G71695.1 |
| 19 | HORVU3Hr1G074940 | Peroxidase superfamily protein | 23.14 | NS | AT4G11290.1 |
| 20 | HORVU1Hr1G016660 | Peroxidase superfamily protein | 21.19 | 4.77 | AT4G33420.1 |
| 21 | HORVU7Hr1G083550 | Peroxidase superfamily protein | 16.63 | 7.56 | AT5G66390.1 |
| 22 | HORVU6Hr1G009360 | Peroxidase superfamily protein | 13.69 | -19.70 | AT5G05340.1 |
| 23 | HORVU2Hr1G075970 | Peroxidase superfamily protein | 13.42 | 12.79 | AT4G08390.3 |
| 24 | HORVU5Hr1G046900 | Peroxidase superfamily protein | 11.53 | NS | AT5G05340.1 |
| 25 | HORVU1Hr1G020690 | Peroxidase superfamily protein | 8.97 | NS | AT5G06730.1 |
| 26 | HORVU5Hr1G097270 | Peroxidase superfamily protein | 8.65 | 7.60 | AT3G49960.1 |
| 27 | HORVU2Hr1G018570 | Peroxidase superfamily protein | 5.78 | NS | AT5G05340.1 |
| 28 | HORVU2Hr1G125110 | Peroxidase superfamily protein | 4.50 | NS | AT1G71695.1 |
| 29 | HORVU6Hr1G021520 | Peroxidase superfamily protein | 3.18 | NS | AT5G05340.1 |
| 30 | HORVU0Hr1G004850 | Peroxidase superfamily protein | -7.47 | NS | AT4G17690.1 |
| 31 | HORVU6Hr1G009500 | Peroxidase superfamily protein | -10.75 | NS | AT5G05340.1 |
| 32 | HORVU7Hr1G098510 | Peroxidase superfamily protein | -10.82 | NS | AT2G18980.1 |
| 33 | HORVU2Hr1G002010 | Peroxidase superfamily protein | -19.94 | NS | AT1G77490.1 |
| 34 | HORVU2Hr1G063460 | Peroxidase superfamily protein | -22.46 | NS | AT4G37530.1 |
| 35 | HORVU6Hr1G093940 | Peroxidase superfamily protein | -26.19 | -7.17 | AT1G77490.1 |

SUPPLEMENTARY TABLE 11

| S No | 19 "Bowman_Up" and "Harrington_Down": | |
|---|---|---|
| 1 | HORVU7Hr1G086340 | AT2G32210.1 |
| 2 | HORVU5Hr1G034830 | AT4G11070.2 |
| 3 | HORVU3Hr1G083680 | AT2G35980.1 |
| 4 | HORVU1Hr1G019770 | AT2G05940.1 |
| 5 | HORVU4Hr1G054770 | AT4G16120.1 |
| 6 | HORVU7Hr1G107330 | AT1G56150.1 |
| 7 | HORVU7Hr1G101740 | AT4G27280.1 |
| 8 | HORVU1Hr1G070250 | AT2G38470.1 |
| 9 | HORVU3Hr1G007290 | AT5G57580.1 |
| 10 | HORVU7Hr1G036380 | AT5G27420.1 |
| 11 | HORVU4Hr1G086660 | AT5G02420.1 |
| 12 | HORVU2Hr1G123390 | AT4G10220.1 |
| 13 | HORVU5Hr1G066260 | AT3G29000.1 |
| 14 | HORVU2Hr1G118060 | AT1G01490.2 |
| 15 | HORVU3Hr1G007420 | AT5G37770.2 |
| 16 | HORVU7Hr1G118020 | AT4G35880.1 |
| 17 | HORVU7Hr1G113850 | AT1G80840.1 |
| 18 | HORVU7Hr1G022050 | AT1G10170.4 |
| 19 | HORVU5Hr1G093860 | AT4G19390.2 |

SUPPLEMENTARY TABLE 12

| S No | 9 "Harrington_Up" and "Bowman_Down": | |
|---|---|---|
| 1 | HORVU5Hr1G124120 | AT4G25420.1 |
| 2 | HORVU4Hr1G012530 | AT5G67390.3 |
| 3 | HORVU6Hr1G009360 | AT5G05340.1 |
| 4 | HORVU2Hr1G102110 | AT5G48485.1 |
| 5 | HORVU4Hr1G018150 | AT4G22756.1 |
| 6 | HORVU4Hr1G074990 | AT5G39660.3 |
| 7 | HORVU2Hr1G113260 | AT3G16150.1 |
| 8 | HORVU2Hr1G053620 | AT1G27320.1 |
| 9 | HORVU5Hr1G124160 | AT2G34420.1 |

SUPPLEMENTARY TABLE 13

| S No | #node1 | node1_id | node2 | node2_id |
|---|---|---|---|---|
| 1 | EMB3136 | AT5G13510 | SAG24 | AT1G66580 |
| 2 | EMB3136 | AT5G13510 | AT4G01310 | AT4G01310 |
| 3 | MTO2 | AT4G29840 | AKHSDH2 | AT4G19710 |

| | | | | |
|---|---|---|---|---|
| 4 | AT4G01310 | AT4G01310 | SAG24 | AT1G66580 |
| 5 | RPE | AT5G61410 | AT3G60750 | AT3G60750 |
| 6 | IAA16 | AT3G04730 | SHY2 | AT1G04240 |
| 7 | AT4G01310 | AT4G01310 | emb2184 | AT1G75350 |
| 8 | EMB3136 | AT5G13510 | emb2184 | AT1G75350 |
| 9 | RBCL | ATCG00490 | PSBA | ATCG00020 |
| 10 | PSAF | AT1G31330 | PSAO | AT1G08380 |
| 11 | HCT | AT5G48930 | CCoAOMT1 | AT4G34050 |
| 12 | NDPK2 | AT5G63310 | AT5G35170 | AT5G35170 |
| 13 | WR3 | AT5G50200 | NRT2:1 | AT1G08090 |
| 14 | ELI3-1 | AT4G37980 | ALDH2C4 | AT3G24503 |
| 15 | HCT | AT5G48930 | TT4 | AT5G13930 |
| 16 | AKHSDH2 | AT4G19710 | BCAT3 | AT3G49680 |
| 17 | PSBO2 | AT3G50820 | PSAF | AT1G31330 |
| 18 | emb2184 | AT1G75350 | SAG24 | AT1G66580 |
| 19 | DAP | AT4G33680 | AKHSDH2 | AT4G19710 |
| 20 | RBCL | ATCG00490 | RPOC1 | ATCG00180 |
| 21 | Lhca5 | AT1G45474 | PSAO | AT1G08380 |
| 22 | GR | AT3G54660 | GSTU19 | AT1G78380 |
| 23 | EXO70F1 | AT5G50380 | RAB11c | AT1G09630 |
| 24 | GR | AT3G54660 | ERD9 | AT1G10370 |
| 25 | GR | AT3G54660 | GSTU18 | AT1G10360 |
| 26 | HCHIB | AT3G12500 | PR4 | AT3G04720 |
| 27 | GR | AT3G54660 | GSTU8 | AT3G09270 |
| 28 | RPOC1 | ATCG00180 | PSBA | ATCG00020 |
| 29 | GA1 | AT4G02780 | GA2 | AT1G79460 |
| 30 | AT3G60750 | AT3G60750 | LOS2 | AT2G36530 |
| 31 | OMT1 | AT5G54160 | ELI3-1 | AT4G37980 |
| 32 | ATGSTF13 | AT3G62760 | GR | AT3G54660 |
| 33 | CPK28 | AT5G66210 | RBOHD | AT5G47910 |
| 34 | AO | AT5G14760 | AKHSDH2 | AT4G19710 |
| 35 | OMT1 | AT5G54160 | CCoAOMT1 | AT4G34050 |
| 36 | OMT1 | AT5G54160 | ALDH2C4 | AT3G24503 |
| 37 | AT5G19890 | AT5G19890 | ELI3-1 | AT4G37980 |
| 38 | TT4 | AT5G13930 | CCoAOMT1 | AT4G34050 |
| 39 | CYP71B11 | AT5G25120 | CYP71B24 | AT3G26230 |
| 40 | RPOC1 | ATCG00180 | NDPK2 | AT5G63310 |
| 41 | AT3G47570 | AT3G47570 | UKL2 | AT3G27190 |
| 42 | ELI3-1 | AT4G37980 | AT1G71695 | AT1G71695 |
| 43 | At3g49670 | AT3G49670 | UKL2 | AT3G27190 |
| 44 | CYP71B35 | AT3G26310 | CYP71B24 | AT3G26230 |

| | | | | |
|---|---|---|---|---|
| 45 | CYP71B34 | AT3G26300 | CYP71B24 | AT3G26230 |
| 46 | GDI2 | AT3G59920 | TPR10 | AT3G04710 |
| 47 | CYP71B11 | AT5G25120 | CYP71B34 | AT3G26300 |
| 48 | CYP71B11 | AT5G25120 | CYP71B35 | AT3G26310 |
| 49 | OMT1 | AT5G54160 | PRX52 | AT5G05340 |
| 50 | CYP71B35 | AT3G26310 | CYP71B34 | AT3G26300 |
| 51 | OMT1 | AT5G54160 | AT5G06730 | AT5G06730 |
| 52 | OMT1 | AT5G54160 | AT1G71695 | AT1G71695 |
| 53 | OMT1 | AT5G54160 | AT2G18140 | AT2G18140 |
| 54 | ELI3-1 | AT4G37980 | AT2G18140 | AT2G18140 |
| 55 | BT2 | AT3G48360 | CYP71B24 | AT3G26230 |
| 56 | CYP71B11 | AT5G25120 | BT2 | AT3G48360 |
| 57 | GAUT4 | AT5G47780 | AT5G19730 | AT5G19730 |
| 58 | OMT1 | AT5G54160 | AT4G11290 | AT4G11290 |
| 59 | OMT1 | AT5G54160 | AT5G19890 | AT5G19890 |
| 60 | BT2 | AT3G48360 | CYP71B34 | AT3G26300 |
| 61 | BT2 | AT3G48360 | CYP71B35 | AT3G26310 |
| 62 | AT5G06730 | AT5G06730 | ELI3-1 | AT4G37980 |
| 63 | ELI3-1 | AT4G37980 | AT4G11290 | AT4G11290 |
| 64 | PRX52 | AT5G05340 | ELI3-1 | AT4G37980 |
| 65 | PSBA | ATCG00020 | PSAF | AT1G31330 |
| 66 | CYP38 | AT3G01480 | emb2184 | AT1G75350 |
| 67 | SERAT1;1 | AT5G56760 | AKHSDH2 | AT4G19710 |
| 68 | PP2A-4 | AT3G58500 | At3g49670 | AT3G49670 |
| 69 | PP2A-4 | AT3G58500 | AT3G47570 | AT3G47570 |
| 70 | AUX1 | AT2G38120 | SHY2 | AT1G04240 |
| 71 | PSBO2 | AT3G50820 | CYP38 | AT3G01480 |
| 72 | RPOC1 | ATCG00180 | SAG24 | AT1G66580 |
| 73 | NDPK2 | AT5G63310 | AT4G01310 | AT4G01310 |
| 74 | PSBO2 | AT3G50820 | AT1G51400 | AT1G51400 |
| 75 | ACP4 | AT4G25050 | AT2G37660 | AT2G37660 |
| 76 | AT1G51400 | AT1G51400 | PSAF | AT1G31330 |
| 77 | PSBO2 | AT3G50820 | PSAO | AT1G08380 |
| 78 | ACP4 | AT4G25050 | emb2184 | AT1G75350 |
| 79 | APE2 | AT5G46110 | PSAO | AT1G08380 |
| 80 | GR | AT3G54660 | CAT1 | AT1G20630 |
| 81 | PSBA | ATCG00020 | PSBO2 | AT3G50820 |
| 82 | AT1G51400 | AT1G51400 | PSAO | AT1G08380 |
| 83 | APE2 | AT5G46110 | PSBO2 | AT3G50820 |
| 84 | APE2 | AT5G46110 | PSAF | AT1G31330 |
| 85 | AT4G01310 | AT4G01310 | AT2G37660 | AT2G37660 |

| | | | | |
|---|---|---|---|---|
| 86 | AT3G60750 | AT3G60750 | UKL2 | AT3G27190 |
| 87 | AT4G16260 | AT4G16260 | PR4 | AT3G04720 |
| 88 | EMB1030 | AT5G22800 | EDD1 | AT3G48110 |
| 89 | PAM68 | AT4G19100 | CYP38 | AT3G01480 |
| 90 | PGRL1A | AT4G22890 | AT1G18060 | AT1G18060 |
| 91 | EMB1030 | AT5G22800 | AKHSDH2 | AT4G19710 |
| 92 | CERK1 | AT3G21630 | LYK5 | AT2G33580 |
| 93 | RPE | AT5G61410 | UKL2 | AT3G27190 |
| 94 | AT2G37660 | AT2G37660 | emb2184 | AT1G75350 |
| 95 | At3g49670 | AT3G49670 | RAB11c | AT1G09630 |
| 96 | AT3G47570 | AT3G47570 | RAB11c | AT1G09630 |
| 97 | TYRDC | AT4G28680 | AAS | AT2G20340 |
| 98 | NRT1:2 | AT1G69850 | NRT2:1 | AT1G08090 |
| 99 | AT1G51110 | AT1G51110 | AT1G18060 | AT1G18060 |
| 100 | LPA2 | AT5G51545 | PAM68 | AT4G19100 |
| 101 | EMB3136 | AT5G13510 | CYP38 | AT3G01480 |
| 102 | PGRL1A | AT4G22890 | PSBO2 | AT3G50820 |
| 103 | AT3G47570 | AT3G47570 | TPR10 | AT3G04710 |
| 104 | At3g49670 | AT3G49670 | TPR10 | AT3G04710 |
| 105 | ACP4 | AT4G25050 | CYP38 | AT3G01480 |
| 106 | AT5G35170 | AT5G35170 | AT4G01310 | AT4G01310 |
| 107 | APE2 | AT5G46110 | AT1G51400 | AT1G51400 |
| 108 | NDPK2 | AT5G63310 | AT2G37660 | AT2G37660 |
| 109 | EMB3136 | AT5G13510 | AT2G37660 | AT2G37660 |
| 110 | AT5G58490 | AT5G58490 | UGD3 | AT5G15490 |
| 111 | NDPK2 | AT5G63310 | EMB3136 | AT5G13510 |
| 112 | AT4G16260 | AT4G16260 | HCHIB | AT3G12500 |
| 113 | NDPK2 | AT5G63310 | AKHSDH2 | AT4G19710 |
| 114 | RPOC1 | ATCG00180 | AT4G01310 | AT4G01310 |
| 115 | Lhca5 | AT1G45474 | PSAF | AT1G31330 |
| 116 | ALN | AT4G04955 | AT2G31050 | AT2G31050 |
| 117 | OSM34 | AT4G11650 | HCHIB | AT3G12500 |
| 118 | EMB3136 | AT5G13510 | ACP4 | AT4G25050 |
| 119 | AT5G35170 | AT5G35170 | AT1G51110 | AT1G51110 |
| 120 | DAP | AT4G33680 | BCAT3 | AT3G49680 |
| 121 | PnsB2 | AT1G64770 | Lhca5 | AT1G45474 |
| 122 | GR | AT3G54660 | MDAR4 | AT3G27820 |
| 123 | CYP38 | AT3G01480 | AT2G42220 | AT2G42220 |
| 124 | KEA3 | AT4G04850 | NHD1 | AT3G19490 |
| 125 | CYP38 | AT3G01480 | AT2G37660 | AT2G37660 |
| 126 | PGRL1A | AT4G22890 | CYP38 | AT3G01480 |

| | | | | |
|---|---|---|---|---|
| 127 | PGRL1A | AT4G22890 | AT1G51110 | AT1G51110 |
| 128 | AKHSDH2 | AT4G19710 | EDD1 | AT3G48110 |
| 129 | ACP4 | AT4G25050 | AT2G42220 | AT2G42220 |
| 130 | RBCL | ATCG00490 | UKL2 | AT3G27190 |
| 131 | AT4G01310 | AT4G01310 | CYP38 | AT3G01480 |
| 132 | RPE | AT5G61410 | AT5G48220 | AT5G48220 |
| 133 | UVR8 | AT5G63860 | TT4 | AT5G13930 |
| 134 | PGRL1A | AT4G22890 | Lhca5 | AT1G45474 |
| 135 | KEA3 | AT4G04850 | ECA3 | AT1G10130 |
| 136 | CYP38 | AT3G01480 | PSAF | AT1G31330 |
| 137 | PSBA | ATCG00020 | PAM68 | AT4G19100 |
| 138 | AT4G39970 | AT4G39970 | AT2G33255 | AT2G33255 |
| 139 | AT1G77090 | AT1G77090 | AT1G18060 | AT1G18060 |
| 140 | PGRL1A | AT4G22890 | AT1G51400 | AT1G51400 |
| 141 | PSBO2 | AT3G50820 | Lhca5 | AT1G45474 |
| 142 | AKHSDH2 | AT4G19710 | At3g49670 | AT3G49670 |
| 143 | GDI2 | AT3G59920 | RAB11c | AT1G09630 |
| 144 | AT5G19850 | AT5G19850 | GR | AT3G54660 |
| 145 | AT4G09040 | AT4G09040 | At3g49670 | AT3G49670 |
| 146 | At3g49670 | AT3G49670 | AT2G28710 | AT2G28710 |
| 147 | YY1 | AT4G06634 | AT3G47570 | AT3G47570 |
| 148 | YY1 | AT4G06634 | At3g49670 | AT3G49670 |
| 149 | PnsB2 | AT1G64770 | AT1G64680 | AT1G64680 |
| 150 | AT3G47570 | AT3G47570 | AT2G28710 | AT2G28710 |
| 151 | AT2G42220 | AT2G42220 | AT1G51110 | AT1G51110 |
| 152 | AT4G28080 | AT4G28080 | AT3G48200 | AT3G48200 |
| 153 | ACP4 | AT4G25050 | At1g72610 | AT1G72610 |
| 154 | AT5G35170 | AT5G35170 | PGRL1A | AT4G22890 |
| 155 | LPA2 | AT5G51545 | CYP38 | AT3G01480 |
| 156 | At3g49670 | AT3G49670 | AT3G47570 | AT3G47570 |
| 157 | AT5G35170 | AT5G35170 | SAG24 | AT1G66580 |
| 158 | CYP38 | AT3G01480 | AT1G51110 | AT1G51110 |
| 159 | RBCL | ATCG00490 | PSAF | AT1G31330 |
| 160 | AT1G77090 | AT1G77090 | AT1G64680 | AT1G64680 |
| 161 | emb2746 | AT5G63420 | SVR1 | AT2G39140 |
| 162 | RPE | AT5G61410 | APE2 | AT5G46110 |
| 163 | AT2G39670 | AT2G39670 | SVR1 | AT2G39140 |
| 164 | AT5G39110 | AT5G39110 | AT4G33720 | AT4G33720 |
| 165 | RBOHD | AT5G47910 | RPM1 | AT3G07040 |
| 166 | ACP4 | AT4G25050 | PSBO2 | AT3G50820 |
| 167 | AT5G47530 | AT5G47530 | AT1G61590 | AT1G61590 |

| | | | | |
|---|---|---|---|---|
| 168 | NRT1:2 | AT1G69850 | ABCG40 | AT1G15520 |
| 169 | AT1G77090 | AT1G77090 | AT1G51110 | AT1G51110 |
| 170 | PGRL1A | AT4G22890 | AT1G64680 | AT1G64680 |
| 171 | YLS9 | AT2G35980 | ABCG40 | AT1G15520 |
| 172 | RPOC1 | ATCG00180 | EMB3136 | AT5G13510 |
| 173 | AKHSDH2 | AT4G19710 | AT3G47570 | AT3G47570 |
| 174 | FRS5 | AT4G38180 | CYP71A25 | AT3G48280 |
| 175 | EMB1030 | AT5G22800 | UKL2 | AT3G27190 |
| 176 | AT5G35170 | AT5G35170 | CYP38 | AT3G01480 |
| 177 | AT5G02070 | AT5G02070 | AKHSDH2 | AT4G19710 |
| 178 | AKHSDH2 | AT4G19710 | AT1G16260 | AT1G16260 |
| 179 | AKHSDH2 | AT4G19710 | WAK2 | AT1G21270 |
| 180 | B120 | AT4G21390 | AKHSDH2 | AT4G19710 |
| 181 | AT5G65600 | AT5G65600 | AKHSDH2 | AT4G19710 |
| 182 | AT5G51770 | AT5G51770 | AKHSDH2 | AT4G19710 |
| 183 | AKHSDH2 | AT4G19710 | AT1G61590 | AT1G61590 |
| 184 | AKHSDH2 | AT4G19710 | CRK2 | AT1G70520 |
| 185 | AT5G12000 | AT5G12000 | AKHSDH2 | AT4G19710 |
| 186 | AKHSDH2 | AT4G19710 | AT2G23450 | AT2G23450 |
| 187 | AKHSDH2 | AT4G19710 | LYK5 | AT2G33580 |
| 188 | AKHSDH2 | AT4G19710 | WAK5 | AT1G21230 |
| 189 | AT5G10530 | AT5G10530 | AKHSDH2 | AT4G19710 |
| 190 | AKHSDH2 | AT4G19710 | CERK1 | AT3G21630 |
| 191 | AT4G29990 | AT4G29990 | AKHSDH2 | AT4G19710 |
| 192 | AKHSDH2 | AT4G19710 | CIPK9 | AT1G01140 |
| 193 | AKHSDH2 | AT4G19710 | AT1G61420 | AT1G61420 |
| 194 | AKHSDH2 | AT4G19710 | CCR3 | AT3G55950 |
| 195 | AKHSDH2 | AT4G19710 | WAK3 | AT1G21240 |
| 196 | PGRL1A | AT4G22890 | emb2184 | AT1G75350 |
| 197 | Lhca5 | AT1G45474 | AT1G18060 | AT1G18060 |
| 198 | FAD6 | AT4G30950 | FATB | AT1G08510 |
| 199 | HCT | AT5G48930 | CYP98A8 | AT1G74540 |
| 200 | CPK28 | AT5G66210 | LYK5 | AT2G33580 |
| 201 | ALDH2C4 | AT3G24503 | PKT3 | AT2G33150 |
| 202 | emb2746 | AT5G63420 | AT2G39670 | AT2G39670 |
| 203 | AT4G09040 | AT4G09040 | AT3G47570 | AT3G47570 |
| 204 | OSM34 | AT4G11650 | DOX1 | AT3G01420 |
| 205 | PR4 | AT3G04720 | PR1 | AT2G14610 |
| 206 | RBCL | ATCG00490 | LOS2 | AT2G36530 |
| 207 | RPOC1 | ATCG00180 | AT5G35170 | AT5G35170 |
| 208 | APE2 | AT5G46110 | PGRL1A | AT4G22890 |

| | | | | |
|---|---|---|---|---|
| 209 | NDPK2 | AT5G63310 | AT2G39670 | AT2G39670 |
| 210 | AT2G42220 | AT2G42220 | AT1G51400 | AT1G51400 |
| 211 | APE2 | AT5G46110 | AT2G42220 | AT2G42220 |
| 212 | PGRL1A | AT4G22890 | AT1G77090 | AT1G77090 |
| 213 | HCHIB | AT3G12500 | PR1 | AT2G14610 |
| 214 | PSBO2 | AT3G50820 | AT1G77090 | AT1G77090 |
| 215 | PSBO2 | AT3G50820 | AT2G42220 | AT2G42220 |
| 216 | AT4G16260 | AT4G16260 | OSM34 | AT4G11650 |
| 217 | PnsB2 | AT1G64770 | AT1G18060 | AT1G18060 |
| 218 | CERK1 | AT3G21630 | RPM1 | AT3G07040 |
| 219 | OSM34 | AT4G11650 | AT2G15220 | AT2G15220 |
| 220 | PnsB2 | AT1G64770 | AT1G51110 | AT1G51110 |
| 221 | UVR8 | AT5G63860 | EMB2730 | AT5G02250 |
| 222 | CGLD27 | AT5G67370 | PGRL1A | AT4G22890 |
| 223 | VSP1 | AT5G24780 | PR4 | AT3G04720 |
| 224 | HCT | AT5G48930 | TYRDC | AT4G28680 |
| 225 | AT5G48220 | AT5G48220 | AKHSDH2 | AT4G19710 |
| 226 | AT3G60750 | AT3G60750 | ALDH2C4 | AT3G24503 |
| 227 | AT4G39970 | AT4G39970 | CYP38 | AT3G01480 |
| 228 | PR1 | AT2G14610 | FMO1 | AT1G19250 |
| 229 | TT4 | AT5G13930 | SHY2 | AT1G04240 |
| 230 | CGLD27 | AT5G67370 | AT1G77090 | AT1G77090 |
| 231 | GA1 | AT4G02780 | AUX1 | AT2G38120 |
| 232 | ALDH2C4 | AT3G24503 | CAT1 | AT1G20630 |
| 233 | MDAR4 | AT3G27820 | UKL2 | AT3G27190 |
| 234 | AT5G35170 | AT5G35170 | UKL2 | AT3G27190 |
| 235 | AT3G22600 | AT3G22600 | YLS9 | AT2G35980 |
| 236 | AT4G28080 | AT4G28080 | CYP38 | AT3G01480 |
| 237 | APE2 | AT5G46110 | CYP38 | AT3G01480 |
| 238 | At3g49670 | AT3G49670 | CYP38 | AT3G01480 |
| 239 | CYP71B37 | AT3G26330 | FD3 | AT2G27510 |
| 240 | AT1G77090 | AT1G77090 | Lhca5 | AT1G45474 |
| 241 | AT4G09040 | AT4G09040 | CYP38 | AT3G01480 |
| 242 | CYP38 | AT3G01480 | Lhca5 | AT1G45474 |
| 243 | AT1G77090 | AT1G77090 | PnsB2 | AT1G64770 |
| 244 | EMB2730 | AT5G02250 | LOS2 | AT2G36530 |
| 245 | AT2G42220 | AT2G42220 | emb2184 | AT1G75350 |
| 246 | RPE | AT5G61410 | PGRL1A | AT4G22890 |
| 247 | CYP38 | AT3G01480 | At1g72610 | AT1G72610 |
| 248 | AT4G13010 | AT4G13010 | ALDH2C4 | AT3G24503 |
| 249 | ALDH2C4 | AT3G24503 | NADK2 | AT1G21640 |

| | | | | |
|---|---|---|---|---|
| 250 | NEET | AT5G51720 | MDAR4 | AT3G27820 |
| 251 | ACP4 | AT4G25050 | AT4G01310 | AT4G01310 |
| 252 | RBOHD | AT5G47910 | CERK1 | AT3G21630 |
| 253 | BT2 | AT3G48360 | UKL2 | AT3G27190 |
| 254 | RBCL | ATCG00490 | AT4G39970 | AT4G39970 |
| 255 | AT4G10540 | AT4G10540 | AT3G47570 | AT3G47570 |
| 256 | AT4G10540 | AT4G10540 | At3g49670 | AT3G49670 |
| 257 | KEA3 | AT4G04850 | GR | AT3G54660 |
| 258 | GR | AT3G54660 | NADK2 | AT1G21640 |
| 259 | CYP38 | AT3G01480 | PnsB2 | AT1G64770 |
| 260 | RPE | AT5G61410 | CLA1 | AT4G15560 |
| 261 | AT5G48220 | AT5G48220 | CYP38 | AT3G01480 |
| 262 | AT5G39110 | AT5G39110 | WAK5 | AT1G21230 |
| 263 | AT5G65600 | AT5G65600 | ABCG40 | AT1G15520 |
| 264 | APE2 | AT5G46110 | ACP4 | AT4G25050 |
| 265 | AT2G42220 | AT2G42220 | AT1G18060 | AT1G18060 |
| 266 | RPE | AT5G61410 | emb2184 | AT1G75350 |
| 267 | WRKY75 | AT5G13080 | YLS9 | AT2G35980 |
| 268 | FD3 | AT2G27510 | CYP89A6 | AT1G64940 |
| 269 | GR | AT3G54660 | LOS2 | AT2G36530 |
| 270 | CNI1 | AT5G27420 | LYK5 | AT2G33580 |
| 271 | RPE | AT5G61410 | AT2G37660 | AT2G37660 |
| 272 | HINT3 | AT5G48545 | LOS2 | AT2G36530 |
| 273 | ELI3-1 | AT4G37980 | AT4G13010 | AT4G13010 |
| 274 | AT1G64680 | AT1G64680 | Lhca5 | AT1G45474 |
| 275 | UKL2 | AT3G27190 | emb2184 | AT1G75350 |
| 276 | OSM34 | AT4G11650 | PR4 | AT3G04720 |
| 277 | B120 | AT4G21390 | CCR3 | AT3G55950 |
| 278 | RPE | AT5G61410 | CYP38 | AT3G01480 |
| 279 | emb2746 | AT5G63420 | EMB1030 | AT5G22800 |
| 280 | NDPK2 | AT5G63310 | CYP38 | AT3G01480 |
| 281 | DAP | AT4G33680 | MTO2 | AT4G29840 |
| 282 | PGRL1A | AT4G22890 | PnsB2 | AT1G64770 |
| 283 | GR | AT3G54660 | BCAT3 | AT3G49680 |
| 284 | FAD6 | AT4G30950 | CYP38 | AT3G01480 |
| 285 | NDPK2 | AT5G63310 | At1g54130 | AT1G54130 |
| 286 | OMT1 | AT5G54160 | HCT | AT5G48930 |
| 287 | PP2A-4 | AT3G58500 | CIPK9 | AT1G01140 |
| 288 | B120 | AT4G21390 | MYB15 | AT3G23250 |
| 289 | AT3G60750 | AT3G60750 | GR | AT3G54660 |
| 290 | AT1G50000 | AT1G50000 | AT1G45110 | AT1G45110 |

| | | | | |
|---|---|---|---|---|
| 291 | DAP | AT4G33680 | GulLO2 | AT2G46750 |
| 292 | DAP | AT4G33680 | AT4G20820 | AT4G20820 |
| 293 | EMB1030 | AT5G22800 | EMB3136 | AT5G13510 |
| 294 | DMR6 | AT5G24530 | CYP71B37 | AT3G26330 |
| 295 | PP2A-4 | AT3G58500 | WAK2 | AT1G21270 |
| 296 | ALDH2C4 | AT3G24503 | HB1 | AT2G16060 |
| 297 | ABCB20 | AT3G55320 | ABCG39 | AT1G66950 |
| 298 | AKHSDH2 | AT4G19710 | AT1G45110 | AT1G45110 |
| 299 | ACP4 | AT4G25050 | At1g54130 | AT1G54130 |
| 300 | At3g49670 | AT3G49670 | AT2G39670 | AT2G39670 |
| 301 | PP2A-4 | AT3G58500 | CERK1 | AT3G21630 |
| 302 | AT5G48960 | AT5G48960 | AT4G01310 | AT4G01310 |
| 303 | MYB15 | AT3G23250 | AT2G42270 | AT2G42270 |
| 304 | AT2G42220 | AT2G42220 | PSAF | AT1G31330 |
| 305 | RBOHD | AT5G47910 | LAC7 | AT3G09220 |
| 306 | AZI1 | AT4G12470 | LTPG1 | AT1G27950 |
| 307 | PP2A-4 | AT3G58500 | WAK5 | AT1G21230 |
| 308 | PP2A-4 | AT3G58500 | AT2G23450 | AT2G23450 |
| 309 | AT5G02070 | AT5G02070 | PP2A-4 | AT3G58500 |
| 310 | PP2A-4 | AT3G58500 | CRK2 | AT1G70520 |
| 311 | AT5G10530 | AT5G10530 | PP2A-4 | AT3G58500 |
| 312 | PP2A-4 | AT3G58500 | WAK3 | AT1G21240 |
| 313 | PP2A-4 | AT3G58500 | AT1G61590 | AT1G61590 |
| 314 | AT5G65600 | AT5G65600 | PP2A-4 | AT3G58500 |
| 315 | PP2A-4 | AT3G58500 | LYK5 | AT2G33580 |
| 316 | AT4G29990 | AT4G29990 | PP2A-4 | AT3G58500 |
| 317 | AT4G39970 | AT4G39970 | AT1G51110 | AT1G51110 |
| 318 | PP2A-4 | AT3G58500 | AT1G16260 | AT1G16260 |
| 319 | PP2A-4 | AT3G58500 | CCR3 | AT3G55950 |
| 320 | AT1G78140 | AT1G78140 | AT1G51110 | AT1G51110 |
| 321 | B120 | AT4G21390 | PP2A-4 | AT3G58500 |
| 322 | AT5G12000 | AT5G12000 | PP2A-4 | AT3G58500 |
| 323 | AT5G51770 | AT5G51770 | PP2A-4 | AT3G58500 |
| 324 | J8 | AT1G80920 | AMI1 | AT1G08980 |
| 325 | PP2A-4 | AT3G58500 | AT1G61420 | AT1G61420 |
| 326 | PAM68 | AT4G19100 | AT1G78915 | AT1G78915 |
| 327 | RBOHD | AT5G47910 | CNI1 | AT5G27420 |
| 328 | LPA2 | AT5G51545 | AT4G01310 | AT4G01310 |
| 329 | APE2 | AT5G46110 | Lhca5 | AT1G45474 |
| 330 | GR | AT3G54660 | AAS | AT2G20340 |
| 331 | EMB2730 | AT5G02250 | AKHSDH2 | AT4G19710 |

| | | | | |
|---|---|---|---|---|
| 332 | DIT2.1 | AT5G64290 | NHD1 | AT3G19490 |
| 333 | ELI3-1 | AT4G37980 | AAS | AT2G20340 |
| 334 | AT2G42220 | AT2G42220 | PSAO | AT1G08380 |
| 335 | LOS2 | AT2G36530 | SAG24 | AT1G66580 |
| 336 | PGRL1A | AT4G22890 | AT1G52870 | AT1G52870 |
| 337 | PGRL1A | AT4G22890 | KEA3 | AT4G04850 |
| 338 | EMB3136 | AT5G13510 | FAD6 | AT4G30950 |
| 339 | AKHSDH2 | AT4G19710 | GR | AT3G54660 |
| 340 | RPE | AT5G61410 | AT2G33255 | AT2G33255 |
| 341 | CPK28 | AT5G66210 | AT3G47570 | AT3G47570 |
| 342 | PKT3 | AT2G33150 | CAT1 | AT1G20630 |
| 343 | AT5G35170 | AT5G35170 | At1g54130 | AT1G54130 |
| 344 | AT4G01310 | AT4G01310 | AT2G42220 | AT2G42220 |
| 345 | CYP94D2 | AT3G56630 | XPL1 | AT3G18000 |
| 346 | RPE | AT5G61410 | PSBO2 | AT3G50820 |
| 347 | RBCL | ATCG00490 | AT2G33255 | AT2G33255 |
| 348 | DIT2.1 | AT5G64290 | APE2 | AT5G46110 |
| 349 | CGLD27 | AT5G67370 | BTS | AT3G18290 |
| 350 | AT1G51110 | AT1G51110 | Lhca5 | AT1G45474 |
| 351 | TYRDC | AT4G28680 | GR | AT3G54660 |
| 352 | FD3 | AT2G27510 | CYP79F2 | AT1G16400 |
| 353 | AT4G20820 | AT4G20820 | AT2G33255 | AT2G33255 |
| 354 | AT4G20820 | AT4G20820 | ALDH2C4 | AT3G24503 |
| 355 | ALDH2C4 | AT3G24503 | GulLO2 | AT2G46750 |
| 356 | GulLO2 | AT2G46750 | AT2G33255 | AT2G33255 |
| 357 | NDPK2 | AT5G63310 | DAP | AT4G33680 |
| 358 | AT5G58490 | AT5G58490 | CYP71B34 | AT3G26300 |
| 359 | NAPRT1 | AT4G36940 | DJ1B | AT1G53280 |
| 360 | TYRDC | AT4G28680 | BCAT3 | AT3G49680 |
| 361 | DAP | AT4G33680 | AT4G26860 | AT4G26860 |
| 362 | BCAT3 | AT3G49680 | AAS | AT2G20340 |
| 363 | RPE | AT5G61410 | AMI1 | AT1G08980 |
| 364 | LOS2 | AT2G36530 | AT1G61420 | AT1G61420 |
| 365 | NDPK2 | AT5G63310 | AT2G47870 | AT2G47870 |
| 366 | TYRDC | AT4G28680 | AT4G13010 | AT4G13010 |
| 367 | AT2G42220 | AT2G42220 | AT2G37660 | AT2G37660 |
| 368 | AT3G47570 | AT3G47570 | CYP38 | AT3G01480 |
| 369 | UKL2 | AT3G27190 | CDS1 | AT1G62430 |
| 370 | AT4G13010 | AT4G13010 | AAS | AT2G20340 |
| 371 | AT4G16260 | AT4G16260 | AT2G15220 | AT2G15220 |
| 372 | AT4G39970 | AT4G39970 | ALDH2C4 | AT3G24503 |

| | | | | |
|---|---|---|---|---|
| 373 | At3g49670 | AT3G49670 | AT2G42270 | AT2G42270 |
| 374 | AT3G47570 | AT3G47570 | AT2G42270 | AT2G42270 |
| 375 | AKHSDH2 | AT4G19710 | AT3G60750 | AT3G60750 |
| 376 | FAD6 | AT4G30950 | ACP4 | AT4G25050 |
| 377 | AO | AT5G14760 | DAP | AT4G33680 |
| 378 | RPE | AT5G61410 | WAK2 | AT1G21270 |
| 379 | LPA2 | AT5G51545 | EMB3136 | AT5G13510 |
| 380 | EDD1 | AT3G48110 | AT1G50000 | AT1G50000 |
| 381 | CYP76C4 | AT2G45550 | UGT85A3 | AT1G22380 |
| 382 | UGT76B1 | AT3G11340 | ILA | AT1G64790 |
| 383 | CLA1 | AT4G15560 | NADK2 | AT1G21640 |
| 384 | KEA3 | AT4G04850 | AT1G64680 | AT1G64680 |
| 385 | PDX2 | AT5G60540 | BCAT3 | AT3G49680 |
| 386 | BCAT3 | AT3G49680 | MDAR4 | AT3G27820 |
| 387 | RBOHD | AT5G47910 | GR | AT3G54660 |
| 388 | NAPRT1 | AT4G36940 | UKL2 | AT3G27190 |
| 389 | AT4G26860 | AT4G26860 | NADK2 | AT1G21640 |
| 390 | AT5G48220 | AT5G48220 | PnsB2 | AT1G64770 |
| 391 | FD3 | AT2G27510 | CYP98A8 | AT1G74540 |
| 392 | CYP71B11 | AT5G25120 | FD3 | AT2G27510 |
| 393 | AKHSDH2 | AT4G19710 | LOS2 | AT2G36530 |
| 394 | CYP71A25 | AT3G48280 | FD3 | AT2G27510 |
| 395 | CYP71B34 | AT3G26300 | FD3 | AT2G27510 |
| 396 | CYP94D2 | AT3G56630 | FD3 | AT2G27510 |
| 397 | CYP71B10 | AT5G57260 | FD3 | AT2G27510 |
| 398 | WRKY75 | AT5G13080 | NAC042 | AT2G43000 |
| 399 | CYP71B24 | AT3G26230 | FD3 | AT2G27510 |
| 400 | FD3 | AT2G27510 | CYP711A1 | AT2G26170 |
| 401 | CYP71B35 | AT3G26310 | FD3 | AT2G27510 |
| 402 | CYP76C4 | AT2G45550 | FD3 | AT2G27510 |
| 403 | HCHIB | AT3G12500 | RPM1 | AT3G07040 |
| 404 | CCoAOMT1 | AT4G34050 | UKL2 | AT3G27190 |
| 405 | LAC7 | AT3G09220 | DOX1 | AT3G01420 |
| 406 | VIT1 | AT2G01770 | ECA3 | AT1G10130 |
| 407 | PAM68 | AT4G19100 | AT1G77090 | AT1G77090 |
| 408 | At1g72610 | AT1G72610 | PSAF | AT1G31330 |
| 409 | AT3G60750 | AT3G60750 | CAT1 | AT1G20630 |
| 410 | RPE | AT5G61410 | GR | AT3G54660 |
| 411 | AT5G35170 | AT5G35170 | ECA3 | AT1G10130 |
| 412 | UKL2 | AT3G27190 | LOS2 | AT2G36530 |
| 413 | CYP38 | AT3G01480 | AT1G77090 | AT1G77090 |

| | | | | |
|---|---|---|---|---|
| 414 | EMB3136 | AT5G13510 | PSBO2 | AT3G50820 |
| 415 | PAM68 | AT4G19100 | PSBO2 | AT3G50820 |
| 416 | KEA3 | AT4G04850 | AT1G77090 | AT1G77090 |
| 417 | ACP4 | AT4G25050 | PGRL1A | AT4G22890 |
| 418 | RPE | AT5G61410 | AT5G58490 | AT5G58490 |
| 419 | AT5G58490 | AT5G58490 | EDD1 | AT3G48110 |
| 420 | GR | AT3G54660 | AT2G47870 | AT2G47870 |
| 421 | HINT3 | AT5G48545 | GSTU19 | AT1G78380 |
| 422 | WRKY75 | AT5G13080 | UGT76B1 | AT3G11340 |
| 423 | TPR10 | AT3G04710 | CAT1 | AT1G20630 |
| 424 | NDPK2 | AT5G63310 | UKL2 | AT3G27190 |
| 425 | RPE | AT5G61410 | AT2G39670 | AT2G39670 |
| 426 | TT4 | AT5G13930 | AUX1 | AT2G38120 |
| 427 | AT4G09040 | AT4G09040 | AT2G42270 | AT2G42270 |
| 428 | CNI1 | AT5G27420 | B120 | AT4G21390 |
| 429 | AT5G58490 | AT5G58490 | CYP79F2 | AT1G16400 |
| 430 | AT5G58490 | AT5G58490 | CYP71A25 | AT3G48280 |
| 431 | AT5G58490 | AT5G58490 | CYP71B10 | AT5G57260 |
| 432 | AT5G58490 | AT5G58490 | CYP71B24 | AT3G26230 |
| 433 | AT5G58490 | AT5G58490 | CYP98A8 | AT1G74540 |
| 434 | AT5G58490 | AT5G58490 | CYP71B35 | AT3G26310 |
| 435 | AT5G58490 | AT5G58490 | CYP71B11 | AT5G25120 |
| 436 | AT5G58490 | AT5G58490 | CYP711A1 | AT2G26170 |
| 437 | AT5G58490 | AT5G58490 | CYP94D2 | AT3G56630 |
| 438 | AT5G65600 | AT5G65600 | B120 | AT4G21390 |
| 439 | AT5G58490 | AT5G58490 | CYP76C4 | AT2G45550 |
| 440 | AT5G58490 | AT5G58490 | CYP71B37 | AT3G26330 |
| 441 | AT5G58490 | AT5G58490 | CYP89A6 | AT1G64940 |
| 442 | RPE | AT5G61410 | AT1G16260 | AT1G16260 |
| 443 | EDD1 | AT3G48110 | AT1G78915 | AT1G78915 |
| 444 | emb1417 | AT4G21190 | emb2184 | AT1G75350 |
| 445 | AKHSDH2 | AT4G19710 | CAT1 | AT1G20630 |
| 446 | CLA1 | AT4G15560 | AT3G60750 | AT3G60750 |
| 447 | HCHIB | AT3G12500 | AT2G15220 | AT2G15220 |
| 448 | AT5G35170 | AT5G35170 | GR | AT3G54660 |
| 449 | At3g49670 | AT3G49670 | CYP71A25 | AT3G48280 |
| 450 | CYP38 | AT3G01480 | AT1G18060 | AT1G18060 |
| 451 | NDPK2 | AT5G63310 | AT1G61420 | AT1G61420 |
| 452 | IAA16 | AT3G04730 | AUX1 | AT2G38120 |
| 453 | AT4G34480 | AT4G34480 | UGT85A3 | AT1G22380 |
| 454 | RR12 | AT2G25180 | SHY2 | AT1G04240 |

| | | | | |
|---|---|---|---|---|
| 455 | EMB3136 | AT5G13510 | PGRL1A | AT4G22890 |
| 456 | CLA1 | AT4G15560 | AT1G51400 | AT1G51400 |
| 457 | TYRDC | AT4G28680 | ABCB20 | AT3G55320 |
| 458 | CPK28 | AT5G66210 | MYB15 | AT3G23250 |
| 459 | AT2G42220 | AT2G42220 | At1g72610 | AT1G72610 |
| 460 | emb2746 | AT5G63420 | AT4G17740 | AT4G17740 |
| 461 | emb2184 | AT1G75350 | PSAF | AT1G31330 |
| 462 | RPE | AT5G61410 | CERK1 | AT3G21630 |
| 463 | PSBO2 | AT3G50820 | AT1G51110 | AT1G51110 |
| 464 | AKHSDH2 | AT4G19710 | UKL2 | AT3G27190 |
| 465 | AT4G39970 | AT4G39970 | UKL2 | AT3G27190 |
| 466 | HINT3 | AT5G48545 | APTX | AT5G01310 |
| 467 | CB5-E | AT5G53560 | AT4G24330 | AT4G24330 |
| 468 | PGRL1A | AT4G22890 | AT2G42220 | AT2G42220 |
| 469 | NADK2 | AT1G21640 | AT1G18060 | AT1G18060 |
| 470 | NAC042 | AT2G43000 | YLS9 | AT2G35980 |
| 471 | AT1G61420 | AT1G61420 | DJ1B | AT1G53280 |
| 472 | CGLD27 | AT5G67370 | CYP38 | AT3G01480 |
| 473 | KEA3 | AT4G04850 | AT3G47570 | AT3G47570 |
| 474 | AKHSDH2 | AT4G19710 | AT1G50000 | AT1G50000 |
| 475 | KEA3 | AT4G04850 | At3g49670 | AT3G49670 |
| 476 | LAP1 | AT2G24200 | AT1G19450 | AT1G19450 |
| 477 | LOS2 | AT2G36530 | RR12 | AT2G25180 |
| 478 | AT3G47570 | AT3G47570 | AT2G39670 | AT2G39670 |
| 479 | RBOHD | AT5G47910 | CAT1 | AT1G20630 |
| 480 | CYP71B34 | AT3G26300 | WAK5 | AT1G21230 |
| 481 | ABCB20 | AT3G55320 | ABCG40 | AT1G15520 |
| 482 | NDPK2 | AT5G63310 | LAP1 | AT2G24200 |
| 483 | FAD6 | AT4G30950 | HB1 | AT2G16060 |
| 484 | PGRL1A | AT4G22890 | GSTU18 | AT1G10360 |
| 485 | EMB1030 | AT5G22800 | CDS1 | AT1G62430 |
| 486 | AT5G35170 | AT5G35170 | AKHSDH2 | AT4G19710 |
| 487 | AKHSDH2 | AT4G19710 | NADK2 | AT1G21640 |
| 488 | RPE | AT5G61410 | AT1G51110 | AT1G51110 |
| 489 | MTO2 | AT4G29840 | LOS2 | AT2G36530 |
| 490 | AKHSDH2 | AT4G19710 | ALDH2C4 | AT3G24503 |
| 491 | AT5G48220 | AT5G48220 | EDD1 | AT3G48110 |
| 492 | RPE | AT5G61410 | CIPK9 | AT1G01140 |
| 493 | RPE | AT5G61410 | AT1G61590 | AT1G61590 |
| 494 | RPE | AT5G61410 | LYK5 | AT2G33580 |
| 495 | RPE | AT5G61410 | B120 | AT4G21390 |

| | | | | |
|---|---|---|---|---|
| 496 | RPE | AT5G61410 | At3g49670 | AT3G49670 |
| 497 | RPE | AT5G61410 | AT1G61420 | AT1G61420 |
| 498 | RPE | AT5G61410 | AT2G23450 | AT2G23450 |
| 499 | RPE | AT5G61410 | AT3G47570 | AT3G47570 |
| 500 | RPE | AT5G61410 | CCR3 | AT3G55950 |
| 501 | RPE | AT5G61410 | AT4G29990 | AT4G29990 |
| 502 | RPE | AT5G61410 | AT5G10530 | AT5G10530 |
| 503 | RPE | AT5G61410 | CRK2 | AT1G70520 |
| 504 | RPE | AT5G61410 | WAK3 | AT1G21240 |
| 505 | WRKY75 | AT5G13080 | MYB15 | AT3G23250 |
| 506 | RPE | AT5G61410 | AT5G12000 | AT5G12000 |
| 507 | RPE | AT5G61410 | WAK5 | AT1G21230 |
| 508 | RPE | AT5G61410 | AT5G02070 | AT5G02070 |
| 509 | RPE | AT5G61410 | AT5G51770 | AT5G51770 |
| 510 | AT5G65600 | AT5G65600 | RPE | AT5G61410 |
| 511 | HSF3 | AT5G16820 | AT2G15130 | AT2G15130 |
| 512 | At3g49670 | AT3G49670 | ECA3 | AT1G10130 |
| 513 | VSP1 | AT5G24780 | HCHIB | AT3G12500 |
| 514 | CYP38 | AT3G01480 | AT1G64680 | AT1G64680 |
| 515 | AT3G47570 | AT3G47570 | ECA3 | AT1G10130 |
| 516 | WAK3 | AT1G21240 | NRT2:1 | AT1G08090 |
| 517 | AT3G60750 | AT3G60750 | DJ1B | AT1G53280 |
| 518 | AT3G27180 | AT3G27180 | At1g54130 | AT1G54130 |
| 519 | AT5G35170 | AT5G35170 | AT3G60750 | AT3G60750 |
| 520 | emb2746 | AT5G63420 | AT3G27180 | AT3G27180 |
| 521 | AT3G47570 | AT3G47570 | NRT2:1 | AT1G08090 |
| 522 | At3g49670 | AT3G49670 | NRT2:1 | AT1G08090 |
| 523 | PDLP7 | AT5G37660 | LYK5 | AT2G33580 |
| 524 | HAK5 | AT4G13420 | KEA3 | AT4G04850 |
| 525 | TT4 | AT5G13930 | PCR2 | AT1G14870 |
| 526 | WR3 | AT5G50200 | EP3 | AT3G54420 |
| 527 | AT4G09040 | AT4G09040 | AT4G01310 | AT4G01310 |
| 528 | AKHSDH2 | AT4G19710 | At1g54130 | AT1G54130 |
| 529 | RBOHD | AT5G47910 | LYK5 | AT2G33580 |
| 530 | CYP71B11 | AT5G25120 | AT3G47570 | AT3G47570 |
| 531 | AT3G47570 | AT3G47570 | CYP89A6 | AT1G64940 |
| 532 | AT3G47570 | AT3G47570 | CYP98A8 | AT1G74540 |
| 533 | AT3G47570 | AT3G47570 | CYP76C4 | AT2G45550 |
| 534 | EMB2730 | AT5G02250 | AT4G17740 | AT4G17740 |
| 535 | At3g49670 | AT3G49670 | CYP711A1 | AT2G26170 |
| 536 | EDD1 | AT3G48110 | SVR1 | AT2G39140 |

| | | | | |
|---|---|---|---|---|
| 537 | At3g49670 | AT3G49670 | CYP71B35 | AT3G26310 |
| 538 | CYP94D2 | AT3G56630 | AT3G47570 | AT3G47570 |
| 539 | At3g49670 | AT3G49670 | CYP71B37 | AT3G26330 |
| 540 | CYP71B10 | AT5G57260 | At3g49670 | AT3G49670 |
| 541 | CYP71B10 | AT5G57260 | AT3G47570 | AT3G47570 |
| 542 | CYP94D2 | AT3G56630 | At3g49670 | AT3G49670 |
| 543 | B120 | AT4G21390 | ABCG40 | AT1G15520 |
| 544 | CYP71A25 | AT3G48280 | AT3G47570 | AT3G47570 |
| 545 | At3g49670 | AT3G49670 | CYP76C4 | AT2G45550 |
| 546 | CYP71B11 | AT5G25120 | At3g49670 | AT3G49670 |
| 547 | AT3G47570 | AT3G47570 | CYP79F2 | AT1G16400 |
| 548 | At3g49670 | AT3G49670 | CYP98A8 | AT1G74540 |
| 549 | AT3G47570 | AT3G47570 | CYP711A1 | AT2G26170 |
| 550 | At3g49670 | AT3G49670 | CYP71B24 | AT3G26230 |
| 551 | AT3G47570 | AT3G47570 | CYP71B34 | AT3G26300 |
| 552 | At3g49670 | AT3G49670 | CYP71B34 | AT3G26300 |
| 553 | AT3G47570 | AT3G47570 | CYP71B24 | AT3G26230 |
| 554 | AT3G47570 | AT3G47570 | CYP71B35 | AT3G26310 |
| 555 | AT3G47570 | AT3G47570 | CYP71B37 | AT3G26330 |
| 556 | At3g49670 | AT3G49670 | CYP79F2 | AT1G16400 |
| 557 | At3g49670 | AT3G49670 | CYP89A6 | AT1G64940 |
| 558 | RPE | AT5G61410 | Lhca5 | AT1G45474 |
| 559 | AT5G39530 | AT5G39530 | PAM68 | AT4G19100 |
| 560 | PAM68 | AT4G19100 | AT4G09040 | AT4G09040 |
| 561 | UVR8 | AT5G63860 | AT1G16260 | AT1G16260 |
| 562 | AUX1 | AT2G38120 | GH3.1 | AT2G14960 |
| 563 | AT3G27180 | AT3G27180 | AMI1 | AT1G08980 |
| 564 | AT5G35170 | AT5G35170 | EDD1 | AT3G48110 |
| 565 | EMB1030 | AT5G22800 | CCoAOMT1 | AT4G34050 |
| 566 | ACP4 | AT4G25050 | BCAT3 | AT3G49680 |
| 567 | HINT3 | AT5G48545 | J8 | AT1G80920 |
| 568 | AKHSDH2 | AT4G19710 | emb2184 | AT1G75350 |
| 569 | AT5G65600 | AT5G65600 | AT3G23550 | AT3G23550 |
| 570 | CYP71B11 | AT5G25120 | AT1G66330 | AT1G66330 |
| 571 | AT1G61420 | AT1G61420 | HMGB2 | AT1G20693 |
| 572 | RPE | AT5G61410 | FAD6 | AT4G30950 |
| 573 | TT4 | AT5G13930 | MYB15 | AT3G23250 |
| 574 | CKA1 | AT5G67380 | TPR10 | AT3G04710 |
| 575 | J8 | AT1G80920 | AT1G50000 | AT1G50000 |
| 576 | SVR1 | AT2G39140 | NADK2 | AT1G21640 |
| 577 | PSBO2 | AT3G50820 | emb2184 | AT1G75350 |

| | | | | |
|---|---|---|---|---|
| 578 | AT5G12000 | AT5G12000 | AT2G39670 | AT2G39670 |
| 579 | AT5G51770 | AT5G51770 | AT2G39670 | AT2G39670 |
| 580 | CLA1 | AT4G15560 | AT3G27180 | AT3G27180 |
| 581 | AT2G39670 | AT2G39670 | LYK5 | AT2G33580 |
| 582 | AT2G39670 | AT2G39670 | AT1G61420 | AT1G61420 |
| 583 | AT2G39670 | AT2G39670 | CRK2 | AT1G70520 |
| 584 | AT5G65600 | AT5G65600 | AT2G39670 | AT2G39670 |
| 585 | AT1G47480 | AT1G47480 | FMO1 | AT1G19250 |
| 586 | AT2G39670 | AT2G39670 | WAK2 | AT1G21270 |
| 587 | CCR3 | AT3G55950 | AT2G39670 | AT2G39670 |
| 588 | AT4G29990 | AT4G29990 | AT2G39670 | AT2G39670 |
| 589 | CERK1 | AT3G21630 | AT2G39670 | AT2G39670 |
| 590 | AT2G39670 | AT2G39670 | WAK5 | AT1G21230 |
| 591 | AT2G39670 | AT2G39670 | AT2G23450 | AT2G23450 |
| 592 | AT2G39670 | AT2G39670 | AT1G16260 | AT1G16260 |
| 593 | AT5G02070 | AT5G02070 | AT2G39670 | AT2G39670 |
| 594 | AT2G39670 | AT2G39670 | AT1G61590 | AT1G61590 |
| 595 | AT2G39670 | AT2G39670 | WAK3 | AT1G21240 |
| 596 | RBOHD | AT5G47910 | PR1 | AT2G14610 |
| 597 | B120 | AT4G21390 | AT2G39670 | AT2G39670 |
| 598 | AT5G10530 | AT5G10530 | AT2G39670 | AT2G39670 |
| 599 | AT2G39670 | AT2G39670 | CIPK9 | AT1G01140 |
| 600 | AT5G12000 | AT5G12000 | TPR10 | AT3G04710 |
| 601 | AT5G02070 | AT5G02070 | TPR10 | AT3G04710 |
| 602 | AT5G51770 | AT5G51770 | TPR10 | AT3G04710 |
| 603 | TPR10 | AT3G04710 | AT1G61590 | AT1G61590 |
| 604 | TPR10 | AT3G04710 | AT2G23450 | AT2G23450 |
| 605 | CB5-E | AT5G53560 | NEET | AT5G51720 |
| 606 | EMB1030 | AT5G22800 | PSAO | AT1G08380 |
| 607 | CCR3 | AT3G55950 | TPR10 | AT3G04710 |
| 608 | AT3G48200 | AT3G48200 | PnsB2 | AT1G64770 |
| 609 | EMB3136 | AT5G13510 | At1g72610 | AT1G72610 |
| 610 | UKL2 | AT3G27190 | ALDH2C4 | AT3G24503 |
| 611 | APE2 | AT5G46110 | AT5G35170 | AT5G35170 |
| 612 | YY1 | AT4G06634 | TPR10 | AT3G04710 |
| 613 | TPR10 | AT3G04710 | AT2G28710 | AT2G28710 |
| 614 | At3g49670 | AT3G49670 | J8 | AT1G80920 |
| 615 | AT5G10530 | AT5G10530 | ABCG40 | AT1G15520 |
| 616 | AT3G47570 | AT3G47570 | J8 | AT1G80920 |
| 617 | PAM68 | AT4G19100 | AT2G42220 | AT2G42220 |
| 618 | AT5G35170 | AT5G35170 | PSBO2 | AT3G50820 |

| | | | | |
|---|---|---|---|---|
| 619 | At3g49670 | AT3G49670 | LOS2 | AT2G36530 |
| 620 | AT3G47570 | AT3G47570 | LOS2 | AT2G36530 |
| 621 | LYK5 | AT2G33580 | NRT2:1 | AT1G08090 |
| 622 | AT1G61420 | AT1G61420 | NRT2:1 | AT1G08090 |
| 623 | CCR3 | AT3G55950 | NRT2:1 | AT1G08090 |
| 624 | NRT2:1 | AT1G08090 | CIPK9 | AT1G01140 |
| 625 | AT5G12000 | AT5G12000 | NRT2:1 | AT1G08090 |
| 626 | AT1G16260 | AT1G16260 | NRT2:1 | AT1G08090 |
| 627 | AT4G29990 | AT4G29990 | NRT2:1 | AT1G08090 |
| 628 | AT2G23450 | AT2G23450 | NRT2:1 | AT1G08090 |
| 629 | WAK2 | AT1G21270 | NRT2:1 | AT1G08090 |
| 630 | CRK2 | AT1G70520 | NRT2:1 | AT1G08090 |
| 631 | CNI1 | AT5G27420 | MYB15 | AT3G23250 |
| 632 | AT5G02070 | AT5G02070 | NRT2:1 | AT1G08090 |
| 633 | AT5G10530 | AT5G10530 | NRT2:1 | AT1G08090 |
| 634 | WAK5 | AT1G21230 | NRT2:1 | AT1G08090 |
| 635 | AT5G51770 | AT5G51770 | NRT2:1 | AT1G08090 |
| 636 | AT5G65600 | AT5G65600 | NRT2:1 | AT1G08090 |
| 637 | CLA1 | AT4G15560 | GA2 | AT1G79460 |
| 638 | AT1G61590 | AT1G61590 | NRT2:1 | AT1G08090 |
| 639 | B120 | AT4G21390 | NRT2:1 | AT1G08090 |
| 640 | AT5G19850 | AT5G19850 | AT1G66330 | AT1G66330 |
| 641 | CERK1 | AT3G21630 | NRT2:1 | AT1G08090 |
| 642 | ATGSTF13 | AT3G62760 | SAG24 | AT1G66580 |
| 643 | AT4G20820 | AT4G20820 | DJ1B | AT1G53280 |
| 644 | CCoAOMT1 | AT4G34050 | CYP98A8 | AT1G74540 |
| 645 | ERD4 | AT1G30360 | AT1G15130 | AT1G15130 |
| 646 | NAPRT1 | AT4G36940 | AKHSDH2 | AT4G19710 |
| 647 | GulLO2 | AT2G46750 | DJ1B | AT1G53280 |
| 648 | emb2184 | AT1G75350 | AT1G45110 | AT1G45110 |
| 649 | At1g54130 | AT1G54130 | AT1G50000 | AT1G50000 |
| 650 | AT4G17740 | AT4G17740 | DJ1B | AT1G53280 |
| 651 | AT2G39670 | AT2G39670 | CDS1 | AT1G62430 |
| 652 | emb2746 | AT5G63420 | CCoAOMT1 | AT4G34050 |
| 653 | AT4G39970 | AT4G39970 | ISA3 | AT4G09020 |
| 654 | NDPK2 | AT5G63310 | AT1G78140 | AT1G78140 |
| 655 | BCAT3 | AT3G49680 | EDD1 | AT3G48110 |
| 656 | SERAT1;1 | AT5G56760 | ADC2 | AT4G34710 |
| 657 | HINT3 | AT5G48545 | EMB1030 | AT5G22800 |
| 658 | HINT3 | AT5G48545 | AT1G50000 | AT1G50000 |
| 659 | AT3G60750 | AT3G60750 | AT1G50000 | AT1G50000 |

| | | | | |
|---|---|---|---|---|
| 660 | AT1G45110 | AT1G45110 | FATB | AT1G08510 |
| 661 | RPE | AT5G61410 | PnsB2 | AT1G64770 |
| 662 | AT5G48220 | AT5G48220 | AT2G04360 | AT2G04360 |
| 663 | TPR10 | AT3G04710 | RAB11c | AT1G09630 |
| 664 | AT4G17740 | AT4G17740 | AT1G50000 | AT1G50000 |
| 665 | AT3G27180 | AT3G27180 | AT2G39670 | AT2G39670 |
| 666 | NDPK2 | AT5G63310 | emb2184 | AT1G75350 |
| 667 | EMB1030 | AT5G22800 | CYP38 | AT3G01480 |
| 668 | AT5G48220 | AT5G48220 | AT1G77090 | AT1G77090 |
| 669 | KEA3 | AT4G04850 | AT3G48200 | AT3G48200 |
| 670 | PSBO2 | AT3G50820 | PnsB2 | AT1G64770 |
| 671 | RPM1 | AT3G07040 | FMO1 | AT1G19250 |
| 672 | NDPK2 | AT5G63310 | SHY2 | AT1G04240 |
| 673 | AKHSDH2 | AT4G19710 | KEA3 | AT4G04850 |
| 674 | PSBO2 | AT3G50820 | AT1G18060 | AT1G18060 |
| 675 | KEA3 | AT4G04850 | ALDH2C4 | AT3G24503 |
| 676 | DAP | AT4G33680 | AT2G38900 | AT2G38900 |
| 677 | PIP1;4 | AT4G00430 | UKL2 | AT3G27190 |
| 678 | NEET | AT5G51720 | AT4G34480 | AT4G34480 |
| 679 | RBCL | ATCG00490 | TT4 | AT5G13930 |
| 680 | AT5G48220 | AT5G48220 | MTO2 | AT4G29840 |
| 681 | EMB3136 | AT5G13510 | AT4G28080 | AT4G28080 |
| 682 | BG1 | AT3G57270 | BG3 | AT3G57240 |
| 683 | GA1 | AT4G02780 | SHY2 | AT1G04240 |
| 684 | AUX1 | AT2G38120 | GA2 | AT1G79460 |
| 685 | FAD6 | AT4G30950 | AT4G01310 | AT4G01310 |
| 686 | CB5-E | AT5G53560 | CAT1 | AT1G20630 |
| 687 | AUX1 | AT2G38120 | RR12 | AT2G25180 |
| 688 | MTO2 | AT4G29840 | EDD1 | AT3G48110 |
| 689 | CLA1 | AT4G15560 | KEA3 | AT4G04850 |
| 690 | AT3G23550 | AT3G23550 | ABCG40 | AT1G15520 |
| 691 | ALN | AT4G04955 | AMI1 | AT1G08980 |
| 692 | RPE | AT5G61410 | AT5G35170 | AT5G35170 |
| 693 | AT4G39970 | AT4G39970 | AT3G51350 | AT3G51350 |
| 694 | AT4G09040 | AT4G09040 | SVR1 | AT2G39140 |
| 695 | AT5G06130 | AT5G06130 | CYP38 | AT3G01480 |
| 696 | AT3G48200 | AT3G48200 | AT1G77090 | AT1G77090 |
| 697 | emb2184 | AT1G75350 | AT1G51400 | AT1G51400 |
| 698 | ABCB20 | AT3G55320 | At3g49670 | AT3G49670 |
| 699 | RPE | AT5G61410 | AT1G64680 | AT1G64680 |
| 700 | RBOHD | AT5G47910 | HB1 | AT2G16060 |

| | | | | |
|---|---|---|---|---|
| 701 | ABCB20 | AT3G55320 | AT3G47570 | AT3G47570 |
| 702 | TT4 | AT5G13930 | CLA1 | AT4G15560 |
| 703 | RPM1 | AT3G07040 | PR1 | AT2G14610 |
| 704 | GSTU8 | AT3G09270 | NAC042 | AT2G43000 |
| 705 | AT4G09040 | AT4G09040 | AT2G37660 | AT2G37660 |
| 706 | AT4G28080 | AT4G28080 | GR | AT3G54660 |
| 707 | BG3 | AT3G57240 | EP3 | AT3G54420 |
| 708 | MRE11 | AT5G54260 | APTX | AT5G01310 |
| 709 | GSTU8 | AT3G09270 | GSTU19 | AT1G78380 |
| 710 | WRKY75 | AT5G13080 | AT3G22600 | AT3G22600 |
| 711 | RPM1 | AT3G07040 | PR4 | AT3G04720 |
| 712 | AT4G26860 | AT4G26860 | AT1G18060 | AT1G18060 |
| 713 | ACP4 | AT4G25050 | Lhca5 | AT1G45474 |
| 714 | FAD6 | AT4G30950 | emb2184 | AT1G75350 |
| 715 | AT3G25727 | AT3G25727 | AT2G13980 | AT2G13980 |
| 716 | AT5G35170 | AT5G35170 | LOS2 | AT2G36530 |
| 717 | MYB15 | AT3G23250 | GH3.1 | AT2G14960 |
| 718 | AT5G10770 | AT5G10770 | PRX52 | AT5G05340 |
| 719 | AT5G38570 | AT5G38570 | AT3G25727 | AT3G25727 |
| 720 | BG3 | AT3G57240 | PR1 | AT2G14610 |
| 721 | AT4G11290 | AT4G11290 | AT3G22600 | AT3G22600 |
| 722 | BG3 | AT3G57240 | AT2G38900 | AT2G38900 |
| 723 | CYP38 | AT3G01480 | AT1G78915 | AT1G78915 |
| 724 | AT5G42500 | AT5G42500 | AT3G22600 | AT3G22600 |
| 725 | AT3G22600 | AT3G22600 | AT1G61590 | AT1G61590 |
| 726 | AT5G48220 | AT5G48220 | AT3G45050 | AT3G45050 |
| 727 | LOS2 | AT2G36530 | CAT1 | AT1G20630 |
| 728 | AT5G42500 | AT5G42500 | AT1G61590 | AT1G61590 |
| 729 | AT5G38570 | AT5G38570 | AT2G13980 | AT2G13980 |
| 730 | FAD6 | AT4G30950 | AT2G42220 | AT2G42220 |
| 731 | YY1 | AT4G06634 | AT1G15130 | AT1G15130 |
| 732 | AT4G33720 | AT4G33720 | WAK5 | AT1G21230 |
| 733 | AT3G06390 | AT3G06390 | AT1G61590 | AT1G61590 |
| 734 | GR | AT3G54660 | UKL2 | AT3G27190 |
| 735 | PSBO2 | AT3G50820 | AT1G64680 | AT1G64680 |
| 736 | EMB3136 | AT5G13510 | AT2G42220 | AT2G42220 |
| 737 | PDX2 | AT5G60540 | NAPRT1 | AT4G36940 |
| 738 | AT3G47570 | AT3G47570 | RPM1 | AT3G07040 |
| 739 | AT5G19850 | AT5G19850 | ACP4 | AT4G25050 |
| 740 | ACP4 | AT4G25050 | XPL1 | AT3G18000 |
| 741 | NHL25 | AT5G36970 | ABCG40 | AT1G15520 |

| | | | | |
|---|---|---|---|---|
| 742 | ACP4 | AT4G25050 | EDD1 | AT3G48110 |
| 743 | AT5G19850 | AT5G19850 | KEA3 | AT4G04850 |
| 744 | CNI1 | AT5G27420 | PCR2 | AT1G14870 |
| 745 | AT4G01310 | AT4G01310 | AT2G39670 | AT2G39670 |
| 746 | AT4G39970 | AT4G39970 | AT1G77090 | AT1G77090 |
| 747 | EDD1 | AT3G48110 | AT2G04360 | AT2G04360 |
| 748 | APE2 | AT5G46110 | AT1G18060 | AT1G18060 |
| 749 | EDD1 | AT3G48110 | AT2G39670 | AT2G39670 |
| 750 | At3g49670 | AT3G49670 | HMGB2 | AT1G20693 |
| 751 | AT3G47570 | AT3G47570 | HMGB2 | AT1G20693 |
| 752 | EMB1030 | AT5G22800 | AT1G78915 | AT1G78915 |
| 753 | ALN | AT4G04955 | AT2G03200 | AT2G03200 |
| 754 | BCAT3 | AT3G49680 | CYP79F2 | AT1G16400 |
| 755 | TT4 | AT5G13930 | HCHIB | AT3G12500 |
| 756 | ACP4 | AT4G25050 | CDS1 | AT1G62430 |
| 757 | RBOHD | AT5G47910 | YLS9 | AT2G35980 |
| 758 | AT5G48220 | AT5G48220 | EMB3136 | AT5G13510 |
| 759 | NHL25 | AT5G36970 | B120 | AT4G21390 |
| 760 | WR3 | AT5G50200 | NRT1:2 | AT1G69850 |
| 761 | FAD6 | AT4G30950 | AT2G04360 | AT2G04360 |
| 762 | AT1G78140 | AT1G78140 | FMO1 | AT1G19250 |
| 763 | AT2G42220 | AT2G42220 | AT1G77090 | AT1G77090 |
| 764 | PSBA | ATCG00020 | AT1G77090 | AT1G77090 |
| 765 | EMB2730 | AT5G02250 | PP2A-4 | AT3G58500 |
| 766 | NDPK2 | AT5G63310 | CDS1 | AT1G62430 |
| 767 | PAM68 | AT4G19100 | PSAF | AT1G31330 |
| 768 | AT5G58490 | AT5G58490 | AKHSDH2 | AT4G19710 |
| 769 | AKHSDH2 | AT4G19710 | J8 | AT1G80920 |
| 770 | PLAT2 | AT2G22170 | AT1G18060 | AT1G18060 |
| 771 | PSBO2 | AT3G50820 | GSTU18 | AT1G10360 |
| 772 | ATGSTF13 | AT3G62760 | AT2G47870 | AT2G47870 |
| 773 | AT1G51400 | AT1G51400 | GSTU18 | AT1G10360 |
| 774 | BG1 | AT3G57270 | AT2G38900 | AT2G38900 |
| 775 | AT5G65600 | AT5G65600 | NHL25 | AT5G36970 |
| 776 | PYL2 | AT2G26040 | ABCG40 | AT1G15520 |
| 777 | YY1 | AT4G06634 | BBX15 | AT1G25440 |
| 778 | GATA9 | AT4G32890 | AT2G28710 | AT2G28710 |
| 779 | AT2G28710 | AT2G28710 | BBX15 | AT1G25440 |
| 780 | RPOC1 | ATCG00180 | AKHSDH2 | AT4G19710 |
| 781 | RPOC1 | ATCG00180 | EMB1030 | AT5G22800 |
| 782 | GATA9 | AT4G32890 | YY1 | AT4G06634 |

| | | | | |
|---|---|---|---|---|
| 783 | AT3G22600 | AT3G22600 | AT3G06390 | AT3G06390 |
| 784 | AT2G36290 | AT2G36290 | MLP165 | AT1G35260 |
| 785 | SVP | AT2G22540 | BBX15 | AT1G25440 |
| 786 | OMT1 | AT5G54160 | TT4 | AT5G13930 |
| 787 | CGLD27 | AT5G67370 | PSBO2 | AT3G50820 |
| 788 | PGRL1A | AT4G22890 | AT2G37660 | AT2G37660 |
| 789 | AT4G11290 | AT4G11290 | AT1G61590 | AT1G61590 |
| 790 | CYP71B10 | AT5G57260 | AT3G19240 | AT3G19240 |
| 791 | AT4G16260 | AT4G16260 | AT2G28710 | AT2G28710 |
| 792 | AT3G48200 | AT3G48200 | AT1G64680 | AT1G64680 |
| 793 | AT4G09040 | AT4G09040 | AMI1 | AT1G08980 |
| 794 | CYP38 | AT3G01480 | AT1G51400 | AT1G51400 |
| 795 | AT4G01310 | AT4G01310 | LOS2 | AT2G36530 |
| 796 | UKL2 | AT3G27190 | At1g54130 | AT1G54130 |
| 797 | EMB1030 | AT5G22800 | AT4G28080 | AT4G28080 |
| 798 | RPE | AT5G61410 | AT1G77090 | AT1G77090 |
| 799 | AT5G39160 | AT5G39160 | PAP22 | AT3G52820 |
| 800 | NDPK2 | AT5G63310 | TT4 | AT5G13930 |
| 801 | AKHSDH2 | AT4G19710 | FD3 | AT2G27510 |
| 802 | FAD6 | AT4G30950 | AT4G28080 | AT4G28080 |
| 803 | AT4G28080 | AT4G28080 | PnsB2 | AT1G64770 |
| 804 | AT5G26330 | AT5G26330 | ALN | AT4G04955 |
| 805 | AT4G26860 | AT4G26860 | AT1G52870 | AT1G52870 |
| 806 | AT1G78915 | AT1G78915 | AT1G51110 | AT1G51110 |
| 807 | MYB15 | AT3G23250 | CYP38 | AT3G01480 |
| 808 | AT2G28710 | AT2G28710 | AT1G15130 | AT1G15130 |
| 809 | EMB3136 | AT5G13510 | UKL2 | AT3G27190 |
| 810 | ACP4 | AT4G25050 | UKL2 | AT3G27190 |
| 811 | GR | AT3G54660 | CYP38 | AT3G01480 |
| 812 | AT5G08390 | AT5G08390 | ECA3 | AT1G10130 |
| 813 | CERK1 | AT3G21630 | YLS9 | AT2G35980 |
| 814 | PAM68 | AT4G19100 | AT4G01310 | AT4G01310 |
| 815 | ACP4 | AT4G25050 | PSAF | AT1G31330 |
| 816 | FAD6 | AT4G30950 | CLA1 | AT4G15560 |
| 817 | UKL2 | AT3G27190 | J8 | AT1G80920 |
| 818 | NHL25 | AT5G36970 | AT3G23550 | AT3G23550 |
| 819 | AT2G39670 | AT2G39670 | AT1G45110 | AT1G45110 |
| 820 | AT5G35170 | AT5G35170 | EMB3136 | AT5G13510 |
| 821 | AT5G48220 | AT5G48220 | ELI3-1 | AT4G37980 |
| 822 | CYP71A25 | AT3G48280 | AT1G19450 | AT1G19450 |
| 823 | AT5G35170 | AT5G35170 | AT1G77090 | AT1G77090 |

| | | | | |
|---|---|---|---|---|
| 824 | ALN | AT4G04955 | UKL2 | AT3G27190 |
| 825 | AKHSDH2 | AT4G19710 | RR12 | AT2G25180 |
| 826 | emb2184 | AT1G75350 | PSAO | AT1G08380 |
| 827 | LPA2 | AT5G51545 | SVR1 | AT2G39140 |
| 828 | AUX1 | AT2G38120 | CYP711A1 | AT2G26170 |
| 829 | RPOC1 | ATCG00180 | UKL2 | AT3G27190 |
| 830 | UKL2 | AT3G27190 | AT3G23550 | AT3G23550 |
| 831 | AT5G19850 | AT5G19850 | AKHSDH2 | AT4G19710 |
| 832 | CGLD27 | AT5G67370 | AT5G39530 | AT5G39530 |
| 833 | AT5G48220 | AT5G48220 | AT2G39670 | AT2G39670 |
| 834 | AT4G13010 | AT4G13010 | KEA3 | AT4G04850 |
| 835 | TT4 | AT5G13930 | PR4 | AT3G04720 |
| 836 | DIT2.1 | AT5G64290 | AT3G60750 | AT3G60750 |
| 837 | AT2G47870 | AT2G47870 | CAT1 | AT1G20630 |
| 838 | AT5G19890 | AT5G19890 | WAK5 | AT1G21230 |
| 839 | CGLD27 | AT5G67370 | AT5G19850 | AT5G19850 |
| 840 | AT5G48220 | AT5G48220 | AT1G51110 | AT1G51110 |
| 841 | AT2G39670 | AT2G39670 | AT2G04360 | AT2G04360 |
| 842 | AT5G58490 | AT5G58490 | SVR1 | AT2G39140 |
| 843 | PP2A-4 | AT3G58500 | UKL2 | AT3G27190 |
| 844 | DMR6 | AT5G24530 | PCR2 | AT1G14870 |
| 845 | RPE | AT5G61410 | LOS2 | AT2G36530 |
| 846 | HINT3 | AT5G48545 | ERD9 | AT1G10370 |
| 847 | HINT3 | AT5G48545 | GSTU8 | AT3G09270 |
| 848 | HINT3 | AT5G48545 | GSTU18 | AT1G10360 |
| 849 | AT1G78915 | AT1G78915 | AT1G77090 | AT1G77090 |
| 850 | RPOC1 | ATCG00180 | LOS2 | AT2G36530 |
| 851 | NAPRT1 | AT4G36940 | LOS2 | AT2G36530 |
| 852 | PSBO2 | AT3G50820 | AT2G37660 | AT2G37660 |
| 853 | AT5G48220 | AT5G48220 | EMB1030 | AT5G22800 |
| 854 | ACP4 | AT4G25050 | AT3G60750 | AT3G60750 |
| 855 | RPM1 | AT3G07040 | YLS9 | AT2G35980 |
| 856 | AT5G58490 | AT5G58490 | At3g49670 | AT3G49670 |
| 857 | AT5G58490 | AT5G58490 | AT3G47570 | AT3G47570 |
| 858 | AT4G17740 | AT4G17740 | CYP38 | AT3G01480 |
| 859 | YY1 | AT4G06634 | ECA3 | AT1G10130 |
| 860 | AT4G28080 | AT4G28080 | AT4G01310 | AT4G01310 |
| 861 | AT3G60750 | AT3G60750 | PKT3 | AT2G33150 |
| 862 | FRS5 | AT4G38180 | bZIP16 | AT2G35530 |
| 863 | BTS | AT3G18290 | VIT1 | AT2G01770 |
| 864 | AT4G01310 | AT4G01310 | AT1G18060 | AT1G18060 |

| | | | | |
|---|---|---|---|---|
| 865 | PSBO2 | AT3G50820 | At1g72610 | AT1G72610 |
| 866 | BG1 | AT3G57270 | UGT85A3 | AT1G22380 |
| 867 | AT3G48200 | AT3G48200 | CYP38 | AT3G01480 |
| 868 | HAM3 | AT4G00150 | BG1 | AT3G57270 |
| 869 | LAC7 | AT3G09220 | AT1G01490 | AT1G01490 |
| 870 | AT4G09040 | AT4G09040 | AT2G39670 | AT2G39670 |
| 871 | HAM3 | AT4G00150 | BG3 | AT3G57240 |
| 872 | AT4G16260 | AT4G16260 | HAM3 | AT4G00150 |
| 873 | CGLD27 | AT5G67370 | FD3 | AT2G27510 |
| 874 | AKHSDH2 | AT4G19710 | LAP1 | AT2G24200 |
| 875 | KEA3 | AT4G04850 | GulLO2 | AT2G46750 |
| 876 | NEET | AT5G51720 | UGT85A3 | AT1G22380 |
| 877 | AT4G20820 | AT4G20820 | KEA3 | AT4G04850 |
| 878 | At3g49670 | AT3G49670 | CAT1 | AT1G20630 |
| 879 | LPA2 | AT5G51545 | emb2184 | AT1G75350 |
| 880 | AT3G47570 | AT3G47570 | CAT1 | AT1G20630 |
| 881 | CGLD27 | AT5G67370 | PAM68 | AT4G19100 |
| 882 | YLS9 | AT2G35980 | PCR2 | AT1G14870 |
| 883 | CYP71A25 | AT3G48280 | ALDH2C4 | AT3G24503 |
| 884 | AT4G28080 | AT4G28080 | AT3G60750 | AT3G60750 |
| 885 | KEA3 | AT4G04850 | AT1G16260 | AT1G16260 |
| 886 | AT4G26860 | AT4G26860 | CYP38 | AT3G01480 |
| 887 | AT4G09040 | AT4G09040 | AT1G78915 | AT1G78915 |
| 888 | NDPK2 | AT5G63310 | AT5G48220 | AT5G48220 |
| 889 | EMB1030 | AT5G22800 | EMB2730 | AT5G02250 |
| 890 | INO80 | AT3G57300 | BT2 | AT3G48360 |
| 891 | AT4G10540 | AT4G10540 | CYP76C4 | AT2G45550 |
| 892 | FMO1 | AT1G19250 | ABCG40 | AT1G15520 |
| 893 | ALDH2C4 | AT3G24503 | LOS2 | AT2G36530 |
